# Integrated transcriptomic analysis identifies Paternally Expressed Gene 10 as a novel candidate biomarker associated with aggressive features in liposarcoma

**DOI:** 10.64898/2026.09.24.754104

**Authors:** Turiello Roberta, Scognamiglio Giosuè, Bonicelli Andrea, Gitano Giorgia, Manco Emanuele, Olivieri Sara, Barbato Anna, Fruggiero Carmine, Salati Massimiliano, Iuliano Antonella, Cerrone Margherita, Arcucci Alessandro, Martorana Anna, De Pietro Giovanna, Santella Emili, Fernanda Picozzi, Cantile Monica, Tafuto Salvatore, Budillon Alfredo, Franco Brunella, Carotenuto Pietro

## Abstract

**Background:** Liposarcoma is a rare cancer, comprising four major histological subtypes: well-differentiated, dedifferentiated, myxoid and pleomorphic liposarcoma. Therapeutic options remain limited for advanced disease, and additional biomarkers that may support biological characterization, diagnosis, or risk stratification are needed.

**Methods:** Bulk RNA sequencing (RNA-seq) was performed on paired tumour and adjacent non-tumour tissues from an analysable subset of 16 patients (22 tumour specimens) within a clinicopathological cohort of 21 patients. Differential gene expression analysis, principal component analysis, gene set enrichment (GSEA) analysis and single-sample GSEA (ssGSEA) were used to define subtype-specific transcriptional programs and imprinted-gene dysregulation. *PEG10* expression was validated by RT-qPCR and RNA fluorescence in situ hybridization (FISH), and its cellular localization was explored by combined FISH and immunofluorescence for P53, vimentin, Ki67, and CD34. *PEG10* expression in liposarcoma was further investigated in an external single-cell RNA-sequencing dataset of dedifferentiated liposarcoma. An exploratory survival analysis was performed in the pan-sarcoma TCGA-SARC cohort.

**Results:** Transcriptomic profiling identified distinct subtype-specific gene expression patterns and broad dysregulation of imprinted genes in liposarcoma compared with adjacent non-tumour tissue. Among all the imprinted genes, PEG10 showed a marked tumour-to-normal expression difference and good discriminatory performance between tumour and normal tissues in the discovery cohort (AUC=0.867). In the TCGA sarcoma cohort high PEG10 expression was associated with shorter overall survival. Immuno-FISH and single-cell analyses revealed that *PEG10* was predominantly detected in tumour and proliferating cell populations. Importantly, *PEG10* expression was higher in high-grade tumours. In bulk RNA-seq data, *PEG10* expression correlated with hypoxia, extracellular matrix remodelling and invasion ssGSEA signatures. In the external single-cell RNA-seq dataset PEG10^high^ tumours showed a reduced immune cell infiltration.

**Conclusions:** This study provides an integrated transcriptomic characterisation of liposarcoma and identifies *PEG10* as a candidate tumour-associated biomarker linked to aggressive disease features. The diagnostic and prognostic value of PEG10 requires validation in larger, independent, subtype-balanced liposarcoma cohorts, and its potential therapeutic relevance requires direct functional investigation.

## Background

Liposarcomas are rare mesenchymal malignancies characterized by adipocytic differentiation. These tumours account for 20% of all soft tissue sarcoma and are characterised by high intra- and inter-tumour heterogeneity (1). Liposarcomas are classified into four subtypes, which in some cases coexist within the same lesion. The subtypes include well-differentiated liposarcoma (WDLPS), also known as atypical lipomatous tumour, dedifferentiated liposarcomas (DDLPS), myxoid liposarcomas (MLPS), and pleomorphic liposarcomas (PLPS) (1,2). These subtypes differ in their histological and molecular features and exhibit distinct clinical responses to the available treatments.

WDLPS, the most common among the subtypes, arises in the retroperitoneum and proximal extremities, and it is characterised by high proliferation of mature adipocytes (3). This subtype is successfully treated with surgical resection and does not have invasive and metastatic potential per se, even though it can show local recurrence. DDLPS can arise within WDLPS and is associated with a substantially greater risk of recurrence and metastatic dissemination, which contribute to poor outcome of patients. Notably, distant metastases, when present, are always composed of dedifferentiated cells (4,5).

DDLPS is characterised by the presence of both lipogenic and undifferentiated populations, with a variable frequency of each. This subtype represents the most aggressive form of liposarcoma, with reported survival rates of 50% and 35% at 5 and 10 years, respectively (6). Both WDLPS and DDLPS are characterized by amplification of chromosome 12q14–15, resulting in *MDM2* and *CDK4* overexpression, in 100% and 90% of cases, respectively (7,8). The activity of both MDM2 and CDK4 promotes the uncontrolled proliferation of tumour cells. Indeed, MDM2 is not only able to inhibit TP53, a known onco-suppressor (9), but it also stimulates the synthesis of nucleotides, needed for cell proliferation (10). In addition to the protumour activity exerted by MDM2, the dysregulation of CDK4, a cyclin dependent kinase, drives the progression from the G1 to S phase of cell cycle (11).

MLPS originates mostly in proximal extremities, and it is characterised by non-adipocytic tumour cells and immature lipoblasts. This tumour often metastasizes to atypical sites, including bone, limbs, and retroperitoneum and is more sensitive to chemotherapy and radiotherapy than WDLPS and DDLPS (1). MLPS is characterised by two specific genomic rearrangements: approximately 95% of MLPS harbor the t(12;16)(q13;p11) translocation, which results in the FUS-DDIT3 fusion gene. Less frequently the t(12;22)(q13;q12) translocation is observed, producing the *EWSR1-DDIT3* fusion gene (12).

PLPS originates most frequently in limbs, rarely in the retroperitoneum. This subtype is the most aggressive type of liposarcoma and it is resistant to both chemotherapy and radiotherapy, with metastasis development in about 40% of patients (1). PLPS lacks specific molecular markers and is characterised by complex karyotypes, harboring multiple chromosomal aberrations (12).

Being a rare and highly heterogeneous type of tumour, diagnosis of liposarcoma is notably challenging, with misdiagnosis occurring in nearly 30% of cases (2,13,14). Liposarcomas are indeed difficult to distinguish from benign adipocytic tumours or other sarcomas (14,15).

In current clinical practice the assessment of MDM2/CDK4 amplification, most commonly by fluorescence in situ hybridization (FISH) is an important ancillary diagnostic tool for both WDLPS and DDLPS; CDK4 amplification or protein expression may provide additional support in selected cases (16).

Therapeutic options for advanced liposarcoma remain limited and vary substantially by histological subtype. MLPS is relatively radiosensitive and can respond to systemic therapy, whereas DDLPS, WDLPS and PLPS generally show more limited or heterogeneous responses to conventional systemic treatments (17).

In this context, the identification of new targets is of high priority, as well as the identification of biomarkers to improve the diagnosis and classification of these tumours.

To characterise the biology of liposarcoma more comprehensively, we integrated bulk RNA sequencing (RNA-seq) with an external single-cell RNA-seq dataset. We focused on Paternally Expressed Gene 10 (*PEG10*), an imprinted gene that showed marked tumour-associated upregulation in our cohort. We then evaluated its relationship with histological grade, MDM2 amplification status, transcriptional programs related to tumour aggressiveness, and the cellular composition of the tumour microenvironment.

Our integrated transcriptomic analysis allowed us to associate *PEG10* expression with common features of tumour aggressiveness and we propose *PEG1*0 as a potential candidate biomarker, requiring independent validation.

## Methods

### Patients and human samples

A clinicopathological cohort of 21 patients with liposarcoma was assembled. All liposarcoma samples were collected before any treatment and were available as formalin-fixed paraffin-embedded (FFPE) tissues. FFPE samples were sliced by standard procedures at 4 µm slice thickness and adhered onto poly-L-lysine-coated slides.

Subsequent tissue cuts were used to perform haematoxylin and eosin staining, immunohistochemistry, RNA fluorescence in situ hybridization (FISH) and RNA extraction, and to construct two tissue microarray (TMA) blocks.

Where applicable, tumour malignancy was scored according to the French Federation of Cancer Centers (FNCLCC) (grade 1, G1: low grade; grade 2, G2: intermediate grade; grade 3, G3; high grade). Samples were analysed via FISH to assess MDM2/CEP12 amplification, and DDIT3 rearrangement, as standard procedure of diagnosis.

### Tissue Microarray design and construction

FFPE blocks from the patients included in our cohort (n=21) were used to construct two TMA blocks. For each FFPE block, 4 cores (1 mm in diameter) were extracted: two representing the tumour tissue and two representing the adjacent normal tissue. Sections of 4 μM thickness were cut from each TMA block and used for subsequential experiments. For quantitative analyses, core-level measurements were aggregated at the patient level before statistical comparison.

### RNA isolation from FFPE tissues

Total RNA was extracted after microscopic dissection of the tumour and matched adjacent non-tumour regions, using the Maxwell® RSC RNA FFPE Kit (Promega, Madison, WI, USA) on the Maxwell® RSC Instruments (Promega, Madison, WI, USA) according to the manufacturer’s instructions (18). Adequate RNA for bulk RNA-seq was available from 16 patients, yielding 22 tumour specimens, as five patients contributed multiple spatially distinct tumour specimens.

### RNAseq library preparation and deep sequencing

For RNA sequencing (RNA-seq), libraries were generated starting from 250 ng of total RNA, using the QuantSeq 3’ mRNA-Seq Library Prep Kit FWD for Illumina (Lexogen GmbH, Wien, Austria), following the manufacturer’s protocol. Quality control of library templates was performed using a High Sensitivity D1000 Screen Tape (Agilent Technologies, Santa Clara, CA, USA) on a TapeStation 4200 (Agilent Technologies, Santa Clara, CA, USA). Sample concentrations were normalised using the Qubit quantification platform (Qubit 2.0 Fluorometer, Life Technologies/Thermo Fisher Scientific, MA, USA), prior to library preparation. Multiplexing allowed for pooling of up to 87 samples into a single lane, to ensure adequate depth of sequencing. The sequencing was carried out in collaboration with Next Generation Diagnostic S.r.l. (Pozzuoli, Naples, Italy), using the NovaSeq 6000 platform with single-end chemistry (SP 100 cycles; Illumina, Cambridge, UK). Libraries were loaded at a concentration of 250 pM, which was previously determined as optimal. An average yield of approximately 4.5 Mb was obtained per sample (18).

### RNA-Sequencing Data Pre-processing and Annotation

Bulk RNA-sequencing raw counts and metadata were processed using the R statistical environment (v.4.5.2). Ensembl gene identifiers were mapped to official HGNC gene symbols using the biomaRt package (v.2.60.0), In cases where gene symbols were unavailable, original Ensembl IDs were retained. Library size distribution was assessed for quality control, and median library sizes were calculated across all samples (19).

### Normalisation and Exploratory Data Analysis

Differential expression analysis was performed using the edgeR package (v.4.2.0). Low-expressed genes were filtered, retaining genes with a minimum count of 5 in at least 30% of the samples. Data normalisation was conducted using the Trimmed Mean of M-values (TMM). For exploratory visualisation, normalised counts were transformed to log2-counts per million (log2-CPM). Principal Component Analysis (PCA) and Multi-Dimensional Scaling (MDS) were used to explore sample-level structure, and hierarchical clustering was visualized with pheatmap package (v1.0.13).

### Differential Expression Analysis

Common, trended, and tagwise biological coefficient of variation (BCV) were estimated using edgeR’s robust empirical Bayes framework, and differential expression was modeled using generalized linear models (GLM) with quasi-likelihood testing. Two primary comparisons were conducted: (1) Tumour (T) vs. Normal (N) samples, and (2) *PEG10* stratification within the tumour subset by categorising into *PEG10*^High^ and *PEG10*^Low^ groups based on the median log2-CPM expression of the *PEG10* gene.

Statistical significance was determined using the Quasi-Likelihood F-test (glmQLFTest). Genes were considered significantly differentially expressed (DEG) if they met a threshold of an absolute log2-fold change ≥ 1 and a Benjamini-Hochberg False Discovery Rate (FDR < 0.05). Volcano plots were generated using the EnhancedVolcano package to visualize the magnitude and significance of gene expression changes.

### Gene Set Enrichment Analysis (GSEA)

Gene Set Enrichment Analysis (GSEA) was performed using the clusterProfiler package (v.4.12.0) to identify transcriptional programs associated with the comparisons of interest. Differentially expressed genes were defined using an absolute log2 fold-change threshold of log2FC > 1 and a Benjamini–Hochberg false discovery rate (FDR) < 0.05. Hallmark and Gene Ontology Biological Process gene sets were retrieved from the Molecular Signatures Database (MSigDB) using the msigdbr package.

For genomic-imprinting analyses, custom gene sets were generated using the human imprinted-gene database available at Geneimprint (https://geneimprint.com/). Genes annotated as imprinted were included in the overall imprinted-gene set and were further classified according to the annotated expressed parental allele into maternally expressed and paternally expressed gene sets. Gene annotations were downloaded from Geneimprint in September 2026.

Enrichment analyses were performed using 10,000 permutations, with a minimum gene-set size of 10 and a maximum of 800 genes. Enrichment results were summarized using normalized enrichment scores (NES), and multiple-testing correction was performed using the Benjamini– Hochberg method. An FDR < 0.05 was considered statistically significant.

### Single-cell data processing

Single-cell RNA sequencing (scRNA-seq) data from a liposarcoma atlas were retrieved from the Gene Expression Omnibus (GSE221493) and processed using Seurat v5 (20) in R version 4.5.2. Pre-computed normalised expression matrices, cell level metadata, and UMAP coordinates from the original study were used to preserve the original cell annotations and embedding (4). To maintain consistency with the source dataset, original UMAP coordinates were extracted and incorporated into the analysis object for visualisation. Patients were stratified into high and low expression groups based on the median patient-level mean *PEG10* expression to investigate its impact on cellular composition. Differences between the two groups were assessed using Wilcoxon rank-sum tests implemented in dittoSeq (21). Single-sample gene set enrichment (ssGSEA) scores were calculated using GSVA (v2.4.9) to derive sample-level pathway scores for the predefined adipocyte-differentiation, angiogenesis, extracellular-matrix remodelling, hypoxia, invasion, and stemness signatures. Pearson correlation was used to test associations between PEG10 expression and ssGSEA scores.

### Real-Time Reverse Transcription Quantitative Polymerase Chain Reaction (RT-qPCR)

Reverse transcription was performed with TaqMan reverse transcription kit (Life Technologies, Paisley, UK). RT-qPCR experiments were performed using TaqMan gene expression assay (ThermoFisher Scientific), following the manufacturer’s instructions. The following primer sets were included: PEG10 (assay ID Hs00248288_s1, cat. 4331182), 18S (Assay ID Hs99999901_s1, cat. 4331182), GAPDH (Assay ID Hs03929097_g1, cat. 4331182).

### Fluorescence in situ hybridization (FISH)

PEG10 mRNA was detected through FISH experiments using the following probe: RNAscope™ 2.5 LS Probe - Hs-PEG10 (Advanced Cell Diagnostics, Bio-Techne, Catalog #1187748-C1). Staining was performed using the LS Multiplex Fluorescent Reagent kit (Bio-Techne, catalog #322800) on the BOND RX Automated Research Stainer (Leica).

Briefly, tissues were incubated with the PEG10-specific probe at room temperature (RT) for 30 minutes. Fixation was performed in 10% neutral buffered formalin (NBF) at RT for 30 minutes, and the enzymatic pretreatment with ACD enzyme was conducted at 40°C for 30 minutes, followed by target retrieval at RT for 10 minutes. Probe hybridization was carried out at 42°C for 2 hours. Signal amplification consisted of three sequential amplification steps at 42°C, for 30 minutes each, with LS rinse steps at RT for 5 minutes, between amplifications. Samples were then incubated for 15 minutes at 42°C with HRP-C2. The signal was developed using TSA 570 fluorophore at RT for 30 minutes and HRP blocking was carried out at 42°C for 15 minutes. Samples were finally washed in wash buffer for 10 minutes at RT.

FISH Diagnostic Assessment of MDM2/CEP12 amplification and DDIT3 rearrangement was conducted following standard clinical practice of Istituto Nazionale per lo Studio e la Cura dei Tumori, IRCCS Fondazione “G. Pascale”.

### Histopathological analysis of tissues

For each FFPE sample, haematoxylin and eosin (HCE) staining was performed using standard protocols; slides were digitized using a ScanScope CS digital scanner (Aperio, Park Center Dr, Vista, CA, USA) at a 40x magnification. Images were annotated by pathologists.

### Immuno-FISH

FISH experiments for PEG10 detection were combined with immunofluorescence for the detection of selected protein of interest. After primary antibody incubation, the RNAscope workflow was performed with the modifications described below, followed by fluorophore development and DAPI nuclear counterstaining. Briefly, samples were first incubated with the primary antibody at RT for 30 minutes, then fixed in 10% NBF for 30 minutes at RT. Enzymatic pretreatment with ACD enzyme was performed at 40°C for 30 minutes, followed by target retrieval at RT for 10 minutes. Hybridization with PEG10-probe was carried out at 42°C for 2 hours and followed by three sequential amplification steps, as described above. Samples were incubated with HRP-C2 for 15 minutes at 42°C, and the signal was then developed using TSA 750 fluorophore at RT for 30 minutes. HRP blocking was then carried out at 42°C for 15 minutes. Subsequently, post-primary incubation and polymer incubation were carried out for 8 minutes each. The signal was developed using TSA 520 at RT for 10 minutes and the nuclear counterstaining was performed incubating the samples with DAPI at RT for 10 minutes.

The following antibodies were used: p53 (DAKO, clone DO-7, catalog #M7001, dilution 1:100, antigen retrieval pH 9), vimentin (Leica, clone V9, catalog #NCL-L-VIM-V9, dilution 1:800, antigen retrieval pH 6), Ki67 (Agilent, clone MIB1, catalog #M724, dilution 1:50, antigen retrieval pH 6), CD34 (Ventana-Roche, clone QBEnd/10, catalog #790-2927, prediluted, antigen retrieval pH 8).

The TMAs contained material from 21 patients, with two tumor cores and two adjacent non-tumor cores per case. Separate TMA sections were used for PEG10 FISH combined with each immunofluorescence marker. Samples with tissue loss, staining failure, or insufficient analysable material were excluded from the corresponding analysis. The number of evaluable patients is reported in each figure or legend.

### Image acquisition and analysis

FISH and Immuno-FISH images were acquired using a Leica DM 500 microscope. Once acquired, the images were analysed in QuPath (version 0.6.0) (22). Cell segmentation was performed using Instanseg (23), which documentation is available at https://github.com/instanseg/instanseg. QuPath object classifiers were used to categorise cells according to PEG10 FISH signal and positivity for either P53, vimentin, Ki67 or CD34.

PEG10 FISH signal intensity and spot counts were quantified in QuPath. Mean fluorescence intensity (MFI) and the number of PEG10 positive spots per cell were summarised at the patient level for group comparison.

### Statistical analysis

Statistical analyses were performed using GraphPad Prism (v10.0.3, GraphPad Software) or R Studio (R version 4.5.2). Exact tests, sample sizes, definitions of the statistical unit, and multiple-comparison procedures are reported in the figure legends. Two-sided P<0.05 was considered statistically significant, unless FDR control is specified.

### Figure preparation

Figures were prepared using Adobe Illustrator (v27.5, Adobe Inc.) and/or GraphPad Prism (v10.0.3, GraphPad Software)

## Results

### Clinicopathological characteristics of patients

A total of 21 patients diagnosed with liposarcoma were included in this study. The most represented subtypes were WDLPS (28.58%, n=6) and MLPS (28.57%, n=6), while DDLPS and PLPS represented 23.80% (n=5) and 19.05% (n=4) of cases, respectively. The characteristics of patients are summarised in Table1 and Fig 1A. HCE images for each sample are included in Supplementary File 1. As per standard clinical practice, all samples were evaluated by FISH for MDM2/CEP12 amplification and DDIT3 rearrangement. MDM2/CEP12 amplification was found in 5/6 patients diagnosed with WDLPS and 4/5 patients with DDLPS, while DDIT3 was rearranged in all 6 patients with MLPS. The single WDLPS and single DDLPS case lacking detectable MDM2 amplification were re-reviewed by a pathologist, confirming the morphology, and retained in the cohort on this basis.

**Figure 1.**
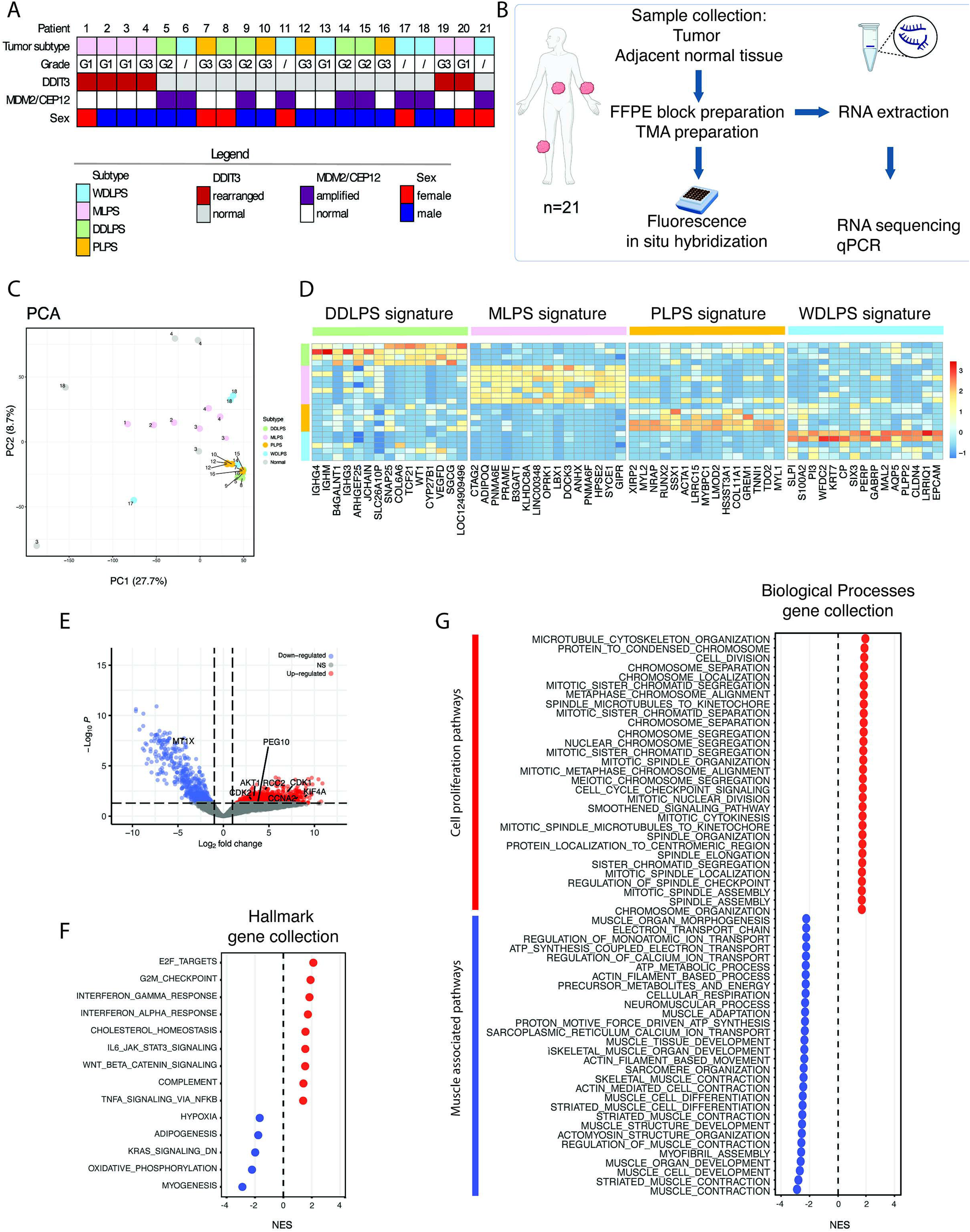
Transcriptomics analysis of liposarcoma tumours and paired adjacent normal tissues. A) Clinicopathological characteristics of the 21 patients included in our cohort: tumour subtype, grade, DDIT3 status and MDM2/CEP12 amplification and sex. B) Experimental workflow C) PCA of bulk RNA-seq data showing the transcriptional separation of liposarcoma samples (n=22) according to histological subtype and from normal tissue (n=5). D) Heatmap showing the subtype-specific gene signatures identified by edgeR. For each liposarcoma subtype, the top 15 upregulated genes were selected from subtype-versus-rest differential expression analysis (FDR<0.05, log2FC>1). Expression values are shown as row-wise z-scores, with samples ordered by histological subtype. E) Volcano plot showing the differentially expressed genes in tumour tissues compared with normal tissues. F) Dot plot showing the enriched gene sets from the Gene Ontology Hallmark gene set, in tumour tissues and normal tissues (FDR<0.05). G) Dot plot showing the enriched gene sets from the Gene Ontology Biological Process gene set, in tumour tissues and normal tissues (FDR<0.05).

**Table 1.**
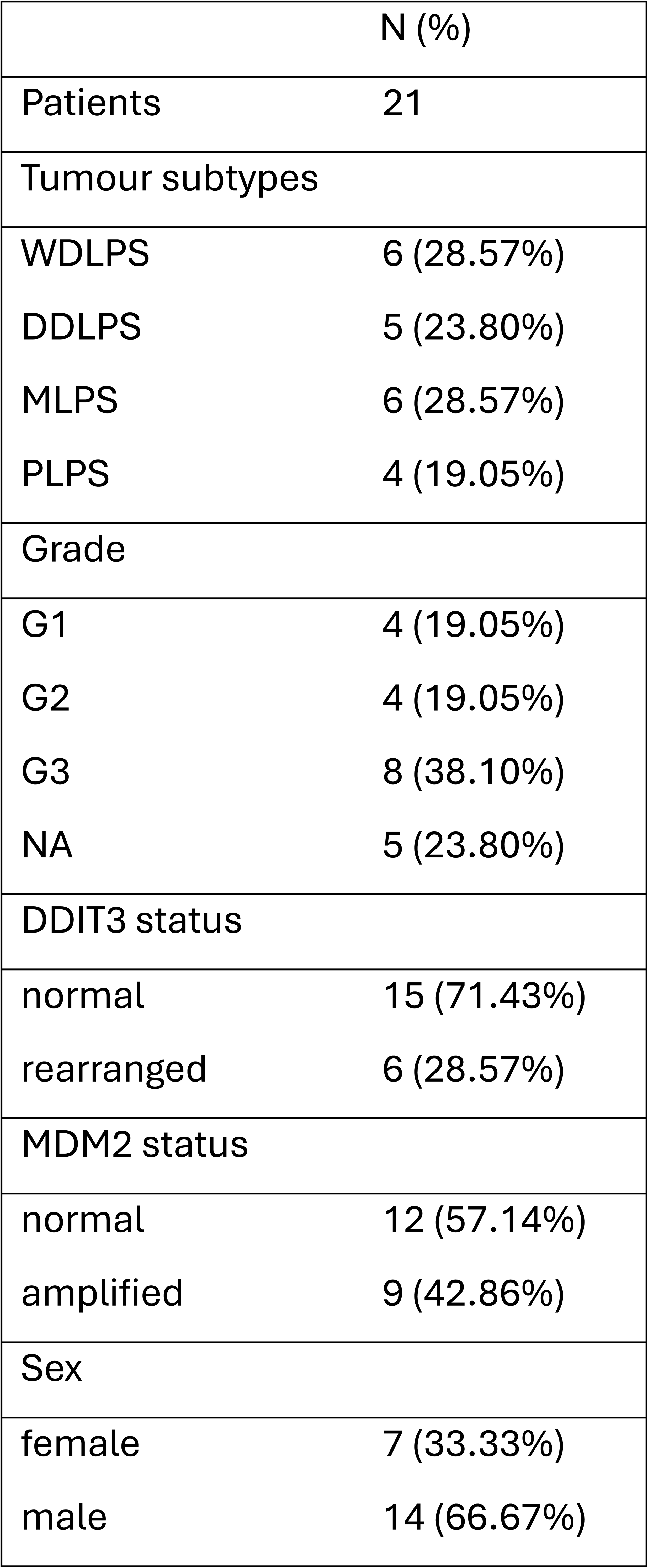
Clinicopathological characteristics of the liposarcoma patients included in the study.

|  | N (%) |
| --- | --- |
| Patients | 21 |
| Tumour subtypes |  |
| WDLPS | 6 (28.57%) |
| DDLPS | 5 (23.80%) |
| MLPS | 6 (28.57%) |
| PLPS | 4 (19.05%) |
| Grade |  |
| G1 | 4 (19.05%) |
| G2 | 4 (19.05%) |
| G3 | 8 (38.10%) |
| NA | 5 (23.80%) |
| DDIT3 status |  |
| normal | 15 (71.43%) |
| rearranged | 6 (28.57%) |
| MDM2 status |  |
| normal | 12 (57.14%) |
| amplified | 9 (42.86%) |
| Sex |  |
| female | 7 (33.33%) |
| male | 14 (66.67%) |

Each tumour, available as FFPE block, was processed to perform bulk RNA sequencing and to prepare TMAs then used for FISH and Immuno-FISH experiments (Fig 1B).

### Subtype-specific transcriptional signatures distinguish liposarcoma histotypes

Aimed to explore the transcriptomic landscape of liposarcoma we performed bulk RNA sequencing on paired tumour and normal tissues. After quality control assessment, the final analytic cohort used for the analyses comprised a total of 16 patients (4 with two independent specimens and 1 with 3 specimens), for a total of 22 samples.

Principal component analysis (PCA) revealed distinct transcriptional profiles across the cohort, with each liposarcoma subtype exhibiting a unique expression pattern that was also clearly separated from normal tissues (Fig. 1C). To identify subtype-specific transcriptional signatures in our cohort, we performed differential expression analysis using edgeR. For each liposarcoma subtype, gene expression was compared with that of the remaining three subtypes (subtype-versus-rest analysis). Genes significantly upregulated in each subtype (FDR<0.05 and log2FC ≥ 1) were ranked and the top 15 genes were selected for each subtype (Fig. 1D). In the DDLPS subtype, the top enriched genes included *COLCAC*, *TCF21*, *WT1* and *VEGFD1*, reflecting the activation of stromal/mesenchymal programs and extracellular matrix remodelling. Interestingly, this subtype exhibited a high expression of immunoglobulin-related transcripts, such as *IGH4*, *IGHM*, *IGHG3* and *JCHAIN,* potentially reflecting increased infiltration of B-cells or plasma cells within this subtype. *B4GALNT1*, a glycosyltransferase frequently overexpressed in cancer, was highly expressed in DDLPS. *B4GALNT1* is located on chromosome 12q, resulting in some cases co-amplified alongside the oncogenes *MDM2* and *CDK4* (*24*), which are located on the recurrently amplified 12q13-15 locus. The amplification of this chromosomal region is a hallmark genetic alteration for both DDLPS and WDLPS. In our cohort, increased *B4GALNT1* expression in DDLPS may reflect co-amplification of this gene together with *MDM2* and *CDK4*.

MLPS subtypes showed a distinct transcriptional profile enriched for adipocytic differentiation genes, such as *ADIPOQ* and *GIPR*. However, because FUS:DDIT3 is known to impair terminal adipogenic differentiation (25–27), the biological basis of this expression pattern more likely reflects intratumoural heterogeneity, variable tumour differentiation or contributions from non-neoplastic adipocytes, than genuine adipocytic maturation within tumour cells. Neuronal-associated genes such as *LBX1* and *DOCK3* were highly expressed, although their functional relevance to this subtype remains unclear. *CTAG2*, often expressed in sarcoma, were also highly expressed in MLPS, in line with what reported in other studies (28,29).

The PLPS transcriptional signature was characterised by genes involved in mesenchymal phenotype, including *RUN*X2 and *GREM1*, as well as extracellular matrix remodelling genes, including *COL11A1* and *LRRC15*. Additional genes involved in cancer progression, including *HS3ST3A1* and *SSX1,* were also enriched in this subtype, Although SSX1 gene is a well-established oncogenic driver in synovial sarcoma (30–32), to our knowledge, its association with PLPS has not been previously reported.

Among the top enriched genes in WDLPS were several genes involved in epithelial differentiation and cell adhesion, including *EPCAM*, *KRT7*, *CLDN4* and *MAL2*. Although these genes were identified as the top enriched in the WDLPS subtype, their expression was mainly driven by two patients, indicating an inter-patient-heterogeneity within this subtype rather than a subtype-wide molecular feature. Whether this represents entrapped normal epithelium or a genuine biological subpopulation warrants further investigations.

Comparing all liposarcoma samples to normal control group, 1590 differentially expressed genes (DEGs) were identified in the liposarcoma group, including 1016 upregulated, and 574 downregulated genes (FDR<0.05 and log2FC ≥ 1) (Table S1, Fig. 1E). Among the most significantly dysregulated genes were the proliferation-associated transcripts *CDK1*, *CDK2*, *CCNA2*, *KIF4A*, *RCC2* and *AKT1* (upregulated), the metal-homeostasis gene MT1X (downregulated), a pattern consistent with a proliferative transcriptional program.

Together, these findings demonstrate that each liposarcoma subtype exhibits a distinct transcriptional program, consistent with its underlying biology.

### Pathway enrichment reveals proliferative and inflammatory programs in liposarcoma

To characterise the molecular features of liposarcoma, we performed Gene Set Enrichment analysis (GSEA). Analysis of the “Hallmark” gene set revealed a significant enrichment of pathways associated with cancer progression and aggressiveness in liposarcoma, including cellular proliferation (E2F targets, G2M checkpoint), inflammation (IL6-JAK-STAT3 signalling, TNFα signalling, interferon response), and WNT-β catenin signalling (NES > 1 or NES < −1 and *P* < 0.05; Fig. 1F; Table S2). Conversely, genes related to myogenesis, adipogenesis, and oxidative phosphorylation were significantly enriched in normal tissues (NES > 1 or NES < − 1 and *P* < 0.05; Fig 1F; Table S2).

GSEA of the Gene Ontology Biological Process gene set collection revealed a positive enrichment of multiple gene sets related to mitosis and mitotic spindle organisation, suggesting enhanced mitotic activity and potentially reflecting centrosome amplification and chromosome instability in these tumours (Fig 1G; Table S2), in agreement with the upregulation of the cell-cycle genes (*CDK1*, *CDK2*, *CCNA2*) identified in the DEG analysis above.

Collectively, these findings show that liposarcoma subtypes are characterised by distinct transcriptional programs, whereas the comparison with adjacent normal tissues reveals the activation of proliferative and inflammatory pathways that define the liposarcoma transcriptome.

### Imprinted gene profiling of liposarcoma reveals distinct expression patterns across liposarcoma subtypes and identifies *PEG10* as a candidate biomarker

To further characterise the transcriptomic landscape of liposarcoma, we next examined the expression of imprinted genes which are frequently deregulated across multiple cancer types, and contribute to cancer initiation and progression (33–37). GSEA revealed a significant enrichment of imprinted genes in liposarcoma samples, compared to the normal counterparts (Fig. 2A). When splitting the imprinted genes in maternally and paternally expressed, only the paternally expressed genes were significantly enriched (Fig. 2A). This finding is in line with other studies describing the transcriptional upregulation of imprinted genes across cancer types (38).

**Figure 2.**
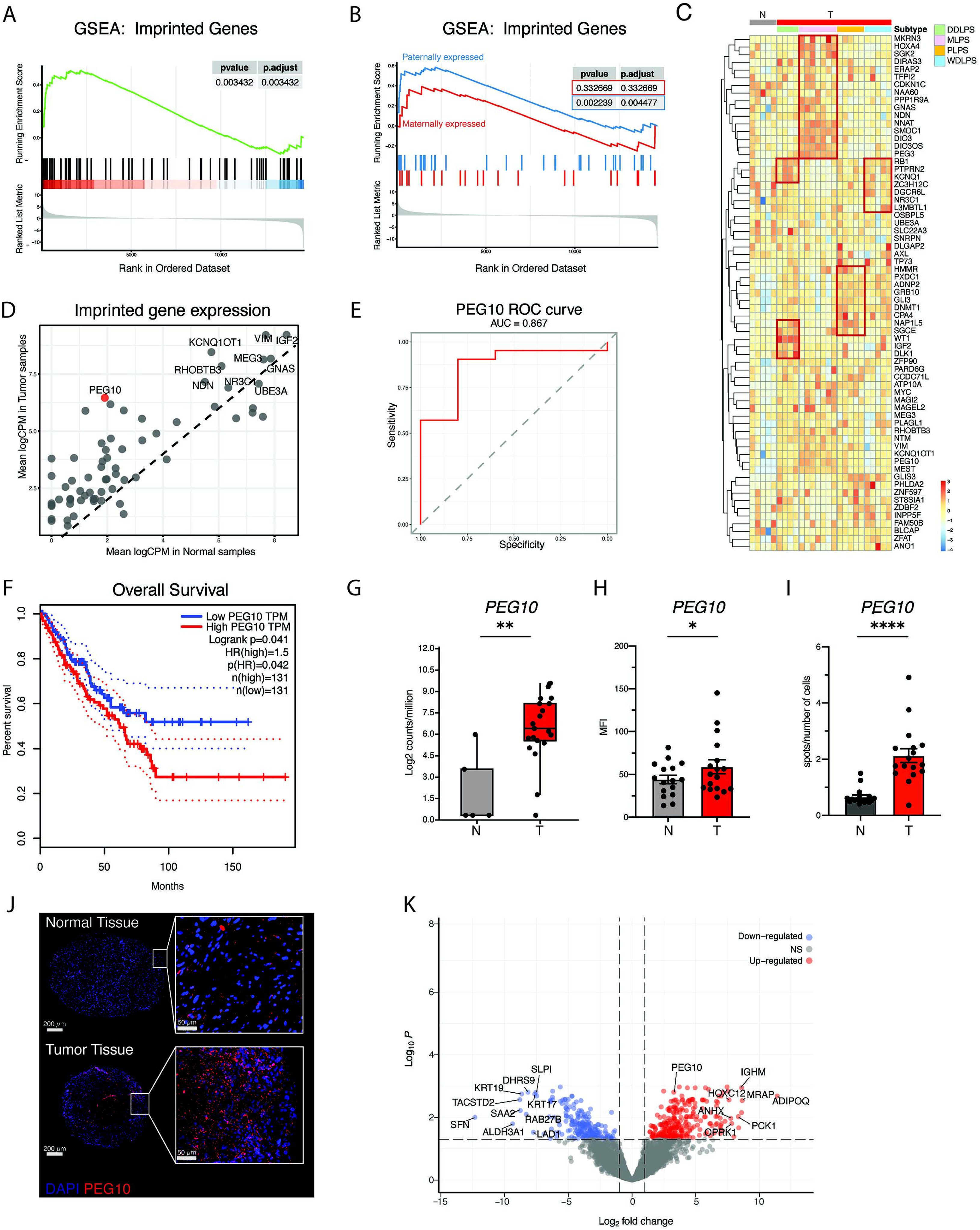
Analysis of imprinted gene expression in liposarcoma and adjacent normal tissues. A) GSEA enrichment plot showing significant enrichment of imprinted gene set in liposarcoma samples (n=22) compared with normal tissues (n=5). B) GSEA enrichment plot showing enrichment of both paternal and maternal expressed genes in liposarcoma samples, compared to normal tissues. C) Heatmap showing the expression of imprinted genes across normal tissues and the different liposarcoma subtypes. Expression values are displayed as row-wise z-scores, with samples grouped according to tissue type and histological subtype. D) Scatterplot comparing the expression of imprinted genes in normal and liposarcoma tissues. Each dot represents an imprinted gene, with expression in normal tissues shown on the x-axis and expression in liposarcoma tissues on the y-axis. The 10 most highly expressed genes are labelled and *PEG10* is highlighted in red. The dashed line indicates equal expression in tumour and normal tissues (x = y). E) ROC curve evaluating the sensitivity and specificity of PEG10 expression in discriminating normal and liposarcoma tissues in the discovery cohort. The area under the curve (AUC) is 0.867. F) Exploratory Kaplan-Meier overall survival in sarcoma patients from TCGA SARC cohort, stratified according to PEG10 expression (n=262, 131 per group). Survival distributions were compared using the log-rank test. TPM: transcripts per million; HR: hazard ratio. G) Box and whiskers showing the expression of PEG10 in normal tissues (n=5) and tumour tissues (n=22), measured by bulk RNA sequencing. *P* value from t-test (*: *p* ≤ 0.05) H) PEG10 expression in tumour (n=18) and normal tissues (n=18) as measured by FISH. Bars indicate the mean values ± SEM of the mean fluorescence intensity (MFI). Dots indicate value from individual patients. I) PEG10 expression in tumour (n=18) and normal tissues (n=18) as measured by FISH. Bars indicate the mean values ± SEM of the number of positive spots divided by the number of total cells. Dots indicate value from individual patients. J) Representative microscopy image showing PEG10 detection following FISH experiments, in normal tissues and tumour tissues. Nuclei were stained with DAPI and are shown in blue, while PEG10 is shown in red. K) Volcano plot showing the differentially expressed genes PEG10^high^ (n=11) and PEG10^low^ (n=11) (FDR<0.05 and log2FC ≥ 1).

Analysis of the imprinted gene expression revealed distinct expression patterns across the different liposarcoma subtypes (Fig. 2C). A subset of imprinted genes was specifically enriched in MLPS, including *MKRN3*, *HOXA4*, *SGK2*, *DIRAS3*, *ERAP2*, *TFPI2*, *CDKN1C*, *NAAC0*, *PPP1RSA*, *GNAS*, *NDN*, *NNAT*, *SMOC1*, *DIO3*, *DIO3OS*, *PEG3.* DDLPS and WDLPS shared a high expression of *RB1*, *PTPRN2*, *KCNQ1*. DDLPS was additionally characterised by the high expression of *NAP1L5*, *SGCE*, *WT1*, *IGF2*, *DLK1*; while WDLPS showed a particularly high expression of ZC3H12C, DGCR6L, NR3C1, L3MBTL1. PLPS showed a high expression of *HMMR*, *PXDC1*, *ADNP2*, *GRB10*, *GLI3*, *DNMT1*, *CPA4*, *NAP1L5*, *SGCE*.

We then ranked the imprinted genes by their expression level in tumour samples and identified the most highly expressed genes overall (Fig. S1A). We next sought to identify which imprinted genes most effectively discriminate tumour samples from normal tissues and identified *PEG10* as the top discriminating gene, showing high expression in tumour samples and low expression in normal tissues (Fig. 2D). Although *PEG10* ranked only tenth among imprinted genes by absolute expression level in tumour samples (Fig. S1A), the nine genes above (*IGF2*, *VIM*, *KCNQ1OT1*, *GNAS*, *MEG3*, *RHOBTB3*, *NDN*, *UBE3A*, *NR3C1*) were also highly expressed in normal tissues (Fig. 2D, Fig. S1A), making them poor discriminators despite their high absolute expression. The analysis of the Receiver Operating Characteristic (ROC) analysis showed good discriminatory performance for PEG10 in distinguishing tumor from normal tissues within the discovery cohort (AUC=0.867; Fig. 2E). *PEG10* also appeared as one of the upregulated genes in the tumour-versus-normal DE analysis (Fig. 1E).

Interestingly, in an exploratory analysis of the TCGA-SARC cohort (39), using GEPIA (40), we found that patients with high *PEG10* expression had significantly poorer overall survival, than those with low *PEG10* expression (log-rank P=0.041, HR=1.5, p(HR)=0.042; n=131 per group; Fig. 2F), with a non-significant trend towards a shorter disease-free survival (log-rank P=0.16, HR=1.3, p(HR)=0.16, n=131 per group; Fig. S1B). Based on these results and given that *PEG10* has previously been associated with disease progression in other cancer types (41), we next sought to further characterise the significance of *PEG10* upregulation in our liposarcoma cohort.

RNA-seq analysis revealed that *PEG10* expression is significantly higher in the tumour microenvironment compared to the normal adjacent compartment (Fig. 1E, Fig 2G). We additionally confirmed this result by measuring *PEG10* expression via RT-qPCR (Fig. S1C).

We further validated this at tissue level by FISH experiments, to specifically detect *PEG10*. The analysis was performed on TMAs, containing samples from liposarcoma patients (n=21), each represented by 2 tissue cores. The TMAs comprised both the tumour tissues and the corresponding normal adjacent tissues. The analysis confirmed the overexpression of *PEG10* in tumour tissues, compared to normal tissues (Fig. 2H-J), consistent with our RNA-sequencing and qPCR results.

In order to gain insights into the role of PEG10 in liposarcoma, we then stratified the 22 tumor specimens into *PEG10*^high^ (n=11) and *PEG10*^low^ (n=11), based on the median *PEG10* expression value from the RNA-seq. Interestingly, most of the MLPS samples (n=7/8, coming from 4 patients, 2 with two independent specimens, and 1 with three independent specimens) were assigned to the *PEG1*0^high^, whereas the other subtypes were distributed across the two groups.

Comparing *PEG10*^high^ and *PEG10*^low^ groups, we identified 470 Differentially Expressed Genes (DEGs), among which 264 were upregulated, and 206 were downregulated in the PEG10^high^ group (FDR<0.05 and log2FC ≥ 1) (Fig. 2K; Table S3), including upregulation of *ADIPOQ* and *PCK1* in the *PEG10*^high^ group, and lower expression of several epithelial markers (*KRT17*, *KRT1S*, *TACSTD2*, *SFN*) in the same group.

We then performed GSEA-Preranked analysis to identify key differences in transcriptional pathways between the two groups. Analysis of the Hallmark gene collection revealed that genes involved in adipogenesis were enriched in *PEG10*^high^ tumours, whereas E2F targets and MYC targets were enriched in *PEG10*^low^ group (Fig. S1E, Table S4), together with genes involved in MTORC1 and P53 signalling. Genes involved in ROS pathway were enriched in *PEG10*^low^ group as well.

Analysis of the Gene Ontology “Biological process” additionally revealed that PEG10^high^ tumours are significantly enriched with genes regulating the metabolism of lipids, including triglycerides and fatty acids (Fig. S1E, Table S4). These results suggest that *PEG10* high expression associates with metabolic shift toward lipid metabolism, while low *PEG10* expression associates with a more stress-responsive phenotype. The concomitant enrichment of ROS and p53 signalling and E2F/MYC targets in *PEG10*^low^ samples may further suggest an increased cellular stress in this group. This analysis additionally revealed that cilia assembly and microtubule-based transport are enriched in *PEG10*^low^ group, suggesting alterations in cytoskeletal organisation and intracellular trafficking.

Taken together, these results demonstrate widespread dysregulation of imprinted genes in liposarcoma, with many imprinted genes upregulated compared to normal tissues. Among the imprinted genes, *PEG10* emerged as a promising candidate tumour biomarker, warranting further validation in independent, subtype-stratified cohorts.

### *PEG10* is expressed in all liposarcoma subtypes and marks high-grade tumours

To further assess the clinical relevance of PEG10 expression in liposarcoma, we investigated its association with histological subtypes, tumour grade and MDM2 amplification.

We first examined whether *PEG10* could discriminate between the different liposarcoma subtypes. In our FISH experiments, the highest expression for *PEG10* was detected in PLPS samples (average MFI: 82.45), followed by MLPS (average MFI: 63.53), while lower levels were measured for WDLPS (average MFI: 55.78) and DDLPS (MFI: 28.87) (Fig. 3A-C), with a similar pattern observed using RNA-FISH spot counts (Fig. 3B). None of these differences reached statistical significance (Fig. 3A-B), suggesting that *PEG10* expression is not restricted to a specific liposarcoma subtype.

**Figure 3.**
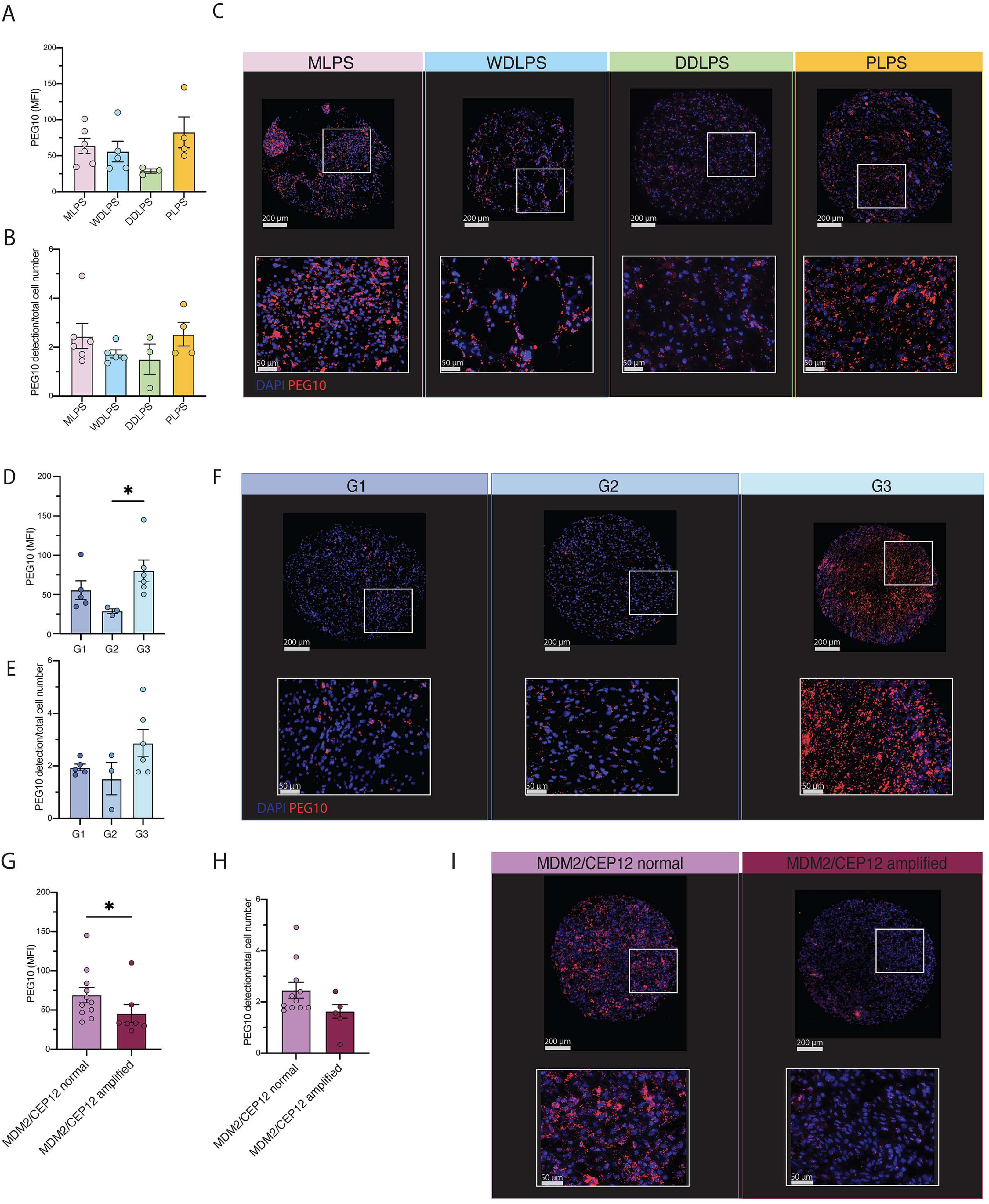
PEG10 expression across liposarcoma subtype, histological grades and MDM2 amplification groups. A) Bar plot showing the mean intensity fluorescence measured for PEG10 in the different tumour subtypes (n=18, MLPS: 6, WDLPS: 5, DDLPS: 3, PLPS: 4), by FISH experiments. Kruskal-Wallis test followed by Dunn’s multiple comparison test did not show any significant difference between groups. B) Bar plot showing number of PEG10 detections, divided by the total number of cells, counted following FISH experiments, in the different tumour subtypes (n=18, MLPS: 6, WDLPS: 5, DDLPS: 3, PLPS: 4). C) Representative microscopy images showing PEG10 detection following FISH experiments, in the four liposarcoma subtypes D) Bar plot showing the mean intensity fluorescence measured for PEG10, following FISH experiments, in grade 1 (G1, n=5), grade 2 (G2, n=3) and grade 3 (G3, n=6) liposarcoma. *P* values from one way Kruskal-Wallis followed by Dunn’s multiple comparison test (*: p ≤ 0.05). E) Bar plot showing number of PEG10 detections, divided by the total number of cells, counted following FISH experiments, in G1 (n=5), G2 (n=3) and G3 (n=6) liposarcoma. F) Representative microscopy images showing PEG10 detection following FISH experiments, in G1, G2 and G3 liposarcoma. G) Bar plot showing the mean intensity fluorescence measured for PEG10, following FISH experiments, in MDM2/CEP12 normal (n=11) and MDM2/CEP12 amplified (n=7) tumours. *P* value from Mann-Whitney test (*: *p* ≤ 0.05).H) Bar plot showing number of PEG10 detections, divided by the total number of cells, counted following FISH experiments, in MDM2/CEP12 normal (n=11) and MDM2/CEP12 amplified (n=7) tumours. I) Representative microscopy images showing PEG10 detection following FISH experiments, in liposarcoma harbouring or lacking MDM2 amplification. A, B, D, E, G, H) Bars indicate the mean values ± SEM. Dots indicate value from individual patients.

We next evaluated whether *PEG10* was associated with tumour grade. PEG10 expression was significantly higher in G3 tumours, compared with G2 (p=0.0147, Fig. 3D), with a numerically higher but not significant difference relative to G1 (Fig. 3D-F). A consistent, non-significant trend was observed for the spot-count metric (Fig. 3E). This grade association was directionally supported by RT-qPCR comparing G1 and G3 tumours (Fig. S2A). Together these results support an association between *PEG10* expression and tumour aggressiveness.

DDLPS and WDLPS are characterised by a systematic amplification of MDM2 oncogene, which is routinely evaluated in clinical practice, for the diagnosis of liposarcoma. We therefore investigated the relationship between *PEG10* expression and MDM2 amplification. We observed a higher PEG10 expression in MDM2/CEP12 normal tumours (MLPS and PLPS; average MFI: 68.88) compared to MDM2/CEP12 amplified tumours (WDLPS and DDLPS; average MFI: 45.54, p=0.0346) (Fig. 3G-I).

Together, these results suggest that PEG10 represents a clinically relevant biomarker in liposarcoma, associated with tumour aggressiveness rather than histological subtypes. *PEG10* expression is broadly detectable across subtypes, increases with tumour grade and associated with MDM2 amplification status.

### *PEG10* expression within the tumour lesion is predominantly localized to malignant and proliferating cells

After confirming the upregulation of *PEG10* expression in liposarcoma, compared with normal tissues, we aimed to determine whether *PEG10* is expressed by tumour cells and/or other cell populations. Therefore, we performed FISH experiments for *PEG10*, combined with immunofluorescence staining (Immuno-FISH) for p53 and vimentin, to assess its co-expression with tumour-associated cellular markers.

Across all the samples, the majority of cells co-expressed *PEG10* and p53 (average 64% of total cells), while 35% of cells expressed only p53 (Fig. 4A-B). Interestingly, cells expressing *PEG10* alone were extremely rare (with an average of 0.07% of total cells) (Fig. 4A-B). These expression patterns were consistent across subtypes (Supplementary Fig. 3A). Specifically, the frequency of *PEG10*+p53+ cells was 68.86% in WDLPS (n=5), 65.17% (n=5) in MLPS, 79.72% (n=4) in PLPS. For DDLPS subtype (n=4) the frequency of double positive cells in this tumour was 42.09% (Supplementary Fig. 3A).

**Figure 4.**
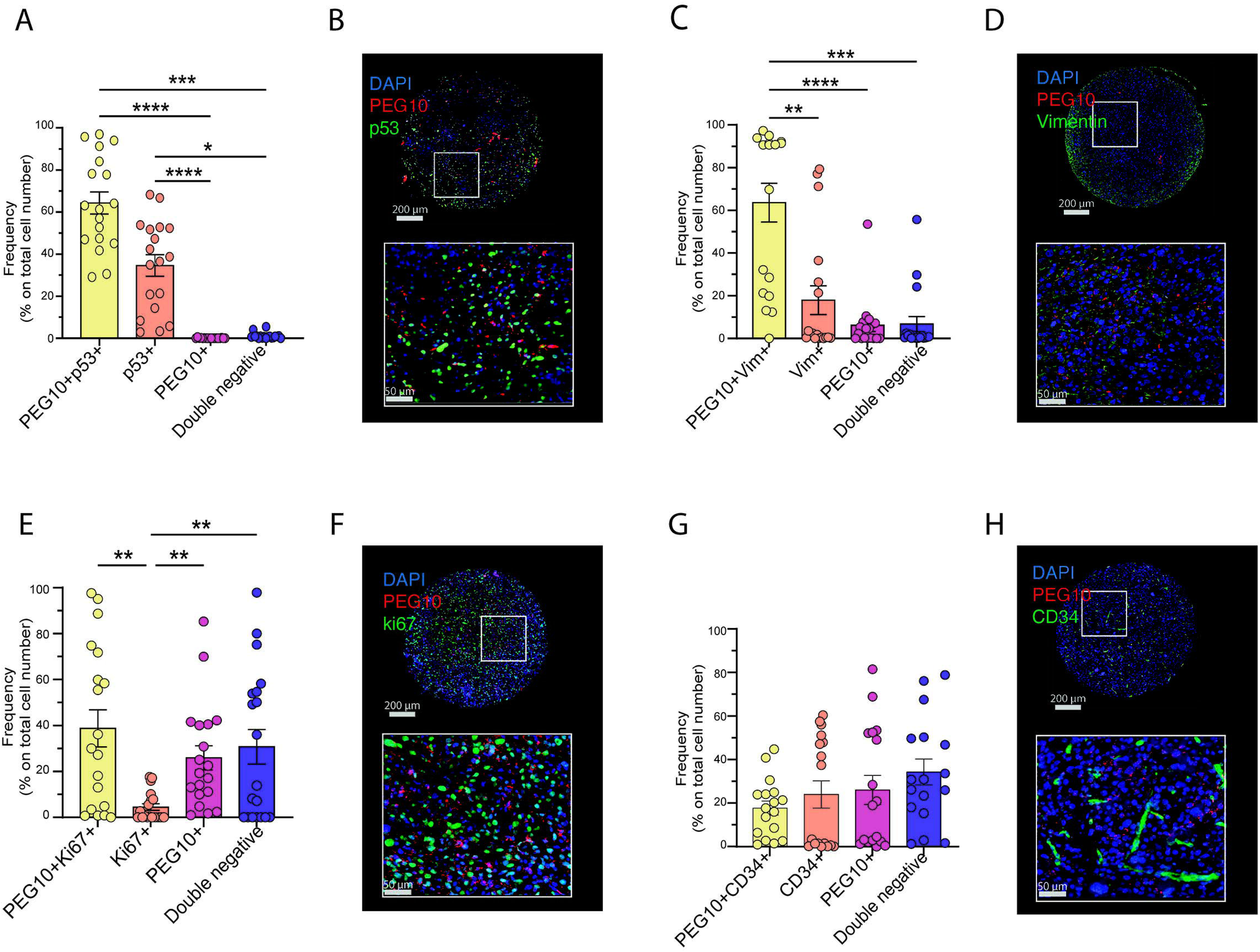
PEG10 is consistently expressed by tumour and proliferating cells. A) Bar plot showing the frequency of cells expressing PEG10, p53, both markers or neither in liposarcoma samples (n=18). *P* values from Friedman test. B) Representative Immuno-FISH images showing the co-expression patterns of PEG10 (red) and p53 (green). C) Bar plot showing the frequency of cells expressing PEG10, vimentin, both markers or neither in liposarcoma samples (n=18). *P* values from Friedman test. D) Representative Immuno-FISH images showing the co-expression patterns of PEG10 (red) and vimentin (Vim) (green), E) Bar plot showing the frequency of cells expressing PEG10, Ki67, both markers or neither in liposarcoma samples (n=19). *P* values from Friedman test. F) Representative Immuno-FISH images showing the co-expression patterns of PEG10 (red) and ki67 (green). G) Bar plot showing the frequency of cells expressing PEG10, CD34, both markers or neither in liposarcoma samples (n=18). No significant difference between groups was observed following Friedman test. H) Representative Immuno-FISH images showing the co-expression patterns of PEG10 (red) and CD34 (green). B, D, F, H) Nuclei were stained with DAPI and are shown in blue. PEG10 was detected using an RNA probe, while the other markers were detected via immunofluorescence staining. A, C, E, G) Bars indicate the mean values ± SEM. Dots indicate value from individual patients. p ≤ 0.05; **: p ≤ 0.01; ***: p ≤ 0.001; ****: p ≤ 0.0001

Consistent with these findings, most of the analysed cells co-expressed both *PEG10* and Vimentin (63.63%), while 17.95% of cells were double negative (Fig. 4C-D). The frequency of cells positive only to *PEG10* and only to Vimentin was 6.24% and 6.76% respectively.

We observed that the frequency of PEG10+Vimentin+ cells was particularly high in MLPS (83.81%, n=5) (Supplementary Fig. 3B), and lower in WDLPS (59.34%, n=5), DDLPS (56.21%, n=4) and PLPS (46.52%, n=4) (Supplementary Fig. 3B).

Together, these results showed that PEG10 expression largely overlapped with p53- and vimentin-positive cell populations, supporting its preferential expression in cells with a tumour-associated phenotype.

We next investigated whether *PEG10* expression is associated with tumour cell proliferation. To address this, we performed Immuno-FISH experiments to analyse the expression of *PEG10* and Ki67, a well-established marker of proliferation. Overall, 38.8% of cells within the tumour samples (n=19) co-expressed Ki67 and *PEG10* (Fig. 4E-F). In contrast, only 4.48% of cells was positive for Ki67 and negative for *PEG10*, whereas 25.95% of cells expressed only PEG10 (Fig. 4E-F). The average percentage of *PEG10*+Ki67+ cells did not significantly differ among subtypes: 34.38% in MLPS (n=6), 45.66% in WDLPS (n=5), 51.77% in DDLPS (n=4), 23.87% in PLPS (n=4) (Supplementary Fig. 3C). Therefore, these results indicate that most of the proliferating cells also express *PEG10*.

Finally, we evaluated PEG10 co-expression with CD34, a marker of adipose-derived progenitor cells and vascular endothelium (42–44). We observed no significant association between *PEG10* and CD34: around 20% of cells co-expressed PEG10 and CD34 and a comparable number of cells expressed only one of them (Fig 4G-H, Supplementary Figure 3D). This finding indicates that PEG10 expression is not preferentially associated with CD34-positive stromal or vascular populations, as well as differentiated adipocytes.

### Single-cell RNA sequencing analysis of DDLPS tumour microenvironment confirms that *PEG10* marks aggressive tumour-cell states and associates with a lower myeloid-cell fraction

Our data suggested a potential relationship between *PEG10* and aggressiveness of liposarcoma. We next sought to further validate our findings at single-cell resolution in an external dataset. Specifically, we used data obtained from the Gene Expression Omnibus (GEO) under accession number GSE221493 (4). The dataset comprises samples from 11 primary untreated DDLPS patients. For each tumour, well-differentiated (WD) and dedifferentiated (DD) components were histologically distinguished by pathologists during the macroscopic examination of the fresh surgical specimen just after surgery and simultaneously submitted to sequencing. The diagnosis of DDLPS had been established in all cases on pre-operative biopsies and later confirmed on the surgical specimen.

We reproduced the original clustering analysis which identified 10 main cell populations (Fig. 5A). Gene expression profiles and expression of canonical markers enabled grouping these clusters into 10 main cellular subfamilies, including tumour cells, lymphoid cells, myeloid cells, cycling cells, adipocytes, mesothelial cells, mast cells, endothelial cells, pericytes, and red blood cells (RBC) (Fig. 5A-C). The analysis confirmed that *PEG10* expression is restricted to tumour cells and cycling cells (Fig. 5B), with no detectable expression in immune cells and stromal cells (Fig. 5B), consistent with our own cohort.

**Figure 5.**
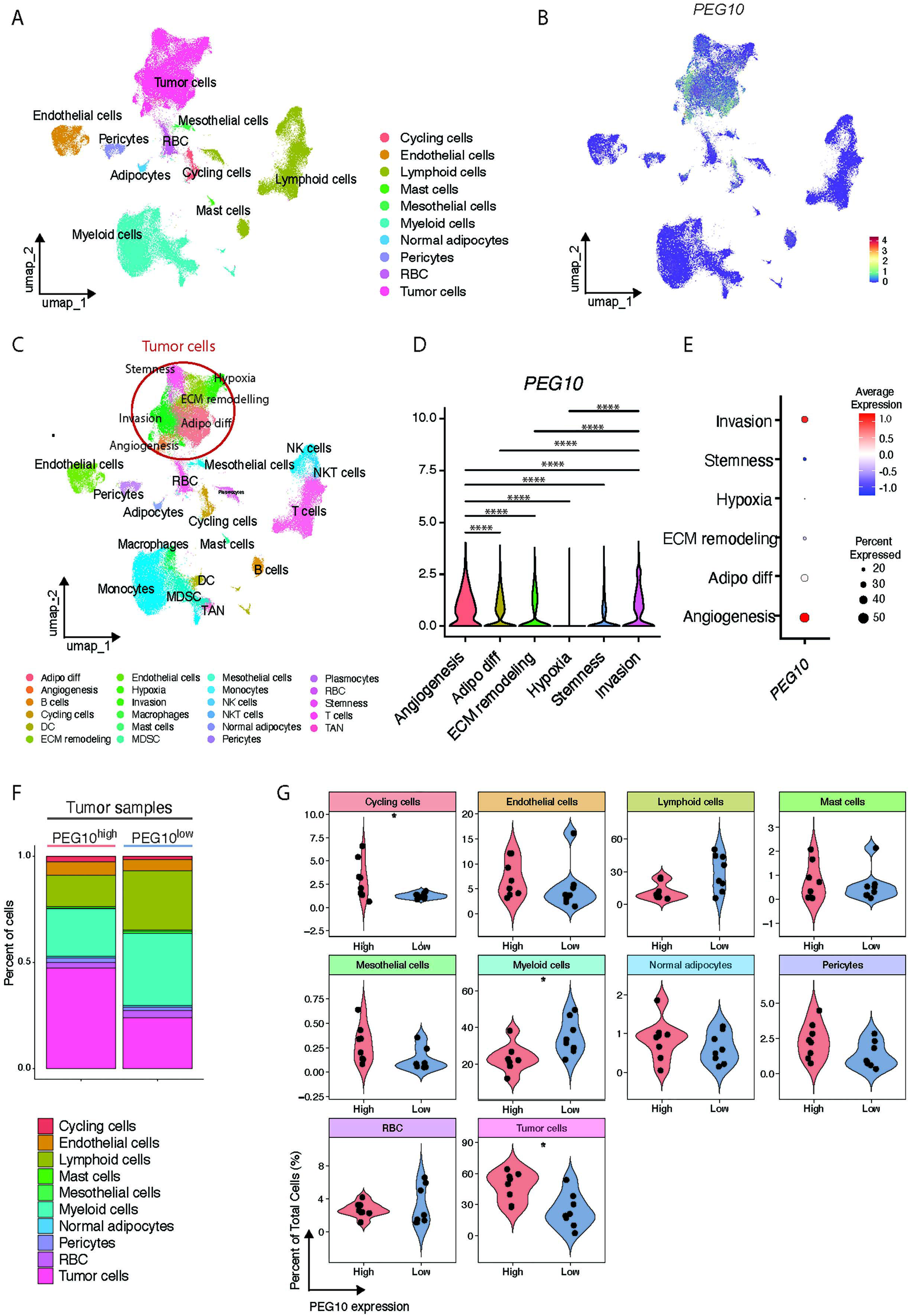
Single-cell distribution of PEG10 expression in an external DDLPS dataset. A) Cellular subsets as identified using scRNA seq data and visualised on weighted-nearest neighbour (wnn) UMAP coordinates. B) UMAP showing the expression of PEG10. C) UMAP showing the cellular subsets as identified by subclustering the populations shown in A. D) Violin plots showing the expression of PEG10 in tumour cell subsets. *P* values are from Wilcoxon test, ****: p ≤ 0.0001. E) Dot plot showing the frequency and average expression of PEG10 in the tumour cell subset. F) Bar plot showing the frequency of cellular subsets, as identified in A, in PEG10^high^ and PEG10^low^ tumours. G) Violin plots showing the frequency of cellular subsets, as identified in A and shown in F, and the statistical significance, when present. Dots indicate values from individual patients.

Tumour cells were further subclustered in six subpopulations, based on the expression of genes associated to stemness, invasion, hypoxia, ECM remodelling, adipocyte differentiation and angiogenesis, as described by Gruel and colleagues (Fig. 5C, Supplementary Table 5) (4). Focusing on the tumour population, we observed that PEG10 is mainly expressed by tumour cells showing the angiogenesis signature (mean expression: 2.027), and tumour cells showing an invasive signature (mean expression 1.902) (Fig 5 D-E). PEG10 expression in the angiogenesis cluster was significantly higher than all the other subtypes (Fig 5D), while its expression in the invasion subcluster was significantly higher than the adipocyte differentiation (mean expression: 1.36), ECM remodelling (mean expression: 1.15), hypoxia (mean expression: 0.899) and stemness (mean expression: 0.807) clusters (Fig 5D). We next asked whether the same functional signatures correlated with *PEG10* in our cohort, using RNA-seq data. *PEG10* significantly correlated with hypoxia (R=0.76, *P*=6.2×10^−6^), invasion (R=0.61, *P*=0.00097) and ECM remodelling (R=0.58, *P*=0.0021) signatures. No correlation was observed between *PEG10* and adipocyte differentiation, angiogenesis or stemness signatures (Supplementary Figure 4A). Notably, although angiogenesis was the single-cell subcluster with the highest PEG10 expression, this association did not replicate at the bulk-tissue level, likely reflecting the dilution of a cell-intrinsic signal by the transcriptomes of other cell populations present in bulk tumour samples; this apparent discrepancy between single-cell and bulk-level associations should be interpreted considering this limitation.

We then used the single-cell RNAseq dataset from Gruel and colleagues to evaluate differences in the frequency of immune cell populations between *PEG10*^high^ and *PEG10*^low^ tumours (as stratified according to the median expression level). Interestingly, we observed a tendency towards a higher frequency of lymphoid cells in *PEG10*^low^ tumours (Fig 5 F-G). The frequency of myeloid cells was significantly higher in *PEG10*^low^ tumours (Fig 5 F-G). When evaluating other cell populations, we found a significantly higher frequency of cycling cells in *PEG10*^high^ tumours (Fig 5 F-G), while no significant differences were observed comparing the other populations (Fig 5 F-G). Because these values represent proportions of total cells, the higher tumour-cell fractions in PEG10^high^ samples may itself partly account for the relatively lower myeloid-cell proportion in this group and further studies are required to clarify whether specific populations are selectively depleted.

This dataset represents an important resource for studying liposarcoma, particularly given the rarity of this tumour, however it is limited to DDLPS and does not include the other liposarcoma subtypes present in our cohort of patients. Collectively, these results support our cohort-level findings: PEG10 is predominantly expressed by tumour cells and proliferating cells, with the highest expression in tumour-cell states associated with invasion and angiogenesis at single-cell resolution, and its expression correlates with invasion, hypoxia and ECM remodelling signatures at the bulk-tissue level. Together these findings underline an association between PEG10 expression and aggressive tumour features.

## Discussion

Liposarcoma is a rare tumour, whose clinical management remains highly challenging. Indeed, the misdiagnosis of this tumour occurs in 30% of cases and the therapeutic options remain limited (1,16,25,45). Therefore, the identification of new therapeutic targets and diagnostic or prognostic biomarkers is urgent.

In our study we analysed the transcriptome of 16 liposarcoma tumours (22 specimens), including all the main four subtypes, ranging from low to high grade, within a broader clinicopathological cohort of 21 patients. Transcriptomics analysis revealed the dysregulation of several pathways involved in cell cycle regulation, indicating an enhanced proliferation rate in tumours compared to normal tissue, as expected. Pathways associated with tumour aggressiveness were enhanced in tumour samples, including cellular proliferation (E2F targets, G2M checkpoint), inflammation (IL6-JAK-STAT3 signalling, TNFα signalling, interferon response), and WNT-β catenin signalling.

To further characterise the liposarcoma transcriptome, we investigated the expression profile of imprinted genes. Dysregulation of imprinted gene expression is represents a hallmark of several malignancies, referred as imprinting disorders (46). Genomic imprinting mediates the expression of specific genes in a parent of origin manner and their expression is often regulated by coordinated epigenetic mechanisms (47,48). Disruption of genomic imprinting causes abnormal expression by either activating an allele that is normally silenced or silencing an allele that is normally expressed (48). In the context of cancer, loss of imprinting has been widely reported (48–52).

Our analysis revealed a broad enrichment of imprinted genes in all liposarcoma subtypes compared with normal tissues, involving mainly the paternally expressed genes. We also identified distinct subtype-specific expression patterns of imprinted genes across the different liposarcoma subtypes. These transcriptional signatures warrant further validation and may ultimately contribute to improved molecular classification of liposarcoma. Moreover, elucidating the functional role of individual imprinted genes could provide valuable mechanistic insights. To the best of our knowledge, our study is the first one providing a systemic characterisation of the expression landscape of imprinted genes in liposarcoma. However, in our study we did not determine whether the observed expression levels reflect loss of imprinting, altered DNA methylation or other regulatory mechanisms. Integrative methylome and allele-specific expression analyses will therefore be required in further studies to clarify the epigenetic basis of imprintome alterations.

Among the imprinted genes, the paternally expressed gene 10, *PEG10*, showed the strongest ability to discriminate tumour from normal tissue, highlighting its potential as a tumour-associated biomarker. Although PEG10 has previously been linked to cancer progression in other tumour types a detailed characterisation of its expression in liposarcoma was lacking. Therefore, we decided to further characterise *PEG10* expression in our cohort in depth, evaluating its potential as biomarker and/or therapeutic target.

*PEG10* gene plays an important role in placental development (55,56) and has more recently been implicated in several neurological disorders, including Angelman syndrome, a severe neurodevelopmental disorder, and amyotrophic lateral sclerosis (56). In the context of adipocyte differentiation PEG10 was found to be upregulated in the early stages of induced differentiation of 3T3-L1 fibroblastic cells (57).

Previous studies have reported that *PEG10* is upregulated in several tumour types, including hepatocellular carcinoma, breast cancer, pancreatic cancer, prostate cancer and colon cancer (41), while loss of *PEG10* imprinting has been observed in breast cancer (50), and high-risk B-cell chronic lymphocytic leukemia (58).

We found that *PEG10* is expressed mainly by tumour cells and by proliferating cells. In our cohort, *PEG10* was over-expressed across all liposarcoma subtypes, with no significant differences among them, as measured by Immuno-FISH experiments. However, when patients were stratified by disease grade, we observed that *PEG10* expression was significantly higher in high-grade tumours. This finding suggests that *PEG10* expression is enhanced in more aggressive tumours. Therefore, aimed to validate our findings in an additional independent cohort, we used single-cell RNA sequencing dataset (4). This analysis not only confirmed our Immuno-FISH results but also allowed for a deeper characterisation of *PEG10* expression: *PEG10* expression is restricted to tumour and proliferating cells, while other populations, including stromal or immune cells, do not show any detectable expression. Additionally, *PEG10* expression was highest in tumour cells displaying an angiogenesis- or invasion-associated signature.

We further found that *PEG10*^high^ tumours, as defined in this single-cell dataset, were characterised by lower myeloid-cell fraction and non-significant trend toward a lower lymphoid-cell fraction, alongside a higher tumour-cell fraction. A relatively higher immune infiltration in *PEG10*^low^ group may be consistent with the enrichment of pathways involved in cellular stress that we observed in our bulk RNA-seq analysis (ROS, p53, pathways). Whether reduced immune infiltration in PEG10^high^ tumours contributes to or merely accompanies increased aggressiveness in this subgroup remains to be demonstrated. Whether PEG10 directly contributes to immune exclusion or is simply a marker of this phenotype needs to be determined in further studies.

Analysis of TCGA SARC dataset further explored the clinical relevance of *PEG10* overexpression, revealing that high *PEG10* expression was significantly associated with a shorter overall survival, and with a trend towards shorter disease-free survival. However, the TCGA-SARC cohort comprises multiple soft-tissue sarcoma histological subtypes in addition to liposarcoma, indicating that the observed association reflects the broader sarcoma cohort rather than liposarcoma specifically.

Taken together, our results strongly suggest that a high expression of *PEG10* is a feature of high aggressiveness of liposarcoma. Indeed, we showed that *PEG10* expression is higher in high-grade tumours, it associates with worse overall survival of patients, and correlates with hypoxia, extracellular matrix remodelling and invasion signatures. *PEG10*^high^ tumours additionally show a reduced immune infiltration.

Other studies reported that *PEG10* expression is associated with poor prognosis of cancer patients. Ge and colleagues performed a meta-analysis to evaluate any correlation between PEG10 and clinicopathological characteristics of patients with solid tumours, within an Asian population (59), finding that PEG10 expression was associated with poor prognosis, higher risk of solid tumour incidence, lower degree of differentiation, increased lymph node metastases and advanced TNM stage (59). In another study, Bang and coworkers analysed PEG10 protein expression in tumour tissues from 218 hepatocellular carcinoma patients undergoing curative resection (60). They show that PEG10 expression is significantly associated with shorter overall survival and progression-free survival and was an independent predictor of shorter progression-free survival (60). Another study demonstrated the role of PEG10 in promoting tumour cell proliferation and invasion in bladder cancer, both *in vitro* and *in vivo (C1)*. The same study additionally showed that PEG10 expression is related to acquired resistance to chemotherapy (61). These previous studies, together with our findings, highlight a potential involvement of PEG10 in tumour progression.

Our RNA-seq analysis additionally revealed that *PEG10*^high^ tumours are characterised by enhanced lipid uptake and metabolism, alongside increased adipogenesis. However, these findings should be interpreted with caution, as the *PEG10*^high^ group was predominantly composed of MLPS (7/11 tumours), whereas the *PEG10*^low^ group was more balanced.

Metabolic reprogramming affecting lipid metabolism has been linked to cancer progression. Indeed, enhanced lipid uptake and biosynthesis are considered to promote tumour proliferation, providing both biosynthetic precursors for membrane formation and metabolic substrates for energy production. Therefore, the modulation of lipid metabolism has been proposed as new potential therapeutic approach for cancer treatment (62). Larger subtype-balanced cohorts will be required to fully uncover the biological effects directly attributable to PEG10, independent of subtype composition, and to elucidate whether its expression is related to metabolic reprogramming in liposarcoma.

We also found that *PEG10* expression is higher in MDM2 non-amplified tumours compared to MDM2-amplified tumours. Although here we cannot yet explain the molecular mechanism behind this finding, the expression of PEG10 in MDM2 non-amplified tumours could serve as additional diagnostic marker. Indeed, the assessment of MDM2 amplification is routine clinical practice, to diagnose WDLPS and DDLPS. The MDM2 non-amplified liposarcomas include all the other subtypes, which are misdiagnosed in a considerable number of cases. In this context, the potential role of PEG10 as diagnostic tool warrants further investigation and validation.

We acknowledge that our study has some limitations. First, the small number of patients, mainly due to the rarity of liposarcoma, which makes the collection of a great number of cases particularly challenging. Second, the PEG10-high/low stratification was skewed toward MLPS, and the resulting transcriptional and pathway differences may partly reflect subtype composition rather than a PEG10-specific effect. Despite these limitations, the results obtained contribute meaningfully to the understanding of an understudied disease, providing valuable insights that warrant further investigations in future studies. The role of *PEG10* in liposarcoma microenvironment still needs to be unravelled and its potential as therapeutic target needs to be investigated.

Collectively, our findings identify genomic-imprinting dysregulation as an underexplored feature of liposarcoma biology and position PEG10 as a promising candidate biomarker associated with aggressive tumor features. These observations provide a foundation for future functional studies and for the development of molecularly informed diagnostic and prognostic strategies.

## Conclusions

In conclusion, our study provides a transcriptomic dataset of liposarcoma and paired adjacent normal tissue that may be used also by others for additional analyses. This resource is particularly valuable, considering the rarity of this tumour. We additionally provide a comprehensive characterisation of the imprinted gene expression landscape in liposarcoma, which will be useful for future investigations into the role of genomic imprinting in liposarcoma. Among the imprinted genes, we found that PEG10 showed good discriminatory performance between liposarcoma and adjacent normal tissues in the discovery cohort. Furthermore, our study revealed that PEG10 marks high-grade liposarcoma and it is associated with aggressive tumour features. Altogether, our results support further evaluation of PEG10 as a candidate biomarker in liposarcoma.

## Supporting information

Table S2_GOBP

Table S4_GOBP

## List of abbreviations

AUC: area under the curve
BCV: biological coefficient of variation
CPM: counts per million
DD: dedifferentiated
DDLPS: dedifferentiated liposarcomas
DEG: differentially expressed genes
ECM: extracellular matrix
FDR: false discovery rate
FFPE: formalin-fixed paraffin embedded
FISH: fluorescence in situ hybridization
G1: grade 1
G2: grade 2
G3: grade 3
GEO: gene expression omnibus
GLM: generalized linear model
GSEA: gene set enrichment analysis
HCE: haematoxylin and eosin staining
HR: hazard ratio
MFI: mean fluorescence intensity
MLPS: myxoid liposarcomas
MSigDB: molecular signatures database
NES: normalised enrichment score
NBF: neutral buffered formalin
PCA: principal component analysis
PEG10: paternally expressed gene 10
PLPS: pleomorphic liposarcomas
RNA-seq: RNA sequencing
ROC: receiver operating characteristic
RT-qPCR: Real-Time Reverse Transcription Quantitative Polymerase Chain Reaction
scRNA-seq: single-cell RNA sequencing
ssGSEA: single-sample Gene Set Enrichment Analysis
TCGA: The Cancer Genome Atlas
TCGA-SARC: TCGA sarcoma cohort
TMA: tissue micro array
TPM: transcripts per million
UMAP: Uniform Manifold Approximation and Projection
WD: well differentiated
WDLPS: well-differentiated liposarcoma

## Declarations

### Ethics approval and consent to participate

The human liposarcoma tissues were collected under approval of the Ethical Committee for Clinical Research at Istituto Nazionale per lo Studio e la Cura dei Tumori, IRCCS Fondazione “G. Pascale”, Naples, Italy (n.40/19OSS). The study protocols conformed to the ethical guidelines of the 1975 Declaration of Helsinki, as per ethical approval given by the institutional review board. Written informed consent was obtained from all participants. Diagnosis of liposarcoma was established on radiological findings and was pathologically proven in all patients.

### Data availability

The datasets used and/or analysed during the current study are being deposited in GEO with a pending accession number.

### Competing interests

The authors declare no competing interests.

### Funding

This study was supported by POR CAMPANIA Region FESR 2014/2020 grants “Dalla Genomica Alla Terapia Di Tumori Rari” and “Campania Oncoterapie” (CUP B61G18000470007); by Cohesion Fund 2021/2027 CAMPANIA Region grant “MOSTRA Project” – (“Promozione di progetti di ricerca, sviluppo sperimentale e innovazione collaborativi nel campo delle malattie rare” - “Studio multi-omico per la diagnosi e la caratterizzazione dei tumori rari”, IntervN.29 - CUPE63C25000270002); by the Italian Association for Cancer Research (AIRC) (Grant IG26414/2021); by Italian Ministry of University and Research (Progetti di Rilevante Interesse Nazionale, PRIN2020 - cod. 2020XBCMHJ; PRIN2022 - cod. 20227RH9M4).

### Author contributions

RT: conceptualization, data acquisition, interpretation, analysis, data curation and visualisation, writing original draft of manuscript, writing-review and editing.

GS: conceptualization, data acquisition, interpretation, analysis, writing-review and editing.

ABo: statistical analyses, analysis of bulk and single-cell transcriptomic data, interpretation, data curation and visualisation.

GG, EM, SO, ABa, MS, AM, MCe, AA, GDP, ES, MCa, ST, FP, ABu: data acquisition, methodology

CF, AI: statistical analyses, analysis of bulk transcriptomic data, data curation and visualisation.

BF, ABu: conceptualization, interpretation, project administration, funding acquisition, writing-review and editing

PC: interpretation, analysis, data curation and visualisation, writing original draft of manuscript, writing-review and editing, project administration, supervision, funding acquisition

All authors read and approved the final manuscript.

## Acknowledgements

We thank the Institutional Biobank (BBI-Pascale) of the INT IRCCS Pascale (Naples, Italy) for providing support and resources for this study. We also acknowledge the Advanced Microscopy and Imaging Core Facility of Telethon Institute of Genetics and Medicine (TIGEM, Pozzuoli, Italy), for providing technical support and the microscopes utilized in our study.

## Supplementary information

Supplementary file 1 – HCE images for all the samples (pdf file).

Table S1. DEG Tumour vs Normal tissues (.xlsx file)

Table S2. GSEA results Tumor vs Normal tissues (.xlsx file)

Table S3. DEG PEG10^high^ vs PEG10^low^ (.xlsx file)

Table S4. GSEA results PEG10^high^ vs PEG10^low^ (.xlsx file)

Table S5. Genes included in the following signatures: stemness, invasion, hypoxia, ECM remodelling, adipocyte differentiation and angiogenesis (.xlsx file)

## Supplementary figures

**Supplementary Figure 1.**
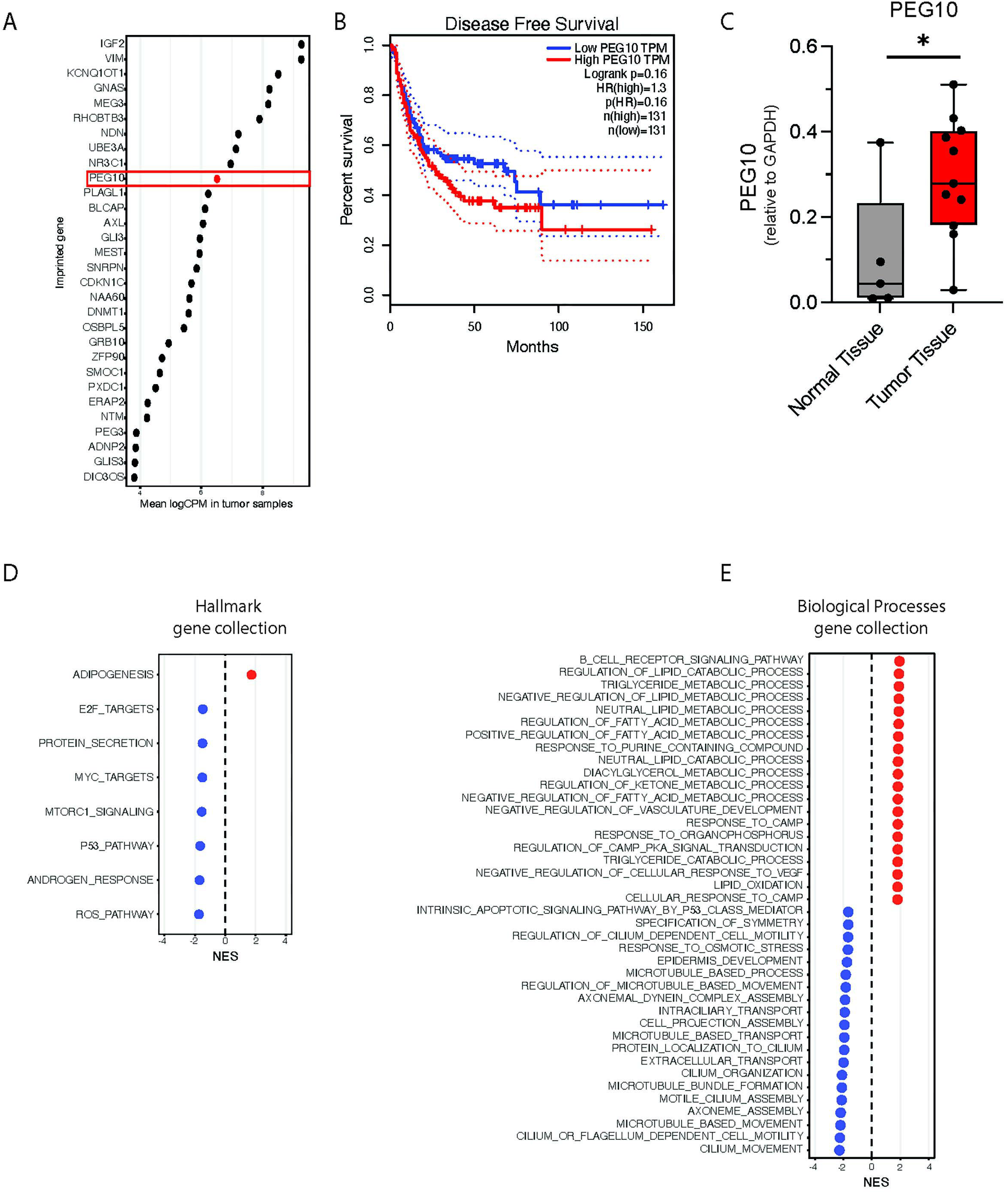
A) Dot plot showing the expression of top 30 imprinted genes in our liposarcoma cohort. B) Kaplan-Meier analysis of disease-free survival in sarcoma patients from TCGA SARC cohort, stratified according to PEG10 expression (n=262, 131 per group). Survival distributions were compared using the log-rank test. TPM: transcripts per million; HR: hazard ratio. C) Box and whiskers showing the expression of *PEG10* in normal tissues (n=5) and tumour tissues (n=11), measured by qPCR. *P* value from t-test (*: *p* ≤ 0.05). D) Dot plot showing the enriched gene sets from the Gene Ontology Hallmark gene set, in PEG10^high^ (n=11) and PEG10^low^ (n=11) tumours (FDR<0.05). E) Dot plot showing the enriched gene sets from the Gene Ontology Biological Process gene set, in PEG10^high^ (n=11) and PEG10^low^ (n=11) tumours.

**Supplementary Figure 2.**
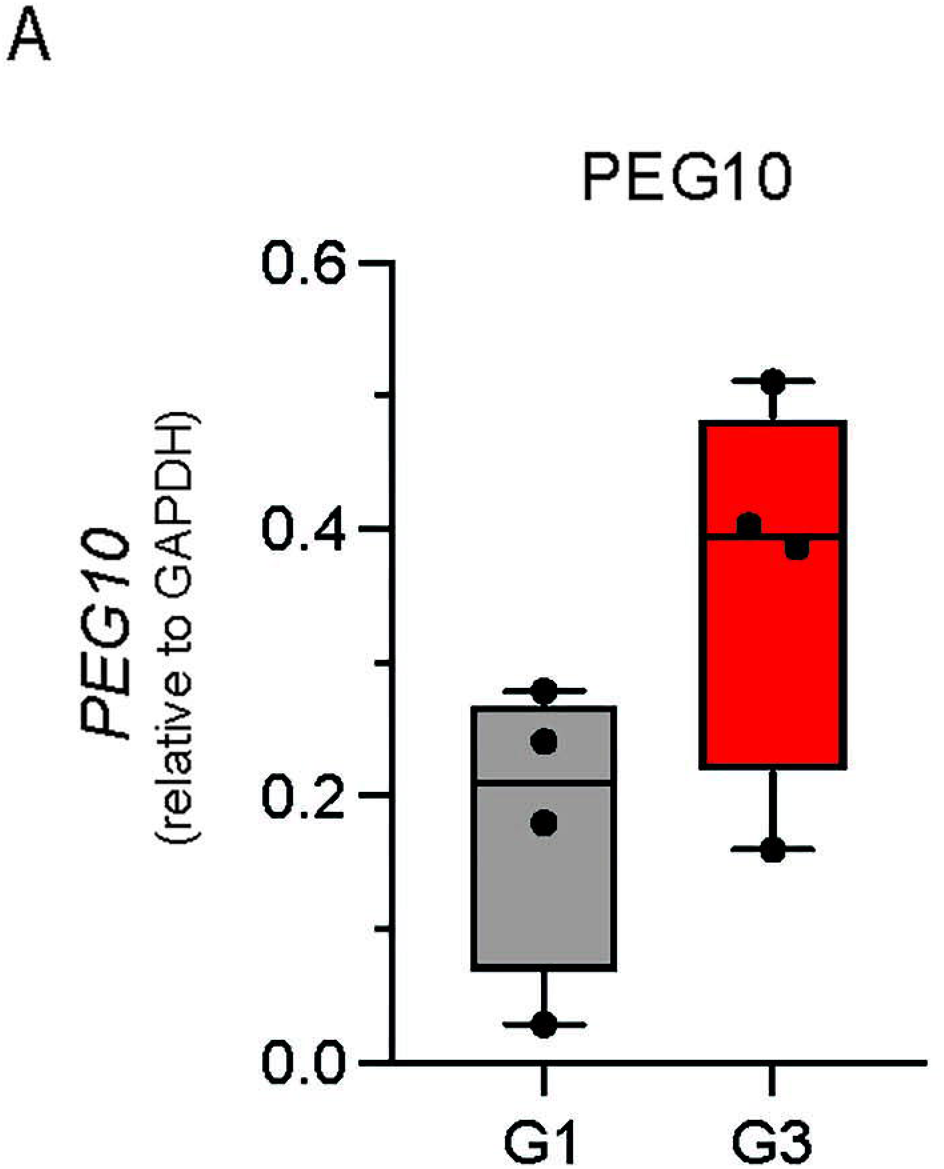
A) Box and whiskers showing the expression of *PEG10* in grade 1 (G1, n=4), and grade 3 (G3, n=4) liposarcoma, measured by qPCR.

**Supplementary Figure 3.**
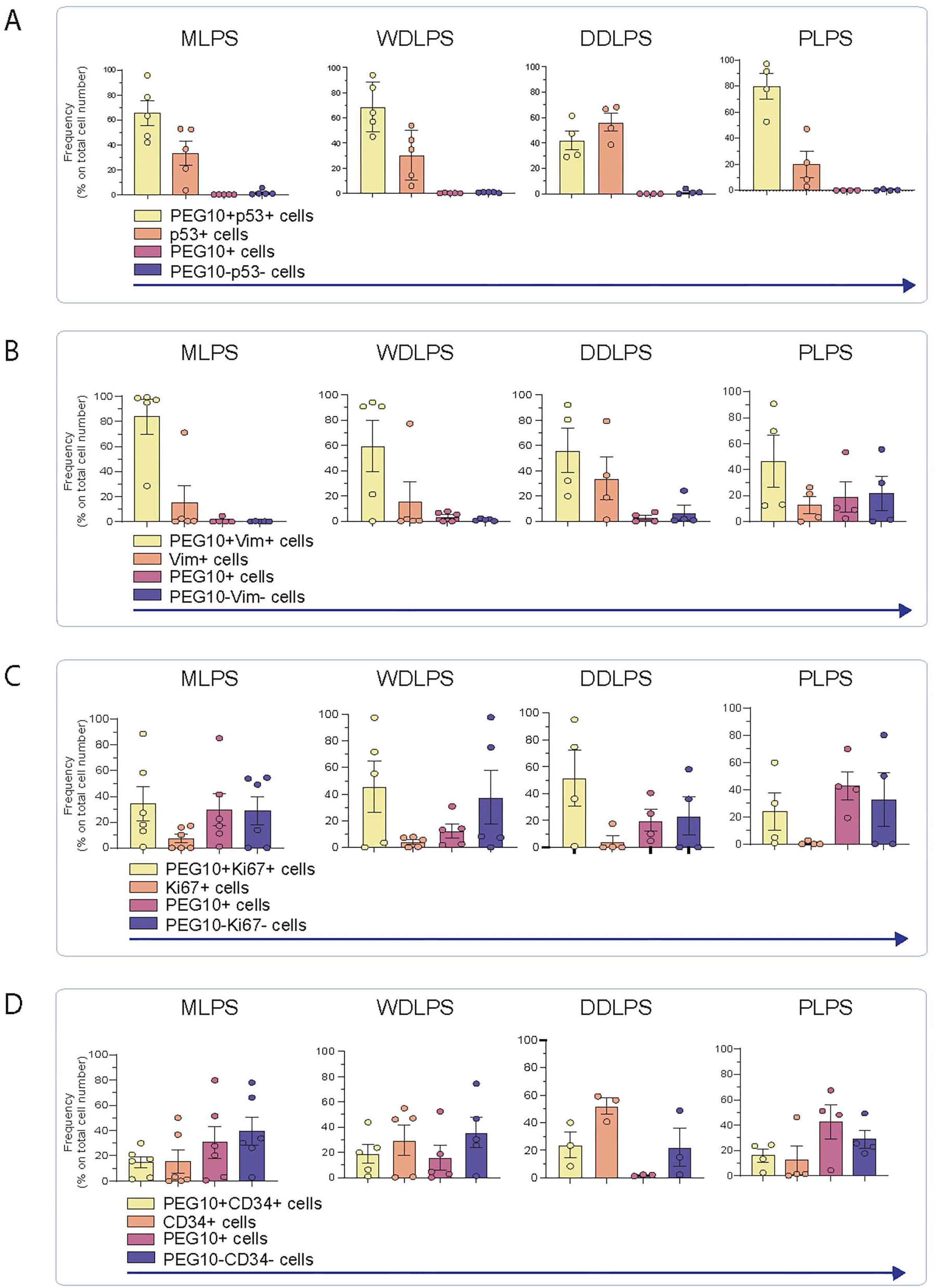
A) Bar plot showing the frequency of cells expressing PEG10, p53, both markers or neither in the different liposarcoma subtypes (MLPS, n=5, WDLPS, n=5, DDLPS, n=4, PLPS, n=4). B) Bar plot showing the frequency of cells expressing PEG10, vimentin (Vim), both markers or neither in the different liposarcoma subtypes (MLPS, n=5, WDLPS, n=5, DDLPS, n=4, PLPS, n=4). C) Bar plot showing the frequency of cells expressing PEG10, Ki67, both markers or neither in the different liposarcoma subtypes (MLPS, n=6, WDLPS, n=5, DDLPS, n=4, PLPS, n=4). D) Bar plot showing the frequency of cells expressing PEG10, CD34, both markers or neither in the different liposarcoma subtypes (MLPS, n=6, WDLPS, n=5, DDLPS, n=3, PLPS, n=4). A-D) Bars indicate the mean values ± SEM. Dots indicate value from individual patients.

**Supplementary Figure 4.**
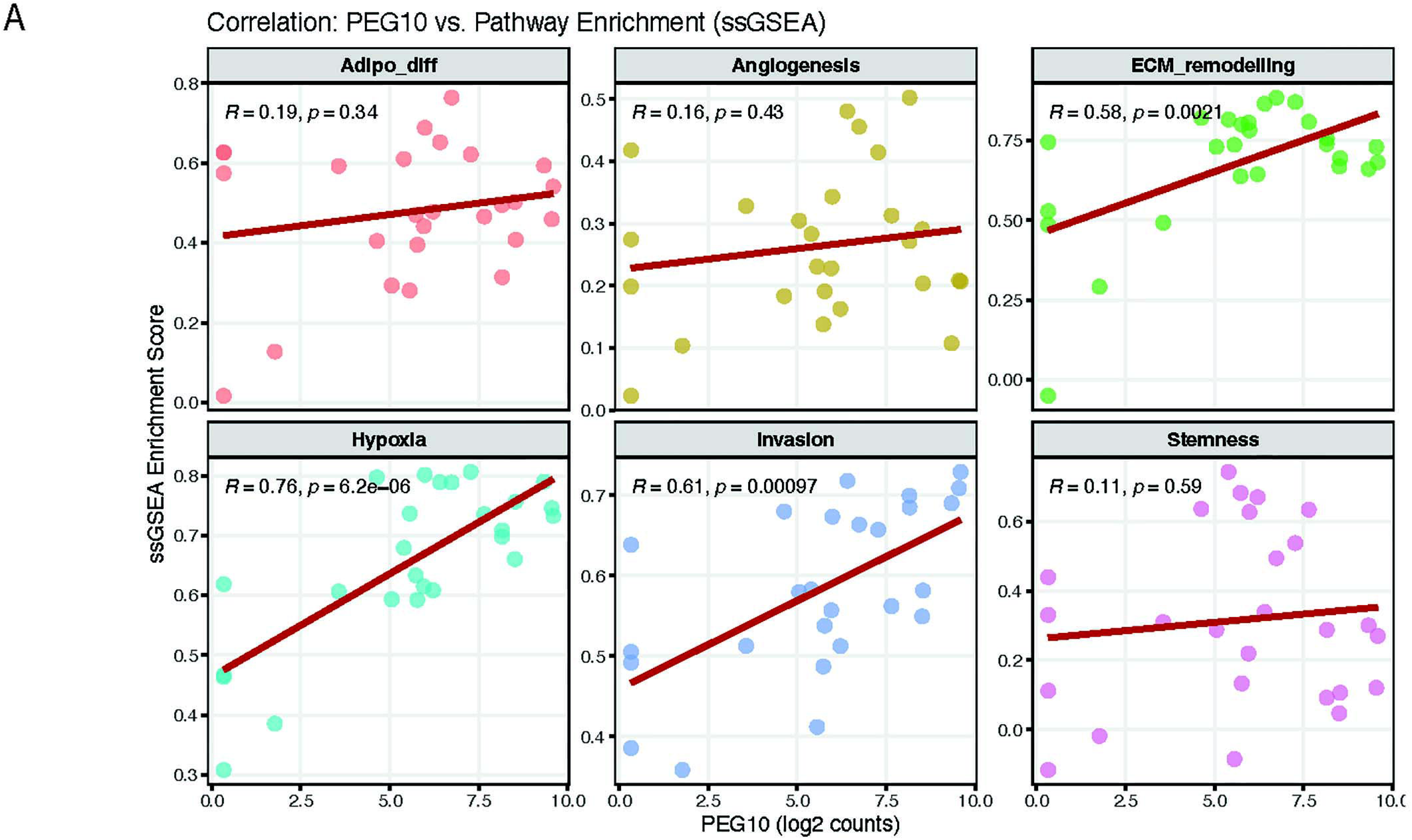
A) Scatter plots showing the correlation between PEG10 expression and single-sample gene set enrichment analysis (ssGSEA) for the following gene sets: adipocyte differentiation (Adipo_diff), angiogenesis, extracellular matrix remodelling (ECM remodelling), hypoxia, invasion and stemness. Each dot represents an individual tumour sample (n=22). The red line indicates the fitted linear regression. Pearson correlation coefficient (R) and corresponding *P* values are shown in each panel.

## Notes

### Competing Interest Statement

The authors have declared no competing interest.

### Summary of Updates

The new versione improved the quality of table files, and supplemental material

