## Supplementary figures and images for "Integrated transcriptomic analysis identifies Paternally Expressed Gene 10 as a novel candidate biomarker associated with aggressive features in liposarcoma"

### Supplemantary File 1

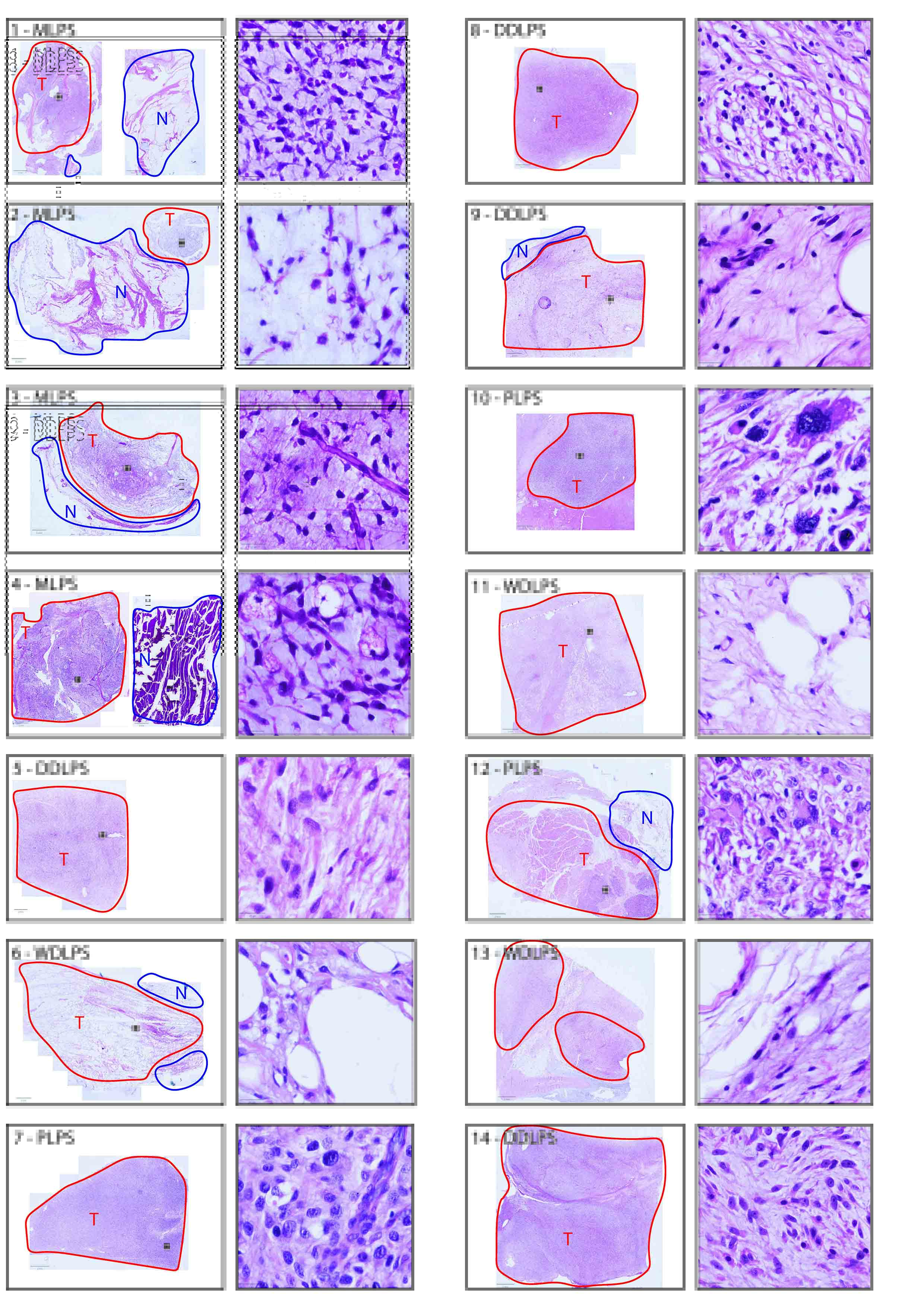

### Supplemantary File 1

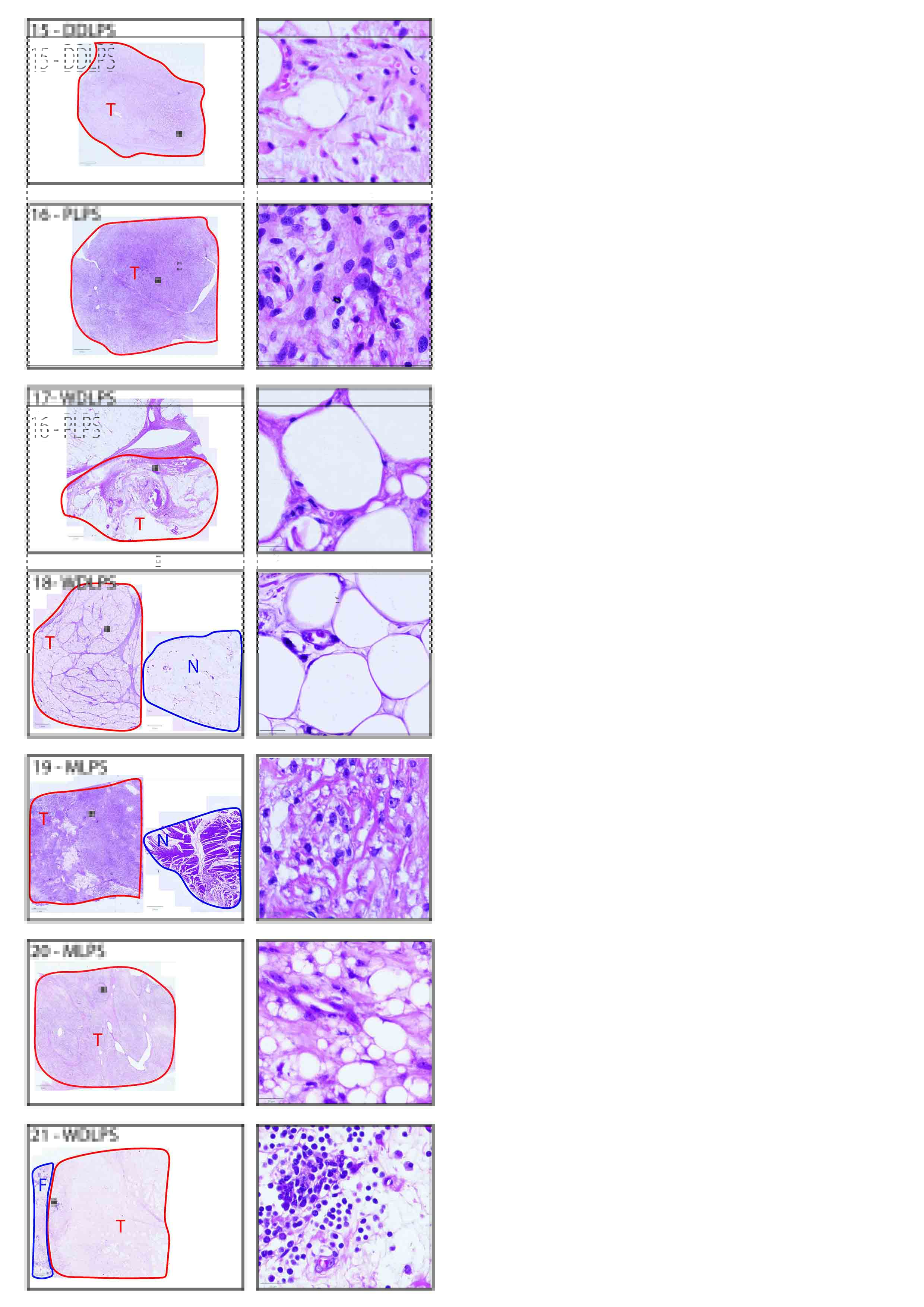
