## Supplementary material for "Integrated transcriptomic analysis identifies Paternally Expressed Gene 10 as a novel candidate biomarker associated with aggressive features in liposarcoma": Table S1

| | A | B | C | D | E | F | G | H |
|---|---|---|---|---|---|---|---|---|
| 1 | gene | ensembl_id | logCPM | logFC | F | PValue | FDR | significant |
| 2 | FITM1 | ENSG00000139914 | 5.643426764 | -7.348545302 | 105.0548355 | 8.44458E-16 | 1.23603E-11 | Yes |
| 3 | SMTNL1 | ENSG00000214872 | 6.657838035 | -9.614828566 | 110.6582801 | 3.16994E-15 | 2.31992E-11 | Yes |
| 4 | PGAM2 | ENSG00000164708 | 7.997400191 | -9.624340725 | 95.27581509 | 7.52065E-15 | 3.66933E-11 | Yes |
| 5 | MIR1.1HG | | 7.097871349 | -8.796027643 | 91.0623094 | 1.79828E-14 | 6.58034E-11 | Yes |
| 6 | TRIM63 | ENSG00000158022 | 7.971688705 | -8.812780221 | 82.41146884 | 3.77554E-13 | 1.10525E-09 | Yes |
| 7 | ATP2A1 | ENSG00000196296 | 9.456350007 | -9.222661887 | 85.95181447 | 1.04081E-12 | 2.53905E-09 | Yes |
| 8 | SGCA | ENSG00000108823 | 6.404001711 | -7.022970686 | 77.13085495 | 1.63241E-12 | 3.41337E-09 | Yes |
| 9 | MYOM1 | ENSG00000101605 | 9.883524636 | -6.454069902 | 93.27876461 | 2.76313E-12 | 5.0555E-09 | Yes |
| 10 | RTN2 | ENSG00000125744 | 6.20125717 | -5.888136731 | 79.88394493 | 3.31027E-12 | 5.3836E-09 | Yes |
| 11 | FHL3 | ENSG00000183386 | 8.094327819 | -6.190591164 | 86.63051191 | 4.47652E-12 | 6.55228E-09 | Yes |
| 12 | ENO3 | ENSG00000108515 | 9.77238978 | -7.910647626 | 85.37360246 | 5.65023E-12 | 7.51841E-09 | Yes |
| 13 | RYR1 | ENSG00000196218 | 7.39495497 | -7.274430539 | 76.07660568 | 6.20963E-12 | 7.5742E-09 | Yes |
| 14 | TNNT3 | ENSG00000130595 | 9.772485799 | -6.631042118 | 86.59487019 | 7.63594E-12 | 8.59749E-09 | Yes |
| 15 | CACNA1S | ENSG00000081248 | 5.024479056 | -7.29788936 | 66.44673543 | 8.25536E-12 | 8.63097E-09 | Yes |
| 16 | AMPD1 | ENSG00000116748 | 6.401512021 | -7.458498563 | 67.62049871 | 8.93666E-12 | 8.7204E-09 | Yes |
| 17 | DHRS7C | ENSG00000184544 | 5.341274381 | -7.473254717 | 65.33234776 | 9.87788E-12 | 9.03641E-09 | Yes |
| 18 | CAV3 | ENSG00000182533 | 4.862831375 | -7.012747354 | 63.71353566 | 1.56116E-11 | 1.34416E-08 | Yes |
| 19 | TRIM7 | ENSG00000146054 | 5.59389828 | -5.959720942 | 67.07692618 | 2.24147E-11 | 1.82269E-08 | Yes |
| 20 | MT1X | ENSG00000187193 | 8.447058473 | -4.330022837 | 80.50314738 | 2.49916E-11 | 1.8304E-08 | Yes |
| 21 | PYGM | ENSG00000068976 | 9.701018977 | -7.553484507 | 77.71974597 | 2.57587E-11 | 1.8304E-08 | Yes |
| 22 | UCP3 | ENSG00000175564 | 5.243592615 | -5.927981789 | 64.66107696 | 2.62611E-11 | 1.8304E-08 | Yes |
| 23 | RPL3L | ENSG00000140986 | 6.109026532 | -7.257173242 | 62.23358034 | 3.28256E-11 | 2.18395E-08 | Yes |
| 24 | MYPN | ENSG00000138347 | 5.009811886 | -6.476810545 | 60.16344445 | 4.28291E-11 | 2.72561E-08 | Yes |
| 25 | STAC3 | ENSG00000185482 | 8.194404865 | -6.60302454 | 72.96891392 | 4.64474E-11 | 2.76463E-08 | Yes |
| 26 | COX7A1 | ENSG00000161281 | 8.699334301 | -6.073912822 | 75.31451167 | 4.72199E-11 | 2.76463E-08 | Yes |
| 27 | HJV | ENSG00000168509 | 5.524677943 | -6.942168405 | 59.55518207 | 5.36115E-11 | 3.01812E-08 | Yes |
| 28 | CLCN1 | ENSG00000188037 | 4.110977767 | -7.024031537 | 64.05726771 | 5.93709E-11 | 3.21856E-08 | Yes |
| 29 | SMYD1 | ENSG00000115593 | 4.907525844 | -6.827513302 | 58.33217671 | 6.39611E-11 | 3.26538E-08 | Yes |
| 30 | MLIP | ENSG00000146147 | 5.180964773 | -6.172512855 | 59.95287707 | 6.46962E-11 | 3.26538E-08 | Yes |
| 31 | SYPL2 | ENSG00000143028 | 6.212642564 | -6.180022497 | 64.47225502 | 6.9552E-11 | 3.38987E-08 | Yes |
| 32 | DUSP26 | ENSG00000133878 | 6.719714116 | -7.083890544 | 62.60675694 | 7.22893E-11 | 3.38987E-08 | Yes |
| 33 | MYL3 | ENSG00000160808 | 8.865932318 | -7.935277503 | 68.51633253 | 7.41106E-11 | 3.38987E-08 | Yes |
| 34 | CASQ1 | ENSG00000143318 | 7.760905572 | -7.609626587 | 64.75647131 | 8.0053E-11 | 3.55071E-08 | Yes |
| 35 | SRL | ENSG00000185739 | 5.683228592 | -6.495900649 | 59.09304962 | 1.02585E-10 | 4.4163E-08 | Yes |
| 36 | OBSCN | ENSG00000154358 | 7.457393877 | -6.131703621 | 67.71150653 | 1.11226E-10 | 4.65149E-08 | Yes |
| 37 | GAMT | ENSG00000130005 | 5.920263103 | -5.147331275 | 64.64906826 | 1.2938E-10 | 5.2604E-08 | Yes |
| 38 | UNC45B | ENSG00000141161 | 4.862693869 | -6.862464323 | 55.54986628 | 1.4448E-10 | 5.48312E-08 | Yes |
| 39 | LDB3 | ENSG00000122367 | 9.354818772 | -7.182943287 | 68.68941433 | 1.45825E-10 | 5.48312E-08 | Yes |
| 40 | FHL1 | ENSG00000022267 | 8.75853884 | -5.228729547 | 70.39788842 | 1.46097E-10 | 5.48312E-08 | Yes |
| 41 | TPM1 | ENSG00000140416 | 9.82262702 | -4.304347629 | 71.22324207 | 1.50289E-10 | 5.49944E-08 | Yes |
| 42 | COX6A2 | ENSG00000156885 | 9.339728036 | -8.562263623 | 64.33059871 | 1.62904E-10 | 5.81567E-08 | Yes |
| 43 | TNNI2 | ENSG00000130598 | 10.70647171 | -8.436507791 | 66.81400623 | 2.2252E-10 | 7.73325E-08 | Yes |
| 44 | TMEM52 | ENSG00000178821 | 6.085850175 | -6.743012045 | 56.74484167 | 2.28768E-10 | 7.73325E-08 | Yes |
| 45 | ALDOA | ENSG00000149925 | 10.58469575 | -4.148504262 | 68.99817485 | 2.34839E-10 | 7.73325E-08 | Yes |
| 46 | ENSG00000268518 | ENSG00000268518 | 4.911443358 | -6.715427905 | 53.78149573 | 2.37751E-10 | 7.73325E-08 | Yes |
| 47 | KCNJ11 | ENSG00000187486 | 4.092477794 | -6.3935507 | 54.33542135 | 2.48057E-10 | 7.89307E-08 | Yes |
| 48 | MAFA | ENSG00000182759 | 3.658770315 | -5.377868813 | 53.39620821 | 2.60453E-10 | 8.11116E-08 | Yes |
| 49 | MYADML2 | ENSG00000185105 | 4.702992841 | -6.815985772 | 51.89707108 | 4.37569E-10 | 1.33431E-07 | Yes |
| 50 | GAPDH | ENSG00000111640 | 12.22495748 | -4.084822891 | 65.40533105 | 4.84009E-10 | 1.41922E-07 | Yes |
| 51 | SYNPO | ENSG00000171992 | 8.085736291 | -4.223422173 | 64.41558408 | 4.84806E-10 | 1.41922E-07 | Yes |
| 52 | TPM2 | ENSG00000198467 | 10.80206932 | -5.181251247 | 64.23751443 | 5.8229E-10 | 1.67117E-07 | Yes |
| 53 | PPP1R27 | ENSG00000182676 | 5.596247081 | -6.833003152 | 51.46939027 | 6.06435E-10 | 1.707E-07 | Yes |
| 54 | PHKG1 | ENSG00000164776 | 5.43445211 | -5.607129153 | 54.22094338 | 6.73438E-10 | 1.83843E-07 | Yes |
| 55 | CACNG1 | ENSG00000108878 | 4.551656429 | -6.464138036 | 50.27508905 | 6.78249E-10 | 1.83843E-07 | Yes |
| 56 | TNNC2 | ENSG00000101470 | 11.8260773 | -8.85602782 | 62.00260133 | 7.20038E-10 | 1.91622E-07 | Yes |
| 57 | SMTNL2 | ENSG00000188176 | 4.453522529 | -5.48487004 | 50.83210274 | 7.71467E-10 | 2.01642E-07 | Yes |
| 58 | COQ8A | ENSG00000163050 | 7.349683719 | -4.899819947 | 60.57710149 | 8.06822E-10 | 2.07183E-07 | Yes |
| 59 | TCAP | ENSG00000173991 | 11.13174362 | -8.000842464 | 61.42818051 | 8.34968E-10 | 2.10714E-07 | Yes |
| 60 | TMEM38A | ENSG00000072954 | 7.536623219 | -6.059506556 | 58.63475729 | 8.67343E-10 | 2.13855E-07 | Yes |
| 61 | SYNPO2L | ENSG00000166317 | 6.073273551 | -6.408818535 | 52.91965117 | 8.76635E-10 | 2.13855E-07 | Yes |
| 62 | LINC01569 | ENSG00000262468 | 3.907988636 | -5.312967105 | 49.35943608 | 9.40042E-10 | 2.25564E-07 | Yes |
| 63 | HSPB2 | ENSG00000170276 | 5.305112896 | -4.5244783 | 55.35333039 | 1.05312E-09 | 2.48621E-07 | Yes |
| 64 | CKM | ENSG00000104879 | 10.62958561 | -8.698539718 | 58.01708425 | 1.26526E-09 | 2.93962E-07 | Yes |
| 65 | FBXO32 | ENSG00000156804 | 7.5400082 | -4.808787976 | 58.27479113 | 1.43291E-09 | 3.2771E-07 | Yes |
| 66 | PKIA | ENSG00000171033 | 6.646308145 | -5.324449041 | 55.41611964 | 1.61327E-09 | 3.63284E-07 | Yes |
| 67 | STRIT1 | ENSG00000240045 | 7.964988773 | -7.426323715 | 53.4837467 | 1.68819E-09 | 3.71519E-07 | Yes |
| 68 | TNNC1 | ENSG00000114854 | 9.330993888 | -8.006666362 | 55.52480078 | 1.70061E-09 | 3.71519E-07 | Yes |
| 69 | CMYA5 | ENSG00000164309 | 7.787391136 | -6.28401586 | 55.55942041 | 1.78611E-09 | 3.8446E-07 | Yes |
| 70 | ADCY2 | ENSG00000078295 | 4.610597467 | -5.23704297 | 48.79550499 | 1.89775E-09 | 4.02571E-07 | Yes |
| 71 | MYH1 | ENSG00000109061 | 5.912338041 | -6.537457059 | 48.59028717 | 2.24755E-09 | 4.69963E-07 | Yes |
| 72 | SMPX | ENSG00000091482 | 5.356835429 | -6.183720212 | 47.44520133 | 2.60853E-09 | 5.37762E-07 | Yes |
| 73 | SLC29A2 | ENSG00000174669 | 5.849207876 | -5.638149874 | 50.38571207 | 2.67667E-09 | 5.44144E-07 | Yes |
| 74 | FBXO40 | ENSG00000163833 | 4.969335367 | -6.537316941 | 45.54062254 | 3.05407E-09 | 6.12361E-07 | Yes |
| 75 | HRC | ENSG00000130528 | 6.257926579 | -5.740935186 | 50.42345924 | 3.37782E-09 | 6.68123E-07 | Yes |
| 76 | TTN | ENSG00000155657 | 12.7410517 | -7.502788928 | 55.5626604 | 3.51165E-09 | 6.85333E-07 | Yes |
| 77 | DUSP13B | ENSG00000079393 | 5.392706384 | -6.367719285 | 46.02512595 | 3.61207E-09 | 6.95656E-07 | Yes |
| 78 | LMOD3 | ENSG00000163380 | 6.076598512 | -6.414420955 | 47.13879995 | 4.29185E-09 | 8.15843E-07 | Yes |
| 79 | EEF1A2 | ENSG00000101210 | 8.716458343 | -7.099260302 | 51.7466678 | 4.68629E-09 | 8.79401E-07 | Yes |
| 80 | TPM3 | ENSG00000143549 | 8.658349073 | -4.325106104 | 53.6730989 | 4.97896E-09 | 9.22494E-07 | Yes |
| 81 | MYF6 | ENSG00000111046 | 4.987603928 | -6.037788758 | 44.66491075 | 5.08983E-09 | 9.31248E-07 | Yes |
| 82 | SRPK3 | ENSG00000184343 | 6.212730498 | -6.277474487 | 47.24530532 | 5.23148E-09 | 9.45349E-07 | Yes |
| 83 | MYO18B | ENSG00000133454 | 4.380795348 | -5.776696865 | 43.33473537 | 6.28011E-09 | 1.121E-06 | Yes |
| 84 | MYOZ1 | ENSG00000177791 | 9.229628508 | -7.382772066 | 50.20616634 | 7.40962E-09 | 1.30668E-06 | Yes |
| 85 | TRIM54 | ENSG00000138100 | 6.668230199 | -6.566958457 | 46.26756227 | 7.90903E-09 | 1.37815E-06 | Yes |
| 86 | FEM1A | ENSG00000141965 | 7.565064078 | -3.982752614 | 51.01364682 | 8.34132E-09 | 1.42147E-06 | Yes |
| 87 | PEBP4 | ENSG00000134020 | 5.719284758 | -5.771318227 | 45.49487769 | 8.35189E-09 | 1.42147E-06 | Yes |
| 88 | MYOM2 | ENSG00000036448 | 7.206183563 | -5.197543608 | 49.44411078 | 8.97254E-09 | 1.49427E-06 | Yes |
| 89 | PRELP | ENSG00000188783 | 5.69247972 | -4.762181236 | 47.39364628 | 8.98381E-09 | 1.49427E-06 | Yes |
| 90 | DES | ENSG00000175084 | 10.3056245 | -6.521724103 | 50.80031292 | 9.30648E-09 | 1.53055E-06 | Yes |
| 91 | KLHL41 | ENSG00000239474 | 8.273100833 | -6.901390406 | 48.39251531 | 9.60536E-09 | 1.56215E-06 | Yes |
| 92 | FXYD1 | ENSG00000266964 | 8.159860305 | -4.219104149 | 50.43854133 | 1.00106E-08 | 1.61016E-06 | Yes |
| 93 | COL4A3 | ENSG00000169031 | 3.290194253 | -5.774220747 | 42.52783962 | 1.15281E-08 | 1.8341E-06 | Yes |
| 94 | ASB2 | ENSG00000100628 | 6.105939312 | -6.090760547 | 44.43331353 | 1.21399E-08 | 1.91066E-06 | Yes |
| 95 | ATP2A2 | ENSG00000174437 | 8.988977799 | -3.802467907 | 49.80821182 | 1.25521E-08 | 1.94141E-06 | Yes |
| 96 | SHISA4 | ENSG00000198892 | 6.606277007 | -4.210525963 | 48.3306617 | 1.26005E-08 | 1.94141E-06 | Yes |
| 97 | S100A1 | ENSG00000160678 | 7.352967786 | -6.159955534 | 46.65325274 | 1.35681E-08 | 2.06871E-06 | Yes |
| 98 | SELENOW | ENSG00000178980 | 9.807996742 | -2.998688004 | 49.45443123 | 1.41203E-08 | 2.11103E-06 | Yes |
| 99 | VHRT | ENSG00000185847 | 7.229296536 | -6.782569357 | 45.00924296 | 1.42641E-08 | 2.11103E-06 | Yes |
| 100 | MYH7 | ENSG00000092054 | 10.48819054 | -8.271947714 | 47.96769994 | 1.42783E-08 | 2.11103E-06 | Yes |
| 101 | LRRC20 | ENSG00000172731 | 5.400957143 | -4.703843717 | 43.96072814 | 1.9616E-08 | 2.87119E-06 | Yes |
| 102 | SPTB | ENSG00000070182 | 6.100233851 | -5.639612236 | 43.06073672 | 2.34253E-08 | 3.39481E-06 | Yes |
| 103 | MYBPC2 | ENSG00000086967 | 8.99507343 | -7.317677354 | 45.29560588 | 2.37447E-08 | 3.40736E-06 | Yes |
| 104 | ENSG00000235475 | ENSG00000235475 | 4.022644514 | -5.67425958 | 38.81548572 | 2.59217E-08 | 3.68365E-06 | Yes |
| 105 | TNNT1 | ENSG00000105048 | 11.90530166 | -7.901062665 | 46.05108911 | 2.93238E-08 | 4.12704E-06 | Yes |
| 106 | CACNG6 | ENSG00000130433 | 4.972835814 | -5.417900104 | 39.46897583 | 3.68259E-08 | 5.13352E-06 | Yes |
| 107 | MAMSTR | ENSG00000176909 | 6.001856134 | -4.885012009 | 42.28553645 | 3.92914E-08 | 5.42554E-06 | Yes |
| 108 | TMOD4 | ENSG00000163157 | 3.660234197 | -5.262653735 | 37.40956879 | 4.14754E-08 | 5.6736E-06 | Yes |
| 109 | CACNB1 | ENSG00000067191 | 5.866075868 | -4.714900852 | 42.04490141 | 4.23139E-08 | 5.7347E-06 | Yes |
| 110 | MB | ENSG00000198125 | 12.60628503 | -8.43806684 | 44.31217142 | 4.54435E-08 | 6.10235E-06 | Yes |
| 111 | MYOZ3 | ENSG00000164591 | 5.515065994 | -5.43192556 | 39.71731264 | 4.80563E-08 | 6.36863E-06 | Yes |
| 112 | TRDN | ENSG00000186439 | 8.983394904 | -6.530292188 | 43.26726567 | 4.82966E-08 | 6.36863E-06 | Yes |
| 113 | MYL6B | ENSG00000196465 | 7.589741913 | -4.080916744 | 43.32903083 | 5.33885E-08 | 6.9772E-06 | Yes |
| 114 | CAPN3 | ENSG00000092529 | 7.064510139 | -3.895567877 | 42.87478138 | 5.69629E-08 | 7.37846E-06 | Yes |
| 115 | ANKRD2 | ENSG00000165887 | 6.24675104 | -5.905500547 | 39.17635726 | 6.81414E-08 | 8.74899E-06 | Yes |
| 116 | ALPK3 | ENSG00000136383 | 5.975355406 | -4.85881623 | 40.19291266 | 7.02528E-08 | 8.94165E-06 | Yes |
| 117 | HPN | ENSG00000105707 | 3.756029195 | -4.656875205 | 35.85485871 | 8.44342E-08 | 1.0654E-05 | Yes |
| 118 | PADI2 | ENSG00000117115 | 6.574915712 | -5.488290171 | 39.155166 | 9.60513E-08 | 1.20067E-05 | Yes |
| 119 | SLC25A4 | ENSG00000151729 | 6.154032772 | -4.377056891 | 39.80082656 | 9.67952E-08 | 1.20067E-05 | Yes |
| 120 | FLNC | ENSG00000128591 | 7.42631797 | -4.200820465 | 40.82823344 | 9.8583E-08 | 1.21257E-05 | Yes |
| 121 | LTK | ENSG00000062524 | 4.429285321 | -5.028363663 | 35.64769158 | 1.10946E-07 | 1.35327E-05 | Yes |
| 122 | ATP1A2 | ENSG00000018625 | 6.291419898 | -5.513116292 | 37.88788115 | 1.21412E-07 | 1.46869E-05 | Yes |
| 123 | KIF1B | ENSG00000054523 | 7.634108463 | -2.648604651 | 40.25989182 | 1.27286E-07 | 1.52712E-05 | Yes |
| 124 | JPH2 | ENSG00000149596 | 6.15223051 | -5.152498972 | 37.956725 | 1.28829E-07 | 1.53307E-05 | Yes |
| 125 | MYO18A | ENSG00000196535 | 5.993543169 | -3.578305096 | 39.10297605 | 1.32064E-07 | 1.55889E-05 | Yes |
| 126 | NEB | ENSG00000183091 | 10.00223912 | -6.741382453 | 39.67600059 | 1.38459E-07 | 1.6213E-05 | Yes |
| 127 | MBNL1.AS1 | | 4.852616426 | -4.3877293 | 36.19623402 | 1.57552E-07 | 1.83023E-05 | Yes |
| 128 | SEMA6C | ENSG00000143434 | 7.588705378 | -4.093110902 | 38.79240972 | 1.7405E-07 | 2.00596E-05 | Yes |
| 129 | PPP1R1A | ENSG00000135447 | 7.328602933 | -5.407424611 | 37.66636196 | 1.89539E-07 | 2.1674E-05 | Yes |
| 130 | SYNPO2 | ENSG00000172403 | 7.987110018 | -3.992111692 | 38.52480154 | 1.94911E-07 | 2.21156E-05 | Yes |
| 131 | PLCD4 | ENSG00000115556 | 4.903913817 | -4.168934824 | 35.76668618 | 2.02176E-07 | 2.27635E-05 | Yes |
| 132 | SLN | ENSG00000170290 | 9.311171323 | -6.930017322 | 37.56753907 | 2.18362E-07 | 2.43982E-05 | Yes |
| 133 | SH2B2 | ENSG00000160999 | 4.397337974 | -4.451817107 | 33.67154697 | 2.69103E-07 | 2.98398E-05 | Yes |
| 134 | CRYAB | ENSG00000109846 | 8.782368042 | -4.418300632 | 37.37051199 | 2.73548E-07 | 3.01047E-05 | Yes |
| 135 | APOD | ENSG00000189058 | 8.351490645 | -3.73166956 | 37.32565477 | 2.78144E-07 | 3.0382E-05 | Yes |
| 136 | SH3BGR | ENSG00000185437 | 5.080252876 | -4.367368911 | 34.71163848 | 2.83317E-07 | 3.07178E-05 | Yes |
| 137 | CCL21 | ENSG00000137077 | 3.85923174 | -4.645253105 | 32.19652368 | 3.12668E-07 | 3.35129E-05 | Yes |
| 138 | IDH2 | ENSG00000182054 | 7.919351399 | -4.294127953 | 36.68446556 | 3.13676E-07 | 3.35129E-05 | Yes |
| 139 | LMOD2 | ENSG00000170807 | 8.943662759 | -6.70180291 | 35.85158927 | 3.44105E-07 | 3.64976E-05 | Yes |
| 140 | COQ10A | ENSG00000135469 | 5.675880576 | -4.002098246 | 35.12038406 | 3.46848E-07 | 3.65238E-05 | Yes |
| 141 | PLEC | ENSG00000178209 | 7.519397714 | -2.598053733 | 36.36277961 | 3.59716E-07 | 3.76083E-05 | Yes |
| 142 | TMEM233 | ENSG00000224982 | 6.495034861 | -4.392787809 | 35.29503695 | 3.74811E-07 | 3.89086E-05 | Yes |
| 143 | ZNF106 | ENSG00000103994 | 8.37541858 | -2.651723729 | 36.09761251 | 3.99279E-07 | 4.10108E-05 | Yes |
| 144 | LRRC2 | ENSG00000163827 | 5.287225164 | -4.495593328 | 33.72465443 | 4.00665E-07 | 4.10108E-05 | Yes |
| 145 | HSPB1 | ENSG00000106211 | 11.95840597 | -2.628008425 | 36.13692657 | 4.10552E-07 | 4.17309E-05 | Yes |
| 146 | SCN1B | ENSG00000105711 | 7.656511304 | -4.433446603 | 35.55637333 | 4.14275E-07 | 4.18189E-05 | Yes |
| 147 | HSPB7 | ENSG00000173641 | 7.83334513 | -5.067701752 | 35.1657836 | 4.43186E-07 | 4.41297E-05 | Yes |
| 148 | SVIL | ENSG00000197321 | 7.073286288 | -3.370438808 | 35.38825442 | 4.43196E-07 | 4.41297E-05 | Yes |
| 149 | MYL11 | ENSG00000180209 | 11.72725293 | -6.875250004 | 35.62539034 | 4.53687E-07 | 4.48112E-05 | Yes |
| 150 | MYOD1 | ENSG00000129152 | 4.432880339 | -5.116176475 | 31.37177634 | 4.56164E-07 | 4.48112E-05 | Yes |
| 151 | MYL2 | ENSG00000111245 | 11.91725984 | -7.903107385 | 35.42987656 | 4.64283E-07 | 4.53047E-05 | Yes |
| 152 | NEURL1 | ENSG00000107954 | 4.833209235 | -5.087676596 | 31.74068726 | 4.85866E-07 | 4.70969E-05 | Yes |
| 153 | NRAP | ENSG00000197893 | 8.776005174 | -6.644907403 | 34.36521285 | 5.16474E-07 | 4.96939E-05 | Yes |
| 154 | BRMS1L | ENSG00000100916 | 4.510035377 | -4.328183354 | 32.03062749 | 5.19448E-07 | 4.96939E-05 | Yes |
| 155 | SLC2A4 | ENSG00000181856 | 5.773404254 | -4.698731136 | 32.90161026 | 5.81565E-07 | 5.52751E-05 | Yes |
| 156 | ACHE | ENSG00000087085 | 5.415085163 | -4.494352783 | 32.57531616 | 6.03621E-07 | 5.70013E-05 | Yes |
| 157 | ACTN2 | ENSG00000077522 | 8.530441647 | -6.300772469 | 33.71458022 | 6.32574E-07 | 5.93524E-05 | Yes |
| 158 | EMC9 | ENSG00000100908 | 4.631205903 | -3.55728503 | 32.09845579 | 6.83604E-07 | 6.37319E-05 | Yes |
| 159 | TNNI1 | ENSG00000159173 | 10.33372955 | -7.068753239 | 33.79727463 | 7.00402E-07 | 6.48847E-05 | Yes |
| 160 | MYH2 | ENSG00000125414 | 10.80590896 | -7.26809479 | 33.72680685 | 7.29512E-07 | 6.71564E-05 | Yes |
| 161 | RPS6KL1 | ENSG00000198208 | 3.137305963 | -3.856749433 | 29.46564583 | 7.63264E-07 | 6.98243E-05 | Yes |
| 162 | NDOR1 | ENSG00000188566 | 3.635601857 | -4.079679086 | 29.78901403 | 8.01305E-07 | 7.28491E-05 | Yes |
| 163 | SYNM | ENSG00000182253 | 5.242144963 | -3.635770066 | 31.93219859 | 8.59999E-07 | 7.77025E-05 | Yes |
| 164 | ATP5MC1 | ENSG00000159199 | 8.042502613 | -2.293340435 | 33.10981293 | 9.18462E-07 | 8.24757E-05 | Yes |
| 165 | WSCD1 | ENSG00000179314 | 4.250778567 | -4.518666866 | 29.51637446 | 9.90051E-07 | 8.83621E-05 | Yes |
| 166 | CAMK2B | ENSG00000058404 | 6.797507546 | -4.228294329 | 32.19215679 | 1.00394E-06 | 8.90589E-05 | Yes |
| 167 | RAPSN | ENSG00000165917 | 4.211061687 | -4.726093347 | 29.16208245 | 1.03198E-06 | 9.09947E-05 | Yes |
| 168 | ALDH2 | ENSG00000111275 | 6.683034859 | -2.428513692 | 32.43788705 | 1.04441E-06 | 9.15392E-05 | Yes |
| 169 | BIN1 | ENSG00000136717 | 8.023474358 | -2.991406885 | 32.51070824 | 1.07748E-06 | 9.38757E-05 | Yes |
| 170 | TMOD1 | ENSG00000136842 | 4.736253999 | -4.130790048 | 30.13787755 | 1.13369E-06 | 9.81885E-05 | Yes |
| 171 | PLPP7 | ENSG00000160539 | 6.637017255 | -3.891261699 | 31.77175975 | 1.1539E-06 | 9.93511E-05 | Yes |
| 172 | AGMAT | ENSG00000116771 | 3.991284013 | -4.101625873 | 29.0918195 | 1.18732E-06 | 0.000101631 | Yes |
| 173 | MIR133A1HG | ENSG00000265142 | 6.118105511 | -5.285380732 | 29.76562315 | 1.48993E-06 | 0.000126791 | Yes |
| 174 | PNMT | ENSG00000141744 | 2.292612503 | -4.394702549 | 27.67099166 | 1.5618E-06 | 0.000132139 | Yes |
| 175 | ENSG00000268061 | ENSG00000268061 | 3.508719694 | -4.212420265 | 27.53129313 | 1.646E-06 | 0.000138462 | Yes |
| 176 | GPR153 | ENSG00000158292 | 6.625118055 | 5.738909594 | 29.90270717 | 1.70669E-06 | 0.000142748 | Yes |
| 177 | ENSG00000223749 | ENSG00000223749 | 5.764872056 | 9.708012934 | 31.61840315 | 1.85548E-06 | 0.000154311 | Yes |
| 178 | PFKM | ENSG00000152556 | 5.855872394 | -3.896815232 | 29.77045309 | 1.86879E-06 | 0.00015454 | Yes |
| 179 | SCN4A | ENSG00000007314 | 6.323447187 | -4.538407689 | 29.63964501 | 1.92722E-06 | 0.000158476 | Yes |
| 180 | DMD | ENSG00000198947 | 6.551030993 | -3.176289419 | 29.94846282 | 2.09418E-06 | 0.000171243 | Yes |
| 181 | DCAF6 | ENSG00000143164 | 6.936357045 | -3.358147457 | 29.9351632 | 2.15144E-06 | 0.000174948 | Yes |
| 182 | PWWP3B | ENSG00000157502 | 2.952086856 | -5.059492918 | 26.58490288 | 2.20391E-06 | 0.000177601 | Yes |
| 183 | TCEA3 | ENSG00000204219 | 6.388462379 | -3.197684473 | 29.71551292 | 2.20833E-06 | 0.000177601 | Yes |
| 184 | MAP3K20 | ENSG00000091436 | 8.007483986 | -2.803174608 | 30.04288196 | 2.2349E-06 | 0.000178755 | Yes |
| 185 | ANK1 | ENSG00000029534 | 7.301903535 | -4.577621967 | 29.59642663 | 2.26383E-06 | 0.000180085 | Yes |
| 186 | BTG3 | ENSG00000154640 | 5.985253246 | 6.080416713 | 29.12059277 | 2.33744E-06 | 0.000184936 | Yes |
| 187 | MEX3D | ENSG00000181588 | 5.373253684 | 9.307189693 | 30.31654442 | 2.61256E-06 | 0.000205592 | Yes |
| 188 | ADSS1 | ENSG00000185100 | 5.774860539 | -3.900562007 | 28.54846324 | 2.7005E-06 | 0.000211375 | Yes |
| 189 | PTK7 | ENSG00000112655 | 6.486963883 | 10.44520391 | 30.32919101 | 2.7667E-06 | 0.000215405 | Yes |
| 190 | AGL | ENSG00000162688 | 6.415652866 | -2.967182817 | 29.00954707 | 2.7905E-06 | 0.000216108 | Yes |
| 191 | AK1 | ENSG00000106992 | 7.625572199 | -2.667478856 | 28.99315203 | 3.03966E-06 | 0.000233214 | Yes |
| 192 | TRIM72 | ENSG00000177238 | 4.742732231 | -4.806474873 | 26.6417866 | 3.04324E-06 | 0.000233214 | Yes |
| 193 | PLAUR | ENSG00000011422 | 7.096484787 | 7.897830429 | 28.783098 | 3.10801E-06 | 0.000236937 | Yes |
| 194 | CSRP3 | ENSG00000129170 | 6.03445045 | -5.089969445 | 27.57012751 | 3.14936E-06 | 0.000238846 | Yes |
| 195 | FBXW5 | ENSG00000159069 | 8.030071424 | -2.151191533 | 28.88859144 | 3.19727E-06 | 0.000240235 | Yes |
| 196 | NDRG2 | ENSG00000165795 | 9.033977981 | -3.753826355 | 28.89008108 | 3.20051E-06 | 0.000240235 | Yes |
| 197 | TRIM55 | ENSG00000147573 | 3.000116476 | -4.489666058 | 25.31985192 | 3.26744E-06 | 0.000244008 | Yes |
| 198 | FAM180B | ENSG00000196666 | 5.200283549 | -4.361337546 | 27.14581218 | 3.40623E-06 | 0.000252627 | Yes |
| 199 | ABLIM2 | ENSG00000163995 | 4.121775222 | -4.402726284 | 26.01448296 | 3.41737E-06 | 0.000252627 | Yes |
| 200 | ADPRHL1 | ENSG00000153531 | 5.644260651 | -4.004436007 | 27.6575563 | 3.45266E-06 | 0.000253953 | Yes |
| 201 | MYBPC1 | ENSG00000196091 | 10.35258168 | -6.486314735 | 28.46134873 | 3.53431E-06 | 0.000257567 | Yes |
| 202 | PRKAA2 | ENSG00000162409 | 5.543754348 | -4.088920922 | 27.4631872 | 3.53973E-06 | 0.000257567 | Yes |
| 203 | ACTA1 | ENSG00000143632 | 12.52584778 | -7.234933265 | 28.56969019 | 3.5546E-06 | 0.000257567 | Yes |
| 204 | ABCC3 | ENSG00000108846 | 5.028140043 | -4.313088652 | 26.86333342 | 3.59073E-06 | 0.000258904 | Yes |
| 205 | SMARCD3 | ENSG00000082014 | 6.802441499 | -2.821659145 | 28.24138728 | 3.66764E-06 | 0.000263153 | Yes |
| 206 | IP6K3 | ENSG00000161896 | 4.044026723 | -4.372485296 | 25.56684132 | 3.96584E-06 | 0.000283161 | Yes |
| 207 | MAPT | ENSG00000186868 | 6.114851509 | -3.498499242 | 27.59972459 | 4.04676E-06 | 0.000286614 | Yes |
| 208 | CAVIN4 | ENSG00000170681 | 4.005680454 | -4.533580599 | 25.30778863 | 4.06843E-06 | 0.000286614 | Yes |
| 209 | ACSS2 | ENSG00000131069 | 5.852567625 | -2.537428565 | 27.69758054 | 4.07295E-06 | 0.000286614 | Yes |
| 210 | GOT2 | ENSG00000125166 | 7.547186823 | -2.894441458 | 27.76059623 | 4.41101E-06 | 0.000308918 | Yes |
| 211 | TXLNB | ENSG00000164440 | 5.580684029 | -4.560443001 | 26.2875651 | 4.83695E-06 | 0.000337136 | Yes |
| 212 | LRRC47 | ENSG00000130764 | 7.593153185 | -2.219952288 | 27.35631877 | 5.06653E-06 | 0.000351464 | Yes |
| 213 | CDH15 | ENSG00000129910 | 3.365366067 | -4.217839118 | 24.31642738 | 5.27319E-06 | 0.000364074 | Yes |
| 214 | GMPR | ENSG00000137198 | 5.314746424 | -4.059177492 | 26.07223427 | 5.36389E-06 | 0.000368597 | Yes |
| 215 | CDA | ENSG00000158825 | 4.307801046 | -3.602378617 | 25.47308684 | 5.45705E-06 | 0.000373247 | Yes |
| 216 | MYOT | ENSG00000120729 | 5.644919078 | -5.018952767 | 25.59097266 | 5.62188E-06 | 0.000382732 | Yes |
| 217 | ASB15 | ENSG00000146809 | 4.298432252 | -5.393148383 | 24.08064759 | 5.71996E-06 | 0.000387607 | Yes |
| 218 | CA3 | ENSG00000164879 | 8.967498428 | -5.324777365 | 26.79259415 | 5.86553E-06 | 0.00039564 | Yes |
| 219 | SNTA1 | ENSG00000101400 | 7.310749262 | -3.25813206 | 26.65367879 | 6.12086E-06 | 0.000410968 | Yes |
| 220 | PPP1R3C | ENSG00000119938 | 4.92423171 | -3.609616485 | 25.57061219 | 6.24543E-06 | 0.000416461 | Yes |
| 221 | PROB1 | ENSG00000228672 | 3.758335209 | -3.853599142 | 24.40557024 | 6.25958E-06 | 0.000416461 | Yes |
| 222 | LINC00205 | ENSG00000223768 | 4.861705535 | 6.249957748 | 26.63110335 | 6.29279E-06 | 0.000416776 | Yes |
| 223 | APOBEC2 | ENSG00000124701 | 3.802410786 | -4.589742551 | 23.72787495 | 6.90418E-06 | 0.000455209 | Yes |
| 224 | NDUFS7 | ENSG00000115286 | 7.413587075 | -2.670979466 | 26.28635357 | 7.01174E-06 | 0.000460228 | Yes |
| 225 | TRIT1 | ENSG00000043514 | 5.205319434 | 5.990979193 | 25.95248386 | 7.14141E-06 | 0.000466647 | Yes |
| 226 | RBFOX1 | ENSG00000078328 | 5.388193614 | -4.276292875 | 25.12093161 | 7.24124E-06 | 0.000471067 | Yes |
| 227 | CRABP2 | ENSG00000143320 | 7.733253738 | 5.97081077 | 25.47313547 | 7.40519E-06 | 0.00047788 | Yes |
| 228 | CORO6 | ENSG00000167549 | 7.188904347 | -3.785916297 | 25.96820377 | 7.41127E-06 | 0.00047788 | Yes |
| 229 | GPX8 | ENSG00000164294 | 4.803094823 | 5.737620021 | 25.66150056 | 7.84853E-06 | 0.000503855 | Yes |
| 230 | P2RX5 | ENSG00000083454 | 4.068739999 | -4.166557012 | 23.81283563 | 7.97987E-06 | 0.00051005 | Yes |
| 231 | HMGB2 | ENSG00000164104 | 4.983795126 | 8.90596762 | 26.41433833 | 8.08151E-06 | 0.0005143 | Yes |
| 232 | POLR2I | ENSG00000105258 | 7.583049946 | -2.393472955 | 25.84777657 | 8.14199E-06 | 0.000515906 | Yes |
| 233 | LIG1 | ENSG00000105486 | 5.692729615 | 5.31241428 | 25.04206135 | 8.29355E-06 | 0.000523244 | Yes |
| 234 | NDUFB10 | ENSG00000140990 | 7.526373026 | -2.619946395 | 25.73764282 | 8.39282E-06 | 0.000527235 | Yes |
| 235 | ZNF559 | ENSG00000188321 | 4.814187582 | 8.717064003 | 26.2092641 | 8.46566E-06 | 0.000527911 | Yes |
| 236 | RAMP1 | ENSG00000132329 | 4.832241442 | -4.273902303 | 24.2665188 | 8.47572E-06 | 0.000527911 | Yes |
| 237 | CAND2 | ENSG00000144712 | 4.863422786 | -3.853293005 | 24.47491657 | 8.61948E-06 | 0.00053459 | Yes |
| 238 | GDF15 | ENSG00000130513 | 6.915350783 | 10.87944579 | 26.38383675 | 8.94294E-06 | 0.000552311 | Yes |
| 239 | YBX3 | ENSG00000060138 | 8.903213922 | -2.648656474 | 25.5306024 | 9.21027E-06 | 0.000566432 | Yes |
| 240 | MYL1 | ENSG00000168530 | 10.33321613 | -6.247295388 | 25.33295034 | 9.54953E-06 | 0.000584839 | Yes |
| 241 | BCL6 | ENSG00000113916 | 7.560204017 | -2.508540022 | 25.31600244 | 9.63844E-06 | 0.000587824 | Yes |
| 242 | NOS1 | ENSG00000089250 | 4.463804649 | -5.311635366 | 22.52265987 | 1.10805E-05 | 0.000672194 | Yes |
| 243 | PTP4A3 | ENSG00000184489 | 6.846983156 | -2.981280479 | 24.7577246 | 1.11137E-05 | 0.000672194 | Yes |
| 244 | TOX2 | ENSG00000124191 | 5.838969867 | 9.784477552 | 25.48687945 | 1.13715E-05 | 0.000684955 | Yes |
| 245 | PLK1 | ENSG00000166851 | 5.459078334 | 9.397106561 | 25.40939725 | 1.14203E-05 | 0.000685079 | Yes |
| 246 | SLC38A4 | ENSG00000139209 | 3.195935571 | -3.910422583 | 22.2980366 | 1.17068E-05 | 0.000699398 | Yes |
| 247 | HMGN1 | ENSG00000205581 | 6.859213519 | 3.681027475 | 24.36269031 | 1.17748E-05 | 0.000700599 | Yes |
| 248 | UQCRC1 | ENSG00000010256 | 7.59186345 | -2.07082594 | 24.14810749 | 1.41837E-05 | 0.000840512 | Yes |
| 249 | LINC01409 | ENSG00000237491 | 3.655658327 | -4.013113159 | 22.03946304 | 1.43585E-05 | 0.000847444 | Yes |
| 250 | CHST14 | ENSG00000169105 | 5.151341185 | 9.081186932 | 24.59730586 | 1.44484E-05 | 0.000849322 | Yes |
| 251 | ATP5F1D | ENSG00000099624 | 6.387673982 | -2.787466313 | 23.87122175 | 1.45977E-05 | 0.000854668 | Yes |
| 252 | IRAK4 | ENSG00000198001 | 4.447385505 | 6.585283742 | 24.131253 | 1.4787E-05 | 0.000862298 | Yes |
| 253 | EZH1 | ENSG00000108799 | 5.808853104 | -2.056471221 | 23.70715132 | 1.51858E-05 | 0.000882041 | Yes |
| 254 | ENSG00000274341 | ENSG00000274341 | 4.536165392 | -3.711410277 | 22.59444079 | 1.58612E-05 | 0.000917629 | Yes |
| 255 | NDUFS3 | ENSG00000213619 | 7.144090282 | -2.18193791 | 23.6791962 | 1.63488E-05 | 0.000942118 | Yes |
| 256 | GBP1 | ENSG00000117228 | 6.451198412 | 5.799373673 | 23.0160953 | 1.67157E-05 | 0.00095948 | Yes |
| 257 | KANSL1.AS1 | | 3.814204041 | -3.21976817 | 22.10779016 | 1.71233E-05 | 0.000979038 | Yes |
| 258 | HBA2 | ENSG00000188536 | 8.717385316 | -3.629664414 | 23.4992724 | 1.76622E-05 | 0.00100592 | Yes |
| 259 | NTMT1 | ENSG00000148335 | 5.175763383 | -3.099632512 | 22.786523 | 1.83166E-05 | 0.001039149 | Yes |
| 260 | PUS1 | ENSG00000177192 | 4.276911552 | 8.162966581 | 23.52310739 | 1.8794E-05 | 0.001061532 | Yes |
| 261 | TXNDC12 | ENSG00000117862 | 5.921274597 | 5.924096147 | 22.76680024 | 1.88562E-05 | 0.001061532 | Yes |
| 262 | PACSIN3 | ENSG00000165912 | 5.946698129 | -2.977328012 | 22.94582497 | 1.91246E-05 | 0.001072516 | Yes |
| 263 | DSN1 | ENSG00000149636 | 5.495648567 | 6.454064179 | 23.08872883 | 1.92126E-05 | 0.001073338 | Yes |
| 264 | TRAFD1 | ENSG00000135148 | 4.908477639 | 7.075857568 | 23.44316311 | 1.93263E-05 | 0.001075587 | Yes |
| 265 | FSD2 | ENSG00000186628 | 4.856076612 | -4.399804029 | 21.82358679 | 2.00914E-05 | 0.001113929 | Yes |
| 266 | AAMDC | ENSG00000087884 | 6.30620593 | -3.197341419 | 22.78538472 | 2.05772E-05 | 0.001136562 | Yes |
| 267 | COL11A2 | ENSG00000204248 | 4.495336023 | -4.042489201 | 21.58755022 | 2.15204E-05 | 0.001184187 | Yes |
| 268 | MAP4 | ENSG00000047849 | 8.274375701 | -1.996395228 | 22.85920317 | 2.20263E-05 | 0.001204382 | Yes |
| 269 | MCM2 | ENSG00000073111 | 6.580166587 | 4.463513041 | 22.29754247 | 2.20519E-05 | 0.001204382 | Yes |
| 270 | C19orf47 | ENSG00000160392 | 6.480090802 | -2.947594649 | 22.62519865 | 2.22299E-05 | 0.00120959 | Yes |
| 271 | AAMP | ENSG00000127837 | 6.62157876 | 3.258924969 | 22.47378416 | 2.2507E-05 | 0.00121571 | Yes |
| 272 | BEST3 | ENSG00000127325 | 3.957061429 | -4.202625327 | 21.008014 | 2.25085E-05 | 0.00121571 | Yes |
| 273 | PDLIM3 | ENSG00000154553 | 7.802072966 | -3.801645703 | 22.6614608 | 2.27559E-05 | 0.001224552 | Yes |
| 274 | CHCHD10 | ENSG00000250479 | 9.008458085 | -2.601506236 | 22.73944141 | 2.30267E-05 | 0.001234588 | Yes |
| 275 | DDN | ENSG00000181418 | 4.340569297 | -4.463764216 | 21.02871131 | 2.33266E-05 | 0.001246102 | Yes |
| 276 | ZBTB18 | ENSG00000179456 | 5.989890387 | -3.054725778 | 22.31797823 | 2.37287E-05 | 0.00126297 | Yes |
| 277 | ANKRD23 | ENSG00000163126 | 4.318701053 | -3.368237373 | 21.43768666 | 2.43875E-05 | 0.001289334 | Yes |
| 278 | ABHD15 | ENSG00000168792 | 4.528930811 | 6.673547819 | 22.64181919 | 2.44002E-05 | 0.001289334 | Yes |
| 279 | PINK1 | ENSG00000158828 | 6.904712687 | -2.612687618 | 22.22412301 | 2.62675E-05 | 0.001383014 | Yes |
| 280 | JPH1 | ENSG00000104369 | 3.711409444 | -4.050041448 | 20.41770053 | 2.722E-05 | 0.001428024 | Yes |
| 281 | NEXN | ENSG00000162614 | 7.461636755 | -3.139579901 | 22.01379871 | 2.84751E-05 | 0.001488538 | Yes |
| 282 | INTS9 | ENSG00000104299 | 4.161703923 | 8.038117436 | 22.00040512 | 3.07357E-05 | 0.001600993 | Yes |
| 283 | YIPF7 | ENSG00000177752 | 2.9213724 | -4.485650811 | 19.68094222 | 3.10917E-05 | 0.001613791 | Yes |
| 284 | ASB5 | ENSG00000164122 | 4.830592077 | -4.574462039 | 20.40541719 | 3.30095E-05 | 0.001707279 | Yes |
| 285 | ENSG00000272754 | ENSG00000272754 | 3.489304556 | -3.424580673 | 19.99208586 | 3.40102E-05 | 0.001752841 | Yes |
| 286 | ENSG00000250899 | ENSG00000250899 | 2.732270609 | -3.626691973 | 19.49910476 | 3.41542E-05 | 0.001754088 | Yes |
| 287 | MEF2C | ENSG00000081189 | 7.331990761 | -3.036927934 | 21.44572764 | 3.45429E-05 | 0.001767845 | Yes |
| 288 | TRIOBP | ENSG00000100106 | 7.565246837 | 3.154875009 | 21.35897305 | 3.51139E-05 | 0.001790808 | Yes |
| 289 | DDX41 | ENSG00000183258 | 6.353921701 | -1.929955154 | 21.2853841 | 3.59446E-05 | 0.001826808 | Yes |
| 290 | XIRP1 | ENSG00000168334 | 6.117834441 | -4.587221439 | 20.81567082 | 3.61218E-05 | 0.001829464 | Yes |
| 291 | CDCA4 | ENSG00000170779 | 5.121841254 | 9.055333758 | 21.75647869 | 3.6306E-05 | 0.001832453 | Yes |
| 292 | NOL4L | ENSG00000197183 | 4.421943099 | 6.5658908 | 21.42218145 | 3.652E-05 | 0.001836919 | Yes |
| 293 | TNNI3 | ENSG00000129991 | 2.982237216 | -3.834375419 | 19.37432942 | 3.68726E-05 | 0.001848303 | Yes |
| 294 | ENSG00000254343 | ENSG00000254343 | 3.152725807 | -3.494906576 | 19.53422605 | 3.70441E-05 | 0.001850564 | Yes |
| 295 | IFNAR2 | ENSG00000159110 | 5.474835155 | 4.243803907 | 20.67869876 | 3.7267E-05 | 0.001855365 | Yes |
| 296 | ITPRIPL1 | ENSG00000198885 | 4.02814066 | -3.380698973 | 20.08922473 | 3.78885E-05 | 0.001875402 | Yes |
| 297 | ACADS | ENSG00000122971 | 5.769083853 | -2.502531 | 20.99267017 | 3.79257E-05 | 0.001875402 | Yes |
| 298 | AFAP1 | ENSG00000196526 | 5.797213985 | 4.67421811 | 20.61531137 | 3.84764E-05 | 0.001896226 | Yes |
| 299 | FXR1 | ENSG00000114416 | 7.384744628 | -2.639382998 | 21.1461556 | 3.86314E-05 | 0.001897477 | Yes |
| 300 | SNRPF | ENSG00000139343 | 7.503339017 | 4.922777738 | 20.76294164 | 3.94828E-05 | 0.001932807 | Yes |
| 301 | NFE2L1 | ENSG00000082641 | 7.203699198 | -2.188595836 | 21.05726128 | 3.9881E-05 | 0.001945792 | Yes |
| 302 | GAL3ST4 | ENSG00000197093 | 4.906061561 | 7.115524512 | 21.18000024 | 4.12781E-05 | 0.002007265 | Yes |
| 303 | MYDGF | ENSG00000074842 | 6.446651897 | 4.879738025 | 20.34836373 | 4.31616E-05 | 0.00209191 | Yes |
| 304 | PHYH | ENSG00000107537 | 6.016879598 | -2.896839541 | 20.62084132 | 4.33762E-05 | 0.00209537 | Yes |
| 305 | RCC2 | ENSG00000179051 | 7.818651378 | 4.58155993 | 20.45572228 | 4.59535E-05 | 0.00221257 | Yes |
| 306 | GOT1 | ENSG00000120053 | 5.789196091 | -2.931498819 | 20.33215759 | 4.7182E-05 | 0.002257464 | Yes |
| 307 | COL4A4 | ENSG00000081052 | 2.55189063 | -3.983263812 | 18.66590489 | 4.71944E-05 | 0.002257464 | Yes |
| 308 | PGM2 | ENSG00000169299 | 4.147200052 | 8.019520686 | 20.51917315 | 5.08987E-05 | 0.002426723 | Yes |
| 309 | PCGF2 | ENSG00000277258 | 6.48178049 | 3.048924435 | 20.12708464 | 5.13749E-05 | 0.002441474 | Yes |
| 310 | IQCN | ENSG00000130518 | 5.092508845 | -3.705814534 | 19.49139396 | 5.58613E-05 | 0.002646092 | Yes |
| 311 | LPAR2 | ENSG00000064547 | 4.014414347 | 7.852818764 | 20.16354141 | 5.6584E-05 | 0.002668482 | Yes |
| 312 | BAHD1 | ENSG00000140320 | 4.105831159 | 7.981749414 | 20.19755166 | 5.66986E-05 | 0.002668482 | Yes |
| 313 | EHBP1L1 | ENSG00000173442 | 7.255725055 | -2.545246623 | 19.98838333 | 5.78409E-05 | 0.002713517 | Yes |
| 314 | PARP16 | ENSG00000138617 | 4.44794358 | 6.592000989 | 20.05941136 | 5.86914E-05 | 0.002744621 | Yes |
| 315 | LGI4 | ENSG00000153902 | 5.403711994 | -3.091853145 | 19.55123388 | 5.98571E-05 | 0.002790219 | Yes |
| 316 | PRR11 | ENSG00000068489 | 5.109275631 | 9.038096177 | 20.24514574 | 6.04508E-05 | 0.002808948 | Yes |
| 317 | SRD5A3 | ENSG00000128039 | 4.425184905 | 8.32817481 | 20.03402485 | 6.1988E-05 | 0.002871261 | Yes |
| 318 | ADAMTSL2 | ENSG00000197859 | 4.644518371 | -3.589182384 | 18.99333114 | 6.31249E-05 | 0.002914699 | Yes |
| 319 | PRRT3.AS1 | | 3.331098892 | -3.344286029 | 18.32818593 | 6.42966E-05 | 0.002959464 | Yes |
| 320 | PPP2R3A | ENSG00000073711 | 5.949392301 | -2.824061721 | 19.49075275 | 6.49158E-05 | 0.002978599 | Yes |
| 321 | TBC1D10A | ENSG00000099992 | 3.927167831 | 6.016511421 | 19.54292831 | 6.68287E-05 | 0.003056788 | Yes |
| 322 | SLC25A11 | ENSG00000108528 | 7.486467527 | -2.403742632 | 19.52797619 | 6.85729E-05 | 0.003126797 | Yes |
| 323 | H2AX | ENSG00000188486 | 3.975412097 | 6.080651348 | 19.46778301 | 6.91148E-05 | 0.003141721 | Yes |
| 324 | HECW2 | ENSG00000138411 | 4.286874711 | 8.172458126 | 19.46222616 | 7.47204E-05 | 0.003386015 | Yes |
| 325 | ODF2 | ENSG00000136811 | 5.332622868 | 4.852104315 | 18.77014063 | 7.60205E-05 | 0.003434298 | Yes |
| 326 | ENSG00000273058 | ENSG00000273058 | 4.098185289 | -3.616002353 | 18.20879474 | 7.70105E-05 | 0.003453936 | Yes |
| 327 | MDH2 | ENSG00000146701 | 7.842190305 | -1.984239119 | 19.22748179 | 7.70345E-05 | 0.003453936 | Yes |
| 328 | CENPW | ENSG00000203760 | 4.487050097 | 8.379900631 | 19.419206 | 7.71631E-05 | 0.003453936 | Yes |
| 329 | ENSG00000273373 | ENSG00000273373 | 4.306113163 | 8.189195104 | 19.33514189 | 7.82592E-05 | 0.003492317 | Yes |
| 330 | CDCA8 | ENSG00000134690 | 4.453199124 | 8.355243921 | 19.3500088 | 7.89136E-05 | 0.003510816 | Yes |
| 331 | MRPL49 | ENSG00000149792 | 5.704530817 | 4.017836508 | 18.62393933 | 8.07113E-05 | 0.003579914 | Yes |
| 332 | SLX4IP | ENSG00000149346 | 3.597197634 | 7.434098503 | 18.9491317 | 8.28865E-05 | 0.003665286 | Yes |
| 333 | LDLRAD3 | ENSG00000179241 | 5.072211062 | 5.855784518 | 18.77703467 | 8.41848E-05 | 0.003711486 | Yes |
| 334 | CDK1 | ENSG00000170312 | 4.900476956 | 7.115570323 | 19.10888561 | 8.48265E-05 | 0.003727818 | Yes |
| 335 | NUP54 | ENSG00000138750 | 4.359404178 | 6.539141728 | 18.99586065 | 8.50646E-05 | 0.003727818 | Yes |
| 336 | ENSG00000274124 | ENSG00000274124 | 2.899450308 | -4.096783778 | 17.27195278 | 8.59292E-05 | 0.003754465 | Yes |
| 337 | SNAI2 | ENSG00000019549 | 4.219219629 | 8.108457897 | 19.02178187 | 8.68389E-05 | 0.003782919 | Yes |
| 338 | EFNA5 | ENSG00000184349 | 6.094461676 | 3.879856585 | 18.4256267 | 8.93313E-05 | 0.003872135 | Yes |
| 339 | TMEM67 | ENSG00000164953 | 4.376045887 | 6.497606811 | 18.85282218 | 8.9416E-05 | 0.003872135 | Yes |
| 340 | ENSG00000271579 | ENSG00000271579 | 2.745311747 | -3.444504617 | 17.2396052 | 8.98218E-05 | 0.003878234 | Yes |
| 341 | SLC19A1 | ENSG00000173638 | 4.493938563 | 8.35691751 | 18.92724082 | 9.16391E-05 | 0.003945061 | Yes |
| 342 | TMEM91 | ENSG00000142046 | 4.068241848 | 5.360726258 | 18.5259079 | 9.23747E-05 | 0.003965071 | Yes |
| 343 | SCAMP5 | ENSG00000198794 | 3.810468174 | 7.643661735 | 18.69461894 | 9.32066E-05 | 0.003989081 | Yes |
| 344 | MEGF6 | ENSG00000162591 | 4.009257452 | -3.25171759 | 17.751756 | 9.47761E-05 | 0.004037036 | Yes |
| 345 | RNF41 | ENSG00000181852 | 5.679315125 | -2.035579169 | 18.47038957 | 9.48788E-05 | 0.004037036 | Yes |
| 346 | APH1A | ENSG00000117362 | 8.000964746 | 2.287929592 | 18.61471777 | 9.5459E-05 | 0.004049952 | Yes |
| 347 | LTO1 | ENSG00000149716 | 5.055950252 | 4.361974443 | 18.10957123 | 9.6093E-05 | 0.004065069 | Yes |
| 348 | CECR2 | ENSG00000099954 | 4.399462608 | -3.751133163 | 17.74626011 | 9.63898E-05 | 0.00406587 | Yes |
| 349 | CRAT | ENSG00000095321 | 6.987792699 | -2.223546144 | 18.50319361 | 9.83214E-05 | 0.004135434 | Yes |
| 350 | PTTG1 | ENSG00000164611 | 5.973721504 | 5.330940828 | 18.07296704 | 9.98145E-05 | 0.004186201 | Yes |
| 351 | TYMS | ENSG00000176890 | 5.866721585 | 4.341360078 | 18.0544808 | 0.000100124 | 0.004187171 | Yes |
| 352 | ENSG00000273007 | ENSG00000273007 | 3.191458708 | -3.742542846 | 17.02428974 | 0.000101553 | 0.004234842 | Yes |
| 353 | HMGCR | ENSG00000113161 | 4.915616105 | 5.543316048 | 18.1848828 | 0.000101869 | 0.004235942 | Yes |
| 354 | ENSG00000251396 | ENSG00000251396 | 3.079828439 | -3.973367478 | 16.92125998 | 0.00010216 | 0.004236019 | Yes |
| 355 | CDK17 | ENSG00000059758 | 4.815364116 | 8.726471997 | 18.66131111 | 0.000103201 | 0.004267112 | Yes |
| 356 | USP13 | ENSG00000058056 | 6.763487159 | -2.845998205 | 18.30904019 | 0.000103856 | 0.004273089 | Yes |
| 357 | ARRB2 | ENSG00000141480 | 7.00665908 | 3.502623149 | 18.21621102 | 0.000104177 | 0.004273089 | Yes |
| 358 | GYS1 | ENSG00000104812 | 6.761273441 | -2.262310592 | 18.32021792 | 0.000104421 | 0.004273089 | Yes |
| 359 | SORBS1 | ENSG00000095637 | 5.921553618 | -2.986822388 | 18.16854885 | 0.000104514 | 0.004273089 | Yes |
| 360 | USP54 | ENSG00000166348 | 6.257978494 | -2.861759465 | 18.21740402 | 0.000105111 | 0.004285541 | Yes |
| 361 | TNIP1 | ENSG00000145901 | 6.424333252 | -2.465530723 | 18.23534621 | 0.000106202 | 0.004317996 | Yes |
| 362 | MRPL13 | ENSG00000172172 | 4.248136899 | 8.141071277 | 18.38882109 | 0.00010921 | 0.004428 | Yes |
| 363 | RNF123 | ENSG00000164068 | 5.791597735 | -2.333769356 | 18.06602347 | 0.000110074 | 0.004450696 | Yes |
| 364 | MT.CO3 | | 14.58334268 | -1.782131056 | 18.27657748 | 0.00011067 | 0.004451533 | Yes |
| 365 | NHSL1 | ENSG00000135540 | 4.760841018 | 8.662073422 | 18.45330683 | 0.000110703 | 0.004451533 | Yes |
| 366 | NRGN | ENSG00000154146 | 5.094578316 | -2.995579821 | 17.80490682 | 0.000111935 | 0.004481766 | Yes |
| 367 | RILP | ENSG00000167705 | 5.063135116 | -2.773492302 | 17.82220837 | 0.000112067 | 0.004481766 | Yes |
| 368 | NUF2 | ENSG00000143228 | 5.598480213 | 9.542646033 | 18.51405039 | 0.000112765 | 0.004497387 | Yes |
| 369 | CCDC28B | ENSG00000160050 | 5.690674442 | -2.698064455 | 17.92391179 | 0.000114025 | 0.004535295 | Yes |
| 370 | ZNF703 | ENSG00000183779 | 7.964710782 | -1.908791982 | 18.08285708 | 0.000116767 | 0.004631753 | Yes |
| 371 | CHRNA1 | ENSG00000138435 | 4.137176319 | -3.876251166 | 17.07127471 | 0.000118349 | 0.00467059 | Yes |
| 372 | PHLDA1 | ENSG00000139289 | 6.075732584 | 4.809049878 | 17.61699273 | 0.000118384 | 0.00467059 | Yes |
| 373 | UTP6 | ENSG00000108651 | 5.112096429 | 3.446124709 | 17.53913068 | 0.000119793 | 0.004713476 | Yes |
| 374 | CAMK2A | ENSG00000070808 | 4.433904409 | -3.703627033 | 17.17846448 | 0.000122244 | 0.004796678 | Yes |
| 375 | GPR155 | ENSG00000163328 | 5.625380554 | -2.716359083 | 17.71830336 | 0.000122563 | 0.004796678 | Yes |
| 376 | ZFAS1 | ENSG00000177410 | 6.988026007 | 3.052617606 | 17.78657752 | 0.000124049 | 0.004841888 | Yes |
| 377 | WIPF3 | ENSG00000122574 | 3.959643541 | -3.689735314 | 16.90606405 | 0.000125314 | 0.004878266 | Yes |
| 378 | ASB8 | ENSG00000177981 | 5.375842179 | -2.470261412 | 17.61383096 | 0.000126389 | 0.004907058 | Yes |
| 379 | YPEL3 | ENSG00000090238 | 7.503645697 | -1.804571187 | 17.81257938 | 0.000128214 | 0.004964746 | Yes |
| 380 | TMED9 | ENSG00000184840 | 8.697082379 | 2.930075483 | 17.7979145 | 0.000129109 | 0.004986195 | Yes |
| 381 | TPX2 | ENSG00000088325 | 5.883409022 | 4.900514337 | 17.35157317 | 0.000130628 | 0.005019126 | Yes |
| 382 | TUBA4A | ENSG00000127824 | 5.859536381 | -3.45456646 | 17.50879223 | 0.000130647 | 0.005019126 | Yes |
| 383 | ZNF710.AS1 | | 4.200138605 | -2.939714159 | 17.09782156 | 0.000131137 | 0.005024745 | Yes |
| 384 | CADM2 | ENSG00000175161 | 2.162398423 | -4.059863644 | 16.38476257 | 0.00013206 | 0.005046915 | Yes |
| 385 | VANGL1 | ENSG00000173218 | 4.564251041 | 8.475428656 | 17.90831569 | 0.000133067 | 0.005072148 | Yes |
| 386 | ENDOG | ENSG00000167136 | 7.003733212 | -2.35209966 | 17.65974603 | 0.000133863 | 0.005089246 | Yes |
| 387 | PGM1 | ENSG00000079739 | 7.434388836 | -2.472284257 | 17.65822145 | 0.000134956 | 0.005117498 | Yes |
| 388 | CKS1B | ENSG00000173207 | 6.642365527 | 4.537840905 | 17.31970312 | 0.000136241 | 0.00515288 | Yes |
| 389 | OSTC | ENSG00000198856 | 6.682635049 | 3.327017249 | 17.45501084 | 0.000136993 | 0.005167968 | Yes |
| 390 | SLC25A35 | ENSG00000125434 | 4.082242757 | 7.961219625 | 17.70519334 | 0.000137846 | 0.005186778 | Yes |
| 391 | ZDHHC8 | ENSG00000099904 | 5.846179116 | 3.945867123 | 17.14476188 | 0.000142956 | 0.005365233 | Yes |
| 392 | SLC35F1 | ENSG00000196376 | 3.033474363 | -3.564414715 | 16.19962072 | 0.000144298 | 0.005401748 | Yes |
| 393 | HCFC1R1 | ENSG00000103145 | 7.139669227 | -2.254028922 | 17.4240059 | 0.000146673 | 0.005476661 | Yes |
| 394 | TM9SF2 | ENSG00000125304 | 7.440449807 | 3.170965726 | 17.35709051 | 0.000147531 | 0.005494667 | Yes |
| 395 | CREB3L1 | ENSG00000157613 | 7.274709737 | 5.26734599 | 17.08423077 | 0.000149456 | 0.005542302 | Yes |
| 396 | AGTRAP | ENSG00000177674 | 5.523854013 | 3.437403269 | 17.02383233 | 0.000149567 | 0.005542302 | Yes |
| 397 | IQGAP3 | ENSG00000183856 | 5.061274351 | 7.279999605 | 17.51555847 | 0.000151664 | 0.005605807 | Yes |
| 398 | CCNB2 | ENSG00000157456 | 4.021640756 | 7.896024263 | 17.42135829 | 0.000152119 | 0.005608488 | Yes |
| 399 | CNOT3 | ENSG00000088038 | 4.158257665 | 5.401568409 | 17.18597802 | 0.000152721 | 0.005616525 | Yes |
| 400 | TBX1 | ENSG00000184058 | 5.673855537 | -3.45957414 | 17.03231434 | 0.000154109 | 0.005653385 | Yes |
| 401 | MOSPD3 | ENSG00000106330 | 5.422248428 | 4.180775741 | 16.88423896 | 0.000154931 | 0.005669312 | Yes |
| 402 | FAHD2B | ENSG00000144199 | 4.005179002 | -3.216232829 | 16.51243267 | 0.00015606 | 0.005696395 | Yes |

| | A | B | C | D | E | F | G | H |
|---|---|---|---|---|---|---|---|---|
| 403 | TMEM14A | ENSG00000096092 | 4.631406623 | 6.783662647 | 17.35832382 | 0.000156896 | 0.005712663 | Yes |
| 404 | CENPK | ENSG00000123219 | 4.232540693 | 8.122542262 | 17.35464064 | 0.000158819 | 0.005768333 | Yes |
| 405 | ENSG00000272829 | ENSG00000272829 | 3.238799188 | -3.338707958 | 16.08521851 | 0.000160464 | 0.005813651 | Yes |
| 406 | RBBP8 | ENSG00000101773 | 5.577100657 | 5.379920309 | 16.81875658 | 0.000162743 | 0.005881656 | Yes |
| 407 | SFT2D1 | ENSG00000198818 | 3.675723566 | 5.751826144 | 17.03422393 | 0.000163483 | 0.005893843 | Yes |
| 408 | EPHB2 | ENSG00000133216 | 5.169013355 | 7.429444405 | 17.28781597 | 0.000165798 | 0.005948327 | Yes |
| 409 | CIPC | ENSG00000198894 | 4.088370787 | -2.775152567 | 16.50955178 | 0.000165807 | 0.005948327 | Yes |
| 410 | LMOD1 | ENSG00000163431 | 4.486396097 | -3.719993985 | 16.39825548 | 0.000167438 | 0.005987675 | Yes |
| 411 | SPEG | ENSG00000072195 | 5.688817397 | -3.082473011 | 16.86269026 | 0.000167722 | 0.005987675 | Yes |
| 412 | ADAR | ENSG00000160710 | 7.453774466 | 3.015525198 | 17.01465881 | 0.000168382 | 0.005996615 | Yes |
| 413 | EEF1A1 | ENSG00000156508 | 8.763454945 | 3.112425014 | 17.06160441 | 0.000169462 | 0.006020427 | Yes |
| 414 | ZNF865 | ENSG00000261221 | 5.632214589 | -2.239826692 | 16.89059985 | 0.000170027 | 0.006025863 | Yes |
| 415 | ZNF658 | ENSG00000274349 | 2.616941295 | -3.060803476 | 15.7525276 | 0.000170895 | 0.006042009 | Yes |
| 416 | ENSG00000275966 | ENSG00000275966 | 4.065033775 | -3.558875706 | 16.1732704 | 0.000175053 | 0.006165543 | Yes |
| 417 | HMGXB3 | ENSG00000113716 | 5.46067897 | 3.23620974 | 16.62230167 | 0.000175232 | 0.006165543 | Yes |
| 418 | HSPB3 | ENSG00000169271 | 4.009325916 | -3.680487034 | 16.01210779 | 0.000182642 | 0.006410868 | Yes |
| 419 | KIF1C | ENSG00000129250 | 8.184103187 | -2.378371812 | 16.82574745 | 0.000185962 | 0.006498981 | Yes |
| 420 | DPY19L4 | ENSG00000156162 | 4.726684567 | 5.113780503 | 16.51908517 | 0.000186228 | 0.006498981 | Yes |
| 421 | TENT4A | ENSG00000112941 | 4.893638383 | 4.727262157 | 16.4267259 | 0.000186484 | 0.006498981 | Yes |
| 422 | SLC39A9 | ENSG00000029364 | 6.129432574 | 3.429104968 | 16.53234347 | 0.000187673 | 0.006524861 | Yes |
| 423 | RABGGTB | ENSG00000137955 | 5.032328928 | 4.782979783 | 16.37391757 | 0.000190579 | 0.006593013 | Yes |
| 424 | CHRAC1 | ENSG00000104472 | 5.88361931 | 3.724837931 | 16.42193995 | 0.000190826 | 0.006593013 | Yes |
| 425 | AKAP6 | ENSG00000151320 | 3.8203276 | -3.223197856 | 15.94504474 | 0.000191017 | 0.006593013 | Yes |
| 426 | MSRB3 | ENSG00000174099 | 5.750273676 | -2.781330504 | 16.55131971 | 0.000191899 | 0.006593013 | Yes |
| 427 | PLAU | ENSG00000122861 | 5.760188693 | 9.709004636 | 17.03628973 | 0.000192338 | 0.006593013 | Yes |
| 428 | ENSG00000271959 | ENSG00000271959 | 1.886403841 | -4.056472753 | 15.7226348 | 0.000192559 | 0.006593013 | Yes |
| 429 | CSTB | ENSG00000160213 | 7.293303024 | 3.717164195 | 16.5907129 | 0.000192786 | 0.006593013 | Yes |
| 430 | RAD23A | ENSG00000179262 | 9.113976557 | -1.743623866 | 16.74094377 | 0.000193294 | 0.006594979 | Yes |
| 431 | CFL2 | ENSG00000165410 | 6.689715986 | -2.357565976 | 16.64803488 | 0.000193888 | 0.006599873 | Yes |
| 432 | CYB5R1 | ENSG00000159348 | 6.618862648 | -2.332665521 | 16.63474831 | 0.000194552 | 0.006607093 | Yes |
| 433 | TMEM39B | ENSG00000121775 | 4.761147466 | 6.077908705 | 16.62165492 | 0.000196523 | 0.006649324 | Yes |
| 434 | MAD2L1BP | ENSG00000124688 | 4.189273132 | 5.440444486 | 16.52634248 | 0.000196704 | 0.006649324 | Yes |
| 435 | LINC00106 | ENSG00000236871 | 6.404498206 | 4.253521337 | 16.34804543 | 0.000197959 | 0.006676313 | Yes |
| 436 | ADAMTS9.AS2 | | 5.33679017 | -3.531009588 | 16.26285492 | 0.000200231 | 0.006726906 | Yes |
| 437 | NAPG | ENSG00000134265 | 4.582623524 | 5.376883131 | 16.4170009 | 0.000200378 | 0.006726906 | Yes |
| 438 | SLC1A5 | ENSG00000105281 | 5.347807336 | 4.215501768 | 16.20536693 | 0.000201644 | 0.006753905 | Yes |
| 439 | CMTM7 | ENSG00000153551 | 6.190914255 | 3.24010421 | 16.36050321 | 0.000202886 | 0.006755409 | Yes |
| 440 | TMEM44 | ENSG00000145014 | 4.010135861 | 6.090966664 | 16.54497291 | 0.000203063 | 0.006755409 | Yes |
| 441 | SEC61G | ENSG00000132432 | 5.81360387 | 4.5986835 | 16.21196579 | 0.000203073 | 0.006755409 | Yes |
| 442 | RGMA | ENSG00000182175 | 5.816396296 | -2.454268855 | 16.40374259 | 0.000205683 | 0.006826723 | Yes |
| 443 | GLIS3 | ENSG00000107249 | 4.708344961 | 8.615027053 | 16.72675156 | 0.000206543 | 0.006828277 | Yes |
| 444 | LHPP | ENSG00000107902 | 4.368579631 | -2.773479562 | 16.05395206 | 0.000206663 | 0.006828277 | Yes |
| 445 | DNAJB6 | ENSG00000105993 | 6.499432925 | -2.116274114 | 16.46495215 | 0.000207343 | 0.006835302 | Yes |
| 446 | CD52 | ENSG00000169442 | 4.844695889 | -3.341279362 | 16.0324246 | 0.000211359 | 0.006952057 | Yes |
| 447 | HBA1 | ENSG00000206172 | 6.724375827 | -3.568198133 | 16.34170488 | 0.000212807 | 0.006983966 | Yes |
| 448 | SIPA1L2 | ENSG00000116991 | 4.789670019 | 6.138763782 | 16.36854758 | 0.000217041 | 0.007107013 | Yes |
| 449 | CKMT2 | ENSG00000131730 | 3.804765427 | -3.583619715 | 15.52551027 | 0.000217983 | 0.007121919 | Yes |
| 450 | ENSG00000257151 | ENSG00000257151 | 5.451511556 | -2.406636687 | 16.1984911 | 0.00021847 | 0.007121919 | Yes |
| 451 | CAPS | ENSG00000105519 | 6.759502743 | 8.942854707 | 16.59803786 | 0.000220815 | 0.007182364 | Yes |
| 452 | GJA1 | ENSG00000152661 | 6.800191526 | 4.439546015 | 16.09215596 | 0.000221465 | 0.007187562 | Yes |
| 453 | TFEB | ENSG00000112561 | 4.622253674 | -2.87251495 | 15.90203443 | 0.000223822 | 0.007247959 | Yes |
| 454 | RPS6KA1 | ENSG00000117676 | 3.815050743 | 7.670518157 | 16.22596034 | 0.000233072 | 0.007509559 | Yes |
| 455 | TWSG1 | ENSG00000128791 | 4.946918133 | 6.271501445 | 16.19028321 | 0.000233323 | 0.007509559 | Yes |
| 456 | POLR2B | ENSG00000047315 | 6.733549576 | 3.352134866 | 16.06022749 | 0.000233439 | 0.007509559 | Yes |
| 457 | MSTO1 | ENSG00000125459 | 4.1527447 | 4.200772585 | 15.78390676 | 0.000236004 | 0.007549004 | Yes |
| 458 | TMSB4X | ENSG00000205542 | 8.54588954 | 4.105102208 | 16.13476095 | 0.000236172 | 0.007549004 | Yes |
| 459 | FUNDC1 | ENSG00000069509 | 4.294826694 | 5.056740754 | 15.94810299 | 0.00023646 | 0.007549004 | Yes |
| 460 | ZNF70 | ENSG00000187792 | 3.912966356 | 5.191946072 | 15.99922477 | 0.000236728 | 0.007549004 | Yes |
| 461 | UBE2C | ENSG00000175063 | 5.873759374 | 5.095380499 | 15.81834625 | 0.000237893 | 0.007560869 | Yes |
| 462 | ENSG00000173727 | ENSG00000173727 | 3.0750619 | -2.716370263 | 15.22593924 | 0.000238134 | 0.007560869 | Yes |
| 463 | PIGL | ENSG00000108474 | 3.364852844 | 7.174664622 | 16.01696727 | 0.000239991 | 0.007602216 | Yes |
| 464 | ERI1 | ENSG00000104626 | 3.917313459 | 7.778784674 | 16.16645991 | 0.000240636 | 0.007602216 | Yes |
| 465 | EXT1 | ENSG00000182197 | 5.928515567 | 4.328784103 | 15.79135107 | 0.000240994 | 0.007602216 | Yes |
| 466 | FAM120C | ENSG00000184083 | 4.973702611 | 5.037387915 | 15.80274109 | 0.000242236 | 0.007624954 | Yes |
| 467 | PLEKHG4 | ENSG00000196155 | 6.398346135 | 3.992616727 | 15.82248422 | 0.000245368 | 0.007706987 | Yes |
| 468 | MICOS13 | ENSG00000174917 | 5.897856768 | -2.168686451 | 15.94608237 | 0.000247526 | 0.00774378 | Yes |
| 469 | METTL26 | ENSG00000130731 | 6.296380905 | -2.127668896 | 15.97956921 | 0.000248057 | 0.00774378 | Yes |
| 470 | AURKB | ENSG00000178999 | 5.642946789 | 9.589867131 | 16.33174977 | 0.000248127 | 0.00774378 | Yes |
| 471 | TMEM143 | ENSG00000161558 | 5.135939171 | -2.763857514 | 15.73162923 | 0.000253162 | 0.007875192 | Yes |
| 472 | UQCRQ | ENSG00000164405 | 8.223507748 | -2.104628091 | 16.0096294 | 0.000253414 | 0.007875192 | Yes |
| 473 | MHENCR | ENSG00000232442 | 3.846618044 | 5.909438049 | 15.91224619 | 0.000254351 | 0.007887587 | Yes |
| 474 | MYLK3 | ENSG00000140795 | 3.74913123 | -4.169502481 | 14.9374702 | 0.000256827 | 0.007947511 | Yes |
| 475 | PDCL | ENSG00000136940 | 3.830899542 | 5.925300941 | 15.81644596 | 0.000264257 | 0.008160181 | Yes |
| 476 | SYTL3 | ENSG00000164674 | 4.401507763 | -3.141225884 | 15.37762097 | 0.000265248 | 0.008161721 | Yes |
| 477 | CENPE | ENSG00000138778 | 4.191395463 | 6.31802908 | 15.88041308 | 0.000265422 | 0.008161721 | Yes |
| 478 | GPSM1 | ENSG00000160360 | 6.156242274 | 4.97558201 | 15.52046703 | 0.000268185 | 0.008229415 | Yes |
| 479 | MAGOHB | ENSG00000111196 | 4.946370875 | 4.724345815 | 15.47099479 | 0.000271714 | 0.008320247 | Yes |
| 480 | CDKN2A | ENSG00000147889 | 8.044124471 | 5.689761024 | 15.5489783 | 0.000275841 | 0.008425874 | Yes |
| 481 | BCAM | ENSG00000187244 | 5.821106795 | -2.065656392 | 15.65841199 | 0.000276339 | 0.008425874 | Yes |
| 482 | ENSG00000268858 | ENSG00000268858 | 3.709753302 | 5.782691162 | 15.6641928 | 0.00027689 | 0.008425874 | Yes |
| 483 | ZNF556 | ENSG00000172000 | 3.673503106 | -3.312304388 | 14.95632927 | 0.000278588 | 0.008459949 | Yes |
| 484 | OAT | ENSG00000065154 | 5.285083242 | 4.979665656 | 15.41243721 | 0.000279813 | 0.008479545 | Yes |
| 485 | IGFBP6 | ENSG00000167779 | 8.140123336 | -2.372398876 | 15.73034796 | 0.000281459 | 0.00849973 | Yes |
| 486 | HBB | ENSG00000244734 | 8.712262451 | -2.790844564 | 15.73572818 | 0.00028164 | 0.00849973 | Yes |
| 487 | ANAPC16 | ENSG00000166295 | 7.427217218 | -1.980805927 | 15.69143305 | 0.000283933 | 0.008550081 | Yes |
| 488 | KIF11 | ENSG00000138160 | 3.901925794 | 7.759180791 | 15.71771928 | 0.000284976 | 0.008550081 | Yes |
| 489 | ENSG00000269318 | ENSG00000269318 | 3.698253387 | -3.195029004 | 14.94055306 | 0.00028554 | 0.008550081 | Yes |
| 490 | LBR | ENSG00000143815 | 5.158992431 | 3.700177223 | 15.31438678 | 0.000286211 | 0.008550081 | Yes |
| 491 | STMN1 | ENSG00000117632 | 7.724734989 | 3.676229021 | 15.60385697 | 0.000286229 | 0.008550081 | Yes |
| 492 | IFT57 | ENSG00000114446 | 5.432278143 | 6.807557734 | 15.66607817 | 0.000290182 | 0.008650502 | Yes |
| 493 | HOXC.AS1 | | 4.375639709 | -2.96430367 | 15.15879519 | 0.000292256 | 0.008694612 | Yes |
| 494 | IFI44L | ENSG00000137959 | 7.309630852 | 4.640144632 | 15.41418218 | 0.000293523 | 0.008714591 | Yes |
| 495 | ZMYM3 | ENSG00000147130 | 5.890363012 | 2.971596461 | 15.38960752 | 0.000294345 | 0.008721315 | Yes |
| 496 | ADSS2 | ENSG00000035687 | 4.344061363 | 6.488736431 | 15.60408776 | 0.000298128 | 0.00881237 | Yes |
| 497 | SS18 | ENSG00000141380 | 6.522420485 | 2.828593497 | 15.44467143 | 0.000298622 | 0.00881237 | Yes |
| 498 | PPP1R21 | ENSG00000162869 | 3.60092123 | 5.655159802 | 15.41449297 | 0.000301675 | 0.008884545 | Yes |
| 499 | ENSG00000221990 | ENSG00000221990 | 3.355386845 | -2.899427139 | 14.73616562 | 0.000302324 | 0.008885771 | Yes |
| 500 | FBP2 | ENSG00000130957 | 4.167056866 | -4.201505554 | 14.69706653 | 0.000303332 | 0.008897523 | Yes |
| 501 | LIN7B | ENSG00000104863 | 4.628427885 | -2.866826853 | 15.13790255 | 0.00030463 | 0.008917728 | Yes |
| 502 | PRKCQ | ENSG00000065675 | 3.971535552 | -4.021170558 | 14.62437929 | 0.000309364 | 0.009038151 | Yes |
| 503 | AKT1 | ENSG00000142208 | 6.606494517 | 3.302341508 | 15.32090624 | 0.000309978 | 0.009038151 | Yes |
| 504 | ENSG00000281325 | ENSG00000281325 | 3.986987162 | 6.062498755 | 15.42556548 | 0.000310968 | 0.009048977 | Yes |
| 505 | HINT2 | ENSG00000137133 | 5.953958117 | -1.853221149 | 15.33991875 | 0.000315294 | 0.009156653 | Yes |
| 506 | SYCP2 | ENSG00000196074 | 4.262799769 | 8.14590562 | 15.50324669 | 0.000318046 | 0.009218309 | Yes |
| 507 | RRAD | ENSG00000166592 | 6.409031477 | -3.382304252 | 15.27654371 | 0.000319035 | 0.009228697 | Yes |
| 508 | SPIN4 | ENSG00000186767 | 5.604863931 | 4.953911533 | 15.0486395 | 0.000322507 | 0.009310717 | Yes |
| 509 | MT.ATP6 | | 13.23292553 | -1.932967571 | 15.38923522 | 0.000325141 | 0.009368273 | Yes |
| 510 | SH3BP1 | ENSG00000100092 | 5.005903708 | 4.147626307 | 14.96831147 | 0.000327808 | 0.009426585 | Yes |
| 511 | AP1AR | ENSG00000138660 | 3.909008959 | 7.757118204 | 15.33928008 | 0.000329473 | 0.009455875 | Yes |
| 512 | PPP2R3B | ENSG00000167393 | 6.675793815 | -2.236565813 | 15.19301204 | 0.000339189 | 0.009715677 | Yes |
| 513 | PYCR1 | ENSG00000183010 | 5.837036888 | 3.589314391 | 14.96144453 | 0.000340195 | 0.009725459 | Yes |
| 514 | EAF1 | ENSG00000144597 | 3.640085648 | 5.700828678 | 15.0955765 | 0.000343198 | 0.009774811 | Yes |
| 515 | CCNC | ENSG00000112237 | 4.942663509 | 3.383527344 | 14.8506178 | 0.000343257 | 0.009774811 | Yes |
| 516 | CLTCL1 | ENSG00000070371 | 3.983366234 | -2.972562819 | 14.60064731 | 0.00034901 | 0.009902538 | Yes |
| 517 | SLC25A22 | ENSG00000177542 | 3.98920164 | 5.279839086 | 15.02121015 | 0.000349131 | 0.009902538 | Yes |
| 518 | RASD2 | ENSG00000100302 | 3.199193092 | -2.931835355 | 14.3206425 | 0.000349772 | 0.009902538 | Yes |
| 519 | MYD88 | ENSG00000172936 | 3.778293816 | 7.6389185 | 15.13806159 | 0.000352762 | 0.009967902 | Yes |
| 520 | MIR34AHG | ENSG00000228526 | 4.015305732 | 7.884844644 | 15.17956811 | 0.000353778 | 0.00997736 | Yes |
| 521 | MT1F | ENSG00000198417 | 3.945367684 | 6.047651818 | 15.03858582 | 0.000360641 | 0.010132187 | Yes |
| 522 | FOXM1 | ENSG00000111206 | 3.96012215 | 7.821398854 | 15.11774324 | 0.000360652 | 0.010132187 | Yes |
| 523 | LTV1 | ENSG00000135521 | 5.494309412 | 4.851992979 | 14.75264395 | 0.000362726 | 0.010163488 | Yes |
| 524 | ARRDC2 | ENSG00000105643 | 6.092134393 | -2.533254054 | 14.96199024 | 0.000363524 | 0.010163488 | Yes |
| 525 | NCS1 | ENSG00000107130 | 3.867597234 | 7.738003302 | 15.07570083 | 0.000364266 | 0.010163488 | Yes |
| 526 | KPNA2 | ENSG00000182481 | 4.179133277 | 8.071573407 | 15.13465804 | 0.000364544 | 0.010163488 | Yes |
| 527 | LINC00174 | ENSG00000179406 | 3.596214924 | 7.391551942 | 14.9713506 | 0.000368384 | 0.010230796 | Yes |
| 528 | IMPA2 | ENSG00000141401 | 5.372502712 | -2.514609005 | 14.83942813 | 0.000369139 | 0.010230796 | Yes |
| 529 | OSR2 | ENSG00000164920 | 6.536252093 | 6.303434361 | 14.7671021 | 0.000369515 | 0.010230796 | Yes |
| 530 | MSS51 | ENSG00000166343 | 3.44604758 | -3.194182828 | 14.22820299 | 0.000369754 | 0.010230796 | Yes |
| 531 | HJURP | ENSG00000123485 | 3.803647937 | 7.646687058 | 15.00911997 | 0.000371112 | 0.010246712 | Yes |
| 532 | ENSG00000261326 | ENSG00000261326 | 4.572784879 | 8.449440092 | 15.14010786 | 0.000371729 | 0.010246712 | Yes |
| 533 | MUL1 | ENSG00000090432 | 5.539268698 | -2.333456448 | 14.81032366 | 0.000378635 | 0.010417443 | Yes |
| 534 | CLPB | ENSG00000162129 | 2.731534419 | -2.844520787 | 13.94780374 | 0.000387693 | 0.010646638 | Yes |
| 535 | GATA6.AS1 | | 3.940662986 | 7.788691135 | 14.91865524 | 0.00038864 | 0.010652654 | Yes |
| 536 | INPP5E | ENSG00000148384 | 4.33200302 | 6.440924021 | 14.86201877 | 0.000396286 | 0.010841954 | Yes |
| 537 | LRRC39 | ENSG00000122477 | 3.95112122 | -3.460463607 | 14.17767487 | 0.000397873 | 0.010865046 | Yes |
| 538 | TLE5 | ENSG00000104964 | 8.726072832 | -1.622239811 | 14.81544231 | 0.000402805 | 0.010979254 | Yes |
| 539 | RHNO1 | ENSG00000171792 | 4.088530866 | 5.393735739 | 14.67041545 | 0.000403912 | 0.010988966 | Yes |
| 540 | NOTCH2 | ENSG00000134250 | 6.434454671 | 4.168059496 | 14.53663839 | 0.000407185 | 0.01105745 | Yes |
| 541 | MXRA5 | ENSG00000101825 | 7.989033254 | 4.940952169 | 14.62582776 | 0.000408036 | 0.011060032 | Yes |
| 542 | CCDC69 | ENSG00000198624 | 6.211697996 | -2.922479655 | 14.65415074 | 0.000409008 | 0.011065894 | Yes |
| 543 | EYA3 | ENSG00000158161 | 5.097615187 | 3.766482663 | 14.40193059 | 0.000413022 | 0.01115389 | Yes |
| 544 | ENSG00000203709 | ENSG00000203709 | 4.0263997 | -2.683770304 | 14.24102448 | 0.000414692 | 0.01117836 | Yes |
| 545 | TUBB | ENSG00000196230 | 10.71558579 | 2.195419516 | 14.72605898 | 0.000419234 | 0.011269381 | Yes |
| 546 | CLPTM1L | ENSG00000049656 | 6.448429779 | 3.238252072 | 14.5434215 | 0.000419609 | 0.011269381 | Yes |
| 547 | FAM120B | ENSG00000112584 | 4.732204211 | 4.186313753 | 14.34855572 | 0.000420538 | 0.011273659 | Yes |
| 548 | CYB5B | ENSG00000103018 | 5.554630475 | 2.880920548 | 14.44982956 | 0.000421541 | 0.011279891 | Yes |
| 549 | WHRN | ENSG00000095397 | 3.187247884 | 6.95395889 | 14.50477317 | 0.000423883 | 0.011301957 | Yes |
| 550 | DCLRE1B | ENSG00000118655 | 3.862908413 | 7.677892627 | 14.67447183 | 0.00042391 | 0.011301957 | Yes |
| 551 | NDUFB9 | ENSG00000147684 | 7.783718551 | -2.227081467 | 14.64284201 | 0.000427689 | 0.011365782 | Yes |
| 552 | FAH | ENSG00000103876 | 5.848245535 | -2.412084992 | 14.53180657 | 0.000427857 | 0.011365782 | Yes |
| 553 | P2RX6 | ENSG00000099957 | 2.812607242 | -3.229132449 | 13.67514371 | 0.00042938 | 0.011385583 | Yes |
| 554 | SLCO5A1 | ENSG00000137571 | 2.622565289 | -3.920575645 | 13.57006953 | 0.000431359 | 0.011417372 | Yes |
| 555 | CD109 | ENSG00000156535 | 5.224447515 | 9.160871113 | 14.80521408 | 0.000434774 | 0.011486968 | Yes |
| 556 | LOC107985020 | ENSG00000274565 | 4.435027671 | 8.338725027 | 14.69210537 | 0.000439147 | 0.011581609 | Yes |
| 557 | POGLUT3 | ENSG00000178202 | 4.547867807 | 5.377450054 | 14.4107557 | 0.000441178 | 0.011593407 | Yes |
| 558 | PHGDH | ENSG00000092621 | 5.529586238 | 3.225435608 | 14.30264042 | 0.000441936 | 0.011593407 | Yes |
| 559 | SPCS2 | ENSG00000118363 | 4.20383715 | 4.964438693 | 14.35703967 | 0.000442158 | 0.011593407 | Yes |
| 560 | MRPL55 | ENSG00000162910 | 6.490268672 | -1.975527022 | 14.50152867 | 0.00044412 | 0.011593407 | Yes |
| 561 | SCN3B | ENSG00000166257 | 2.583366525 | -3.077513825 | 13.55506076 | 0.000444578 | 0.011593407 | Yes |
| 562 | KIF4A | ENSG00000090889 | 4.999186326 | 8.928925277 | 14.72220634 | 0.000445366 | 0.011593407 | Yes |
| 563 | SCRN1 | ENSG00000136193 | 4.736871781 | 5.299081168 | 14.34208916 | 0.000446044 | 0.011593407 | Yes |
| 564 | FILIP1 | ENSG00000118407 | 6.1553034 | -2.483916095 | 14.44971143 | 0.00044639 | 0.011593407 | Yes |
| 565 | CNOT9 | ENSG00000144580 | 5.907009865 | 3.771416069 | 14.2800857 | 0.000446723 | 0.011593407 | Yes |
| 566 | AFAP1L1 | ENSG00000157510 | 4.484977567 | -2.705835942 | 14.16993049 | 0.000449512 | 0.01163905 | Yes |
| 567 | PAN2 | ENSG00000135473 | 4.937757876 | 5.747449156 | 14.38823046 | 0.000450072 | 0.01163905 | Yes |
| 568 | EXTL1 | ENSG00000158008 | 2.281935743 | -3.545483167 | 13.44454572 | 0.000455448 | 0.0117573 | Yes |
| 569 | APP | ENSG00000142192 | 8.662981749 | 2.175822426 | 14.47897236 | 0.000457836 | 0.01179621 | Yes |
| 570 | MKNK2 | ENSG00000099875 | 8.83077436 | -2.057784446 | 14.48338589 | 0.000458597 | 0.01179621 | Yes |
| 571 | SRSF9 | ENSG00000111786 | 7.673966994 | 2.642891054 | 14.43534077 | 0.000459373 | 0.01179621 | Yes |
| 572 | MADCAM1 | ENSG00000099866 | 2.31964399 | -3.217604777 | 13.42047106 | 0.00046142 | 0.011828027 | Yes |
| 573 | CDR2L | ENSG00000109089 | 4.442832524 | 5.365629916 | 14.29381849 | 0.000463356 | 0.011856882 | Yes |
| 574 | USP49 | ENSG00000164663 | 3.804034549 | 7.665032187 | 14.43594724 | 0.000465062 | 0.011879767 | Yes |
| 575 | MAP9 | ENSG00000164114 | 5.585865961 | 4.568389734 | 14.09805794 | 0.000472206 | 0.012041261 | Yes |
| 576 | UCKL1 | ENSG00000198276 | 5.748364352 | 3.17467484 | 14.14668804 | 0.000476919 | 0.012140286 | Yes |
| 577 | NEMP1 | ENSG00000166881 | 3.429141161 | 7.213764941 | 14.25468871 | 0.000480955 | 0.012197041 | Yes |
| 578 | SLC39A11 | ENSG00000133195 | 4.979964577 | 6.341684742 | 14.3317732 | 0.000481161 | 0.012197041 | Yes |
| 579 | FOXK1 | ENSG00000164916 | 6.20214295 | 2.596817051 | 14.21933083 | 0.000481649 | 0.012197041 | Yes |
| 580 | ADIRF | ENSG00000148671 | 7.700042921 | -2.545822451 | 14.30743173 | 0.00048679 | 0.012289358 | Yes |
| 581 | KICS2 | ENSG00000174206 | 4.229402546 | 8.100972332 | 14.39252377 | 0.000486973 | 0.012289358 | Yes |
| 582 | EML4 | ENSG00000143924 | 5.855143575 | 3.352443966 | 14.08870737 | 0.000488037 | 0.012294994 | Yes |
| 583 | ZWILCH | ENSG00000174442 | 4.85861703 | 3.640963838 | 13.96330712 | 0.000491292 | 0.012355736 | Yes |
| 584 | NDUFB2 | ENSG00000090266 | 7.133895688 | -1.807615764 | 14.26735157 | 0.000493162 | 0.012370865 | Yes |
| 585 | CD99L2 | ENSG00000102181 | 5.939294768 | -2.285571836 | 14.18745313 | 0.000493584 | 0.012370865 | Yes |
| 586 | ST20 | ENSG00000180953 | 3.404876314 | 7.226183541 | 14.18331752 | 0.000495448 | 0.012396356 | Yes |
| 587 | ARHGAP11A | ENSG00000198826 | 4.61506662 | 8.52960679 | 14.38587636 | 0.000499081 | 0.012465961 | Yes |
| 588 | RRM2 | ENSG00000171848 | 4.675911568 | 8.591187062 | 14.38747972 | 0.000500119 | 0.012470597 | Yes |
| 589 | CNKSR1 | ENSG00000142675 | 4.226077412 | -2.710586531 | 13.82780855 | 0.000503889 | 0.01253944 | Yes |
| 590 | MCRIP2 | ENSG00000172366 | 4.485118042 | -2.657041844 | 13.88627338 | 0.000504988 | 0.01253944 | Yes |
| 591 | GPC1 | ENSG00000063660 | 7.07024529 | -2.352631772 | 14.19469843 | 0.00050545 | 0.01253944 | Yes |
| 592 | MAD2L1 | ENSG00000164109 | 3.780822161 | 7.62650997 | 14.21027901 | 0.000506891 | 0.012553927 | Yes |
| 593 | EFHD2 | ENSG00000142634 | 6.663758614 | 3.247313153 | 14.07687868 | 0.000510122 | 0.01261259 | Yes |
| 594 | FYCO1 | ENSG00000163820 | 4.343192903 | -2.510337521 | 13.83741925 | 0.000513094 | 0.012664684 | Yes |
| 595 | SIRT2 | ENSG00000068903 | 5.623744293 | -2.193713103 | 14.05950884 | 0.000514272 | 0.012672391 | Yes |
| 596 | SLC25A5 | ENSG00000005022 | 6.001392991 | 2.70431875 | 14.01370088 | 0.000517316 | 0.012710974 | Yes |
| 597 | AICDA | ENSG00000111732 | 3.026910017 | -3.408523755 | 13.27925876 | 0.000517575 | 0.012710974 | Yes |
| 598 | CSNK1G1 | ENSG00000169118 | 4.524505582 | 4.563095825 | 13.83254334 | 0.000527415 | 0.012930938 | Yes |
| 599 | PAPSS1 | ENSG00000138801 | 5.367004293 | 3.462740239 | 13.82905638 | 0.000529369 | 0.012957149 | Yes |
| 600 | MYH7B | ENSG00000078814 | 4.367130387 | -3.552387011 | 13.58205769 | 0.000533452 | 0.013035287 | Yes |
| 601 | STT3A | ENSG00000134910 | 6.177952312 | 2.780634293 | 13.9412605 | 0.000535429 | 0.01306178 | Yes |
| 602 | BMP1 | ENSG00000168487 | 7.038898315 | 2.857763549 | 13.99954169 | 0.000537528 | 0.013091187 | Yes |
| 603 | NDUFV2 | ENSG00000178127 | 7.124709722 | -1.900004712 | 14.027827 | 0.000542007 | 0.013178343 | Yes |
| 604 | SUGP1 | ENSG00000105705 | 5.328078596 | -2.045486107 | 13.88827155 | 0.000545485 | 0.0132409 | Yes |
| 605 | CASP7 | ENSG00000165806 | 4.261673251 | 4.618252251 | 13.76415785 | 0.000547528 | 0.013258707 | Yes |
| 606 | FSCN1 | ENSG00000075618 | 8.745569319 | 2.757920947 | 14.017455 | 0.000548223 | 0.013258707 | Yes |
| 607 | CCNA2 | ENSG00000145386 | 4.249889743 | 8.14399497 | 14.09316374 | 0.000548936 | 0.013258707 | Yes |
| 608 | ADCY1 | ENSG00000164742 | 4.572011336 | 8.478874819 | 14.12683546 | 0.000550848 | 0.013282979 | Yes |
| 609 | NDUFS8 | ENSG00000110717 | 6.372976835 | -2.186721231 | 13.92616329 | 0.000555611 | 0.013364373 | Yes |
| 610 | COP1 | ENSG00000143207 | 4.74717677 | 4.802091067 | 13.72053341 | 0.000556397 | 0.013364373 | Yes |
| 611 | CDK7 | ENSG00000134058 | 3.473613548 | 7.268653756 | 13.90067853 | 0.000556963 | 0.013364373 | Yes |
| 612 | ZNF404 | ENSG00000176222 | 3.782916628 | 7.610776577 | 13.94876956 | 0.000561666 | 0.013454421 | Yes |
| 613 | SRSF1 | ENSG00000136450 | 6.095616635 | 2.822197918 | 13.80791917 | 0.000562554 | 0.013454421 | Yes |
| 614 | UQCRHL | ENSG00000233954 | 3.337979496 | -2.800079861 | 13.29361197 | 0.000564145 | 0.013456479 | Yes |
| 615 | ENSG00000283013 | ENSG00000283013 | 3.977256485 | -2.873866525 | 13.46827188 | 0.000564761 | 0.013456479 | Yes |
| 616 | STIL | ENSG00000123473 | 3.615622431 | 7.447679241 | 13.90076181 | 0.000565398 | 0.013456479 | Yes |
| 617 | ENSG00000260563 | ENSG00000260563 | 4.640966655 | 4.139156193 | 13.61915634 | 0.000566863 | 0.013469448 | Yes |
| 618 | ZNF254 | ENSG00000213096 | 3.074962699 | 6.84642088 | 13.74574413 | 0.000569241 | 0.01350401 | Yes |
| 619 | PHETA2 | ENSG00000177096 | 4.992712395 | 4.739595478 | 13.63843598 | 0.000571421 | 0.013526587 | Yes |
| 620 | TRAPPC13 | ENSG00000113597 | 3.78476014 | 5.090941336 | 13.75582182 | 0.000572135 | 0.013526587 | Yes |
| 621 | ATP10A | ENSG00000206190 | 3.507444858 | 7.318248617 | 13.84086527 | 0.000572965 | 0.013526587 | Yes |
| 622 | GOLM1 | ENSG00000135052 | 5.950045602 | 4.540994771 | 13.63321033 | 0.000575125 | 0.013555723 | Yes |
| 623 | WDR35 | ENSG00000118965 | 2.933295133 | 6.674196183 | 13.668195 | 0.000576619 | 0.013569088 | Yes |
| 624 | PCDH19 | ENSG00000165194 | 4.122834768 | 7.987261166 | 13.93527282 | 0.000578981 | 0.013597557 | Yes |
| 625 | MIR1915HG | ENSG00000204682 | 3.33657389 | 7.139977296 | 13.77257746 | 0.000579687 | 0.013597557 | Yes |
| 626 | PLK3 | ENSG00000173846 | 4.668546424 | 4.358286331 | 13.56958534 | 0.000582241 | 0.013635623 | Yes |
| 627 | CCNB1 | ENSG00000134057 | 4.102174445 | 6.227200072 | 13.83311539 | 0.000587074 | 0.013726846 | Yes |
| 628 | UNC50 | ENSG00000115446 | 4.282392788 | 4.515413889 | 13.56546904 | 0.000589636 | 0.013759458 | Yes |
| 629 | IFI6 | ENSG00000126709 | 8.963131944 | 2.966693756 | 13.82920212 | 0.000591203 | 0.013759458 | Yes |
| 630 | ZBTB41 | ENSG00000177888 | 4.118349686 | 4.875613062 | 13.63099208 | 0.000591289 | 0.013759458 | Yes |
| 631 | CDCA7L | ENSG00000164649 | 3.901000083 | 5.111759898 | 13.67381987 | 0.000593028 | 0.013778017 | Yes |
| 632 | TBPL1 | ENSG00000028839 | 3.47798725 | 7.310396359 | 13.73666194 | 0.000597091 | 0.013835922 | Yes |
| 633 | BTN3A3 | ENSG00000111801 | 5.236473082 | 3.304172618 | 13.5300343 | 0.000597411 | 0.013835922 | Yes |
| 634 | SRPX | ENSG00000101955 | 7.537379457 | 4.225919121 | 13.66969737 | 0.00060301 | 0.013936116 | Yes |
| 635 | EZH2 | ENSG00000106462 | 6.247958148 | 2.552738967 | 13.66613062 | 0.000603641 | 0.013936116 | Yes |
| 636 | ELOVL6 | ENSG00000170522 | 4.257617835 | 4.963994492 | 13.56854185 | 0.00060838 | 0.013972129 | Yes |
| 637 | HSF1 | ENSG00000185122 | 6.418788637 | -1.768793603 | 13.70492365 | 0.000610063 | 0.013972129 | Yes |
| 638 | SATB1 | ENSG00000182568 | 5.461549504 | -2.194424096 | 13.61697371 | 0.000610838 | 0.013972129 | Yes |
| 639 | USP2 | ENSG00000036672 | 3.351372413 | -3.193120218 | 13.05264938 | 0.000611439 | 0.013972129 | Yes |
| 640 | ZMYM1 | ENSG00000197056 | 2.927599247 | 6.666662156 | 13.52328033 | 0.000611483 | 0.013972129 | Yes |
| 641 | PABPC1L | ENSG00000101104 | 5.693372394 | 3.732556036 | 13.49275861 | 0.000612183 | 0.013972129 | Yes |
| 642 | ARL4D | ENSG00000175906 | 5.426244388 | 5.84609006 | 13.57645283 | 0.000612256 | 0.013972129 | Yes |
| 643 | MANF | ENSG00000145050 | 5.859373802 | 3.398108214 | 13.52883221 | 0.000612838 | 0.013972129 | Yes |
| 644 | TPI1 | ENSG00000111669 | 8.332973867 | -1.793518988 | 13.72630138 | 0.000617828 | 0.014063985 | Yes |
| 645 | SURF4 | ENSG00000148248 | 6.572281839 | 2.689780359 | 13.62277588 | 0.000619097 | 0.014070989 | Yes |
| 646 | NECAP2 | ENSG00000157191 | 5.759532396 | 3.713714723 | 13.461809 | 0.000622034 | 0.014115836 | Yes |
| 647 | FMO4 | ENSG00000076258 | 3.807719092 | 7.664970557 | 13.69566558 | 0.000623783 | 0.014133606 | Yes |
| 648 | OSBPL6 | ENSG00000079156 | 5.314672166 | -2.562462255 | 13.51616742 | 0.000627109 | 0.014187 | Yes |
| 649 | GPATCH3 | ENSG00000198746 | 4.858436364 | -2.321465239 | 13.44084398 | 0.000633837 | 0.014317079 | Yes |
| 650 | TRIM5 | ENSG00000132256 | 3.57900747 | 4.380785169 | 13.36771325 | 0.000641147 | 0.01445989 | Yes |
| 651 | LRRC41 | ENSG00000132128 | 6.885600047 | 3.129200658 | 13.51329497 | 0.000647385 | 0.014562818 | Yes |
| 652 | DUSP12 | ENSG00000081721 | 4.007292041 | 4.680277254 | 13.38433812 | 0.000647701 | 0.014562818 | Yes |
| 653 | ANO10 | ENSG00000160746 | 4.480078448 | 4.229500784 | 13.29719737 | 0.000649586 | 0.014573344 | Yes |
| 654 | RPS7 | ENSG00000171863 | 7.390213462 | 3.315219256 | 13.5269728 | 0.00065016 | 0.014573344 | Yes |
| 655 | B3GNT9 | ENSG00000237172 | 3.268560382 | 7.068164557 | 13.45752149 | 0.000654355 | 0.014643337 | Yes |
| 656 | POPDC3 | ENSG00000132429 | 3.654936545 | -3.586797691 | 12.90785793 | 0.000655284 | 0.014643337 | Yes |
| 657 | AP1G2.AS1 | | 3.566641115 | 7.373160364 | 13.50590355 | 0.000658709 | 0.014697443 | Yes |
| 658 | ZHX1 | ENSG00000165156 | 5.452486809 | 3.935733526 | 13.27941178 | 0.00066144 | 0.014735914 | Yes |
| 659 | C4orf33 | ENSG00000151470 | 3.536007205 | 7.364142088 | 13.47739882 | 0.000665869 | 0.014812053 | Yes |
| 660 | ADAM8 | ENSG00000151651 | 4.311915766 | 6.428261269 | 13.52544061 | 0.000670948 | 0.014890095 | Yes |
| 661 | ZNF140 | ENSG00000196387 | 4.045867817 | 4.720888989 | 13.30209243 | 0.000671412 | 0.014890095 | Yes |
| 662 | BPHL | ENSG00000137274 | 4.101739942 | 4.217552406 | 13.20218359 | 0.000676933 | 0.014969754 | Yes |
| 663 | XPO6 | ENSG00000169180 | 6.176996877 | 3.268446712 | 13.329586 | 0.00067705 | 0.014969754 | Yes |
| 664 | EBF4 | ENSG00000088881 | 3.544535709 | -2.972081061 | 12.90986021 | 0.000678959 | 0.014989327 | Yes |
| 665 | MOCOS | ENSG00000075643 | 4.712331532 | 4.997428253 | 13.25634887 | 0.000682599 | 0.015046997 | Yes |
| 666 | SF3B4 | ENSG00000143368 | 6.301389291 | 3.928028031 | 13.24137948 | 0.000691949 | 0.01523017 | Yes |
| 667 | SOCS4 | ENSG00000180008 | 3.738572616 | 4.984379062 | 13.26603723 | 0.000695053 | 0.015254142 | Yes |
| 668 | XIRP2 | ENSG00000163092 | 9.32816044 | -4.568927682 | 13.41667837 | 0.000696096 | 0.015254142 | Yes |
| 669 | TACC3 | ENSG00000013810 | 6.214091784 | 3.830019511 | 13.22389502 | 0.000696503 | 0.015254142 | Yes |
| 670 | E2F1 | ENSG00000101412 | 5.64130172 | 4.735516748 | 13.15825736 | 0.000697222 | 0.015254142 | Yes |
| 671 | SUSD6 | ENSG00000100647 | 5.074434225 | 4.810790413 | 13.15969726 | 0.000698249 | 0.015254142 | Yes |
| 672 | UCN | ENSG00000163794 | 4.139940358 | -2.809075159 | 13.0028186 | 0.000704186 | 0.015353473 | Yes |
| 673 | ARF4 | ENSG00000168374 | 6.576663331 | 2.632796996 | 13.3066247 | 0.000704894 | 0.015353473 | Yes |
| 674 | CACNA2D1 | ENSG00000153956 | 5.007557712 | -2.724852159 | 13.1632533 | 0.000708942 | 0.015386872 | Yes |
| 675 | LASP1 | ENSG00000002834 | 8.221957021 | 2.436442722 | 13.36685747 | 0.000709415 | 0.015386872 | Yes |
| 676 | LIMCH1 | ENSG00000064042 | 6.856573707 | -2.057785994 | 13.34105137 | 0.000710588 | 0.015386872 | Yes |
| 677 | CDCA2 | ENSG00000184661 | 3.798285447 | 7.651072632 | 13.36792173 | 0.000710799 | 0.015386872 | Yes |
| 678 | GOLT1B | ENSG00000111711 | 4.663872925 | 4.17441733 | 13.07597722 | 0.000711768 | 0.015386872 | Yes |
| 679 | CDCA3 | ENSG00000111665 | 4.105557013 | 7.988202989 | 13.41289406 | 0.000712735 | 0.015386872 | Yes |
| 680 | COMP | ENSG00000105664 | 6.625734862 | -2.738078376 | 13.28833082 | 0.000718216 | 0.015482362 | Yes |
| 681 | GOLGA7 | ENSG00000147533 | 5.129772545 | 3.282425482 | 13.06219871 | 0.000723076 | 0.015564209 | Yes |
| 682 | UQCRFS1 | ENSG00000169021 | 5.642345714 | -2.311013629 | 13.19566953 | 0.000729232 | 0.015673673 | Yes |
| 683 | APEH | ENSG00000164062 | 6.426299638 | -1.893678438 | 13.22627888 | 0.000739668 | 0.015874669 | Yes |
| 684 | ZEB1.AS1 | | 3.137346185 | 6.913091199 | 13.11935697 | 0.000740762 | 0.01587487 | Yes |
| 685 | SLC39A8 | ENSG00000138821 | 3.700923429 | 7.549923273 | 13.21193995 | 0.000751777 | 0.016087372 | Yes |
| 686 | GTF2H3 | ENSG00000111358 | 5.562920652 | 3.152308209 | 13.01306334 | 0.000753574 | 0.01610229 | Yes |
| 687 | SLC2A1 | ENSG00000117394 | 5.521353831 | 9.465971558 | 13.38022025 | 0.000762079 | 0.016259778 | Yes |
| 688 | RABIF | ENSG00000183155 | 3.775611542 | 5.842520997 | 13.1257524 | 0.000763166 | 0.016259778 | Yes |
| 689 | TUSC3 | ENSG00000104723 | 5.092870225 | 4.032886168 | 12.89691543 | 0.000770202 | 0.016385828 | Yes |
| 690 | HACD3 | ENSG00000074696 | 6.642101747 | 2.699004368 | 13.07342111 | 0.000775741 | 0.016463145 | Yes |
| 691 | SPAG5 | ENSG00000076382 | 4.278975124 | 8.175751857 | 13.22473964 | 0.000776086 | 0.016463145 | Yes |
| 692 | NME7 | ENSG00000143156 | 4.337900603 | 4.704436695 | 12.8951008 | 0.000787497 | 0.016666963 | Yes |
| 693 | ENSG00000273576 | ENSG00000273576 | 3.333288732 | 7.113965967 | 13.00992764 | 0.000789083 | 0.016666963 | Yes |
| 694 | SNHG32 | ENSG00000204387 | 6.905975845 | 2.303892477 | 13.05914391 | 0.000790236 | 0.016666963 | Yes |
| 695 | CEP170B | ENSG00000099814 | 6.945116725 | 3.21057097 | 13.02401626 | 0.000790263 | 0.016666963 | Yes |
| 696 | RPUSD4 | ENSG00000165526 | 6.707513371 | -1.903947129 | 13.07215399 | 0.000791388 | 0.016666963 | Yes |
| 697 | TRMT11 | ENSG00000066651 | 3.024274289 | 6.769598552 | 12.92168738 | 0.000792762 | 0.016671921 | Yes |
| 698 | ZBTB1 | ENSG00000126804 | 4.50139748 | -2.432031639 | 12.83394718 | 0.000794401 | 0.016682425 | Yes |
| 699 | ENSG00000272369 | ENSG00000272369 | 2.615724141 | -3.28589345 | 12.23256289 | 0.000805125 | 0.016883409 | Yes |
| 700 | NDUFA13 | ENSG00000186010 | 7.708377735 | -1.61506788 | 13.05176604 | 0.000807833 | 0.016903142 | Yes |
| 701 | RIPK2 | ENSG00000104312 | 4.077207757 | 4.832554032 | 12.86880434 | 0.000808376 | 0.016903142 | Yes |
| 702 | CCNQ | ENSG00000262919 | 4.633327881 | -2.654591317 | 12.78648612 | 0.000811359 | 0.016941324 | Yes |
| 703 | QTRT2 | ENSG00000151576 | 3.074089499 | 6.834660864 | 12.83425268 | 0.000827324 | 0.017250059 | Yes |
| 704 | CDK4 | ENSG00000135446 | 9.373979263 | 4.334234632 | 12.96655969 | 0.000831737 | 0.017317405 | Yes |
| 705 | CARMIL2 | ENSG00000159753 | 2.355529665 | -3.067134104 | 12.11057491 | 0.000842923 | 0.017505716 | Yes |
| 706 | CDCA5 | ENSG00000146670 | 3.152853403 | 6.91525596 | 12.80667575 | 0.000843173 | 0.017505716 | Yes |
| 707 | RTL5 | ENSG00000242732 | 3.866563493 | 6.044564867 | 12.89046857 | 0.000848142 | 0.017583922 | Yes |
| 708 | TMEM63B | ENSG00000137216 | 3.131314046 | 6.910838359 | 12.78425349 | 0.000850725 | 0.017612536 | Yes |
| 709 | H2AZ1 | ENSG00000164032 | 7.693511419 | 2.570305472 | 12.86464063 | 0.000864024 | 0.017839938 | Yes |
| 710 | ENSG00000232224 | ENSG00000232224 | 2.661343546 | -2.877210589 | 12.13415467 | 0.000864147 | 0.017839938 | Yes |
| 711 | MAPK11 | ENSG00000185386 | 4.491540375 | -2.599440191 | 12.61122078 | 0.000865453 | 0.017841744 | Yes |
| 712 | LINC01133 | ENSG00000224259 | 6.142442032 | 4.441673911 | 12.65936266 | 0.000867319 | 0.01785506 | Yes |
| 713 | GOLPH3 | ENSG00000113384 | 5.412897176 | 3.311246402 | 12.63614627 | 0.000873732 | 0.017961818 | Yes |
| 714 | NRM | ENSG00000137404 | 4.724372736 | 4.304974104 | 12.56079764 | 0.000887052 | 0.018155891 | Yes |
| 715 | FGD1 | ENSG00000102302 | 3.965830265 | 7.807309123 | 12.8446677 | 0.000887073 | 0.018155891 | Yes |
| 716 | ABCC10 | ENSG00000124574 | 4.85485992 | 4.171688036 | 12.55694128 | 0.000887139 | 0.018155891 | Yes |
| 717 | HELZ2 | ENSG00000130589 | 6.320914054 | 3.119714692 | 12.69624463 | 0.000888134 | 0.018155891 | Yes |
| 718 | COX7C | ENSG00000127184 | 8.96247861 | -1.777370437 | 12.82755726 | 0.000889714 | 0.01816282 | Yes |
| 719 | CRYZ | ENSG00000116791 | 4.74339864 | 4.460454375 | 12.54995076 | 0.000894958 | 0.018229275 | Yes |
| 720 | RCOR2 | ENSG00000167771 | 3.546007824 | 7.356982677 | 12.74928675 | 0.00089546 | 0.018229275 | Yes |
| 721 | HMGCS1 | ENSG00000112972 | 6.128916259 | 2.745351038 | 12.65726959 | 0.000905022 | 0.018398343 | Yes |
| 722 | UQCR10 | ENSG00000184076 | 7.280148909 | -1.774462981 | 12.75672288 | 0.000907083 | 0.018414658 | Yes |
| 723 | SHCBP1 | ENSG00000171241 | 4.154484039 | 8.041443015 | 12.80584381 | 0.000912689 | 0.018502805 | Yes |
| 724 | TMEM14C | ENSG00000111843 | 3.991171396 | 5.263278527 | 12.63172107 | 0.000917268 | 0.018569925 | Yes |
| 725 | SEMA4D | ENSG00000187764 | 4.287463975 | -2.69650611 | 12.37627843 | 0.000936241 | 0.018926321 | Yes |
| 726 | LONP1 | ENSG00000196365 | 6.911722829 | -1.645805767 | 12.66491265 | 0.000938581 | 0.018926321 | Yes |
| 727 | DYNLL1 | ENSG00000088986 | 8.135654375 | 2.031383176 | 12.6806039 | 0.000938752 | 0.018926321 | Yes |
| 728 | HNRNPH1 | ENSG00000169045 | 9.151357771 | 1.835404643 | 12.68687196 | 0.000941877 | 0.018952055 | Yes |
| 729 | MMP14 | ENSG00000157227 | 9.325291626 | 2.967308122 | 12.67960402 | 0.000942618 | 0.018952055 | Yes |
| 730 | DOCK5 | ENSG00000147459 | 5.342850901 | 3.426676704 | 12.43907264 | 0.000944716 | 0.01896519 | Yes |
| 731 | ENSG00000214770 | ENSG00000214770 | 2.710114448 | -2.579650046 | 11.98108428 | 0.000945862 | 0.01896519 | Yes |
| 732 | RRP12 | ENSG00000052749 | 5.275710477 | -2.047293121 | 12.53620332 | 0.000947621 | 0.018974467 | Yes |
| 733 | AP3S1 | ENSG00000177879 | 5.543750974 | 3.043686346 | 12.46398671 | 0.000950499 | 0.019006079 | Yes |
| 734 | CARF | ENSG00000138380 | 3.217825435 | 4.400767347 | 12.39799805 | 0.000958442 | 0.019126338 | Yes |
| 735 | MACROD1 | ENSG00000133315 | 5.948892773 | -2.533704959 | 12.54645106 | 0.000959126 | 0.019126338 | Yes |
| 736 | KAZN | ENSG00000189337 | 5.98577098 | 3.364322095 | 12.42156069 | 0.00097688 | 0.019432014 | Yes |
| 737 | PLA2G12A | ENSG00000123739 | 5.124215591 | 3.133702913 | 12.34895421 | 0.000978538 | 0.019432014 | Yes |
| 738 | CDC42 | ENSG00000070831 | 7.639262606 | 2.779030076 | 12.5525109 | 0.000979384 | 0.019432014 | Yes |
| 739 | TUBA1B | ENSG00000123416 | 8.799018216 | 2.334301741 | 12.5835065 | 0.000979765 | 0.019432014 | Yes |
| 740 | OCIAD2 | ENSG00000145247 | 4.084175892 | 7.973029999 | 12.61414808 | 0.000983215 | 0.019467145 | Yes |
| 741 | ALPL | ENSG00000162551 | 3.462058855 | 7.270735396 | 12.50398202 | 0.000984501 | 0.019467145 | Yes |
| 742 | PLN | ENSG00000198523 | 4.913221417 | -3.584428953 | 12.2681119 | 0.000985527 | 0.019467145 | Yes |
| 743 | GYG1 | ENSG00000163754 | 5.119793044 | -2.611751241 | 12.37927096 | 0.000992951 | 0.019587368 | Yes |
| 744 | SEPTIN6 | ENSG00000125354 | 7.076876934 | 2.892756203 | 12.48839732 | 0.000994469 | 0.019590914 | Yes |
| 745 | LRR1 | ENSG00000165501 | 3.759257189 | 5.906433167 | 12.44154536 | 0.001013233 | 0.01993372 | Yes |
| 746 | CCN5 | ENSG00000064205 | 7.196127412 | -2.735607381 | 12.45757766 | 0.001019563 | 0.020031333 | Yes |
| 747 | SLC71A1 | ENSG00000156875 | 3.637350475 | 5.688440685 | 12.35878059 | 0.001037962 | 0.020362995 | Yes |
| 748 | RABL2B | ENSG00000079974 | 6.621986099 | 3.003620135 | 12.34778255 | 0.001039787 | 0.020362995 | Yes |
| 749 | PTDSS2 | ENSG00000174915 | 5.491758245 | -2.024200557 | 12.33248917 | 0.001041125 | 0.020362995 | Yes |
| 750 | ARMCX5 | ENSG00000125962 | 3.536645763 | 5.599310811 | 12.33178186 | 0.001042392 | 0.020362995 | Yes |
| 751 | RNF181 | ENSG00000168894 | 6.987625497 | -1.740795343 | 12.40883186 | 0.0010434 | 0.020362995 | Yes |
| 752 | SIPA1L3 | ENSG00000105738 | 4.253425394 | 3.46167095 | 12.13145745 | 0.001044869 | 0.020364519 | Yes |
| 753 | GSDME | ENSG00000105928 | 4.642818317 | 5.987386056 | 12.37760436 | 0.001048703 | 0.020412057 | Yes |
| 754 | PPFIA4 | ENSG00000143847 | 3.969654803 | -3.042831801 | 11.99434003 | 0.001055937 | 0.02052556 | Yes |
| 755 | ENSG00000256618 | ENSG00000256618 | 6.410587186 | -1.993709324 | 12.35215378 | 0.001057593 | 0.020530492 | Yes |
| 756 | SERP1 | ENSG00000120742 | 6.771156097 | 2.579752757 | 12.32653198 | 0.001061236 | 0.020573928 | Yes |
| 757 | CPTP | ENSG00000224051 | 5.491678155 | -2.082282827 | 12.27404561 | 0.001065574 | 0.020630691 | Yes |
| 758 | CIT | ENSG00000122966 | 3.702283182 | 7.538235241 | 12.35089137 | 0.00106839 | 0.020657889 | Yes |
| 759 | TRIB3 | ENSG00000101255 | 3.761919808 | 5.020380439 | 12.21313979 | 0.001078706 | 0.020829847 | Yes |
| 760 | CCDC102A | ENSG00000135736 | 3.412649118 | 7.212834895 | 12.26778254 | 0.001081322 | 0.02085284 | Yes |
| 761 | CDK2 | ENSG00000123374 | 6.801552835 | 3.029470371 | 12.25070129 | 0.001087714 | 0.020923345 | Yes |
| 762 | MICALL2 | ENSG00000164877 | 4.948948131 | 3.60333154 | 12.07324759 | 0.001089359 | 0.020923345 | Yes |
| 763 | IMPDH1 | ENSG00000106348 | 5.316926311 | 3.429786168 | 12.09980458 | 0.001089948 | 0.020923345 | Yes |
| 764 | AJUBA | ENSG00000129474 | 3.304111771 | 4.68359125 | 12.12883483 | 0.001090696 | 0.020923345 | Yes |
| 765 | PACRGL | ENSG00000163138 | 3.119968819 | 5.109335189 | 12.13127152 | 0.001093424 | 0.020948237 | Yes |
| 766 | ROBO3 | ENSG00000154134 | 4.649721749 | 4.275030996 | 12.05010531 | 0.001101847 | 0.0210747 | Yes |
| 767 | ANKRD9 | ENSG00000156381 | 7.29111699 | -2.16312157 | 12.27585031 | 0.001103942 | 0.0210747 | Yes |
| 768 | MAPK12 | ENSG00000188130 | 6.218849318 | -2.690603803 | 12.21941678 | 0.001104777 | 0.0210747 | Yes |
| 769 | NCOA5 | ENSG00000124160 | 3.884180621 | 3.991570113 | 12.02065396 | 0.001105785 | 0.0210747 | Yes |
| 770 | TEAD4 | ENSG00000197905 | 4.456171085 | -2.615048842 | 11.99207664 | 0.001121777 | 0.021351691 | Yes |
| 771 | LRP11 | ENSG00000120256 | 5.413801983 | 3.548712355 | 12.02838432 | 0.00112572 | 0.021398917 | Yes |
| 772 | DCPS | ENSG00000110063 | 3.873928891 | 4.582278305 | 12.0490912 | 0.001128243 | 0.021404176 | Yes |
| 773 | FIS1 | ENSG00000214253 | 7.093438974 | -1.453761827 | 12.22306796 | 0.001128921 | 0.021404176 | Yes |
| 774 | DAPK2 | ENSG00000035664 | 5.643373936 | -1.969584462 | 12.12939671 | 0.00113929 | 0.021547854 | Yes |
| 775 | MYBPH | ENSG00000133055 | 4.85963351 | -3.505434466 | 11.93988077 | 0.001139444 | 0.021547854 | Yes |
| 776 | COX5B | ENSG00000135940 | 8.048419632 | -2.070089101 | 12.20558204 | 0.001143839 | 0.021603054 | Yes |
| 777 | GPT2 | ENSG00000166123 | 5.775666137 | -2.672155876 | 12.09931877 | 0.001146742 | 0.021629984 | Yes |
| 778 | ALKBH7 | ENSG00000125652 | 6.286570224 | -1.593240933 | 12.15293396 | 0.001149051 | 0.02164564 | Yes |
| 779 | FAIM | ENSG00000158234 | 3.091571873 | 6.809089197 | 12.03710926 | 0.00115301 | 0.021652037 | Yes |
| 780 | SLC7A11 | ENSG00000151012 | 3.483333465 | 7.301874632 | 12.12797156 | 0.001153652 | 0.021652037 | Yes |
| 781 | LOXL2 | ENSG00000134013 | 6.46162023 | 4.186207037 | 12.01717955 | 0.001153829 | 0.021652037 | Yes |
| 782 | COLGALT1 | ENSG00000130309 | 7.533471393 | 2.796929935 | 12.14602487 | 0.00115686 | 0.021681132 | Yes |
| 783 | AOPEP | ENSG00000148120 | 4.913719046 | -2.272626873 | 12.00409049 | 0.001161883 | 0.021747418 | Yes |
| 784 | ENSG00000243444 | ENSG00000243444 | 5.125077131 | -3.075945668 | 11.96146445 | 0.001168399 | 0.02184145 | Yes |
| 785 | LAPTM4A | ENSG00000068697 | 9.779828351 | 1.814513617 | 12.14263728 | 0.001180304 | 0.022017696 | Yes |
| 786 | CHROMR | ENSG00000223960 | 2.870931785 | 6.585077662 | 11.933125 | 0.001180836 | 0.022017696 | Yes |
| 787 | LIX1L | ENSG00000271601 | 5.276961807 | 3.961672417 | 11.88949009 | 0.001185756 | 0.022055643 | Yes |
| 788 | SORL1 | ENSG00000137642 | 4.690396086 | 5.4712913 | 12.00716008 | 0.001185884 | 0.022055643 | Yes |
| 789 | MAPK7 | ENSG00000166484 | 5.829402198 | 4.152123938 | 11.90046079 | 0.001192373 | 0.022148181 | Yes |
| 790 | MID1 | ENSG00000101871 | 6.198499308 | 2.982998436 | 11.98186848 | 0.001196891 | 0.022203921 | Yes |
| 791 | CD86 | ENSG00000114013 | 3.647451782 | 7.452894227 | 12.05755294 | 0.001200101 | 0.022235291 | Yes |
| 792 | SNX4 | ENSG00000114520 | 5.146044911 | 3.062062711 | 11.86084638 | 0.001207812 | 0.022348517 | Yes |
| 793 | TXNDC5 | ENSG00000239264 | 3.312758983 | 7.106728984 | 11.98163284 | 0.001209266 | 0.022348517 | Yes |
| 794 | LINC02693 | ENSG00000212719 | 4.383640147 | 4.746500151 | 11.87065101 | 0.001215715 | 0.022439374 | Yes |
| 795 | PIAS2 | ENSG00000078043 | 6.371900823 | -1.465113298 | 11.99757421 | 0.001228593 | 0.022648515 | Yes |
| 796 | ARHGEF3 | ENSG00000163947 | 5.403975122 | 4.458194385 | 11.80403179 | 0.001232551 | 0.022692884 | Yes |
| 797 | QNG1 | ENSG00000165118 | 3.704301143 | 5.784026167 | 11.93607634 | 0.001244354 | 0.022881426 | Yes |
| 798 | NDUFB11 | ENSG00000147123 | 7.936567969 | -1.484162702 | 11.99850118 | 0.001246959 | 0.022900554 | Yes |
| 799 | ZNF572 | ENSG00000180938 | 2.890592871 | 6.62401068 | 11.80461766 | 0.001252017 | 0.022964619 | Yes |
| 800 | LDLR | ENSG00000130164 | 6.572672514 | 3.675484943 | 11.85261711 | 0.001258004 | 0.023045558 | Yes |
| 801 | TMSB10 | ENSG00000034510 | 10.16325775 | 2.028432545 | 11.97704753 | 0.00126506 | 0.023129373 | Yes |
| 802 | NIBAN2 | ENSG00000136830 | 7.158193953 | 3.262738552 | 11.89918606 | 0.001265739 | 0.023129373 | Yes |
| 803 | STRA6 | ENSG00000137868 | 4.936860589 | 8.862085227 | 12.06128981 | 0.001274738 | 0.023264772 | Yes |
| 804 | IDH1 | ENSG00000138413 | 7.115806769 | 2.861452928 | 11.88800572 | 0.001278622 | 0.023306593 | Yes |

| | A | B | C | D | E | F | G | H |
|---|---|---|---|---|---|---|---|---|
| 805 | ENSG00000257303 | ENSG00000257303 | 2.671490611 | -3.063549694 | 11.24605453 | 0.001284543 | 0.02336234 | Yes |
| 806 | CDC14B | ENSG00000081377 | 3.365756344 | 7.169583936 | 11.84862843 | 0.001284873 | 0.02336234 | Yes |
| 807 | NDEL1 | ENSG00000166579 | 6.044758458 | 2.999179351 | 11.79529876 | 0.001287302 | 0.023362715 | Yes |
| 808 | GINS1 | ENSG00000101003 | 3.220173243 | 7.00998124 | 11.81442573 | 0.001288086 | 0.023362715 | Yes |
| 809 | HSDL1 | ENSG00000103160 | 4.719386059 | -2.457562809 | 11.72285081 | 0.001290517 | 0.02337784 | Yes |
| 810 | SAMD8 | ENSG00000156671 | 3.324314116 | 4.543594825 | 11.71737282 | 0.001294225 | 0.023416034 | Yes |
| 811 | SKP2 | ENSG00000145604 | 4.941038957 | 3.501870331 | 11.65574251 | 0.001304233 | 0.023567971 | Yes |
| 812 | PAK4 | ENSG00000130669 | 5.74185978 | 2.759249597 | 11.74145042 | 0.001310492 | 0.023603308 | Yes |
| 813 | FLVCR1 | ENSG00000162769 | 3.223035934 | 5.286625685 | 11.72681357 | 0.001310833 | 0.023603308 | Yes |
| 814 | DGKA | ENSG00000065357 | 5.416986065 | 3.340906429 | 11.67713655 | 0.001313328 | 0.023603308 | Yes |
| 815 | ENSG00000269936 | ENSG00000269936 | 5.180710855 | -2.625117874 | 11.7203296 | 0.001315275 | 0.023603308 | Yes |
| 816 | EMC7 | ENSG00000134153 | 4.653085074 | 4.175549172 | 11.63243016 | 0.001316318 | 0.023603308 | Yes |
| 817 | CKLF | ENSG00000217555 | 4.714386916 | 4.029160613 | 11.62616342 | 0.001317387 | 0.023603308 | Yes |
| 818 | MTMR4 | ENSG00000108389 | 5.306315888 | 3.136784892 | 11.66802221 | 0.001317956 | 0.023603308 | Yes |
| 819 | EPHA2 | ENSG00000142627 | 3.089760059 | 6.859069276 | 11.73008185 | 0.001319089 | 0.023603308 | Yes |
| 820 | RMI2 | ENSG00000175643 | 3.402450416 | 7.22156519 | 11.78915914 | 0.001322335 | 0.023632504 | Yes |
| 821 | AP5Z1 | ENSG00000242802 | 7.308131186 | 2.520016559 | 11.81363908 | 0.001328547 | 0.023714564 | Yes |
| 822 | PPOX | ENSG00000143224 | 4.485981662 | 3.463646061 | 11.58355513 | 0.00133108 | 0.023720689 | Yes |
| 823 | GALNT10 | ENSG00000164574 | 5.920236906 | 3.69778162 | 11.66348819 | 0.001333269 | 0.023720689 | Yes |
| 824 | ENSG00000248559 | ENSG00000248559 | 4.228084145 | -2.571945171 | 11.55379618 | 0.001333752 | 0.023720689 | Yes |
| 825 | CBX8 | ENSG00000141570 | 3.42625007 | 7.220092011 | 11.76456573 | 0.001335958 | 0.023731081 | Yes |
| 826 | CPEB2 | ENSG00000137449 | 4.187567026 | 8.079733796 | 11.85724499 | 0.001348708 | 0.023928535 | Yes |
| 827 | OLFML2B | ENSG00000162745 | 5.00405184 | 7.174012047 | 11.86456308 | 0.001353613 | 0.023986477 | Yes |
| 828 | SLC6A9 | ENSG00000196517 | 4.055378472 | 4.81158961 | 11.63116066 | 0.001361778 | 0.024071643 | Yes |
| 829 | NAXD | ENSG00000213995 | 4.381982449 | 3.770081495 | 11.53013434 | 0.001363342 | 0.024071643 | Yes |
| 830 | AGAP2.AS1 | | 4.7224068 | 8.628534605 | 11.87861806 | 0.00136448 | 0.024071643 | Yes |
| 831 | NEK2 | ENSG00000117650 | 3.214785489 | 7.005836862 | 11.67686176 | 0.001364997 | 0.024071643 | Yes |
| 832 | HPSE | ENSG00000173083 | 2.87526056 | -2.693637462 | 11.19025612 | 0.001377654 | 0.024255313 | Yes |
| 833 | DIP2B | ENSG00000066084 | 4.797526628 | 3.440612648 | 11.51578316 | 0.001381037 | 0.024255313 | Yes |
| 834 | SNRPA1 | ENSG00000131876 | 4.647886094 | 3.31105921 | 11.50486549 | 0.001382291 | 0.024255313 | Yes |
| 835 | KIAA0753 | ENSG00000198920 | 4.332850492 | 3.043045811 | 11.48526775 | 0.001382867 | 0.024255313 | Yes |
| 836 | KIF22 | ENSG00000079616 | 6.196574355 | -2.032975626 | 11.69837155 | 0.001383698 | 0.024255313 | Yes |
| 837 | MLKL | ENSG00000168404 | 4.067522745 | 4.382376566 | 11.53547025 | 0.001387156 | 0.024276487 | Yes |
| 838 | MRPL46 | ENSG00000259494 | 5.29748668 | -2.01331571 | 11.63356798 | 0.001388223 | 0.024276487 | Yes |
| 839 | FAM111B | ENSG00000189057 | 3.353915639 | 7.122319951 | 11.64415341 | 0.001396005 | 0.024331369 | Yes |
| 840 | PRANCR | ENSG00000257815 | 4.376369531 | 5.625551148 | 11.66174793 | 0.001397256 | 0.024331369 | Yes |
| 841 | NDUFV3 | ENSG00000160194 | 7.009213637 | -1.710808222 | 11.70729906 | 0.001397382 | 0.024331369 | Yes |
| 842 | CMPK2 | ENSG00000134326 | 3.500089609 | 7.307762669 | 11.66939415 | 0.001398932 | 0.024331369 | Yes |
| 843 | TSPAN5 | ENSG00000168785 | 4.277523709 | 8.171129853 | 11.77709576 | 0.001399673 | 0.024331369 | Yes |
| 844 | PCDHB14 | ENSG00000120327 | 3.15885109 | 6.911202862 | 11.59218035 | 0.001404731 | 0.024390331 | Yes |
| 845 | CAD | ENSG00000084774 | 5.200814192 | 4.180074527 | 11.47373499 | 0.001416644 | 0.024568038 | Yes |
| 846 | CFAP119 | ENSG00000196118 | 3.795756626 | 7.664306267 | 11.68350794 | 0.001420725 | 0.024603392 | Yes |
| 847 | NIFK | ENSG00000155438 | 5.142233611 | 3.020927033 | 11.4789702 | 0.001424607 | 0.024603392 | Yes |
| 848 | SDCBP2 | ENSG00000125775 | 3.839754695 | 7.696289657 | 11.67909322 | 0.001425798 | 0.024603392 | Yes |
| 849 | ZNF528 | ENSG00000167555 | 3.77380558 | 5.145276711 | 11.5645266 | 0.001427078 | 0.024603392 | Yes |
| 850 | BNIP1 | ENSG00000113734 | 3.589559157 | 5.650399294 | 11.5929422 | 0.001427088 | 0.024603392 | Yes |
| 851 | STXBP4 | ENSG00000166263 | 4.334304385 | 4.035371233 | 11.40334513 | 0.001447499 | 0.024869567 | Yes |
| 852 | NUP42 | ENSG00000136243 | 4.065987592 | 4.450358586 | 11.44167911 | 0.001448677 | 0.024869567 | Yes |
| 853 | ADHFE1 | ENSG00000147576 | 3.404337855 | -2.514267289 | 11.20190913 | 0.001449388 | 0.024869567 | Yes |
| 854 | TBK1 | ENSG00000183735 | 3.746329556 | 5.804902112 | 11.57610639 | 0.00145129 | 0.024869567 | Yes |
| 855 | GPC2 | ENSG00000213420 | 4.730251941 | 4.229182999 | 11.40824173 | 0.00145233 | 0.024869567 | Yes |
| 856 | MYH10 | ENSG00000133026 | 7.47876483 | 2.477228858 | 11.60703408 | 0.001452721 | 0.024869567 | Yes |
| 857 | MAPK13 | ENSG00000156711 | 3.985311954 | 7.847057454 | 11.64794889 | 0.001455649 | 0.024890583 | Yes |
| 858 | STX6 | ENSG00000135823 | 5.169154397 | 3.027073015 | 11.42311294 | 0.00146019 | 0.024939086 | Yes |
| 859 | DDOST | ENSG00000244038 | 6.291518452 | 3.064605251 | 11.50031315 | 0.001469375 | 0.025066707 | Yes |
| 860 | HNRNPF | ENSG00000169813 | 7.700747391 | 2.968835354 | 11.56790053 | 0.001474367 | 0.025103279 | Yes |
| 861 | KLHL15 | ENSG00000174010 | 3.664871194 | 7.516399206 | 11.57294686 | 0.001475887 | 0.025103279 | Yes |
| 862 | ZNF493 | ENSG00000196268 | 4.596384825 | 3.48603902 | 11.34950387 | 0.001477409 | 0.025103279 | Yes |
| 863 | CCDC9B | ENSG00000188549 | 3.9451009 | -2.762564565 | 11.24682786 | 0.001483007 | 0.025103279 | Yes |
| 864 | GLI3 | ENSG00000106571 | 6.46679299 | 3.316367617 | 11.47812597 | 0.001483142 | 0.025103279 | Yes |
| 865 | C20orf96 | ENSG00000196476 | 5.285500508 | 3.315114774 | 11.38485293 | 0.00148346 | 0.025103279 | Yes |
| 866 | CCDC120 | ENSG00000147144 | 3.864374066 | 5.938829717 | 11.53576129 | 0.001485166 | 0.025103279 | Yes |
| 867 | LAMB1 | ENSG00000091136 | 6.426177627 | 2.672402675 | 11.49970267 | 0.001487049 | 0.025103279 | Yes |
| 868 | EME1 | ENSG00000154920 | 2.861509384 | 6.592166539 | 11.3925587 | 0.001490355 | 0.025103279 | Yes |
| 869 | SMO | ENSG00000128602 | 4.887975827 | 3.570608645 | 11.34329875 | 0.001491148 | 0.025103279 | Yes |
| 870 | EDA2R | ENSG00000131080 | 2.980097558 | 6.711395274 | 11.41412355 | 0.001491705 | 0.025103279 | Yes |
| 871 | BASP1 | ENSG00000176788 | 8.025458926 | 3.049262602 | 11.54699284 | 0.001492099 | 0.025103279 | Yes |
| 872 | PCLAF | ENSG00000166803 | 5.239207313 | 4.859352569 | 11.36528378 | 0.001496316 | 0.025145329 | Yes |
| 873 | ATG9A | ENSG00000198925 | 6.570615074 | -1.979913382 | 11.52151154 | 0.001501014 | 0.025195348 | Yes |
| 874 | ENSG00000259529 | ENSG00000259529 | 3.27894287 | 7.051035616 | 11.45478131 | 0.001505245 | 0.025237423 | Yes |
| 875 | RAB1A | ENSG00000138069 | 6.587878328 | 2.469718564 | 11.48199415 | 0.001508904 | 0.025259483 | Yes |
| 876 | ATAD2 | ENSG00000156802 | 4.77186784 | 4.080713475 | 11.31470392 | 0.001510012 | 0.025259483 | Yes |
| 877 | IGFBP3 | ENSG00000146674 | 7.360791853 | 3.388770051 | 11.47475636 | 0.001517401 | 0.025354105 | Yes |
| 878 | ASF1B | ENSG00000105011 | 5.960504021 | 5.133700164 | 11.32736082 | 0.001524562 | 0.025409092 | Yes |
| 879 | DTNA | ENSG00000134769 | 5.019528616 | -2.655285635 | 11.35646686 | 0.001525691 | 0.025409092 | Yes |
| 880 | ZCCHC4 | ENSG00000168228 | 3.057366666 | 6.769589308 | 11.37021101 | 0.00152743 | 0.025409092 | Yes |
| 881 | ORAI1 | ENSG00000276045 | 5.301668407 | -2.189018432 | 11.39577941 | 0.001532746 | 0.025409092 | Yes |
| 882 | SLCO2A1 | ENSG00000174640 | 3.701097887 | 7.550864078 | 11.48737765 | 0.00153308 | 0.025409092 | Yes |
| 883 | MFN1 | ENSG00000171109 | 4.342362256 | 4.200208346 | 11.28146251 | 0.00153409 | 0.025409092 | Yes |
| 884 | NSUN2 | ENSG00000037474 | 6.121998647 | 2.611138949 | 11.40559639 | 0.001536656 | 0.025409092 | Yes |
| 885 | TMCC3 | ENSG00000057704 | 4.888954465 | -2.187336758 | 11.34952058 | 0.001536825 | 0.025409092 | Yes |
| 886 | XPO5 | ENSG00000124571 | 4.593937275 | 3.417393644 | 11.25845564 | 0.001537249 | 0.025409092 | Yes |
| 887 | ZSCAN9 | ENSG00000137185 | 4.108661454 | 4.045907682 | 11.26511665 | 0.001538051 | 0.025409092 | Yes |
| 888 | PCK2 | ENSG00000100889 | 3.363435145 | 5.382372935 | 11.37145475 | 0.001542436 | 0.025438145 | Yes |
| 889 | CRYBG1 | ENSG00000112297 | 5.603088582 | 3.702482902 | 11.29965485 | 0.001544512 | 0.025438145 | Yes |
| 890 | ATXN7L2 | ENSG00000162650 | 3.980552719 | 4.718460384 | 11.32729168 | 0.001546136 | 0.025438145 | Yes |
| 891 | TTYH3 | ENSG00000136295 | 7.596506975 | 3.195003998 | 11.44520308 | 0.001546762 | 0.025438145 | Yes |
| 892 | ENSG00000272760 | ENSG00000272760 | 3.219988649 | 6.985444016 | 11.35416575 | 0.001563875 | 0.025670205 | Yes |
| 893 | WDR41 | ENSG00000164253 | 4.947164441 | 2.866183348 | 11.25239065 | 0.00156438 | 0.025670205 | Yes |
| 894 | RAMAC | ENSG00000169612 | 2.958386848 | 6.700235254 | 11.29132116 | 0.001571346 | 0.025755645 | Yes |
| 895 | RAI14 | ENSG00000039560 | 6.721870947 | 3.828972441 | 11.32545045 | 0.001576756 | 0.025815412 | Yes |
| 896 | SBNO1.AS1 | | 3.43117592 | 7.25347952 | 11.36248022 | 0.001587501 | 0.025962293 | Yes |
| 897 | NDUFB3 | ENSG00000119013 | 5.338046916 | 2.938560156 | 11.24628665 | 0.001589612 | 0.025967809 | Yes |
| 898 | SND1.IT1 | | 4.989428625 | 3.383266194 | 11.19644618 | 0.001594535 | 0.026019181 | Yes |
| 899 | MOB1A | ENSG00000114978 | 5.046127429 | 3.07445245 | 11.19979668 | 0.001601658 | 0.026106313 | Yes |
| 900 | SERPINE2 | ENSG00000135919 | 8.629576518 | 3.661323016 | 11.3762974 | 0.001605511 | 0.026137655 | Yes |
| 901 | TMX3 | ENSG00000166479 | 5.078365911 | 3.22169648 | 11.18509097 | 0.001609834 | 0.026137655 | Yes |
| 902 | KDELR3 | ENSG00000100196 | 4.285340968 | 5.144577851 | 11.26963601 | 0.001610543 | 0.026137655 | Yes |
| 903 | RIPK1 | ENSG00000137275 | 4.979020906 | 3.204666306 | 11.17572461 | 0.001611858 | 0.026137655 | Yes |
| 904 | TMEM147 | ENSG00000105677 | 7.354126525 | -1.542803082 | 11.37526651 | 0.00161251 | 0.026137655 | Yes |
| 905 | RNGTT | ENSG00000111880 | 4.751651809 | -2.0887382 | 11.22571255 | 0.001615207 | 0.026152425 | Yes |
| 906 | ZNF594 | ENSG00000180626 | 3.001755718 | 4.965476399 | 11.18889864 | 0.001619961 | 0.026200414 | Yes |
| 907 | ARHGAP23 | ENSG00000275832 | 5.760226506 | 3.167388634 | 11.21950073 | 0.001624242 | 0.026217174 | Yes |
| 908 | ANKRD13C | ENSG00000118454 | 3.637449184 | 4.565748497 | 11.19866647 | 0.00162458 | 0.026217174 | Yes |
| 909 | FSTL1 | ENSG00000163430 | 9.750645961 | 2.915173233 | 11.3678525 | 0.001628073 | 0.026244607 | Yes |
| 910 | MKS1 | ENSG00000011143 | 4.511038845 | 3.103500175 | 11.1086225 | 0.001638822 | 0.026347325 | Yes |
| 911 | ARL6IP6 | ENSG00000177917 | 3.031452584 | 6.779602414 | 11.20734691 | 0.001639827 | 0.026347325 | Yes |
| 912 | PCSK1N | ENSG00000102109 | 4.16881658 | 8.053578724 | 11.38339078 | 0.001641345 | 0.026347325 | Yes |
| 913 | CRTC3 | ENSG00000140577 | 5.274816372 | 2.762132984 | 11.17289813 | 0.00164251 | 0.026347325 | Yes |
| 914 | PPP3CB | ENSG00000107758 | 6.499049357 | -2.008169866 | 11.30433278 | 0.001643445 | 0.026347325 | Yes |
| 915 | ZNF710 | ENSG00000140548 | 3.022471059 | 6.78458751 | 11.18753144 | 0.001654582 | 0.026487293 | Yes |
| 916 | GTF2H2 | ENSG00000145736 | 3.563319087 | 5.691880042 | 11.2423441 | 0.001655795 | 0.026487293 | Yes |
| 917 | BUB1B | ENSG00000156970 | 3.917025359 | 7.77397532 | 11.32057721 | 0.001664123 | 0.026520726 | Yes |
| 918 | ZKSCAN4 | ENSG00000187626 | 3.146981614 | 5.152189934 | 11.15760315 | 0.001664967 | 0.026520726 | Yes |
| 919 | HOMER1 | ENSG00000152413 | 5.015919177 | -2.426307475 | 11.16545812 | 0.001665011 | 0.026520726 | Yes |
| 920 | MT.CO1 | | 10.36527614 | -1.719118803 | 11.32089534 | 0.001665494 | 0.026520726 | Yes |
| 921 | ENSG00000227398 | ENSG00000227398 | 2.322901791 | -2.735321646 | 10.66349258 | 0.001666945 | 0.026520726 | Yes |
| 922 | JRKL | ENSG00000183340 | 3.037096966 | 6.81642967 | 11.16835388 | 0.001672493 | 0.026580112 | Yes |
| 923 | CENPM | ENSG00000100162 | 3.602421195 | 7.447310071 | 11.24763659 | 0.001686794 | 0.026778306 | Yes |
| 924 | FAM83D | ENSG00000101447 | 3.928345042 | 7.806767073 | 11.2838241 | 0.001692795 | 0.026818036 | Yes |
| 925 | GMNN | ENSG00000112312 | 4.47481984 | 4.515206692 | 11.07601956 | 0.001692961 | 0.026818036 | Yes |
| 926 | TWF1 | ENSG00000151239 | 4.076515795 | 4.05425491 | 11.02836097 | 0.001705594 | 0.026963913 | Yes |
| 927 | CTBS | ENSG00000117151 | 3.887176295 | 4.535045644 | 11.08246907 | 0.001705854 | 0.026963913 | Yes |
| 928 | RP2 | ENSG00000102218 | 3.247218699 | 7.053120806 | 11.1550964 | 0.001711423 | 0.027013272 | Yes |
| 929 | ENSG00000231856 | ENSG00000231856 | 2.219430966 | -2.486170161 | 10.59821145 | 0.001712668 | 0.027013272 | Yes |
| 930 | TBC1D17 | ENSG00000104946 | 5.557327027 | -2.098872754 | 11.15467953 | 0.001715215 | 0.02702433 | Yes |
| 931 | FKBPL | ENSG00000204315 | 3.058208184 | 6.816337832 | 11.1007112 | 0.00172204 | 0.027102687 | Yes |
| 932 | SPC24 | ENSG00000161888 | 3.996321028 | 7.877333402 | 11.23074115 | 0.00173681 | 0.027305786 | Yes |
| 933 | SCAMP1.AS1 | | 3.06614878 | 6.84631008 | 11.08139837 | 0.001740448 | 0.027333627 | Yes |
| 934 | ATP6V1B1 | ENSG00000116039 | 3.043359181 | 6.802706659 | 11.05540749 | 0.0017543 | 0.027492416 | Yes |
| 935 | STARD3NL | ENSG00000010270 | 3.952251506 | 4.182189995 | 10.97458268 | 0.001757226 | 0.027492416 | Yes |
| 936 | IGDCC3 | ENSG00000174498 | 3.680626882 | 7.51758807 | 11.15884805 | 0.001758587 | 0.027492416 | Yes |
| 937 | ELAC1 | ENSG00000141642 | 3.210915624 | 6.986455709 | 11.08013029 | 0.001759269 | 0.027492416 | Yes |
| 938 | KCTD13 | ENSG00000174943 | 4.482508692 | 4.046148974 | 10.95856065 | 0.00175995 | 0.027492416 | Yes |
| 939 | PHTF1 | ENSG00000116793 | 3.737476194 | 5.839025954 | 11.11806853 | 0.001764888 | 0.027540159 | Yes |
| 940 | SERPINH1 | ENSG00000149257 | 7.446974698 | 2.57205842 | 11.12643013 | 0.001778455 | 0.027722308 | Yes |
| 941 | ASCC2 | ENSG00000100325 | 6.659681045 | 2.612250089 | 11.08739998 | 0.001784203 | 0.027782323 | Yes |
| 942 | MYH8 | ENSG00000133020 | 2.951175955 | -3.270394336 | 10.56680877 | 0.001787787 | 0.027808546 | Yes |
| 943 | WASF1 | ENSG00000112290 | 3.017022214 | 6.779887867 | 11.00249013 | 0.001791866 | 0.027842401 | Yes |
| 944 | SIN3B | ENSG00000127511 | 7.770952884 | 2.254762701 | 11.10186514 | 0.001806686 | 0.028042907 | Yes |
| 945 | LURAP1L | ENSG00000153714 | 3.028000947 | 6.796125828 | 10.98367519 | 0.00180879 | 0.028045822 | Yes |
| 946 | DYNLT2B | ENSG00000213123 | 4.288136981 | 8.191380342 | 11.16110824 | 0.00181181 | 0.028049834 | Yes |
| 947 | NPIPA1 | ENSG00000183426 | 3.333011086 | 7.126166622 | 11.03250632 | 0.001812881 | 0.028049834 | Yes |
| 948 | NIPSNAP3A | ENSG00000136783 | 4.538885107 | 3.341485131 | 10.87153549 | 0.001819553 | 0.02812334 | Yes |
| 949 | MDK | ENSG00000110492 | 8.276317553 | 3.199919517 | 11.07219183 | 0.001826973 | 0.028161538 | Yes |
| 950 | ANKRD13C.DT | | 4.923090639 | 3.768378082 | 10.87934138 | 0.001827892 | 0.028161538 | Yes |
| 951 | COPB1 | ENSG00000129083 | 6.185287601 | 2.450930766 | 11.00942006 | 0.001829168 | 0.028161538 | Yes |
| 952 | APTR | ENSG00000214293 | 3.054466541 | 5.024734087 | 10.91962144 | 0.001829721 | 0.028161538 | Yes |
| 953 | TUBA1C | ENSG00000167553 | 7.663986606 | 3.207256518 | 11.0484248 | 0.001832474 | 0.028174287 | Yes |
| 954 | TM9SF4 | ENSG00000101337 | 4.7849746 | 4.06375758 | 10.86209169 | 0.001840932 | 0.028261814 | Yes |
| 955 | AFG2A | ENSG00000145375 | 4.658458173 | 3.095071369 | 10.84939397 | 0.001843511 | 0.028261814 | Yes |
| 956 | KIF14 | ENSG00000118193 | 3.478053469 | 7.30245404 | 11.0187944 | 0.001843959 | 0.028261814 | Yes |
| 957 | FANK1 | ENSG00000203780 | 4.798007745 | 5.607359566 | 10.97671772 | 0.001846042 | 0.028264142 | Yes |
| 958 | NIPSNAP2 | ENSG00000146729 | 6.202662153 | -1.886080491 | 11.01431689 | 0.00185212 | 0.028327568 | Yes |
| 959 | FAM3C | ENSG00000196937 | 5.812624956 | 3.273679265 | 10.91041927 | 0.001856838 | 0.028370074 | Yes |
| 960 | UNC5B | ENSG00000107731 | 5.435439726 | 3.781928424 | 10.86294369 | 0.001859702 | 0.028384205 | Yes |
| 961 | S100A11 | ENSG00000163191 | 9.975704274 | 2.612136027 | 11.05198403 | 0.001863194 | 0.02840788 | Yes |
| 962 | PTPN18 | ENSG00000072135 | 6.54785538 | -1.536051327 | 11.01304309 | 0.001866121 | 0.028422911 | Yes |
| 963 | ASIC1 | ENSG00000110881 | 4.030414383 | 4.670543414 | 10.87685516 | 0.001875136 | 0.02853053 | Yes |
| 964 | LNCTAM34A | ENSG00000234546 | 3.410876365 | 7.221475402 | 10.95193516 | 0.001888251 | 0.028700231 | Yes |
| 965 | TIGD1 | ENSG00000221944 | 4.163734361 | 4.896044013 | 10.86072952 | 0.001903946 | 0.028908768 | Yes |
| 966 | CEP15 | ENSG00000114405 | 3.105267968 | 6.880279898 | 10.87604161 | 0.001907052 | 0.028925931 | Yes |
| 967 | NR1D1 | ENSG00000126368 | 5.200886284 | -2.027105153 | 10.85759831 | 0.00193017 | 0.029246276 | Yes |
| 968 | ADAM17 | ENSG00000151694 | 6.565534886 | 2.366312848 | 10.90428816 | 0.00193279 | 0.029255677 | Yes |
| 969 | WRAP53 | ENSG00000141499 | 3.215979926 | 5.248698033 | 10.82331391 | 0.001937699 | 0.02925721 | Yes |
| 970 | TRIP6 | ENSG00000087077 | 6.041082366 | 2.505551556 | 10.86333418 | 0.001939227 | 0.02925721 | Yes |
| 971 | WASHC2C.1 | | 4.213490269 | 3.472346628 | 10.71231699 | 0.001940378 | 0.02925721 | Yes |
| 972 | ROR2 | ENSG00000169071 | 4.032162193 | 7.907477946 | 10.9713581 | 0.001940886 | 0.02925721 | Yes |
| 973 | BBS12 | ENSG00000181004 | 3.003217745 | 6.776485814 | 10.80454905 | 0.001952399 | 0.029400472 | Yes |
| 974 | EIF3G | ENSG00000130811 | 7.87620362 | -1.426787044 | 10.92930255 | 0.001955217 | 0.029402104 | Yes |
| 975 | CUL9 | ENSG00000112659 | 4.739182268 | 2.725629581 | 10.72747189 | 0.001958146 | 0.029402104 | Yes |
| 976 | SLC30A1 | ENSG00000170385 | 4.435407895 | 3.085749094 | 10.6996596 | 0.001958533 | 0.029402104 | Yes |
| 977 | C5orf15 | ENSG00000113583 | 6.630091995 | 2.825569218 | 10.85199038 | 0.00196703 | 0.029499408 | Yes |
| 978 | SAMD4B | ENSG00000179134 | 7.450222614 | 1.806776559 | 10.89246084 | 0.001974777 | 0.029585276 | Yes |
| 979 | PKMYT1 | ENSG00000127564 | 4.000430483 | 5.267511789 | 10.81281541 | 0.001982823 | 0.029648215 | Yes |
| 980 | DOCK11 | ENSG00000147251 | 4.63095215 | -2.379675933 | 10.7248856 | 0.001983835 | 0.029648215 | Yes |
| 981 | BEND5 | ENSG00000162373 | 2.995290645 | 6.709995197 | 10.75494279 | 0.001985055 | 0.029648215 | Yes |
| 982 | C17orf58 | ENSG00000186665 | 6.997312471 | 4.25147232 | 10.78029628 | 0.001991308 | 0.029711294 | Yes |
| 983 | TMEM138 | ENSG00000149483 | 5.512114773 | 3.143191481 | 10.72597661 | 0.001997925 | 0.029779151 | Yes |
| 984 | SLA | ENSG00000155926 | 3.783993838 | 7.578134892 | 10.86292998 | 0.002001492 | 0.029779151 | Yes |
| 985 | UBAP2L | ENSG00000143569 | 6.239688755 | 2.085166472 | 10.81520638 | 0.00200196 | 0.029779151 | Yes |
| 986 | GBP4 | ENSG00000162654 | 4.460046125 | 4.486526793 | 10.68592605 | 0.002007446 | 0.029830443 | Yes |
| 987 | POLR2J | ENSG00000005075 | 7.936060736 | -1.330713211 | 10.84982681 | 0.00202338 | 0.030003421 | Yes |
| 988 | MYO19 | ENSG00000278259 | 4.883647783 | 3.33541734 | 10.64757716 | 0.002025512 | 0.030003421 | Yes |
| 989 | DUSP3 | ENSG00000108861 | 7.216952091 | -1.830155971 | 10.83181867 | 0.002027986 | 0.030003421 | Yes |
| 990 | STK3 | ENSG00000104375 | 5.591776247 | -2.18582561 | 10.76525208 | 0.002029335 | 0.030003421 | Yes |
| 991 | SERTAD3 | ENSG00000167565 | 3.002366183 | 6.740088808 | 10.70898742 | 0.00202998 | 0.030003421 | Yes |
| 992 | LSM6 | ENSG00000164167 | 3.700109389 | 4.168357724 | 10.64402958 | 0.002032608 | 0.030003421 | Yes |
| 993 | CD47 | ENSG00000196776 | 6.408506609 | 2.42882556 | 10.77711984 | 0.002033435 | 0.030003421 | Yes |
| 994 | ARMC5 | ENSG00000140691 | 3.869889095 | -2.445476409 | 10.55149791 | 0.002038563 | 0.03004879 | Yes |
| 995 | ELMOD2 | ENSG00000179387 | 4.166015692 | 4.166019473 | 10.62481256 | 0.002045354 | 0.030118557 | Yes |
| 996 | NDUFV1 | ENSG00000167792 | 7.920800964 | -1.698753886 | 10.81613926 | 0.002051683 | 0.030181386 | Yes |
| 997 | ARL4A | ENSG00000122644 | 3.995990206 | 6.120567529 | 10.79379842 | 0.002053781 | 0.030181918 | Yes |
| 998 | CASP9 | ENSG00000132906 | 3.8861837 | 4.591316977 | 10.65734903 | 0.002059104 | 0.030229798 | Yes |
| 999 | ERLIN1 | ENSG00000107566 | 5.19973482 | 2.217034366 | 10.67166841 | 0.002061544 | 0.030235297 | Yes |
| 1000 | NANS | ENSG00000095380 | 5.496000803 | 3.75050132 | 10.62941604 | 0.002063744 | 0.030237263 | Yes |
| 1001 | TCF19 | ENSG00000137310 | 5.252329049 | 4.710656281 | 10.62135495 | 0.00206646 | 0.030246782 | Yes |
| 1002 | RNF138 | ENSG00000134758 | 4.039053749 | 3.589083274 | 10.56233442 | 0.002071503 | 0.030268067 | Yes |
| 1003 | ZNF784 | ENSG00000179922 | 4.994870455 | -2.182041729 | 10.67054394 | 0.002072051 | 0.030268067 | Yes |
| 1004 | CTSL | ENSG00000135047 | 6.386280879 | 2.797828855 | 10.71223598 | 0.002078429 | 0.030330979 | Yes |
| 1005 | RAB8B | ENSG00000166128 | 4.16125558 | 4.057017762 | 10.57313749 | 0.002085314 | 0.030401139 | Yes |
| 1006 | CD276 | ENSG00000103855 | 7.787145085 | 2.219531894 | 10.76314817 | 0.002088619 | 0.030419019 | Yes |
| 1007 | TIMM50 | ENSG00000105197 | 6.276626193 | -1.42846981 | 10.74010598 | 0.002090851 | 0.030421265 | Yes |
| 1008 | PALB2 | ENSG00000083093 | 2.750557605 | 6.459624797 | 10.58483061 | 0.002097254 | 0.030484115 | Yes |
| 1009 | EHHADH | ENSG00000113790 | 3.060504635 | 6.788936383 | 10.63836079 | 0.002101153 | 0.0304895 | Yes |
| 1010 | MCM4 | ENSG00000104738 | 6.199127137 | 2.632958542 | 10.68192819 | 0.00210179 | 0.0304895 | Yes |
| 1011 | GGT5 | ENSG00000099998 | 4.622431591 | -2.611515407 | 10.56980187 | 0.00211045 | 0.030542913 | Yes |
| 1012 | DCLK2 | ENSG00000170390 | 4.296925702 | 4.651766405 | 10.58891581 | 0.002113613 | 0.030542913 | Yes |
| 1013 | FABP3 | ENSG00000121769 | 3.844785928 | -2.775645248 | 10.4332991 | 0.002115057 | 0.030542913 | Yes |
| 1014 | C9orf72 | ENSG00000147894 | 3.322666274 | 7.125497408 | 10.67525829 | 0.002115105 | 0.030542913 | Yes |
| 1015 | TEX30 | ENSG00000151287 | 3.75392546 | 5.001477643 | 10.63869805 | 0.002115906 | 0.030542913 | Yes |
| 1016 | STMP1 | ENSG00000243317 | 6.374188867 | -1.404675616 | 10.71015743 | 0.002121076 | 0.030587383 | Yes |
| 1017 | FOXD1 | ENSG00000251493 | 3.042211515 | 6.802845636 | 10.60988797 | 0.002129665 | 0.030623606 | Yes |
| 1018 | FN1 | ENSG00000115414 | 11.57434289 | 3.89517314 | 10.74066095 | 0.002130628 | 0.030623606 | Yes |
| 1019 | SMC4 | ENSG00000113810 | 6.371419125 | 2.905599561 | 10.64957621 | 0.002130965 | 0.030623606 | Yes |
| 1020 | HELLS | ENSG00000119969 | 5.848743575 | 2.912804399 | 10.61407735 | 0.002131957 | 0.030623606 | Yes |
| 1021 | TPBG | ENSG00000146242 | 3.509703262 | 7.350312287 | 10.67116688 | 0.002147239 | 0.030812876 | Yes |
| 1022 | DNAI4 | ENSG00000152763 | 4.317704444 | 8.225359136 | 10.75556722 | 0.002153416 | 0.030851238 | Yes |
| 1023 | ASF1A | ENSG00000111875 | 2.97131719 | 6.725513591 | 10.57101015 | 0.002154128 | 0.030851238 | Yes |
| 1024 | MPZL2 | ENSG00000149573 | 5.975533014 | 7.306999434 | 10.70878508 | 0.002164485 | 0.03096927 | Yes |
| 1025 | UGCG | ENSG00000148154 | 5.769450935 | 4.397041746 | 10.50924199 | 0.002176111 | 0.031105064 | Yes |
| 1026 | IGFN1 | ENSG00000163395 | 5.511089843 | -3.053682543 | 10.56129525 | 0.00218035 | 0.031105064 | Yes |
| 1027 | ZNF263 | ENSG00000006194 | 4.511593713 | 2.741978116 | 10.46657625 | 0.002180351 | 0.031105064 | Yes |
| 1028 | MYH11 | ENSG00000133392 | 6.165778248 | -2.927817326 | 10.60092547 | 0.002188322 | 0.031133281 | Yes |
| 1029 | PRELID3B | ENSG00000101166 | 5.00420605 | 2.767031364 | 10.4954415 | 0.00218883 | 0.031133281 | Yes |
| 1030 | EXOC2 | ENSG00000112685 | 3.624205663 | 4.063293129 | 10.46702761 | 0.002189255 | 0.031133281 | Yes |
| 1031 | HAUS2 | ENSG00000137814 | 4.462200087 | 3.947214201 | 10.45954889 | 0.002190837 | 0.031133281 | Yes |
| 1032 | TK1 | ENSG00000167900 | 4.514286075 | 4.270188287 | 10.47132163 | 0.002193005 | 0.03113386 | Yes |
| 1033 | ENSG00000276855 | ENSG00000276855 | 2.985492152 | 6.723825746 | 10.52218422 | 0.002200501 | 0.03121001 | Yes |
| 1034 | ATP5MJ | ENSG00000156411 | 8.156463213 | -1.805858776 | 10.64609617 | 0.002208554 | 0.031278136 | Yes |
| 1035 | ZNF562 | ENSG00000171466 | 5.967405279 | 2.503423193 | 10.55805888 | 0.002209578 | 0.031278136 | Yes |
| 1036 | LINC02803 | ENSG00000227740 | 3.140232757 | 5.07193073 | 10.48859418 | 0.0022089 | 0.031407896 | Yes |
| 1037 | HSD11B2 | ENSG00000176387 | 6.664429533 | 3.935514776 | 10.51914963 | 0.002226509 | 0.031449816 | Yes |
| 1038 | MAP4K5 | ENSG00000012983 | 5.941339925 | 2.386684852 | 10.54127286 | 0.002228152 | 0.031449816 | Yes |
| 1039 | HSCB | ENSG00000100209 | 2.966421285 | 4.910005287 | 10.44925781 | 0.002235163 | 0.03149722 | Yes |
| 1040 | PEG10 | ENSG00000242265 | 7.431748076 | 3.771819127 | 10.55959873 | 0.002235814 | 0.03149722 | Yes |
| 1041 | ENSG00000268218 | ENSG00000268218 | 1.697309714 | -3.183265529 | 10.08955812 | 0.002246335 | 0.031614999 | Yes |
| 1042 | MCTP2 | ENSG00000140563 | 3.189450316 | 6.979055267 | 10.50941643 | 0.002252248 | 0.031667774 | Yes |
| 1043 | HIP1R | ENSG00000130787 | 3.453296739 | 5.519447173 | 10.50516239 | 0.002257727 | 0.031714352 | Yes |
| 1044 | YRDC | ENSG00000196449 | 3.997361515 | 4.235688357 | 10.40653291 | 0.002261894 | 0.031742419 | Yes |
| 1045 | MPZL1 | ENSG00000197965 | 6.289161398 | 2.54528392 | 10.51847978 | 0.002264666 | 0.031750881 | Yes |
| 1046 | OSGIN1 | ENSG00000140961 | 3.072667428 | -2.683152844 | 10.14032546 | 0.002276618 | 0.031887895 | Yes |
| 1047 | NPTXR | ENSG00000221890 | 5.806930942 | 4.386850688 | 10.40078313 | 0.002284794 | 0.031921703 | Yes |
| 1048 | LINC01138 | ENSG00000274020 | 3.277225743 | 6.97914258 | 10.47625511 | 0.002285008 | 0.031921703 | Yes |
| 1049 | CROCC | ENSG00000058453 | 5.110242237 | 3.246655098 | 10.38961087 | 0.002285574 | 0.031921703 | Yes |
| 1050 | ARAP1 | ENSG00000186635 | 7.236080022 | 2.504572023 | 10.53164287 | 0.002291062 | 0.031947163 | Yes |
| 1051 | ZNF155 | ENSG00000204920 | 3.299245875 | 7.109544593 | 10.48843521 | 0.002291762 | 0.031947163 | Yes |
| 1052 | SLC37A4 | ENSG00000137700 | 4.846002963 | -2.051220894 | 10.42476434 | 0.00229937 | 0.032008714 | Yes |
| 1053 | FKBP3 | ENSG00000100442 | 6.222794663 | -1.889962203 | 10.5105559 | 0.002301868 | 0.032008714 | Yes |
| 1054 | NONO | ENSG00000147140 | 7.740127301 | 2.388093673 | 10.53338228 | 0.002302738 | 0.032008714 | Yes |
| 1055 | TMEM216 | ENSG00000187049 | 3.341568853 | 7.139197439 | 10.46219212 | 0.002322294 | 0.032249918 | Yes |
| 1056 | SAMD9 | ENSG00000205413 | 4.876616491 | 4.027901174 | 10.33226923 | 0.002329218 | 0.032289336 | Yes |
| 1057 | TADA2B | ENSG00000173011 | 4.802223753 | -2.005255493 | 10.3929352 | 0.002329544 | 0.032289336 | Yes |
| 1058 | CNIH3 | ENSG00000143786 | 2.820947842 | 6.522135944 | 10.34954773 | 0.002338938 | 0.032371299 | Yes |
| 1059 | ENO1 | ENSG00000074800 | 8.711423422 | 2.150150411 | 10.50953413 | 0.00234177 | 0.032371299 | Yes |
| 1060 | TMEM62 | ENSG00000137842 | 5.18252378 | -1.954723674 | 10.41476616 | 0.002342092 | 0.032371299 | Yes |
| 1061 | CAPZA1 | ENSG00000116489 | 7.25597772 | 2.321002083 | 10.47957222 | 0.002347147 | 0.032410558 | Yes |
| 1062 | ZNF429 | ENSG00000197013 | 3.102484721 | 6.831373636 | 10.38869744 | 0.002351209 | 0.032436046 | Yes |
| 1063 | FAM13B | ENSG00000031003 | 4.235271013 | 3.577566749 | 10.27417452 | 0.002362168 | 0.032556545 | Yes |
| 1064 | ENSG00000274383 | ENSG00000274383 | 4.35315771 | 3.176544079 | 10.26843206 | 0.002367795 | 0.032603399 | Yes |
| 1065 | ISG20L2 | ENSG00000143319 | 5.194389274 | 2.635359116 | 10.32392855 | 0.002382829 | 0.032773778 | Yes |
| 1066 | DUOX2 | ENSG00000140279 | 4.327624207 | 8.22125033 | 10.51600174 | 0.002384647 | 0.032773778 | Yes |
| 1067 | PPARD | ENSG00000112033 | 4.67127709 | 3.315625315 | 10.26623613 | 0.002388048 | 0.03278973 | Yes |
| 1068 | PRIMPOL | ENSG00000164306 | 3.158644428 | 6.875719447 | 10.33864285 | 0.00241068 | 0.033052256 | Yes |
| 1069 | MASTL | ENSG00000120539 | 2.856481998 | 6.566435984 | 10.28810936 | 0.002411683 | 0.033052256 | Yes |
| 1070 | ZNF766 | ENSG00000196214 | 3.507166235 | 4.212176778 | 10.2533175 | 0.002424143 | 0.033173084 | Yes |
| 1071 | GDPD5 | ENSG00000158555 | 4.13215335 | -2.003880836 | 10.22813009 | 0.002425032 | 0.033173084 | Yes |
| 1072 | INCA1 | ENSG00000196388 | 3.386610937 | -2.367776629 | 10.08100009 | 0.002432781 | 0.033231648 | Yes |
| 1073 | B4GAT1 | ENSG00000174684 | 5.963249611 | -1.867460635 | 10.37189202 | 0.002433854 | 0.033231648 | Yes |
| 1074 | CRTC2 | ENSG00000160741 | 3.081687604 | 6.812154677 | 10.30495408 | 0.002436293 | 0.03323394 | Yes |
| 1075 | LEMD2 | ENSG00000161904 | 6.281186588 | 2.90069091 | 10.3185502 | 0.002456477 | 0.033478075 | Yes |
| 1076 | TRIL | ENSG00000255690 | 5.613254666 | 4.798892133 | 10.22530314 | 0.002466079 | 0.033577675 | Yes |
| 1077 | RFK | ENSG00000135002 | 4.562908792 | 3.730481085 | 10.18741396 | 0.002470741 | 0.033609889 | Yes |
| 1078 | ITPRIP | ENSG00000148841 | 4.779333405 | 3.414575939 | 10.18333324 | 0.00248256 | 0.03373071 | Yes |
| 1079 | MSH6 | ENSG00000116062 | 5.614817748 | 2.304423388 | 10.27434621 | 0.002484346 | 0.03373071 | Yes |
| 1080 | SH2D4A | ENSG00000104611 | 3.67987246 | 7.541202591 | 10.35493933 | 0.002486537 | 0.03373071 | Yes |
| 1081 | SSU72 | ENSG00000160075 | 7.533017977 | -1.313438285 | 10.35564485 | 0.002497529 | 0.033820968 | Yes |
| 1082 | MZB1 | ENSG00000170476 | 3.653559304 | -3.253609772 | 9.98905508 | 0.002497811 | 0.033820968 | Yes |
| 1083 | FERMT3 | ENSG00000149781 | 3.405012446 | 7.210346954 | 10.29804164 | 0.002505101 | 0.033882096 | Yes |
| 1084 | OSBPL1A | ENSG00000141447 | 5.64798393 | -1.759583963 | 10.29135747 | 0.002506956 | 0.033882096 | Yes |
| 1085 | TMEM167A | ENSG00000174695 | 5.987753836 | 2.481724361 | 10.26749165 | 0.002511108 | 0.033906909 | Yes |
| 1086 | LOC105372295 | ENSG00000279529 | 3.074678685 | 6.822489396 | 10.22700028 | 0.002523286 | 0.034039943 | Yes |
| 1087 | TNFAIP6 | ENSG00000123610 | 5.229620549 | 4.796346683 | 10.16749224 | 0.002529983 | 0.034098853 | Yes |
| 1088 | EXOSC8 | ENSG00000120699 | 5.322620022 | 2.09920677 | 10.21713672 | 0.002533992 | 0.034121475 | Yes |
| 1089 | SCARF2 | ENSG00000244486 | 3.227329428 | 5.214640336 | 10.20463457 | 0.002536758 | 0.034127317 | Yes |
| 1090 | PTPRH | ENSG00000080031 | 4.248577606 | 8.146363971 | 10.36308196 | 0.002539801 | 0.034136882 | Yes |
| 1091 | DFFB | ENSG00000169598 | 3.674718135 | 5.704094652 | 10.25441464 | 0.00254976 | 0.034239294 | Yes |
| 1092 | ZBTB16 | ENSG00000109906 | 5.892419831 | -2.613133555 | 10.24173892 | 0.00255381 | 0.034262249 | Yes |
| 1093 | C4orf46 | ENSG00000205208 | 4.636070886 | 3.875024148 | 10.10801268 | 0.002565334 | 0.0343383 | Yes |
| 1094 | SP140L | ENSG00000185404 | 4.833943586 | 2.951949607 | 10.12001425 | 0.002566077 | 0.0343383 | Yes |
| 1095 | CYP4F3 | ENSG00000186529 | 3.496111658 | -3.125476186 | 9.913749575 | 0.002566516 | 0.0343383 | Yes |
| 1096 | LOC124900372 | ENSG00000261971 | 2.933427286 | 6.644368229 | 10.15238392 | 0.002576059 | 0.034434495 | Yes |
| 1097 | NUP93 | ENSG00000102900 | 6.938121246 | 2.357063342 | 10.24790162 | 0.002584349 | 0.034513787 | Yes |
| 1098 | CENPQ | ENSG00000031691 | 2.815167817 | 6.551269893 | 10.12632629 | 0.002588654 | 0.034539767 | Yes |
| 1099 | TLE3 | ENSG00000140332 | 5.515044228 | 2.488707718 | 10.16065191 | 0.002595487 | 0.03457038 | Yes |
| 1100 | SLC7A1 | ENSG00000139514 | 4.651881535 | 3.573386648 | 10.07901007 | 0.002595672 | 0.03457038 | Yes |
| 1101 | ACOX3 | ENSG00000087008 | 4.831930399 | 2.796569771 | 10.09539263 | 0.002598985 | 0.034583046 | Yes |
| 1102 | MCM5 | ENSG00000100297 | 5.792640766 | 2.887500886 | 10.15097012 | 0.002608921 | 0.034683725 | Yes |
| 1103 | CEP152 | ENSG00000103995 | 5.238397658 | 2.862460098 | 10.09841294 | 0.002626382 | 0.034884169 | Yes |
| 1104 | ENSG00000213888 | ENSG00000213888 | 2.651375825 | 6.324467086 | 10.03939363 | 0.00264534 | 0.035085464 | Yes |
| 1105 | ZNF202 | ENSG00000166261 | 3.404358897 | 4.612566342 | 10.08902524 | 0.002646331 | 0.035085464 | Yes |
| 1106 | S100A2 | ENSG00000196754 | 8.476098044 | 10.73965612 | 10.32915812 | 0.002654822 | 0.035166184 | Yes |
| 1107 | PTN | ENSG00000105894 | 5.582260926 | 4.269159799 | 10.04815104 | 0.002666046 | 0.03528293 | Yes |
| 1108 | ABT1 | ENSG00000146109 | 6.066107282 | 2.601595272 | 10.12372766 | 0.002674872 | 0.035367756 | Yes |
| 1109 | GNL1 | ENSG00000204590 | 6.956181992 | 2.545112469 | 10.15592275 | 0.002686373 | 0.03548776 | Yes |
| 1110 | ZIC4 | ENSG00000174963 | 3.431500209 | 7.22327844 | 10.12367169 | 0.002705752 | 0.03563911 | Yes |
| 1111 | POLR2G | ENSG00000168002 | 6.733691258 | 1.851413135 | 10.14400359 | 0.002707347 | 0.03563911 | Yes |
| 1112 | TC2N | ENSG00000165929 | 3.92611667 | 7.801030145 | 10.18505905 | 0.002707599 | 0.03563911 | Yes |
| 1113 | FBXO8 | ENSG00000164117 | 3.085406432 | 6.862944686 | 10.07207967 | 0.002709335 | 0.03563911 | Yes |
| 1114 | DHX34 | ENSG00000134815 | 3.786507154 | 4.234456285 | 10.00329416 | 0.002710004 | 0.03563911 | Yes |
| 1115 | CILP | ENSG00000138615 | 5.636448113 | -2.314983869 | 10.09375276 | 0.002719613 | 0.035717351 | Yes |
| 1116 | PBX4 | ENSG00000105717 | 3.157948385 | 6.944166631 | 10.07407864 | 0.002720834 | 0.035717351 | Yes |
| 1117 | FBXO5 | ENSG00000112029 | 2.766982271 | 6.49857564 | 9.999353689 | 0.002729024 | 0.035784076 | Yes |
| 1118 | CADPS | ENSG00000163618 | 2.573805559 | -2.931875863 | 9.63327855 | 0.002731643 | 0.035784076 | Yes |
| 1119 | ZNF143.AS1 | | 2.800054006 | 6.513476312 | 9.998293628 | 0.002733251 | 0.035784076 | Yes |
| 1120 | ZNF300 | ENSG00000145908 | 4.639081123 | 3.469966621 | 9.954465186 | 0.002744611 | 0.035900689 | Yes |
| 1121 | MDGA1 | ENSG00000112139 | 5.668447963 | 3.621173677 | 9.992976729 | 0.002752531 | 0.035972137 | Yes |
| 1122 | NGRN | ENSG00000182768 | 4.993639424 | -1.986517824 | 10.02713382 | 0.002760071 | 0.0360385 | Yes |
| 1123 | PON2 | ENSG00000105854 | 5.74851915 | 2.803249758 | 10.01449875 | 0.002773234 | 0.036178096 | Yes |
| 1124 | CKMT2.AS1 | | 3.801644098 | 5.04240277 | 10.01504171 | 0.002787066 | 0.036326168 | Yes |
| 1125 | ZNF169 | ENSG00000175787 | 2.360255676 | 5.957161193 | 9.855151374 | 0.002791303 | 0.03634901 | Yes |
| 1126 | BLOC1S4 | ENSG00000186222 | 4.643556315 | 4.065543162 | 9.91666472 | 0.002801177 | 0.036425667 | Yes |
| 1127 | ZNF793 | ENSG00000188227 | 5.389516253 | -2.267027965 | 10.01202115 | 0.002802166 | 0.036425667 | Yes |
| 1128 | IFIT1 | ENSG00000185745 | 4.361424536 | 4.389676586 | 9.928382517 | 0.00280561 | 0.036438083 | Yes |
| 1129 | ZBTB34 | ENSG00000177125 | 3.683727736 | 4.387672958 | 9.935432973 | 0.002814205 | 0.036517302 | Yes |
| 1130 | CTDP1 | ENSG00000060069 | 4.609910867 | 3.310676794 | 9.893586212 | 0.002818331 | 0.036535653 | Yes |
| 1131 | ERRFI1 | ENSG00000116285 | 5.301490657 | 3.346398679 | 9.927054477 | 0.002820611 | 0.036535653 | Yes |
| 1132 | ADAMTS7 | ENSG00000136378 | 5.1051405 | 3.97057397 | 9.89448805 | 0.002839814 | 0.036742055 | Yes |
| 1133 | IRX1 | ENSG00000170549 | 5.124150494 | 5.396305366 | 9.945619461 | 0.002841566 | 0.036742055 | Yes |
| 1134 | HCN3 | ENSG00000143630 | 3.609196095 | 4.872237993 | 9.953598601 | 0.002844374 | 0.036745901 | Yes |
| 1135 | FUOM | ENSG00000148803 | 3.070040644 | 5.042356755 | 9.92120853 | 0.002847599 | 0.036755125 | Yes |
| 1136 | PGAP1 | ENSG00000197121 | 5.606181466 | 2.991680393 | 9.932177868 | 0.002852814 | 0.036771232 | Yes |
| 1137 | COMMD6 | ENSG00000188243 | 7.20043076 | 2.52424343 | 10.02410229 | 0.002855897 | 0.036771232 | Yes |
| 1138 | ATP1A1 | ENSG00000163399 | 7.862776965 | 2.6424961 | 10.03620198 | 0.002856601 | 0.036771232 | Yes |
| 1139 | ENSG00000272086 | ENSG00000272086 | 3.575821704 | 5.651649597 | 9.985338268 | 0.00286029 | 0.036771232 | Yes |
| 1140 | IDNK | ENSG00000148057 | 2.825092333 | 6.530788929 | 9.898572338 | 0.002861635 | 0.036771232 | Yes |
| 1141 | UBA1 | ENSG00000130985 | 8.894376877 | 1.803171903 | 10.0479638 | 0.002863921 | 0.036771232 | Yes |
| 1142 | REX1BD | ENSG00000006015 | 6.983946591 | -1.426242046 | 10.02409184 | 0.00287396 | 0.036824866 | Yes |
| 1143 | PFDN2 | ENSG00000143256 | 6.995344674 | 2.492678654 | 10.0031045 | 0.002875311 | 0.036824866 | Yes |
| 1144 | FBXL7 | ENSG00000183580 | 3.899358148 | 3.845371766 | 9.841526019 | 0.002875645 | 0.036824866 | Yes |
| 1145 | SLC31A1 | ENSG00000136868 | 4.6563459 | 3.756731466 | 9.847709719 | 0.002882093 | 0.036875177 | Yes |
| 1146 | GSTM5 | ENSG00000134201 | 3.562150424 | -3.062690244 | 9.67517718 | 0.002888433 | 0.036914199 | Yes |
| 1147 | GPC6 | ENSG00000183098 | 5.570629638 | 4.139693406 | 9.868915021 | 0.002890187 | 0.036914199 | Yes |
| 1148 | SGO2 | ENSG00000163535 | 4.368932931 | 3.972659649 | 9.837912699 | 0.002893907 | 0.03692948 | Yes |
| 1149 | TMEM243 | ENSG00000135185 | 4.199255388 | 3.239901811 | 9.813737821 | 0.002897793 | 0.036940092 | Yes |
| 1150 | KIF23 | ENSG00000137807 | 3.264785321 | 7.076283376 | 9.947704521 | 0.002899786 | 0.036940092 | Yes |
| 1151 | NOTCH1 | ENSG00000148400 | 4.276785208 | 4.288032107 | 9.845217001 | 0.00290551 | 0.036944384 | Yes |
| 1152 | CHPF | ENSG00000123989 | 6.618463935 | 2.115125451 | 9.971913681 | 0.002908709 | 0.036944384 | Yes |
| 1153 | RCL1 | ENSG00000120158 | 4.855482018 | -2.125646489 | 9.893810507 | 0.002909093 | 0.036944384 | Yes |
| 1154 | PHTF2 | ENSG00000006576 | 6.041448763 | -2.083250239 | 9.963334905 | 0.00291078 | 0.036944384 | Yes |
| 1155 | CCT6B | ENSG00000132141 | 2.983102272 | -2.457722077 | 9.617929131 | 0.002913985 | 0.036944384 | Yes |
| 1156 | PFAS | ENSG00000178921 | 4.765144814 | 3.113139416 | 9.827290654 | 0.002915359 | 0.036944384 | Yes |
| 1157 | MOB1B | ENSG00000173542 | 2.79936892 | 6.431830886 | 9.839224886 | 0.002917791 | 0.036944384 | Yes |
| 1158 | LHX2 | ENSG00000106689 | 4.674329869 | 6.08681318 | 9.976841247 | 0.002924598 | 0.036965988 | Yes |
| 1159 | ENSG00000167747 | ENSG00000167747 | 5.547345019 | 3.008643354 | 9.872562779 | 0.002925004 | 0.036965988 | Yes |
| 1160 | XNDC1N | ENSG00000254469 | 2.894849128 | 6.600229089 | 9.857978937 | 0.002928357 | 0.036965988 | Yes |
| 1161 | USP45 | ENSG00000123552 | 2.462538195 | 6.105760741 | 9.775457303 | 0.002929599 | 0.036965988 | Yes |
| 1162 | CLYBL | ENSG00000125246 | 3.606726065 | -2.663086684 | 9.676822569 | 0.002937994 | 0.037039986 | Yes |
| 1163 | NCSTN | ENSG00000162736 | 5.42952819 | 3.402163491 | 9.838787018 | 0.002941397 | 0.037050974 | Yes |
| 1164 | RIMOC1 | ENSG00000205765 | 3.026926058 | 5.029617407 | 9.837760381 | 0.002953722 | 0.037174232 | Yes |
| 1165 | NAT9 | ENSG00000109065 | 4.791831303 | 2.929458132 | 9.797590724 | 0.0029629 | 0.037217857 | Yes |
| 1166 | SPTLC3 | ENSG00000172296 | 3.360583682 | 7.180811959 | 9.912130351 | 0.002962929 | 0.037217857 | Yes |
| 1167 | LDHD | ENSG00000166816 | 4.450596046 | -2.95141064 | 9.762301947 | 0.002964817 | 0.037217857 | Yes |
| 1168 | RNASET2 | ENSG00000026297 | 6.653219832 | 2.891522425 | 9.899823806 | 0.002978608 | 0.037335547 | Yes |
| 1169 | CTDNEP1 | ENSG00000175826 | 8.332785079 | -1.5462349 | 9.954787405 | 0.002981714 | 0.037335547 | Yes |
| 1170 | PTPN2 | ENSG00000175354 | 5.189961529 | 3.031041134 | 9.805618164 | 0.002981844 | 0.037335547 | Yes |
| 1171 | BTN2A1 | ENSG00000112763 | 3.853220984 | 3.49432074 | 9.737246274 | 0.002990604 | 0.037391496 | Yes |
| 1172 | CLN5 | ENSG00000102805 | 4.351891364 | 4.06923074 | 9.76789249 | 0.002992279 | 0.037391496 | Yes |
| 1173 | TAF2 | ENSG00000064313 | 5.397364897 | 2.421188281 | 9.833585761 | 0.002993976 | 0.037391496 | Yes |
| 1174 | AZIN1 | ENSG00000155096 | 5.860069022 | 2.44213331 | 9.860351805 | 0.00299711 | 0.037398719 | Yes |
| 1175 | PMAIP1 | ENSG00000141682 | 3.259465606 | 7.053962175 | 9.862843376 | 0.003006908 | 0.037483159 | Yes |
| 1176 | CRNDE | ENSG00000245694 | 4.870016307 | 4.239141858 | 9.763160965 | 0.003011134 | 0.037483159 | Yes |
| 1177 | ERO1B | ENSG00000086619 | 3.224527111 | 4.425286835 | 9.775397638 | 0.003016339 | 0.037483159 | Yes |
| 1178 | ZBTB48 | ENSG00000204859 | 4.326661635 | -2.056657535 | 9.763909011 | 0.003016435 | 0.037483159 | Yes |
| 1179 | WDR45B | ENSG00000141580 | 6.50850068 | 2.531993144 | 9.86902939 | 0.003025078 | 0.037483159 | Yes |
| 1180 | ACO2 | ENSG00000100412 | 7.293629748 | -1.71373103 | 9.909317381 | 0.003026719 | 0.037483159 | Yes |
| 1181 | KNL1 | ENSG00000137812 | 3.618415412 | 4.883815773 | 9.8142181 | 0.003027954 | 0.037483159 | Yes |
| 1182 | DCBLD2 | ENSG00000057019 | 6.011649303 | 3.035004099 | 9.820228086 | 0.003032208 | 0.037483159 | Yes |
| 1183 | VMP1 | ENSG00000062716 | 7.400210636 | 2.846917295 | 9.886493202 | 0.003032697 | 0.037483159 | Yes |
| 1184 | CFAP36 | ENSG00000163001 | 4.627631411 | 4.142296808 | 9.742322111 | 0.003032951 | 0.037483159 | Yes |
| 1185 | PNMA1 | ENSG00000176903 | 5.355876793 | 2.973176787 | 9.779880823 | 0.00303447 | 0.037483159 | Yes |
| 1186 | TDP1 | ENSG00000042088 | 4.318244427 | 3.331321672 | 9.717718072 | 0.003034607 | 0.037483159 | Yes |
| 1187 | ZNF585B | ENSG00000245680 | 2.919831829 | 6.670022345 | 9.781904716 | 0.00304437 | 0.03757204 | Yes |
| 1188 | NLRX1 | ENSG00000160703 | 4.470368896 | -2.479731461 | 9.720962276 | 0.003066071 | 0.037807982 | Yes |
| 1189 | POLR2L | ENSG00000177700 | 8.434600804 | -1.363807141 | 9.886486612 | 0.003073265 | 0.0378648 | Yes |
| 1190 | ENSG00000251867 | ENSG00000251867 | 2.69764834 | 6.364971079 | 9.710135107 | 0.003077854 | 0.037889446 | Yes |
| 1191 | S1PR3 | ENSG00000213694 | 3.247509112 | 7.041832776 | 9.804129056 | 0.003084114 | 0.037914256 | Yes |
| 1192 | CEP55 | ENSG00000138180 | 3.116372962 | 6.891452722 | 9.781923401 | 0.003087425 | 0.037914256 | Yes |
| 1193 | DNAAF2 | ENSG00000165506 | 2.697261649 | 6.380471608 | 9.704975657 | 0.003088556 | 0.037914256 | Yes |
| 1194 | LOXL1 | ENSG00000129038 | 7.013354535 | 3.44707366 | 9.811748942 | 0.003094045 | 0.037914256 | Yes |
| 1195 | CPXM1 | ENSG00000088882 | 5.026566582 | 5.292664677 | 9.751047413 | 0.003095096 | 0.037914256 | Yes |
| 1196 | GABARAPL1 | ENSG00000139112 | 6.839210905 | -1.780763557 | 9.847157901 | 0.0030994 | 0.037914256 | Yes |
| 1197 | GPRIN1 | ENSG00000169258 | 3.028673461 | 6.801048158 | 9.760001359 | 0.003100494 | 0.037914256 | Yes |
| 1198 | CDH2 | ENSG00000170558 | 4.5188223 | 5.279246636 | 9.775127882 | 0.003101733 | 0.037914256 | Yes |
| 1199 | RYR3 | ENSG00000198838 | 3.20569776 | -2.987455406 | 9.465944851 | 0.003103182 | 0.037914256 | Yes |
| 1200 | NR3C2 | ENSG00000151623 | 1.949024884 | -2.900774065 | 9.326568218 | 0.003108871 | 0.037918148 | Yes |
| 1201 | ACKR2 | ENSG00000144648 | 3.588853229 | -2.782275127 | 9.554989377 | 0.003110347 | 0.037918148 | Yes |
| 1202 | TRRAP | ENSG00000196367 | 2.301678941 | 5.907640084 | 9.609246389 | 0.003111273 | 0.037918148 | Yes |
| 1203 | MEX3B | ENSG00000183496 | 3.019381228 | 6.784551131 | 9.734616818 | 0.003132616 | 0.038146508 | Yes |
| 1204 | ZNF439 | ENSG00000171291 | 2.936324358 | 6.698553888 | 9.717352461 | 0.00313962 | 0.038156088 | Yes |
| 1205 | LRRC24 | ENSG00000254402 | 3.703338236 | 7.493489101 | 9.814132645 | 0.003142025 | 0.038156088 | Yes |
| 1206 | INMT | ENSG00000241644 | 4.021785611 | -2.617754223 | 9.604134756 | 0.003142408 | 0.038156088 | Yes |

| | A | B | C | D | E | F | G | H |
|---|---|---|---|---|---|---|---|---|
| 1207 | PI16 | ENSG00000164530 | 4.95564205 | -2.68738421 | 9.70589264 | 0.00314383 | 0.038156088 | Yes |
| 1208 | SNAPC4 | ENSG00000165684 | 5.661760153 | 2.645847376 | 9.727364679 | 0.003151973 | 0.038196067 | Yes |
| 1209 | PCYOX1L | ENSG00000145882 | 3.295749542 | 5.347757031 | 9.732413309 | 0.003152343 | 0.038196067 | Yes |
| 1210 | ATP6V1C1 | ENSG00000155097 | 5.417836588 | 2.378825204 | 9.716893554 | 0.003158212 | 0.038235521 | Yes |
| 1211 | PPIA | ENSG00000196262 | 7.781890624 | 1.582825439 | 9.809806545 | 0.003166136 | 0.038299775 | Yes |
| 1212 | ZNF341 | ENSG00000131061 | 4.027237278 | -2.283454945 | 9.601003107 | 0.00318123 | 0.038450586 | Yes |
| 1213 | GK | ENSG00000198814 | 2.995226922 | 6.760995395 | 9.686956498 | 0.003195428 | 0.038590325 | Yes |
| 1214 | PPP1R18 | ENSG00000146112 | 8.155750988 | 2.351908185 | 9.784433434 | 0.003200354 | 0.038617953 | Yes |
| 1215 | ACKR1 | ENSG00000213088 | 5.39322234 | -2.116940947 | 9.712027658 | 0.003206711 | 0.038662794 | Yes |
| 1216 | EIF5A2 | ENSG00000163577 | 3.03632908 | 4.974967204 | 9.64702575 | 0.003211489 | 0.038674443 | Yes |
| 1217 | GAP43 | ENSG00000172020 | 4.856225567 | 8.771658694 | 9.857290519 | 0.003213226 | 0.038674443 | Yes |
| 1218 | STC2 | ENSG00000113739 | 4.172333751 | 4.936308318 | 9.676907333 | 0.003215604 | 0.038674443 | Yes |
| 1219 | NFAM1 | ENSG00000235568 | 3.093899543 | 6.845511317 | 9.682157083 | 0.003219026 | 0.038683817 | Yes |
| 1220 | ADCY7 | ENSG00000121281 | 3.977815888 | 3.452902934 | 9.574557045 | 0.003224233 | 0.038714605 | Yes |
| 1221 | REEP4 | ENSG00000168476 | 6.717846663 | 2.612667059 | 9.725094899 | 0.003232821 | 0.038736268 | Yes |
| 1222 | RASL11B | ENSG00000128045 | 2.945420969 | 6.670764904 | 9.645721906 | 0.003236275 | 0.038736268 | Yes |
| 1223 | PFKFB3 | ENSG00000170525 | 6.135030636 | -2.138842159 | 9.725716627 | 0.0032363 | 0.038736268 | Yes |
| 1224 | ARL2 | ENSG00000213465 | 5.905098393 | -1.541742602 | 9.722722206 | 0.003239397 | 0.038736268 | Yes |
| 1225 | UMAD1 | ENSG00000219545 | 3.687004559 | 4.305646126 | 9.615008287 | 0.00324104 | 0.038736268 | Yes |
| 1226 | OGFR | ENSG00000060491 | 6.707202051 | 2.101868297 | 9.729065119 | 0.003241916 | 0.038736268 | Yes |
| 1227 | RAB2B | ENSG00000129472 | 5.557931459 | 3.216280904 | 9.627719869 | 0.003254031 | 0.038848341 | Yes |
| 1228 | DIAPH3 | ENSG00000139734 | 3.408282319 | 7.224662394 | 9.703726867 | 0.003256604 | 0.038848341 | Yes |
| 1229 | ISOC1 | ENSG00000066583 | 4.665931392 | 3.413205076 | 9.564365641 | 0.003275086 | 0.038972083 | Yes |
| 1230 | MANSC1 | ENSG00000111261 | 2.658171006 | 6.358093782 | 9.570922754 | 0.003276703 | 0.038972083 | Yes |
| 1231 | YWHAZ | ENSG00000164924 | 8.253290222 | 1.692581431 | 9.735333192 | 0.003277555 | 0.038972083 | Yes |
| 1232 | FGGY | ENSG00000172456 | 5.002729626 | -2.672819268 | 9.617356245 | 0.003279084 | 0.038972083 | Yes |
| 1233 | RHEBL1 | ENSG00000167550 | 2.558282152 | 6.236030726 | 9.548942614 | 0.00328029 | 0.038972083 | Yes |
| 1234 | C21orf58 | ENSG00000160298 | 5.252189228 | 2.698500824 | 9.604553341 | 0.003286801 | 0.038979198 | Yes |
| 1235 | SDHD | ENSG00000204370 | 5.636179443 | -1.828229318 | 9.674805769 | 0.003287636 | 0.038979198 | Yes |
| 1236 | DCK | ENSG00000156136 | 3.071184592 | 6.80212388 | 9.628281859 | 0.003288878 | 0.038979198 | Yes |
| 1237 | SLC75A1 | ENSG00000109736 | 6.804223595 | 2.107184218 | 9.682832804 | 0.00331289 | 0.039232021 | Yes |
| 1238 | SLC22A15 | ENSG00000163393 | 2.537376552 | 6.196290467 | 9.513856996 | 0.003323348 | 0.039300551 | Yes |
| 1239 | KLHL18 | ENSG00000114648 | 4.228788529 | -2.452603233 | 9.515556003 | 0.003325962 | 0.039300551 | Yes |
| 1240 | ZNF628 | ENSG00000197483 | 2.442627987 | -2.27409426 | 9.250945661 | 0.003326732 | 0.039300551 | Yes |
| 1241 | C15orf39 | ENSG00000167173 | 4.871454757 | 2.64423841 | 9.551055275 | 0.003331118 | 0.039320621 | Yes |
| 1242 | SMOX | ENSG00000088826 | 4.259990489 | 5.035695787 | 9.598055266 | 0.003339742 | 0.039346941 | Yes |
| 1243 | KLF9 | ENSG00000119138 | 5.94460587 | -2.047401118 | 9.647906909 | 0.003341133 | 0.039346941 | Yes |
| 1244 | APOBEC3D | ENSG00000243811 | 3.04249565 | 6.793788286 | 9.591836936 | 0.003341412 | 0.039346941 | Yes |
| 1245 | VSIG10L | ENSG00000186806 | 2.60074528 | 6.247892478 | 9.502164899 | 0.003353781 | 0.039413013 | Yes |
| 1246 | ZFHX3.AS1 | | 2.339237854 | 5.919605712 | 9.447580771 | 0.003354496 | 0.039413013 | Yes |
| 1247 | MSRB2 | ENSG00000148450 | 7.243390977 | -1.629036068 | 9.67471633 | 0.003355101 | 0.039413013 | Yes |
| 1248 | ENSG00000237753 | ENSG00000237753 | 3.255425399 | -2.410583543 | 9.340449917 | 0.00339292 | 0.039814631 | Yes |
| 1249 | GINS4 | ENSG00000147536 | 3.117514527 | 5.176794066 | 9.538778828 | 0.00339473 | 0.039814631 | Yes |
| 1250 | RFWD3 | ENSG00000168411 | 5.64547784 | 2.922736319 | 9.545281952 | 0.003401725 | 0.039845557 | Yes |
| 1251 | ZNF575 | ENSG00000176472 | 2.933803024 | -2.209264597 | 9.292925209 | 0.003402811 | 0.039845557 | Yes |
| 1252 | TMEM45A | ENSG00000181458 | 5.765082045 | 4.099930918 | 9.501169328 | 0.003422767 | 0.040041159 | Yes |
| 1253 | KLHL35 | ENSG00000149243 | 3.073426536 | 6.832951994 | 9.54206492 | 0.003424987 | 0.040041159 | Yes |
| 1254 | VCAM1 | ENSG00000162692 | 5.670402819 | 4.054783518 | 9.491863243 | 0.003432763 | 0.040100038 | Yes |
| 1255 | MGAT2 | ENSG00000168282 | 3.317928015 | 5.37468785 | 9.542521701 | 0.003436551 | 0.040104797 | Yes |
| 1256 | ZNF8.DT | | 3.73512591 | 4.446430404 | 9.494885362 | 0.00343865 | 0.040104797 | Yes |
| 1257 | SDF2 | ENSG00000132581 | 5.625587601 | 2.20117929 | 9.53634343 | 0.003455141 | 0.040214372 | Yes |
| 1258 | SPECC1 | ENSG00000128487 | 3.776593819 | 7.633294874 | 9.611901399 | 0.003455712 | 0.040214372 | Yes |
| 1259 | FAM111A.DT | | 2.280634594 | 5.860731947 | 9.373010359 | 0.003456287 | 0.040214372 | Yes |
| 1260 | FTSJ1 | ENSG00000068438 | 4.858023163 | 2.89053829 | 9.458516011 | 0.003462558 | 0.040228362 | Yes |
| 1261 | NIPAL2 | ENSG00000104361 | 3.733838552 | 4.043470431 | 9.443233703 | 0.003467734 | 0.040228362 | Yes |
| 1262 | SLC39A6 | ENSG00000141424 | 6.328845551 | 2.350927368 | 9.556803696 | 0.00347026 | 0.040228362 | Yes |
| 1263 | RAD51AP1 | ENSG00000111247 | 2.993139674 | 6.756367248 | 9.499959152 | 0.003474354 | 0.040228362 | Yes |
| 1264 | C5 | ENSG00000106804 | 2.807601304 | 6.506579239 | 9.463658423 | 0.003475473 | 0.040228362 | Yes |
| 1265 | BDH2 | ENSG00000164039 | 4.13275143 | 4.040449828 | 9.436025599 | 0.003476361 | 0.040228362 | Yes |
| 1266 | SCLT1 | ENSG00000151466 | 3.670182391 | 4.322501255 | 9.46117287 | 0.003476729 | 0.040228362 | Yes |
| 1267 | NTN4 | ENSG00000074527 | 4.2988763 | 4.207662093 | 9.441002971 | 0.003480308 | 0.040237967 | Yes |
| 1268 | TMEM229B | ENSG00000198133 | 3.766900757 | 7.637283332 | 9.588716647 | 0.003491851 | 0.040332674 | Yes |
| 1269 | FHAD1 | ENSG00000142621 | 4.883718578 | 8.816059573 | 9.663305817 | 0.003498711 | 0.040332674 | Yes |
| 1270 | CDS1 | ENSG00000163624 | 5.367952227 | 9.314142266 | 9.682679087 | 0.003499283 | 0.040332674 | Yes |
| 1271 | ENSG00000259865 | ENSG00000259865 | 2.648107199 | 6.326559217 | 9.420101343 | 0.003501473 | 0.040332674 | Yes |
| 1272 | ENSG00000276101 | ENSG00000276101 | 2.296090676 | 5.857602276 | 9.343857624 | 0.003502277 | 0.040332674 | Yes |
| 1273 | SESN3 | ENSG00000149212 | 5.069004493 | 2.919023157 | 9.430243248 | 0.003525558 | 0.04054822 | Yes |
| 1274 | NEK6 | ENSG00000119408 | 5.152446417 | 2.819858297 | 9.437237633 | 0.003528811 | 0.04054822 | Yes |
| 1275 | ENSG00000272572 | ENSG00000272572 | 2.885388287 | 6.608573003 | 9.44311687 | 0.003531581 | 0.04054822 | Yes |
| 1276 | E2F2 | ENSG00000007968 | 3.325183443 | 7.14117544 | 9.511582407 | 0.003532075 | 0.04054822 | Yes |
| 1277 | KDELR2 | ENSG00000136240 | 7.413194585 | 2.50224525 | 9.544400251 | 0.003536547 | 0.040567746 | Yes |
| 1278 | CBARP | ENSG00000099625 | 3.208853094 | 5.318166791 | 9.460111223 | 0.003544475 | 0.040607421 | Yes |
| 1279 | FBN2 | ENSG00000138829 | 4.365737942 | 8.26126909 | 9.602353328 | 0.003545555 | 0.040607421 | Yes |
| 1280 | RPA3 | ENSG00000106399 | 4.835523346 | 3.626664507 | 9.392580136 | 0.003551213 | 0.040640421 | Yes |
| 1281 | PROSER1 | ENSG00000120685 | 4.031052468 | 3.737527663 | 9.371517307 | 0.003555749 | 0.040654581 | Yes |
| 1282 | GEMIN2 | ENSG00000092208 | 2.844122475 | 6.576602355 | 9.42212205 | 0.003558005 | 0.040654581 | Yes |
| 1283 | RGS16 | ENSG00000143333 | 4.261868758 | -2.338244072 | 9.375894028 | 0.00356528 | 0.040705935 | Yes |
| 1284 | UQCR11 | ENSG00000127540 | 8.332354761 | -1.700694599 | 9.543959094 | 0.00357116 | 0.040741281 | Yes |
| 1285 | ENSG00000254615 | ENSG00000254615 | 4.511237008 | 3.946757692 | 9.373486918 | 0.003574846 | 0.040751579 | Yes |
| 1286 | BRIP1 | ENSG00000136492 | 3.658178768 | 4.519932376 | 9.409919803 | 0.003582285 | 0.040797963 | Yes |
| 1287 | CPNE8 | ENSG00000139117 | 3.493269697 | 4.191970405 | 9.381384745 | 0.00358707 | 0.040797963 | Yes |
| 1288 | DDX20 | ENSG00000064703 | 3.652212028 | 3.564891979 | 9.335528673 | 0.003591344 | 0.040797963 | Yes |
| 1289 | ST6GALNAC3 | ENSG00000184005 | 2.910771011 | 6.645464214 | 9.411097487 | 0.003591588 | 0.040797963 | Yes |
| 1290 | ZNF571 | ENSG00000180479 | 2.349893426 | 5.953706296 | 9.304487162 | 0.003592852 | 0.040797963 | Yes |
| 1291 | TPT1 | ENSG00000133112 | 11.65425104 | -1.60698952 | 9.532461915 | 0.003605444 | 0.040909214 | Yes |
| 1292 | DNMT3B | ENSG00000088305 | 2.918379382 | 6.611543488 | 9.392197911 | 0.003614541 | 0.040965877 | Yes |
| 1293 | PTH2R | ENSG00000144407 | 3.719834616 | 7.542129616 | 9.500867499 | 0.003616036 | 0.040965877 | Yes |
| 1294 | ZNF551 | ENSG00000204519 | 3.469238617 | 4.627113677 | 9.391543461 | 0.003624978 | 0.041035426 | Yes |
| 1295 | ZNF747.DT | | 2.519798011 | 6.175350059 | 9.315442017 | 0.003633735 | 0.041102769 | Yes |
| 1296 | ACAD10 | ENSG00000111271 | 5.898155827 | 2.518453947 | 9.423274952 | 0.003641516 | 0.041150692 | Yes |
| 1297 | YPEL2 | ENSG00000175155 | 5.829322971 | -1.540264677 | 9.455569544 | 0.003643595 | 0.041150692 | Yes |
| 1298 | GALNT12 | ENSG00000119514 | 4.812754401 | 2.926164236 | 9.340273866 | 0.003649418 | 0.041184679 | Yes |
| 1299 | CKAP4 | ENSG00000136026 | 8.275899354 | 1.766464917 | 9.489205072 | 0.003653316 | 0.041196912 | Yes |
| 1300 | SPATA18 | ENSG00000163071 | 4.165988312 | 8.060751506 | 9.513996178 | 0.003663866 | 0.041259575 | Yes |
| 1301 | SOX6 | ENSG00000110693 | 5.222883414 | -2.421680358 | 9.394066054 | 0.003664511 | 0.041259575 | Yes |
| 1302 | ARHGAP18 | ENSG00000146376 | 6.278350048 | 2.459897278 | 9.42048638 | 0.003679546 | 0.041397011 | Yes |
| 1303 | COL1A1 | ENSG00000108821 | 13.56660174 | 3.458056301 | 9.484681787 | 0.003682624 | 0.041399827 | Yes |
| 1304 | ARHGEF26 | ENSG00000114790 | 3.773282256 | 4.48509893 | 9.34101442 | 0.003691706 | 0.041470067 | Yes |
| 1305 | SLC41A2 | ENSG00000136052 | 2.678534158 | 6.336769306 | 9.295534358 | 0.003708943 | 0.041631752 | Yes |
| 1306 | LINC02983 | ENSG00000234432 | 2.496231318 | -2.60215278 | 9.010815526 | 0.00371861 | 0.041708276 | Yes |
| 1307 | ALDH16A1 | ENSG00000161618 | 5.873051453 | 3.040136827 | 9.352146735 | 0.003726009 | 0.041734092 | Yes |
| 1308 | POC5 | ENSG00000152359 | 2.29206497 | 5.896330642 | 9.214855331 | 0.003729024 | 0.041734092 | Yes |
| 1309 | ATL2 | ENSG00000119787 | 4.918315505 | 2.822374246 | 9.299564802 | 0.003729466 | 0.041734092 | Yes |
| 1310 | ZFAND4 | ENSG00000172671 | 3.195999264 | 4.342538922 | 9.288927542 | 0.003746945 | 0.041897653 | Yes |
| 1311 | BAZ1A.AS1 | | 2.847479372 | 6.557482882 | 9.299018276 | 0.003757743 | 0.041979541 | Yes |
| 1312 | RHBDL1 | ENSG00000103269 | 3.341771799 | 4.619776119 | 9.303608929 | 0.00376076 | 0.041979541 | Yes |
| 1313 | KCTD15 | ENSG00000153885 | 4.251964741 | 3.663837442 | 9.24844973 | 0.003765168 | 0.041979541 | Yes |
| 1314 | FILIP1L | ENSG00000168386 | 6.188806399 | -1.887665338 | 9.389760631 | 0.00376574 | 0.041979541 | Yes |
| 1315 | MCPH1.DT | | 3.283387372 | 5.326307449 | 9.33063729 | 0.003771062 | 0.041994774 | Yes |
| 1316 | EYA2 | ENSG00000064655 | 5.332107317 | 6.163460091 | 9.369355438 | 0.003773125 | 0.041994774 | Yes |
| 1317 | SRP9 | ENSG00000143742 | 4.586866571 | 3.288971637 | 9.249846954 | 0.003776728 | 0.041994774 | Yes |
| 1318 | LMCD1 | ENSG00000071282 | 6.301228425 | -1.743190949 | 9.386919074 | 0.003778583 | 0.041994774 | Yes |
| 1319 | TPM4 | ENSG00000167460 | 9.180413208 | 2.958610694 | 9.407584056 | 0.003790239 | 0.042092362 | Yes |
| 1320 | ZBTB25 | ENSG00000089775 | 4.671997586 | 2.747149916 | 9.239834234 | 0.003810069 | 0.0422805 | Yes |
| 1321 | GADD45G | ENSG00000130222 | 5.664779959 | -2.330031366 | 9.324507171 | 0.003829295 | 0.042461659 | Yes |
| 1322 | TRUB1 | ENSG00000165832 | 4.315909686 | 3.373254686 | 9.201717217 | 0.003841815 | 0.042532969 | Yes |
| 1323 | LBX2.AS1 | | 2.924928801 | 6.667438214 | 9.265202136 | 0.003842305 | 0.042532969 | Yes |
| 1324 | SRP14.DT | | 2.954524026 | 6.655435789 | 9.262290357 | 0.003844443 | 0.042532969 | Yes |
| 1325 | CHD3 | ENSG00000170004 | 7.698868225 | 1.967387285 | 9.362501279 | 0.003851901 | 0.042583289 | Yes |
| 1326 | CDCP1 | ENSG00000163814 | 4.749015406 | 8.670727477 | 9.43395857 | 0.003855721 | 0.042590278 | Yes |
| 1327 | TRO | ENSG00000067445 | 6.758446532 | 3.288674911 | 9.310688719 | 0.003858353 | 0.042590278 | Yes |
| 1328 | TRAF1 | ENSG00000056558 | 3.577086165 | 4.833420892 | 9.26014762 | 0.003876291 | 0.042741698 | Yes |
| 1329 | ZNF736 | ENSG00000234444 | 3.375872539 | 3.940218139 | 9.190647908 | 0.00387791 | 0.042741698 | Yes |
| 1330 | B4GALT1 | ENSG00000086062 | 7.531263479 | 2.127118918 | 9.339633337 | 0.003884807 | 0.042785488 | Yes |
| 1331 | COPS7A | ENSG00000111652 | 5.174765096 | 2.675108206 | 9.221709093 | 0.003902377 | 0.04294669 | Yes |
| 1332 | ZBTB26 | ENSG00000171448 | 2.50277861 | 6.111473529 | 9.14754815 | 0.003907702 | 0.042954571 | Yes |
| 1333 | DMKN | ENSG00000161249 | 4.78193151 | 8.692625298 | 9.401505904 | 0.003912794 | 0.042954571 | Yes |
| 1334 | ABHD13 | ENSG00000139826 | 3.560284034 | 3.838694681 | 9.162938902 | 0.003915963 | 0.042954571 | Yes |
| 1335 | IP6K2 | ENSG00000068745 | 6.593452903 | 2.144672142 | 9.298077626 | 0.003919196 | 0.042954571 | Yes |
| 1336 | ENSG00000272733 | ENSG00000272733 | 3.527572957 | 5.544255658 | 9.267544539 | 0.003919233 | 0.042954571 | Yes |
| 1337 | CRY1 | ENSG00000008405 | 4.18290288 | 3.431261284 | 9.154347734 | 0.003920701 | 0.042954571 | Yes |
| 1338 | IL20RB | ENSG00000174564 | 4.578054332 | 8.497924957 | 9.382192549 | 0.003929138 | 0.043014806 | Yes |
| 1339 | POLD4 | ENSG00000175482 | 5.401790843 | -1.708497136 | 9.257733147 | 0.003944858 | 0.04311247 | Yes |
| 1340 | EVI2A | ENSG00000126860 | 3.989057093 | 7.866035642 | 9.33165596 | 0.003945948 | 0.04311247 | Yes |
| 1341 | CNIH2 | ENSG00000174871 | 2.868661896 | 6.56547032 | 9.192013504 | 0.003947266 | 0.04311247 | Yes |
| 1342 | RAB35.AS1 | | 3.20383908 | 6.986076226 | 9.244310128 | 0.003949841 | 0.04311247 | Yes |
| 1343 | TMEM37 | ENSG00000171227 | 4.9854772 | -2.19830849 | 9.21740501 | 0.003956565 | 0.043153687 | Yes |
| 1344 | CNN3 | ENSG00000117519 | 7.32990258 | 2.546147759 | 9.282957562 | 0.003968842 | 0.043223855 | Yes |
| 1345 | ELAVL1 | ENSG00000066044 | 6.63530127 | 2.301106517 | 9.26790115 | 0.003969686 | 0.043223855 | Yes |
| 1346 | CELSR3 | ENSG00000008300 | 3.128489958 | 4.417465538 | 9.162833228 | 0.003971858 | 0.043223855 | Yes |
| 1347 | CALHM6 | ENSG00000188820 | 3.736739978 | 7.593516033 | 9.292271589 | 0.003975627 | 0.043232724 | Yes |
| 1348 | ENSG00000215790 | ENSG00000215790 | 3.367101297 | 5.380794691 | 9.21650006 | 0.003980938 | 0.043258349 | Yes |
| 1349 | FAM234B | ENSG00000084444 | 3.156568313 | 5.107578062 | 9.178986897 | 0.003998807 | 0.043420278 | Yes |
| 1350 | ZNF12 | ENSG00000164631 | 4.511282851 | 4.02170672 | 9.128994292 | 0.004002181 | 0.043420855 | Yes |
| 1351 | AFMID | ENSG00000183077 | 5.176906316 | 2.862740575 | 9.156659278 | 0.004007465 | 0.043420855 | Yes |
| 1352 | MAPRE1 | ENSG00000101367 | 4.126669346 | 3.748336527 | 9.111700564 | 0.0040098 | 0.043420855 | Yes |
| 1353 | ZNF668 | ENSG00000167394 | 4.478789623 | -2.13353934 | 9.148079267 | 0.004012628 | 0.043420855 | Yes |
| 1354 | DEPTOR | ENSG00000155792 | 3.826527426 | -2.48040937 | 9.055521195 | 0.004013693 | 0.043420855 | Yes |
| 1355 | DPF3 | ENSG00000205683 | 3.361089387 | -2.727637261 | 8.961241675 | 0.004024129 | 0.043501605 | Yes |
| 1356 | GABBR2 | ENSG00000136928 | 2.518548333 | -2.96354703 | 8.824900445 | 0.004027598 | 0.043506971 | Yes |
| 1357 | DAP | ENSG00000112977 | 6.019315919 | 2.823829555 | 9.193171747 | 0.004033358 | 0.043537062 | Yes |
| 1358 | ENSG00000273338 | ENSG00000273338 | 4.475885727 | 5.80751662 | 9.233763447 | 0.004037417 | 0.043547805 | Yes |
| 1359 | SLC19A2 | ENSG00000117479 | 2.649031284 | 6.336963422 | 9.108892843 | 0.004040303 | 0.043547805 | Yes |
| 1360 | SLC15A2 | ENSG00000163406 | 4.18810973 | 6.325044771 | 9.259055826 | 0.004045131 | 0.043567761 | Yes |
| 1361 | MINDY2 | ENSG00000128923 | 5.416336013 | 2.913694227 | 9.145563808 | 0.004050692 | 0.043574213 | Yes |
| 1362 | EED | ENSG00000074266 | 4.197908627 | 4.130778823 | 9.10576654 | 0.004051684 | 0.043574213 | Yes |
| 1363 | STMN3 | ENSG00000197457 | 5.163251114 | 3.06167653 | 9.121838845 | 0.004060551 | 0.043630984 | Yes |
| 1364 | TNFRSF21 | ENSG00000146072 | 6.510581433 | 3.339900073 | 9.183066676 | 0.004062925 | 0.043630984 | Yes |
| 1365 | NMRAL1 | ENSG00000153406 | 4.439483569 | 3.049178538 | 9.079293734 | 0.004071449 | 0.043690472 | Yes |
| 1366 | AOX1 | ENSG00000138356 | 4.215396524 | 5.502405339 | 9.196666756 | 0.004075196 | 0.043694608 | Yes |
| 1367 | ZNF780A | ENSG00000197782 | 3.081800402 | 4.27326804 | 9.095082748 | 0.004077805 | 0.043694608 | Yes |
| 1368 | COPA | ENSG00000122218 | 7.846880989 | 2.000451157 | 9.232304792 | 0.004084829 | 0.043737849 | Yes |
| 1369 | SNX7 | ENSG00000162627 | 5.142332511 | 3.593685658 | 9.092137829 | 0.004095346 | 0.043818405 | Yes |
| 1370 | ZNF185 | ENSG00000147394 | 3.330628584 | 7.154919534 | 9.179485519 | 0.004101932 | 0.043847435 | Yes |
| 1371 | GRIN2D | ENSG00000105464 | 4.282687573 | 8.181670038 | 9.265022178 | 0.00410405 | 0.043847435 | Yes |
| 1372 | ATG16L2 | ENSG00000168010 | 6.015980533 | 2.453304266 | 9.162594042 | 0.004110707 | 0.043886515 | Yes |
| 1373 | CDC25C | ENSG00000158402 | 2.685013408 | 6.379218409 | 9.070769716 | 0.004123121 | 0.043986972 | Yes |
| 1374 | CENPH | ENSG00000153044 | 3.561833244 | 5.637880075 | 9.158881744 | 0.004132276 | 0.044043633 | Yes |
| 1375 | PNMA8A | ENSG00000182013 | 3.156470828 | 6.903442839 | 9.132858288 | 0.004135536 | 0.044043633 | Yes |
| 1376 | SLC12A7 | ENSG00000113504 | 5.660771782 | 2.518711537 | 9.127730899 | 0.00413746 | 0.044043633 | Yes |
| 1377 | CBX4 | ENSG00000141582 | 6.768439471 | 2.430834767 | 9.163751165 | 0.004162639 | 0.044279471 | Yes |
| 1378 | KLHL34 | ENSG00000185915 | 3.609280688 | -3.572944591 | 8.837170984 | 0.004178065 | 0.044345709 | Yes |
| 1379 | TOMM7 | ENSG00000196683 | 8.713028751 | -1.745389971 | 9.191652382 | 0.004179048 | 0.044345709 | Yes |
| 1380 | PSMB9 | ENSG00000240065 | 6.006944905 | 3.221189153 | 9.099060061 | 0.004179204 | 0.044345709 | Yes |
| 1381 | LRP1 | ENSG00000123384 | 8.820940039 | 2.261186687 | 9.187661164 | 0.004180985 | 0.044345709 | Yes |
| 1382 | ZNF557 | ENSG00000130544 | 4.14256603 | 3.073826321 | 9.005528514 | 0.004188329 | 0.044374027 | Yes |
| 1383 | TBX18 | ENSG00000112837 | 3.410598616 | 7.237560748 | 9.141120592 | 0.004189718 | 0.044374027 | Yes |
| 1384 | TROAP | ENSG00000135451 | 4.671177401 | 3.46485999 | 9.021893517 | 0.00419851 | 0.044410952 | Yes |
| 1385 | SLC16A10 | ENSG00000112394 | 3.788071885 | -2.273926437 | 8.962570606 | 0.004200745 | 0.044410952 | Yes |
| 1386 | EMB | ENSG00000170571 | 3.305601039 | 5.403444253 | 9.095793814 | 0.004202307 | 0.044410952 | Yes |
| 1387 | RNF32.DT | | 2.648830897 | 6.266232929 | 9.003897359 | 0.004220232 | 0.04453801 | Yes |
| 1388 | RFLNB | ENSG00000183688 | 7.70967193 | 2.729751999 | 9.148860016 | 0.004223102 | 0.04453801 | Yes |
| 1389 | GNG12 | ENSG00000172380 | 5.849567649 | 2.353805203 | 9.095650745 | 0.004227382 | 0.04453801 | Yes |
| 1390 | UBAC1 | ENSG00000130560 | 6.175121834 | -1.98691919 | 9.129559938 | 0.004228376 | 0.04453801 | Yes |
| 1391 | ENSG00000258472 | ENSG00000258472 | 3.692076028 | 4.153183316 | 9.016014556 | 0.004231042 | 0.04453801 | Yes |
| 1392 | EID2B | ENSG00000176401 | 3.344213412 | 4.496222185 | 9.038769718 | 0.004232587 | 0.04453801 | Yes |
| 1393 | SMCR8 | ENSG00000176994 | 4.345404752 | 3.18996201 | 8.985074566 | 0.004244699 | 0.044633373 | Yes |
| 1394 | IGIP | ENSG00000182700 | 3.815840989 | 5.045459702 | 9.074238928 | 0.00424913 | 0.044647893 | Yes |
| 1395 | UTP3 | ENSG00000132467 | 5.172062662 | 2.905359642 | 9.024765312 | 0.004253273 | 0.044654467 | Yes |
| 1396 | DZIP1 | ENSG00000134874 | 5.326285441 | 3.070110726 | 9.027029007 | 0.004257606 | 0.044654467 | Yes |
| 1397 | SH3YL1 | ENSG00000035115 | 5.257299017 | 2.481582838 | 9.040883891 | 0.004258908 | 0.044654467 | Yes |
| 1398 | FIRRM | ENSG00000000460 | 3.433855791 | 4.127172382 | 8.998638132 | 0.004263324 | 0.04466519 | Yes |
| 1399 | PROCR | ENSG00000101000 | 5.453240245 | 4.459729958 | 9.007802292 | 0.004266034 | 0.04466519 | Yes |
| 1400 | KHDC1 | ENSG00000135314 | 2.710638233 | 6.410038442 | 8.997237938 | 0.004273262 | 0.044690185 | Yes |
| 1401 | PTPN13 | ENSG00000163629 | 3.741966099 | 3.999926123 | 8.984005784 | 0.004274527 | 0.044690185 | Yes |
| 1402 | ZNF845 | ENSG00000213799 | 2.464152203 | 6.109713284 | 8.951591357 | 0.004278314 | 0.044697846 | Yes |
| 1403 | AQP5 | ENSG00000161798 | 5.344735045 | 9.287695179 | 9.214569483 | 0.004290805 | 0.044748412 | Yes |
| 1404 | DUSP15 | ENSG00000149599 | 2.634253215 | 6.267354803 | 8.967914245 | 0.004291287 | 0.044748412 | Yes |
| 1405 | SCAMP4 | ENSG00000227500 | 6.933610208 | 1.724049944 | 9.108838436 | 0.004294004 | 0.044748412 | Yes |
| 1406 | NUP107 | ENSG00000111581 | 7.641242443 | 3.632175452 | 9.093546629 | 0.004295383 | 0.044748412 | Yes |
| 1407 | RRAGD | ENSG00000025039 | 6.14940531 | -1.822950864 | 9.083628411 | 0.00431836 | 0.044905594 | Yes |
| 1408 | USP35 | ENSG00000118369 | 2.445246095 | 6.031725251 | 8.919604828 | 0.004318859 | 0.044905594 | Yes |
| 1409 | SNCAIP | ENSG00000064692 | 3.671624912 | 4.904978639 | 9.026177385 | 0.004322001 | 0.044905594 | Yes |
| 1410 | MTERF3 | ENSG00000156469 | 3.526332027 | 5.517463037 | 9.049814682 | 0.004322742 | 0.044905594 | Yes |
| 1411 | PFN1 | ENSG00000108518 | 7.576920586 | 2.180536786 | 9.099095674 | 0.00432588 | 0.044906315 | Yes |
| 1412 | HDLBP | ENSG00000115677 | 8.122845912 | -1.371684153 | 9.109877991 | 0.004329752 | 0.044914656 | Yes |
| 1413 | C1orf21 | ENSG00000116667 | 5.831397441 | -1.919202682 | 9.060165775 | 0.004341992 | 0.044998808 | Yes |
| 1414 | FAM111A | ENSG00000166801 | 6.062103592 | 2.419180388 | 9.043199359 | 0.004344622 | 0.044998808 | Yes |
| 1415 | ZNF582.DT | | 2.682955301 | 4.606221547 | 8.931993309 | 0.004347087 | 0.044998808 | Yes |
| 1416 | ZNF792 | ENSG00000180884 | 2.551633173 | 6.229624953 | 8.932986669 | 0.004350414 | 0.045001417 | Yes |
| 1417 | FBLN7 | ENSG00000144152 | 2.990219681 | 6.724879885 | 8.99365961 | 0.00436246 | 0.045094161 | Yes |
| 1418 | MYH15 | ENSG00000144821 | 3.728536706 | 7.529309319 | 9.075053492 | 0.00437001 | 0.045138151 | Yes |
| 1419 | SGTA | ENSG00000104969 | 6.256943019 | -1.88710928 | 9.058093927 | 0.004372884 | 0.045138151 | Yes |
| 1420 | NETO2 | ENSG00000171208 | 3.601771407 | 4.867001917 | 8.994468549 | 0.004378958 | 0.045146281 | Yes |
| 1421 | DPY19L1 | ENSG00000173852 | 4.717797155 | 2.949953321 | 8.936173289 | 0.00437984 | 0.045146281 | Yes |
| 1422 | ITGAL | ENSG00000005844 | 3.105957967 | 6.817103447 | 8.988881088 | 0.004394412 | 0.045234626 | Yes |
| 1423 | PHLPP2 | ENSG00000040199 | 2.586661307 | 6.246791374 | 8.913174992 | 0.004395379 | 0.045234626 | Yes |
| 1424 | SOCS1 | ENSG00000185338 | 3.4549283 | 7.263447146 | 9.034431709 | 0.004401188 | 0.045234626 | Yes |
| 1425 | AMH | ENSG00000104899 | 4.330108567 | 4.319900165 | 8.932221786 | 0.00440364 | 0.045234626 | Yes |
| 1426 | NDC80 | ENSG00000080986 | 3.000642675 | 6.771846462 | 8.978785174 | 0.004403863 | 0.045234626 | Yes |
| 1427 | FGD6 | ENSG00000180263 | 3.285940001 | 7.092315125 | 9.011533359 | 0.004411608 | 0.045258161 | Yes |
| 1428 | ATP1B4 | ENSG00000101892 | 2.590289766 | -3.149293179 | 8.637226604 | 0.004412338 | 0.045258161 | Yes |
| 1429 | ERP44 | ENSG00000023318 | 6.065592226 | 2.382466596 | 9.007272552 | 0.004418252 | 0.045287085 | Yes |
| 1430 | PSRC1 | ENSG00000134222 | 3.179236186 | 6.978331314 | 8.990608396 | 0.004428587 | 0.045361247 | Yes |
| 1431 | DVL1 | ENSG00000107404 | 7.081242208 | -1.609302491 | 9.045727543 | 0.004434002 | 0.045384956 | Yes |
| 1432 | SPART | ENSG00000133104 | 6.02330747 | 2.545655718 | 8.990772693 | 0.00443816 | 0.04539577 | Yes |
| 1433 | LRRC59 | ENSG00000108829 | 7.255317066 | 2.225843408 | 9.032023819 | 0.004446599 | 0.045450323 | Yes |
| 1434 | CDKN2B | ENSG00000147883 | 4.913528329 | 5.233603222 | 8.945571141 | 0.00445617 | 0.045516371 | Yes |
| 1435 | CBX5 | ENSG00000094916 | 7.169108355 | 1.407424803 | 9.020453261 | 0.004481883 | 0.045718161 | Yes |
| 1436 | LGR4 | ENSG00000205213 | 3.773483805 | 4.481338334 | 8.915123624 | 0.004483169 | 0.045718161 | Yes |
| 1437 | CCDC40 | ENSG00000141519 | 3.252873119 | 5.302306627 | 8.945712658 | 0.004485296 | 0.045718161 | Yes |
| 1438 | YWHAG | ENSG00000170027 | 7.252345206 | -1.587012667 | 9.015659866 | 0.004498966 | 0.045825583 | Yes |
| 1439 | PDIA3 | ENSG00000167004 | 7.715819985 | 2.350661133 | 9.009457326 | 0.004504259 | 0.045847596 | Yes |
| 1440 | DENND6B | ENSG00000205593 | 4.858789616 | -2.22489236 | 8.916329141 | 0.004517966 | 0.045955151 | Yes |
| 1441 | ANKRD42 | ENSG00000137494 | 3.265807395 | 5.279911999 | 8.926964005 | 0.004523372 | 0.045973516 | Yes |
| 1442 | SNRPD1 | ENSG00000167088 | 7.155230585 | 2.143197578 | 8.991688427 | 0.004526053 | 0.045973516 | Yes |
| 1443 | CGAS | ENSG00000164430 | 2.817525503 | 6.51987421 | 8.884112849 | 0.004532145 | 0.045994363 | Yes |
| 1444 | COPG1 | ENSG00000181789 | 6.545388829 | 1.632689273 | 8.979808613 | 0.00453439 | 0.045994363 | Yes |
| 1445 | RAC2 | ENSG00000128340 | 4.781293611 | 3.592248534 | 8.856741349 | 0.004538531 | 0.04599958 | Yes |
| 1446 | DDHD1 | ENSG00000100523 | 4.975642863 | 2.75064385 | 8.876002547 | 0.00454119 | 0.04599958 | Yes |
| 1447 | ENSG00000270127 | ENSG00000270127 | 3.440717962 | 7.216355926 | 8.958392796 | 0.004545311 | 0.046009483 | Yes |
| 1448 | SALL4 | ENSG00000101115 | 2.922510177 | 6.65462578 | 8.889110776 | 0.004557845 | 0.046104477 | Yes |
| 1449 | ABL1 | ENSG00000097007 | 7.471696018 | 2.195762382 | 8.979237587 | 0.004561595 | 0.04611054 | Yes |
| 1450 | BAMBI | ENSG00000095739 | 2.355321964 | 5.914317151 | 8.781447852 | 0.004569273 | 0.046111566 | Yes |
| 1451 | ZBTB21 | ENSG00000173276 | 3.51946826 | 4.114737972 | 8.844900685 | 0.004576791 | 0.046111566 | Yes |
| 1452 | PSMB8.AS1 | | 4.142416431 | 3.898651406 | 8.830294678 | 0.004577139 | 0.046111566 | Yes |
| 1453 | EFNA3 | ENSG00000143590 | 2.759299384 | 6.45817134 | 8.852364568 | 0.004581698 | 0.046111566 | Yes |
| 1454 | BST1 | ENSG00000109743 | 2.763100296 | 6.481121637 | 8.855164504 | 0.004582249 | 0.046111566 | Yes |
| 1455 | MILR1 | ENSG00000271605 | 3.400578028 | 4.636223609 | 8.875866936 | 0.004582558 | 0.046111566 | Yes |
| 1456 | TEDC2 | ENSG00000162062 | 2.67821937 | 6.328695336 | 8.833918569 | 0.004583749 | 0.046111566 | Yes |
| 1457 | WDR4 | ENSG00000160193 | 4.952860188 | 3.127484406 | 8.840038438 | 0.004595589 | 0.046168319 | Yes |
| 1458 | ENSG00000271971 | ENSG00000271971 | 3.026858682 | 4.994328858 | 8.859338518 | 0.004595699 | 0.046168319 | Yes |
| 1459 | ERI2 | ENSG00000196678 | 3.205001816 | 4.37978327 | 8.844540237 | 0.004601683 | 0.046196732 | Yes |
| 1460 | TNFRSF10D | ENSG00000173530 | 2.899170965 | 4.886284364 | 8.840890783 | 0.004608207 | 0.046230516 | Yes |
| 1461 | PRKAG2 | ENSG00000106617 | 3.755996504 | 4.201685964 | 8.829119937 | 0.004618335 | 0.046300392 | Yes |
| 1462 | ESM1 | ENSG00000164283 | 4.953747203 | 3.612299657 | 8.813861178 | 0.004640999 | 0.046446354 | Yes |
| 1463 | TMED3 | ENSG00000166557 | 5.858038566 | 2.374921772 | 8.889358496 | 0.004641746 | 0.046446354 | Yes |
| 1464 | NAT14 | ENSG00000090971 | 5.038010116 | 3.242879713 | 8.819175204 | 0.004644673 | 0.046446354 | Yes |
| 1465 | CACNA1A | ENSG00000141837 | 5.228049094 | 2.521057253 | 8.847760263 | 0.004645587 | 0.046446354 | Yes |
| 1466 | EPSTI1 | ENSG00000133106 | 4.38066828 | 4.74702914 | 8.840514266 | 0.004648965 | 0.046448401 | Yes |
| 1467 | FZD2 | ENSG00000180340 | 3.567348813 | 4.73389127 | 8.850025793 | 0.004658222 | 0.046509138 | Yes |
| 1468 | TP53 | ENSG00000141510 | 5.743369475 | 2.619385006 | 8.866792853 | 0.0046614 | 0.046509139 | Yes |
| 1469 | RND3 | ENSG00000115963 | 5.589995226 | 3.038138783 | 8.841237684 | 0.004667479 | 0.046538068 | Yes |
| 1470 | TUBA8 | ENSG00000183785 | 3.704075316 | -2.839337968 | 8.688967448 | 0.004674426 | 0.046575609 | Yes |
| 1471 | UST | ENSG00000111962 | 3.936179341 | 3.920420641 | 8.782074099 | 0.004681615 | 0.046615427 | Yes |
| 1472 | BRI3BP | ENSG00000184992 | 4.751958463 | 3.145654289 | 8.789066356 | 0.004684792 | 0.046615427 | Yes |
| 1473 | RFX2 | ENSG00000087903 | 5.12464596 | 2.969337106 | 8.807992866 | 0.004689869 | 0.046629332 | Yes |
| 1474 | FAM24B | ENSG00000213185 | 2.70387822 | 6.402695564 | 8.793244725 | 0.00469256 | 0.046629332 | Yes |
| 1475 | NUP160 | ENSG00000030066 | 5.432035809 | 2.173685433 | 8.847119513 | 0.004696872 | 0.046640509 | Yes |
| 1476 | IQCC | ENSG00000160051 | 2.724607354 | 6.395715308 | 8.785832398 | 0.004706563 | 0.046680455 | Yes |
| 1477 | ZNF793.AS1 | | 3.26893581 | 7.069486428 | 8.866158829 | 0.004707273 | 0.046680455 | Yes |
| 1478 | ASAH2B | ENSG00000204147 | 2.30600236 | 5.869163613 | 8.707809425 | 0.004715335 | 0.046726659 | Yes |
| 1479 | PDHX | ENSG00000110435 | 5.385675999 | -1.931809222 | 8.856703933 | 0.004719924 | 0.046726659 | Yes |
| 1480 | MRPS14 | ENSG00000120333 | 4.392837913 | 3.894744107 | 8.766161593 | 0.004721509 | 0.046726659 | Yes |
| 1481 | SFTA1P | ENSG00000225383 | 2.969632153 | 6.725973687 | 8.812281959 | 0.004740298 | 0.046855762 | Yes |
| 1482 | LOC124901151 | ENSG00000248367 | 2.973065186 | 6.685002352 | 8.806431856 | 0.00474216 | 0.046855762 | Yes |
| 1483 | PLXNC1 | ENSG00000136040 | 5.034176467 | 4.820605469 | 8.782841554 | 0.004746824 | 0.046855762 | Yes |
| 1484 | GNAI3 | ENSG00000065135 | 6.880014881 | 2.248171472 | 8.878303662 | 0.004747359 | 0.046855762 | Yes |
| 1485 | PERP | ENSG00000112378 | 6.539237969 | 10.50257945 | 9.007787624 | 0.004751422 | 0.046864262 | Yes |
| 1486 | ENSG00000224032 | ENSG00000224032 | 5.620346063 | 2.875320123 | 8.804417346 | 0.004763052 | 0.046947336 | Yes |
| 1487 | ADGRB2 | ENSG00000121753 | 3.139371519 | 6.901968266 | 8.814782622 | 0.004779152 | 0.047074324 | Yes |
| 1488 | NVL | ENSG00000143748 | 4.48889408 | 2.966992909 | 8.731216794 | 0.004788366 | 0.047133365 | Yes |
| 1489 | SPINDOC | ENSG00000168005 | 3.39150756 | 4.591872623 | 8.774759691 | 0.004794223 | 0.047159304 | Yes |
| 1490 | QSOX2 | ENSG00000165661 | 3.141099887 | 4.341848783 | 8.742307147 | 0.004813047 | 0.047215547 | Yes |
| 1491 | TUB | ENSG00000166402 | 3.309887775 | 7.097137911 | 8.82006678 | 0.004813286 | 0.047215547 | Yes |
| 1492 | ZNF614 | ENSG00000142556 | 2.48567097 | 6.1229858 | 8.700115153 | 0.00481363 | 0.047215547 | Yes |
| 1493 | ABHD4 | ENSG00000100439 | 4.88385318 | 3.312556682 | 8.73338994 | 0.00481573 | 0.047215547 | Yes |
| 1494 | FOXN2 | ENSG00000170802 | 2.853766574 | 4.784349457 | 8.733202818 | 0.00481607 | 0.047215547 | Yes |
| 1495 | NEMP2 | ENSG00000189362 | 2.661791328 | 6.339532247 | 8.723077237 | 0.004828686 | 0.04726925 | Yes |
| 1496 | UBP1 | ENSG00000153560 | 6.21428974 | 1.84257598 | 8.827878456 | 0.004829976 | 0.04726925 | Yes |
| 1497 | ENSG00000116883 | ENSG00000116883 | 3.035092047 | 6.779325557 | 8.777130132 | 0.004831236 | 0.04726925 | Yes |
| 1498 | CMIP | ENSG00000153815 | 7.695731201 | 1.664959814 | 8.854539261 | 0.00484005 | 0.047278169 | Yes |
| 1499 | KIF18B | ENSG00000186185 | 3.453122496 | 7.2867938 | 8.825855505 | 0.004841594 | 0.047278169 | Yes |
| 1500 | PTGER4 | ENSG00000171522 | 4.822751551 | 3.420370003 | 8.718955299 | 0.004841838 | 0.047278169 | Yes |
| 1501 | SKA1 | ENSG00000154839 | 3.369142387 | 5.410300085 | 8.783557435 | 0.004853315 | 0.047358647 | Yes |
| 1502 | LRIF1 | ENSG00000121931 | 3.709354954 | 4.96477021 | 8.774178949 | 0.004860007 | 0.04739235 | Yes |
| 1503 | VILL | ENSG00000136059 | 3.356686858 | 7.188864993 | 8.802752644 | 0.004871809 | 0.047475808 | Yes |
| 1504 | CYB5RL | ENSG00000215883 | 2.551511965 | 6.177752471 | 8.679292814 | 0.004877746 | 0.047498637 | Yes |
| 1505 | GLI2 | ENSG00000074047 | 4.016604604 | 6.134379133 | 8.826412369 | 0.00488485 | 0.047498637 | Yes |
| 1506 | NDUFA5 | ENSG00000128609 | 7.177547361 | -1.822380953 | 8.830337543 | 0.004886218 | 0.047498637 | Yes |
| 1507 | ARHGEF1 | ENSG00000076928 | 6.907454845 | 2.543392749 | 8.810118443 | 0.004887132 | 0.047498637 | Yes |
| 1508 | SLC16A5 | ENSG00000170190 | 2.270476847 | 5.86945116 | 8.628832867 | 0.004894094 | 0.047534743 | Yes |
| 1509 | PTGES | ENSG00000148344 | 7.880696904 | 4.38073246 | 8.789582964 | 0.004900807 | 0.047562198 | Yes |
| 1510 | INTS14 | ENSG00000138614 | 2.637421527 | 6.34391077 | 8.690131106 | 0.004904541 | 0.047562198 | Yes |
| 1511 | KLHL5 | ENSG00000109790 | 4.881806428 | 3.402833827 | 8.690822777 | 0.004910082 | 0.047562198 | Yes |
| 1512 | MT.ND4 | | 13.5398367 | -1.24428122 | 8.839048342 | 0.004911907 | 0.047562198 | Yes |
| 1513 | TTK | ENSG00000112742 | 2.73183639 | 6.433587482 | 8.698003906 | 0.004913168 | 0.047562198 | Yes |
| 1514 | PBXIP1 | ENSG00000163346 | 6.424352471 | -1.298911186 | 8.801932131 | 0.004927238 | 0.047637194 | Yes |
| 1515 | PNMA6A | ENSG00000235961 | 4.517437744 | -2.692817553 | 8.681745319 | 0.004927424 | 0.047637194 | Yes |
| 1516 | DUSP2 | ENSG00000158050 | 3.798403599 | 5.867983017 | 8.785413726 | 0.004934764 | 0.047669455 | Yes |
| 1517 | NUSAP1 | ENSG00000137804 | 5.193255344 | 3.34793133 | 8.691790071 | 0.004937275 | 0.047669455 | Yes |
| 1518 | SPATA13 | ENSG00000182957 | 2.90821803 | 6.644110069 | 8.710695538 | 0.004944388 | 0.047696293 | Yes |
| 1519 | NBPF12 | ENSG00000268043 | 3.53655144 | 7.288659672 | 8.77743948 | 0.004949415 | 0.047696293 | Yes |
| 1520 | CDKN3 | ENSG00000100526 | 3.616171541 | 5.702844063 | 8.765910574 | 0.004949831 | 0.047696293 | Yes |
| 1521 | GNPNAT1 | ENSG00000100522 | 5.029043138 | 3.245147522 | 8.677734892 | 0.00495706 | 0.047734529 | Yes |
| 1522 | APH1B | ENSG00000138613 | 3.124853657 | 4.327448638 | 8.674661584 | 0.004963034 | 0.047760638 | Yes |
| 1523 | NDUFA3 | ENSG00000170906 | 6.634924263 | -1.678539965 | 8.784066867 | 0.004972024 | 0.047783031 | Yes |
| 1524 | CPNE1 | ENSG00000214078 | 7.324591157 | 1.53179195 | 8.790803757 | 0.004973604 | 0.047783031 | Yes |
| 1525 | B3GALNT1 | ENSG00000169255 | 3.702854738 | 3.875506419 | 8.647614596 | 0.004977936 | 0.047783031 | Yes |
| 1526 | NDUFB8 | ENSG00000166136 | 7.148021783 | -1.415485815 | 8.789298541 | 0.00498034 | 0.047783031 | Yes |
| 1527 | SUCO | ENSG00000094975 | 5.57861589 | 2.665090108 | 8.711062955 | 0.004982754 | 0.047783031 | Yes |
| 1528 | PDIA4 | ENSG00000155660 | 5.519321615 | 2.630152856 | 8.708350311 | 0.004984948 | 0.047783031 | Yes |
| 1529 | CTSB | ENSG00000164733 | 9.562653544 | 2.012255875 | 8.79120137 | 0.005005303 | 0.047946743 | Yes |
| 1530 | RAP1B | ENSG00000127314 | 6.565785436 | 2.114162028 | 8.752397438 | 0.005011722 | 0.047967983 | Yes |
| 1531 | HMBOX1 | ENSG00000147421 | 4.161829256 | 3.632192354 | 8.624583911 | 0.005018717 | 0.047967983 | Yes |
| 1532 | MDH1 | ENSG00000014641 | 7.826832326 | -1.596238186 | 8.77808226 | 0.005020522 | 0.047967983 | Yes |
| 1533 | SLC36A4 | ENSG00000180773 | 3.076734801 | 5.04638796 | 8.674830429 | 0.005021991 | 0.047967983 | Yes |
| 1534 | ENSG00000226067 | ENSG00000226067 | 3.912972888 | 5.148123015 | 8.713343375 | 0.005025894 | 0.047967983 | Yes |
| 1535 | DZIP1L | ENSG00000158163 | 4.46465903 | 3.615032221 | 8.625865193 | 0.005031494 | 0.047967983 | Yes |
| 1536 | HROB | ENSG00000125319 | 2.589085058 | 6.251188543 | 8.622112836 | 0.005033016 | 0.047967983 | Yes |
| 1537 | ENSG00000273650 | ENSG00000273650 | 2.834829157 | 6.533300336 | 8.658252291 | 0.005033738 | 0.047967983 | Yes |
| 1538 | RNPEP | ENSG00000176393 | 6.153315896 | 2.061966889 | 8.727537275 | 0.005041471 | 0.047993427 | Yes |
| 1539 | HIBADH | ENSG00000106049 | 4.035525739 | 4.005512327 | 8.62498359 | 0.005046193 | 0.047993427 | Yes |
| 1540 | MKI67 | ENSG00000148773 | 5.410106736 | 3.782895363 | 8.646191784 | 0.005046245 | 0.047993427 | Yes |
| 1541 | BCAR3 | ENSG00000137936 | 3.279414406 | 7.059054546 | 8.702135965 | 0.005072576 | 0.048156025 | Yes |
| 1542 | SRCAP | ENSG00000080603 | 3.014334975 | 4.960956841 | 8.639748638 | 0.005076761 | 0.048156025 | Yes |
| 1543 | LYN | ENSG00000254087 | 5.714706626 | 3.590685231 | 8.648056243 | 0.005078247 | 0.048156025 | Yes |
| 1544 | ARHGAP42 | ENSG00000165895 | 3.175549047 | 4.467581876 | 8.634004798 | 0.005078266 | 0.048156025 | Yes |
| 1545 | EPM2A.DT | | 2.514092943 | 6.127640451 | 8.584497059 | 0.005082346 | 0.048156025 | Yes |
| 1546 | SPRED1 | ENSG00000166068 | 6.585901243 | 1.962897464 | 8.724046007 | 0.005083546 | 0.048156025 | Yes |
| 1547 | LPAR5 | ENSG00000184574 | 2.40929454 | 6.045328059 | 8.570805555 | 0.005088297 | 0.048156025 | Yes |
| 1548 | SPOCD1 | ENSG00000134668 | 5.395328133 | 7.660599621 | 8.799559873 | 0.005089661 | 0.048156025 | Yes |
| 1549 | ARFIP1 | ENSG00000164144 | 4.597881491 | 2.556227837 | 8.607175694 | 0.00510166 | 0.048218513 | Yes |
| 1550 | HIF1A | ENSG00000100644 | 6.050716088 | 2.536921532 | 8.68592333 | 0.005103188 | 0.048218513 | Yes |
| 1551 | SLC45A4 | ENSG00000022567 | 4.869855453 | 2.961417039 | 8.611872309 | 0.005106148 | 0.048218513 | Yes |
| 1552 | DLX3 | ENSG00000064195 | 4.344416586 | 8.245509802 | 8.77643489 | 0.005110494 | 0.048228435 | Yes |
| 1553 | SSH3 | ENSG00000172830 | 4.226562486 | 4.245192572 | 8.604928193 | 0.005118641 | 0.048274197 | Yes |
| 1554 | ADPGK | ENSG00000159322 | 5.340959639 | 2.632183165 | 8.636159981 | 0.005129679 | 0.048316483 | Yes |
| 1555 | ITGB3BP | ENSG00000142856 | 4.477598563 | 3.667214989 | 8.585126516 | 0.005129727 | 0.048316483 | Yes |
| 1556 | FMO1 | ENSG00000010932 | 4.171529284 | 8.018163333 | 8.751406917 | 0.005137277 | 0.04833588 | Yes |
| 1557 | GPR183 | ENSG00000169508 | 3.594506285 | 7.406036571 | 8.705516069 | 0.005138391 | 0.04833588 | Yes |
| 1558 | HLA.DQA1.1 | | 3.950863061 | 6.049571196 | 8.701662801 | 0.005155775 | 0.048437388 | Yes |
| 1559 | RMI1 | ENSG00000178966 | 3.138233337 | 5.165312459 | 8.627972722 | 0.0051558 | 0.048437388 | Yes |
| 1560 | MMP16 | ENSG00000156103 | 5.753057292 | 4.450471834 | 8.591294564 | 0.005182667 | 0.048642231 | Yes |
| 1561 | FAM83H | ENSG00000180921 | 4.206211765 | 8.106989342 | 8.736598167 | 0.005184251 | 0.048642231 | Yes |
| 1562 | EBNA1BP2 | ENSG00000117395 | 5.68983159 | 2.269253985 | 8.640916144 | 0.005187923 | 0.0486455 | Yes |
| 1563 | CCZ1B | ENSG00000146574 | 3.748720452 | 4.995217188 | 8.621245759 | 0.005218107 | 0.0488972 | Yes |
| 1564 | CD300A | ENSG00000167851 | 2.891779975 | 6.624204654 | 8.586936091 | 0.005229284 | 0.048934676 | Yes |
| 1565 | C2CD4B | ENSG00000205502 | 2.658774677 | -2.486230881 | 8.334985599 | 0.005233042 | 0.048934676 | Yes |
| 1566 | BFAR | ENSG00000103429 | 5.90910571 | -1.670651615 | 8.651887293 | 0.005238092 | 0.048934676 | Yes |
| 1567 | MUC1 | ENSG00000185499 | 6.219796469 | 4.983115124 | 8.572093968 | 0.00524154 | 0.048934676 | Yes |
| 1568 | HPGD | ENSG00000164120 | 5.110843332 | 5.816710827 | 8.628533131 | 0.005244317 | 0.048934676 | Yes |
| 1569 | TEFM | ENSG00000172171 | 2.660966761 | 4.570798856 | 8.5227383 | 0.005248444 | 0.048934676 | Yes |
| 1570 | COL11A1 | ENSG00000060718 | 4.995822297 | 8.92630749 | 8.745584785 | 0.00525292 | 0.048934676 | Yes |
| 1571 | ZNF230 | ENSG00000159882 | 2.182811131 | 5.726295378 | 8.458703648 | 0.005253505 | 0.048934676 | Yes |
| 1572 | IMPA1 | ENSG00000133731 | 3.287688482 | 3.963140211 | 8.534596253 | 0.005254049 | 0.048934676 | Yes |
| 1573 | SDCBP2.AS1 | | 2.488435058 | 6.131516978 | 8.512992721 | 0.005256915 | 0.048934676 | Yes |
| 1574 | CRPPA | ENSG00000214960 | 4.053128831 | -2.295109278 | 8.513528686 | 0.005258881 | 0.048934676 | Yes |
| 1575 | HEXB | ENSG00000049860 | 6.944354402 | 2.955818357 | 8.62866889 | 0.005290678 | 0.049199269 | Yes |
| 1576 | CALML4 | ENSG00000129007 | 2.762451444 | 6.453662926 | 8.535805479 | 0.005303396 | 0.049274192 | Yes |
| 1577 | GORAB | ENSG00000120370 | 4.139702058 | 4.359856385 | 8.533414074 | 0.005311897 | 0.049274192 | Yes |
| 1578 | EIF2S3 | ENSG00000130741 | 6.549875318 | 2.065943947 | 8.624075802 | 0.005313845 | 0.049274192 | Yes |
| 1579 | ENSG00000237975 | ENSG00000237975 | 2.646365814 | 6.330788268 | 8.515118496 | 0.005316395 | 0.049274192 | Yes |
| 1580 | SERPINB8 | ENSG00000166401 | 3.027603991 | 6.784124849 | 8.568316214 | 0.005319504 | 0.049274192 | Yes |
| 1581 | PAXIP1 | ENSG00000157212 | 3.751752089 | 3.282046658 | 8.481148408 | 0.005320387 | 0.049274192 | Yes |
| 1582 | CLDN4 | ENSG00000189143 | 5.339011955 | 9.284774149 | 8.727633197 | 0.005322299 | 0.049274192 | Yes |
| 1583 | ZNF22 | ENSG00000165512 | 5.406293424 | 2.314994548 | 8.559899904 | 0.005345685 | 0.049443161 | Yes |
| 1584 | CD99 | ENSG00000002586 | 8.530203325 | 2.413709114 | 8.637697103 | 0.005347306 | 0.049443161 | Yes |
| 1585 | TLR6 | ENSG00000174130 | 2.433455803 | 6.048203105 | 8.462085404 | 0.005355854 | 0.04949093 | Yes |
| 1586 | SH3BP5 | ENSG00000131370 | 6.657714082 | 2.174596473 | 8.601581077 | 0.005371412 | 0.049603377 | Yes |
| 1587 | GNB1 | ENSG00000078369 | 8.049864311 | 2.012860355 | 8.618980426 | 0.00538797 | 0.049700732 | Yes |
| 1588 | ZNF841 | ENSG00000197608 | 2.400901082 | 6.032512785 | 8.447000403 | 0.005388745 | 0.049700732 | Yes |
| 1589 | ZNF416 | ENSG00000083817 | 2.333521161 | 5.95750075 | 8.42945139 | 0.00540757 | 0.049816136 | Yes |
| 1590 | MNS1 | ENSG00000138587 | 5.816342295 | 3.993913508 | 8.506785738 | 0.005408064 | 0.049816136 | Yes |
| 1591 | ZFYVE26 | ENSG00000072121 | 5.316363775 | 2.912304789 | 8.508197412 | 0.005414048 | 0.049839887 | Yes |
