## Supplementary material for "Integrated transcriptomic analysis identifies Paternally Expressed Gene 10 as a novel candidate biomarker associated with aggressive features in liposarcoma": Table S2_H

| ID | Description | setSize | enrichmentScore | NES | pvalue | p.adjust | qvalue | rank | leading_edge | core_enrichment |
|---|---|---|---|---|---|---|---|---|---|---|
| HALLMARK_MYOGENESIS | HALLMARK_MYOGENESIS | 189 | -0.882005061 | -2.885735582 | 1E-10 | 1.66667E-09 | 1.12281E-09 | 681 | tags=44%, list=5%, signal=42% | FOXO4/MRAS/SYNGR2/SSPN/MAPRE3/ITGA7/LPIN1/FABP3/MYH8/MYH11/DTNA/TEAD4/MYBPH/DAPK2/PPFIA4/MAPK12/SIRT2/CKMT2/SPEG/CHRNA1/SORBS1/CRAT/MEF2C/NOS1/PTP4A3/MYL1/[illegible] |
| HALLMARK_OXIDATIVE_PHOSPHORYLATIO | HALLMARK_OXIDATIVE_PHOSPHORYLATION | 199 | -0.668908514 | -2.200475348 | 1E-10 | 1.66667E-09 | 1.12281E-09 | 1463 | tags=40%, list=10%, signal=36% | VDAC1/NDUFS1/GPI/NDUFA7/ATP5PO/SLC25A12/BCKDHA/SUCLA2/MRPL15/ATP5MC3/ATP5MF/ATP5F1A/NDUFA1/PDHA1/NNT/IDH3A/HADHB/NDUFB4/UQCRC2/OGDH/ATP5F1B/ECHS1/TIMM8B/[illegible] |
| HALLMARK_E2F_TARGETS | HALLMARK_E2F_TARGETS | 200 | 0.624947522 | 2.117320633 | 1E-10 | 1.66667E-09 | 1.12281E-09 | 2153 | tags=40%, list=15%, signal=35% | HMGB2/PLK1/LIG1/MCM2/H2AX/CDCA8/CDK1/PTTG1/CKS1B/CCNB2/AURKB/CENPE/STMN1/CDKN2A/LBR/KPNA2/TUBB/KIF4A/DCLRE1B/CNOT9/PAN2/RRM2/MAD2L1/SRSF1/EZH2/CDCA3/TACC3/[illegible] |
| HALLMARK_G2M_CHECKPOINT | HALLMARK_G2M_CHECKPOINT | 197 | 0.567046876 | 1.917033738 | 3.16452E-09 | 3.95565E-08 | 2.66486E-08 | 3666 | tags=50%, list=25%, signal=38% | PLK1/MCM2/H2AX/CDK1/ODF2/EFNA5/PTTG1/CKS1B/TPX2/CCNB2/TENT4A/AURKB/UBE2C/CENPE/STMN1/KIF11/SS18/LBR/KPNA2/KIF4A/NOTCH2/MAD2L1/CCNA2/SRSF1/STIL/EZH2/TACC3/E2F1/[illegible] |
| HALLMARK_INTERFERON_GAMMA_RESPON | HALLMARK_INTERFERON_GAMMA_RESPONSE | 177 | 0.553322252 | 1.842056104 | 6.9178E-08 | 6.9178E-07 | 4.66041E-07 | 3815 | tags=40%, list=26%, signal=30% | TRAFD1/IFNAR2/ADAR/IFI44L/MYD88/CASP7/RIPK2/HELZ2/CD86/CMPK2/RIPK1/GBP4/ARL4A/NUP93/TNFAIP6/IFIT1/PTPN2/OGFR/ISOC1/VCAM1/PSMB9/SOCS1/EPSTI1/HIF1A/OAS3/PSMB8/[illegible] |
| HALLMARK_KRAS_SIGNALING_DN | HALLMARK_KRAS_SIGNALING_DN | 106 | -0.659336353 | -1.990248465 | 1.93146E-06 | 1.60955E-05 | 1.08433E-05 | 526 | tags=14%, list=4%, signal=14% | FGGY/ZBTB16/P2RX6/TNNI3/NOS1/MYOT/PNMT/SMPX/MYH7/CACNG1/CKM/COQ8A/SYNPO/GAMT/RYR1 |
| HALLMARK_INTERFERON_ALPHA_RESPONSE | HALLMARK_INTERFERON_ALPHA_RESPONSE | 91 | 0.576482609 | 1.736337044 | 3.19668E-05 | 0.000228334 | 0.000153825 | 2777 | tags=41%, list=19%, signal=33% | TRAFD1/ADAR/IFI44L/TRIM5/RIPK2/HELZ2/CMPK2/CD47/GBP4/SAMD9/OGFR/PSMB9/PROCR/EPSTI1/PSMB8/UBE2L6/RSAD2/SAMD9L/PNPT1/CXCL10/B2M/TDRD7/PSME1/IFIT3/CSF1/PARP9/[illegible] |
| HALLMARK_ADIPOGENESIS | HALLMARK_ADIPOGENESIS | 196 | -0.537558345 | -1.766961831 | 8.42836E-05 | 0.000526772 | 0.000354878 | 1407 | tags=26%, list=10%, signal=23% | ATP5PO/ARAF/BCKDHA/GPAM/MRPL15/GHITM/CMPK1/IDH3A/COQ5/ECHS1/DHRS7B/CYC1/CIDEA/MIGA2/COQ9/SAMM50/NDUFAB1/COX8A/GPX3/SUCLG1/SPARCL1/ACADM/MGLL/GPX4/CMBL/[illegible] |
| HALLMARK_HYPOXIA | HALLMARK_HYPOXIA | 180 | -0.51066768 | -1.66098086 | 0.001036139 | 0.00575633 | 0.003877948 | 957 | tags=15%, list=7%, signal=14% | HK1/LARGE1/UGP2/PPARGC1A/GALK1/FOS/FOXO3/CDKN1C/SLC2A5/HDLBP/RRAGD/PFKFB3/SLC37A4/DTNA/PPFIA4/CCN5/TPI1/GPC1/PGM1/GYS1/PPP1R3C/PHKG1/GAPDH/ALDOA/PYGM/ENO3/[illegible] |
| HALLMARK_CHOLESTEROL_HOMEOSTASIS | HALLMARK_CHOLESTEROL_HOMEOSTASIS | 70 | 0.541988297 | 1.559660048 | 0.003762138 | 0.018810691 | 0.012672465 | 4180 | tags=57%, list=29%, signal=41% | PLAUR/GPX8/HMGCR/HMGCS1/TRIB3/LDLR/CPEB2/S100A11/ERRFI1/HSD17B7/CHKA/ALCAM/CXCL16/SC5D/GNAI1/LGMN/MAL2/PLSCR1/FBXO6/STARD4/ANXA5/CYP51A1/ALDOC/ACTG1/ANTXR2/[illegible] |
| HALLMARK_IL6_JAK_STAT3_SIGNALING | HALLMARK_IL6_JAK_STAT3_SIGNALING | 71 | 0.532856908 | 1.538488051 | 0.004672897 | 0.021240442 | 0.01430935 | 3809 | tags=41%, list=26%, signal=30% | MYD88/PTPN2/TNFRSF21/SOCS1/CBL/CXCL1/TLR2/CXCL10/OSMR/IL10RB/CSF1/CD44/IL2RA/STAM2/IL3RA/EBI3/IFNGR2/PIK3R5/TNFRSF12A/CSF2RB/IL15RA/SOCS3/CSF2RA/STAT1/CXCL11/IL12RB1/[illegible] |
| HALLMARK_WNT_BETA_CATENIN_SIGNALIN | HALLMARK_WNT_BETA_CATENIN_SIGNALING | 39 | 0.599713375 | 1.539740456 | 0.007352941 | 0.030637255 | 0.020639835 | 3024 | tags=41%, list=21%, signal=33% | SKP2/ADAM17/PPARD/NOTCH1/NCSTN/TP53/LEF1/NKD1/GNAI1/DKK1/MAML1/DLL1/DVL2/HDAC2/RBPJ/WNT5B |
| HALLMARK_COMPLEMENT | HALLMARK_COMPLEMENT | 166 | 0.426736823 | 1.410298892 | 0.012258188 | 0.046856481 | 0.031566472 | 4892 | tags=42%, list=33%, signal=28% | PLAUR/CASP7/RABIF/MMP14/CTSL/FN1/CASP9/PSMB9/LRP1/PFN1/GNAI3/CTSB/LYN/GMFB/CXCL1/PLA2G7/GNB2/TNFAIP3/LGMN/PDGFB/TIMP1/PRDM4/PRKCD/PLSCR1/GNB4/MMP15/CP/[illegible] |
| HALLMARK_TNFA_SIGNALING_VIA_NFKB | HALLMARK_TNFA_SIGNALING_VIA_NFKB | 182 | 0.416790727 | 1.39549101 | 0.013119815 | 0.046856481 | 0.031566472 | 3960 | tags=34%, list=27%, signal=25% | BTG3/PLAUR/PHLDA1/PLAU/RIPK2/LDLR/YRDC/TNFAIP6/B4GALT1/TRAF1/PTGER4/DUSP2/GPR183/SERPINB8/CXCL1/TLR2/PLPP3/PHLDA2/ID2/CXCL10/MXD1/TNFAIP3/CSF1/FUT4/CD44/[illegible] |
| HALLMARK_MYC_TARGETS_V1 | HALLMARK_MYC_TARGETS_V1 | 200 | 0.389970141 | 1.321217858 | 0.027143136 | 0.09047712 | 0.060953007 | 4052 | tags=30%, list=28%, signal=22% | MCM2/TYMS/KPNA2/MAD2L1/CCNA2/SRSF1/CDK4/H2AZ1/AP3S1/CDK2/CAD/SNRPA1/MCM4/MCM5/PPIA/SNRPD1/IARS1/HNRNPU/CDC45/FAM120A/PCNA/CSTF2/RANBP1/XPOT/SNRPD3/[illegible] |
| HALLMARK_FATTY_ACID_METABOLISM | HALLMARK_FATTY_ACID_METABOLISM | 145 | -0.429422882 | -1.358121196 | 0.038437035 | 0.118246946 | 0.0796611 | 1581 | tags=24%, list=11%, signal=22% | HMGCL/D2HGDH/UROS/ETFDH/EPHX1/GLUL/PPARA/SUCLA2/PDHA1/HADHB/ECHS1/HSDL2/CIDEA/VNN1/LGALS1/GSTZ1/ECI1/ACADVL/SUCLG1/ADSL/ACADM/MGLL/MLYCD/SDHA/GPD1/MDH1/[illegible] |
| HALLMARK_MITOTIC_SPINDLE | HALLMARK_MITOTIC_SPINDLE | 198 | 0.381804157 | 1.291482751 | 0.040203962 | 0.118246946 | 0.0796611 | 2990 | tags=28%, list=20%, signal=23% | PLK1/CDK1/TPX2/CCNB2/CENPE/KIF11/KIF4A/NOTCH2/FSCN1/CDC42/MID1/ARHGEF3/MYH10/NEK2/WASF1/SMC4/FBXO5/KIF23/MAPRE1/ABL1/FGD6/NDC80/NUSAP1/TTK/ARHGEF2/CDC42EP4/[illegible] |
| HALLMARK_ANGIOGENESIS | HALLMARK_ANGIOGENESIS | 28 | 0.584282885 | 1.408919232 | 0.043999343 | 0.122220398 | 0.082337952 | 2560 | tags=43%, list=17%, signal=35% | APP/SLCO2A1/FSTL1/TNFRSF21/POSTN/VAV2/COL3A1/STC1/TIMP1/FGFR1/VEGFA/COL5A2 |
| HALLMARK_UNFOLDED_PROTEIN_RESPONS | HALLMARK_UNFOLDED_PROTEIN_RESPONSE | 111 | 0.413804091 | 1.286983821 | 0.071762287 | 0.188848125 | 0.127224 | 4794 | tags=36%, list=33%, signal=24% | H2AX/CKS1B/SERP1/KDELR3/EXOC2/IFIT1/YWHAZ/STC2/EIF2AK3/PDIA6/IARS1/CALR/LSM1/EXOSC9/XPOT/TATDN2/SEC11A/EIF4A3/PSAT1/NPM1/VEGFA/SLC7A5/ALDH18A1/MTHFD2/HYOU1/XBP1/[illegible] |
| HALLMARK_ANDROGEN_RESPONSE | HALLMARK_ANDROGEN_RESPONSE | 91 | 0.430396499 | 1.296332923 | 0.078540852 | 0.196352131 | 0.13227933 | 5678 | tags=47%, list=39%, signal=29% | HMGCR/AKT1/HMGCS1/CDC14B/B4GALT1/GNAI3/HPGD/CCND3/INSIG1/B2M/KRT19/APPBP2/FADS1/PGM3/MAP7/MERTK/CAMKK2/LMAN1/UBE2J1/GSR/ACTN1/SLC26A2/XRCC5/PLPP1/ELOVL5/[illegible] |
| HALLMARK_EPITHELIAL_MESENCHYMAL_TR | HALLMARK_EPITHELIAL_MESENCHYMAL_TRANSIT | 191 | 0.363755907 | 1.226432329 | 0.08653473 | 0.206035072 | 0.138802575 | 2593 | tags=28%, list=18%, signal=23% | PLAUR/SNAI2/GJA1/MXRA5/NOTCH2/BMP1/MMP14/COLGALT1/LOXL2/BASP1/IGFBP3/SERPINE2/FSTL1/SERPINH1/FN1/LOXL1/CDH2/FBN2/VCAM1/COL1A1/TPM4/COPA/LRP1/COL11A1/COL5A1/[illegible] |
| HALLMARK_UV_RESPONSE_DN | HALLMARK_UV_RESPONSE_DN | 143 | 0.38329318 | 1.239689403 | 0.096408318 | 0.219109813 | 0.147610821 | 5406 | tags=38%, list=37%, signal=24% | SNAI2/TENT4A/GJA1/NOTCH2/LDLR/SLC7A1/COL1A1/RND3/CACNA1A/FZD2/COL11A1/MMP16/CITED2/PLPP3/VAV2/PIK3CD/COL3A1/INSIG1/CDON/ADGRL2/RUNX1/WDR37/GCNT1/ANXA2/RGS4/[illegible] |
| HALLMARK_PI3K_AKT_MTOR_SIGNALING | HALLMARK_PI3K_AKT_MTOR_SIGNALING | 99 | 0.398498559 | 1.21751164 | 0.137353434 | 0.298594421 | 0.201158347 | 1741 | tags=22%, list=12%, signal=20% | IRAK4/CDK1/RPS6KA1/AKT1/MYD88/SLC2A1/E2F1/CDK4/PLA2G12A/CDK2/TRIB3/PAK4/TBK1/RIPK1/SLA/PFN1/CALR/SFN/ARPC3/DAPP1/NGF/CLTC |
| HALLMARK_APICAL_JUNCTION | HALLMARK_APICAL_JUNCTION | 173 | -0.367406578 | -1.190883212 | 0.14395673 | 0.299909854 | 0.202044533 | 417 | tags=8%, list=3%, signal=8% | CLDN5/MAPK11/MADCAM1/CADM2/SPEG/CRAT/NEXN/CDH15/ACTA1/MAP3K20/ACTN2/NRAP/FLNC/GAMT |
| HALLMARK_NOTCH_SIGNALING | HALLMARK_NOTCH_SIGNALING | 32 | 0.500177044 | 1.239795195 | 0.163045281 | 0.316854758 | 0.213460047 | 4789 | tags=44%, list=33%, signal=30% | APH1A/NOTCH2/PPARD/NOTCH1/DLL1/HES1/FZD7/WNT2/SAP30/WNT5A/TCF7L2/JAG1/PSENEN/DTX4 |
| HALLMARK_GLYCOLYSIS | HALLMARK_GLYCOLYSIS | 182 | -0.358031425 | -1.165911813 | 0.164764474 | 0.316854758 | 0.213460047 | 568 | tags=10%, list=4%, signal=10% | MDH1/HDLBP/RRAGD/SLC37A4/HOMER1/NDUFV3/PPFIA4/TPI1/GPC1/FBP2/LHPP/GYS1/MDH2/GOT1/GOT2/AGL/ALDOA/PGAM2 |
| HALLMARK_PANCREAS_BETA_CELLS | HALLMARK_PANCREAS_BETA_CELLS | 19 | 0.557747834 | 1.240515095 | 0.174321994 | 0.322818507 | 0.217477731 | 2669 | tags=42%, list=18%, signal=34% | SRP9/PAX6/SEC11A/ELP4/DPP4/ABCC8/ISL1/MAFB |
| HALLMARK_APICAL_SURFACE | HALLMARK_APICAL_SURFACE | 36 | 0.480928175 | 1.21559291 | 0.180945969 | 0.323117801 | 0.217679361 | 1512 | tags=33%, list=10%, signal=30% | PLAUR/EFNA5/APP/SRPX/CRYBG1/CROCC/MDGA1/DCBLD2/B4GALT1/LYN/NCOA6/EPHB4 |
| HALLMARK_ALLOGRAFT_REJECTION | HALLMARK_ALLOGRAFT_REJECTION | 149 | 0.350576559 | 1.139035549 | 0.199426934 | 0.343839542 | 0.23163927 | 3842 | tags=34%, list=26%, signal=25% | IFNAR2/CDKN2A/AKT1/RIPK2/CD86/CD47/SOCS1/ITGAL/HIF1A/LYN/TLR6/TLR2/CCND3/B2M/TIMP1/CD28/CSF1/ABCE1/BCAT1/GCNT1/IL2RA/ELF4/PRF1/IL18/TAP1/TLR3/NPM1/ITGB2/NLRP3/[illegible] |
| HALLMARK_IL2_STAT5_SIGNALING | HALLMARK_IL2_STAT5_SIGNALING | 177 | -0.340688888 | -1.10820099 | 0.237021717 | 0.395036194 | 0.266129647 | 715 | tags=7%, list=5%, signal=7% | GPX4/SYNGR2/CDKN1C/RRAGD/RGS16/GABARAPL1/PHTF2/FAH/PLEC/CAPN3/S100A1/SLC29A2/ENO3 |
| HALLMARK_ESTROGEN_RESPONSE_EARLY | HALLMARK_ESTROGEN_RESPONSE_EARLY | 176 | 0.325657594 | 1.084460466 | 0.269519954 | 0.434709603 | 0.292856996 | 3900 | tags=32%, list=27%, signal=24% | RBBP8/GJA1/ADCY1/SLC2A1/KAZN/UGCG/TPBG/ISG20L2/PMAIP1/STC2/SLC39A6/B4GALT1/ZNF185/SLC19A2/PTGES/SH3BP5/MUC1/RHOBTB3/ELF3/RARA/SCARB1/NHERF1/SFN/KRT19/CD44/[illegible] |
| HALLMARK_HEME_METABOLISM | HALLMARK_HEME_METABOLISM | 163 | -0.338913882 | -1.088543561 | 0.278393641 | 0.434990065 | 0.293045938 | 779 | tags=12%, list=5%, signal=12% | BLVRB/HAGH/RBM38/OPTN/MARCHF8/FOXO3/TMCC2/ATP6V0A1/GLRX5/UBAC1/ACKR1/RCL1/BCAM/HBB/RAD23A/RNF123/NFE2L1/EZH1/ANK1/SPTB |
| HALLMARK_UV_RESPONSE_UP | HALLMARK_UV_RESPONSE_UP | 142 | -0.344188989 | -1.086515878 | 0.290048801 | 0.439467881 | 0.296062572 | 1005 | tags=13%, list=7%, signal=12% | FKBP4/GPX3/GRINA/HSPA2/SPR/CLTB/FOS/CDC34/PRKACA/CDKN1C/RAB27A/RRAD/CYB5R1/TUBA4A/PDLIM3/SLC25A4/SELENOW/ALDOA |
| HALLMARK_MTORC1_SIGNALING | HALLMARK_MTORC1_SIGNALING | 197 | 0.315044842 | 1.0650823 | 0.298843817 | 0.439476202 | 0.296068178 | 3777 | tags=38%, list=26%, signal=29% | PLK1/MCM2/HMGCR/SLC1A5/CYB5B/PHGDH/RRM2/ELOVL6/SLC2A1/HMGCS1/SERP1/TRIB3/SLC7A11/IDH1/LDLR/RAB1A/TBK1/SERPINH1/MCM4/SLA/ENO1/PPIA/SRD5A1/DDX39A/CALR/PSMG1/[illegible] |
| HALLMARK_BILE_ACID_METABOLISM | HALLMARK_BILE_ACID_METABOLISM | 82 | -0.371564759 | -1.076924149 | 0.324014968 | 0.442853739 | 0.298343572 | 1485 | tags=16%, list=10%, signal=14% | PEX11G/AR/SLC67A1/PEX12/ABCA5/RXRA/GNPAT/OPTN/MLYCD/NR3C2/PHYH/PFKM/IDH2 |
| HALLMARK_XENOBIOTIC_METABOLISM | HALLMARK_XENOBIOTIC_METABOLISM | 153 | -0.332219395 | -1.059410123 | 0.325816649 | 0.442853739 | 0.298343572 | 1056 | tags=12%, list=7%, signal=12% | PDLIM5/DCXR/ITIH4/VNN1/BLVRB/SAR1B/ATOH8/GSTM4/TNFRSF1A/GABARAPL1/ACO2/LONP1/FAH/PINK1/CDA/ABCC3/NDRG2/ALDH2/ATP2A2 |
| HALLMARK_KRAS_SIGNALING_UP | HALLMARK_KRAS_SIGNALING_UP | 165 | 0.318847377 | 1.052791787 | 0.327711767 | 0.442853739 | 0.298343572 | 4346 | tags=33%, list=30%, signal=23% | PLAUR/EPHB2/PLAU/ADAM8/CBX8/IGFBP3/PCSK1N/ADAM17/MPZL2/PRELID3B/YRDC/DCBLD2/TRAF1/CBL/PSMB8/FBXO4/EVI5/ID2/SNAP25/CXCL10/TNFAIP3/TMEM176B/NGF/MAP7/LY96/CLEC4A/[illegible] |
| HALLMARK_INFLAMMATORY_RESPONSE | HALLMARK_INFLAMMATORY_RESPONSE | 151 | 0.316429141 | 1.031336191 | 0.378790769 | 0.498408906 | 0.33577021 | 3516 | tags=34%, list=24%, signal=26% | PLAUR/RIPK2/MMP14/LDLR/TPBG/TNFAIP6/SLC7A1/SLC31A1/DCBLD2/GNAI3/PTGER4/HIF1A/LYN/GPR183/ITGB8/TLR2/PCDH7/CXCL10/MXD1/OSMR/TIMP1/FPR1/CSF1/CHST2/TNFRSF9/SGMS2/[illegible] |
| HALLMARK_APOPTOSIS | HALLMARK_APOPTOSIS | 143 | 0.315523254 | 1.020500377 | 0.403213611 | 0.516940526 | 0.34825467 | 5691 | tags=43%, list=39%, signal=27% | BTG3/HMGB2/APP/CASP7/CDK2/CASP9/PMAIP1/DAP/BNIP3L/LEF1/RARA/SC5D/CASP6/SLC20A1/TIMP1/CD44/PRF1/BCL2L11/TOP2A/IL18/DAP3/TAP1/MMP2/TIMP2/PLAT/GSR/BAX/TNFRSF12A/[illegible] |
| HALLMARK_MYC_TARGETS_V2 | HALLMARK_MYC_TARGETS_V2 | 57 | 0.365843307 | 1.015012804 | 0.437511017 | 0.546888771 | 0.36843033 | 1950 | tags=19%, list=13%, signal=17% | PLK1/PUS1/SLC19A1/CDK4/MCM4/MCM5/DUSP2/PLK4/HSPD1/BYSL/PRMT3 |
| HALLMARK_PROTEIN_SECRETION | HALLMARK_PROTEIN_SECRETION | 94 | 0.316903461 | 0.960626788 | 0.526579926 | 0.642170641 | 0.432620221 | 5058 | tags=37%, list=35%, signal=25% | NAPG/SCRN1/AP3S1/COPB1/ATP6V1B1/ATP1A1/CLN5/ARFIP1/SNX2/COPB2/CLTC/BET1/VPS45/LMAN1/AP3B1/MON2/CD63/PPT1/VPS4B/TMX1/STX12/VAMP4/ATP7A/SCAMP1/GALC/YKT6/CTSC/[illegible] |
| HALLMARK_ESTROGEN_RESPONSE_LATE | HALLMARK_ESTROGEN_RESPONSE_LATE | 167 | 0.287856218 | 0.951955704 | 0.558997896 | 0.665473686 | 0.448319115 | 3891 | tags=34%, list=27%, signal=25% | BTG3/RBBP8/STIL/MAPK13/MDK/TPBG/PERP/PTGES/DUSP2/ST14/UGDH/ACOX2/BATF/LAMC2/MEST/ID2/SCARB1/PLK4/NHERF1/SFN/KRT19/CD44/FOXC1/KIF20A/IGFBP4/NXT1/AREG/MICB/CDC20/[illegible] |
| HALLMARK_SPERMATOGENESIS | HALLMARK_SPERMATOGENESIS | 77 | 0.306087546 | 0.897308423 | 0.6481615 | 0.740040502 | 0.498553602 | 3332 | tags=31%, list=23%, signal=24% | CDK1/CCNB2/EZH2/NEK2/PCSK1N/ARL4A/CNIH2/CDKN3/TTK/PSMG1/RPL39L/SPATA6/MAP7/STAM2/KIF2C/GMCL1/CLGN/NCAPH/ACRBP/AURKA/ACE/MTOR/PGS1/PCSK4 |
| HALLMARK_TGF_BETA_SIGNALING | HALLMARK_TGF_BETA_SIGNALING | 51 | 0.32942679 | 0.893820075 | 0.651235642 | 0.740040502 | 0.498553602 | 5105 | tags=33%, list=35%, signal=22% | BCAR3/TRIM33/WWTR1/ID2/SLC20A1/ACVR1/SKI/IFNGR2/RAB31/MAP3K7/CTNNB1/CDK9/PPP1R15A/BMPR1A/XIAP/HDAC1/ID3 |
| HALLMARK_COAGULATION | HALLMARK_COAGULATION | 95 | -0.292170834 | -0.868292817 | 0.704240588 | 0.782489543 | 0.52715085 | 776 | tags=8%, list=5%, signal=8% | CRIP2/RAPGEF3/MSRB2/COMP/SIRT2/SH2B2/HPN/S100A1 |
| HALLMARK_PEROXISOME | HALLMARK_PEROXISOME | 90 | -0.274756262 | -0.809856669 | 0.81347487 | 0.865398798 | 0.583005506 | 1176 | tags=10%, list=8%, signal=9% | CLN8/GNPAT/CNBP/ABCB4/MLYCD/FIS1/CRAT/IDH2/SLC25A4 |
| HALLMARK_HEDGEHOG_SIGNALING | HALLMARK_HEDGEHOG_SIGNALING | 31 | -0.317932239 | -0.759002555 | 0.806207419 | 0.865398798 | 0.583005506 | 156 | tags=6%, list=1%, signal=6% | AMOT/ACHE |
| HALLMARK_DNA_REPAIR | HALLMARK_DNA_REPAIR | 148 | 0.250073055 | 0.812127268 | 0.898854962 | 0.936307252 | 0.630775412 | 2815 | tags=19%, list=19%, signal=15% | LIG1/TYMS/GTF2H3/GSDME/CMPK2/POLR2G/SNAPC4/RPA3/TP53/ITPA/PCNA/CCNO/ZWINT/AAAS/POLD3/CETN2/ERCC3/ALYREF/CLP1/SRSF6/PRIM1/RALA/ELOA/DGUOK/STX3/POLR1C/SDCBP/[illegible] |
| HALLMARK_P53_PATHWAY | HALLMARK_P53_PATHWAY | 190 | 0.240833304 | 0.811524723 | 0.929719264 | 0.948693127 | 0.63911958 | 1706 | tags=15%, list=12%, signal=13% | TRAFD1/CDKN2A/APP/PLK3/TRIB3/SLC7A11/DGKA/EPHA2/SERTAD3/NOTCH1/IP6K2/PROCR/SLC19A2/SOCS1/PERP/TP53/CDKN2B/ABHD4/ST14/NOL8/CCND3/MXD1/PCNA/SFN/MDM2/PVT1/CLCA2/[illegible] |
| HALLMARK_REACTIVE_OXYGEN_SPECIES_PA | HALLMARK_REACTIVE_OXYGEN_SPECIES_PATHWA | 44 | -0.185740539 | -0.479999764 | 1 | 1 | 0.673684211 | 1257 | tags=16%, list=9%, signal=15% | PRDX1/PRDX2/NDUFB4/GPX3/OXSR1/NDUFA6/GPX4 |
