## Supplementary material for "Integrated transcriptomic analysis identifies Paternally Expressed Gene 10 as a novel candidate biomarker associated with aggressive features in liposarcoma": Table S2_GOBP

| ID | Description | setSize | enrichmentScore | NES | pvalue | p.adjust | qvalue | rank | leading_edge | core_enrichment |
|---|---|---|---|---|---|---|---|---|---|---|
| GOBP_MUSCLE_CONTRACTION | GOBP_MUSCLE_CONTRACTION | 328 | -0.8261119 | -2.8736224 | 1E-10 | 1.5981E-08 | 1.4627E-08 | 485 | [illegible] | [illegible] |
| GOBP_MUSCLE_SYSTEM_PROCESS | GOBP_MUSCLE_SYSTEM_PROCESS | 412 | -0.7975523 | -2.8371924 | 1E-10 | 1.5981E-08 | 1.4627E-08 | 788 | [illegible] | [illegible] |
| GOBP_STRIATED_MUSCLE_CONTRACTION | GOBP_STRIATED_MUSCLE_CONTRACTION | 206 | -0.8408535 | -2.7899027 | 1E-10 | 1.5981E-08 | 1.4627E-08 | 531 | [illegible] | [illegible] |
| GOBP_MUSCLE_CELL_DEVELOPMENT | GOBP_MUSCLE_CELL_DEVELOPMENT | 172 | -0.830985 | -2.6869718 | 1E-10 | 1.5981E-08 | 1.4627E-08 | 518 | [illegible] | [illegible] |
| GOBP_MUSCLE_ORGAN_DEVELOPMENT | GOBP_MUSCLE_ORGAN_DEVELOPMENT | 309 | -0.7576993 | -2.6203229 | 1E-10 | 1.5981E-08 | 1.4627E-08 | 807 | [illegible] | [illegible] |
| GOBP_CELLULAR_COMPONENT_ASSEMBLY_INVOLVED_IN_MORPHOGE | GOBP_CELLULAR_COMPONENT_ASSEMBLY_INVOLVED_IN_MORPHO | 111 | -0.8592384 | -2.6007039 | 1E-10 | 1.5981E-08 | 1.4627E-08 | 481 | [illegible] | [illegible] |
| GOBP_REGULATION_OF_MUSCLE_SYSTEM_PROCESS | GOBP_REGULATION_OF_MUSCLE_SYSTEM_PROCESS | 196 | -0.7867095 | -2.585763 | 1E-10 | 1.5981E-08 | 1.4627E-08 | 788 | [illegible] | [illegible] |
| GOBP_REGULATION_OF_MUSCLE_CONTRACTION | GOBP_REGULATION_OF_MUSCLE_CONTRACTION | 135 | -0.826939 | -2.5840316 | 1E-10 | 1.5981E-08 | 1.4627E-08 | 442 | [illegible] | [illegible] |
| GOBP_MYOFIBRIL_ASSEMBLY | GOBP_MYOFIBRIL_ASSEMBLY | 69 | -0.9166686 | -2.5748415 | 1E-10 | 1.5981E-08 | 1.4627E-08 | 481 | [illegible] | [illegible] |
| GOBP_CARDIAC_MUSCLE_CONTRACTION | GOBP_CARDIAC_MUSCLE_CONTRACTION | 118 | -0.8386486 | -2.5712121 | 1E-10 | 1.5981E-08 | 1.4627E-08 | 427 | [illegible] | [illegible] |
| GOBP_ACTOMYOSIN_STRUCTURE_ORGANIZATION | GOBP_ACTOMYOSIN_STRUCTURE_ORGANIZATION | 190 | -0.7775892 | -2.5476186 | 1E-10 | 1.5981E-08 | 1.4627E-08 | 481 | [illegible] | [illegible] |
| GOBP_MUSCLE_STRUCTURE_DEVELOPMENT | GOBP_MUSCLE_STRUCTURE_DEVELOPMENT | 646 | -0.6836145 | -2.5331251 | 1E-10 | 1.5981E-08 | 1.4627E-08 | 836 | [illegible] | [illegible] |
| GOBP_REGULATION_OF_STRIATED_MUSCLE_CONTRACTION | GOBP_REGULATION_OF_STRIATED_MUSCLE_CONTRACTION | 83 | -0.8616609 | -2.4952269 | 1E-10 | 1.5981E-08 | 1.4627E-08 | 442 | [illegible] | [illegible] |
| GOBP_STRIATED_MUSCLE_CELL_DIFFERENTIATION | GOBP_STRIATED_MUSCLE_CELL_DIFFERENTIATION | 301 | -0.7213265 | -2.4862503 | 1E-10 | 1.5981E-08 | 1.4627E-08 | 836 | [illegible] | [illegible] |
| GOBP_MUSCLE_CELL_DIFFERENTIATION | GOBP_MUSCLE_CELL_DIFFERENTIATION | 398 | -0.6985343 | -2.4789506 | 1E-10 | 1.5981E-08 | 1.4627E-08 | 788 | [illegible] | [illegible] |
| GOBP_MULTICELLULAR_ORGANISMAL_MOVEMENT | GOBP_MULTICELLULAR_ORGANISMAL_MOVEMENT | 57 | -0.9088804 | -2.4648633 | 1E-10 | 1.5981E-08 | 1.4627E-08 | 485 | [illegible] | [illegible] |
| GOBP_HEART_PROCESS | GOBP_HEART_PROCESS | 196 | -0.7490592 | -2.4620136 | 1E-10 | 1.5981E-08 | 1.4627E-08 | 427 | [illegible] | [illegible] |
| GOBP_ACTIN_MEDIATED_CELL_CONTRACTION | GOBP_ACTIN_MEDIATED_CELL_CONTRACTION | 99 | -0.8202619 | -2.4425733 | 1E-10 | 1.5981E-08 | 1.4627E-08 | 485 | [illegible] | [illegible] |
| GOBP_REGULATION_OF_HEART_CONTRACTION | GOBP_REGULATION_OF_HEART_CONTRACTION | 157 | -0.7601955 | -2.4296751 | 1E-10 | 1.5981E-08 | 1.4627E-08 | 778 | [illegible] | [illegible] |
| GOBP_SKELETAL_MUSCLE_CONTRACTION | GOBP_SKELETAL_MUSCLE_CONTRACTION | 46 | -0.9319242 | -2.4296443 | 1E-10 | 1.5981E-08 | 1.4627E-08 | 485 | [illegible] | [illegible] |
| GOBP_SARCOMERE_ORGANIZATION | GOBP_SARCOMERE_ORGANIZATION | 45 | -0.9315382 | -2.4171692 | 1E-10 | 1.5981E-08 | 1.4627E-08 | 442 | [illegible] | [illegible] |
| GOBP_REGULATION_OF_SYSTEM_PROCESS | GOBP_REGULATION_OF_SYSTEM_PROCESS | 404 | -0.6711542 | -2.3845466 | 1E-10 | 1.5981E-08 | 1.4627E-08 | 788 | [illegible] | [illegible] |
| GOBP_SKELETAL_MUSCLE_ORGAN_DEVELOPMENT | GOBP_SKELETAL_MUSCLE_ORGAN_DEVELOPMENT | 165 | -0.738118 | -2.3743384 | 1E-10 | 1.5981E-08 | 1.4627E-08 | 989 | [illegible] | [illegible] |
| GOBP_ACTIN_FILAMENT_BASED_MOVEMENT | GOBP_ACTIN_FILAMENT_BASED_MOVEMENT | 120 | -0.7712493 | -2.3711595 | 1E-10 | 1.5981E-08 | 1.4627E-08 | 485 | [illegible] | [illegible] |
| GOBP_REGULATION_OF_BLOOD_CIRCULATION | GOBP_REGULATION_OF_BLOOD_CIRCULATION | 193 | -0.7213345 | -2.3690145 | 1E-10 | 1.5981E-08 | 1.4627E-08 | 778 | [illegible] | [illegible] |
| GOBP_MUSCLE_TISSUE_DEVELOPMENT | GOBP_MUSCLE_TISSUE_DEVELOPMENT | 365 | -0.670805 | -2.3595713 | 1E-10 | 1.5981E-08 | 1.4627E-08 | 820 | [illegible] | [illegible] |
| GOBP_REGULATION_OF_CARDIAC_MUSCLE_CONTRACTION | GOBP_REGULATION_OF_CARDIAC_MUSCLE_CONTRACTION | 66 | -0.8424786 | -2.3470067 | 1E-10 | 1.5981E-08 | 1.4627E-08 | 756 | [illegible] | [illegible] |
| GOBP_SARCOPLASMIC_RETICULUM_CALCIUM_ION_TRANSPORT | GOBP_SARCOPLASMIC_RETICULUM_CALCIUM_ION_TRANSPORT | 41 | -0.9169799 | -2.3403826 | 1E-10 | 1.5981E-08 | 1.4627E-08 | 756 | [illegible] | [illegible] |
| GOBP_REGULATION_OF_MONOATOMIC_ION_TRANSMEMBRANE_TRAN | GOBP_REGULATION_OF_MONOATOMIC_ION_TRANSMEMBRANE_TF | 204 | -0.7015685 | -2.3222627 | 1E-10 | 1.5981E-08 | 1.4627E-08 | 469 | [illegible] | [illegible] |
| GOBP_NEUROMUSCULAR_PROCESS | GOBP_NEUROMUSCULAR_PROCESS | 141 | -0.7361322 | -2.3160314 | 1E-10 | 1.5981E-08 | 1.4627E-08 | 556 | [illegible] | [illegible] |
| GOBP_MUSCLE_ADAPTATION | GOBP_MUSCLE_ADAPTATION | 92 | -0.7835282 | -2.3055204 | 1E-10 | 1.5981E-08 | 1.4627E-08 | 836 | [illegible] | [illegible] |
| GOBP_CELLULAR_RESPIRATION | GOBP_CELLULAR_RESPIRATION | 281 | -0.6637023 | -2.2772187 | 1E-10 | 1.5981E-08 | 1.4627E-08 | 1323 | [illegible] | [illegible] |
| GOBP_ACTIN_FILAMENT_BASED_PROCESS | GOBP_ACTIN_FILAMENT_BASED_PROCESS | 712 | -0.6081161 | -2.2710927 | 1E-10 | 1.5981E-08 | 1.4627E-08 | 699 | [illegible] | [illegible] |
| GOBP_ATP_METABOLIC_PROCESS | GOBP_ATP_METABOLIC_PROCESS | 201 | -0.6868528 | -2.268558 | 1E-10 | 1.5981E-08 | 1.4627E-08 | 1040 | [illegible] | [illegible] |
| GOBP_GENERATION_OF_PRECURSOR_METABOLITES_AND_ENERGY | GOBP_GENERATION_OF_PRECURSOR_METABOLITES_AND_ENERGY | 406 | -0.6378362 | -2.2663924 | 1E-10 | 1.5981E-08 | 1.4627E-08 | 1323 | [illegible] | [illegible] |
| GOBP_REGULATION_OF_CALCIUM_ION_TRANSPORT | GOBP_REGULATION_OF_CALCIUM_ION_TRANSPORT | 169 | -0.6997933 | -2.2565845 | 1E-10 | 1.5981E-08 | 1.4627E-08 | 464 | [illegible] | [illegible] |
| GOBP_REGULATION_OF_MONOATOMIC_ION_TRANSPORT | GOBP_REGULATION_OF_MONOATOMIC_ION_TRANSPORT | 292 | -0.6502757 | -2.2388204 | 1E-10 | 1.5981E-08 | 1.4627E-08 | 469 | [illegible] | [illegible] |
| GOBP_REGULATION_OF_TRANSMEMBRANE_TRANSPORT | GOBP_REGULATION_OF_TRANSMEMBRANE_TRANSPORT | 312 | -0.6317459 | -2.1864216 | 1E-10 | 1.5981E-08 | 1.4627E-08 | 871 | [illegible] | [illegible] |
| GOBP_CALCIUM_ION_TRANSPORT | GOBP_CALCIUM_ION_TRANSPORT | 303 | -0.629774 | -2.1741383 | 1E-10 | 1.5981E-08 | 1.4627E-08 | 756 | [illegible] | [illegible] |
| GOBP_CIRCULATORY_SYSTEM_PROCESS | GOBP_CIRCULATORY_SYSTEM_PROCESS | 456 | -0.5813547 | -2.0870569 | 1E-10 | 1.5981E-08 | 1.4627E-08 | 716 | [illegible] | [illegible] |
| GOBP_MONOATOMIC_CATION_TRANSPORT | GOBP_MONOATOMIC_CATION_TRANSPORT | 707 | -0.5439722 | -2.0301773 | 1E-10 | 1.5981E-08 | 1.4627E-08 | 1095 | [illegible] | [illegible] |
| GOBP_SUPRAMOLECULAR_FIBER_ORGANIZATION | GOBP_SUPRAMOLECULAR_FIBER_ORGANIZATION | 677 | -0.5306466 | -1.9741369 | 1E-10 | 1.5981E-08 | 1.4627E-08 | 663 | [illegible] | [illegible] |
| GOBP_MONOATOMIC_ION_TRANSMEMBRANE_TRANSPORT | GOBP_MONOATOMIC_ION_TRANSMEMBRANE_TRANSPORT | 656 | -0.527155 | -1.9556347 | 1E-10 | 1.5981E-08 | 1.4627E-08 | 756 | [illegible] | [illegible] |
| GOBP_METAL_ION_TRANSPORT | GOBP_METAL_ION_TRANSPORT | 561 | -0.5268205 | -1.9286478 | 1E-10 | 1.5981E-08 | 1.4627E-08 | 756 | [illegible] | [illegible] |
| GOBP_MONOATOMIC_ION_TRANSPORT | GOBP_MONOATOMIC_ION_TRANSPORT | 808 | -0.5115412 | -1.926624 | 1E-10 | 1.5981E-08 | 1.4627E-08 | 1095 | [illegible] | [illegible] |
| GOBP_CELL_DIVISION | GOBP_CELL_DIVISION | 571 | 0.48308737 | 1.83030966 | 1E-10 | 1.5981E-08 | 1.4627E-08 | 3173 | [illegible] | [illegible] |
| GOBP_TRANSMEMBRANE_TRANSPORT | GOBP_TRANSMEMBRANE_TRANSPORT | 1056 | -0.4529879 | -1.7361295 | 1E-10 | 1.5981E-08 | 1.4627E-08 | 1184 | [illegible] | [illegible] |
| GOBP_CALCIUM_ION_TRANSMEMBRANE_TRANSPORT | GOBP_CALCIUM_ION_TRANSMEMBRANE_TRANSPORT | 229 | -0.6461665 | -2.1685524 | 1.3103E-10 | 2.0504E-08 | 1.8767E-08 | 696 | [illegible] | [illegible] |
| GOBP_PROTON_MOTIVE_FORCE_DRIVEN_ATP_SYNTHESIS | GOBP_PROTON_MOTIVE_FORCE_DRIVEN_ATP_SYNTHESIS | 65 | -0.8296402 | -2.3028558 | 1.9303E-10 | 2.9589E-08 | 2.7082E-08 | 1564 | [illegible] | [illegible] |
| GOBP_MEMBRANELESS_ORGANELLE_ASSEMBLY | GOBP_MEMBRANELESS_ORGANELLE_ASSEMBLY | 420 | -0.5526465 | -1.9710053 | 2.7634E-10 | 4.1512E-08 | 3.7995E-08 | 503 | [illegible] | [illegible] |
| GOBP_CYTOSKELETON_ORGANIZATION | GOBP_CYTOSKELETON_ORGANIZATION | 1276 | -0.4249529 | -1.6475088 | 3.0871E-10 | 4.5466E-08 | 4.1614E-08 | 744 | [illegible] | [illegible] |
| GOBP_OXIDATIVE_PHOSPHORYLATION | GOBP_OXIDATIVE_PHOSPHORYLATION | 123 | -0.7210728 | -2.226329 | 4.0704E-10 | 5.8793E-08 | 5.3813E-08 | 1319 | [illegible] | [illegible] |
| GOBP_REGULATION_OF_CALCIUM_ION_TRANSMEMBRANE_TRANSPOR | GOBP_REGULATION_OF_CALCIUM_ION_TRANSMEMBRANE_TRANSF | 110 | -0.7345796 | -2.2182286 | 8.6658E-10 | 1.2281E-07 | 1.1241E-07 | 756 | [illegible] | [illegible] |
| GOBP_RELEASE_OF_SEQUESTERED_CALCIUM_ION_INTO_CYTOSOL_BY_ | GOBP_RELEASE_OF_SEQUESTERED_CALCIUM_ION_INTO_CYTOSOL_E | 36 | -0.896719 | -2.2305106 | 9.7644E-10 | 1.3582E-07 | 1.2431E-07 | 756 | [illegible] | [illegible] |
| GOBP_CARDIAC_MUSCLE_TISSUE_DEVELOPMENT | GOBP_CARDIAC_MUSCLE_TISSUE_DEVELOPMENT | 196 | -0.6339367 | -2.0836282 | 1.1511E-09 | 1.572E-07 | 1.4389E-07 | 820 | [illegible] | [illegible] |
| GOBP_PURINE_CONTAINING_COMPOUND_METABOLIC_PROCESS | GOBP_PURINE_CONTAINING_COMPOUND_METABOLIC_PROCESS | 480 | -0.5177065 | -1.8700741 | 1.7715E-09 | 2.376E-07 | 2.1747E-07 | 1431 | [illegible] | [illegible] |
| GOBP_ELECTRON_TRANSPORT_CHAIN | GOBP_ELECTRON_TRANSPORT_CHAIN | 101 | -0.7483305 | -2.2388536 | 1.8793E-09 | 2.4764E-07 | 2.2666E-07 | 1312 | [illegible] | [illegible] |
| GOBP_SEQUESTERING_OF_CALCIUM_ION | GOBP_SEQUESTERING_OF_CALCIUM_ION | 106 | -0.7249306 | -2.1783201 | 4.9585E-09 | 6.4212E-07 | 5.8773E-07 | 756 | [illegible] | [illegible] |
| GOBP_SKELETAL_MUSCLE_ADAPTATION | GOBP_SKELETAL_MUSCLE_ADAPTATION | 17 | -0.9675969 | -2.0579447 | 5.9382E-09 | 7.5596E-07 | 6.9193E-07 | 166 | [illegible] | [illegible] |
| GOBP_ATP_SYNTHESIS_COUPLED_ELECTRON_TRANSPORT | GOBP_ATP_SYNTHESIS_COUPLED_ELECTRON_TRANSPORT | 81 | -0.7716763 | -2.2289423 | 6.3359E-09 | 7.9315E-07 | 7.2596E-07 | 1312 | [illegible] | [illegible] |
| GOBP_RIBOSE_PHOSPHATE_METABOLIC_PROCESS | GOBP_RIBOSE_PHOSPHATE_METABOLIC_PROCESS | 325 | -0.5674478 | -1.972871 | 6.7474E-09 | 8.3081E-07 | 7.6043E-07 | 1040 | [illegible] | [illegible] |
| GOBP_REGULATION_OF_MUSCLE_ADAPTATION | GOBP_REGULATION_OF_MUSCLE_ADAPTATION | 71 | -0.7785763 | -2.1959589 | 7.8129E-09 | 9.465E-07 | 8.6632E-07 | 836 | [illegible] | [illegible] |
| GOBP_MICROTUBULE_CYTOSKELETON_ORGANIZATION_INVOLVED_IN_ | GOBP_MICROTUBULE_CYTOSKELETON_ORGANIZATION_INVOLVED_ | 161 | 0.5820211 | 1.91956702 | 8.0491E-09 | 9.5963E-07 | 8.7834E-07 | 2960 | [illegible] | [illegible] |
| GOBP_CHROMOSOME_SEGREGATION | GOBP_CHROMOSOME_SEGREGATION | 397 | 0.48969252 | 1.79907652 | 9.0512E-09 | 1.0622E-06 | 9.7226E-07 | 2630 | [illegible] | [illegible] |
| GOBP_MUSCLE_ORGAN_MORPHOGENESIS | GOBP_MUSCLE_ORGAN_MORPHOGENESIS | 67 | -0.795999 | -2.2236293 | 1.0161E-08 | 1.1741E-06 | 1.0747E-06 | 400 | [illegible] | [illegible] |
| GOBP_CARDIAC_MUSCLE_CELL_CONTRACTION | GOBP_CARDIAC_MUSCLE_CELL_CONTRACTION | 70 | -0.7809327 | -2.1987979 | 1.0439E-08 | 1.1879E-06 | 1.0873E-06 | 427 | [illegible] | [illegible] |
| GOBP_STRIATED_MUSCLE_ADAPTATION | GOBP_STRIATED_MUSCLE_ADAPTATION | 36 | -0.8795744 | -2.1878652 | 1.0597E-08 | 1.1879E-06 | 1.0873E-06 | 166 | [illegible] | [illegible] |
| GOBP_CELL_CYCLE_PROCESS | GOBP_CELL_CYCLE_PROCESS | 1162 | 0.38556973 | 1.53483907 | 1.1986E-08 | 0.000001324 | 1.2118E-06 | 2745 | [illegible] | [illegible] |
| GOBP_NUCLEOSIDE_TRIPHOSPHATE_METABOLIC_PROCESS | GOBP_NUCLEOSIDE_TRIPHOSPHATE_METABOLIC_PROCESS | 245 | -0.5940427 | -2.0047821 | 1.446E-08 | 0.000001574 | 1.4407E-06 | 1040 | [illegible] | [illegible] |
| GOBP_HEART_DEVELOPMENT | GOBP_HEART_DEVELOPMENT | 515 | -0.4971559 | -1.8074818 | 1.5219E-08 | 1.6329E-06 | 1.4946E-06 | 826 | [illegible] | [illegible] |
| GOBP_POSITIVE_REGULATION_OF_MONOATOMIC_ION_TRANSPORT | GOBP_POSITIVE_REGULATION_OF_MONOATOMIC_ION_TRANSPORT | 131 | -0.6849671 | -2.1327046 | 2.4154E-08 | 2.5552E-06 | 2.3387E-06 | 469 | [illegible] | [illegible] |
| GOBP_NUCLEOTIDE_METABOLIC_PROCESS | GOBP_NUCLEOTIDE_METABOLIC_PROCESS | 420 | -0.5234021 | -1.8667058 | 2.4908E-08 | 2.5984E-06 | 2.3783E-06 | 1040 | [illegible] | [illegible] |
| GOBP_ATP_BIOSYNTHETIC_PROCESS | GOBP_ATP_BIOSYNTHETIC_PROCESS | 90 | -0.7437634 | -2.1854716 | 2.8715E-08 | 2.9545E-06 | 2.7042E-06 | 1564 | [illegible] | [illegible] |
| GOBP_REGULATION_OF_RELEASE_OF_SEQUESTERED_CALCIUM_ION_IN | GOBP_REGULATION_OF_RELEASE_OF_SEQUESTERED_CALCIUM_ION | 31 | -0.8964273 | -2.1537529 | 3.2253E-08 | 3.2737E-06 | 2.9964E-06 | 756 | [illegible] | [illegible] |
| GOBP_CALCIUM_ION_HOMEOSTASIS | GOBP_CALCIUM_ION_HOMEOSTASIS | 232 | -0.6080643 | -2.0437568 | 3.6917E-08 | 3.6971E-06 | 3.3839E-06 | 696 | [illegible] | [illegible] |
| GOBP_NUCLEAR_CHROMOSOME_SEGREGATION | GOBP_NUCLEAR_CHROMOSOME_SEGREGATION | 292 | 0.49768132 | 1.77469746 | 4.9485E-08 | 4.8905E-06 | 4.4762E-06 | 2630 | [illegible] | [illegible] |
| GOBP_MYOTUBE_DIFFERENTIATION | GOBP_MYOTUBE_DIFFERENTIATION | 147 | -0.6612151 | -2.0914859 | 5.163E-08 | 5.0362E-06 | 4.6096E-06 | 600 | [illegible] | [illegible] |
| GOBP_MITOTIC_CELL_CYCLE | GOBP_MITOTIC_CELL_CYCLE | 839 | 0.40109813 | 1.56552194 | 5.2769E-08 | 5.0814E-06 | 4.6509E-06 | 2642 | [illegible] | [illegible] |
| GOBP_CALCIUM_ION_TRANSMEMBRANE_IMPORT_INTO_CYTOSOL | GOBP_CALCIUM_ION_TRANSMEMBRANE_IMPORT_INTO_CYTOSOL | 135 | -0.6805364 | -2.1265506 | 5.4008E-08 | 5.1086E-06 | 4.6759E-06 | 756 | [illegible] | [illegible] |
| GOBP_CALCIUM_MEDIATED_SIGNALING | GOBP_CALCIUM_MEDIATED_SIGNALING | 175 | -0.6222186 | -2.0187005 | 5.4412E-08 | 5.1086E-06 | 4.6759E-06 | 771 | [illegible] | [illegible] |
| GOBP_CELLULAR_HOMEOSTASIS | GOBP_CELLULAR_HOMEOSTASIS | 620 | -0.4687186 | -1.7294825 | 5.9253E-08 | 5.4944E-06 | 0.000005029 | 966 | [illegible] | [illegible] |
| GOBP_CHROMOSOME_ORGANIZATION | GOBP_CHROMOSOME_ORGANIZATION | 506 | 0.44476676 | 1.6707459 | 7.6543E-08 | 7.0111E-06 | 6.4172E-06 | 3493 | [illegible] | [illegible] |
| GOBP_REGULATION_OF_CARDIAC_MUSCLE_CONTRACTION_BY_CALCIU | GOBP_REGULATION_OF_CARDIAC_MUSCLE_CONTRACTION_BY_CAL( | 30 | -0.887707 | -2.117679 | 9.0036E-08 | 8.0507E-06 | 7.3687E-06 | 756 | [illegible] | [illegible] |
| GOBP_MITOTIC_CELL_CYCLE_PROCESS | GOBP_MITOTIC_CELL_CYCLE_PROCESS | 720 | 0.41229801 | 1.5921612 | 8.9535E-08 | 8.0507E-06 | 7.3687E-06 | 2642 | [illegible] | [illegible] |
| GOBP_INORGANIC_ION_HOMEOSTASIS | GOBP_INORGANIC_ION_HOMEOSTASIS | 394 | -0.5133453 | -1.8207545 | 9.9617E-08 | 8.8026E-06 | 8.0569E-06 | 850 | [illegible] | [illegible] |
| GOBP_MAINTENANCE_OF_LOCATION | GOBP_MAINTENANCE_OF_LOCATION | 180 | -0.6178796 | -2.0118213 | 1.1793E-07 | 0.000010299 | 9.4269E-06 | 756 | [illegible] | [illegible] |
| GOBP_ACTIN_MYOSIN_FILAMENT_SLIDING | GOBP_ACTIN_MYOSIN_FILAMENT_SLIDING | 15 | -0.9589984 | -1.9860325 | 1.486E-07 | 0.000012829 | 0.000011742 | 485 | [illegible] | [illegible] |
| GOBP_REGULATION_OF_ACTIN_FILAMENT_BASED_MOVEMENT | GOBP_REGULATION_OF_ACTIN_FILAMENT_BASED_MOVEMENT | 37 | -0.8665476 | -2.1652451 | 1.6584E-07 | 0.000014155 | 0.000012956 | 427 | [illegible] | [illegible] |
| GOBP_ACTIN_FILAMENT_ORGANIZATION | GOBP_ACTIN_FILAMENT_ORGANIZATION | 400 | -0.5062606 | -1.7980375 | 2.3632E-07 | 0.000019944 | 0.000018255 | 695 | [illegible] | [illegible] |
| GOBP_POSITIVE_REGULATION_OF_MONOATOMIC_ION_TRANSMEMBR | GOBP_POSITIVE_REGULATION_OF_MONOATOMIC_ION_TRANSMEM | 84 | -0.7251197 | -2.1044437 | 3.8389E-07 | 0.000031934 | 0.000029229 | 469 | [illegible] | [illegible] |
| GOBP_ANATOMICAL_STRUCTURE_FORMATION_INVOLVED_IN_MORPH | GOBP_ANATOMICAL_STRUCTURE_FORMATION_INVOLVED_IN_MOR | 1016 | -0.4087028 | -1.5619063 | 3.869E-07 | 0.000031934 | 0.000029229 | 807 | [illegible] | [illegible] |
| GOBP_DNA_METABOLIC_PROCESS | GOBP_DNA_METABOLIC_PROCESS | 905 | 0.38479793 | 1.50996692 | 4.4233E-07 | 0.000036113 | 0.000033053 | 3810 | [illegible] | [illegible] |
| GOBP_REGULATION_OF_ACTIN_FILAMENT_BASED_PROCESS | GOBP_REGULATION_OF_ACTIN_FILAMENT_BASED_PROCESS | 341 | -0.5223125 | -1.8244355 | 5.1019E-07 | 0.000041205 | 0.000037714 | 641 | [illegible] | [illegible] |
| GOBP_MITOCHONDRIAL_ELECTRON_TRANSPORT_NADH_TO_UBIQUIN( | GOBP_MITOCHONDRIAL_ELECTRON_TRANSPORT_NADH_TO_UBIQU | 43 | -0.821578 | -2.1162968 | 5.2161E-07 | 0.000041679 | 0.000038148 | 1540 | [illegible] | [illegible] |
| GOBP_CELL_CYCLE | GOBP_CELL_CYCLE | 1455 | 0.35256876 | 1.42015907 | 6.3271E-07 | 0.000050024 | 0.000045786 | 3029 | [illegible] | [illegible] |
| GOBP_MITOTIC_SISTER_CHROMATID_SEGREGATION | GOBP_MITOTIC_SISTER_CHROMATID_SEGREGATION | 196 | 0.53106927 | 1.80161551 | 6.4528E-07 | 0.000050486 | 0.00004621 | 2630 | [illegible] | [illegible] |
| GOBP_REGULATION_OF_SKELETAL_MUSCLE_CONTRACTION | GOBP_REGULATION_OF_SKELETAL_MUSCLE_CONTRACTION | 17 | -0.9400359 | -1.9993263 | 8.4591E-07 | 0.000065501 | 0.000059953 | 176 | [illegible] | [illegible] |
| GOBP_PROTON_TRANSMEMBRANE_TRANSPORT | GOBP_PROTON_TRANSMEMBRANE_TRANSPORT | 138 | -0.6391786 | -2.0030574 | 1.0702E-06 | 0.000082027 | 0.000075078 | 1481 | [illegible] | [illegible] |
| GOBP_MONOATOMIC_ION_HOMEOSTASIS | GOBP_MONOATOMIC_ION_HOMEOSTASIS | 449 | -0.4803752 | -1.7233599 | 1.0952E-06 | 0.000083093 | 0.000076054 | 815 | [illegible] | [illegible] |
| GOBP_CARDIAC_MUSCLE_TISSUE_MORPHOGENESIS | GOBP_CARDIAC_MUSCLE_TISSUE_MORPHOGENESIS | 49 | -0.7921232 | -2.0956178 | 1.4934E-06 | 0.00011106 | 0.00010165 | 400 | [illegible] | [illegible] |
| GOBP_MITOTIC_NUCLEAR_DIVISION | GOBP_MITOTIC_NUCLEAR_DIVISION | 269 | 0.48877032 | 1.72638641 | 1.4807E-06 | 0.00011106 | 0.00010165 | 2485 | [illegible] | [illegible] |
| GOBP_ACTION_POTENTIAL | GOBP_ACTION_POTENTIAL | 113 | -0.6623535 | -2.0119549 | 1.9286E-06 | 0.00014201 | 0.00012998 | 756 | [illegible] | [illegible] |
| GOBP_DNA_REPAIR | GOBP_DNA_REPAIR | 588 | 0.40999638 | 1.55684103 | 1.9977E-06 | 0.00014568 | 0.00013334 | 3805 | [illegible] | [illegible] |
| GOBP_NUCLEOBASE_CONTAINING_SMALL_MOLECULE_METABOLIC_PR | GOBP_NUCLEOBASE_CONTAINING_SMALL_MOLECULE_METABOLIC_ | 576 | -0.4504758 | -1.6512419 | 2.1838E-06 | 0.00015772 | 0.00014436 | 1447 | [illegible] | [illegible] |
| GOBP_POSITIVE_REGULATION_OF_TRANSMEMBRANE_TRANSPORT | GOBP_POSITIVE_REGULATION_OF_TRANSMEMBRANE_TRANSPORT | 128 | -0.6468631 | -2.0065983 | 2.6594E-06 | 0.00019024 | 0.00017412 | 469 | [illegible] | [illegible] |
| GOBP_REGULATION_OF_RELEASE_OF_SEQUESTERED_CALCIUM_ION_IN | GOBP_REGULATION_OF_RELEASE_OF_SEQUESTERED_CALCIUM_ION | 63 | -0.7465583 | -2.0645731 | 2.7325E-06 | 0.00019362 | 0.00017722 | 427 | [illegible] | [illegible] |
| GOBP_MYOTUBE_CELL_DEVELOPMENT | GOBP_MYOTUBE_CELL_DEVELOPMENT | 35 | -0.8316848 | -2.0594599 | 2.8273E-06 | 0.00019846 | 0.00018165 | 464 | [illegible] | [illegible] |
| GOBP_NERVOUS_SYSTEM_PROCESS | GOBP_NERVOUS_SYSTEM_PROCESS | 697 | -0.4313946 | -1.6082707 | 3.3998E-06 | 0.00023644 | 0.00021641 | 748 | [illegible] | [illegible] |
| GOBP_VENTRICULAR_CARDIAC_MUSCLE_TISSUE_DEVELOPMENT | GOBP_VENTRICULAR_CARDIAC_MUSCLE_TISSUE_DEVELOPMENT | 51 | -0.7723216 | -2.0560472 | 3.7179E-06 | 0.00025619 | 0.00023449 | 272 | [illegible] | [illegible] |
| GOBP_CHROMOSOME_LOCALIZATION | GOBP_CHROMOSOME_LOCALIZATION | 119 | 0.57105881 | 1.80171142 | 3.8047E-06 | 0.00025979 | 0.00023778 | 2728 | [illegible] | [illegible] |
| GOBP_CARDIAC_VENTRICLE_MORPHOGENESIS | GOBP_CARDIAC_VENTRICLE_MORPHOGENESIS | 57 | -0.7522351 | -2.0400447 | 4.0187E-06 | 0.00027193 | 0.00024889 | 272 | [illegible] | [illegible] |
| GOBP_REGULATION_OF_HEART_RATE | GOBP_REGULATION_OF_HEART_RATE | 78 | -0.7117709 | -2.0409688 | 4.6473E-06 | 0.00031106 | 0.00028471 | 778 | [illegible] | [illegible] |
| GOBP_CARDIAC_CELL_DEVELOPMENT | GOBP_CARDIAC_CELL_DEVELOPMENT | 76 | -0.7106508 | -2.0246164 | 4.6797E-06 | 0.00031106 | 0.00028471 | 518 | [illegible] | [illegible] |
| GOBP_PYRUVATE_METABOLIC_PROCESS | GOBP_PYRUVATE_METABOLIC_PROCESS | 107 | -0.6427729 | -1.9337814 | 5.5516E-06 | 0.00036337 | 0.00033258 | 836 | [illegible] | [illegible] |
| GOBP_REGULATION_OF_MEMBRANE_POTENTIAL | GOBP_REGULATION_OF_MEMBRANE_POTENTIAL | 297 | -0.5111728 | -1.7621071 | 5.5634E-06 | 0.00036337 | 0.00033258 | 799 | [illegible] | [illegible] |
| GOBP_POSITIVE_REGULATION_OF_CHROMOSOME_SEPARATION | GOBP_POSITIVE_REGULATION_OF_CHROMOSOME_SEPARATION | 31 | 0.74261293 | 1.83086894 | 6.2621E-06 | 0.00040547 | 0.00037112 | 2002 | [illegible] | [illegible] |
| GOBP_ORGANELLE_FISSION | GOBP_ORGANELLE_FISSION | 428 | 0.43068646 | 1.59325655 | 6.8868E-06 | 0.00044211 | 0.00040466 | 3056 | [illegible] | [illegible] |
| GOBP_CELL_CYCLE_CHECKPOINT_SIGNALING | GOBP_CELL_CYCLE_CHECKPOINT_SIGNALING | 191 | 0.51000454 | 1.72122934 | 6.9527E-06 | 0.00044256 | 0.00040507 | 2499 | [illegible] | [illegible] |
| GOBP_MITOTIC_SISTER_CHROMATID_SEPARATION | GOBP_MITOTIC_SISTER_CHROMATID_SEPARATION | 69 | 0.62576528 | 1.80468359 | 0.000007575 | 0.00047486 | 0.00043463 | 2002 | [illegible] | [illegible] |
| GOBP_REGULATION_OF_CELL_CYCLE_PROCESS | GOBP_REGULATION_OF_CELL_CYCLE_PROCESS | 678 | 0.39148117 | 1.50470546 | 7.5866E-06 | 0.00047486 | 0.00043463 | 3815 | [illegible] | [illegible] |
| GOBP_SISTER_CHROMATID_SEGREGATION | GOBP_SISTER_CHROMATID_SEGREGATION | 234 | 0.4870593 | 1.69046708 | 7.8781E-06 | 0.00048903 | 0.0004476 | 2485 | [illegible] | [illegible] |
| GOBP_CARBOHYDRATE_CATABOLIC_PROCESS | GOBP_CARBOHYDRATE_CATABOLIC_PROCESS | 142 | -0.6118238 | -1.9274399 | 0.000008078 | 0.00049732 | 0.0004552 | 836 | [illegible] | [illegible] |
| GOBP_REGULATION_OF_SODIUM_ION_TRANSMEMBRANE_TRANSPORT | GOBP_REGULATION_OF_SODIUM_ION_TRANSMEMBRANE_TRANSP( | 32 | -0.8276548 | -2.0071764 | 8.4325E-06 | 0.00051493 | 0.00047131 | 236 | [illegible] | [illegible] |
| GOBP_REGULATION_OF_TRANSPORT | GOBP_REGULATION_OF_TRANSPORT | 1221 | -0.3720304 | -1.4386851 | 8.5043E-06 | 0.00051513 | 0.00047149 | 1118 | [illegible] | [illegible] |
| GOBP_MITOTIC_SPINDLE_ORGANIZATION | GOBP_MITOTIC_SPINDLE_ORGANIZATION | 131 | 0.55570353 | 1.77908929 | 0.000008988 | 0.00054007 | 0.00049432 | 2960 | [illegible] | [illegible] |
| GOBP_METAPHASE_CHROMOSOME_ALIGNMENT | GOBP_METAPHASE_CHROMOSOME_ALIGNMENT | 105 | 0.57894288 | 1.79864627 | 9.4223E-06 | 0.00056167 | 0.00051409 | 2728 | [illegible] | [illegible] |
| GOBP_DNA_DAMAGE_RESPONSE | GOBP_DNA_DAMAGE_RESPONSE | 892 | 0.37192234 | 1.45801234 | 0.000010082 | 0.00059628 | 0.00054577 | 3766 | [illegible] | [illegible] |
| GOBP_CELL_CYCLE_PHASE_TRANSITION | GOBP_CELL_CYCLE_PHASE_TRANSITION | 521 | 0.4081873 | 1.53584442 | 0.000010789 | 0.00063307 | 0.00057945 | 2255 | [illegible] | [illegible] |
| GOBP_MUSCLE_HYPERTROPHY | GOBP_MUSCLE_HYPERTROPHY | 76 | -0.6987338 | -1.9906655 | 0.00001096 | 0.00063816 | 0.0005841 | 836 | [illegible] | [illegible] |
| GOBP_RIBOSE_PHOSPHATE_BIOSYNTHETIC_PROCESS | GOBP_RIBOSE_PHOSPHATE_BIOSYNTHETIC_PROCESS | 157 | -0.5876993 | -1.8783568 | 0.000011228 | 0.00063889 | 0.00058477 | 1447 | [illegible] | [illegible] |
| GOBP_HEART_MORPHOGENESIS | GOBP_HEART_MORPHOGENESIS | 226 | -0.5426912 | -1.8176281 | 0.000011158 | 0.00063889 | 0.00058477 | 826 | [illegible] | [illegible] |
| GOBP_SPINDLE_ORGANIZATION | GOBP_SPINDLE_ORGANIZATION | 198 | 0.50343769 | 1.7088765 | 0.000011118 | 0.00063889 | 0.00058477 | 2960 | [illegible] | [illegible] |
| GOBP_CARDIAC_MUSCLE_CELL_DIFFERENTIATION | GOBP_CARDIAC_MUSCLE_CELL_DIFFERENTIATION | 105 | -0.6447652 | -1.9339395 | 0.000011475 | 0.00064804 | 0.00059314 | 1056 | [illegible] | [illegible] |
| GOBP_VENTRICULAR_CARDIAC_MUSCLE_TISSUE_MORPHOGENESIS | GOBP_VENTRICULAR_CARDIAC_MUSCLE_TISSUE_MORPHOGENESIS | 38 | -0.7992794 | -2.0069728 | 0.000011716 | 0.00065673 | 0.0006011 | 272 | [illegible] | [illegible] |
| GOBP_CHEMICAL_HOMEOSTASIS | GOBP_CHEMICAL_HOMEOSTASIS | 780 | -0.4093634 | -1.5377204 | 0.000012428 | 0.00069143 | 0.00063286 | 1194 | [illegible] | [illegible] |
| GOBP_PROTEIN_LOCALIZATION_TO_CONDENSED_CHROMOSOME | GOBP_PROTEIN_LOCALIZATION_TO_CONDENSED_CHROMOSOME | 15 | 0.86836566 | 1.84793823 | 0.000012704 | 0.00070162 | 0.00064218 | 985 | [illegible] | [illegible] |
| GOBP_PURINE_CONTAINING_COMPOUND_BIOSYNTHETIC_PROCESS | GOBP_PURINE_CONTAINING_COMPOUND_BIOSYNTHETIC_PROCESS | 228 | -0.5434167 | -1.8223558 | 0.000013651 | 0.00074839 | 0.000685 | 1584 | [illegible] | [illegible] |
| GOBP_REGULATION_OF_ATP_DEPENDENT_ACTIVITY | GOBP_REGULATION_OF_ATP_DEPENDENT_ACTIVITY | 31 | -0.8334196 | -2.0023709 | 0.000014139 | 0.00076403 | 0.00069931 | 614 | [illegible] | [illegible] |
| GOBP_NUCLEOSIDE_PHOSPHATE_CATABOLIC_PROCESS | GOBP_NUCLEOSIDE_PHOSPHATE_CATABOLIC_PROCESS | 159 | -0.5786651 | -1.8507229 | 0.000014131 | 0.00076403 | 0.00069931 | 1015 | [illegible] | [illegible] |
| GOBP_RIBONUCLEOTIDE_CATABOLIC_PROCESS | GOBP_RIBONUCLEOTIDE_CATABOLIC_PROCESS | 116 | -0.6381779 | -1.95023 | 0.000016426 | 0.000875 | 0.00080088 | 865 | [illegible] | [illegible] |
| GOBP_ATTACHMENT_OF_SPINDLE_MICROTUBULES_TO_KINETOCHORE | GOBP_ATTACHMENT_OF_SPINDLE_MICROTUBULES_TO_KINETOCHO | 56 | 0.6503685 | 1.80047601 | 0.000016409 | 0.000875 | 0.00080088 | 1797 | [illegible] | [illegible] |
| GOBP_CARDIAC_MUSCLE_CELL_ACTION_POTENTIAL | GOBP_CARDIAC_MUSCLE_CELL_ACTION_POTENTIAL | 67 | -0.7129646 | -1.9916721 | 0.000017227 | 0.00089856 | 0.00082244 | 756 | [illegible] | [illegible] |
| GOBP_NUCLEOSIDE_DIPHOSPHATE_CATABOLIC_PROCESS | GOBP_NUCLEOSIDE_DIPHOSPHATE_CATABOLIC_PROCESS | 89 | -0.670881 | -1.9685645 | 0.000017217 | 0.00089856 | 0.00082244 | 836 | [illegible] | [illegible] |
| GOBP_CHROMOSOME_SEPARATION | GOBP_CHROMOSOME_SEPARATION | 86 | 0.59914346 | 1.79470429 | 0.000017223 | 0.00089856 | 0.00082244 | 3026 | [illegible] | [illegible] |
| GOBP_REGULATION_OF_MYOTUBE_DIFFERENTIATION | GOBP_REGULATION_OF_MYOTUBE_DIFFERENTIATION | 35 | -0.8049426 | -1.9932397 | 0.000018194 | 0.00093599 | 0.00085671 | 836 | [illegible] | [illegible] |
| GOBP_MEMBRANE_DEPOLARIZATION_DURING_ACTION_POTENTIAL | GOBP_MEMBRANE_DEPOLARIZATION_DURING_ACTION_POTENTIAI | 26 | -0.8581737 | -1.9859514 | 0.00001809 | 0.00093599 | 0.00085671 | 405 | [illegible] | [illegible] |
| GOBP_REPRODUCTIVE_PROCESS | GOBP_REPRODUCTIVE_PROCESS | 1083 | 0.35109904 | 1.39299299 | 0.000018853 | 0.00096332 | 0.00088171 | 4000 | [illegible] | [illegible] |
| GOBP_DOUBLE_STRAND_BREAK_REPAIR | GOBP_DOUBLE_STRAND_BREAK_REPAIR | 296 | 0.45656791 | 1.63134972 | 0.000019766 | 0.00100315 | 0.00091817 | 4689 | [illegible] | [illegible] |
| GOBP_PURINE_CONTAINING_COMPOUND_CATABOLIC_PROCESS | GOBP_PURINE_CONTAINING_COMPOUND_CATABOLIC_PROCESS | 138 | -0.5993237 | -1.8781604 | 0.000023335 | 0.0011763 | 0.00107666 | 1015 | [illegible] | [illegible] |
| GOBP_CALCIUM_ION_IMPORT | GOBP_CALCIUM_ION_IMPORT | 57 | -0.7202115 | -1.9531973 | 0.00002646 | 0.00132492 | 0.00121268 | 464 | [illegible] | [illegible] |
| GOBP_REGULATION_OF_STRIATED_MUSCLE_CELL_DIFFERENTIATION | GOBP_REGULATION_OF_STRIATED_MUSCLE_CELL_DIFFERENTIATIOI | 69 | -0.6988422 | -1.9629865 | 0.000028652 | 0.00141898 | 0.00129878 | 836 | [illegible] | [illegible] |
| GOBP_ORGANELLE_ASSEMBLY | GOBP_ORGANELLE_ASSEMBLY | 958 | -0.3813872 | -1.4515528 | 0.000028716 | 0.00141898 | 0.00129878 | 511 | [illegible] | [illegible] |
| GOBP_REGULATION_OF_CARDIAC_MUSCLE_CONTRACTION_BY_REGUL | GOBP_REGULATION_OF_CARDIAC_MUSCLE_CONTRACTION_BY_REG | 23 | -0.8715625 | -1.967997 | 0.000029201 | 0.00143353 | 0.00131209 | 756 | [illegible] | [illegible] |
| GOBP_PYRIDINE_CONTAINING_COMPOUND_CATABOLIC_PROCESS | GOBP_PYRIDINE_CONTAINING_COMPOUND_CATABOLIC_PROCESS | 90 | -0.6624597 | -1.9465692 | 0.000032112 | 0.00156618 | 0.00143351 | 1006 | [illegible] | [illegible] |
| GOBP_REGULATION_OF_MUSCLE_CELL_DIFFERENTIATION | GOBP_REGULATION_OF_MUSCLE_CELL_DIFFERENTIATION | 121 | -0.6097434 | -1.8764183 | 0.000033367 | 0.00161691 | 0.00147994 | 836 | [illegible] | [illegible] |
| GOBP_MEIOSIS_I_CELL_CYCLE_PROCESS | GOBP_MEIOSIS_I_CELL_CYCLE_PROCESS | 96 | 0.57378513 | 1.75761014 | 0.000040784 | 0.00196364 | 0.00179729 | 3244 | [illegible] | [illegible] |
| GOBP_CILIUM_ORGANIZATION | GOBP_CILIUM_ORGANIZATION | 380 | 0.4292033 | 1.56966292 | 0.000041627 | 0.00199147 | 0.00182277 | 3794 | [illegible] | [illegible] |
| GOBP_MITOTIC_METAPHASE_CHROMOSOME_ALIGNMENT | GOBP_MITOTIC_METAPHASE_CHROMOSOME_ALIGNMENT | 64 | 0.61824774 | 1.755134 | 0.000042488 | 0.0020198 | 0.0018487 | 2630 | [illegible] | [illegible] |
| GOBP_REGULATION_OF_MITOTIC_SISTER_CHROMATID_SEGREGATION | GOBP_REGULATION_OF_MITOTIC_SISTER_CHROMATID_SEGREGATI | 62 | 0.62593947 | 1.76714564 | 0.000048169 | 0.00227543 | 0.00208268 | 1697 | [illegible] | [illegible] |
| GOBP_RESPONSE_TO_MUSCLE_STRETCH | GOBP_RESPONSE_TO_MUSCLE_STRETCH | 25 | -0.8436057 | -1.9388271 | 0.00005192 | 0.00243733 | 0.00223086 | 251 | [illegible] | [illegible] |
| GOBP_CARDIAC_CHAMBER_MORPHOGENESIS | GOBP_CARDIAC_CHAMBER_MORPHOGENESIS | 111 | -0.6069754 | -1.8371657 | 0.000053986 | 0.00251856 | 0.00230521 | 309 | [illegible] | [illegible] |
| GOBP_GLUCAN_METABOLIC_PROCESS | GOBP_GLUCAN_METABOLIC_PROCESS | 72 | -0.6748469 | -1.9078744 | 0.000057131 | 0.00264883 | 0.00242445 | 1052 | [illegible] | [illegible] |
| GOBP_MEIOTIC_CELL_CYCLE_PROCESS | GOBP_MEIOTIC_CELL_CYCLE_PROCESS | 158 | 0.50509359 | 1.65988927 | 0.000059578 | 0.0027286 | 0.00249745 | 4213 | [illegible] | [illegible] |
| GOBP_DNA_RECOMBINATION | GOBP_DNA_RECOMBINATION | 289 | 0.44826624 | 1.59581196 | 0.000059498 | 0.0027286 | 0.00249745 | 5187 | [illegible] | [illegible] |
| GOBP_REGULATION_OF_MEMBRANE_REPOLARIZATION | GOBP_REGULATION_OF_MEMBRANE_REPOLARIZATION | 28 | -0.8160576 | -1.9205812 | 0.000060064 | 0.00273418 | 0.00250256 | 756 | [illegible] | [illegible] |
| GOBP_VOLUNTARY_SKELETAL_MUSCLE_CONTRACTION | GOBP_VOLUNTARY_SKELETAL_MUSCLE_CONTRACTION | 5 | -0.9892017 | -1.6798325 | 0.000061003 | 0.00276022 | 0.0025264 | 164 | [illegible] | [illegible] |
| GOBP_CARBOHYDRATE_METABOLIC_PROCESS | GOBP_CARBOHYDRATE_METABOLIC_PROCESS | 450 | -0.4391701 | -1.5758647 | 0.000062523 | 0.00281205 | 0.00257384 | 1052 | [illegible] | [illegible] |
| GOBP_REGULATION_OF_CELL_CYCLE | GOBP_REGULATION_OF_CELL_CYCLE | 971 | 0.34868981 | 1.37292521 | 0.000064513 | 0.00288425 | 0.00263992 | 3815 | [illegible] | [illegible] |
| GOBP_CARDIAC_CONDUCTION | GOBP_CARDIAC_CONDUCTION | 70 | -0.6767469 | -1.905452 | 0.00006567 | 0.00290147 | 0.00265568 | 756 | [illegible] | [illegible] |
| GOBP_REGULATION_OF_SKELETAL_MUSCLE_ADAPTATION | GOBP_REGULATION_OF_SKELETAL_MUSCLE_ADAPTATION | 10 | -0.9532664 | -1.8293974 | 0.000065356 | 0.00290147 | 0.00265568 | 162 | [illegible] | [illegible] |
| GOBP_PYRIDINE_CONTAINING_COMPOUND_METABOLIC_PROCESS | GOBP_PYRIDINE_CONTAINING_COMPOUND_METABOLIC_PROCESS | 153 | -0.5670094 | -1.8054131 | 0.000070194 | 0.00308322 | 0.00282204 | 836 | [illegible] | [illegible] |
| GOBP_REGULATION_OF_CARDIAC_MUSCLE_CELL_MEMBRANE_REPOLA | GOBP_REGULATION_OF_CARDIAC_MUSCLE_CELL_MEMBRANE_REP( | 20 | -0.8732796 | -1.9155782 | 0.000078619 | 0.00336247 | 0.00307763 | 756 | [illegible] | [illegible] |
| GOBP_TRANSITION_BETWEEN_FAST_AND_SLOW_FIBER | GOBP_TRANSITION_BETWEEN_FAST_AND_SLOW_FIBER | 6 | -0.9876355 | -1.731549 | 0.000078374 | 0.00336247 | 0.00307763 | 149 | [illegible] | [illegible] |
| GOBP_MITOTIC_CYTOKINESIS | GOBP_MITOTIC_CYTOKINESIS | 97 | 0.56131342 | 1.72265902 | 0.00007879 | 0.00336247 | 0.00307763 | 2419 | [illegible] | [illegible] |
| GOBP_SMOOTHENED_SIGNALING_PATHWAY | GOBP_SMOOTHENED_SIGNALING_PATHWAY | 133 | 0.53453719 | 1.71536518 | 0.000077236 | 0.00336247 | 0.00307763 | 3106 | [illegible] | [illegible] |
| GOBP_POSITIVE_REGULATION_OF_ACTIN_FILAMENT_BASED_MOVEME | GOBP_POSITIVE_REGULATION_OF_ACTIN_FILAMENT_BASED_MOVE | 5 | -0.9884356 | -1.6785315 | 0.000077924 | 0.00336247 | 0.00307763 | 63 | [illegible] | [illegible] |
| GOBP_REGULATION_OF_TRANSMEMBRANE_TRANSPORTER_ACTIVITY | GOBP_REGULATION_OF_TRANSMEMBRANE_TRANSPORTER_ACTIVIT | 75 | -0.6710672 | -1.9092645 | 0.000081378 | 0.00339572 | 0.00310807 | 835 | [illegible] | [illegible] |
| GOBP_CARDIAC_MYOFIBRIL_ASSEMBLY | GOBP_CARDIAC_MYOFIBRIL_ASSEMBLY | 17 | -0.894998 | -1.903537 | 0.000080726 | 0.00339572 | 0.00310807 | 356 | [illegible] | [illegible] |
| GOBP_REGULATION_OF_VENTRICULAR_CARDIAC_MUSCLE_CELL_MEMI | GOBP_REGULATION_OF_VENTRICULAR_CARDIAC_MUSCLE_CELL_ME | 19 | -0.8737137 | -1.8933123 | 0.000081027 | 0.00339572 | 0.00310807 | 756 | [illegible] | [illegible] |
| GOBP_POSITIVE_REGULATION_OF_MITOTIC_SISTER_CHROMATID_SEPA | GOBP_POSITIVE_REGULATION_OF_MITOTIC_SISTER_CHROMATID_SI | 21 | 0.78195376 | 1.78610637 | 0.000080646 | 0.00339572 | 0.00310807 | 2002 | [illegible] | [illegible] |
| GOBP_T_TUBULE_ORGANIZATION | GOBP_T_TUBULE_ORGANIZATION | 6 | -0.9873551 | -1.7310573 | 0.000085539 | 0.00354964 | 0.00324895 | 190 | [illegible] | [illegible] |
| GOBP_REGULATION_OF_NUCLEAR_DIVISION | GOBP_REGULATION_OF_NUCLEAR_DIVISION | 127 | 0.51948405 | 1.65485125 | 0.000096631 | 0.0039879 | 0.00365008 | 2002 | [illegible] | [illegible] |
| GOBP_GLUCOSE_METABOLIC_PROCESS | GOBP_GLUCOSE_METABOLIC_PROCESS | 168 | -0.5473999 | -1.7642298 | 0.00010011 | 0.00409251 | 0.00374583 | 1050 | [illegible] | [illegible] |
| GOBP_REGULATION_OF_CELL_CYCLE_PHASE_TRANSITION | GOBP_REGULATION_OF_CELL_CYCLE_PHASE_TRANSITION | 425 | 0.40838627 | 1.51138269 | 0.00010026 | 0.00409251 | 0.00374583 | 2255 | [illegible] | [illegible] |
| GOBP_SMALL_MOLECULE_METABOLIC_PROCESS | GOBP_SMALL_MOLECULE_METABOLIC_PROCESS | 1353 | -0.3488815 | -1.3578499 | 0.00010337 | 0.00419674 | 0.00384123 | 1249 | [illegible] | [illegible] |
| GOBP_MONOSACCHARIDE_CATABOLIC_PROCESS | GOBP_MONOSACCHARIDE_CATABOLIC_PROCESS | 42 | -0.7536596 | -1.9334541 | 0.00010544 | 0.00423954 | 0.00388041 | 391 | [illegible] | [illegible] |
| GOBP_RECOMBINATIONAL_REPAIR | GOBP_RECOMBINATIONAL_REPAIR | 178 | 0.49276715 | 1.65051639 | 0.00010555 | 0.00423954 | 0.00388041 | 4689 | [illegible] | [illegible] |
| GOBP_POTASSIUM_ION_TRANSPORT | GOBP_POTASSIUM_ION_TRANSPORT | 128 | -0.5860246 | -1.8178747 | 0.00010858 | 0.00433818 | 0.00397068 | 1035 | [illegible] | [illegible] |
| GOBP_REGULATION_OF_THE_FORCE_OF_HEART_CONTRACTION | GOBP_REGULATION_OF_THE_FORCE_OF_HEART_CONTRACTION | 20 | -0.8685014 | -1.9050971 | 0.00011678 | 0.00461716 | 0.00422604 | 427 | [illegible] | [illegible] |
| GOBP_NUCLEOTIDE_BIOSYNTHETIC_PROCESS | GOBP_NUCLEOTIDE_BIOSYNTHETIC_PROCESS | 192 | -0.5234765 | -1.7188338 | 0.0001168 | 0.00461716 | 0.00422604 | 1418 | [illegible] | [illegible] |
| GOBP_NUCLEOSIDE_DIPHOSPHATE_METABOLIC_PROCESS | GOBP_NUCLEOSIDE_DIPHOSPHATE_METABOLIC_PROCESS | 116 | -0.6050331 | -1.8489416 | 0.00012135 | 0.00474863 | 0.00434636 | 836 | [illegible] | [illegible] |
| GOBP_CIRCULATORY_SYSTEM_DEVELOPMENT | GOBP_CIRCULATORY_SYSTEM_DEVELOPMENT | 984 | -0.3723059 | -1.4205406 | 0.00012139 | 0.00474863 | 0.00434636 | 826 | [illegible] | [illegible] |
| GOBP_REGULATION_OF_MUSCLE_ORGAN_DEVELOPMENT | GOBP_REGULATION_OF_MUSCLE_ORGAN_DEVELOPMENT | 33 | -0.7784081 | -1.8984086 | 0.00012884 | 0.005014 | 0.00458926 | 380 | [illegible] | [illegible] |
| GOBP_CARDIOCYTE_DIFFERENTIATION | GOBP_CARDIOCYTE_DIFFERENTIATION | 135 | -0.5831175 | -1.8221343 | 0.00013224 | 0.00511976 | 0.00468606 | 1056 | [illegible] | [illegible] |
| GOBP_POSITIVE_REGULATION_OF_CELL_CYCLE_PROCESS | GOBP_POSITIVE_REGULATION_OF_CELL_CYCLE_PROCESS | 222 | 0.46144686 | 1.59016595 | 0.00013441 | 0.00517732 | 0.00473875 | 3805 | [illegible] | [illegible] |
| GOBP_RELAXATION_OF_MUSCLE | GOBP_RELAXATION_OF_MUSCLE | 30 | -0.8053714 | -1.9212623 | 0.00013785 | 0.00527505 | 0.0048282 | 696 | [illegible] | [illegible] |
| GOBP_POSITIVE_REGULATION_OF_MUSCLE_CELL_DIFFERENTIATION | GOBP_POSITIVE_REGULATION_OF_MUSCLE_CELL_DIFFERENTIATION | 55 | -0.7061533 | -1.9053613 | 0.00013836 | 0.00527505 | 0.0048282 | 650 | [illegible] | [illegible] |
| GOBP_REGULATION_OF_MITOTIC_NUCLEAR_DIVISION | GOBP_REGULATION_OF_MITOTIC_NUCLEAR_DIVISION | 112 | 0.54146705 | 1.69473692 | 0.00014039 | 0.00532557 | 0.00487444 | 2002 | [illegible] | [illegible] |
| GOBP_MEIOTIC_CHROMOSOME_SEGREGATION | GOBP_MEIOTIC_CHROMOSOME_SEGREGATION | 66 | 0.6143815 | 1.75579761 | 0.00014165 | 0.00534641 | 0.00489351 | 2969 | [illegible] | [illegible] |
| GOBP_NUCLEOSIDE_TRIPHOSPHATE_BIOSYNTHETIC_PROCESS | GOBP_NUCLEOSIDE_TRIPHOSPHATE_BIOSYNTHETIC_PROCESS | 113 | -0.6012054 | -1.8262123 | 0.00014408 | 0.00541093 | 0.00495256 | 1447 | [illegible] | [illegible] |
| GOBP_NEGATIVE_REGULATION_OF_CHROMOSOME_ORGANIZATION | GOBP_NEGATIVE_REGULATION_OF_CHROMOSOME_ORGANIZATION | 98 | 0.54769778 | 1.68405732 | 0.00014817 | 0.00553691 | 0.00506787 | 2246 | [illegible] | [illegible] |
| GOBP_ENERGY_RESERVE_METABOLIC_PROCESS | GOBP_ENERGY_RESERVE_METABOLIC_PROCESS | 82 | -0.6425746 | -1.861664 | 0.00016071 | 0.00597556 | 0.00546936 | 1052 | [illegible] | [illegible] |
| GOBP_SMALL_MOLECULE_CATABOLIC_PROCESS | GOBP_SMALL_MOLECULE_CATABOLIC_PROCESS | 273 | -0.4743651 | -1.6208763 | 0.00017056 | 0.0063107 | 0.00577612 | 1157 | [illegible] | [illegible] |
| GOBP_SPINDLE_ASSEMBLY | GOBP_SPINDLE_ASSEMBLY | 133 | 0.5218813 | 1.6747516 | 0.00017879 | 0.00658268 | 0.00602506 | 2630 | [illegible] | [illegible] |
| GOBP_MICROTUBULE_BASED_PROCESS | GOBP_MICROTUBULE_BASED_PROCESS | 834 | 0.34793977 | 1.35714694 | 0.00019233 | 0.00704683 | 0.00644988 | 3035 | [illegible] | [illegible] |
| GOBP_REGULATION_OF_IMMUNE_SYSTEM_PROCESS | GOBP_REGULATION_OF_IMMUNE_SYSTEM_PROCESS | 1233 | 0.32769645 | 1.3082035 | 0.00020183 | 0.00735904 | 0.00673565 | 4218 | [illegible] | [illegible] |
| GOBP_POINTED_END_ACTIN_FILAMENT_CAPPING | GOBP_POINTED_END_ACTIN_FILAMENT_CAPPING | 7 | -0.9684755 | -1.7425454 | 0.00020646 | 0.0074915 | 0.00685689 | 320 | [illegible] | [illegible] |
| GOBP_POSITIVE_REGULATION_OF_STRIATED_MUSCLE_CELL_DIFFEREN | GOBP_POSITIVE_REGULATION_OF_STRIATED_MUSCLE_CELL_DIFFER | 30 | -0.7940178 | -1.8941779 | 0.00023444 | 0.00837235 | 0.00766312 | 650 | [illegible] | [illegible] |
| GOBP_POSITIVE_REGULATION_OF_CALCIUM_ION_TRANSPORT | GOBP_POSITIVE_REGULATION_OF_CALCIUM_ION_TRANSPORT | 69 | -0.6687867 | -1.8785632 | 0.0002352 | 0.00837235 | 0.00766312 | 464 | [illegible] | [illegible] |
| GOBP_SODIUM_ION_TRANSMEMBRANE_TRANSPORT | GOBP_SODIUM_ION_TRANSMEMBRANE_TRANSPORT | 100 | -0.6150044 | -1.8384441 | 0.00023336 | 0.00837235 | 0.00766312 | 394 | [illegible] | [illegible] |
| GOBP_REGULATION_OF_SUPRAMOLECULAR_FIBER_ORGANIZATION | GOBP_REGULATION_OF_SUPRAMOLECULAR_FIBER_ORGANIZATION | 337 | -0.4551574 | -1.5882363 | 0.00023315 | 0.00837235 | 0.00766312 | 641 | [illegible] | [illegible] |
| GOBP_MEMBRANE_REPOLARIZATION | GOBP_MEMBRANE_REPOLARIZATION | 39 | -0.7484781 | -1.8916842 | 0.0002551 | 0.00903804 | 0.00827242 | 756 | [illegible] | [illegible] |
| GOBP_ENDOPLASMIC_RETICULUM_ORGANIZATION | GOBP_ENDOPLASMIC_RETICULUM_ORGANIZATION | 93 | -0.6155001 | -1.8151455 | 0.00026032 | 0.00909765 | 0.00832698 | 845 | [illegible] | [illegible] |
| GOBP_POSITIVE_REGULATION_OF_MITOCHONDRIAL_CALCIUM_ION_C( | GOBP_POSITIVE_REGULATION_OF_MITOCHONDRIAL_CALCIUM_ION | 9 | -0.9500185 | -1.7885277 | 0.00026013 | 0.00909765 | 0.00832698 | 7 | [illegible] | [illegible] |
| GOBP_ORGANOPHOSPHATE_METABOLIC_PROCESS | GOBP_ORGANOPHOSPHATE_METABOLIC_PROCESS | 892 | -0.3698289 | -1.4012583 | 0.00026042 | 0.00909765 | 0.00832698 | 1054 | [illegible] | [illegible] |
| GOBP_MEIOTIC_CELL_CYCLE | GOBP_MEIOTIC_CELL_CYCLE | 226 | 0.45101221 | 1.5593082 | 0.00027473 | 0.00955314 | 0.00874388 | 3309 | [illegible] | [illegible] |
| GOBP_POSITIVE_REGULATION_OF_HEART_RATE | GOBP_POSITIVE_REGULATION_OF_HEART_RATE | 19 | -0.8544827 | -1.8516394 | 0.00028383 | 0.00982409 | 0.00899188 | 394 | [illegible] | [illegible] |
| GOBP_NEGATIVE_REGULATION_OF_NUCLEAR_DIVISION | GOBP_NEGATIVE_REGULATION_OF_NUCLEAR_DIVISION | 65 | 0.59062545 | 1.68337799 | 0.00028782 | 0.00991642 | 0.00907639 | 1697 | [illegible] | [illegible] |
| GOBP_ATTACHMENT_OF_MITOTIC_SPINDLE_MICROTUBULES_TO_KINE | GOBP_ATTACHMENT_OF_MITOTIC_SPINDLE_MICROTUBULES_TO_KI | 26 | 0.72257499 | 1.71918335 | 0.0002907 | 0.00997024 | 0.00912565 | 2485 | [illegible] | [illegible] |
| GOBP_GLUCOSE_CATABOLIC_PROCESS | GOBP_GLUCOSE_CATABOLIC_PROCESS | 23 | -0.8360503 | -1.8878099 | 0.00029375 | 0.01001743 | 0.00916885 | 391 | [illegible] | [illegible] |
| GOBP_NEGATIVE_REGULATION_OF_CELL_CYCLE_PROCESS | GOBP_NEGATIVE_REGULATION_OF_CELL_CYCLE_PROCESS | 288 | 0.4307667 | 1.53315818 | 0.00029475 | 0.01001743 | 0.00916885 | 2745 | [illegible] | [illegible] |
| GOBP_VENTRICULAR_CARDIAC_MUSCLE_CELL_MEMBRANE_REPOLARIZ | GOBP_VENTRICULAR_CARDIAC_MUSCLE_CELL_MEMBRANE_REPOLA | 25 | -0.81442 | -1.8717509 | 0.00029774 | 0.01007344 | 0.00922011 | 756 | [illegible] | [illegible] |
| GOBP_MEMBRANE_DEPOLARIZATION_DURING_CARDIAC_MUSCLE_CEL | GOBP_MEMBRANE_DEPOLARIZATION_DURING_CARDIAC_MUSCLE_ | 18 | -0.8640677 | -1.8535757 | 0.00030631 | 0.01030394 | 0.00943108 | 405 | [illegible] | [illegible] |
| GOBP_ORGANOPHOSPHATE_CATABOLIC_PROCESS | GOBP_ORGANOPHOSPHATE_CATABOLIC_PROCESS | 201 | -0.5100294 | -1.6845403 | 0.00030729 | 0.01030394 | 0.00943108 | 1015 | [illegible] | [illegible] |
| GOBP_MUSCLE_CELL_CELLULAR_HOMEOSTASIS | GOBP_MUSCLE_CELL_CELLULAR_HOMEOSTASIS | 18 | -0.862976 | -1.8512337 | 0.00032519 | 0.01085569 | 0.0099361 | 319 | [illegible] | [illegible] |
| GOBP_POSITIVE_REGULATION_OF_SKELETAL_MUSCLE_FIBER_DEVELOP | GOBP_POSITIVE_REGULATION_OF_SKELETAL_MUSCLE_FIBER_DEVEL | 5 | -0.980462 | -1.6649909 | 0.00033277 | 0.0110594 | 0.01012255 | 166 | [illegible] | [illegible] |
| GOBP_POSITIVE_REGULATION_OF_TRANSPORTER_ACTIVITY | GOBP_POSITIVE_REGULATION_OF_TRANSPORTER_ACTIVITY | 29 | -0.7800385 | -1.8489725 | 0.00033693 | 0.01114837 | 0.01020398 | 469 | [illegible] | [illegible] |
| GOBP_POSITIVE_REGULATION_OF_MUSCLE_CONTRACTION | GOBP_POSITIVE_REGULATION_OF_MUSCLE_CONTRACTION | 35 | -0.7440056 | -1.8423443 | 0.00035794 | 0.01179159 | 0.01079271 | 408 | [illegible] | [illegible] |
| GOBP_CARDIAC_MUSCLE_CELL_ACTION_POTENTIAL_INVOLVED_IN_CO | GOBP_CARDIAC_MUSCLE_CELL_ACTION_POTENTIAL_INVOLVED_IN_ | 47 | -0.697681 | -1.8280999 | 0.00036175 | 0.01186508 | 0.01085998 | 405 | [illegible] | [illegible] |
| GOBP_PROTEIN_LOCALIZATION_TO_CHROMOSOME_CENTROMERIC_RE | GOBP_PROTEIN_LOCALIZATION_TO_CHROMOSOME_CENTROMERIC_ | 29 | 0.69956 | 1.70209995 | 0.00036452 | 0.01188089 | 0.01087445 | 1413 | [illegible] | [illegible] |
| GOBP_MITOTIC_SPINDLE_ASSEMBLY | GOBP_MITOTIC_SPINDLE_ASSEMBLY | 73 | 0.57697639 | 1.68469978 | 0.0003654 | 0.01188089 | 0.01087445 | 3460 | [illegible] | [illegible] |
| GOBP_MICROTUBULE_CYTOSKELETON_ORGANIZATION | GOBP_MICROTUBULE_CYTOSKELETON_ORGANIZATION | 578 | 0.37080945 | 1.40679633 | 0.00038412 | 0.01243578 | 0.01138233 | 2630 | [illegible] | [illegible] |
| GOBP_MYOBLAST_DIFFERENTIATION | GOBP_MYOBLAST_DIFFERENTIATION | 90 | -0.6176901 | -1.8150184 | 0.00039977 | 0.01284892 | 0.01176047 | 586 | [illegible] | [illegible] |
| GOBP_NEGATIVE_REGULATION_OF_SIGNAL_TRANSDUCTION_IN_ABSEN | GOBP_NEGATIVE_REGULATION_OF_SIGNAL_TRANSDUCTION_IN_AB | 27 | 0.71158323 | 1.7073983 | 0.0004003 | 0.01284892 | 0.01176047 | 753 | [illegible] | [illegible] |
| GOBP_RESPONSE_TO_TEMPERATURE_STIMULUS | GOBP_RESPONSE_TO_TEMPERATURE_STIMULUS | 147 | -0.549568 | -1.7383357 | 0.00040321 | 0.01288714 | 0.01179546 | 1220 | [illegible] | [illegible] |
| GOBP_MEMBRANE_DEPOLARIZATION | GOBP_MEMBRANE_DEPOLARIZATION | 68 | -0.6459108 | -1.8090075 | 0.0004111 | 0.01308368 | 0.01197535 | 405 | [illegible] | [illegible] |
| GOBP_REGULATION_OF_CHROMOSOME_SEGREGATION | GOBP_REGULATION_OF_CHROMOSOME_SEGREGATION | 133 | 0.51014917 | 1.63710242 | 0.00041528 | 0.01316116 | 0.01204627 | 2579 | [illegible] | [illegible] |
| GOBP_MONOCARBOXYLIC_ACID_METABOLIC_PROCESS | GOBP_MONOCARBOXYLIC_ACID_METABOLIC_PROCESS | 489 | -0.413002 | -1.4942971 | 0.00041991 | 0.01325173 | 0.01212917 | 1054 | [illegible] | [illegible] |
| GOBP_POSITIVE_REGULATION_OF_CYTOSKELETON_ORGANIZATION | GOBP_POSITIVE_REGULATION_OF_CYTOSKELETON_ORGANIZATION | 168 | -0.5239641 | -1.6886981 | 0.00043278 | 0.01360094 | 0.01244879 | 641 | [illegible] | [illegible] |
| GOBP_REGULATION_OF_CARDIAC_MUSCLE_CELL_ACTION_POTENTIAL_ | GOBP_REGULATION_OF_CARDIAC_MUSCLE_CELL_ACTION_POTENTI/ | 7 | -0.9594036 | -1.7262227 | 0.00044383 | 0.01389005 | 0.01271342 | 79 | [illegible] | [illegible] |
| GOBP_REGULATION_OF_CARDIAC_CONDUCTION | GOBP_REGULATION_OF_CARDIAC_CONDUCTION | 16 | -0.8653162 | -1.8181024 | 0.0004696 | 0.01463555 | 0.01339576 | 684 | [illegible] | [illegible] |
| GOBP_REGULATION_OF_SPINDLE_CHECKPOINT | GOBP_REGULATION_OF_SPINDLE_CHECKPOINT | 31 | 0.67624131 | 1.66723358 | 0.00047339 | 0.01469265 | 0.01344802 | 1697 | [illegible] | [illegible] |
| GOBP_D_GLUCOSE_IMPORT | GOBP_D_GLUCOSE_IMPORT | 59 | -0.6757139 | -1.8473595 | 0.00048496 | 0.01492827 | 0.01366368 | 1235 | [illegible] | [illegible] |
| GOBP_EPITHELIAL_CELL_DIFFERENTIATION | GOBP_EPITHELIAL_CELL_DIFFERENTIATION | 487 | 0.38042297 | 1.42500395 | 0.00048422 | 0.01492827 | 0.01366368 | 4256 | [illegible] | [illegible] |
| GOBP_VENTRICULAR_CARDIAC_MUSCLE_CELL_ACTION_POTENTIAL | GOBP_VENTRICULAR_CARDIAC_MUSCLE_CELL_ACTION_POTENTIAL | 32 | -0.7641386 | -1.8531409 | 0.00049944 | 0.01531126 | 0.01401423 | 756 | [illegible] | [illegible] |
| GOBP_CELLULAR_RESPONSE_TO_CAFFEINE | GOBP_CELLULAR_RESPONSE_TO_CAFFEINE | 10 | -0.929375 | -1.7835478 | 0.00051676 | 0.01577797 | 0.0144414 | 756 | [illegible] | [illegible] |
| GOBP_AXONEME_ASSEMBLY | GOBP_AXONEME_ASSEMBLY | 57 | 0.5961699 | 1.65820926 | 0.00054558 | 0.01659054 | 0.01518514 | 4878 | [illegible] | [illegible] |
| GOBP_MICROTUBULE_BUNDLE_FORMATION | GOBP_MICROTUBULE_BUNDLE_FORMATION | 85 | 0.54469982 | 1.63064255 | 0.00056177 | 0.01701388 | 0.01557262 | 4704 | [illegible] | [illegible] |
| GOBP_MICROTUBULE_ORGANIZING_CENTER_ORGANIZATION | GOBP_MICROTUBULE_ORGANIZING_CENTER_ORGANIZATION | 148 | 0.49138253 | 1.60049798 | 0.00056805 | 0.01706647 | 0.01562075 | 3483 | [illegible] | [illegible] |
| GOBP_NEGATIVE_REGULATION_OF_CELL_CYCLE | GOBP_NEGATIVE_REGULATION_OF_CELL_CYCLE | 352 | 0.40019488 | 1.45406195 | 0.00056652 | 0.01706647 | 0.01562075 | 2745 | [illegible] | [illegible] |
| GOBP_CELLULAR_RESPONSE_TO_PURINE_CONTAINING_COMPOUND | GOBP_CELLULAR_RESPONSE_TO_PURINE_CONTAINING_COMPOUNE | 12 | -0.9023865 | -1.7796767 | 0.00058254 | 0.01743223 | 0.01595553 | 756 | [illegible] | [illegible] |
| GOBP_REGULATION_OF_SISTER_CHROMATID_SEGREGATION | GOBP_REGULATION_OF_SISTER_CHROMATID_SEGREGATION | 109 | 0.52705601 | 1.64539454 | 0.0006068 | 0.01808613 | 0.01655404 | 2246 | [illegible] | [illegible] |
| GOBP_POLYSACCHARIDE_METABOLIC_PROCESS | GOBP_POLYSACCHARIDE_METABOLIC_PROCESS | 85 | -0.6018073 | -1.7489683 | 0.00063941 | 0.01898252 | 0.0173745 | 1052 | [illegible] | [illegible] |
| GOBP_NEGATIVE_REGULATION_OF_STRIATED_MUSCLE_CONTRACTION | GOBP_NEGATIVE_REGULATION_OF_STRIATED_MUSCLE_CONTRACTI | 7 | -0.9538716 | -1.7162692 | 0.00066033 | 0.0193848 | 0.0177427 | 160 | [illegible] | [illegible] |
| GOBP_REGULATION_OF_ACTIN_FILAMENT_ORGANIZATION | GOBP_REGULATION_OF_ACTIN_FILAMENT_ORGANIZATION | 232 | -0.4889163 | -1.6432902 | 0.00065723 | 0.0193848 | 0.0177427 | 641 | [illegible] | [illegible] |
| GOBP_MITOTIC_CELL_CYCLE_PHASE_TRANSITION | GOBP_MITOTIC_CELL_CYCLE_PHASE_TRANSITION | 423 | 0.38753941 | 1.43377027 | 0.0006607 | 0.0193848 | 0.0177427 | 2255 | [illegible] | [illegible] |
| GOBP_SPINDLE_ELONGATION | GOBP_SPINDLE_ELONGATION | 12 | 0.84051915 | 1.70728005 | 0.00067261 | 0.01965756 | 0.01799235 | 648 | [illegible] | [illegible] |
| GOBP_NEGATIVE_REGULATION_OF_POTASSIUM_ION_TRANSPORT | GOBP_NEGATIVE_REGULATION_OF_POTASSIUM_ION_TRANSPORT | 22 | -0.8136304 | -1.8179466 | 0.00067648 | 0.01968461 | 0.01801711 | 966 | [illegible] | [illegible] |
| GOBP_POSITIVE_REGULATION_OF_CELL_CYCLE | GOBP_POSITIVE_REGULATION_OF_CELL_CYCLE | 277 | 0.4236139 | 1.50045615 | 0.00067878 | 0.01968461 | 0.01801711 | 3805 | [illegible] | [illegible] |
| GOBP_NEGATIVE_REGULATION_OF_SUBSTRATE_ADHESION_DEPENDEN | GOBP_NEGATIVE_REGULATION_OF_SUBSTRATE_ADHESION_DEPENI | 13 | 0.81761791 | 1.68818182 | 0.00070098 | 0.0202501 | 0.0185347 | 1146 | [illegible] | [illegible] |
| GOBP_POSITIVE_REGULATION_OF_CELL_POPULATION_PROLIFERATION | GOBP_POSITIVE_REGULATION_OF_CELL_POPULATION_PROLIFERATI | 676 | 0.35294469 | 1.35666103 | 0.00071595 | 0.02060345 | 0.01885812 | 4208 | [illegible] | [illegible] |
| GOBP_CAMP_PKA_SIGNAL_TRANSDUCTION | GOBP_CAMP_PKA_SIGNAL_TRANSDUCTION | 51 | -0.6688123 | -1.7804885 | 0.00072246 | 0.0207115 | 0.01895701 | 908 | [illegible] | [illegible] |
| GOBP_DEFENSE_RESPONSE_TO_BACTERIUM | GOBP_DEFENSE_RESPONSE_TO_BACTERIUM | 165 | 0.47579325 | 1.57372299 | 0.00072584 | 0.02072917 | 0.01897318 | 3780 | [illegible] | [illegible] |
| GOBP_EPITHELIUM_DEVELOPMENT | GOBP_EPITHELIUM_DEVELOPMENT | 894 | 0.33889703 | 1.32936039 | 0.00073153 | 0.02081263 | 0.01904957 | 4238 | [illegible] | [illegible] |
| GOBP_POSITIVE_REGULATION_OF_CATION_CHANNEL_ACTIVITY | GOBP_POSITIVE_REGULATION_OF_CATION_CHANNEL_ACTIVITY | 17 | -0.8542389 | -1.8168479 | 0.00074451 | 0.02102258 | 0.01924174 | 469 | [illegible] | [illegible] |
| GOBP_IMMUNE_EFFECTOR_PROCESS | GOBP_IMMUNE_EFFECTOR_PROCESS | 477 | 0.37348887 | 1.39475427 | 0.00074232 | 0.02102258 | 0.01924174 | 4406 | [illegible] | [illegible] |
| GOBP_MAINTENANCE_OF_ORGANELLE_LOCATION | GOBP_MAINTENANCE_OF_ORGANELLE_LOCATION | 11 | -0.9024553 | -1.7447358 | 0.00076213 | 0.02138286 | 0.0195715 | 7 | [illegible] | [illegible] |
| GOBP_SKELETAL_MUSCLE_THIN_FILAMENT_ASSEMBLY | GOBP_SKELETAL_MUSCLE_THIN_FILAMENT_ASSEMBLY | 5 | -0.9728721 | -1.652102 | 0.00076296 | 0.02138286 | 0.0195715 | 185 | [illegible] | [illegible] |
| GOBP_POSITIVE_REGULATION_OF_BLOOD_CIRCULATION | GOBP_POSITIVE_REGULATION_OF_BLOOD_CIRCULATION | 27 | -0.7904354 | -1.8437904 | 0.00077491 | 0.02163697 | 0.01980408 | 716 | [illegible] | [illegible] |
| GOBP_CELL_PROJECTION_ASSEMBLY | GOBP_CELL_PROJECTION_ASSEMBLY | 547 | 0.36689008 | 1.38527993 | 0.00079314 | 0.02206395 | 0.02019489 | 3794 | [illegible] | [illegible] |
| GOBP_REGULATION_OF_CAMP_PKA_SIGNAL_TRANSDUCTION | GOBP_REGULATION_OF_CAMP_PKA_SIGNAL_TRANSDUCTION | 40 | -0.7238034 | -1.8385452 | 0.00081794 | 0.02266983 | 0.02074945 | 908 | [illegible] | [illegible] |
| GOBP_SKELETAL_MUSCLE_CELL_DIFFERENTIATION | GOBP_SKELETAL_MUSCLE_CELL_DIFFERENTIATION | 65 | -0.6359175 | -1.7651343 | 0.00082928 | 0.02281582 | 0.02088307 | 956 | [illegible] | [illegible] |
| GOBP_ESTABLISHMENT_OF_MITOTIC_SPINDLE_LOCALIZATION | GOBP_ESTABLISHMENT_OF_MITOTIC_SPINDLE_LOCALIZATION | 37 | 0.65431254 | 1.66787809 | 0.00082737 | 0.02281582 | 0.02088307 | 2592 | [illegible] | [illegible] |
| GOBP_REGULATION_OF_SODIUM_ION_TRANSPORT | GOBP_REGULATION_OF_SODIUM_ION_TRANSPORT | 57 | -0.6578255 | -1.784008 | 0.00083588 | 0.02291338 | 0.02097236 | 394 | [illegible] | [illegible] |
| GOBP_SUBSTRATE_ADHESION_DEPENDENT_CELL_SPREADING | GOBP_SUBSTRATE_ADHESION_DEPENDENT_CELL_SPREADING | 94 | 0.52949668 | 1.61349863 | 0.0008511 | 0.0232459 | 0.02127672 | 3300 | [illegible] | [illegible] |
| GOBP_TYPE_2_MITOPHAGY | GOBP_TYPE_2_MITOPHAGY | 15 | -0.8617515 | -1.7846396 | 0.00085435 | 0.02325007 | 0.02128054 | 1726 | [illegible] | [illegible] |
| GOBP_REGULATION_OF_POTASSIUM_ION_TRANSPORT | GOBP_REGULATION_OF_POTASSIUM_ION_TRANSPORT | 54 | -0.6702223 | -1.8021434 | 0.00085797 | 0.02326441 | 0.02129366 | 1035 | [illegible] | [illegible] |
| GOBP_MITOTIC_SPINDLE_ELONGATION | GOBP_MITOTIC_SPINDLE_ELONGATION | 10 | 0.84897203 | 1.6537034 | 0.00086388 | 0.02334022 | 0.02136305 | 648 | [illegible] | [illegible] |
| GOBP_NEGATIVE_REGULATION_OF_GENE_EXPRESSION | GOBP_NEGATIVE_REGULATION_OF_GENE_EXPRESSION | 1007 | 0.3281961 | 1.29549466 | 0.00088448 | 0.02381127 | 0.02179419 | 4413 | [illegible] | [illegible] |
| GOBP_GLYCOLYTIC_PROCESS_THROUGH_FRUCTOSE_6_PHOSPHATE | GOBP_GLYCOLYTIC_PROCESS_THROUGH_FRUCTOSE_6_PHOSPHATE | 22 | -0.8077122 | -1.8047231 | 0.00089588 | 0.02403189 | 0.02199613 | 765 | [illegible] | [illegible] |
| GOBP_CARDIAC_MUSCLE_CELL_MEMBRANE_REPOLARIZATION | GOBP_CARDIAC_MUSCLE_CELL_MEMBRANE_REPOLARIZATION | 31 | -0.7547797 | -1.8134309 | 0.00090919 | 0.02430223 | 0.02224357 | 756 | [illegible] | [illegible] |
| GOBP_REGULATION_OF_ANATOMICAL_STRUCTURE_SIZE | GOBP_REGULATION_OF_ANATOMICAL_STRUCTURE_SIZE | 398 | -0.4170389 | -1.4799829 | 0.0009406 | 0.02505263 | 0.0229304 | 431 | [illegible] | [illegible] |
| GOBP_SKELETAL_MYOFIBRIL_ASSEMBLY | GOBP_SKELETAL_MYOFIBRIL_ASSEMBLY | 8 | -0.9504176 | -1.756564 | 0.00096374 | 0.0255499 | 0.02338554 | 481 | [illegible] | [illegible] |
| GOBP_MONOSACCHARIDE_METABOLIC_PROCESS | GOBP_MONOSACCHARIDE_METABOLIC_PROCESS | 220 | -0.4848201 | -1.6158195 | 0.00096607 | 0.0255499 | 0.02338554 | 1050 | [illegible] | [illegible] |
| GOBP_EMBRYO_DEVELOPMENT | GOBP_EMBRYO_DEVELOPMENT | 940 | 0.32659961 | 1.2838734 | 0.00097105 | 0.02559133 | 0.02342347 | 4201 | [illegible] | [illegible] |

| | | | | | | | | | | |
|---|---|---|---|---|---|---|---|---|---|---|
| GOBP_GLYCOPROTEIN_METABOLIC_PROCESS | GOBP_GLYCOPROTEIN_METABOLIC_PROCESS | 315 | 0.40141399 | 1.44424418 | 0.00099624 | 0.02616359 | 0.02394725 | 4602 | tags=43%, list=31%, signal= | CHST14/GAL3ST4/SRD5A3/PHLDA [illegible] |
| GOBP_REGULATION_OF_MYOBLAST_DIFFERENTIATION | GOBP_REGULATION_OF_MYOBLAST_DIFFERENTIATION | 64 | -0.6386369 | -1.77119 | 0.00104522 | 0.02724735 | 0.0249392 | 586 | tags=20%, list=4%, signal=2 | HIF1AN/MBNL3/RBM24/DPF3/M [illegible] |
| GOBP_RESPONSE_TO_ISCHEMIA | GOBP_RESPONSE_TO_ISCHEMIA | 51 | -0.6592757 | -1.7551005 | 0.00104355 | 0.02724735 | 0.0249392 | 1398 | tags=25%, list=10%, signal= | EEF2K/ROCK2/CPEB4/SQSTM1/EG [illegible] |
| GOBP_MITOCHONDRIAL_RESPIRATORY_CHAIN_COMPLEX_ASSEMBLY | GOBP_MITOCHONDRIAL_RESPIRATORY_CHAIN_COMPLEX_ASSEMBL | 93 | -0.5839133 | -1.7219942 | 0.00105159 | 0.02724735 | 0.0249392 | 1131 | tags=20%, list=8%, signal=1 | NDUFS1/TMEM186/NDUFAF3/PE [illegible] |
| GOBP_AMINOGLYCAN_METABOLIC_PROCESS | GOBP_AMINOGLYCAN_METABOLIC_PROCESS | 58 | 0.59010835 | 1.64594762 | 0.00105202 | 0.02724735 | 0.0249392 | 3437 | tags=53%, list=23%, signal= | EXT1/B3GNT9/CTBS/TNFAIP6/HE [illegible] |
| GOBP_POSITIVE_REGULATION_OF_METAPHASE_ANAPHASE_TRANSITIO | GOBP_POSITIVE_REGULATION_OF_METAPHASE_ANAPHASE_TRANSI | 17 | 0.76069799 | 1.66253706 | 0.00106277 | 0.02743115 | 0.02510743 | 2002 | tags=53%, list=14%, signal= | PLK1/MAD2L1BP/UBE2C/SKA1/D [illegible] |
| GOBP_REGULATION_OF_SPINDLE_ORGANIZATION | GOBP_REGULATION_OF_SPINDLE_ORGANIZATION | 46 | 0.61752901 | 1.64195123 | 0.00106767 | 0.02746322 | 0.02513679 | 2419 | tags=37%, list=17%, signal= | PLK1/TPX2/MAP9/STIL/TACC3/SP [illegible] |
| GOBP_RESPONSE_TO_METAL_ION | GOBP_RESPONSE_TO_METAL_ION | 242 | -0.4594169 | -1.549224 | 0.00108516 | 0.02781779 | 0.02546132 | 756 | tags=10%, list=5%, signal=1 | CASQ2/FOS/GSK3A/SMPD1/RYR3/ [illegible] |
| GOBP_MITOCHONDRIAL_ELECTRON_TRANSPORT_CYTOCHROME_C_TO | GOBP_MITOCHONDRIAL_ELECTRON_TRANSPORT_CYTOCHROME_C_ | 15 | -0.8576417 | -1.7761284 | 0.00108913 | 0.02782465 | 0.0254676 | 998 | tags=53%, list=7%, signal=5 | COX8A/COX6B1/COX5A/COX4I1/ [illegible] |
| GOBP_CARDIAC_CHAMBER_DEVELOPMENT | GOBP_CARDIAC_CHAMBER_DEVELOPMENT | 150 | -0.5242972 | -1.6639948 | 0.00110836 | 0.02821987 | 0.02582934 | 400 | tags=11%, list=3%, signal=1 | EGLN1/GREB1L/GSK3A/XIRP2/TB [illegible] |
| GOBP_NUCLEOSIDE_PHOSPHATE_BIOSYNTHETIC_PROCESS | GOBP_NUCLEOSIDE_PHOSPHATE_BIOSYNTHETIC_PROCESS | 279 | -0.4498837 | -1.541244 | 0.00113951 | 0.02891502 | 0.0264656 | 1418 | tags=23%, list=10%, signal= | NDUFA7/TGFB1/HACD1/ATP5PO/ [illegible] |
| GOBP_ACTIN_FILAMENT_DEPOLYMERIZATION | GOBP_ACTIN_FILAMENT_DEPOLYMERIZATION | 52 | -0.6686423 | -1.786257 | 0.0011482 | 0.02903751 | 0.02657772 | 431 | tags=21%, list=3%, signal=2 | CARMIL2/LMOD1/CFL2/TMOD1/ [illegible] |
| GOBP_NUCLEAR_TRANSCRIBED_MRNA_CATABOLIC_PROCESS | GOBP_NUCLEAR_TRANSCRIBED_MRNA_CATABOLIC_PROCESS | 141 | 0.48386273 | 1.56555076 | 0.00115914 | 0.02921574 | 0.02674085 | 2415 | tags=23%, list=16%, signal= | SNRPF/CNOT3/TENT4A/ERI1/MA [illegible] |
| GOBP_REGULATION_OF_PROTEIN_DEPOLYMERIZATION | GOBP_REGULATION_OF_PROTEIN_DEPOLYMERIZATION | 85 | -0.5888314 | -1.7112577 | 0.00118394 | 0.029741 | 0.02722161 | 431 | tags=15%, list=3%, signal=1 | CARMIL2/LMOD1/CFL2/TMOD1/ [illegible] |
| GOBP_ORGANIC_ACID_METABOLIC_PROCESS | GOBP_ORGANIC_ACID_METABOLIC_PROCESS | 699 | -0.3739611 | -1.3950783 | 0.00122299 | 0.03061967 | 0.02802585 | 1054 | tags=13%, list=7%, signal=1 | ACOT9/AKT2/AKR1B1/DCXR/ABC [illegible] |
| GOBP_REGULATION_OF_RESPONSE_TO_STRESS | GOBP_REGULATION_OF_RESPONSE_TO_STRESS | 1200 | 0.31756692 | 1.26607144 | 0.00125405 | 0.03129291 | 0.02864206 | 4345 | tags=31%, list=30%, signal= | PLAUR/HMGB2/PLK1/IRAK4/TRA [illegible] |
| GOBP_MITOTIC_CELL_CYCLE_CHECKPOINT_SIGNALING | GOBP_MITOTIC_CELL_CYCLE_CHECKPOINT_SIGNALING | 140 | 0.48613901 | 1.57293813 | 0.00125965 | 0.03132847 | 0.02867461 | 2246 | tags=28%, list=15%, signal= | PLK1/CDCA8/CDK1/NUF2/RBBP8/ [illegible] |
| GOBP_RESPONSE_TO_HEAT | GOBP_RESPONSE_TO_HEAT | 91 | -0.5809633 | -1.7082028 | 0.00127527 | 0.03161227 | 0.02893437 | 1099 | tags=19%, list=8%, signal=1 | FGF1/IER5/MAPKAPK2/HSPA2/EI [illegible] |
| GOBP_POSITIVE_REGULATION_OF_MYOBLAST_DIFFERENTIATION | GOBP_POSITIVE_REGULATION_OF_MYOBLAST_DIFFERENTIATION | 41 | -0.7009078 | -1.7889079 | 0.00128625 | 0.0317796 | 0.02908752 | 586 | tags=22%, list=4%, signal=2 | HIF1AN/RBM24/DPF3/MEF2C/CS [illegible] |
| GOBP_POSITIVE_REGULATION_OF_CHROMOSOME_SEGREGATION | GOBP_POSITIVE_REGULATION_OF_CHROMOSOME_SEGREGATION | 29 | 0.67468121 | 1.64156733 | 0.00129128 | 0.03179934 | 0.02910559 | 2579 | tags=55%, list=18%, signal= | RCC2/CDCA8/CDK1/MAD2L1BP/ [illegible] |
| GOBP_GLYCOLYTIC_PROCESS_THROUGH_GLUCOSE_6_PHOSPHATE | GOBP_GLYCOLYTIC_PROCESS_THROUGH_GLUCOSE_6_PHOSPHATE | 20 | -0.8161654 | -1.7902957 | 0.00130784 | 0.03200528 | 0.02929408 | 391 | tags=40%, list=3%, signal=3 | HK1/GALK1/TPI1/PFKM/GAPDH/ [illegible] |
| GOBP_CALCIUM_ION_IMPORT_INTO_SARCOPLASMIC_RETICULUM | GOBP_CALCIUM_ION_IMPORT_INTO_SARCOPLASMIC_RETICULUM | 6 | -0.9687736 | -1.6984797 | 0.00130816 | 0.03200528 | 0.02929408 | 427 | tags=83%, list=3%, signal=8 | PLN/SLN/ATP2A2/STRIT1/ATP2A1 |
| GOBP_HOMEOSTATIC_PROCESS | GOBP_HOMEOSTATIC_PROCESS | 1369 | -0.3314675 | -1.2913093 | 0.00131537 | 0.03207717 | 0.02935988 | 971 | tags=8%, list=7%, signal=89 | CNBP/OXSR1/ATP6V0E2/HK1/GR [illegible] |
| GOBP_MITOCHONDRIAL_ELECTRON_TRANSPORT_UBIQUINOL_TO_CYT | GOBP_MITOCHONDRIAL_ELECTRON_TRANSPORT_UBIQUINOL_TO_C | 12 | -0.8860428 | -1.747444 | 0.00132148 | 0.03212168 | 0.02940063 | 1195 | tags=75%, list=8%, signal=6 | UQCRB/UQCRC2/CYC1/UQCR11/ [illegible] |
| GOBP_RESPONSE_TO_CAFFEINE | GOBP_RESPONSE_TO_CAFFEINE | 14 | -0.8582504 | -1.7535382 | 0.00134235 | 0.03231534 | 0.02957788 | 756 | tags=36%, list=5%, signal=3 | CASQ2/RYR3/TMEM38A/CACNA1 [illegible] |
| GOBP_REGULATION_OF_UBIQUITIN_PROTEIN_LIGASE_ACTIVITY | GOBP_REGULATION_OF_UBIQUITIN_PROTEIN_LIGASE_ACTIVITY | 10 | 0.83683277 | 1.63005746 | 0.00134002 | 0.03231534 | 0.02957788 | 2002 | tags=70%, list=14%, signal= | UBE2C/CDKN2A/MAD2L1/CDC14 [illegible] |
| GOBP_REGULATION_OF_RESPONSE_TO_BIOTIC_STIMULUS | GOBP_REGULATION_OF_RESPONSE_TO_BIOTIC_STIMULUS | 461 | 0.36968204 | 1.37734383 | 0.00133657 | 0.03231534 | 0.02957788 | 4657 | tags=34%, list=32%, signal= | HMGB2/IRAK4/TRAFD1/SLC19A1/ [illegible] |
| GOBP_POSITIVE_REGULATION_OF_CHEMOKINE_PRODUCTION | GOBP_POSITIVE_REGULATION_OF_CHEMOKINE_PRODUCTION | 51 | 0.59226561 | 1.60746528 | 0.00139533 | 0.03337694 | 0.03054955 | 4259 | tags=55%, list=29%, signal= | MYD88/APP/RIPK2/ADAM17/CS/ [illegible] |
| GOBP_DNA_REPLICATION | GOBP_DNA_REPLICATION | 298 | 0.40948228 | 1.46479436 | 0.00139428 | 0.03337694 | 0.03054955 | 3788 | tags=31%, list=26%, signal= | LIG1/MCM2/CDK1/TYMS/RBBP8/ [illegible] |
| GOBP_POSITIVE_REGULATION_OF_ACTIN_FILAMENT_BUNDLE_ASSEMB | GOBP_POSITIVE_REGULATION_OF_ACTIN_FILAMENT_BUNDLE_ASSE | 56 | -0.6346023 | -1.7182297 | 0.0014212 | 0.03388778 | 0.03101712 | 630 | tags=14%, list=4%, signal=1 | ARHGEF5/FHOD1/RAPGEF3/LIM [illegible] |
| GOBP_NADH_DEHYDROGENASE_COMPLEX_ASSEMBLY | GOBP_NADH_DEHYDROGENASE_COMPLEX_ASSEMBLY | 35 | -0.7110078 | -1.7606334 | 0.00144593 | 0.03436836 | 0.03145698 | 1447 | tags=34%, list=10%, signal= | NDUFS1/TMEM186/NDUFAF3/N [illegible] |
| GOBP_REGULATION_OF_ACTIN_FILAMENT_LENGTH | GOBP_REGULATION_OF_ACTIN_FILAMENT_LENGTH | 133 | -0.5272599 | -1.6450091 | 0.00146115 | 0.03462059 | 0.03168785 | 431 | tags=11%, list=3%, signal=1 | CCR7/CARMIL2/LMOD1/CFL2/TM [illegible] |
| GOBP_CAVEOLIN_MEDIATED_ENDOCYTOSIS | GOBP_CAVEOLIN_MEDIATED_ENDOCYTOSIS | 10 | -0.9108596 | -1.7480153 | 0.00152711 | 0.03606955 | 0.03301407 | 36 | tags=20%, list=0%, signal=2 | MAPK3/CAV3 |
| GOBP_NEGATIVE_REGULATION_OF_NUCLEOTIDE_CATABOLIC_PROCESS | GOBP_NEGATIVE_REGULATION_OF_NUCLEOTIDE_CATABOLIC_PROC | 17 | -0.8322992 | -1.7701851 | 0.00154753 | 0.03632337 | 0.03324639 | 836 | tags=29%, list=6%, signal=2 | SIRT6/PPARA/HDAC4/PRKACA/T [illegible] |
| GOBP_TRANSMISSION_OF_NERVE_IMPULSE | GOBP_TRANSMISSION_OF_NERVE_IMPULSE | 43 | -0.6817539 | -1.756125 | 0.00154379 | 0.03632337 | 0.03324639 | 304 | tags=12%, list=2%, signal=1 | CHRNA1/SCN1B/PLEC/ATP1A2/C [illegible] |
| GOBP_ESTABLISHMENT_OF_GOLGI_LOCALIZATION | GOBP_ESTABLISHMENT_OF_GOLGI_LOCALIZATION | 8 | 0.86634219 | 1.60225595 | 0.00155445 | 0.03637229 | 0.03329116 | 1456 | tags=50%, list=10%, signal= | CDC42/YWHAZ/COPG1/NHERF1 |
| GOBP_PROTEIN_LOCALIZATION_TO_CHROMOSOME | GOBP_PROTEIN_LOCALIZATION_TO_CHROMOSOME | 100 | 0.51978486 | 1.59985568 | 0.00159041 | 0.03709815 | 0.03395553 | 4269 | tags=37%, list=29%, signal= | PLK1/RCC2/H2AX/CDK1/AURKB/ [illegible] |
| GOBP_NEGATIVE_REGULATION_OF_MONOATOMIC_ION_TRANSPORT | GOBP_NEGATIVE_REGULATION_OF_MONOATOMIC_ION_TRANSPOR | 74 | -0.6088948 | -1.7268024 | 0.0016401 | 0.03813859 | 0.03490784 | 1463 | tags=28%, list=10%, signal= | VDAC1/PPP3CC/ADRA2A/TGFB1/ [illegible] |
| GOBP_REGULATION_OF_ATTACHMENT_OF_SPINDLE_MICROTUBULES_T | GOBP_REGULATION_OF_ATTACHMENT_OF_SPINDLE_MICROTUBULE | 23 | 0.70624774 | 1.63764374 | 0.00167257 | 0.03877377 | 0.03548921 | 386 | tags=39%, list=3%, signal=3 | RCC2/CDCA8/CDK1/AURKB/ZWIL [illegible] |
| GOBP_REGULATION_OF_CHROMOSOME_ORGANIZATION | GOBP_REGULATION_OF_CHROMOSOME_ORGANIZATION | 239 | 0.41621513 | 1.4502154 | 0.00169156 | 0.03909332 | 0.03578169 | 3277 | tags=28%, list=22%, signal= | PLK1/HECW2/CDCA8/CDK1/NUF2/ [illegible] |
| GOBP_POSITIVE_REGULATION_OF_MITOTIC_NUCLEAR_DIVISION | GOBP_POSITIVE_REGULATION_OF_MITOTIC_NUCLEAR_DIVISION | 34 | 0.64949703 | 1.62711611 | 0.00171869 | 0.0395232 | 0.03617515 | 2002 | tags=38%, list=14%, signal= | PLK1/MAD2L1BP/UBE2C/SKA1/N [illegible] |
| GOBP_SIGNAL_TRANSDUCTION_IN_RESPONSE_TO_DNA_DAMAGE | GOBP_SIGNAL_TRANSDUCTION_IN_RESPONSE_TO_DNA_DAMAGE | 194 | 0.4430604 | 1.49978874 | 0.00172069 | 0.0395232 | 0.03617515 | 3788 | tags=35%, list=26%, signal= | PLK1/H2AX/CDK1/SNAI2/CDKN2A [illegible] |
| GOBP_NEGATIVE_REGULATION_OF_MUSCLE_CONTRACTION | GOBP_NEGATIVE_REGULATION_OF_MUSCLE_CONTRACTION | 20 | -0.805839 | -1.7676442 | 0.00185587 | 0.04249836 | 0.03889829 | 160 | tags=25%, list=1%, signal=2 | STUB1/BIN1/ATP1A2/TNNT1/AT [illegible] |
| GOBP_REGULATION_OF_PROTEIN_CONTAINING_COMPLEX_DISASSEMB | GOBP_REGULATION_OF_PROTEIN_CONTAINING_COMPLEX_DISASSE | 122 | -0.5296403 | -1.6327408 | 0.00196516 | 0.04486411 | 0.04106364 | 1261 | tags=20%, list=9%, signal=1 | GSN/MAP6D1/EPS8/PLEKHH2/TE [illegible] |
| GOBP_REGULATION_OF_DEFENSE_RESPONSE | GOBP_REGULATION_OF_DEFENSE_RESPONSE | 669 | 0.34443431 | 1.32219624 | 0.001974 | 0.04492946 | 0.04112345 | 4453 | tags=33%, list=30%, signal= | HMGB2/IRAK4/TRAFD1/GBP1/SL [illegible] |
| GOBP_LYMPHOCYTE_ACTIVATION | GOBP_LYMPHOCYTE_ACTIVATION | 650 | 0.34314029 | 1.31356353 | 0.00199255 | 0.04521453 | 0.04138437 | 4406 | tags=33%, list=30%, signal= | EPHB2/CMTM7/TWSG1/CDKN2A/ [illegible] |
| GOBP_DETECTION_OF_MECHANICAL_STIMULUS | GOBP_DETECTION_OF_MECHANICAL_STIMULUS | 39 | -0.6979458 | -1.7639703 | 0.00204904 | 0.04635647 | 0.04242958 | 190 | tags=18%, list=1%, signal=1 | PKD1L3/PKD1/CSRP3/HPN/TTN/T [illegible] |
| GOBP_NEGATIVE_REGULATION_OF_POTASSIUM_ION_TRANSMEMBRA | GOBP_NEGATIVE_REGULATION_OF_POTASSIUM_ION_TRANSMEMB | 15 | -0.8445838 | -1.7490863 | 0.00205794 | 0.04641791 | 0.04248581 | 160 | tags=20%, list=1%, signal=2 | OXSR1/BIN1/CAV3 |
| GOBP_NEGATIVE_REGULATION_OF_PROTEIN_CONTAINING_COMPLEX_ | GOBP_NEGATIVE_REGULATION_OF_PROTEIN_CONTAINING_COMPL | 76 | -0.6034597 | -1.7192334 | 0.00207006 | 0.04655158 | 0.04260815 | 431 | tags=14%, list=3%, signal=1 | CARMIL2/LMOD1/TMOD1/SVIL/ [illegible] |
| GOBP_RESPONSE_TO_STIMULUS_INVOLVED_IN_REGULATION_OF_MUS | GOBP_RESPONSE_TO_STIMULUS_INVOLVED_IN_REGULATION_OF_M | 9 | -0.9185417 | -1.7292688 | 0.0020837 | 0.04662034 | 0.0426711 | 836 | tags=56%, list=6%, signal=5 | HDAC4/DAG1/FBXO32/CASQ1/T [illegible] |
| GOBP_PERICYTE_CELL_DIFFERENTIATION | GOBP_PERICYTE_CELL_DIFFERENTIATION | 10 | 0.82860489 | 1.61403045 | 0.00209174 | 0.04662034 | 0.0426711 | 1497 | tags=30%, list=10%, signal= | EPHA2/NOTCH1/PDGFB |
| GOBP_PROTEIN_DNA_COVALENT_CROSS_LINKING_REPAIR | GOBP_PROTEIN_DNA_COVALENT_CROSS_LINKING_REPAIR | 10 | 0.82811701 | 1.61308013 | 0.00209174 | 0.04662034 | 0.0426711 | 1167 | tags=40%, list=8%, signal=3 | FAM111B/BRIP1/FAM111A/TRAI [illegible] |
| GOBP_REGULATION_OF_SKELETAL_MUSCLE_FIBER_DEVELOPMENT | GOBP_REGULATION_OF_SKELETAL_MUSCLE_FIBER_DEVELOPMENT | 8 | -0.9357319 | -1.7294218 | 0.00216655 | 0.04814494 | 0.04406654 | 166 | tags=38%, list=1%, signal=3 | MYOD1/MYF6/LMOD3 |
| GOBP_POLYSACCHARIDE_CATABOLIC_PROCESS | GOBP_POLYSACCHARIDE_CATABOLIC_PROCESS | 28 | -0.742167 | -1.7466807 | 0.00217979 | 0.0482961 | 0.0442049 | 616 | tags=25%, list=4%, signal=2 | PPP1R3B/GABARAPL1/PGM1/AG [illegible] |
| GOBP_POSITIVE_REGULATION_OF_PROTEIN_POLYMERIZATION | GOBP_POSITIVE_REGULATION_OF_PROTEIN_POLYMERIZATION | 80 | -0.5794546 | -1.6676825 | 0.00219364 | 0.0484601 | 0.044355 | 641 | tags=12%, list=4%, signal=1 | CCR7/ARL2/CARMIL2/LMOD1/PI [illegible] |
| GOBP_REGULATION_OF_CYTOSOLIC_CALCIUM_ION_CONCENTRATION | GOBP_REGULATION_OF_CYTOSOLIC_CALCIUM_ION_CONCENTRATIO | 32 | -0.7263257 | -1.7614393 | 0.00235969 | 0.0516725 | 0.04729528 | 427 | tags=16%, list=3%, signal=1 | PLN/MYH7B/HRC/CAV3/RYR1 |
| GOBP_POSITIVE_REGULATION_OF_ATP_DEPENDENT_ACTIVITY | GOBP_POSITIVE_REGULATION_OF_ATP_DEPENDENT_ACTIVITY | 19 | -0.7987004 | -1.7307607 | 0.00235479 | 0.0516725 | 0.04729528 | 9 | tags=16%, list=0%, signal=1 | SETMAR/TNNT3/ATP2A1 |
| GOBP_SEXUAL_REPRODUCTION | GOBP_SEXUAL_REPRODUCTION | 774 | 0.33476902 | 1.29921686 | 0.00235305 | 0.0516725 | 0.04729528 | 3952 | tags=34%, list=27%, signal= | HMGB2/PLK1/H2AX/CDK1/ODF2/ [illegible] |
| GOBP_RESPONSE_TO_INACTIVITY | GOBP_RESPONSE_TO_INACTIVITY | 7 | -0.9374699 | -1.6867581 | 0.00237595 | 0.0518566 | 0.04746378 | 836 | tags=57%, list=6%, signal=5 | HDAC4/DAG1/FBXO32/CASQ1 |
| GOBP_CARBOHYDRATE_DERIVATIVE_CATABOLIC_PROCESS | GOBP_CARBOHYDRATE_DERIVATIVE_CATABOLIC_PROCESS | 216 | -0.4672485 | -1.5559631 | 0.00238191 | 0.0518566 | 0.04746378 | 865 | tags=11%, list=6%, signal=1 | PDE7A/PDE5A/HDAC4/NEU3/GA [illegible] |
| GOBP_ENUCLEATE_ERYTHROCYTE_MATURATION | GOBP_ENUCLEATE_ERYTHROCYTE_MATURATION | 5 | 0.92721293 | 1.5564958 | 0.00245342 | 0.05325898 | 0.04874736 | 887 | tags=60%, list=6%, signal=5 | NEMP1/DIAPH3/RAC2 |
| GOBP_REGULATION_OF_IMMUNE_RESPONSE | GOBP_REGULATION_OF_IMMUNE_RESPONSE | 737 | 0.33443025 | 1.29312526 | 0.00248131 | 0.05370929 | 0.04915953 | 4257 | tags=30%, list=29%, signal= | HMGB2/IRAK4/TRAFD1/GBP1/SL [illegible] |
| GOBP_NEGATIVE_REGULATION_OF_MUSCLE_HYPERTROPHY | GOBP_NEGATIVE_REGULATION_OF_MUSCLE_HYPERTROPHY | 31 | -0.7238154 | -1.7390362 | 0.00256402 | 0.05521033 | 0.05053342 | 784 | tags=23%, list=5%, signal=2 | TNFRSF1A/GSK3A/STUB1/PI16/M [illegible] |
| GOBP_CELLULAR_RESPONSE_TO_VIRUS | GOBP_CELLULAR_RESPONSE_TO_VIRUS | 48 | 0.59174279 | 1.58942468 | 0.00256536 | 0.05521033 | 0.05053342 | 2658 | tags=38%, list=18%, signal= | IFNAR2/ADAR/EXT1/IFI6/GSDME/ [illegible] |
| GOBP_NEGATIVE_REGULATION_OF_CALCIUM_ION_TRANSPORT | GOBP_NEGATIVE_REGULATION_OF_CALCIUM_ION_TRANSPORT | 38 | -0.6912227 | -1.735645 | 0.00258128 | 0.05539429 | 0.05070179 | 461 | tags=42%, list=3%, signal=4 | VDAC1/PPP3CC/ADRA2A/TGFB1/ [illegible] |
| GOBP_NEGATIVE_REGULATION_OF_ACTIN_FILAMENT_DEPOLYMERIZAT | GOBP_NEGATIVE_REGULATION_OF_ACTIN_FILAMENT_DEPOLYMERI | 40 | -0.6882891 | -1.7483349 | 0.00268324 | 0.05741825 | 0.0525543 | 431 | tags=22%, list=3%, signal=2 | CARMIL2/LMOD1/TMOD1/SVIL/ [illegible] |
| GOBP_NEGATIVE_REGULATION_OF_MONOATOMIC_ION_TRANSMEMB | GOBP_NEGATIVE_REGULATION_OF_MONOATOMIC_ION_TRANSMEM | 53 | -0.653983 | -1.7523522 | 0.00269595 | 0.05752638 | 0.05265327 | 461 | tags=26%, list=3%, signal=2 | VDAC1/PPP3CC/TGFB1/CALM1/G [illegible] |
| GOBP_REGULATION_OF_CALCIUM_ION_IMPORT | GOBP_REGULATION_OF_CALCIUM_ION_IMPORT | 32 | -0.7208165 | -1.7480788 | 0.00272624 | 0.05800787 | 0.05309397 | 464 | tags=25%, list=3%, signal=2 | HOMER1/PPP3CB/PLN/UCN/P2R [illegible] |
| GOBP_CELL_PROJECTION_ORGANIZATION | GOBP_CELL_PROJECTION_ORGANIZATION | 1335 | 0.3079394 | 1.23473727 | 0.0027481 | 0.05830784 | 0.05336853 | 4280 | tags=32%, list=29%, signal= | PTK7/CRABP2/TRIOBP/RCC2/TB [illegible] |
| GOBP_REGULATION_OF_METAPHASE_PLATE_CONGRESSION | GOBP_REGULATION_OF_METAPHASE_PLATE_CONGRESSION | 15 | 0.75862556 | 1.6144042 | 0.00276583 | 0.05851866 | 0.05356149 | 1261 | tags=33%, list=9%, signal=3 | CDCA8/CDK1/AURKB/SPAG5/HN [illegible] |
| GOBP_RELAXATION_OF_CARDIAC_MUSCLE | GOBP_RELAXATION_OF_CARDIAC_MUSCLE | 17 | -0.8176606 | -1.7390509 | 0.00285329 | 0.06003103 | 0.05494575 | 696 | tags=35%, list=5%, signal=3 | ATP1B1/PLN/AKAP6/ATP1A2/AT [illegible] |
| GOBP_PROTEIN_LOCALIZATION_TO_CILIARY_TRANSITION_ZONE | GOBP_PROTEIN_LOCALIZATION_TO_CILIARY_TRANSITION_ZONE | 20 | 0.70970485 | 1.60311192 | 0.00284585 | 0.06003103 | 0.05494575 | 1794 | tags=40%, list=12%, signal= | TMEM67/MKS1/TMEM216/B9D2/ [illegible] |
| GOBP_REGULATION_OF_MITOTIC_CELL_CYCLE | GOBP_REGULATION_OF_MITOTIC_CELL_CYCLE | 471 | 0.36382306 | 1.35740504 | 0.00287115 | 0.06023803 | 0.05513521 | 2260 | tags=23%, list=15%, signal= | BTG3/PLK1/RCC2/HECW2/CDCA8 [illegible] |
| GOBP_LEUKOCYTE_MEDIATED_IMMUNITY | GOBP_LEUKOCYTE_MEDIATED_IMMUNITY | 292 | 0.39256667 | 1.39986584 | 0.00291486 | 0.06098473 | 0.05581866 | 4875 | tags=37%, list=33%, signal= | IRAK4/ARRB2/MYD88/TUBB/BTN [illegible] |
| GOBP_KINETOCHORE_ORGANIZATION | GOBP_KINETOCHORE_ORGANIZATION | 31 | 0.64413177 | 1.58806937 | 0.00298317 | 0.06224056 | 0.05696811 | 3666 | tags=61%, list=25%, signal= | CENPW/NUF2/CENPK/CENPE/HJ [illegible] |
| GOBP_REGULATION_OF_SUBSTRATE_ADHESION_DEPENDENT_CELL_SP | GOBP_REGULATION_OF_SUBSTRATE_ADHESION_DEPENDENT_CELL_ | 53 | 0.58164343 | 1.59091183 | 0.00300198 | 0.06228692 | 0.05701055 | 1189 | tags=23%, list=8%, signal=2 | GBP1/TRIOBP/RCC2/EFNA5/AP1A [illegible] |
| GOBP_REGULATION_OF_INNATE_IMMUNE_RESPONSE | GOBP_REGULATION_OF_INNATE_IMMUNE_RESPONSE | 387 | 0.37211078 | 1.36394137 | 0.00299645 | 0.06228692 | 0.05701055 | 4657 | tags=34%, list=32%, signal= | HMGB2/IRAK4/TRAFD1/SLC19A1/ [illegible] |
| GOBP_REGULATION_OF_POTASSIUM_ION_TRANSMEMBRANE_TRANSP | GOBP_REGULATION_OF_POTASSIUM_ION_TRANSMEMBRANE_TRAN | 42 | -0.673859 | -1.7287318 | 0.00310579 | 0.06426333 | 0.05881953 | 1178 | tags=19%, list=8%, signal=1 | KCNS3/ABCC9/OXSR1/ATP1B1/A [illegible] |
| GOBP_SENSORY_PERCEPTION_OF_LIGHT_STIMULUS | GOBP_SENSORY_PERCEPTION_OF_LIGHT_STIMULUS | 101 | 0.50496823 | 1.55812167 | 0.00315516 | 0.06510551 | 0.05959036 | 4186 | tags=40%, list=29%, signal= | RABGGTB/PDCL/OAT/EYA3/WHR [illegible] |
| GOBP_MONONUCLEAR_CELL_DIFFERENTIATION | GOBP_MONONUCLEAR_CELL_DIFFERENTIATION | 441 | 0.36266354 | 1.3451507 | 0.00323085 | 0.06648463 | 0.06085266 | 4196 | tags=33%, list=29%, signal= | CMTM7/CDKN2A/MYD88/NOTCH [illegible] |
| GOBP_REGULATION_OF_ACTOMYOSIN_STRUCTURE_ORGANIZATION | GOBP_REGULATION_OF_ACTOMYOSIN_STRUCTURE_ORGANIZATION | 88 | -0.5676018 | -1.6630746 | 0.00325358 | 0.06676958 | 0.06111348 | 630 | tags=14%, list=4%, signal=1 | AMOT/INPP5K/ARHGEF5/FHOD1/ [illegible] |
| GOBP_NEGATIVE_REGULATION_OF_NUCLEOTIDE_METABOLIC_PROCES | GOBP_NEGATIVE_REGULATION_OF_NUCLEOTIDE_METABOLIC_PRO | 23 | -0.7751447 | -1.7502845 | 0.00328491 | 0.06722871 | 0.06153371 | 836 | tags=26%, list=6%, signal=2 | PPARA/HDAC4/ATPSIF1/PRKACA/ [illegible] |
| GOBP_POSITIVE_REGULATION_OF_PROTEIN_DEUBIQUITINATION | GOBP_POSITIVE_REGULATION_OF_PROTEIN_DEUBIQUITINATION | 5 | -0.9569163 | -1.6250063 | 0.00331688 | 0.0676986 | 0.0619638 | 285 | tags=40%, list=2%, signal=3 | NOP53/TNIP1 |
| GOBP_REGULATION_OF_MITOTIC_CENTROSOME_SEPARATION | GOBP_REGULATION_OF_MITOTIC_CENTROSOME_SEPARATION | 8 | 0.85059647 | 1.57313503 | 0.00337739 | 0.06856094 | 0.06275308 | 386 | tags=38%, list=3%, signal=3 | KIF11/MAP9/NEK2 |
| GOBP_EPIDERMIS_DEVELOPMENT | GOBP_EPIDERMIS_DEVELOPMENT | 233 | 0.4178426 | 1.44951739 | 0.00336834 | 0.06856094 | 0.06275308 | 5149 | tags=51%, list=35%, signal= | CRABP2/TRIOBP/EXT1/CD109/W [illegible] |
| GOBP_G_PROTEIN_COUPLED_RECEPTOR_INTERNALIZATION | GOBP_G_PROTEIN_COUPLED_RECEPTOR_INTERNALIZATION | 9 | 0.8270084 | 1.5709547 | 0.00339684 | 0.06876998 | 0.06294441 | 2076 | tags=67%, list=14%, signal= | ARRB2/APP/GTF2H2/UBQLN2/AP [illegible] |
| GOBP_REGULATION_OF_MEMBRANE_DEPOLARIZATION | GOBP_REGULATION_OF_MEMBRANE_DEPOLARIZATION | 41 | -0.6742741 | -1.7209313 | 0.00342482 | 0.06880915 | 0.06298027 | 394 | tags=12%, list=3%, signal=1 | SCN3B/GOT1/SCN1B/CAV3/FHL1 |
| GOBP_PYRIMIDINE_NUCLEOSIDE_TRIPHOSPHATE_BIOSYNTHETIC_PRO | GOBP_PYRIMIDINE_NUCLEOSIDE_TRIPHOSPHATE_BIOSYNTHETIC_P | 17 | 0.73022667 | 1.59594073 | 0.00342626 | 0.06880915 | 0.06298027 | 2528 | tags=53%, list=17%, signal= | TYMS/UCKL1/TBPL1/NME7/CAD/ [illegible] |
| GOBP_HEMOPOIESIS | GOBP_HEMOPOIESIS | 807 | 0.32715719 | 1.27290379 | 0.003416 | 0.06880915 | 0.06298027 | 4196 | tags=32%, list=29%, signal= | HMGB2/TMEM91/ADAR/CMTM7 [illegible] |
| GOBP_SODIUM_ION_TRANSPORT | GOBP_SODIUM_ION_TRANSPORT | 139 | -0.5118832 | -1.6064161 | 0.00358423 | 0.07178973 | 0.06570836 | 736 | tags=9%, list=5%, signal=99 | NDUFA9/ATP1B1/ATP1B4/SCN3B [illegible] |
| GOBP_REGULATION_OF_DNA_METABOLIC_PROCESS | GOBP_REGULATION_OF_DNA_METABOLIC_PROCESS | 451 | 0.37074001 | 1.37768343 | 0.00362615 | 0.07243627 | 0.06630013 | 5191 | tags=38%, list=35%, signal= | PLK1/MCM2/H2AX/CDK1/CHRAC [illegible] |
| GOBP_MESANGIAL_CELL_DIFFERENTIATION | GOBP_MESANGIAL_CELL_DIFFERENTIATION | 7 | 0.87326819 | 1.56916678 | 0.00364431 | 0.07260597 | 0.06645545 | 1497 | tags=29%, list=10%, signal= | NOTCH1/PDGFB |
| GOBP_MONOSACCHARIDE_BIOSYNTHETIC_PROCESS | GOBP_MONOSACCHARIDE_BIOSYNTHETIC_PROCESS | 81 | -0.5745581 | -1.6595776 | 0.00376356 | 0.07361484 | 0.06737886 | 1041 | tags=23%, list=7%, signal=2 | AKR1B1/NNMT/ATF4/ZNF692/PP [illegible] |
| GOBP_HOMOLOGOUS_CHROMOSOME_SEGREGATION | GOBP_HOMOLOGOUS_CHROMOSOME_SEGREGATION | 40 | 0.62171608 | 1.60844165 | 0.00376282 | 0.07361484 | 0.06737886 | 3244 | tags=48%, list=22%, signal= | PLK1/PTTG1/SYCP2/SGO2/KNL1/ [illegible] |
| GOBP_NEGATIVE_REGULATION_OF_PROTEIN_CONTAINING_COMPLEX_ | GOBP_NEGATIVE_REGULATION_OF_PROTEIN_CONTAINING_COMPL | 127 | -0.5106244 | -1.5838094 | 0.00374408 | 0.07361484 | 0.06737886 | 431 | tags=11%, list=3%, signal=1 | CARMIL2/CPTP/HSF1/SIRT2/LMO [illegible] |
| GOBP_POSITIVE_REGULATION_OF_CELL_CYCLE_PHASE_TRANSITION | GOBP_POSITIVE_REGULATION_OF_CELL_CYCLE_PHASE_TRANSITION | 101 | 0.50125671 | 1.54666944 | 0.00371195 | 0.07361484 | 0.06737886 | 2002 | tags=28%, list=14%, signal= | PLK1/RCC2/CDK1/MAD2L1BP/UB [illegible] |
| GOBP_POSITIVE_REGULATION_OF_CYTOKINE_PRODUCTION | GOBP_POSITIVE_REGULATION_OF_CYTOKINE_PRODUCTION | 384 | 0.37465902 | 1.37226184 | 0.0037552 | 0.07361484 | 0.06737886 | 4273 | tags=35%, list=29%, signal= | HMGB2/SCAMP5/EPHB2/KPNA2/ [illegible] |
| GOBP_REGULATION_OF_CELL_CELL_ADHESION | GOBP_REGULATION_OF_CELL_CELL_ADHESION | 385 | 0.37317507 | 1.3671818 | 0.00375753 | 0.07361484 | 0.06737886 | 4408 | tags=35%, list=30%, signal= | PLAUR/EFNA5/TWSG1/CDKN2A/ [illegible] |
| GOBP_CELL_ACTIVATION | GOBP_CELL_ACTIVATION | 893 | 0.32157358 | 1.26109425 | 0.00371747 | 0.07361484 | 0.06737886 | 4218 | tags=31%, list=29%, signal= | EPHB2/CMTM7/TWSG1/RPS6KA1 [illegible] |
| GOBP_RETINA_HOMEOSTASIS | GOBP_RETINA_HOMEOSTASIS | 32 | 0.64030588 | 1.5855891 | 0.00384358 | 0.07459729 | 0.06827809 | 1794 | tags=34%, list=12%, signal= | WHRN/SLC2A1/BBS12/ARAP1/CR [illegible] |
| GOBP_GLYCOSAMINOGLYCAN_METABOLIC_PROCESS | GOBP_GLYCOSAMINOGLYCAN_METABOLIC_PROCESS | 50 | 0.58239432 | 1.57552603 | 0.00383142 | 0.07459729 | 0.06827809 | 3437 | tags=48%, list=23%, signal= | EXT1/TNFAIP6/HEXB/UGDH/HS3 [illegible] |
| GOBP_MEMBRANE_LIPID_METABOLIC_PROCESS | GOBP_MEMBRANE_LIPID_METABOLIC_PROCESS | 169 | 0.45289057 | 1.50481422 | 0.00383509 | 0.07459729 | 0.06827809 | 4405 | tags=41%, list=30%, signal= | GAL3ST4/TM9SF2/PIGL/ELOVL6/ [illegible] |
| GOBP_POSITIVE_REGULATION_OF_PROGRAMMED_NECROTIC_CELL_DE | GOBP_POSITIVE_REGULATION_OF_PROGRAMMED_NECROTIC_CELL_ | 7 | 0.86552572 | 1.55525442 | 0.00393586 | 0.07619135 | 0.06973712 | 1430 | tags=43%, list=10%, signal= | RIPK1/TP53/CASP6 |
| GOBP_LEUKOCYTE_DIFFERENTIATION | GOBP_LEUKOCYTE_DIFFERENTIATION | 526 | 0.35259764 | 1.32722341 | 0.00397662 | 0.07678249 | 0.07027818 | 4196 | tags=32%, list=29%, signal= | CMTM7/CDKN2A/LBR/MYD88/C [illegible] |
| GOBP_REGULATION_OF_CARDIAC_MUSCLE_CELL_ACTION_POTENTIAL | GOBP_REGULATION_OF_CARDIAC_MUSCLE_CELL_ACTION_POTENTI | 27 | -0.7417502 | -1.730226 | 0.00407176 | 0.07823139 | 0.07160435 | 160 | tags=15%, list=1%, signal=1 | CALM1/BIN1/ATP2A2/CAV3 |
| GOBP_REGULATION_OF_MITOTIC_CELL_CYCLE_PHASE_TRANSITION | GOBP_REGULATION_OF_MITOTIC_CELL_CYCLE_PHASE_TRANSITION | 328 | 0.37982946 | 1.37163379 | 0.00407249 | 0.07823139 | 0.07160435 | 2255 | tags=23%, list=15%, signal= | PLK1/RCC2/HECW2/CDCA8/CDK1 [illegible] |
| GOBP_POSITIVE_REGULATION_OF_MITOTIC_SISTER_CHROMATID_SEG | GOBP_POSITIVE_REGULATION_OF_MITOTIC_SISTER_CHROMATID_S | 8 | 0.83956464 | 1.55273223 | 0.0041116 | 0.0787812 | 0.07210758 | 1697 | tags=62%, list=12%, signal= | CDK1/MAD2L1BP/AURKB/SKA1/ [illegible] |
| GOBP_SEX_DIFFERENTIATION | GOBP_SEX_DIFFERENTIATION | 203 | 0.42948641 | 1.46180624 | 0.00412817 | 0.07889741 | 0.07221395 | 4000 | tags=32%, list=27%, signal= | HMGB2/SYCP2/CNOT9/MMP14/ [illegible] |
| GOBP_SKELETAL_MUSCLE_TISSUE_REGENERATION | GOBP_SKELETAL_MUSCLE_TISSUE_REGENERATION | 36 | -0.6860969 | -1.7066065 | 0.00415242 | 0.07915942 | 0.07245376 | 257 | tags=22%, list=2%, signal=2 | LARGE1/DAG1/XIRP1/P2RX5/MY [illegible] |
| GOBP_CARDIAC_VENTRICLE_DEVELOPMENT | GOBP_CARDIAC_VENTRICLE_DEVELOPMENT | 113 | -0.5297398 | -1.6091297 | 0.00424902 | 0.08038882 | 0.07357902 | 400 | tags=12%, list=3%, signal=1 | EGLN1/GREB1L/GSK3A/XIRP2/TN [illegible] |
| GOBP_REGULATION_OF_DNA_TEMPLATED_DNA_REPLICATION_INITIAT | GOBP_REGULATION_OF_DNA_TEMPLATED_DNA_REPLICATION_INIT | 19 | 0.70892381 | 1.58735473 | 0.00424195 | 0.08038882 | 0.07357902 | 2151 | tags=47%, list=15%, signal= | MCM2/CDK2/GMNN/MCM4/MC [illegible] |
| GOBP_HETEROCHROMATIN_ORGANIZATION | GOBP_HETEROCHROMATIN_ORGANIZATION | 172 | 0.44347152 | 1.47723358 | 0.00424383 | 0.08038882 | 0.07357902 | 4808 | tags=45%, list=33%, signal= | PCGF2/BAHD1/H2AX/CDK1/ERI1/ [illegible] |
| GOBP_OXYGEN_TRANSPORT | GOBP_OXYGEN_TRANSPORT | 6 | -0.9528012 | -1.6704764 | 0.00428072 | 0.08078519 | 0.0739418 | 340 | tags=67%, list=2%, signal=6 | HBB/HBA1/HBA2/MB |
| GOBP_PROTEIN_LOCALIZATION_TO_CILIUM | GOBP_PROTEIN_LOCALIZATION_TO_CILIUM | 77 | 0.52547757 | 1.54886512 | 0.00433135 | 0.08121911 | 0.07433897 | 5167 | tags=53%, list=35%, signal= | TMEM67/INPP5E/WDR35/ARF4/ [illegible] |
| GOBP_REGULATION_OF_PROTEIN_POLYMERIZATION | GOBP_REGULATION_OF_PROTEIN_POLYMERIZATION | 175 | -0.4765296 | -1.5460333 | 0.00433616 | 0.08121911 | 0.07433897 | 641 | tags=10%, list=4%, signal=9 | CCR7/MAPRE3/ARL2/CARMIL2/L [illegible] |
| GOBP_NUCLEIC_ACID_CATABOLIC_PROCESS | GOBP_NUCLEIC_ACID_CATABOLIC_PROCESS | 365 | 0.37119848 | 1.35380048 | 0.004329 | 0.08121911 | 0.07433897 | 2426 | tags=19%, list=17%, signal= | MEX3D/INTS9/SNRPF/CNOT3/TE [illegible] |
| GOBP_ANTIFUNGAL_HUMORAL_RESPONSE | GOBP_ANTIFUNGAL_HUMORAL_RESPONSE | 6 | 0.88854795 | 1.54625834 | 0.00435793 | 0.08142394 | 0.07452645 | 184 | tags=17%, list=1%, signal=1 | APP |
| GOBP_POSITIVE_REGULATION_OF_TYPE_II_INTERFERON_PRODUCTION | GOBP_POSITIVE_REGULATION_OF_TYPE_II_INTERFERON_PRODUCTI | 57 | 0.5625817 | 1.56478577 | 0.0043691 | 0.08143008 | 0.07453207 | 3686 | tags=44%, list=25%, signal= | APP/RIPK2/CD276/PSMB9/ABL1/ [illegible] |
| GOBP_NEGATIVE_REGULATION_OF_CELL_ADHESION_MEDIATED_BY_I | GOBP_NEGATIVE_REGULATION_OF_CELL_ADHESION_MEDIATED_BY | 13 | 0.7697462 | 1.58933842 | 0.00439461 | 0.08150094 | 0.07459692 | 1126 | tags=31%, list=8%, signal=2 | SNAI2/EPHA2/MUC1/CYP1B1 |
| GOBP_POSITIVE_REGULATION_OF_IMMUNE_SYSTEM_PROCESS | GOBP_POSITIVE_REGULATION_OF_IMMUNE_SYSTEM_PROCESS | 859 | 0.31907472 | 1.2479892 | 0.00438885 | 0.08150094 | 0.07459692 | 4269 | tags=31%, list=29%, signal= | HMGB2/IRAK4/GBP1/SLC19A1/A [illegible] |
| GOBP_CAMERA_TYPE_EYE_MORPHOGENESIS | GOBP_CAMERA_TYPE_EYE_MORPHOGENESIS | 100 | 0.49712731 | 1.53011757 | 0.00445682 | 0.08224867 | 0.07528132 | 4055 | tags=42%, list=28%, signal= | HMGN1/EFNA5/EPHB2/NOTCH2/ [illegible] |
| GOBP_NEGATIVE_REGULATION_OF_MITOTIC_CELL_CYCLE_PHASE_TRA | GOBP_NEGATIVE_REGULATION_OF_MITOTIC_CELL_CYCLE_PHASE_T | 174 | 0.44000929 | 1.46789441 | 0.0044496 | 0.08224867 | 0.07528132 | 2745 | tags=28%, list=19%, signal= | PLK1/CDCA8/CDK1/NUF2/RBBP8/ [illegible] |
| GOBP_NEURAL_NUCLEUS_DEVELOPMENT | GOBP_NEURAL_NUCLEUS_DEVELOPMENT | 49 | -0.6355119 | -1.6812916 | 0.00447269 | 0.08233921 | 0.07536419 | 989 | tags=24%, list=7%, signal=2 | PITX2/LARGE1/ARX/COX6B1/MA [illegible] |
| GOBP_POSITIVE_REGULATION_OF_IMMUNE_RESPONSE | GOBP_POSITIVE_REGULATION_OF_IMMUNE_RESPONSE | 591 | 0.34205293 | 1.29937971 | 0.0045045 | 0.08272209 | 0.07571463 | 4218 | tags=31%, list=29%, signal= | HMGB2/IRAK4/GBP1/SLC19A1/A [illegible] |
| GOBP_CILIUM_MOVEMENT | GOBP_CILIUM_MOVEMENT | 145 | 0.4661595 | 1.51357715 | 0.00475285 | 0.08706992 | 0.07969416 | 4930 | tags=50%, list=34%, signal= | VANGL1/NME7/CFAP119/DNAI4/ [illegible] |
| GOBP_POSITIVE_REGULATION_OF_SPINDLE_CHECKPOINT | GOBP_POSITIVE_REGULATION_OF_SPINDLE_CHECKPOINT | 20 | 0.7022433 | 1.58625745 | 0.00490119 | 0.08914922 | 0.08159731 | 1298 | tags=35%, list=9%, signal=3 | CDCA8/AURKB/MAD2L1/DYNLL1/ [illegible] |
| GOBP_CELLULAR_RESPONSE_TO_HYDROGEN_PEROXIDE | GOBP_CELLULAR_RESPONSE_TO_HYDROGEN_PEROXIDE | 53 | 0.57100976 | 1.56182657 | 0.00488315 | 0.08914922 | 0.08159731 | 1986 | tags=26%, list=14%, signal= | PCGF2/CDK1/EZH2/MAPK7/MAP [illegible] |
| GOBP_CELL_MOTILITY | GOBP_CELL_MOTILITY | 1389 | 0.29728675 | 1.19485009 | 0.00490196 | 0.08914922 | 0.08159731 | 3699 | tags=27%, list=25%, signal= | PTK7/GDF15/HMGB2/IRAK4/AA [illegible] |
| GOBP_TYPE_II_INTERFERON_PRODUCTION | GOBP_TYPE_II_INTERFERON_PRODUCTION | 87 | 0.51186545 | 1.5361783 | 0.00495504 | 0.08942651 | 0.08185112 | 4406 | tags=41%, list=30%, signal= | APP/RIPK2/CD47/CD276/IL20RB/ [illegible] |
| GOBP_CELL_ACTIVATION_INVOLVED_IN_IMMUNE_RESPONSE | GOBP_CELL_ACTIVATION_INVOLVED_IN_IMMUNE_RESPONSE | 251 | 0.40483123 | 1.41929035 | 0.00493389 | 0.08942651 | 0.08185112 | 5426 | tags=42%, list=37%, signal= | MYD88/NOTCH2/APP/RIPK2/SNX [illegible] |
| GOBP_GLYCOPROTEIN_BIOSYNTHETIC_PROCESS | GOBP_GLYCOPROTEIN_BIOSYNTHETIC_PROCESS | 255 | 0.40170999 | 1.40992503 | 0.00496032 | 0.08942651 | 0.08185112 | 4602 | tags=44%, list=31%, signal= | CHST14/GAL3ST4/SRD5A3/PHLD [illegible] |
| GOBP_MICROTUBULE_BASED_MOVEMENT | GOBP_MICROTUBULE_BASED_MOVEMENT | 363 | 0.36968294 | 1.34670774 | 0.00496483 | 0.08942651 | 0.08185112 | 3834 | tags=34%, list=26%, signal= | VANGL1/CENPE/IFT57/KIF11/KIF [illegible] |
| GOBP_EMBRYONIC_MORPHOGENESIS | GOBP_EMBRYONIC_MORPHOGENESIS | 497 | 0.35570631 | 1.3343478 | 0.00505902 | 0.09090506 | 0.08320441 | 4224 | tags=36%, list=29%, signal= | PTK7/CRABP2/TRIOBP/PCGF2/EP [illegible] |
| GOBP_REGULATION_OF_EXTRINSIC_APOPTOTIC_SIGNALING_PATHWAY | GOBP_REGULATION_OF_EXTRINSIC_APOPTOTIC_SIGNALING_PATHW | 38 | 0.60800727 | 1.55957298 | 0.00508388 | 0.09091679 | 0.08321516 | 753 | tags=24%, list=5%, signal=2 | SNAI2/AKT1/EYA3/IFI6/SRPX/MA [illegible] |
| GOBP_NEGATIVE_REGULATION_OF_GENE_EXPRESSION_EPIGENETIC | GOBP_NEGATIVE_REGULATION_OF_GENE_EXPRESSION_EPIGENETIC | 191 | 0.43000155 | 1.45122489 | 0.00507713 | 0.09091679 | 0.08321516 | 4808 | tags=45%, list=33%, signal= | PCGF2/BAHD1/H2AX/ERI1/LBR/E [illegible] |
| GOBP_POSITIVE_REGULATION_OF_INTERLEUKIN_13_PRODUCTION | GOBP_POSITIVE_REGULATION_OF_INTERLEUKIN_13_PRODUCTION | 15 | 0.74038893 | 1.57559548 | 0.00522434 | 0.09320668 | 0.08531106 | 2808 | tags=60%, list=19%, signal= | IL20RB/RARA/TRAF3IP2/IL20RA/ [illegible] |
| GOBP_CALCIUM_ION_TRANSMEMBRANE_TRANSPORT_VIA_HIGH_VOL | GOBP_CALCIUM_ION_TRANSMEMBRANE_TRANSPORT_VIA_HIGH_V | 17 | -0.8127518 | -1.7286106 | 0.00530578 | 0.09443535 | 0.08643565 | 405 | tags=41%, list=3%, signal=4 | UBR3/CACNA2D1/NOS1/BIN1/CA [illegible] |
| GOBP_REGULATION_OF_TYPE_2_MITOPHAGY | GOBP_REGULATION_OF_TYPE_2_MITOPHAGY | 12 | -0.8582448 | -1.692621 | 0.00540865 | 0.09603877 | 0.08790324 | 1726 | tags=75%, list=12%, signal= | PRKN/HUWE1/VDAC1/VPS13C/C [illegible] |
| GOBP_NEGATIVE_REGULATION_OF_PROTEIN_IMPORT_INTO_NUCLEUS | GOBP_NEGATIVE_REGULATION_OF_PROTEIN_IMPORT_INTO_NUCLE | 19 | -0.7867166 | -1.7047921 | 0.00549904 | 0.09741338 | 0.08916141 | 130 | tags=16%, list=1%, signal=1 | SIRT6/APOD/PKIA |
| GOBP_NEGATIVE_REGULATION_OF_TRANSMEMBRANE_TRANSPORT | GOBP_NEGATIVE_REGULATION_OF_TRANSMEMBRANE_TRANSPORT | 79 | -0.5657209 | -1.6265841 | 0.00556917 | 0.09819256 | 0.08987458 | 1582 | tags=25%, list=11%, signal= | PRKCB/OAZ2/VDAC1/PPP3CC/TG [illegible] |
| GOBP_MRNA_CATABOLIC_PROCESS | GOBP_MRNA_CATABOLIC_PROCESS | 261 | 0.39736934 | 1.39828034 | 0.00556107 | 0.09819256 | 0.08987458 | 2426 | tags=20%, list=17%, signal= | MEX3D/SNRPF/CNOT3/TENT4A/ [illegible] |
| GOBP_REGULATION_OF_SA_NODE_CELL_ACTION_POTENTIAL | GOBP_REGULATION_OF_SA_NODE_CELL_ACTION_POTENTIAL | 4 | 0.93243882 | 1.49747815 | 0.00571094 | 0.10045635 | 0.0919466 | 833 | tags=50%, list=6%, signal=4 | HCN3/TBX18 |
| GOBP_AMINOGLYCAN_CATABOLIC_PROCESS | GOBP_AMINOGLYCAN_CATABOLIC_PROCESS | 23 | 0.66731624 | 1.54736957 | 0.00578871 | 0.10158649 | 0.09298101 | 3308 | tags=61%, list=23%, signal= | CTBS/HEXB/OVGP1/CEMIP/CHI3 [illegible] |
| GOBP_REGULATION_OF_CELL_COMMUNICATION_BY_ELECTRICAL_COU | GOBP_REGULATION_OF_CELL_COMMUNICATION_BY_ELECTRICAL_C | 17 | -0.8027201 | -1.7072746 | 0.00586428 | 0.10267281 | 0.09397531 | 756 | tags=24%, list=5%, signal=2 | CALM1/CASQ2/TRDN/HRC |
| GOBP_REGULATION_OF_MUSCLE_HYPERTROPHY | GOBP_REGULATION_OF_MUSCLE_HYPERTROPHY | 53 | -0.6284536 | -1.6839461 | 0.0059387 | 0.10373381 | 0.09494644 | 784 | tags=26%, list=5%, signal=2 | EDN1/ROCK2/PPARA/IGFBP5/FO [illegible] |
| GOBP_CENTRIOLE_ASSEMBLY | GOBP_CENTRIOLE_ASSEMBLY | 45 | 0.58186184 | 1.53954464 | 0.00620583 | 0.10814841 | 0.09898706 | 3941 | tags=49%, list=27%, signal= | STIL/CDK2/KIAA0753/CEP152/PO [illegible] |
| GOBP_CELL_ADHESION_MOLECULE_PRODUCTION | GOBP_CELL_ADHESION_MOLECULE_PRODUCTION | 6 | 0.87656997 | 1.52541416 | 0.00624637 | 0.10860295 | 0.0994031 | 1775 | tags=67%, list=12%, signal= | GOLPH3/NOTCH1/PPIA/GCNT1 |
| GOBP_EPIDERMAL_CELL_DIFFERENTIATION | GOBP_EPIDERMAL_CELL_DIFFERENTIATION | 138 | 0.46059347 | 1.48672686 | 0.00626424 | 0.10866209 | 0.09945723 | 5202 | tags=52%, list=36%, signal= | TRIOBP/CD109/WHRN/EZH2/KA [illegible] |
| GOBP_REGULATION_OF_CALCIUM_ION_IMPORT_INTO_SARCOPLASMIC | GOBP_REGULATION_OF_CALCIUM_ION_IMPORT_INTO_SARCOPLASM | 4 | -0.9711611 | -1.5826216 | 0.00630318 | 0.10908573 | 0.09984499 | 427 | tags=100%, list=3%, signal= | SLN/STRIT1/ATP2A1 |
| GOBP_DNA_TEMPLATED_DNA_REPLICATION | GOBP_DNA_TEMPLATED_DNA_REPLICATION | 165 | 0.44489832 | 1.47153559 | 0.00637066 | 0.11 | 0.10068181 | 4026 | tags=38%, list=28%, signal= | LIG1/MCM2/RBBP8/CHRAC1/DY [illegible] |
| GOBP_RESPONSE_TO_MUSCLE_INACTIVITY | GOBP_RESPONSE_TO_MUSCLE_INACTIVITY | 6 | -0.9376137 | -1.6438493 | 0.00641437 | 0.11050073 | 0.10114012 | 836 | tags=67%, list=6%, signal=6 | HDAC4/DAG1/FBXO32/CASQ1 |
| GOBP_POSITIVE_REGULATION_OF_MONOCYTE_CHEMOTACTIC_PROTEI | GOBP_POSITIVE_REGULATION_OF_MONOCYTE_CHEMOTACTIC_PRO | 12 | 0.76704049 | 1.55802867 | 0.00644291 | 0.11073849 | 0.10135773 | 2959 | tags=58%, list=20%, signal= | OAS3/OAS1/CLEC7A/SYK/MCOLN [illegible] |
| GOBP_PROTEIN_POLYMERIZATION | GOBP_PROTEIN_POLYMERIZATION | 253 | -0.4248094 | -1.4392127 | 0.00647249 | 0.11099289 | 0.10159059 | 756 | tags=9%, list=5%, signal=99 | CASQ2/GPX4/CCR7/COBL/MAPRE [illegible] |
| GOBP_CARBOHYDRATE_BIOSYNTHETIC_PROCESS | GOBP_CARBOHYDRATE_BIOSYNTHETIC_PROCESS | 149 | -0.4856559 | -1.5392456 | 0.0065387 | 0.11161867 | 0.10216335 | 1052 | tags=22%, list=7%, signal=2 | INPP5K/AKT2/AKR1B1/PPP1CA/E [illegible] |
| GOBP_MESANGIAL_CELL_DEVELOPMENT | GOBP_MESANGIAL_CELL_DEVELOPMENT | 6 | 0.87355065 | 1.52015992 | 0.0065369 | 0.11161867 | 0.10216335 | 1497 | tags=33%, list=10%, signal= | NOTCH1/PDGFB |
| GOBP_PROTEIN_N_LINKED_GLYCOSYLATION | GOBP_PROTEIN_N_LINKED_GLYCOSYLATION | 83 | 0.51294452 | 1.52966762 | 0.00657414 | 0.11196911 | 0.10248411 | 5893 | tags=61%, list=40%, signal= | SRD5A3/OSTC/STT3A/SLC39A8/T [illegible] |
| GOBP_ZINC_ION_IMPORT_ACROSS_PLASMA_MEMBRANE | GOBP_ZINC_ION_IMPORT_ACROSS_PLASMA_MEMBRANE | 8 | 0.83149326 | 1.53780461 | 0.00660793 | 0.11228995 | 0.10277777 | 701 | tags=50%, list=5%, signal=4 | SLC39A11/SLC39A8/SLC30A1/SLC [illegible] |
| GOBP_PROTEIN_CONTAINING_COMPLEX_DISASSEMBLY | GOBP_PROTEIN_CONTAINING_COMPLEX_DISASSEMBLY | 244 | -0.4274586 | -1.4421994 | 0.00665994 | 0.1129183 | 0.10335289 | 930 | tags=15%, list=6%, signal=1 | CAPG/FLII/CALM1/PEX12/CHMP6 [illegible] |
| GOBP_REGULATION_OF_VENTRICULAR_CARDIAC_MUSCLE_CELL_MEM | GOBP_REGULATION_OF_VENTRICULAR_CARDIAC_MUSCLE_CELL_ME | 6 | -0.9443426 | -1.6556466 | 0.006705 | 0.11342631 | 0.10381788 | 394 | tags=50%, list=3%, signal=4 | GPD1L/SCN3B/CAV3 |
| GOBP_DIAPEDESIS | GOBP_DIAPEDESIS | 6 | -0.9357589 | -1.6405974 | 0.00673509 | 0.11348253 | 0.10386933 | 377 | tags=67%, list=3%, signal=6 | PECAM1/CRKL/CD99L2/TRIM55 |
| GOBP_APPENDAGE_DEVELOPMENT | GOBP_APPENDAGE_DEVELOPMENT | 167 | 0.44147965 | 1.46369681 | 0.00673854 | 0.11348253 | 0.10386933 | 4216 | tags=41%, list=29%, signal= | CRABP2/EXT1/OSR2/NOTCH2/SL [illegible] |
| GOBP_POSITIVE_REGULATION_OF_MAPK_CASCADE | GOBP_POSITIVE_REGULATION_OF_MAPK_CASCADE | 350 | 0.36952736 | 1.34218288 | 0.0067623 | 0.11362774 | 0.10400224 | 3589 | tags=26%, list=25%, signal= | GDF15/MYDGF/LPAR2/ARRB2/I [illegible] |
| GOBP_CYTOKINE_MEDIATED_SIGNALING_PATHWAY | GOBP_CYTOKINE_MEDIATED_SIGNALING_PATHWAY | 384 | 0.36075833 | 1.32134783 | 0.00683371 | 0.11457147 | 0.10486602 | 4433 | tags=34%, list=30%, signal= | IRAK4/IFNAR2/ADAR/TMSB4X/E [illegible] |
| GOBP_RESPONSE_TO_INTERFERON_ALPHA | GOBP_RESPONSE_TO_INTERFERON_ALPHA | 17 | 0.70776401 | 1.54684765 | 0.00700825 | 0.11714171 | 0.10721854 | 2090 | tags=35%, list=14%, signal= | IFNAR2/ADAR/OAS1/EIF2AK2/KL [illegible] |
| GOBP_EYE_MORPHOGENESIS | GOBP_EYE_MORPHOGENESIS | 125 | 0.4645909 | 1.47668592 | 0.00701821 | 0.11714171 | 0.10721854 | 4055 | tags=41%, list=28%, signal= | HMGN1/EFNA5/EPHB2/NOTCH2/ [illegible] |
| GOBP_POSITIVE_REGULATION_OF_STRIATED_MUSCLE_CELL_APOPTOTI | GOBP_POSITIVE_REGULATION_OF_STRIATED_MUSCLE_CELL_APOPT | 11 | -0.871834 | -1.6855348 | 0.00705738 | 0.11748257 | 0.10753052 | 305 | tags=36%, list=2%, signal=3 | ADRB2/CAMK2A/LTK/ATP2A2 |
| GOBP_INTERLEUKIN_13_PRODUCTION | GOBP_INTERLEUKIN_13_PRODUCTION | 19 | 0.69291447 | 1.55150813 | 0.00706991 | 0.11748257 | 0.10753052 | 3189 | tags=63%, list=22%, signal= | IL20RB/TNFRSF21/LEF1/RARA/T [illegible] |
| GOBP_SUBSTANTIA_NIGRA_DEVELOPMENT | GOBP_SUBSTANTIA_NIGRA_DEVELOPMENT | 37 | -0.6660595 | -1.664285 | 0.00710959 | 0.11783557 | 0.10785362 | 844 | tags=24%, list=6%, signal=2 | COX6B1/MAOB/ATP5PF/SIRT2/N [illegible] |
| GOBP_PROTEIN_DEPOLYMERIZATION | GOBP_PROTEIN_DEPOLYMERIZATION | 108 | -0.517486 | -1.559075 | 0.00713822 | 0.11783557 | 0.10785362 | 431 | tags=12%, list=3%, signal=1 | CARMIL2/LMOD1/CFL2/TMOD1/ [illegible] |
| GOBP_CONNECTIVE_TISSUE_DEVELOPMENT | GOBP_CONNECTIVE_TISSUE_DEVELOPMENT | 259 | 0.39272867 | 1.38093702 | 0.00713578 | 0.11783557 | 0.10785362 | 3997 | tags=33%, list=27%, signal= | ABHD15/SNAI2/EXT1/OSR2/BMP [illegible] |
| GOBP_NEGATIVE_REGULATION_OF_CALCIUM_ION_TRANSMEMBRANE_ | GOBP_NEGATIVE_REGULATION_OF_CALCIUM_ION_TRANSMEMBRA | 27 | -0.7212426 | -1.6823893 | 0.00715929 | 0.11792425 | 0.10793479 | 1463 | tags=44%, list=10%, signal= | VDAC1/PPP3CC/TGFB1/CALM1/G [illegible] |
| GOBP_REGULATION_OF_DOUBLE_STRAND_BREAK_REPAIR_VIA_HOMO | GOBP_REGULATION_OF_DOUBLE_STRAND_BREAK_REPAIR_VIA_HO | 76 | 0.51760757 | 1.52227769 | 0.00720072 | 0.11834706 | 0.10832178 | 4836 | tags=45%, list=33%, signal= | PLK1/SKP2/RMI2/KLHL15/WRAP [illegible] |
| GOBP_NEGATIVE_REGULATION_OF_ACTIN_FILAMENT_POLYMERIZATIO | GOBP_NEGATIVE_REGULATION_OF_ACTIN_FILAMENT_POLYMERIZA | 51 | -0.6158649 | -1.6395338 | 0.00728041 | 0.1193956 | 0.1092815 | 431 | tags=18%, list=3%, signal=1 | CARMIL2/LMOD1/TMOD1/SVIL/ [illegible] |
| GOBP_MYOBLAST_FUSION | GOBP_MYOBLAST_FUSION | 80 | -0.5548193 | -1.5967815 | 0.00734967 | 0.12000726 | 0.10984135 | 236 | tags=8%, list=2%, signal=79 | DYRK1B/NOS1/MYOD1/RPL3L/C [illegible] |
| GOBP_REGULATION_OF_CELL_ADHESION | GOBP_REGULATION_OF_CELL_ADHESION | 632 | 0.33105564 | 1.26523787 | 0.00733391 | 0.12000726 | 0.10984135 | 4408 | tags=34%, list=30%, signal= | PLAUR/GBP1/TRIOBP/RCC2/SNA [illegible] |
| GOBP_POSITIVE_REGULATION_OF_POTASSIUM_ION_TRANSPORT | GOBP_POSITIVE_REGULATION_OF_POTASSIUM_ION_TRANSPORT | 24 | -0.7406385 | -1.6828809 | 0.00738007 | 0.12024237 | 0.11005654 | 696 | tags=17%, list=5%, signal=1 | ATP1B1/AKAP6/ACTN2/FHL1 |
| GOBP_POSITIVE_REGULATION_OF_TYPE_2_IMMUNE_RESPONSE | GOBP_POSITIVE_REGULATION_OF_TYPE_2_IMMUNE_RESPONSE | 13 | 0.74644512 | 1.54122736 | 0.00742537 | 0.12045774 | 0.11025367 | 2762 | tags=62%, list=19%, signal= | CD86/RSAD2/RARA/IDO1/IL18/T [illegible] |
| GOBP_REGULATION_OF_HETEROCHROMATIN_ORGANIZATION | GOBP_REGULATION_OF_HETEROCHROMATIN_ORGANIZATION | 13 | 0.74585059 | 1.5399998 | 0.00742537 | 0.12045774 | 0.11025367 | 896 | tags=31%, list=6%, signal=2 | CDK1/ERI1/CDK2/ERI2 |
| GOBP_BLASTOCYST_GROWTH | GOBP_BLASTOCYST_GROWTH | 21 | 0.67119623 | 1.53311862 | 0.0074615 | 0.12078306 | 0.11055143 | 1906 | tags=33%, list=13%, signal= | NDEL1/GINS1/PALB2/GINS4/SAL [illegible] |
| GOBP_REGULATION_OF_SKELETAL_MUSCLE_TISSUE_DEVELOPMENT | GOBP_REGULATION_OF_SKELETAL_MUSCLE_TISSUE_DEVELOPMENT | 18 | -0.7894046 | -1.6934102 | 0.0075188 | 0.12144878 | 0.11116075 | 252 | tags=39%, list=2%, signal=3 | TGFB1/MTM1/MEF2C/YBX3/MY [illegible] |
| GOBP_POSITIVE_REGULATION_OF_SODIUM_ION_TRANSMEMBRANE_T | GOBP_POSITIVE_REGULATION_OF_SODIUM_ION_TRANSMEMBRAN | 14 | -0.8378332 | -1.7118228 | 0.00754498 | 0.12161018 | 0.11130848 | 696 | tags=29%, list=5%, signal=2 | ATP1B1/NOS1/SCN1B/FXYD1 |
| GOBP_REGULATION_OF_SKELETAL_MUSCLE_CELL_DIFFERENTIATION | GOBP_REGULATION_OF_SKELETAL_MUSCLE_CELL_DIFFERENTIATIO | 20 | -0.774551 | -1.6990127 | 0.0076149 | 0.12247439 | 0.11209948 | 797 | tags=40%, list=5%, signal=3 | NR1D2/SIX1/RBM24/SIRT2/GPC1 [illegible] |
| GOBP_TOLL_SIGNALING_PATHWAY | GOBP_TOLL_SIGNALING_PATHWAY | 12 | 0.75822016 | 1.54011264 | 0.00764159 | 0.12264106 | 0.11225203 | 184 | tags=25%, list=1%, signal=2 | IRAK4/MYD88/APP |
| GOBP_POSITIVE_REGULATION_OF_RESPONSE_TO_BIOTIC_STIMULUS | GOBP_POSITIVE_REGULATION_OF_RESPONSE_TO_BIOTIC_STIMULU | 338 | 0.36908552 | 1.33550683 | 0.00776937 | 0.12442592 | 0.11388569 | 4657 | tags=36%, list=32%, signal= | HMGB2/IRAK4/SLC19A1/ARRB2/ [illegible] |
| GOBP_INTRACILIARY_TRANSPORT | GOBP_INTRACILIARY_TRANSPORT | 51 | 0.56424995 | 1.53142812 | 0.00783426 | 0.12519817 | 0.11459253 | 3687 | tags=45%, list=25%, signal= | IFT57/WDR35/NME7/DYNLL1/R [illegible] |
| GOBP_MITOCHONDRIAL_CALCIUM_ION_HOMEOSTASIS | GOBP_MITOCHONDRIAL_CALCIUM_ION_HOMEOSTASIS | 22 | -0.7541826 | -1.6851185 | 0.00802354 | 0.12767961 | 0.11686376 | 1023 | tags=14%, list=7%, signal=1 | C19orf12/SMDT1/ATP2A1 |
| GOBP_RESPONSE_TO_OXYGEN_RADICAL | GOBP_RESPONSE_TO_OXYGEN_RADICAL | 22 | -0.7520903 | -1.6804435 | 0.00802354 | 0.12767961 | 0.11686376 | 101 | tags=18%, list=1%, signal=1 | PARK7/PRDX2/MB/UCP3 |
| GOBP_DEOXYRIBONUCLEOTIDE_BIOSYNTHETIC_PROCESS | GOBP_DEOXYRIBONUCLEOTIDE_BIOSYNTHETIC_PROCESS | 16 | 0.71499503 | 1.5421642 | 0.00805827 | 0.1279612 | 0.1171215 | 3775 | tags=69%, list=26%, signal= | TYMS/RRM2/TBPL1/CMPK2/DCK [illegible] |
| GOBP_TRNA_PROCESSING | GOBP_TRNA_PROCESSING | 137 | 0.45303906 | 1.45987256 | 0.00817025 | 0.12946564 | 0.11849849 | 2087 | tags=21%, list=14%, signal= | TRIT1/PUS1/AKT1/TRMT11/QTR [illegible] |
| GOBP_POSITIVE_REGULATION_OF_MITOTIC_CELL_CYCLE_PHASE_TRAN | GOBP_POSITIVE_REGULATION_OF_MITOTIC_CELL_CYCLE_PHASE_TR | 85 | 0.50620225 | 1.51539418 | 0.00821768 | 0.1299431 | 0.11893551 | 2002 | tags=29%, list=14%, signal= | PLK1/RCC2/CDK1/MAD2L1BP/UB [illegible] |
| GOBP_REGULATION_OF_MEMBRANE_DEPOLARIZATION_DURING_ACTI | GOBP_REGULATION_OF_MEMBRANE_DEPOLARIZATION_DURING_A | 7 | -0.9233443 | -1.6613424 | 0.00827498 | 0.13057438 | 0.11951332 | 36 | tags=29%, list=0%, signal=2 | ANK3/CAV3 |
| GOBP_SKELETAL_MUSCLE_ATROPHY | GOBP_SKELETAL_MUSCLE_ATROPHY | 3 | -0.9834594 | -1.5156281 | 0.00841346 | 0.13248115 | 0.12125856 | 100 | tags=100%, list=1%, signal= | ASB2/TRIM63 |
| GOBP_REGULATION_OF_ACTION_POTENTIAL | GOBP_REGULATION_OF_ACTION_POTENTIAL | 44 | -0.6597988 | -1.705949 | 0.00853208 | 0.13406786 | 0.12271086 | 405 | tags=14%, list=3%, signal=1 | CASQ2/CACNA2D1/NOS1/BIN1/A [illegible] |
| GOBP_CELL_COMMUNICATION_INVOLVED_IN_CARDIAC_CONDUCTION | GOBP_CELL_COMMUNICATION_INVOLVED_IN_CARDIAC_CONDUCTI | 45 | -0.6259294 | -1.6241711 | 0.00856939 | 0.13437302 | 0.12299017 | 756 | tags=20%, list=5%, signal=1 | CASQ2/ATP1B1/PRKACA/CACNA [illegible] |
| GOBP_DETECTION_OF_MUSCLE_STRETCH | GOBP_DETECTION_OF_MUSCLE_STRETCH | 7 | -0.9230889 | -1.6608829 | 0.00859325 | 0.1344665 | 0.12307573 | 190 | tags=57%, list=1%, signal=5 | CSRP3/TTN/TCAP/CAV3 |
| GOBP_RESPONSE_TO_ACTIVITY | GOBP_RESPONSE_TO_ACTIVITY | 66 | -0.5799431 | -1.615626 | 0.00866196 | 0.13525977 | 0.1238018 | 805 | tags=17%, list=5%, signal=1 | PPARGC1A/ADSL/FOS/DAG1/FND [illegible] |
| GOBP_EXTRACELLULAR_TRANSPORT | GOBP_EXTRACELLULAR_TRANSPORT | 36 | 0.60121145 | 1.5245202 | 0.00880162 | 0.13715561 | 0.12553704 | 4924 | tags=69%, list=34%, signal= | VANGL1/NME7/DNAAF2/CCDC40 [illegible] |
| GOBP_MYOSIN_FILAMENT_ORGANIZATION | GOBP_MYOSIN_FILAMENT_ORGANIZATION | 5 | -0.9496827 | -1.6127224 | 0.00892857 | 0.13866425 | 0.12691789 | 481 | tags=60%, list=3%, signal=5 | MYH11/TTN/TCAP |
| GOBP_MULTI_MULTICELLULAR_ORGANISM_PROCESS | GOBP_MULTI_MULTICELLULAR_ORGANISM_PROCESS | 145 | 0.44633557 | 1.44921068 | 0.00893536 | 0.13866425 | 0.12691789 | 2523 | tags=26%, list=17%, signal= | SLC19A1/GJA1/AKT1/SLC2A1/CA [illegible] |
| GOBP_SHORT_CHAIN_FATTY_ACID_METABOLIC_PROCESS | GOBP_SHORT_CHAIN_FATTY_ACID_METABOLIC_PROCESS | 12 | -0.8447961 | -1.6660976 | 0.00901442 | 0.13932148 | 0.12751944 | 279 | tags=42%, list=2%, signal=4 | ACOT9/CRAT/PHYH/ACADS/ACSS [illegible] |
| GOBP_REGULATION_OF_ACTIN_FILAMENT_POLYMERIZATION | GOBP_REGULATION_OF_ACTIN_FILAMENT_POLYMERIZATION | 113 | -0.510016 | -1.5492168 | 0.00901481 | 0.13932148 | 0.12751944 | 431 | tags=11%, list=3%, signal=1 | CCR7/CARMIL2/LMOD1/TMOD1/ [illegible] |
| GOBP_DENSE_CORE_GRANULE_TRANSPORT | GOBP_DENSE_CORE_GRANULE_TRANSPORT | 8 | -0.9029559 | -1.6688451 | 0.00908521 | 0.13994296 | 0.12808828 | 313 | tags=38%, list=2%, signal=3 | PPFIA2/KIF1C/KIF1B |
| GOBP_MEIOTIC_CELL_CYCLE_PHASE_TRANSITION | GOBP_MEIOTIC_CELL_CYCLE_PHASE_TRANSITION | 13 | 0.74037301 | 1.52868993 | 0.00909229 | 0.13994296 | 0.12808828 | 2717 | tags=77%, list=19%, signal= | CCNB2/PKMYT1/KNL1/CDC25C/N [illegible] |
| GOBP_POSITIVE_REGULATION_OF_TYPE_2_MITOPHAGY | GOBP_POSITIVE_REGULATION_OF_TYPE_2_MITOPHAGY | 10 | -0.876369 | -1.6818248 | 0.00917712 | 0.14065488 | 0.12873988 | 1726 | tags=80%, list=12%, signal= | PRKN/HUWE1/VDAC1/CDC37/AT [illegible] |
| GOBP_AZOLE_TRANSMEMBRANE_TRANSPORT | GOBP_AZOLE_TRANSMEMBRANE_TRANSPORT | 11 | 0.76803604 | 1.52960341 | 0.00919472 | 0.14065488 | 0.12873988 | 2373 | tags=55%, list=16%, signal= | SLC7A1/SLC19A2/SLC25A19/SLC4 [illegible] |
| GOBP_SIGNAL_TRANSDUCTION_BY_P53_CLASS_MEDIATOR | GOBP_SIGNAL_TRANSDUCTION_BY_P53_CLASS_MEDIATOR | 184 | 0.42214024 | 1.41717088 | 0.00917969 | 0.14065488 | 0.12873988 | 4369 | tags=36%, list=30%, signal= | SNAI2/AURKB/CDKN2A/AKT1/FO [illegible] |
| GOBP_RESPONSE_TO_REACTIVE_OXYGEN_SPECIES | GOBP_RESPONSE_TO_REACTIVE_OXYGEN_SPECIES | 156 | -0.4636439 | -1.4800373 | 0.00923594 | 0.1409982 | 0.12905413 | 857 | tags=12%, list=6%, signal=1 | COA8/KPNA4/PDK2/FOS/MAPK8/ [illegible] |
| GOBP_NUCLEAR_TRANSPORT | GOBP_NUCLEAR_TRANSPORT | 308 | 0.37504856 | 1.34637274 | 0.00925553 | 0.14101077 | 0.12906563 | 4067 | tags=32%, list=28%, signal= | CDK1/NUP54/ADAR/CDKN2A/MA [illegible] |
| GOBP_CARDIAC_CONDUCTION_SYSTEM_DEVELOPMENT | GOBP_CARDIAC_CONDUCTION_SYSTEM_DEVELOPMENT | 30 | 0.62315447 | 1.52736538 | 0.00928382 | 0.1411554 | 0.12919801 | 2412 | tags=30%, list=16%, signal= | GJA1/NOTCH2/CDC42/NOTCH1/ [illegible] |
| GOBP_NEGATIVE_REGULATION_OF_DOUBLE_STRAND_BREAK_REPAIR_ | GOBP_NEGATIVE_REGULATION_OF_DOUBLE_STRAND_BREAK_REPA | 24 | 0.64895566 | 1.51998853 | 0.00934278 | 0.14176494 | 0.12975592 | 4110 | tags=50%, list=28%, signal= | PLK1/RMI2/KLHL15/ABL1/CGAS/ [illegible] |
| GOBP_REGULATION_OF_FIBRINOLYSIS | GOBP_REGULATION_OF_FIBRINOLYSIS | 8 | 0.82136191 | 1.5190672 | 0.00939794 | 0.14231443 | 0.13025886 | 96 | tags=25%, list=1%, signal=2 | PLAUR/PLAU |
| GOBP_HOMOLOGOUS_RECOMBINATION | GOBP_HOMOLOGOUS_RECOMBINATION | 52 | 0.55639813 | 1.51570783 | 0.00941751 | 0.14232377 | 0.1302674 | 3208 | tags=37%, list=22%, signal= | RBBP8/SYCP2/EME1/RAD51AP1/ [illegible] |
| GOBP_REGULATION_OF_SKELETAL_MUSCLE_TISSUE_REGENERATION | GOBP_REGULATION_OF_SKELETAL_MUSCLE_TISSUE_REGENERATIO | 7 | -0.9185896 | -1.6527874 | 0.00954806 | 0.14400696 | 0.13180801 | 227 | tags=57%, list=2%, signal=5 | P2RX5/MYOD1/CAPN3/MYOZ1 |
| GOBP_MUSCLE_HYPERTROPHY_IN_RESPONSE_TO_STRESS | GOBP_MUSCLE_HYPERTROPHY_IN_RESPONSE_TO_STRESS | 21 | -0.7584549 | -1.6790781 | 0.00972185 | 0.14633426 | 0.13393816 | 85 | tags=29%, list=1%, signal=2 | HDAC4/CAMTA2/MYH7/ATP2A2/ [illegible] |
| GOBP_DOUBLE_STRAND_BREAK_REPAIR_VIA_SINGLE_STRAND_ANNEA | GOBP_DOUBLE_STRAND_BREAK_REPAIR_VIA_SINGLE_STRAND_AN | 6 | 0.85561784 | 1.48895309 | 0.00987798 | 0.14838698 | 0.135817 | 1996 | tags=50%, list=14%, signal= | RBBP8/RNF138/HELQ |
| GOBP_POSITIVE_REGULATION_OF_UBIQUITIN_PROTEIN_TRANSFERASE | GOBP_POSITIVE_REGULATION_OF_UBIQUITIN_PROTEIN_TRANSFER | 9 | 0.7960113 | 1.51207373 | 0.01019052 | 0.15277641 | 0.1398346 | 2002 | tags=44%, list=14%, signal= | UBE2C/CDC14B/FZR1/CDC20 |
| GOBP_DNA_STRAND_ELONGATION_INVOLVED_IN_DNA_REPLICATION | GOBP_DNA_STRAND_ELONGATION_INVOLVED_IN_DNA_REPLICATI | 15 | 0.71537023 | 1.52235407 | 0.01029502 | 0.15312061 | 0.14014963 | 2406 | tags=47%, list=16%, signal= | LIG1/GINS1/MCM4/PCNA/POLD3 [illegible] |
| GOBP_INNER_CELL_MASS_CELL_PROLIFERATION | GOBP_INNER_CELL_MASS_CELL_PROLIFERATION | 15 | 0.71504102 | 1.52165349 | 0.01029502 | 0.15312061 | 0.14014963 | 1849 | tags=40%, list=13%, signal= | NDEL1/GINS1/PALB2/GINS4/SAL [illegible] |
| GOBP_CHROMOSOME_ORGANIZATION_INVOLVED_IN_MEIOTIC_CELL_ | GOBP_CHROMOSOME_ORGANIZATION_INVOLVED_IN_MEIOTIC_CE | 40 | 0.57922255 | 1.49850665 | 0.01025466 | 0.15312061 | 0.14014963 | 3244 | tags=42%, list=22%, signal= | SYCP2/SMC4/BRIP1/RNF212/NCA [illegible] |
| GOBP_TRNA_MODIFICATION | GOBP_TRNA_MODIFICATION | 96 | 0.48298652 | 1.47947717 | 0.01028655 | 0.15312061 | 0.14014963 | 2087 | tags=23%, list=14%, signal= | TRIT1/PUS1/AKT1/QTRT2/QNG1/ [illegible] |
| GOBP_RESPONSE_TO_MACROPHAGE_COLONY_STIMULATING_FACTOR | GOBP_RESPONSE_TO_MACROPHAGE_COLONY_STIMULATING_FACT | 14 | 0.7255063 | 1.52364502 | 0.01037218 | 0.15389618 | 0.14085951 | 1609 | tags=36%, list=11%, signal= | ALPL/PTPN2/TLR2/DOK1/CSF1 |
| GOBP_POSITIVE_REGULATION_OF_MEGAKARYOCYTE_DIFFERENTIATIO | GOBP_POSITIVE_REGULATION_OF_MEGAKARYOCYTE_DIFFERENTIAT | 10 | 0.77836682 | 1.51617227 | 0.01039655 | 0.15389618 | 0.14085951 | 1526 | tags=30%, list=10%, signal= | HMGB2/SCIN/RCOR1 |
| GOBP_DEVELOPMENT_OF_PRIMARY_SEXUAL_CHARACTERISTICS | GOBP_DEVELOPMENT_OF_PRIMARY_SEXUAL_CHARACTERISTICS | 171 | 0.42953131 | 1.42861442 | 0.01040864 | 0.15389618 | 0.14085951 | 4000 | tags=32%, list=27%, signal= | HMGB2/MMP14/IDH1/BASP1/T [illegible] |
| GOBP_APPENDAGE_MORPHOGENESIS | GOBP_APPENDAGE_MORPHOGENESIS | 133 | 0.45220831 | 1.45116637 | 0.01043841 | 0.15391393 | 0.14087575 | 4196 | tags=42%, list=29%, signal= | CRABP2/EXT1/OSR2/NOTCH2/FL [illegible] |
| GOBP_OSSIFICATION | GOBP_OSSIFICATION | 350 | 0.36126987 | 1.31219035 | 0.01045082 | 0.15391393 | 0.14087575 | 3159 | tags=27%, list=22%, signal= | SNAI2/CREB3L1/ADAR/TWSG1/E [illegible] |
| GOBP_REGULATION_OF_HEMOPOIESIS | GOBP_REGULATION_OF_HEMOPOIESIS | 357 | 0.36025464 | 1.31082449 | 0.01049599 | 0.15427663 | 0.14120772 | 4433 | tags=35%, list=30%, signal= | HMGB2/CDKN2A/NOTCH2/ADA [illegible] |
| GOBP_RESPONSE_TO_ABIOTIC_STIMULUS | GOBP_RESPONSE_TO_ABIOTIC_STIMULUS | 904 | -0.3313323 | -1.2562144 | 0.01065002 | 0.15623494 | 0.14300014 | 850 | tags=7%, list=6%, signal=79 | ATF4/EIF2B3/ACADVL/EGLN1/T [illegible] |
| GOBP_REGULATION_OF_CAVEOLIN_MEDIATED_ENDOCYTOSIS | GOBP_REGULATION_OF_CAVEOLIN_MEDIATED_ENDOCYTOSIS | 6 | -0.9291685 | -1.629043 | 0.01090443 | 0.15942097 | 0.14591628 | 36 | tags=33%, list=0%, signal=3 | ITSN1/CAV3 |
| GOBP_REGULATION_OF_CHROMOSOME_CONDENSATION | GOBP_REGULATION_OF_CHROMOSOME_CONDENSATION | 15 | 0.71283482 | 1.51695855 | 0.01090965 | 0.15942097 | 0.14591628 | 2986 | tags=53%, list=20%, signal= | CCNB1/SMC4/NCAPD2/SMC2/N [illegible] |
| GOBP_REGULATION_OF_ATRIAL_CARDIAC_MUSCLE_CELL_MEMBRANE_ | GOBP_REGULATION_OF_ATRIAL_CARDIAC_MUSCLE_CELL_MEMBRA | 8 | -0.8968354 | -1.6575332 | 0.01096491 | 0.15960747 | 0.14608699 | 394 | tags=50%, list=3%, signal=4 | SCN2B/SCN3B/SCN1B/FHL1 |
| GOBP_CYTOKINE_PRODUCTION | GOBP_CYTOKINE_PRODUCTION | 629 | 0.32644127 | 1.24710555 | 0.01096303 | 0.15960747 | 0.14608699 | 4333 | tags=32%, list=30%, signal= | HMGB2/GBP1/SNAI2/SCAMP5/A [illegible] |
| GOBP_VITAMIN_TRANSMEMBRANE_TRANSPORT | GOBP_VITAMIN_TRANSMEMBRANE_TRANSPORT | 16 | 0.70448257 | 1.51949001 | 0.01100263 | 0.15981456 | 0.14627653 | 2207 | tags=44%, list=15%, signal= | SLC19A1/STRA6/SLC19A2/SCARB [illegible] |
| GOBP_DEFENSE_RESPONSE_TO_GRAM_POSITIVE_BACTERIUM | GOBP_DEFENSE_RESPONSE_TO_GRAM_POSITIVE_BACTERIUM | 56 | 0.54782263 | 1.51658868 | 0.01102169 | 0.15981456 | 0.14627653 | 4185 | tags=38%, list=29%, signal= | HMGB2/MYD88/APP/RIPK2/EPH [illegible] |
| GOBP_CALCIUM_ION_TRANSPORT_INTO_CYTOSOL | GOBP_CALCIUM_ION_TRANSPORT_INTO_CYTOSOL | 20 | -0.7657543 | -1.6797168 | 0.01115039 | 0.16060356 | 0.14699869 | 405 | tags=25%, list=3%, signal=2 | CACNA2D1/JPH1/NOS1/P2RX5/J [illegible] |
| GOBP_T_HELPER_1_TYPE_IMMUNE_RESPONSE | GOBP_T_HELPER_1_TYPE_IMMUNE_RESPONSE | 50 | 0.55418717 | 1.49921847 | 0.01114594 | 0.16060356 | 0.14699869 | 5151 | tags=50%, list=35%, signal= | RIPK2/PSMB9/PSMB8/TMEM98/ [illegible] |
| GOBP_NEGATIVE_REGULATION_OF_EXTRINSIC_APOPTOTIC_SIGNALING | GOBP_NEGATIVE_REGULATION_OF_EXTRINSIC_APOPTOTIC_SIGNALI | 82 | 0.49965684 | 1.48588407 | 0.01114359 | 0.16060356 | 0.14699869 | 1470 | tags=17%, list=10%, signal= | HMGB2/SNAI2/AKT1/EYA3/IFI6/ [illegible] |
| GOBP_RESPONSE_TO_MECHANICAL_STIMULUS | GOBP_RESPONSE_TO_MECHANICAL_STIMULUS | 177 | -0.4501591 | -1.4620632 | 0.01116164 | 0.16060356 | 0.14699869 | 784 | tags=11%, list=5%, signal=1 | TNFRSF1A/FOS/MAPK8/DAG1/U [illegible] |
| GOBP_NEGATIVE_REGULATION_OF_SUPRAMOLECULAR_FIBER_ORGAN | GOBP_NEGATIVE_REGULATION_OF_SUPRAMOLECULAR_FIBER_ORG | 137 | -0.4766522 | -1.4925591 | 0.01118379 | 0.16061468 | 0.14700887 | 431 | tags=9%, list=3%, signal=99 | RAB40B/CARMIL2/LMOD1/TMO [illegible] |
| GOBP_OPTIC_NERVE_DEVELOPMENT | GOBP_OPTIC_NERVE_DEVELOPMENT | 9 | 0.79166874 | 1.50382478 | 0.01122434 | 0.16088933 | 0.14726026 | 401 | tags=33%, list=3%, signal=3 | EPHB2/EXT1/GLI3 |
| GOBP_NUCLEAR_EXPORT | GOBP_NUCLEAR_EXPORT | 160 | 0.4347032 | 1.43285873 | 0.01124881 | 0.16093295 | 0.14730019 | 4453 | tags=35%, list=30%, signal= | ADAR/CDKN2A/MAGOHB/LTV1/ [illegible] |
| GOBP_REGULATION_OF_MRNA_CATABOLIC_PROCESS | GOBP_REGULATION_OF_MRNA_CATABOLIC_PROCESS | 192 | 0.41210952 | 1.39164118 | 0.01131928 | 0.16163332 | 0.14794122 | 2426 | tags=21%, list=17%, signal= | MEX3D/CNOT3/TENT4A/MAGOH [illegible] |
| GOBP_POSITIVE_REGULATION_OF_TRANSPORT | GOBP_POSITIVE_REGULATION_OF_TRANSPORT | 641 | -0.3502304 | -1.296305 | 0.01139974 | 0.16247331 | 0.14871006 | 906 | tags=8%, list=6%, signal=89 | HIP1/OSBP/DGKD/ATF4/ABCB4/ [illegible] |
| GOBP_REGULATION_OF_NUCLEAR_TRANSCRIBED_MRNA_CATABOLIC_ | GOBP_REGULATION_OF_NUCLEAR_TRANSCRIBED_MRNA_CATABOLI | 16 | 0.70261608 | 1.51546421 | 0.01146753 | 0.16313002 | 0.14931114 | 2415 | tags=44%, list=16%, signal= | MAGOHB/DHX34/MAGOH/ALYRE [illegible] |
| GOBP_ACTION_POTENTIAL_PROPAGATION | GOBP_ACTION_POTENTIAL_PROPAGATION | 10 | -0.8691075 | -1.6678895 | 0.01162435 | 0.16313397 | 0.14931475 | 140 | tags=40%, list=1%, signal=4 | NTRK3/SCN1B/ATP1A2/CLCN1 |
| GOBP_REGULATION_OF_CELL_COMMUNICATION_BY_ELECTRICAL_COU | GOBP_REGULATION_OF_CELL_COMMUNICATION_BY_ELECTRICAL_C | 11 | -0.8522341 | -1.647642 | 0.0116602 | 0.16313397 | 0.14931475 | 104 | tags=27%, list=1%, signal=2 | CALM1/TRDN/HRC |
| GOBP_STRIATED_MUSCLE_ATROPHY | GOBP_STRIATED_MUSCLE_ATROPHY | 4 | -0.9623347 | -1.568238 | 0.01166089 | 0.16313397 | 0.14931475 | 100 | tags=75%, list=1%, signal=7 | GSN/ASB2/TRIM63 |
| GOBP_REGULATION_OF_CHROMATIN_ORGANIZATION | GOBP_REGULATION_OF_CHROMATIN_ORGANIZATION | 30 | 0.61678864 | 1.51176258 | 0.01160477 | 0.16313397 | 0.14931475 | 1261 | tags=20%, list=9%, signal=1 | CDK1/ERI1/CDK2/ERI2/MKI67/H [illegible] |
| GOBP_RESPONSE_TO_PROTOZOAN | GOBP_RESPONSE_TO_PROTOZOAN | 21 | 0.65839245 | 1.50387275 | 0.01158914 | 0.16313397 | 0.14931475 | 3001 | tags=38%, list=21%, signal= | GBP1/MYD88/GBP4/BATF/CLEC7 [illegible] |
| GOBP_POSITIVE_REGULATION_OF_PROTEIN_IMPORT_INTO_NUCLEUS | GOBP_POSITIVE_REGULATION_OF_PROTEIN_IMPORT_INTO_NUCLE | 35 | 0.59624702 | 1.5028857 | 0.01166329 | 0.16313397 | 0.14931475 | 2170 | tags=29%, list=15%, signal= | CDK1/GLI3/SMO/PRKCD/PRKD1/ [illegible] |
| GOBP_REGULATION_OF_ANAPHASE_PROMOTING_COMPLEX_DEPENDE | GOBP_REGULATION_OF_ANAPHASE_PROMOTING_COMPLEX_DEPEN | 6 | 0.84995898 | 1.47910549 | 0.01162115 | 0.16313397 | 0.14931475 | 2002 | tags=83%, list=14%, signal= | PLK1/AKT1/CDK2/FZR1/CDC20 |
| GOBP_DNA_INTEGRITY_CHECKPOINT_SIGNALING | GOBP_DNA_INTEGRITY_CHECKPOINT_SIGNALING | 128 | 0.4551757 | 1.45199644 | 0.01157056 | 0.16313397 | 0.14931475 | 2499 | tags=27%, list=17%, signal= | PLK1/H2AX/CDK1/RHNO1/E2F1/ [illegible] |
| GOBP_ACTIVATION_OF_IMMUNE_RESPONSE | GOBP_ACTIVATION_OF_IMMUNE_RESPONSE | 471 | 0.34464251 | 1.2858434 | 0.0115135 | 0.16313397 | 0.14931475 | 4187 | tags=31%, list=29%, signal= | IRAK4/GBP1/SLC19A1/ARRB2/A [illegible] |
| GOBP_POSITIVE_REGULATION_OF_POTASSIUM_ION_TRANSMEMBRAN | GOBP_POSITIVE_REGULATION_OF_POTASSIUM_ION_TRANSMEMBR | 17 | -0.7842472 | -1.6679853 | 0.01172857 | 0.16357627 | 0.14971958 | 696 | tags=18%, list=5%, signal=1 | ATP1B1/AKAP6/FHL1 |
| GOBP_MYELOID_LEUKOCYTE_MEDIATED_IMMUNITY | GOBP_MYELOID_LEUKOCYTE_MEDIATED_IMMUNITY | 86 | 0.49780352 | 1.49114556 | 0.01175174 | 0.16357627 | 0.14971958 | 3780 | tags=34%, list=26%, signal= | IRAK4/MYD88/SNX4/ADAM17/P [illegible] |
| GOBP_SPHINGOLIPID_METABOLIC_PROCESS | GOBP_SPHINGOLIPID_METABOLIC_PROCESS | 125 | 0.45480982 | 1.44559712 | 0.01176024 | 0.16357627 | 0.14971958 | 4275 | tags=42%, list=29%, signal= | TM9SF2/ELOVL6/HACD3/SAMD8/ [illegible] |
| GOBP_POSITIVE_REGULATION_OF_MUSCLE_ORGAN_DEVELOPMENT | GOBP_POSITIVE_REGULATION_OF_MUSCLE_ORGAN_DEVELOPMENT | 16 | -0.7875855 | -1.6547836 | 0.01183432 | 0.16369719 | 0.14983026 | 252 | tags=38%, list=2%, signal=3 | MTM1/MRTFB/MEF2C/MYOD1/ [illegible] |
| GOBP_ACTIN_POLYMERIZATION_OR_DEPOLYMERIZATION | GOBP_ACTIN_POLYMERIZATION_OR_DEPOLYMERIZATION | 165 | -0.4540172 | -1.4604582 | 0.01182082 | 0.16369719 | 0.14983026 | 641 | tags=10%, list=4%, signal=9 | CCR7/COBL/MSRB2/CARMIL2/L [illegible] |
| GOBP_CELL_CELL_ADHESION | GOBP_CELL_CELL_ADHESION | 712 | 0.31765984 | 1.22485602 | 0.01181361 | 0.16369719 | 0.14983026 | 4408 | tags=33%, list=30%, signal= | PTK7/PLAUR/EFNA5/TWSG1/EXT [illegible] |
| GOBP_DTTP_BIOSYNTHETIC_PROCESS | GOBP_DTTP_BIOSYNTHETIC_PROCESS | 5 | 0.87600388 | 1.47053208 | 0.01194291 | 0.16459298 | 0.15065017 | 390 | tags=60%, list=3%, signal=5 | TYMS/TBPL1/CMPK2 |
| GOBP_POSITIVE_REGULATION_OF_NECROPTOTIC_PROCESS | GOBP_POSITIVE_REGULATION_OF_NECROPTOTIC_PROCESS | 5 | 0.87536917 | 1.4694666 | 0.01194291 | 0.16459298 | 0.15065017 | 1430 | tags=40%, list=10%, signal= | RIPK1/CASP6 |
| GOBP_REPRODUCTIVE_SYSTEM_DEVELOPMENT | GOBP_REPRODUCTIVE_SYSTEM_DEVELOPMENT | 226 | 0.39687452 | 1.37213513 | 0.01201734 | 0.16531541 | 0.1513114 | 4000 | tags=30%, list=27%, signal= | HMGB2/SYCP2/MMP14/STRA6/I [illegible] |
| GOBP_POSITIVE_REGULATION_OF_TUMOR_NECROSIS_FACTOR_SUPERF | GOBP_POSITIVE_REGULATION_OF_TUMOR_NECROSIS_FACTOR_SUP | 105 | 0.47041217 | 1.46146557 | 0.01205266 | 0.16549824 | 0.15147874 | 3589 | tags=36%, list=25%, signal= | EPHB2/MYD88/APP/ADAM8/RIP [illegible] |
| GOBP_GLOMERULAR_MESANGIAL_CELL_DEVELOPMENT | GOBP_GLOMERULAR_MESANGIAL_CELL_DEVELOPMENT | 5 | 0.87462859 | 1.46822341 | 0.0122342 | 0.16768441 | 0.15347972 | 1497 | tags=40%, list=10%, signal= | NOTCH1/PDGFB |
| GOBP_RESPONSE_TO_INTERLEUKIN_17 | GOBP_RESPONSE_TO_INTERLEUKIN_17 | 17 | 0.69246984 | 1.5134216 | 0.01245912 | 0.17014625 | 0.15573302 | 3316 | tags=47%, list=23%, signal= | SRSF1/NOTCH1/CXCL10/IKBKE/T [illegible] |
| GOBP_T_CELL_ACTIVATION | GOBP_T_CELL_ACTIVATION | 464 | 0.34308118 | 1.27814323 | 0.01244988 | 0.17014625 | 0.15573302 | 4580 | tags=35%, list=31%, signal= | TWSG1/CDKN2A/AKT1/ADAM8/ [illegible] |
| GOBP_NEGATIVE_REGULATION_OF_EXTRINSIC_APOPTOTIC_SIGNALING | GOBP_NEGATIVE_REGULATION_OF_EXTRINSIC_APOPTOTIC_SIGNALI | 25 | 0.63513858 | 1.4985424 | 0.01250812 | 0.17050545 | 0.15606179 | 1470 | tags=24%, list=10%, signal= | HMGB2/FAIM/ITPRIP/ARHGEF2/ [illegible] |
| GOBP_NEGATIVE_REGULATION_OF_CYTOPLASMIC_TRANSLATIONAL_IN | GOBP_NEGATIVE_REGULATION_OF_CYTOPLASMIC_TRANSLATIONAL | 4 | -0.9592489 | -1.5632094 | 0.01260637 | 0.17153336 | 0.15700263 | 35 | tags=50%, list=0%, signal=5 | EPRS1/GAPDH |
| GOBP_NEGATIVE_REGULATION_OF_EPITHELIAL_CELL_DIFFERENTIATIO | GOBP_NEGATIVE_REGULATION_OF_EPITHELIAL_CELL_DIFFERENTIA | 31 | 0.61236115 | 1.50974076 | 0.01263088 | 0.17155612 | 0.15702345 | 4055 | tags=48%, list=28%, signal= | EZH2/SMO/NOTCH1/S1PR3/SPRE [illegible] |
| GOBP_DNA_STRAND_ELONGATION | GOBP_DNA_STRAND_ELONGATION | 36 | 0.59044959 | 1.49723084 | 0.01269465 | 0.17211106 | 0.15753138 | 3139 | tags=31%, list=21%, signal= | LIG1/DCLRE1B/GINS1/MCM4/PC [illegible] |
| GOBP_POSITIVE_REGULATION_OF_ORGANELLE_ASSEMBLY | GOBP_POSITIVE_REGULATION_OF_ORGANELLE_ASSEMBLY | 81 | 0.49706262 | 1.47509815 | 0.01276208 | 0.17271346 | 0.15808276 | 2419 | tags=27%, list=17%, signal= | PAN2/FSCN1/STIL/SPAG5/SNX4/ [illegible] |
| GOBP_QUINONE_BIOSYNTHETIC_PROCESS | GOBP_QUINONE_BIOSYNTHETIC_PROCESS | 18 | -0.7722958 | -1.6567089 | 0.01308828 | 0.17405318 | 0.15930899 | 1155 | tags=28%, list=8%, signal=2 | COQ5/STARD7/COQ9/NDUFA9/C [illegible] |
| GOBP_MUSCLE_ATROPHY | GOBP_MUSCLE_ATROPHY | 7 | -0.9064473 | -1.6309402 | 0.01304901 | 0.17405318 | 0.15930899 | 686 | tags=57%, list=5%, signal=5 | GSN/FOXO3/ASB2/TRIM63 |
| GOBP_ENDOPLASMIC_RETICULUM_CALCIUM_ION_HOMEOSTASIS | GOBP_ENDOPLASMIC_RETICULUM_CALCIUM_ION_HOMEOSTASIS | 24 | -0.7175785 | -1.6304837 | 0.01291513 | 0.17405318 | 0.15930899 | 79 | tags=17%, list=1%, signal=1 | WFS1/GRINA/ATP2A2/ATP2A1 |
| GOBP_RESPONSE_TO_ATP | GOBP_RESPONSE_TO_ATP | 24 | -0.7169251 | -1.6289992 | 0.0131787 | 0.17405318 | 0.15930899 | 531 | tags=21%, list=4%, signal=2 | ABCC9/RYR3/P2RX6/P2RX5/KCN [illegible] |
| GOBP_DETECTION_OF_ABIOTIC_STIMULUS | GOBP_DETECTION_OF_ABIOTIC_STIMULUS | 78 | -0.5402429 | -1.5491204 | 0.01317552 | 0.17405318 | 0.15930899 | 190 | tags=6%, list=1%, signal=69 | CSRP3/HPN/TTN/TCAP/CAV3 |
| GOBP_REGULATION_OF_ACTIN_FILAMENT_BUNDLE_ASSEMBLY | GOBP_REGULATION_OF_ACTIN_FILAMENT_BUNDLE_ASSEMBLY | 91 | -0.5212024 | -1.5324881 | 0.01315501 | 0.17405318 | 0.15930899 | 630 | tags=12%, list=4%, signal=1 | AMOT/INPP5K/ARHGEF5/FHOD1/ [illegible] |
| GOBP_DNA_DOUBLE_STRAND_BREAK_PROCESSING | GOBP_DNA_DOUBLE_STRAND_BREAK_PROCESSING | 18 | 0.68332052 | 1.51200412 | 0.0129465 | 0.17405318 | 0.15930899 | 2406 | tags=33%, list=16%, signal= | RBBP8/RNF138/BRIP1/NBN/HEL [illegible] |
| GOBP_REGULATORY_NCRNA_MEDIATED_HETEROCHROMATIN_FORMA | GOBP_REGULATORY_NCRNA_MEDIATED_HETEROCHROMATIN_FOR | 17 | 0.69101908 | 1.5102509 | 0.01308207 | 0.17405318 | 0.15930899 | 1899 | tags=41%, list=13%, signal= | ERI1/EZH2/ERI2/SPOCD1/HNRN [illegible] |
| GOBP_EMBRYONIC_EYE_MORPHOGENESIS | GOBP_EMBRYONIC_EYE_MORPHOGENESIS | 27 | 0.62688566 | 1.5041719 | 0.01313413 | 0.17405318 | 0.15930899 | 1543 | tags=33%, list=11%, signal= | STRA6/FBN2/CITED2/FRS2/FOXL [illegible] |
| GOBP_SIGNALING_RECEPTOR_LIGAND_PRECURSOR_PROCESSING | GOBP_SIGNALING_RECEPTOR_LIGAND_PRECURSOR_PROCESSING | 37 | 0.58739372 | 1.49729839 | 0.013169 | 0.17405318 | 0.15930899 | 4725 | tags=51%, list=32%, signal= | PCSK1N/ADAM17/CTSL/ERO1B/P [illegible] |
| GOBP_FEMALE_GAMETE_GENERATION | GOBP_FEMALE_GAMETE_GENERATION | 112 | 0.46378213 | 1.45159099 | 0.0130988 | 0.17405318 | 0.15930899 | 3022 | tags=30%, list=21%, signal= | PLK1/CCNB2/AKT1/SYCP2/CCNB [illegible] |
| GOBP_NEGATIVE_REGULATION_OF_CELL_ADHESION_MOLECULE_PROD | GOBP_NEGATIVE_REGULATION_OF_CELL_ADHESION_MOLECULE_P | 3 | 0.94679988 | 1.44449321 | 0.0131855 | 0.17405318 | 0.15930899 | 643 | tags=33%, list=4%, signal=3 | NOTCH1 |
| GOBP_NEGATIVE_REGULATION_OF_MITOTIC_CELL_CYCLE | GOBP_NEGATIVE_REGULATION_OF_MITOTIC_CELL_CYCLE | 216 | 0.40106453 | 1.37787774 | 0.01295642 | 0.17405318 | 0.15930899 | 3059 | tags=30%, list=21%, signal= | BTG3/PLK1/CDCA8/CDK1/NUF2/ [illegible] |
| GOBP_REGULATION_OF_CELLULAR_COMPONENT_SIZE | GOBP_REGULATION_OF_CELLULAR_COMPONENT_SIZE | 301 | -0.3976831 | -1.3707243 | 0.01318418 | 0.17405318 | 0.15930899 | 431 | tags=8%, list=3%, signal=89 | PLXNA4/AKT1S1/CCR7/DEPTOR/ [illegible] |
| GOBP_ZYMOSTEROL_METABOLIC_PROCESS | GOBP_ZYMOSTEROL_METABOLIC_PROCESS | 14 | 0.71761658 | 1.50707571 | 0.01327029 | 0.17486513 | 0.16005216 | 1145 | tags=36%, list=8%, signal=3 | HMGCR/LBR/HMGCS1/MSMO1/ [illegible] |
| GOBP_REGULATION_OF_ATTACHMENT_OF_MITOTIC_SPINDLE_MICROT | GOBP_REGULATION_OF_ATTACHMENT_OF_MITOTIC_SPINDLE_MIC | 10 | 0.77080809 | 1.5014487 | 0.013367 | 0.17554972 | 0.16067875 | 1261 | tags=40%, list=9%, signal=3 | CDCA8/CDK1/AURKB/HNRNPU |

| | | | | | | | | | | |
|---|---|---|---|---|---|---|---|---|---|---|
| GOBP_HISTONE_MRNA_METABOLIC_PROCESS | GOBP_HISTONE_MRNA_METABOLIC_PROCESS | 32 | 0.60589418 | 1.50037542 | 0.01336898 | 0.17554972 | 0.16067875 | 1994 | tags=28%, list=14%, signal= | [illegible] |
| GOBP_NEGATIVE_REGULATION_OF_VASCULAR_ASSOCIATED_SMOOTH_ | GOBP_NEGATIVE_REGULATION_OF_VASCULAR_ASSOCIATED_SMOO | 8 | -0.8885409 | -1.6422033 | 0.01347118 | 0.17658293 | 0.16162444 | 252 | tags=25%, list=2%, signal=2 | MEF2C/TPM1 |
| GOBP_POSITIVE_REGULATION_OF_SODIUM_ION_TRANSPORT | GOBP_POSITIVE_REGULATION_OF_SODIUM_ION_TRANSPORT | 27 | -0.7002763 | -1.6334829 | 0.01355152 | 0.17732661 | 0.16230513 | 696 | tags=19%, list=5%, signal=1 | [illegible] |
| GOBP_REGULATION_OF_CARDIAC_MUSCLE_CELL_MEMBRANE_POTEN | GOBP_REGULATION_OF_CARDIAC_MUSCLE_CELL_MEMBRANE_POTI | 7 | -0.9042125 | -1.6269192 | 0.01368555 | 0.17876899 | 0.16362532 | 427 | tags=57%, list=3%, signal=5 | PLN/TRDN/ATP2A2/FXYD1 |
| GOBP_CELL_COMMUNICATION_BY_ELECTRICAL_COUPLING | GOBP_CELL_COMMUNICATION_BY_ELECTRICAL_COUPLING | 28 | -0.6929174 | -1.6307723 | 0.01374396 | 0.17887045 | 0.16371818 | 756 | tags=21%, list=5%, signal=2 | [illegible] |
| GOBP_EMBRYONIC_APPENDAGE_MORPHOGENESIS | GOBP_EMBRYONIC_APPENDAGE_MORPHOGENESIS | 109 | 0.46366764 | 1.447505 | 0.01378026 | 0.17887045 | 0.16371818 | 4196 | tags=44%, list=29%, signal= | [illegible] |
| GOBP_INTEGRIN_MEDIATED_SIGNALING_PATHWAY | GOBP_INTEGRIN_MEDIATED_SIGNALING_PATHWAY | 108 | 0.46332363 | 1.44543945 | 0.0137572 | 0.17887045 | 0.16371818 | 4380 | tags=36%, list=30%, signal= | [illegible] |
| GOBP_CHROMATIN_ORGANIZATION | GOBP_CHROMATIN_ORGANIZATION | 740 | 0.31410943 | 1.21487777 | 0.01378858 | 0.17887045 | 0.16371818 | 3845 | tags=27%, list=26%, signal= | [illegible] |
| GOBP_DOUBLE_STRAND_BREAK_REPAIR_VIA_BREAK_INDUCED_REPLI | GOBP_DOUBLE_STRAND_BREAK_REPAIR_VIA_BREAK_INDUCED_REI | 11 | 0.74750711 | 1.48871845 | 0.01408868 | 0.18153488 | 0.16615691 | 2151 | tags=64%, list=15%, signal= | [illegible] |
| GOBP_CELL_CYCLE_DNA_REPLICATION_INITIATION | GOBP_CELL_CYCLE_DNA_REPLICATION_INITIATION | 6 | 0.84241366 | 1.46597507 | 0.01409064 | 0.18153488 | 0.16615691 | 2151 | tags=83%, list=15%, signal= | [illegible] |
| GOBP_POSITIVE_REGULATION_OF_ENDOTHELIAL_CELL_MATRIX_ADHE | GOBP_POSITIVE_REGULATION_OF_ENDOTHELIAL_CELL_MATRIX_AD | 6 | 0.84169583 | 1.4647259 | 0.01409064 | 0.18153488 | 0.16615691 | 1347 | tags=33%, list=9%, signal=3 | PLPP3/LEF1 |
| GOBP_REGULATION_OF_ENDOTHELIAL_CELL_MATRIX_ADHESION | GOBP_REGULATION_OF_ENDOTHELIAL_CELL_MATRIX_ADHESION | 6 | 0.84169583 | 1.4647259 | 0.01409064 | 0.18153488 | 0.16615691 | 1347 | tags=33%, list=9%, signal=3 | PLPP3/LEF1 |
| GOBP_POSITIVE_REGULATION_OF_SYNAPTIC_TRANSMISSION_CHOLINE | GOBP_POSITIVE_REGULATION_OF_SYNAPTIC_TRANSMISSION_CHOLI | 2 | 0.97557224 | 1.39665669 | 0.01414564 | 0.18193128 | 0.16651973 | 184 | tags=50%, list=1%, signal=4 | APP |
| GOBP_PYRIMIDINE_RIBONUCLEOSIDE_TRIPHOSPHATE_BIOSYNTHETIC_ | GOBP_PYRIMIDINE_RIBONUCLEOSIDE_TRIPHOSPHATE_BIOSYNTHET | 13 | 0.72434047 | 1.49558665 | 0.01424458 | 0.182794 | 0.16730937 | 2528 | tags=46%, list=17%, signal= | [illegible] |
| GOBP_POSITIVE_REGULATION_OF_RESPONSE_TO_EXTERNAL_STIMULU | GOBP_POSITIVE_REGULATION_OF_RESPONSE_TO_EXTERNAL_STIMU | 520 | 0.3361286 | 1.26392756 | 0.01426139 | 0.182794 | 0.16730937 | 3589 | tags=28%, list=25%, signal= | [illegible] |
| GOBP_PYRIMIDINE_NUCLEOSIDE_TRIPHOSPHATE_METABOLIC_PROCES | GOBP_PYRIMIDINE_NUCLEOSIDE_TRIPHOSPHATE_METABOLIC_PRO | 21 | 0.65235923 | 1.4900919 | 0.01428798 | 0.18282289 | 0.16733581 | 3554 | tags=52%, list=24%, signal= | [illegible] |
| GOBP_RESPONSE_TO_FOLLICLE_STIMULATING_HORMONE | GOBP_RESPONSE_TO_FOLLICLE_STIMULATING_HORMONE | 11 | 0.74583787 | 1.48539403 | 0.01438529 | 0.18375493 | 0.1681889 | 1519 | tags=36%, list=10%, signal= | EFNA5/NOTCH1/SRD5A1/POR |
| GOBP_RESPONSE_TO_PURINE_CONTAINING_COMPOUND | GOBP_RESPONSE_TO_PURINE_CONTAINING_COMPOUND | 111 | -0.4965888 | -1.5030524 | 0.01468683 | 0.18726789 | 0.17140427 | 1052 | tags=14%, list=7%, signal=1 | [illegible] |
| GOBP_VACUOLAR_PROTON_TRANSPORTING_V_TYPE_ATPASE_COMPLE | GOBP_VACUOLAR_PROTON_TRANSPORTING_V_TYPE_ATPASE_COMI | 5 | 0.86525642 | 1.4524905 | 0.01471017 | 0.18726789 | 0.17140427 | 480 | tags=40%, list=3%, signal=3 | TM9SF4/ATP6V1B1 |
| GOBP_CELL_CYCLE_G1_S_PHASE_TRANSITION | GOBP_CELL_CYCLE_G1_S_PHASE_TRANSITION | 236 | 0.38776913 | 1.34866359 | 0.01481921 | 0.18833681 | 0.17238264 | 3309 | tags=28%, list=23%, signal= | [illegible] |
| GOBP_T_FOLLICULAR_HELPER_CELL_DIFFERENTIATION | GOBP_T_FOLLICULAR_HELPER_CELL_DIFFERENTIATION | 7 | 0.81721809 | 1.46845094 | 0.0148688 | 0.18857724 | 0.1726027 | 982 | tags=29%, list=7%, signal=2 | TBK1/GPR183 |
| GOBP_RESPONSE_TO_OXYGEN_CONTAINING_COMPOUND | GOBP_RESPONSE_TO_OXYGEN_CONTAINING_COMPOUND | 1250 | -0.3122263 | -1.2084685 | 0.01488834 | 0.18857724 | 0.1726027 | 927 | tags=10%, list=6%, signal=1 | [illegible] |
| GOBP_GLUCAN_BIOSYNTHETIC_PROCESS | GOBP_GLUCAN_BIOSYNTHETIC_PROCESS | 35 | -0.651273 | -1.6127152 | 0.01492903 | 0.18877427 | 0.17278304 | 1052 | tags=40%, list=7%, signal=3 | [illegible] |
| GOBP_RESPONSE_TO_CALCIUM_ION | GOBP_RESPONSE_TO_CALCIUM_ION | 103 | -0.5046079 | -1.5116692 | 0.01500979 | 0.1891586 | 0.17313481 | 756 | tags=12%, list=5%, signal=1 | [illegible] |
| GOBP_MEMBRANE_LIPID_BIOSYNTHETIC_PROCESS | GOBP_MEMBRANE_LIPID_BIOSYNTHETIC_PROCESS | 125 | 0.44945372 | 1.42857294 | 0.01498483 | 0.1891586 | 0.17313481 | 4235 | tags=38%, list=29%, signal= | [illegible] |
| GOBP_V_D_J_RECOMBINATION | GOBP_V_D_J_RECOMBINATION | 17 | 0.68611774 | 1.49953882 | 0.01510668 | 0.19006077 | 0.17396057 | 1347 | tags=18%, list=9%, signal=1 | HMGB2/LIG1/LEF1 |
| GOBP_REGULATION_OF_IMMATURE_T_CELL_PROLIFERATION | GOBP_REGULATION_OF_IMMATURE_T_CELL_PROLIFERATION | 7 | 0.8155386 | 1.46543307 | 0.01516035 | 0.19023542 | 0.17412042 | 285 | tags=29%, list=2%, signal=2 | CDKN2A/RIPK2 |
| GOBP_MULTICELLULAR_ORGANISMAL_REPRODUCTIVE_PROCESS | GOBP_MULTICELLULAR_ORGANISMAL_REPRODUCTIVE_PROCESS | 674 | 0.3197747 | 1.22879031 | 0.01517122 | 0.19023542 | 0.17412042 | 3336 | tags=27%, list=23%, signal= | [illegible] |
| GOBP_EPHRIN_RECEPTOR_SIGNALING_PATHWAY | GOBP_EPHRIN_RECEPTOR_SIGNALING_PATHWAY | 47 | 0.55579056 | 1.4859783 | 0.01524866 | 0.19088777 | 0.17471751 | 2575 | tags=34%, list=18%, signal= | [illegible] |
| GOBP_NEUROMUSCULAR_SYNAPTIC_TRANSMISSION | GOBP_NEUROMUSCULAR_SYNAPTIC_TRANSMISSION | 18 | -0.7587274 | -1.6276023 | 0.01531607 | 0.19141262 | 0.1751979 | 994 | tags=39%, list=7%, signal=3 | [illegible] |
| GOBP_REGULATION_OF_RELAXATION_OF_CARDIAC_MUSCLE | GOBP_REGULATION_OF_RELAXATION_OF_CARDIAC_MUSCLE | 6 | -0.9201847 | -1.6132923 | 0.01539448 | 0.19207303 | 0.17580237 | 427 | tags=67%, list=3%, signal=6 | PDE4D/PLN/AKAP6/HRC |
| GOBP_REGULATION_OF_LYMPHOCYTE_ACTIVATION | GOBP_REGULATION_OF_LYMPHOCYTE_ACTIVATION | 406 | 0.34698527 | 1.27849517 | 0.01542952 | 0.19219098 | 0.17591032 | 4406 | tags=34%, list=30%, signal= | [illegible] |
| GOBP_MOLTING_CYCLE | GOBP_MOLTING_CYCLE | 85 | 0.48898571 | 1.46385382 | 0.01552228 | 0.19302622 | 0.17667481 | 5243 | tags=59%, list=36%, signal= | [illegible] |
| GOBP_NEGATIVE_REGULATION_OF_ATP_DEPENDENT_ACTIVITY | GOBP_NEGATIVE_REGULATION_OF_ATP_DEPENDENT_ACTIVITY | 11 | -0.8438366 | -1.6314069 | 0.01564897 | 0.19428005 | 0.17782242 | 614 | tags=45%, list=4%, signal=4 | OXA1L/PLN/TNNI3/SLN/TNNC1 |
| GOBP_POSITIVE_REGULATION_OF_CELL_CYCLE_G1_S_PHASE_TRANSITI | GOBP_POSITIVE_REGULATION_OF_CELL_CYCLE_G1_S_PHASE_TRANS | 52 | 0.54187248 | 1.4761379 | 0.01569585 | 0.19454047 | 0.17806078 | 4069 | tags=42%, list=28%, signal= | [illegible] |
| GOBP_POSITIVE_REGULATION_OF_CALCIUM_ION_TRANSMEMBRANE_ | GOBP_POSITIVE_REGULATION_OF_CALCIUM_ION_TRANSMEMBRAN | 38 | -0.6337437 | -1.5913165 | 0.01584979 | 0.19612487 | 0.17951096 | 461 | tags=13%, list=3%, signal=1 | [illegible] |
| GOBP_REGULATION_OF_VITAMIN_METABOLIC_PROCESS | GOBP_REGULATION_OF_VITAMIN_METABOLIC_PROCESS | 7 | 0.8140027 | 1.46267322 | 0.01588921 | 0.19628927 | 0.17966144 | 2562 | tags=57%, list=18%, signal= | SNAI2/RDH10/PRMT3/SNAI1 |
| GOBP_HISTONE_MRNA_CATABOLIC_PROCESS | GOBP_HISTONE_MRNA_CATABOLIC_PROCESS | 26 | 0.62554282 | 1.48831999 | 0.01600523 | 0.19708525 | 0.18038999 | 1584 | tags=27%, list=11%, signal= | [illegible] |
| GOBP_CELLULAR_RESPONSE_TO_MACROPHAGE_COLONY_STIMULATIN | GOBP_CELLULAR_RESPONSE_TO_MACROPHAGE_COLONY_STIMULAT | 12 | 0.73084881 | 1.48451538 | 0.01603236 | 0.19708525 | 0.18038999 | 1609 | tags=33%, list=11%, signal= | PTPN2/TLR2/DOK1/CSF1 |
| GOBP_POSITIVE_REGULATION_OF_UBIQUITIN_PROTEIN_LIGASE_ACTIV | GOBP_POSITIVE_REGULATION_OF_UBIQUITIN_PROTEIN_LIGASE_AC | 5 | 0.86250474 | 1.44787131 | 0.01602097 | 0.19708525 | 0.18038999 | 2002 | tags=80%, list=14%, signal= | UBE2C/CDC14B/FZR1/CDC20 |
| GOBP_INTERLEUKIN_6_PRODUCTION | GOBP_INTERLEUKIN_6_PRODUCTION | 117 | 0.45371042 | 1.42828364 | 0.01625709 | 0.19952123 | 0.18261962 | 4833 | tags=42%, list=33%, signal= | [illegible] |
| GOBP_PURKINJE_MYOCYTE_TO_VENTRICULAR_CARDIAC_MUSCLE_CELL | GOBP_PURKINJE_MYOCYTE_TO_VENTRICULAR_CARDIAC_MUSCLE_C | 6 | -0.9166858 | -1.6071579 | 0.01635664 | 0.20041552 | 0.18343815 | 756 | tags=50%, list=5%, signal=4 | GJA5/CASQ2/SCN1B |
| GOBP_IMMUNE_RESPONSE_REGULATING_SIGNALING_PATHWAY | GOBP_IMMUNE_RESPONSE_REGULATING_SIGNALING_PATHWAY | 457 | 0.3400161 | 1.26515891 | 0.01641069 | 0.20075029 | 0.18374456 | 4187 | tags=32%, list=29%, signal= | [illegible] |
| GOBP_REGULATION_OF_DNA_RECOMBINATION | GOBP_REGULATION_OF_DNA_RECOMBINATION | 117 | 0.45297645 | 1.42597309 | 0.01644612 | 0.20085666 | 0.18384192 | 4837 | tags=42%, list=33%, signal= | [illegible] |
| GOBP_POSITIVE_REGULATION_OF_TORC2_SIGNALING | GOBP_POSITIVE_REGULATION_OF_TORC2_SIGNALING | 8 | 0.79811359 | 1.47607061 | 0.01659325 | 0.20232446 | 0.18518538 | 2094 | tags=38%, list=14%, signal= | AKT1/TBK1/EP300 |
| GOBP_NEGATIVE_REGULATION_OF_GENE_SILENCING_BY_REGULATOR | GOBP_NEGATIVE_REGULATION_OF_GENE_SILENCING_BY_REGULAT | 14 | 0.70840748 | 1.48773556 | 0.01662599 | 0.20239517 | 0.18525011 | 896 | tags=29%, list=6%, signal=2 | ADAR/ERI1/ELAVL1/ERI2 |
| GOBP_HAIR_CELL_DIFFERENTIATION | GOBP_HAIR_CELL_DIFFERENTIATION | 40 | 0.56983767 | 1.47422701 | 0.01674927 | 0.20290935 | 0.18572072 | 4864 | tags=57%, list=33%, signal= | [illegible] |
| GOBP_PROTEIN_O_LINKED_GLYCOSYLATION_VIA_FUCOSE | GOBP_PROTEIN_O_LINKED_GLYCOSYLATION_VIA_FUCOSE | 8 | 0.79696858 | 1.47395297 | 0.01674009 | 0.20290935 | 0.18572072 | 1666 | tags=38%, list=11%, signal= | POFUT2/POFUT1/B3GLCT |
| GOBP_REGULATION_OF_MITOCHONDRIAL_ATP_SYNTHESIS_COUPLED_ | GOBP_REGULATION_OF_MITOCHONDRIAL_ATP_SYNTHESIS_COUPLE | 6 | 0.83638163 | 1.45547808 | 0.0167054 | 0.20290935 | 0.18572072 | 223 | tags=33%, list=2%, signal=3 | CDK1/CCNB1 |
| GOBP_POSITIVE_REGULATION_OF_CREB_TRANSCRIPTION_FACTOR_ACT | GOBP_POSITIVE_REGULATION_OF_CREB_TRANSCRIPTION_FACTOR_ | 6 | 0.83598604 | 1.45478968 | 0.01685067 | 0.20380897 | 0.18654414 | 592 | tags=33%, list=4%, signal=3 | ADCY1/CRTC2 |
| GOBP_PROTEIN_O_LINKED_GLYCOSYLATION | GOBP_PROTEIN_O_LINKED_GLYCOSYLATION | 127 | 0.44779708 | 1.42648762 | 0.01692981 | 0.20443696 | 0.18711893 | 3865 | tags=39%, list=26%, signal= | [illegible] |
| GOBP_HEPATOCYTE_GROWTH_FACTOR_RECEPTOR_SIGNALING_PATHW | GOBP_HEPATOCYTE_GROWTH_FACTOR_RECEPTOR_SIGNALING_PAT | 13 | 0.71797715 | 1.48244793 | 0.01697227 | 0.20462072 | 0.18728713 | 2206 | tags=38%, list=15%, signal= | [illegible] |
| GOBP_POSITIVE_REGULATION_OF_HYDROGEN_PEROXIDE_MEDIATED_ | GOBP_POSITIVE_REGULATION_OF_HYDROGEN_PEROXIDE_MEDIATE | 4 | -0.952819 | -1.552731 | 0.01701859 | 0.20485042 | 0.18749737 | 686 | tags=100%, list=5%, signal= | FOXO3/PAWR/ENDOG |
| GOBP_POSITIVE_REGULATION_OF_PROTEIN_K63_LINKED_UBIQUITINA | GOBP_POSITIVE_REGULATION_OF_PROTEIN_K63_LINKED_UBIQUITI | 6 | 0.83450228 | 1.45220762 | 0.0171412 | 0.20599605 | 0.18854595 | 285 | tags=17%, list=2%, signal=1 | RIPK2 |
| GOBP_INNER_MITOCHONDRIAL_MEMBRANE_ORGANIZATION | GOBP_INNER_MITOCHONDRIAL_MEMBRANE_ORGANIZATION | 44 | -0.6064921 | -1.5681213 | 0.0172043 | 0.20642413 | 0.18893777 | 1974 | tags=34%, list=13%, signal= | [illegible] |
| GOBP_REGULATION_OF_SIGNAL_TRANSDUCTION_BY_P53_CLASS_MED | GOBP_REGULATION_OF_SIGNAL_TRANSDUCTION_BY_P53_CLASS_M | 115 | 0.45463876 | 1.42844415 | 0.01727375 | 0.20692687 | 0.18939792 | 4613 | tags=37%, list=32%, signal= | [illegible] |
| GOBP_STORE_OPERATED_CALCIUM_ENTRY | GOBP_STORE_OPERATED_CALCIUM_ENTRY | 21 | -0.7312288 | -1.6188047 | 0.01755334 | 0.20927476 | 0.19154692 | 1153 | tags=33%, list=8%, signal=3 | [illegible] |
| GOBP_SYNCYTIUM_FORMATION | GOBP_SYNCYTIUM_FORMATION | 91 | -0.5099524 | -1.4994099 | 0.01754001 | 0.20927476 | 0.19154692 | 618 | tags=8%, list=4%, signal=79 | [illegible] |
| GOBP_REGULATION_OF_RESPONSE_TO_EXTERNAL_STIMULUS | GOBP_REGULATION_OF_RESPONSE_TO_EXTERNAL_STIMULUS | 869 | 0.30465844 | 1.19245748 | 0.01754386 | 0.20927476 | 0.19154692 | 4278 | tags=32%, list=29%, signal= | [illegible] |
| GOBP_REGULATION_OF_SPONTANEOUS_SYNAPTIC_TRANSMISSION | GOBP_REGULATION_OF_SPONTANEOUS_SYNAPTIC_TRANSMISSION | 5 | 0.85945279 | 1.44274806 | 0.01762307 | 0.20977319 | 0.19200312 | 184 | tags=20%, list=1%, signal=2 | APP |
| GOBP_MYELOID_LEUKOCYTE_DIFFERENTIATION | GOBP_MYELOID_LEUKOCYTE_DIFFERENTIATION | 204 | 0.39828902 | 1.35644383 | 0.01771305 | 0.21051061 | 0.19267808 | 2808 | tags=25%, list=19%, signal= | [illegible] |
| GOBP_NEGATIVE_REGULATION_OF_STRIATED_MUSCLE_CELL_DIFFEREI | GOBP_NEGATIVE_REGULATION_OF_STRIATED_MUSCLE_CELL_DIFFE | 34 | -0.6471219 | -1.5894364 | 0.0177471 | 0.21055299 | 0.19271687 | 836 | tags=24%, list=6%, signal=2 | [illegible] |
| GOBP_RETROGRADE_NEURONAL_DENSE_CORE_VESICLE_TRANSPORT | GOBP_RETROGRADE_NEURONAL_DENSE_CORE_VESICLE_TRANSPOR | 4 | -0.9519068 | -1.5512446 | 0.01796407 | 0.21055299 | 0.19271687 | 313 | tags=50%, list=2%, signal=4 | KIF1C/KIF1B |
| GOBP_RESPONSE_TO_ALKALOID | GOBP_RESPONSE_TO_ALKALOID | 70 | -0.5439074 | -1.5314284 | 0.01794639 | 0.21055299 | 0.19271687 | 782 | tags=14%, list=5%, signal=1 | [illegible] |
| GOBP_MATING_BEHAVIOR | GOBP_MATING_BEHAVIOR | 24 | 0.62917291 | 1.47365321 | 0.01788015 | 0.21055299 | 0.19271687 | 3090 | tags=33%, list=21%, signal= | [illegible] |
| GOBP_SPINAL_CORD_PATTERNING | GOBP_SPINAL_CORD_PATTERNING | 11 | 0.738452 | 1.47068449 | 0.01794454 | 0.21055299 | 0.19271687 | 1663 | tags=36%, list=11%, signal= | GLI3/SMO/GLI2/GDF11 |
| GOBP_OSTEOBLAST_DIFFERENTIATION | GOBP_OSTEOBLAST_DIFFERENTIATION | 200 | 0.39902094 | 1.35553873 | 0.01785014 | 0.21055299 | 0.19271687 | 4143 | tags=35%, list=28%, signal= | [illegible] |
| GOBP_IMMUNE_RESPONSE_REGULATING_CELL_SURFACE_RECEPTOR_S | GOBP_IMMUNE_RESPONSE_REGULATING_CELL_SURFACE_RECEPTOI | 289 | 0.36843717 | 1.31162331 | 0.01796891 | 0.21055299 | 0.19271687 | 3854 | tags=30%, list=26%, signal= | [illegible] |
| GOBP_ENDOMEMBRANE_SYSTEM_ORGANIZATION | GOBP_ENDOMEMBRANE_SYSTEM_ORGANIZATION | 520 | -0.3559472 | -1.2951952 | 0.01791101 | 0.21055299 | 0.19271687 | 928 | tags=8%, list=6%, signal=89 | [illegible] |
| GOBP_CELL_MIGRATION | GOBP_CELL_MIGRATION | 1237 | 0.29245458 | 1.16758752 | 0.01779524 | 0.21055299 | 0.19271687 | 3172 | tags=24%, list=22%, signal= | [illegible] |
| GOBP_ETHER_CATABOLIC_PROCESS | GOBP_ETHER_CATABOLIC_PROCESS | 4 | 0.89710376 | 1.44073075 | 0.01801142 | 0.21072241 | 0.19287194 | 1153 | tags=25%, list=8%, signal=2 | PLA2G7 |
| GOBP_MONOSACCHARIDE_TRANSMEMBRANE_TRANSPORT | GOBP_MONOSACCHARIDE_TRANSMEMBRANE_TRANSPORT | 86 | -0.5139353 | -1.4985126 | 0.01821773 | 0.21280468 | 0.19477781 | 1235 | tags=17%, list=8%, signal=1 | [illegible] |
| GOBP_REGULATION_OF_TRNA_METABOLIC_PROCESS | GOBP_REGULATION_OF_TRNA_METABOLIC_PROCESS | 6 | 0.83230394 | 1.44838206 | 0.01830331 | 0.21347232 | 0.1953889 | 409 | tags=33%, list=3%, signal=3 | AKT1/NSUN2 |
| GOBP_LIPOPOLYSACCHARIDE_MEDIATED_SIGNALING_PATHWAY | GOBP_LIPOPOLYSACCHARIDE_MEDIATED_SIGNALING_PATHWAY | 54 | 0.53377022 | 1.46526745 | 0.01833741 | 0.21353841 | 0.19544939 | 4786 | tags=43%, list=33%, signal= | [illegible] |
| GOBP_SPERMATID_NUCLEUS_DIFFERENTIATION | GOBP_SPERMATID_NUCLEUS_DIFFERENTIATION | 16 | 0.68723894 | 1.48229744 | 0.01844104 | 0.21383836 | 0.19572393 | 2525 | tags=38%, list=17%, signal= | [illegible] |
| GOBP_PROTEIN_AUTO_ADP_RIBOSYLATION | GOBP_PROTEIN_AUTO_ADP_RIBOSYLATION | 12 | 0.72476893 | 1.47216579 | 0.01842973 | 0.21383836 | 0.19572393 | 2250 | tags=17%, list=15%, signal= | PARP16/PARP11 |
| GOBP_PATTERN_SPECIFICATION_INVOLVED_IN_KIDNEY_DEVELOPMEN | GOBP_PATTERN_SPECIFICATION_INVOLVED_IN_KIDNEY_DEVELOPM | 6 | 0.83173338 | 1.44738916 | 0.01844858 | 0.21383836 | 0.19572393 | 630 | tags=33%, list=4%, signal=3 | FOXD1/IRX1 |
| GOBP_ACTIVATION_OF_INNATE_IMMUNE_RESPONSE | GOBP_ACTIVATION_OF_INNATE_IMMUNE_RESPONSE | 271 | 0.37283598 | 1.31785779 | 0.01847804 | 0.21384992 | 0.19573451 | 3575 | tags=30%, list=24%, signal= | [illegible] |
| GOBP_REGULATION_OF_FC_RECEPTOR_MEDIATED_STIMULATORY_SIG | GOBP_REGULATION_OF_FC_RECEPTOR_MEDIATED_STIMULATORY_ | 11 | 0.73787993 | 1.46954518 | 0.01853774 | 0.21421075 | 0.19606477 | 1645 | tags=36%, list=11%, signal= | CD47/LYN/PTPRJ/PLSCR1 |
| GOBP_NEGATIVE_REGULATION_OF_GROWTH | GOBP_NEGATIVE_REGULATION_OF_GROWTH | 184 | -0.4309498 | -1.4084042 | 0.0186399 | 0.21506037 | 0.19684243 | 716 | tags=8%, list=5%, signal=89 | [illegible] |
| GOBP_NEGATIVE_REGULATION_OF_COAGULATION | GOBP_NEGATIVE_REGULATION_OF_COAGULATION | 32 | 0.59555498 | 1.47477246 | 0.01888369 | 0.21739737 | 0.19898145 | 3597 | tags=50%, list=25%, signal= | [illegible] |
| GOBP_TUMOR_NECROSIS_FACTOR_SUPERFAMILY_CYTOKINE_PRODUCT | GOBP_TUMOR_NECROSIS_FACTOR_SUPERFAMILY_CYTOKINE_PRODI | 156 | 0.42583264 | 1.39721719 | 0.01890034 | 0.21739737 | 0.19898145 | 4453 | tags=40%, list=30%, signal= | [illegible] |
| GOBP_NEGATIVE_ADAPTATION_OF_SIGNALING_PATHWAY | GOBP_NEGATIVE_ADAPTATION_OF_SIGNALING_PATHWAY | 13 | 0.71315813 | 1.47249783 | 0.01894226 | 0.2175464 | 0.19911786 | 2076 | tags=46%, list=14%, signal= | [illegible] |
| GOBP_REGULATION_OF_CELL_POPULATION_PROLIFERATION | GOBP_REGULATION_OF_CELL_POPULATION_PROLIFERATION | 1268 | 0.29037899 | 1.16111162 | 0.01917302 | 0.21986038 | 0.20123582 | 4055 | tags=29%, list=28%, signal= | [illegible] |
| GOBP_POSITIVE_REGULATION_OF_INTERLEUKIN_6_PRODUCTION | GOBP_POSITIVE_REGULATION_OF_INTERLEUKIN_6_PRODUCTION | 79 | 0.49059004 | 1.45020777 | 0.01922728 | 0.22014651 | 0.20149771 | 3589 | tags=35%, list=25%, signal= | [illegible] |
| GOBP_TISSUE_REMODELING | GOBP_TISSUE_REMODELING | 140 | 0.43199641 | 1.39775581 | 0.0193475 | 0.22118576 | 0.20244893 | 4115 | tags=39%, list=28%, signal= | [illegible] |
| GOBP_NEGATIVE_REGULATION_OF_PROTEIN_POLYMERIZATION | GOBP_NEGATIVE_REGULATION_OF_PROTEIN_POLYMERIZATION | 61 | -0.5611253 | -1.5419821 | 0.01937716 | 0.22118825 | 0.20245121 | 431 | tags=15%, list=3%, signal=1 | [illegible] |
| GOBP_CELL_COMMUNICATION_BY_ELECTRICAL_COUPLING_INVOLVED_ | GOBP_CELL_COMMUNICATION_BY_ELECTRICAL_COUPLING_INVOLV | 22 | -0.7148014 | -1.5971267 | 0.01952394 | 0.22145116 | 0.20269184 | 696 | tags=23%, list=5%, signal=2 | [illegible] |
| GOBP_POSITIVE_REGULATION_OF_VASOCONSTRICTION | GOBP_POSITIVE_REGULATION_OF_VASOCONSTRICTION | 17 | -0.7501502 | -1.5954657 | 0.01954761 | 0.22145116 | 0.20269184 | 2 | tags=12%, list=0%, signal=1 | AVPR1A/SMTNL1 |
| GOBP_REGULATION_OF_SKELETAL_MUSCLE_CONTRACTION_BY_REGUL | GOBP_REGULATION_OF_SKELETAL_MUSCLE_CONTRACTION_BY_REG | 5 | -0.9325296 | -1.5835935 | 0.01945153 | 0.22145116 | 0.20269184 | 172 | tags=60%, list=1%, signal=5 | GSTO1/DMD/CASQ1 |
| GOBP_INDOLE_CONTAINING_COMPOUND_CATABOLIC_PROCESS | GOBP_INDOLE_CONTAINING_COMPOUND_CATABOLIC_PROCESS | 8 | 0.79061515 | 1.46220263 | 0.0195301 | 0.22145116 | 0.20269184 | 2463 | tags=50%, list=17%, signal= | AFMID/TDO2/IDO1/IL4I1 |
| GOBP_POSITIVE_REGULATION_OF_OXIDOREDUCTASE_ACTIVITY | GOBP_POSITIVE_REGULATION_OF_OXIDOREDUCTASE_ACTIVITY | 4 | 0.89378231 | 1.43539657 | 0.01947577 | 0.22145116 | 0.20269184 | 575 | tags=25%, list=4%, signal=2 | RFK |
| GOBP_SPHINGOID_METABOLIC_PROCESS | GOBP_SPHINGOID_METABOLIC_PROCESS | 18 | 0.66745394 | 1.47689566 | 0.01965372 | 0.22231791 | 0.20348517 | 3972 | tags=61%, list=27%, signal= | [illegible] |
| GOBP_SPINDLE_LOCALIZATION | GOBP_SPINDLE_LOCALIZATION | 55 | 0.528629 | 1.45747461 | 0.01975179 | 0.22309129 | 0.20419304 | 2592 | tags=35%, list=18%, signal= | [illegible] |
| GOBP_POSITIVE_REGULATION_OF_TRANSLATIONAL_ELONGATION | GOBP_POSITIVE_REGULATION_OF_TRANSLATIONAL_ELONGATION | 5 | 0.85701608 | 1.4386576 | 0.01980775 | 0.22338738 | 0.20446405 | 703 | tags=20%, list=5%, signal=1 | EIF5A2 |
| GOBP_MIDBRAIN_DEVELOPMENT | GOBP_MIDBRAIN_DEVELOPMENT | 72 | -0.5355312 | -1.5140118 | 0.02011754 | 0.226541 | 0.20735052 | 844 | tags=14%, list=6%, signal=1 | [illegible] |
| GOBP_REGULATION_OF_SMOOTHENED_SIGNALING_PATHWAY | GOBP_REGULATION_OF_SMOOTHENED_SIGNALING_PATHWAY | 82 | 0.4850572 | 1.44246751 | 0.02027768 | 0.22800244 | 0.20868816 | 3986 | tags=37%, list=27%, signal= | [illegible] |
| GOBP_HEART_TRABECULA_FORMATION | GOBP_HEART_TRABECULA_FORMATION | 12 | -0.8170651 | -1.6114068 | 0.02043269 | 0.22896938 | 0.20957319 | 807 | tags=25%, list=6%, signal=2 | HEY2/EGLN1/CAV3 |
| GOBP_CELLULAR_RESPONSE_TO_ALKALOID | GOBP_CELLULAR_RESPONSE_TO_ALKALOID | 27 | -0.6804866 | -1.587321 | 0.02045513 | 0.22896938 | 0.20957319 | 782 | tags=22%, list=5%, signal=2 | [illegible] |
| GOBP_LEUKOCYTE_PROLIFERATION | GOBP_LEUKOCYTE_PROLIFERATION | 247 | 0.37676559 | 1.31762191 | 0.02041221 | 0.22896938 | 0.20957319 | 4888 | tags=39%, list=33%, signal= | [illegible] |
| GOBP_TRICARBOXYLIC_ACID_CYCLE | GOBP_TRICARBOXYLIC_ACID_CYCLE | 30 | -0.6590751 | -1.5722638 | 0.02065491 | 0.23086167 | 0.21130518 | 1352 | tags=40%, list=9%, signal=3 | [illegible] |
| GOBP_DNA_DAMAGE_RESPONSE_SIGNAL_TRANSDUCTION_BY_P53_CL | GOBP_DNA_DAMAGE_RESPONSE_SIGNAL_TRANSDUCTION_BY_P53_ | 75 | 0.49415093 | 1.4500277 | 0.02070953 | 0.2309382 | 0.21137523 | 4349 | tags=44%, list=30%, signal= | [illegible] |
| GOBP_REGULATION_OF_CYTOSKELETON_ORGANIZATION | GOBP_REGULATION_OF_CYTOSKELETON_ORGANIZATION | 475 | -0.3593753 | -1.2963274 | 0.020754 | 0.2309382 | 0.21137523 | 716 | tags=8%, list=5%, signal=79 | [illegible] |
| GOBP_ADAPTIVE_IMMUNE_RESPONSE | GOBP_ADAPTIVE_IMMUNE_RESPONSE | 346 | 0.35244651 | 1.27983531 | 0.02074014 | 0.2309382 | 0.21137523 | 4875 | tags=38%, list=33%, signal= | [illegible] |
| GOBP_2_OXOGLUTARATE_METABOLIC_PROCESS | GOBP_2_OXOGLUTARATE_METABOLIC_PROCESS | 16 | -0.7566021 | -1.5896847 | 0.02085094 | 0.23167373 | 0.21204845 | 466 | tags=44%, list=3%, signal=4 | [illegible] |
| GOBP_POSITIVE_REGULATION_OF_INTERFERON_BETA_PRODUCTION | GOBP_POSITIVE_REGULATION_OF_INTERFERON_BETA_PRODUCTIOI | 42 | 0.55789218 | 1.45607362 | 0.0209514 | 0.23244603 | 0.21275533 | 5017 | tags=43%, list=34%, signal= | [illegible] |
| GOBP_REGULATION_OF_CELL_ACTIVATION | GOBP_REGULATION_OF_CELL_ACTIVATION | 503 | 0.33274714 | 1.24866095 | 0.02098347 | 0.23245844 | 0.21276669 | 4344 | tags=34%, list=30%, signal= | [illegible] |
| GOBP_MYELOID_DENDRITIC_CELL_CHEMOTAXIS | GOBP_MYELOID_DENDRITIC_CELL_CHEMOTAXIS | 5 | -0.9311811 | -1.5813037 | 0.02104592 | 0.23246455 | 0.21277228 | 847 | tags=60%, list=6%, signal=5 | ARHGEF5/CCR7/CCL21 |
| GOBP_TISSUE_REGENERATION | GOBP_TISSUE_REGENERATION | 55 | -0.5656826 | -1.5263398 | 0.02102313 | 0.23246455 | 0.21277228 | 257 | tags=16%, list=2%, signal=1 | [illegible] |
| GOBP_MITOCHONDRION_LOCALIZATION | GOBP_MITOCHONDRION_LOCALIZATION | 49 | -0.5843644 | -1.5459772 | 0.02118644 | 0.23278871 | 0.21306898 | 743 | tags=16%, list=5%, signal=1 | [illegible] |
| GOBP_SNRNA_3_END_PROCESSING | GOBP_SNRNA_3_END_PROCESSING | 22 | 0.63486461 | 1.4623961 | 0.02123583 | 0.23278871 | 0.21306898 | 1975 | tags=27%, list=13%, signal= | [illegible] |
| GOBP_VENTRICULAR_CARDIAC_MUSCLE_CELL_DEVELOPMENT | GOBP_VENTRICULAR_CARDIAC_MUSCLE_CELL_DEVELOPMENT | 8 | 0.78721012 | 1.45590519 | 0.02129222 | 0.23278871 | 0.21306898 | 380 | tags=38%, list=3%, signal=3 | CDK1/CCNB1/MYH10 |
| GOBP_SPINAL_CORD_DEVELOPMENT | GOBP_SPINAL_CORD_DEVELOPMENT | 70 | 0.50035217 | 1.44715755 | 0.02125 | 0.23278871 | 0.21306898 | 2945 | tags=34%, list=20%, signal= | [illegible] |
| GOBP_HEMATOPOIETIC_STEM_CELL_MIGRATION_TO_BONE_MARROW | GOBP_HEMATOPOIETIC_STEM_CELL_MIGRATION_TO_BONE_MARR | 5 | 0.85555575 | 1.43620616 | 0.0212642 | 0.23278871 | 0.21306898 | 126 | tags=20%, list=1%, signal=2 | EXT1 |
| GOBP_HEMATOPOIETIC_PROGENITOR_CELL_DIFFERENTIATION | GOBP_HEMATOPOIETIC_PROGENITOR_CELL_DIFFERENTIATION | 110 | 0.4511221 | 1.41102498 | 0.0211542 | 0.23278871 | 0.21306898 | 4884 | tags=39%, list=33%, signal= | [illegible] |
| GOBP_REGULATION_OF_DNA_BINDING_TRANSCRIPTION_FACTOR_ACT | GOBP_REGULATION_OF_DNA_BINDING_TRANSCRIPTION_FACTOR_A | 137 | 0.43422348 | 1.39924127 | 0.02128064 | 0.23278871 | 0.21306898 | 4453 | tags=37%, list=30%, signal= | [illegible] |
| GOBP_REGULATION_OF_MEMBRANE_DEPOLARIZATION_DURING_CARI | GOBP_REGULATION_OF_MEMBRANE_DEPOLARIZATION_DURING_C | 5 | -0.930577 | -1.5802777 | 0.0213648 | 0.23324271 | 0.21348452 | 36 | tags=40%, list=0%, signal=4 | ANK3/CAV3 |
| GOBP_FEMALE_SEX_DIFFERENTIATION | GOBP_FEMALE_SEX_DIFFERENTIATION | 95 | 0.46719209 | 1.42867165 | 0.0214325 | 0.23364221 | 0.21385018 | 3997 | tags=33%, list=27%, signal= | [illegible] |
| GOBP_ACTIN_FILAMENT_POLYMERIZATION | GOBP_ACTIN_FILAMENT_POLYMERIZATION | 139 | -0.45537 | -1.4290637 | 0.02150538 | 0.23409693 | 0.21426638 | 641 | tags=10%, list=4%, signal=1 | [illegible] |
| GOBP_REGULATION_OF_D_GLUCOSE_IMPORT | GOBP_REGULATION_OF_D_GLUCOSE_IMPORT | 43 | -0.6021751 | -1.5511385 | 0.02175472 | 0.23646847 | 0.21643703 | 1235 | tags=16%, list=8%, signal=1 | [illegible] |
| GOBP_MODIFIED_AMINO_ACID_BIOSYNTHETIC_PROCESS | GOBP_MODIFIED_AMINO_ACID_BIOSYNTHETIC_PROCESS | 37 | -0.6243505 | -1.5600665 | 0.02181907 | 0.23682196 | 0.21676057 | 927 | tags=22%, list=6%, signal=2 | [illegible] |
| GOBP_ORGANELLE_INHERITANCE | GOBP_ORGANELLE_INHERITANCE | 16 | 0.68119048 | 1.46925157 | 0.0218503 | 0.23682196 | 0.21676057 | 687 | tags=25%, list=5%, signal=2 | PLK1/CDK1/PLK3/YWHAZ |
| GOBP_RESPONSE_TO_AMINE | GOBP_RESPONSE_TO_AMINE | 34 | 0.58370643 | 1.4622979 | 0.02203532 | 0.23848317 | 0.21828106 | 3090 | tags=32%, list=21%, signal= | [illegible] |
| GOBP_MOLTING_CYCLE_PROCESS | GOBP_MOLTING_CYCLE_PROCESS | 76 | 0.49162396 | 1.44586021 | 0.02214221 | 0.23929521 | 0.21902431 | 5003 | tags=55%, list=34%, signal= | [illegible] |
| GOBP_CYTOKINESIS | GOBP_CYTOKINESIS | 168 | 0.41183148 | 1.3672345 | 0.02227343 | 0.24002252 | 0.21969001 | 2439 | tags=26%, list=17%, signal= | [illegible] |
| GOBP_LYMPHOCYTE_DIFFERENTIATION | GOBP_LYMPHOCYTE_DIFFERENTIATION | 359 | 0.3482405 | 1.26747682 | 0.02224511 | 0.24002252 | 0.21969001 | 4184 | tags=33%, list=29%, signal= | [illegible] |
| GOBP_CYTOSKELETON_DEPENDENT_CYTOKINESIS | GOBP_CYTOSKELETON_DEPENDENT_CYTOKINESIS | 119 | 0.44611405 | 1.40750612 | 0.02231046 | 0.24007712 | 0.21973999 | 2439 | tags=28%, list=17%, signal= | [illegible] |
| GOBP_DEVELOPMENTAL_PROCESS_INVOLVED_IN_REPRODUCTION | GOBP_DEVELOPMENTAL_PROCESS_INVOLVED_IN_REPRODUCTION | 717 | 0.31129146 | 1.20161624 | 0.02249891 | 0.24175865 | 0.22127907 | 4000 | tags=30%, list=27%, signal= | [illegible] |
| GOBP_EMBRYONIC_SKELETAL_SYSTEM_DEVELOPMENT | GOBP_EMBRYONIC_SKELETAL_SYSTEM_DEVELOPMENT | 109 | 0.45057173 | 1.4066214 | 0.02290503 | 0.24577095 | 0.22495148 | 4201 | tags=37%, list=29%, signal= | [illegible] |
| GOBP_NEGATIVE_REGULATION_OF_PROTEIN_LOCALIZATION_TO_CELL_ | GOBP_NEGATIVE_REGULATION_OF_PROTEIN_LOCALIZATION_TO_CE | 12 | -0.8134725 | -1.6043216 | 0.02313702 | 0.24704935 | 0.2261216 | 151 | tags=25%, list=1%, signal=2 | GPM6B/ACTN2/CAV3 |
| GOBP_POSITIVE_REGULATION_OF_SMOOTHENED_SIGNALING_PATHW | GOBP_POSITIVE_REGULATION_OF_SMOOTHENED_SIGNALING_PATH | 33 | 0.58484309 | 1.4579489 | 0.02310784 | 0.24704935 | 0.2261216 | 2817 | tags=33%, list=19%, signal= | [illegible] |
| GOBP_RNA_METHYLATION | GOBP_RNA_METHYLATION | 86 | 0.47648093 | 1.42727483 | 0.02313625 | 0.24704935 | 0.2261216 | 4241 | tags=34%, list=29%, signal= | [illegible] |
| GOBP_REGULATION_OF_APOPTOTIC_SIGNALING_PATHWAY | GOBP_REGULATION_OF_APOPTOTIC_SIGNALING_PATHWAY | 332 | 0.35140421 | 1.2693583 | 0.02315574 | 0.24704935 | 0.2261216 | 3626 | tags=27%, list=25%, signal= | [illegible] |
| GOBP_POSITIVE_REGULATION_OF_NUCLEAR_DIVISION | GOBP_POSITIVE_REGULATION_OF_NUCLEAR_DIVISION | 43 | 0.55397563 | 1.4531157 | 0.02319986 | 0.24716903 | 0.22623113 | 2002 | tags=33%, list=14%, signal= | [illegible] |
| GOBP_PRONEPHROS_DEVELOPMENT | GOBP_PRONEPHROS_DEVELOPMENT | 5 | 0.85170334 | 1.4297392 | 0.02330323 | 0.24721834 | 0.22627627 | 167 | tags=20%, list=1%, signal=2 | OSR2 |
| GOBP_INDOLE_CONTAINING_COMPOUND_BIOSYNTHETIC_PROCESS | GOBP_INDOLE_CONTAINING_COMPOUND_BIOSYNTHETIC_PROCESS | 1 | -0.9891364 | -1.3206131 | 0.02329254 | 0.24721834 | 0.22627627 | 161 | tags=100%, list=1%, signal=99% | |
| GOBP_SEROTONIN_BIOSYNTHETIC_PROCESS | GOBP_SEROTONIN_BIOSYNTHETIC_PROCESS | 1 | -0.9891364 | -1.3206131 | 0.02329254 | 0.24721834 | 0.22627627 | 161 | tags=100%, list=1%, signal=99% | |
| GOBP_POSITIVE_REGULATION_OF_CAMP_PKA_SIGNAL_TRANSDUCTIO | GOBP_POSITIVE_REGULATION_OF_CAMP_PKA_SIGNAL_TRANSDUCT | 12 | -0.8121682 | -1.6017493 | 0.0234375 | 0.24829205 | 0.22725902 | 408 | tags=25%, list=3%, signal=2 | UCN/TTN/MYOM1 |
| GOBP_INNER_EAR_AUDITORY_RECEPTOR_CELL_DIFFERENTIATION | GOBP_INNER_EAR_AUDITORY_RECEPTOR_CELL_DIFFERENTIATION | 32 | 0.58898891 | 1.45851292 | 0.02372995 | 0.2510361 | 0.22977062 | 4864 | tags=56%, list=33%, signal= | [illegible] |
| GOBP_REGULATION_OF_PROTEIN_DEUBIQUITINATION | GOBP_REGULATION_OF_PROTEIN_DEUBIQUITINATION | 11 | -0.8299477 | -1.6045553 | 0.02393372 | 0.25256749 | 0.23117228 | 2022 | tags=45%, list=14%, signal= | [illegible] |
| GOBP_KETONE_BIOSYNTHETIC_PROCESS | GOBP_KETONE_BIOSYNTHETIC_PROCESS | 39 | -0.6095685 | -1.5406077 | 0.02394196 | 0.25256749 | 0.23117228 | 1155 | tags=18%, list=8%, signal=1 | [illegible] |
| GOBP_POSITIVE_REGULATION_OF_DENDRITIC_CELL_ANTIGEN_PROCES | GOBP_POSITIVE_REGULATION_OF_DENDRITIC_CELL_ANTIGEN_PRO | 6 | -0.9072253 | -1.5905715 | 0.02405388 | 0.252887 | 0.23146473 | 641 | tags=50%, list=4%, signal=4 | SLC11A1/CCR7/CCL21 |
| GOBP_NOTCH_RECEPTOR_PROCESSING | GOBP_NOTCH_RECEPTOR_PROCESSING | 9 | 0.76276127 | 1.44891321 | 0.02407325 | 0.252887 | 0.23146473 | 2211 | tags=56%, list=15%, signal= | [illegible] |
| GOBP_REGULATION_OF_T_CELL_ACTIVATION | GOBP_REGULATION_OF_T_CELL_ACTIVATION | 300 | 0.36132383 | 1.29424911 | 0.02406739 | 0.252887 | 0.23146473 | 4406 | tags=36%, list=30%, signal= | [illegible] |
| GOBP_AXONEMAL_DYNEIN_COMPLEX_ASSEMBLY | GOBP_AXONEMAL_DYNEIN_COMPLEX_ASSEMBLY | 18 | 0.65958267 | 1.45947866 | 0.0241772 | 0.25362418 | 0.23213946 | 4619 | tags=83%, list=32%, signal= | [illegible] |
| GOBP_NEURON_FATE_DETERMINATION | GOBP_NEURON_FATE_DETERMINATION | 3 | 0.92691234 | 1.41415161 | 0.0242733 | 0.25398803 | 0.23247249 | 306 | tags=33%, list=2%, signal=3 | CDC42 |
| GOBP_NEGATIVE_REGULATION_OF_MYOBLAST_PROLIFERATION | GOBP_NEGATIVE_REGULATION_OF_MYOBLAST_PROLIFERATION | 1 | -0.9887948 | -1.320157 | 0.02427951 | 0.25398803 | 0.23247249 | 166 | tags=100%, list=1%, signal=99% | |
| GOBP_VASCULAR_PROCESS_IN_CIRCULATORY_SYSTEM | GOBP_VASCULAR_PROCESS_IN_CIRCULATORY_SYSTEM | 206 | -0.4134155 | -1.3716885 | 0.02443457 | 0.25525462 | 0.23363178 | 642 | tags=7%, list=4%, signal=79 | [illegible] |
| GOBP_DETOXIFICATION_OF_CADMIUM_ION | GOBP_DETOXIFICATION_OF_CADMIUM_ION | 4 | 0.88562434 | 1.42229503 | 0.02460097 | 0.25663592 | 0.23489607 | 514 | tags=50%, list=4%, signal=4 | SLC39A8/SLC30A1 |
| GOBP_MUSCLE_CELL_FATE_COMMITMENT | GOBP_MUSCLE_CELL_FATE_COMMITMENT | 12 | -0.8103492 | -1.5981618 | 0.02463942 | 0.25668059 | 0.23493696 | 309 | tags=42%, list=2%, signal=4 | [illegible] |
| GOBP_NEGATIVE_REGULATION_OF_VASCULAR_ASSOCIATED_SMOOTH_ | GOBP_NEGATIVE_REGULATION_OF_VASCULAR_ASSOCIATED_SMOO | 19 | -0.7285606 | -1.5787698 | 0.02474567 | 0.25743037 | 0.23562322 | 252 | tags=21%, list=2%, signal=2 | CNN1/RGS5/MEF2C/TPM1 |
| GOBP_TELOMERE_ORGANIZATION | GOBP_TELOMERE_ORGANIZATION | 162 | 0.41333203 | 1.36447623 | 0.02491376 | 0.25882054 | 0.23689564 | 4027 | tags=29%, list=28%, signal= | [illegible] |
| GOBP_EPITHELIAL_CELL_PROLIFERATION | GOBP_EPITHELIAL_CELL_PROLIFERATION | 321 | 0.35223158 | 1.26912869 | 0.02509692 | 0.2603632 | 0.23830762 | 3589 | tags=28%, list=25%, signal= | [illegible] |
| GOBP_CHONDROITIN_SULFATE_PROTEOGLYCAN_METABOLIC_PROCESS | GOBP_CHONDROITIN_SULFATE_PROTEOGLYCAN_METABOLIC_PROC | 34 | 0.5784204 | 1.44905535 | 0.02523129 | 0.26103608 | 0.23892349 | 4999 | tags=59%, list=34%, signal= | [illegible] |
| GOBP_POSTSYNAPSE_ASSEMBLY | GOBP_POSTSYNAPSE_ASSEMBLY | 73 | 0.49231901 | 1.43751067 | 0.02520706 | 0.26103608 | 0.23892349 | 4146 | tags=38%, list=28%, signal= | [illegible] |
| GOBP_P38MAPK_CASCADE | GOBP_P38MAPK_CASCADE | 49 | -0.5752787 | -1.5219403 | 0.02542373 | 0.26266524 | 0.24041464 | 542 | tags=12%, list=4%, signal=1 | [illegible] |
| GOBP_FLAVIN_CONTAINING_COMPOUND_METABOLIC_PROCESS | GOBP_FLAVIN_CONTAINING_COMPOUND_METABOLIC_PROCESS | 4 | 0.88470507 | 1.42081871 | 0.02547957 | 0.26288059 | 0.24061175 | 1223 | tags=50%, list=8%, signal=4 | RFK/SLC52A2 |
| GOBP_T_HELPER_1_CELL_DIFFERENTIATION | GOBP_T_HELPER_1_CELL_DIFFERENTIATION | 33 | 0.58133826 | 1.44921175 | 0.02561956 | 0.26367133 | 0.24133551 | 5151 | tags=55%, list=35%, signal= | [illegible] |
| GOBP_REGULATION_OF_EXTRINSIC_APOPTOTIC_SIGNALING_PATHWAY | GOBP_REGULATION_OF_EXTRINSIC_APOPTOTIC_SIGNALING_PATHW | 138 | 0.42530082 | 1.37280745 | 0.02562642 | 0.26367133 | 0.24133551 | 5037 | tags=37%, list=34%, signal= | [illegible] |
| GOBP_REGULATION_OF_DNA_DAMAGE_RESPONSE_SIGNAL_TRANSDU | GOBP_REGULATION_OF_DNA_DAMAGE_RESPONSE_SIGNAL_TRANSI | 37 | 0.56591144 | 1.44253889 | 0.02566267 | 0.26368307 | 0.24134625 | 4259 | tags=46%, list=29%, signal= | [illegible] |
| GOBP_ODONTOGENESIS | GOBP_ODONTOGENESIS | 89 | 0.4703892 | 1.41828992 | 0.02572584 | 0.26397098 | 0.24160977 | 4115 | tags=42%, list=28%, signal= | [illegible] |
| GOBP_NEGATIVE_REGULATION_OF_INTRACELLULAR_TRANSPORT | GOBP_NEGATIVE_REGULATION_OF_INTRACELLULAR_TRANSPORT | 54 | -0.5583564 | -1.5013501 | 0.02595884 | 0.26563603 | 0.24313378 | 427 | tags=9%, list=3%, signal=99 | PLN/APOD/CRYAB/SLN/PKIA |
| GOBP_RESPONSE_TO_MOLECULE_OF_FUNGAL_ORIGIN | GOBP_RESPONSE_TO_MOLECULE_OF_FUNGAL_ORIGIN | 5 | 0.84621639 | 1.42052834 | 0.02592485 | 0.26563603 | 0.24313378 | 2003 | tags=80%, list=14%, signal= | MYD88/BTK/CLEC7A/SYK |
| GOBP_REGULATION_OF_INTERLEUKIN_6_MEDIATED_SIGNALING_PATH | GOBP_REGULATION_OF_INTERLEUKIN_6_MEDIATED_SIGNALING_PA | 4 | 0.88433308 | 1.42022129 | 0.02606531 | 0.26636264 | 0.24379884 | 655 | tags=50%, list=4%, signal=4 | RIPK1/PTPN2 |
| GOBP_STRIATED_MUSCLE_CELL_APOPTOTIC_PROCESS | GOBP_STRIATED_MUSCLE_CELL_APOPTOTIC_PROCESS | 37 | -0.6160371 | -1.539294 | 0.02623192 | 0.26697554 | 0.24435982 | 799 | tags=14%, list=5%, signal=1 | [illegible] |
| GOBP_MRNA_METHYLGUANOSINE_CAP_DECAPPING | GOBP_MRNA_METHYLGUANOSINE_CAP_DECAPPING | 23 | 0.62297339 | 1.4445476 | 0.02621 | 0.26697554 | 0.24435982 | 2361 | tags=35%, list=16%, signal= | [illegible] |
| GOBP_SENSORY_ORGAN_MORPHOGENESIS | GOBP_SENSORY_ORGAN_MORPHOGENESIS | 212 | 0.38632552 | 1.32363216 | 0.0261811 | 0.26697554 | 0.24435982 | 4058 | tags=40%, list=28%, signal= | [illegible] |
| GOBP_VASCULOGENESIS_INVOLVED_IN_CORONARY_VASCULAR_MORPI | GOBP_VASCULOGENESIS_INVOLVED_IN_CORONARY_VASCULAR_MO | 4 | 0.8838018 | 1.41936807 | 0.02635818 | 0.26789753 | 0.24520371 | 950 | tags=50%, list=6%, signal=4 | NOTCH1/SPRED1 |
| GOBP_RESPONSE_TO_CORTICOSTEROID | GOBP_RESPONSE_TO_CORTICOSTEROID | 113 | -0.4745923 | -1.4416145 | 0.02640052 | 0.26796523 | 0.24526567 | 734 | tags=10%, list=5%, signal=9 | [illegible] |
| GOBP_SPINAL_CORD_ASSOCIATION_NEURON_DIFFERENTIATION | GOBP_SPINAL_CORD_ASSOCIATION_NEURON_DIFFERENTIATION | 3 | 0.92398611 | 1.40968718 | 0.02652083 | 0.26882312 | 0.24605089 | 615 | tags=33%, list=4%, signal=3 | MDGA1 |
| GOBP_HEMIDESMOSOME_ASSEMBLY | GOBP_HEMIDESMOSOME_ASSEMBLY | 5 | -0.9228044 | -1.5670785 | 0.02678571 | 0.27041331 | 0.24750637 | 135 | tags=40%, list=1%, signal=4 | DST/PLEC |
| GOBP_REGULATION_OF_ATPASE_COUPLED_CALCIUM_TRANSMEMBRAI | GOBP_REGULATION_OF_ATPASE_COUPLED_CALCIUM_TRANSMEMB | 5 | -0.9220035 | -1.5657184 | 0.02678571 | 0.27041331 | 0.24750637 | 427 | tags=80%, list=3%, signal=7 | PLN/SLN/STRIT1/ATP2A1 |
| GOBP_NEGATIVE_REGULATION_OF_DNA_METABOLIC_PROCESS | GOBP_NEGATIVE_REGULATION_OF_DNA_METABOLIC_PROCESS | 112 | 0.4442908 | 1.3905851 | 0.02675898 | 0.27041331 | 0.24750637 | 4232 | tags=36%, list=29%, signal= | [illegible] |
| GOBP_DNA_REPLICATION_INITIATION | GOBP_DNA_REPLICATION_INITIATION | 41 | 0.55090701 | 1.43250308 | 0.02690199 | 0.27122259 | 0.24824709 | 3614 | tags=44%, list=25%, signal= | [illegible] |
| GOBP_CILIUM_OR_FLAGELLUM_DEPENDENT_CELL_MOTILITY | GOBP_CILIUM_OR_FLAGELLUM_DEPENDENT_CELL_MOTILITY | 122 | 0.43611348 | 1.38049048 | 0.02712958 | 0.27315047 | 0.25001166 | 4930 | tags=47%, list=34%, signal= | [illegible] |
| GOBP_RESPONSE_TO_DIETARY_EXCESS | GOBP_RESPONSE_TO_DIETARY_EXCESS | 23 | 0.62133642 | 1.44075179 | 0.02717479 | 0.27323939 | 0.25009305 | 3120 | tags=39%, list=21%, signal= | [illegible] |
| GOBP_GLUCOSE_IMPORT_IN_RESPONSE_TO_INSULIN_STIMULUS | GOBP_GLUCOSE_IMPORT_IN_RESPONSE_TO_INSULIN_STIMULUS | 6 | -0.9034137 | -1.583889 | 0.02726106 | 0.27337498 | 0.25021715 | 1222 | tags=50%, list=8%, signal=4 | SGCB/TRARG1/SLC2A4 |
| GOBP_PROTEIN_REFOLDING | GOBP_PROTEIN_REFOLDING | 28 | -0.6584576 | -1.5496717 | 0.02723339 | 0.27337498 | 0.25021715 | 939 | tags=21%, list=6%, signal=2 | [illegible] |
| GOBP_EMBRYONIC_FORELIMB_MORPHOGENESIS | GOBP_EMBRYONIC_FORELIMB_MORPHOGENESIS | 27 | 0.59953181 | 1.43853809 | 0.0274175 | 0.27457714 | 0.25131747 | 3983 | tags=48%, list=27%, signal= | [illegible] |
| GOBP_REGULATION_OF_RELAXATION_OF_MUSCLE | GOBP_REGULATION_OF_RELAXATION_OF_MUSCLE | 12 | -0.8056201 | -1.5888352 | 0.02764423 | 0.27647912 | 0.25305834 | 427 | tags=33%, list=3%, signal=3 | PLN/AKAP6/SLN/HRC |
| GOBP_INFLAMMATORY_RESPONSE | GOBP_INFLAMMATORY_RESPONSE | 673 | 0.31201746 | 1.19887181 | 0.02769963 | 0.27666481 | 0.2532283 | 4366 | tags=34%, list=30%, signal= | [illegible] |
| GOBP_PYRIMIDINE_NUCLEOSIDE_METABOLIC_PROCESS | GOBP_PYRIMIDINE_NUCLEOSIDE_METABOLIC_PROCESS | 10 | 0.74158556 | 1.44452644 | 0.02777365 | 0.27693464 | 0.25347527 | 3651 | tags=80%, list=25%, signal= | [illegible] |
| GOBP_CHOLESTEROL_BIOSYNTHETIC_PROCESS_VIA_DESMOSTEROL | GOBP_CHOLESTEROL_BIOSYNTHETIC_PROCESS_VIA_DESMOSTEROL | 19 | 0.6449134 | 1.4440287 | 0.02780833 | 0.27693464 | 0.25347527 | 1391 | tags=37%, list=10%, signal= | [illegible] |
| GOBP_TISSUE_HOMEOSTASIS | GOBP_TISSUE_HOMEOSTASIS | 194 | 0.39097887 | 1.32348931 | 0.02783726 | 0.27693464 | 0.25347527 | 4273 | tags=38%, list=29%, signal= | [illegible] |
| GOBP_GLANDULAR_EPITHELIAL_CELL_DIFFERENTIATION | GOBP_GLANDULAR_EPITHELIAL_CELL_DIFFERENTIATION | 11 | 0.72170269 | 1.43732694 | 0.02788077 | 0.27700057 | 0.25353561 | 1361 | tags=36%, list=9%, signal=3 | EXT1/NOTCH1/RARB/RARA |
| GOBP_LIPID_DROPLET_FORMATION | GOBP_LIPID_DROPLET_FORMATION | 17 | -0.7367563 | -1.5669787 | 0.02792516 | 0.27707514 | 0.25360387 | 1011 | tags=29%, list=7%, signal=2 | [illegible] |
| GOBP_LEUKOCYTE_CELL_CELL_ADHESION | GOBP_LEUKOCYTE_CELL_CELL_ADHESION | 327 | 0.34896758 | 1.25896417 | 0.02800491 | 0.2774998 | 0.25399256 | 4558 | tags=37%, list=31%, signal= | [illegible] |
| GOBP_NEGATIVE_REGULATION_OF_CYTOSKELETON_ORGANIZATION | GOBP_NEGATIVE_REGULATION_OF_CYTOSKELETON_ORGANIZATION | 136 | -0.4517853 | -1.4121875 | 0.02813624 | 0.27843384 | 0.25484748 | 431 | tags=9%, list=3%, signal=99 | [illegible] |
| GOBP_RNA_EXPORT_FROM_NUCLEUS | GOBP_RNA_EXPORT_FROM_NUCLEUS | 86 | 0.4712496 | 1.41160463 | 0.02827763 | 0.27920019 | 0.25554891 | 4453 | tags=41%, list=30%, signal= | [illegible] |
| GOBP_POSITIVE_REGULATION_OF_CHROMOSOME_ORGANIZATION | GOBP_POSITIVE_REGULATION_OF_CHROMOSOME_ORGANIZATION | 98 | 0.45525231 | 1.39980663 | 0.02828802 | 0.27920019 | 0.25554891 | 4027 | tags=34%, list=28%, signal= | [illegible] |
| GOBP_QUINONE_METABOLIC_PROCESS | GOBP_QUINONE_METABOLIC_PROCESS | 21 | -0.7048641 | -1.5604381 | 0.02835539 | 0.27939754 | 0.25572954 | 1155 | tags=24%, list=8%, signal=2 | [illegible] |
| GOBP_INTRACELLULAR_PROTEIN_TRANSPORT | GOBP_INTRACELLULAR_PROTEIN_TRANSPORT | 602 | 0.31830928 | 1.21159313 | 0.02838242 | 0.27939754 | 0.25572954 | 4103 | tags=29%, list=28%, signal= | [illegible] |
| GOBP_NEGATIVE_REGULATION_OF_MUSCLE_CELL_DIFFERENTIATION | GOBP_NEGATIVE_REGULATION_OF_MUSCLE_CELL_DIFFERENTIATIO | 55 | -0.5519734 | -1.4893491 | 0.02849801 | 0.28016831 | 0.25643501 | 836 | tags=16%, list=6%, signal=1 | [illegible] |
| GOBP_NEGATIVE_REGULATION_OF_CYTOSOLIC_CALCIUM_ION_CONCE | GOBP_NEGATIVE_REGULATION_OF_CYTOSOLIC_CALCIUM_ION_CON | 9 | -0.8496838 | -1.5996351 | 0.02878366 | 0.28256819 | 0.2586316 | 263 | tags=44%, list=2%, signal=4 | KCNK3/GOT1/NOS1/ATP1A2 |
| GOBP_INTRINSIC_APOPTOTIC_SIGNALING_PATHWAY_IN_RESPONSE_TC | GOBP_INTRINSIC_APOPTOTIC_SIGNALING_PATHWAY_IN_RESPONSE_ | 112 | 0.44166115 | 1.38235459 | 0.02881737 | 0.28256819 | 0.2586316 | 3316 | tags=30%, list=23%, signal= | [illegible] |
| GOBP_PROTEIN_DE_ADP_RIBOSYLATION | GOBP_PROTEIN_DE_ADP_RIBOSYLATION | 6 | -0.9013748 | -1.5803143 | 0.02918538 | 0.28580359 | 0.26159293 | 421 | tags=50%, list=3%, signal=4 | ADPRH/MACROD1/ADPRHL1 |
| GOBP_REGULATION_OF_RIBOSOME_BIOGENESIS | GOBP_REGULATION_OF_RIBOSOME_BIOGENESIS | 11 | 0.71899214 | 1.43192867 | 0.02951209 | 0.28862667 | 0.26417686 | 2750 | tags=36%, list=19%, signal= | AFG2A/RBM10/KAT2B/METTL18 |
| GOBP_L_TRYPTOPHAN_CATABOLIC_PROCESS | GOBP_L_TRYPTOPHAN_CATABOLIC_PROCESS | 6 | 0.81279004 | 1.41442381 | 0.02963393 | 0.28944145 | 0.26492262 | 2463 | tags=67%, list=17%, signal= | AFMID/TDO2/IDO1/IL4I1 |
| GOBP_NEGATIVE_REGULATION_OF_TROPHOBLAST_CELL_MIGRATION | GOBP_NEGATIVE_REGULATION_OF_TROPHOBLAST_CELL_MIGRATIC | 5 | 0.84022242 | 1.41046639 | 0.02971162 | 0.28982337 | 0.26527219 | 1528 | tags=60%, list=10%, signal= | GJA1/CALR/TIMP1 |
| GOBP_TUMOR_NECROSIS_FACTOR_MEDIATED_SIGNALING_PATHWAY | GOBP_TUMOR_NECROSIS_FACTOR_MEDIATED_SIGNALING_PATHW | 106 | 0.44646983 | 1.38875901 | 0.02986459 | 0.29093765 | 0.26629208 | 4649 | tags=41%, list=32%, signal= | [illegible] |
| GOBP_MITOCHONDRIAL_RESPIRASOME_ASSEMBLY | GOBP_MITOCHONDRIAL_RESPIRASOME_ASSEMBLY | 7 | -0.8805495 | -1.5843431 | 0.03023552 | 0.291901 | 0.26717382 | 484 | tags=43%, list=3%, signal=4 | RAB5IF/STMP1/COX7A1 |
| GOBP_TOLL_LIKE_RECEPTOR_9_SIGNALING_PATHWAY | GOBP_TOLL_LIKE_RECEPTOR_9_SIGNALING_PATHWAY | 17 | 0.65877498 | 1.43978009 | 0.03021336 | 0.291901 | 0.26717382 | 3274 | tags=41%, list=22%, signal= | [illegible] |
| GOBP_REGULATION_OF_OLIGODENDROCYTE_DIFFERENTIATION | GOBP_REGULATION_OF_OLIGODENDROCYTE_DIFFERENTIATION | 30 | 0.58328686 | 1.42964898 | 0.03017241 | 0.291901 | 0.26717382 | 4314 | tags=57%, list=29%, signal= | [illegible] |
| GOBP_CELLULAR_RESPONSE_TO_NICOTINE | GOBP_CELLULAR_RESPONSE_TO_NICOTINE | 6 | 0.81199815 | 1.41304576 | 0.03006973 | 0.291901 | 0.26717382 | 1423 | tags=17%, list=10%, signal= | B2M |
| GOBP_RNA_LOCALIZATION | GOBP_RNA_LOCALIZATION | 193 | 0.38950258 | 1.31738766 | 0.030185 | 0.291901 | 0.26717382 | 2365 | tags=22%, list=16%, signal= | [illegible] |
| GOBP_MORPHOGENESIS_OF_AN_EPITHELIUM | GOBP_MORPHOGENESIS_OF_AN_EPITHELIUM | 414 | 0.33649636 | 1.24211081 | 0.03008776 | 0.291901 | 0.26717382 | 4273 | tags=36%, list=29%, signal= | [illegible] |
| GOBP_POSITIVE_REGULATION_OF_SIGNALING | GOBP_POSITIVE_REGULATION_OF_SIGNALING | 1469 | 0.28113396 | 1.1327685 | 0.03007519 | 0.291901 | 0.26717382 | 4196 | tags=29%, list=29%, signal= | [illegible] |
| GOBP_GLUTAMINE_METABOLIC_PROCESS | GOBP_GLUTAMINE_METABOLIC_PROCESS | 13 | 0.69576213 | 1.43657932 | 0.03030762 | 0.2922215 | 0.26746717 | 3346 | tags=38%, list=23%, signal= | PHGDH/CAD/PFAS/CPS1/SIRT4 |
| GOBP_ESTABLISHMENT_OF_SPINDLE_ORIENTATION | GOBP_ESTABLISHMENT_OF_SPINDLE_ORIENTATION | 41 | 0.54760563 | 1.42391862 | 0.03050034 | 0.29370266 | 0.26882286 | 2592 | tags=37%, list=18%, signal= | [illegible] |
| GOBP_PROTEIN_REPAIR | GOBP_PROTEIN_REPAIR | 7 | -0.8800877 | -1.5835123 | 0.03055379 | 0.29384059 | 0.2689491 | 1267 | tags=57%, list=9%, signal=5 | PARK7/MSRB1/MSRB2/MSRB3 |
| GOBP_GERANYLGERANYL_DIPHOSPHATE_METABOLIC_PROCESS | GOBP_GERANYLGERANYL_DIPHOSPHATE_METABOLIC_PROCESS | 9 | 0.75182665 | 1.42814221 | 0.03071924 | 0.29505402 | 0.27005974 | 294 | tags=22%, list=2%, signal=2 | HMGCR/HMGCS1 |
| GOBP_NEGATIVE_REGULATION_OF_CELL_ACTIVATION | GOBP_NEGATIVE_REGULATION_OF_CELL_ACTIVATION | 182 | 0.39617826 | 1.32944876 | 0.03083188 | 0.29575765 | 0.27070377 | 5124 | tags=39%, list=35%, signal= | [illegible] |
| GOBP_CRISTAE_FORMATION | GOBP_CRISTAE_FORMATION | 19 | -0.7137334 | -1.5466397 | 0.03106956 | 0.29765751 | 0.27244269 | 1974 | tags=37%, list=13%, signal= | [illegible] |
| GOBP_POSITIVE_REGULATION_OF_MYOTUBE_DIFFERENTIATION | GOBP_POSITIVE_REGULATION_OF_MYOTUBE_DIFFERENTIATION | 12 | -0.7998416 | -1.5774389 | 0.03125 | 0.29900478 | 0.27367583 | 586 | tags=42%, list=4%, signal=4 | [illegible] |
| GOBP_REGULATION_OF_CELL_CYCLE_CHECKPOINT | GOBP_REGULATION_OF_CELL_CYCLE_CHECKPOINT | 56 | 0.51284226 | 1.41974924 | 0.03131561 | 0.29925129 | 0.27390146 | 2246 | tags=30%, list=15%, signal= | [illegible] |
| GOBP_PYRIMIDINE_CONTAINING_COMPOUND_METABOLIC_PROCESS | GOBP_PYRIMIDINE_CONTAINING_COMPOUND_METABOLIC_PROCE | 65 | 0.49589592 | 1.41338353 | 0.03142198 | 0.29950568 | 0.2741343 | 3697 | tags=42%, list=25%, signal= | [illegible] |
| GOBP_CEREBRAL_CORTEX_DEVELOPMENT | GOBP_CEREBRAL_CORTEX_DEVELOPMENT | 108 | 0.44264245 | 1.38091997 | 0.03141848 | 0.29950568 | 0.2741343 | 3023 | tags=28%, list=21%, signal= | [illegible] |
| GOBP_DEOXYRIBONUCLEOSIDE_METABOLIC_PROCESS | GOBP_DEOXYRIBONUCLEOSIDE_METABOLIC_PROCESS | 10 | 0.73568391 | 1.43303069 | 0.03163523 | 0.30107078 | 0.27556682 | 3554 | tags=70%, list=24%, signal= | [illegible] |
| GOBP_MITOCHONDRION_ORGANIZATION | GOBP_MITOCHONDRION_ORGANIZATION | 387 | -0.3629315 | -1.2843871 | 0.03166634 | 0.30107078 | 0.27556682 | 1312 | tags=14%, list=9%, signal=1 | [illegible] |
| GOBP_COLLAGEN_METABOLIC_PROCESS | GOBP_COLLAGEN_METABOLIC_PROCESS | 84 | 0.46858296 | 1.39994338 | 0.03179825 | 0.30194263 | 0.27636482 | 2906 | tags=36%, list=20%, signal= | [illegible] |
| GOBP_NEUROMUSCULAR_JUNCTION_DEVELOPMENT | GOBP_NEUROMUSCULAR_JUNCTION_DEVELOPMENT | 56 | -0.5431872 | -1.470717 | 0.03196454 | 0.30237485 | 0.27676042 | 922 | tags=14%, list=6%, signal=1 | [illegible] |
| GOBP_NEGATIVE_REGULATION_OF_REPRODUCTIVE_PROCESS | GOBP_NEGATIVE_REGULATION_OF_REPRODUCTIVE_PROCESS | 40 | 0.54900489 | 1.42033054 | 0.03196035 | 0.30237485 | 0.27676042 | 3623 | tags=45%, list=25%, signal= | [illegible] |
| GOBP_PARACELLULAR_TRANSPORT | GOBP_PARACELLULAR_TRANSPORT | 3 | 0.9156626 | 1.39698836 | 0.03191489 | 0.30237485 | 0.27676042 | 966 | tags=33%, list=7%, signal=3 | CLDN4 |
| GOBP_PROTEIN_DEPHOSPHORYLATION | GOBP_PROTEIN_DEPHOSPHORYLATION | 75 | -0.5169114 | -1.4706731 | 0.03236682 | 0.30388402 | 0.27814175 | 1033 | tags=24%, list=7%, signal=2 | [illegible] |
| GOBP_NEUTROPHIL_MEDIATED_IMMUNITY | GOBP_NEUTROPHIL_MEDIATED_IMMUNITY | 28 | 0.59101432 | 1.42801741 | 0.032274 | 0.30388402 | 0.27814175 | 3780 | tags=43%, list=26%, signal= | [illegible] |
| GOBP_REPRODUCTIVE_BEHAVIOR | GOBP_REPRODUCTIVE_BEHAVIOR | 30 | 0.58141594 | 1.42506332 | 0.03232759 | 0.30388402 | 0.27814175 | 3487 | tags=33%, list=24%, signal= | [illegible] |
| GOBP_REGULATION_OF_MAP_KINASE_ACTIVITY | GOBP_REGULATION_OF_MAP_KINASE_ACTIVITY | 60 | 0.50349226 | 1.41300774 | 0.0322467 | 0.30388402 | 0.27814175 | 3780 | tags=35%, list=26%, signal= | [illegible] |
| GOBP_RNA_3_END_PROCESSING | GOBP_RNA_3_END_PROCESSING | 92 | 0.45736233 | 1.38879046 | 0.03228765 | 0.30388402 | 0.27814175 | 4882 | tags=36%, list=33%, signal= | [illegible] |
| GOBP_CELLULAR_COMPONENT_DISASSEMBLY | GOBP_CELLULAR_COMPONENT_DISASSEMBLY | 422 | -0.3559648 | -1.2700171 | 0.03221477 | 0.30388402 | 0.27814175 | 950 | tags=9%, list=6%, signal=99 | [illegible] |
| GOBP_NEGATIVE_REGULATION_OF_CELL_SIZE | GOBP_NEGATIVE_REGULATION_OF_CELL_SIZE | 9 | -0.8446367 | -1.5901333 | 0.03249768 | 0.30473167 | 0.27891759 | 757 | tags=44%, list=5%, signal=4 | AKT1S1/DEPTOR/UCN/CAV3 |
| GOBP_REGULATION_BY_VIRUS_OF_VIRAL_PROTEIN_LEVELS_IN_HOST_ | GOBP_REGULATION_BY_VIRUS_OF_VIRAL_PROTEIN_LEVELS_IN_HO | 8 | 0.76906379 | 1.42234447 | 0.03259912 | 0.30530172 | 0.27943936 | 1298 | tags=38%, list=9%, signal=3 | HACD3/IFIT1/DYNLT1 |
| GOBP_LIPID_BIOSYNTHETIC_PROCESS | GOBP_LIPID_BIOSYNTHETIC_PROCESS | 572 | 0.31862281 | 1.20738398 | 0.03267834 | 0.30566255 | 0.27976962 | 4356 | tags=35%, list=30%, signal= | [illegible] |
| GOBP_PURINE_RIBONUCLEOTIDE_SALVAGE | GOBP_PURINE_RIBONUCLEOTIDE_SALVAGE | 7 | -0.8763674 | -1.5768185 | 0.03309994 | 0.30678793 | 0.28079966 | 187 | tags=43%, list=1%, signal=4 | APRT/ADSS1/AMPD1 |
| GOBP_PRIMARY_AMINO_COMPOUND_BIOSYNTHETIC_PROCESS | GOBP_PRIMARY_AMINO_COMPOUND_BIOSYNTHETIC_PROCESS | 3 | -0.9574622 | -1.4755633 | 0.03305288 | 0.30678793 | 0.28079966 | 180 | tags=67%, list=1%, signal=6 | AGMAT/ALDH2 |
| GOBP_PROTEIN_EXIT_FROM_ENDOPLASMIC_RETICULUM | GOBP_PROTEIN_EXIT_FROM_ENDOPLASMIC_RETICULUM | 54 | 0.51562538 | 1.41545753 | 0.03318198 | 0.30678793 | 0.28079966 | 4242 | tags=39%, list=29%, signal= | [illegible] |
| GOBP_OLIGOPEPTIDE_IMPORT_ACROSS_PLASMA_MEMBRANE | GOBP_OLIGOPEPTIDE_IMPORT_ACROSS_PLASMA_MEMBRANE | 4 | 0.87351748 | 1.40285166 | 0.03294772 | 0.30678793 | 0.28079966 | 799 | tags=50%, list=5%, signal=4 | SLC7A11/SLC15A2 |
| GOBP_INTESTINAL_STEM_CELL_HOMEOSTASIS | GOBP_INTESTINAL_STEM_CELL_HOMEOSTASIS | 5 | 0.83475 | 1.40127993 | 0.03320711 | 0.30678793 | 0.28079966 | 1586 | tags=60%, list=11%, signal= | TMEM63B/LGR4/LPCAT3 |
| GOBP_CARDIAC_SEPTUM_DEVELOPMENT | GOBP_CARDIAC_SEPTUM_DEVELOPMENT | 99 | 0.44869503 | 1.38116362 | 0.03315528 | 0.30678793 | 0.28079966 | 4644 | tags=39%, list=32%, signal= | [illegible] |
| GOBP_NEURAL_TUBE_DEVELOPMENT | GOBP_NEURAL_TUBE_DEVELOPMENT | 140 | 0.41695982 | 1.34910384 | 0.03300455 | 0.30678793 | 0.28079966 | 3742 | tags=34%, list=26%, signal= | [illegible] |
| GOBP_ALPHA_BETA_T_CELL_ACTIVATION | GOBP_ALPHA_BETA_T_CELL_ACTIVATION | 164 | 0.40541001 | 1.34000674 | 0.0329036 | 0.30678793 | 0.28079966 | 4721 | tags=40%, list=32%, signal= | [illegible] |
| GOBP_CARTILAGE_DEVELOPMENT | GOBP_CARTILAGE_DEVELOPMENT | 173 | 0.39924383 | 1.33147439 | 0.03316622 | 0.30678793 | 0.28079966 | 3997 | tags=36%, list=27%, signal= | [illegible] |
| GOBP_RESPIRATORY_SYSTEM_DEVELOPMENT | GOBP_RESPIRATORY_SYSTEM_DEVELOPMENT | 187 | 0.39187976 | 1.31913272 | 0.03293705 | 0.30678793 | 0.28079966 | 5064 | tags=41%, list=35%, signal= | [illegible] |
| GOBP_POSITIVE_REGULATION_OF_STRESS_FIBER_ASSEMBLY | GOBP_POSITIVE_REGULATION_OF_STRESS_FIBER_ASSEMBLY | 47 | -0.5673402 | -1.4865743 | 0.03332545 | 0.30750306 | 0.28145422 | 847 | tags=15%, list=6%, signal=1 | [illegible] |
| GOBP_RESPONSE_TO_COLD | GOBP_RESPONSE_TO_COLD | 38 | -0.598856 | -1.5037143 | 0.03340649 | 0.30787254 | 0.28179239 | 1280 | tags=32%, list=9%, signal=2 | [illegible] |
| GOBP_ANTIGEN_PROCESSING_AND_PRESENTATION_OF_EXOGENOUS_ | GOBP_ANTIGEN_PROCESSING_AND_PRESENTATION_OF_EXOGENOU | 13 | 0.69106294 | 1.42687665 | 0.03348992 | 0.30826325 | 0.28215001 | 1488 | tags=31%, list=10%, signal= | CTSL/UNC93B1/B2M/LGMN |
| GOBP_NEGATIVE_REGULATION_OF_MICROTUBULE_POLYMERIZATION | GOBP_NEGATIVE_REGULATION_OF_MICROTUBULE_POLYMERIZATI | 10 | 0.73231441 | 1.42646728 | 0.03356602 | 0.30858551 | 0.28244497 | 1905 | tags=40%, list=13%, signal= | STMN1/MAPRE1/EML2/CLIP3 |
| GOBP_POSITIVE_REGULATION_OF_DEFENSE_RESPONSE | GOBP_POSITIVE_REGULATION_OF_DEFENSE_RESPONSE | 416 | 0.33398573 | 1.23293068 | 0.03366792 | 0.30914396 | 0.28295611 | 3589 | tags=28%, list=25%, signal= | [illegible] |
| GOBP_REGULATION_OF_CYTOKINESIS | GOBP_REGULATION_OF_CYTOKINESIS | 79 | 0.47315216 | 1.39866054 | 0.03373844 | 0.30941318 | 0.28320253 | 3614 | tags=35%, list=25%, signal= | [illegible] |
| GOBP_PYRIMIDINE_DEOXYRIBONUCLEOSIDE_TRIPHOSPHATE_METABO | GOBP_PYRIMIDINE_DEOXYRIBONUCLEOSIDE_TRIPHOSPHATE_META | 7 | 0.78346442 | 1.40779931 | 0.03381924 | 0.30977601 | 0.28353462 | 390 | tags=43%, list=3%, signal=4 | TYMS/TBPL1/CMPK2 |
| GOBP_NUCLEAR_TRANSCRIBED_MRNA_CATABOLIC_PROCESS_DEADEN | GOBP_NUCLEAR_TRANSCRIBED_MRNA_CATABOLIC_PROCESS_DEAD | 64 | 0.49458025 | 1.40405624 | 0.03398577 | 0.31056721 | 0.2842588 | 2204 | tags=22%, list=15%, signal= | [illegible] |
| GOBP_DNA_TEMPLATED_DNA_REPLICATION_MAINTENANCE_OF_FIDEL | GOBP_DNA_TEMPLATED_DNA_REPLICATION_MAINTENANCE_OF_FI | 63 | 0.49548418 | 1.40374592 | 0.03398832 | 0.31056721 | 0.2842588 | 4403 | tags=43%, list=30%, signal= | [illegible] |
| GOBP_POSITIVE_REGULATION_OF_ATTACHMENT_OF_MITOTIC_SPINDL | GOBP_POSITIVE_REGULATION_OF_ATTACHMENT_OF_MITOTIC_SPIN | 8 | 0.76671381 | 1.4179983 | 0.03406755 | 0.31064845 | 0.28433316 | 2485 | tags=50%, list=17%, signal= | CDCA8/AURKB/HNRNPU/KAT2B |
| GOBP_ZINC_ION_TRANSPORT | GOBP_ZINC_ION_TRANSPORT | 25 | 0.60094972 | 1.41787739 | 0.03411306 | 0.31064845 | 0.28433316 | 3478 | tags=52%, list=24%, signal= | [illegible] |
| GOBP_CELL_SURFACE_RECEPTOR_SIGNALING_PATHWAY_VIA_STAT | GOBP_CELL_SURFACE_RECEPTOR_SIGNALING_PATHWAY_VIA_STAT | 107 | 0.44175977 | 1.37545721 | 0.03416264 | 0.31064845 | 0.28433316 | 5189 | tags=46%, list=35%, signal= | [illegible] |
| GOBP_B_CELL_ACTIVATION | GOBP_B_CELL_ACTIVATION | 231 | 0.37168099 | 1.2889368 | 0.03414249 | 0.31064845 | 0.28433316 | 4297 | tags=34%, list=29%, signal= | [illegible] |
| GOBP_GAS_TRANSPORT | GOBP_GAS_TRANSPORT | 8 | -0.8574817 | -1.5847995 | 0.03446115 | 0.31261446 | 0.28613262 | 340 | tags=62%, list=2%, signal=6 | AQP1/HBB/HBA1/HBA2/MB |
| GOBP_ANTIFUNGAL_INNATE_IMMUNE_RESPONSE | GOBP_ANTIFUNGAL_INNATE_IMMUNE_RESPONSE | 12 | 0.69792826 | 1.41764646 | 0.03446209 | 0.31261446 | 0.28613262 | 3490 | tags=42%, list=24%, signal= | [illegible] |
| GOBP_REGULATION_OF_SKELETAL_MUSCLE_CONTRACTION_BY_CALCIU | GOBP_REGULATION_OF_SKELETAL_MUSCLE_CONTRACTION_BY_CAL | 8 | -0.8570754 | -1.5840486 | 0.03477444 | 0.31345066 | 0.28689799 | 172 | tags=50%, list=1%, signal=4 | DMPK/GSTO1/DMD/CASQ1 |
| GOBP_NEGATIVE_REGULATION_OF_NUCLEOCYTOPLASMIC_TRANSPORT | GOBP_NEGATIVE_REGULATION_OF_NUCLEOCYTOPLASMIC_TRANSP | 30 | -0.6315149 | -1.5065172 | 0.03476071 | 0.31345066 | 0.28689799 | 130 | tags=10%, list=1%, signal=1 | PARK7/APOD/PKIA |
| GOBP_REGULATION_OF_SYSTEMIC_ARTERIAL_BLOOD_PRESSURE_BY_N | GOBP_REGULATION_OF_SYSTEMIC_ARTERIAL_BLOOD_PRESSURE_B | 3 | -0.9546494 | -1.4712286 | 0.03485577 | 0.31345066 | 0.28689799 | 19 | tags=67%, list=0%, signal=6 | ADRB2/TPM1 |
| GOBP_INORGANIC_ION_IMPORT_ACROSS_PLASMA_MEMBRANE | GOBP_INORGANIC_ION_IMPORT_ACROSS_PLASMA_MEMBRANE | 76 | -0.5104289 | -1.4541923 | 0.03485496 | 0.31345066 | 0.28689799 | 696 | tags=12%, list=5%, signal=1 | [illegible] |
| GOBP_HYALURONAN_METABOLIC_PROCESS | GOBP_HYALURONAN_METABOLIC_PROCESS | 24 | 0.60657062 | 1.42071398 | 0.03479381 | 0.31345066 | 0.28689799 | 3308 | tags=54%, list=23%, signal= | [illegible] |
| GOBP_MICROTUBULE_NUCLEATION | GOBP_MICROTUBULE_NUCLEATION | 50 | 0.5212861 | 1.41021262 | 0.03483107 | 0.31345066 | 0.28689799 | 2627 | tags=24%, list=18%, signal= | [illegible] |
| GOBP_ISOPENTENYL_DIPHOSPHATE_BIOSYNTHETIC_PROCESS | GOBP_ISOPENTENYL_DIPHOSPHATE_BIOSYNTHETIC_PROCESS | 6 | 0.80606764 | 1.40272544 | 0.03471819 | 0.31345066 | 0.28689799 | 294 | tags=33%, list=2%, signal=3 | HMGCR/HMGCS1 |
| GOBP_MRNA_TRANSPORT | GOBP_MRNA_TRANSPORT | 124 | 0.42613705 | 1.35270021 | 0.03488813 | 0.31345066 | 0.28689799 | 4453 | tags=40%, list=30%, signal= | [illegible] |
| GOBP_SECRETORY_GRANULE_LOCALIZATION | GOBP_SECRETORY_GRANULE_LOCALIZATION | 20 | -0.7002087 | -1.5359395 | 0.03508295 | 0.31437029 | 0.28773972 | 597 | tags=25%, list=4%, signal=2 | [illegible] |
| GOBP_DNA_CONFORMATION_CHANGE | GOBP_DNA_CONFORMATION_CHANGE | 16 | 0.66272323 | 1.42941978 | 0.03517744 | 0.31437029 | 0.28773972 | 3493 | tags=62%, list=24%, signal= | [illegible] |
| GOBP_SKIN_MORPHOGENESIS | GOBP_SKIN_MORPHOGENESIS | 10 | 0.72955351 | 1.42108935 | 0.03519976 | 0.31437029 | 0.28773972 | 2387 | tags=40%, list=16%, signal= | ERRFI1/COL1A1/VPS33B/COL1A2 |
| GOBP_PHOTORECEPTOR_CELL_MAINTENANCE | GOBP_PHOTORECEPTOR_CELL_MAINTENANCE | 24 | 0.60637839 | 1.42026374 | 0.03511598 | 0.31437029 | 0.28773972 | 1794 | tags=38%, list=12%, signal= | [illegible] |
| GOBP_NEGATIVE_REGULATION_OF_EXOCYTOSIS | GOBP_NEGATIVE_REGULATION_OF_EXOCYTOSIS | 30 | 0.57811158 | 1.41696426 | 0.03514589 | 0.31437029 | 0.28773972 | 1014 | tags=23%, list=7%, signal=2 | [illegible] |
| GOBP_NEGATIVE_REGULATION_OF_HETEROCHROMATIN_ORGANIZATI | GOBP_NEGATIVE_REGULATION_OF_HETEROCHROMATIN_ORGANIZ | 7 | 0.78224499 | 1.40560813 | 0.03542274 | 0.31543093 | 0.28871051 | 896 | tags=29%, list=6%, signal=2 | ERI1/ERI2 |
| GOBP_RNA_GUANINE_N7_METHYLATION | GOBP_RNA_GUANINE_N7_METHYLATION | 6 | 0.80430614 | 1.39966006 | 0.03544451 | 0.31543093 | 0.28871051 | 874 | tags=33%, list=6%, signal=3 | RAMAC/WDR4 |
| GOBP_REGULATION_OF_LEUKOCYTE_PROLIFERATION | GOBP_REGULATION_OF_LEUKOCYTE_PROLIFERATION | 190 | 0.38740677 | 1.30761922 | 0.03543614 | 0.31543093 | 0.28871051 | 4989 | tags=41%, list=34%, signal= | [illegible] |
| GOBP_ADULT_HEART_DEVELOPMENT | GOBP_ADULT_HEART_DEVELOPMENT | 9 | -0.8399771 | -1.5813611 | 0.0355927 | 0.316181 | 0.28939704 | 85 | tags=33%, list=1%, signal=3 | MEF2D/MYH7/TCAP |
| GOBP_TRANSDIFFERENTIATION | GOBP_TRANSDIFFERENTIATION | 4 | -0.9362555 | -1.5257389 | 0.03561298 | 0.316181 | 0.28939704 | 263 | tags=50%, list=2%, signal=4 | ASNSD1/GOT1 |
| GOBP_RNA_MODIFICATION | GOBP_RNA_MODIFICATION | 171 | 0.39774183 | 1.32288308 | 0.03565921 | 0.31621765 | 0.28943059 | 2087 | tags=20%, list=14%, signal= | [illegible] |
| GOBP_RESPONSE_TO_MONOAMINE | GOBP_RESPONSE_TO_MONOAMINE | 43 | 0.53765471 | 1.41030481 | 0.03574497 | 0.31660436 | 0.28978453 | 4636 | tags=49%, list=32%, signal= | [illegible] |
| GOBP_RESPONSE_TO_RADIATION | GOBP_RESPONSE_TO_RADIATION | 348 | 0.3416262 | 1.24128821 | 0.03621567 | 0.32039561 | 0.29325462 | 3791 | tags=28%, list=26%, signal= | [illegible] |
| GOBP_ORGANOPHOSPHATE_BIOSYNTHETIC_PROCESS | GOBP_ORGANOPHOSPHATE_BIOSYNTHETIC_PROCESS | 535 | -0.3405714 | -1.2413834 | 0.03628804 | 0.32065817 | 0.29349494 | 1053 | tags=13%, list=7%, signal=1 | [illegible] |
| GOBP_SPHINGOLIPID_MEDIATED_SIGNALING_PATHWAY | GOBP_SPHINGOLIPID_MEDIATED_SIGNALING_PATHWAY | 15 | 0.6694665 | 1.42466797 | 0.03641672 | 0.32141712 | 0.2941896 | 4046 | tags=53%, list=28%, signal= | [illegible] |
| GOBP_NEUTROPHIL_DIFFERENTIATION | GOBP_NEUTROPHIL_DIFFERENTIATION | 10 | 0.72689238 | 1.41590576 | 0.03653646 | 0.32194544 | 0.29467317 | 2157 | tags=40%, list=15%, signal= | LBR/FAM3C/LEF1/INPP5D |
| GOBP_RESPONSE_TO_BACTERIUM | GOBP_RESPONSE_TO_BACTERIUM | 447 | 0.32790954 | 1.2179625 | 0.0365623 | 0.32194544 | 0.29467317 | 3878 | tags=28%, list=26%, signal= | [illegible] |
| GOBP_TRNA_WOBBLE_BASE_MODIFICATION | GOBP_TRNA_WOBBLE_BASE_MODIFICATION | 29 | 0.58095841 | 1.41353032 | 0.03676834 | 0.32338059 | 0.29598675 | 2760 | tags=34%, list=19%, signal= | [illegible] |
| GOBP_REGULATION_OF_SARCOMERE_ORGANIZATION | GOBP_REGULATION_OF_SARCOMERE_ORGANIZATION | 6 | -0.8932476 | -1.5660655 | 0.03688262 | 0.32400624 | 0.2965594 | 356 | tags=67%, list=2%, signal=6 | EDN1/MYLK3/ANKRD23/CAV3 |
| GOBP_SHORT_CHAIN_FATTY_ACID_CATABOLIC_PROCESS | GOBP_SHORT_CHAIN_FATTY_ACID_CATABOLIC_PROCESS | 5 | -0.9116877 | -1.5482005 | 0.0369898 | 0.32431749 | 0.29684428 | 260 | tags=60%, list=2%, signal=5 | PCCA/PHYH/ACADS |
| GOBP_POSITIVE_REGULATION_OF_CHROMOSOME_CONDENSATION | GOBP_POSITIVE_REGULATION_OF_CHROMOSOME_CONDENSATION | 8 | 0.76154291 | 1.408435 | 0.03700441 | 0.32431749 | 0.29684428 | 2986 | tags=75%, list=20%, signal= | [illegible] |

| | | | | | | | | | | |
|---|---|---|---|---|---|---|---|---|---|---|
| GOBP_GOLGI_LOCALIZATION | GOBP_GOLGI_LOCALIZATION | 13 | 0.68488671 | 1.41412425 | 0.03712684 | 0.32501128 | 0.2974793 | 1456 | tags=31%, list=10%, signal=[illegible] | CDC42/YWHAZ/COPG1/NHERF1 |
| GOBP_SATELLITE_CELL_ACTIVATION_INVOLVED_IN_SKELETAL_MUSCLE | GOBP_SATELLITE_CELL_ACTIVATION_INVOLVED_IN_SKELETAL_MUSC | 5 | -0.911326 | -1.5475863 | 0.03730867 | 0.32546981 | 0.29789899 | 257 | tags=60%, list=2%, signal=[illegible] | SOX15/XIRP1/CAPN3 |
| GOBP_OPTIC_NERVE_MORPHOGENESIS | GOBP_OPTIC_NERVE_MORPHOGENESIS | 3 | 0.90990831 | 1.38820927 | 0.03730896 | 0.32546981 | 0.29789899 | 401 | tags=67%, list=3%, signal=[illegible] | EPHB2/GLI3 |
| GOBP_POSITIVE_REGULATION_OF_MITOTIC_CELL_CYCLE | GOBP_POSITIVE_REGULATION_OF_MITOTIC_CELL_CYCLE | 116 | 0.43148616 | 1.35698513 | 0.03730921 | 0.32546981 | 0.29789899 | 2002 | tags=25%, list=14%, signal=[illegible] | [illegible] |
| GOBP_POSITIVE_REGULATION_OF_CILIUM_ASSEMBLY | GOBP_POSITIVE_REGULATION_OF_CILIUM_ASSEMBLY | 28 | 0.58331911 | 1.40942412 | 0.03737856 | 0.32569649 | 0.29810646 | 2130 | tags=32%, list=15%, signal=[illegible] | [illegible] |
| GOBP_POSITIVE_REGULATION_OF_T_CELL_PROLIFERATION | GOBP_POSITIVE_REGULATION_OF_T_CELL_PROLIFERATION | 69 | 0.48465426 | 1.39772469 | 0.03759131 | 0.32667711 | 0.29900402 | 4490 | tags=45%, list=31%, signal=[illegible] | [illegible] |
| GOBP_INNATE_IMMUNE_RESPONSE_ACTIVATING_CELL_SURFACE_RECE | GOBP_INNATE_IMMUNE_RESPONSE_ACTIVATING_CELL_SURFACE_R | 77 | 0.47145974 | 1.38964552 | 0.03753835 | 0.32667711 | 0.29900402 | 3681 | tags=32%, list=25%, signal=[illegible] | [illegible] |
| GOBP_POSITIVE_REGULATION_OF_PROTEIN_LOCALIZATION_TO_NUCLE | GOBP_POSITIVE_REGULATION_OF_PROTEIN_LOCALIZATION_TO_NU | 87 | 0.45993346 | 1.38032328 | 0.03762158 | 0.32667711 | 0.29900402 | 2424 | tags=24%, list=17%, signal=[illegible] | [illegible] |
| GOBP_POSITIVE_REGULATION_OF_MONOCYTE_CHEMOTAXIS | GOBP_POSITIVE_REGULATION_OF_MONOCYTE_CHEMOTAXIS | 15 | 0.66751908 | 1.42052374 | 0.03795329 | 0.32917685 | 0.301292 | 1919 | tags=40%, list=13%, signal=[illegible] | [illegible] |
| GOBP_VIRAL_PROCESS | GOBP_VIRAL_PROCESS | 320 | 0.34570755 | 1.24503137 | 0.03810882 | 0.33014461 | 0.30217778 | 2662 | tags=19%, list=18%, signal=[illegible] | [illegible] |
| GOBP_REGULATION_OF_PHOSPHOLIPID_CATABOLIC_PROCESS | GOBP_REGULATION_OF_PHOSPHOLIPID_CATABOLIC_PROCESS | 5 | 0.82759127 | 1.38926269 | 0.03815904 | 0.33019883 | 0.30222741 | 1561 | tags=80%, list=11%, signal=[illegible] | IDH1/LDLR/SCARB1/PRKCD |
| GOBP_INTESTINAL_LIPID_ABSORPTION | GOBP_INTESTINAL_LIPID_ABSORPTION | 9 | 0.74121903 | 1.40799235 | 0.03839905 | 0.33151184 | 0.30342919 | 1586 | tags=44%, list=11%, signal=[illegible] | LDLR/UGCG/LIMA1/LPCAT3 |
| GOBP_MESONEPHROS_DEVELOPMENT | GOBP_MESONEPHROS_DEVELOPMENT | 87 | 0.45875254 | 1.37677916 | 0.03835566 | 0.33151184 | 0.30342919 | 3481 | tags=36%, list=24%, signal=[illegible] | [illegible] |
| GOBP_NUCLEAR_TRANSCRIBED_MRNA_POLY_A_TAIL_SHORTENING | GOBP_NUCLEAR_TRANSCRIBED_MRNA_POLY_A_TAIL_SHORTENING | 33 | 0.56483481 | 1.40807047 | 0.03868051 | 0.33279416 | 0.30460289 | 1268 | tags=21%, list=9%, signal=[illegible] | [illegible] |
| GOBP_CELL_SURFACE_PATTERN_RECOGNITION_RECEPTOR_SIGNALING | GOBP_CELL_SURFACE_PATTERN_RECOGNITION_RECEPTOR_SIGNALI | 66 | 0.48636258 | 1.38994136 | 0.03864648 | 0.33279416 | 0.30460289 | 3681 | tags=33%, list=25%, signal=[illegible] | [illegible] |
| GOBP_MYELOID_CELL_ACTIVATION_INVOLVED_IN_IMMUNE_RESPONS | GOBP_MYELOID_CELL_ACTIVATION_INVOLVED_IN_IMMUNE_RESPC | 77 | 0.47010075 | 1.38563984 | 0.03862119 | 0.33279416 | 0.30460289 | 4179 | tags=39%, list=29%, signal=[illegible] | [illegible] |
| GOBP_NEUROGENESIS | GOBP_NEUROGENESIS | 1389 | 0.28011598 | 1.12583761 | 0.03874883 | 0.33300055 | 0.3047918 | 4303 | tags=32%, list=29%, signal=[illegible] | [illegible] |
| GOBP_CELLULAR_RESPONSE_TO_CALCIUM_ION | GOBP_CELLULAR_RESPONSE_TO_CALCIUM_ION | 60 | -0.5298143 | -1.4520983 | 0.03882598 | 0.33328218 | 0.30504957 | 734 | tags=13%, list=5%, signal=[illegible] | [illegible] |
| GOBP_NUCLEOSIDE_SALVAGE | GOBP_NUCLEOSIDE_SALVAGE | 12 | 0.69219434 | 1.40599961 | 0.03895715 | 0.33402641 | 0.30573075 | 356 | tags=25%, list=2%, signal=[illegible] | PGM2/UCKL1/QNG1 |
| GOBP_MACROPHAGE_COLONY_STIMULATING_FACTOR_SIGNALING_PA | GOBP_MACROPHAGE_COLONY_STIMULATING_FACTOR_SIGNALING_ | 7 | 0.77673934 | 1.39571508 | 0.03921283 | 0.33551617 | 0.30709431 | 1609 | tags=43%, list=11%, signal=[illegible] | PTPN2/DOK1/CSF1 |
| GOBP_BIOLOGICAL_PROCESS_INVOLVED_IN_INTERSPECIES_INTERACTI | GOBP_BIOLOGICAL_PROCESS_INVOLVED_IN_INTERSPECIES_INTERAC | 1306 | 0.28160703 | 1.12830079 | 0.03922024 | 0.33551617 | 0.30709431 | 3465 | tags=24%, list=24%, signal=[illegible] | [illegible] |
| GOBP_INTERLEUKIN_10_PRODUCTION | GOBP_INTERLEUKIN_10_PRODUCTION | 45 | 0.52957209 | 1.40119153 | 0.03930357 | 0.33584653 | 0.30739669 | 4401 | tags=49%, list=30%, signal=[illegible] | [illegible] |
| GOBP_NEUROTRANSMITTER_REUPTAKE | GOBP_NEUROTRANSMITTER_REUPTAKE | 20 | -0.6949379 | -1.5243778 | 0.03943432 | 0.3365809 | 0.30806884 | 1835 | tags=50%, list=13%, signal=[illegible] | [illegible] |
| GOBP_L_GLUTAMATE_CATABOLIC_PROCESS | GOBP_L_GLUTAMATE_CATABOLIC_PROCESS | 4 | -0.9314511 | -1.5179095 | 0.03971005 | 0.33784186 | 0.30922299 | 263 | tags=75%, list=2%, signal=[illegible] | GLUL/GOT1/GOT2 |
| GOBP_REGULATION_OF_T_HELPER_1_TYPE_IMMUNE_RESPONSE | GOBP_REGULATION_OF_T_HELPER_1_TYPE_IMMUNE_RESPONSE | 23 | 0.60703684 | 1.40759399 | 0.039717 | 0.33784186 | 0.30922299 | 4297 | tags=52%, list=29%, signal=[illegible] | [illegible] |
| GOBP_REGULATION_OF_RECEPTOR_SIGNALING_PATHWAY_VIA_STAT | GOBP_REGULATION_OF_RECEPTOR_SIGNALING_PATHWAY_VIA_STA | 63 | 0.491596 | 1.39273041 | 0.03965304 | 0.33784186 | 0.30922299 | 5189 | tags=57%, list=35%, signal=[illegible] | [illegible] |
| GOBP_EMBRYONIC_ORGAN_MORPHOGENESIS | GOBP_EMBRYONIC_ORGAN_MORPHOGENESIS | 229 | 0.3687523 | 1.27591073 | 0.03977048 | 0.3379141 | 0.30928911 | 4256 | tags=39%, list=29%, signal=[illegible] | [illegible] |
| GOBP_REGULATION_OF_DEVELOPMENT_HETEROCHRONIC | GOBP_REGULATION_OF_DEVELOPMENT_HETEROCHRONIC | 6 | 0.79898817 | 1.39040569 | 0.0399477 | 0.3390364 | 0.31031634 | 2946 | tags=83%, list=20%, signal=[illegible] | [illegible] |
| GOBP_NEURON_NEURON_SYNAPTIC_TRANSMISSION | GOBP_NEURON_NEURON_SYNAPTIC_TRANSMISSION | 7 | -0.8671848 | -1.5602965 | 0.04042011 | 0.33966188 | 0.31088883 | 690 | tags=43%, list=5%, signal=[illegible] | VDAC1/VDAC3/KIF1B |
| GOBP_NEGATIVE_REGULATION_OF_MYOTUBE_DIFFERENTIATION | GOBP_NEGATIVE_REGULATION_OF_MYOTUBE_DIFFERENTIATION | 12 | -0.7838913 | -1.5459818 | 0.04026442 | 0.33966188 | 0.31088883 | 201 | tags=33%, list=1%, signal=[illegible] | HDAC4/TRIM72/PLPP7/RPL3L |
| GOBP_POSITIVE_REGULATION_OF_CALCIUM_ION_IMPORT | GOBP_POSITIVE_REGULATION_OF_CALCIUM_ION_IMPORT | 19 | -0.7009337 | -1.5189031 | 0.04014297 | 0.33966188 | 0.31088883 | 461 | tags=26%, list=3%, signal=[illegible] | [illegible] |
| GOBP_NEGATIVE_REGULATION_OF_SMOOTH_MUSCLE_CELL_PROLIFER | GOBP_NEGATIVE_REGULATION_OF_SMOOTH_MUSCLE_CELL_PROLII | 41 | -0.5801373 | -1.4806685 | 0.04008641 | 0.33966188 | 0.31088883 | 805 | tags=15%, list=5%, signal=[illegible] | [illegible] |
| GOBP_ANTIGEN_PROCESSING_AND_PRESENTATION_OF_EXOGENOUS_ | GOBP_ANTIGEN_PROCESSING_AND_PRESENTATION_OF_EXOGENOU | 26 | 0.59162307 | 1.40761656 | 0.04050302 | 0.33966188 | 0.31088883 | 2780 | tags=38%, list=19%, signal=[illegible] | [illegible] |
| GOBP_CELLULAR_RESPONSE_TO_OXIDISED_LOW_DENSITY_LIPOPROTEI | GOBP_CELLULAR_RESPONSE_TO_OXIDISED_LOW_DENSITY_LIPOPRC | 10 | 0.72203582 | 1.40644572 | 0.04054656 | 0.33966188 | 0.31088883 | 1049 | tags=30%, list=7%, signal=[illegible] | AKT1/MYD88/TLR6 |
| GOBP_EPIDERMIS_MORPHOGENESIS | GOBP_EPIDERMIS_MORPHOGENESIS | 23 | 0.6062781 | 1.40583462 | 0.04019939 | 0.33966188 | 0.31088883 | 3983 | tags=57%, list=27%, signal=[illegible] | [illegible] |
| GOBP_PHOSPHATIDYLCHOLINE_METABOLIC_PROCESS | GOBP_PHOSPHATIDYLCHOLINE_METABOLIC_PROCESS | 51 | 0.51539574 | 1.39883314 | 0.04056407 | 0.33966188 | 0.31088883 | 3115 | tags=41%, list=21%, signal=[illegible] | [illegible] |
| GOBP_POSTSYNAPTIC_MEMBRANE_ASSEMBLY | GOBP_POSTSYNAPTIC_MEMBRANE_ASSEMBLY | 6 | 0.79732472 | 1.38751093 | 0.04052876 | 0.33966188 | 0.31088883 | 2369 | tags=83%, list=16%, signal=[illegible] | [illegible] |
| GOBP_MODIFIED_AMINO_ACID_METABOLIC_PROCESS | GOBP_MODIFIED_AMINO_ACID_METABOLIC_PROCESS | 139 | -0.4358787 | -1.3678954 | 0.04048071 | 0.33966188 | 0.31088883 | 1015 | tags=15%, list=7%, signal=[illegible] | [illegible] |
| GOBP_THYMOCYTE_MIGRATION | GOBP_THYMOCYTE_MIGRATION | 1 | 0.98305548 | 1.31351696 | 0.04030788 | 0.33966188 | 0.31088883 | 249 | tags=100%, list=2%, signal=[illegible] | ADAM8 |
| GOBP_PROTEIN_LOCALIZATION_TO_ORGANELLE | GOBP_PROTEIN_LOCALIZATION_TO_ORGANELLE | 809 | 0.29697536 | 1.15565961 | 0.04046624 | 0.33966188 | 0.31088883 | 4592 | tags=31%, list=31%, signal=[illegible] | [illegible] |
| GOBP_NUCLEAR_HISTONE_MRNA_CATABOLIC_PROCESS | GOBP_NUCLEAR_HISTONE_MRNA_CATABOLIC_PROCESS | 12 | 0.69053094 | 1.40262088 | 0.04075517 | 0.34012453 | 0.31131229 | 1584 | tags=42%, list=11%, signal=[illegible] | [illegible] |
| GOBP_MITOTIC_CYTOKINETIC_PROCESS | GOBP_MITOTIC_CYTOKINETIC_PROCESS | 37 | 0.54871835 | 1.39871277 | 0.04068884 | 0.34012453 | 0.31131229 | 2419 | tags=30%, list=17%, signal=[illegible] | [illegible] |
| GOBP_NEGATIVE_REGULATION_OF_NEURON_DIFFERENTIATION | GOBP_NEGATIVE_REGULATION_OF_NEURON_DIFFERENTIATION | 47 | 0.52258957 | 1.39721113 | 0.04072085 | 0.34012453 | 0.31131229 | 3263 | tags=43%, list=22%, signal=[illegible] | [illegible] |
| GOBP_POTASSIUM_ION_IMPORT_ACROSS_PLASMA_MEMBRANE | GOBP_POTASSIUM_ION_IMPORT_ACROSS_PLASMA_MEMBRANE | 29 | -0.6334689 | -1.5015495 | 0.04114808 | 0.34226273 | 0.31326937 | 696 | tags=21%, list=5%, signal=[illegible] | [illegible] |
| GOBP_MICROGLIA_DIFFERENTIATION | GOBP_MICROGLIA_DIFFERENTIATION | 13 | 0.67950399 | 1.40301022 | 0.04106683 | 0.34226273 | 0.31326937 | 2218 | tags=46%, list=15%, signal=[illegible] | [illegible] |
| GOBP_GOLGI_DISASSEMBLY | GOBP_GOLGI_DISASSEMBLY | 6 | 0.79522278 | 1.38385312 | 0.04110982 | 0.34226273 | 0.31326937 | 234 | tags=33%, list=2%, signal=[illegible] | [illegible] |
| GOBP_STRESS_FIBER_ASSEMBLY | GOBP_STRESS_FIBER_ASSEMBLY | 100 | -0.4688386 | -1.4015082 | 0.04136885 | 0.34371843 | 0.31460175 | 847 | tags=11%, list=6%, signal=[illegible] | [illegible] |
| GOBP_HYALURONAN_CATABOLIC_PROCESS | GOBP_HYALURONAN_CATABOLIC_PROCESS | 14 | 0.67054448 | 1.408219 | 0.04194631 | 0.34774694 | 0.318289 | 3308 | tags=64%, list=23%, signal=[illegible] | [illegible] |
| GOBP_FC_RECEPTOR_MEDIATED_STIMULATORY_SIGNALING_PATHWA | GOBP_FC_RECEPTOR_MEDIATED_STIMULATORY_SIGNALING_PATHV | 32 | 0.56670294 | 1.40332618 | 0.04194519 | 0.34774694 | 0.318289 | 4531 | tags=47%, list=31%, signal=[illegible] | [illegible] |
| GOBP_REGULATION_OF_SPINDLE_ASSEMBLY | GOBP_REGULATION_OF_SPINDLE_ASSEMBLY | 33 | 0.56177589 | 1.40044493 | 0.04202947 | 0.34801169 | 0.31853133 | 4667 | tags=48%, list=32%, signal=[illegible] | [illegible] |
| GOBP_NEGATIVE_REGULATION_OF_INNATE_IMMUNE_RESPONSE | GOBP_NEGATIVE_REGULATION_OF_INNATE_IMMUNE_RESPONSE | 74 | 0.473485 | 1.38565429 | 0.0420712 | 0.34801169 | 0.31853133 | 3248 | tags=28%, list=22%, signal=[illegible] | [illegible] |
| GOBP_KIDNEY_EPITHELIUM_DEVELOPMENT | GOBP_KIDNEY_EPITHELIUM_DEVELOPMENT | 122 | 0.4228564 | 1.33852601 | 0.04211725 | 0.34801169 | 0.31853133 | 4238 | tags=39%, list=29%, signal=[illegible] | [illegible] |
| GOBP_TRANSITION_METAL_ION_TRANSPORT | GOBP_TRANSITION_METAL_ION_TRANSPORT | 77 | 0.46780961 | 1.37888662 | 0.04223064 | 0.34856524 | 0.31903798 | 3872 | tags=36%, list=26%, signal=[illegible] | [illegible] |
| GOBP_CERAMIDE_METABOLIC_PROCESS | GOBP_CERAMIDE_METABOLIC_PROCESS | 78 | 0.46706456 | 1.37825833 | 0.04236061 | 0.34899928 | 0.31943526 | 4275 | tags=44%, list=29%, signal=[illegible] | [illegible] |
| GOBP_RESPONSE_TO_OXIDATIVE_STRESS | GOBP_RESPONSE_TO_OXIDATIVE_STRESS | 368 | -0.3597585 | -1.2676988 | 0.04237616 | 0.34899928 | 0.31943526 | 1035 | tags=12%, list=7%, signal=[illegible] | [illegible] |
| GOBP_DIAPHRAGM_DEVELOPMENT | GOBP_DIAPHRAGM_DEVELOPMENT | 9 | 0.73606583 | 1.39820353 | 0.04253434 | 0.34953546 | 0.31992602 | 3242 | tags=56%, list=22%, signal=[illegible] | [illegible] |
| GOBP_VIRAL_LIFE_CYCLE | GOBP_VIRAL_LIFE_CYCLE | 241 | 0.36350816 | 1.26830431 | 0.04249012 | 0.34953546 | 0.31992602 | 2662 | tags=20%, list=18%, signal=[illegible] | [illegible] |
| GOBP_PLASMA_MEMBRANE_RAFT_ASSEMBLY | GOBP_PLASMA_MEMBRANE_RAFT_ASSEMBLY | 9 | -0.8301914 | -1.5629383 | 0.04271123 | 0.35060556 | 0.32090546 | 36 | tags=22%, list=0%, signal=[illegible] | [illegible] |
| GOBP_REGULATION_OF_COAGULATION | GOBP_REGULATION_OF_COAGULATION | 50 | 0.51530725 | 1.39403831 | 0.04284222 | 0.35100199 | 0.32126832 | 3320 | tags=40%, list=23%, signal=[illegible] | [illegible] |
| GOBP_MEIOTIC_CHROMOSOME_CONDENSATION | GOBP_MEIOTIC_CHROMOSOME_CONDENSATION | 6 | 0.79269714 | 1.37945798 | 0.04285299 | 0.35100199 | 0.32126832 | 2986 | tags=83%, list=20%, signal=[illegible] | [illegible] |
| GOBP_POSITIVE_REGULATION_OF_CD4_POSITIVE_ALPHA_BETA_T_CEL | GOBP_POSITIVE_REGULATION_OF_CD4_POSITIVE_ALPHA_BETA_T_C | 29 | 0.57397661 | 1.39654289 | 0.04319868 | 0.35306343 | 0.32315513 | 4140 | tags=45%, list=28%, signal=[illegible] | [illegible] |
| GOBP_HINDBRAIN_DEVELOPMENT | GOBP_HINDBRAIN_DEVELOPMENT | 115 | 0.42626293 | 1.33928923 | 0.04318438 | 0.35306343 | 0.32315513 | 4102 | tags=33%, list=28%, signal=[illegible] | [illegible] |
| GOBP_NITRIC_OXIDE_TRANSPORT | GOBP_NITRIC_OXIDE_TRANSPORT | 5 | -0.9056266 | -1.5379078 | 0.04336735 | 0.35367225 | 0.32371238 | 340 | tags=100%, list=2%, signal=[illegible] | [illegible] |
| GOBP_REGULATION_OF_GLYCOLYTIC_PROCESS | GOBP_REGULATION_OF_GLYCOLYTIC_PROCESS | 45 | -0.5637471 | -1.4628194 | 0.04332302 | 0.35367225 | 0.32371238 | 836 | tags=13%, list=6%, signal=[illegible] | [illegible] |
| GOBP_U2_TYPE_PRESPLICEOSOME_ASSEMBLY | GOBP_U2_TYPE_PRESPLICEOSOME_ASSEMBLY | 25 | 0.59015174 | 1.39240071 | 0.04369721 | 0.35566464 | 0.32553599 | 1584 | tags=28%, list=11%, signal=[illegible] | [illegible] |
| GOBP_EMBRYO_IMPLANTATION | GOBP_EMBRYO_IMPLANTATION | 50 | 0.51451836 | 1.39190414 | 0.04371299 | 0.35566464 | 0.32553599 | 2523 | tags=28%, list=17%, signal=[illegible] | [illegible] |
| GOBP_NEGATIVE_REGULATION_OF_TRANSPORT | GOBP_NEGATIVE_REGULATION_OF_TRANSPORT | 312 | -0.36825 | -1.2744836 | 0.04375371 | 0.35566464 | 0.32553599 | 1185 | tags=12%, list=8%, signal=[illegible] | [illegible] |
| GOBP_REGULATION_OF_TOLL_SIGNALING_PATHWAY | GOBP_REGULATION_OF_TOLL_SIGNALING_PATHWAY | 6 | 0.79148345 | 1.3773459 | 0.04386984 | 0.35622313 | 0.32604716 | 184 | tags=17%, list=1%, signal=[illegible] | [illegible] |
| GOBP_REGULATION_OF_STORE_OPERATED_CALCIUM_CHANNEL_ACTIV | GOBP_REGULATION_OF_STORE_OPERATED_CALCIUM_CHANNEL_AC | 7 | -0.8643029 | -1.5551112 | 0.04392107 | 0.35625394 | 0.32607537 | 30 | tags=29%, list=0%, signal=[illegible] | [illegible] |
| GOBP_RENAL_SYSTEM_DEVELOPMENT | GOBP_RENAL_SYSTEM_DEVELOPMENT | 272 | 0.35370618 | 1.25028551 | 0.04405901 | 0.35698731 | 0.32674661 | 5166 | tags=41%, list=35%, signal=[illegible] | [illegible] |
| GOBP_SPINDLE_ASSEMBLY_INVOLVED_IN_FEMALE_MEIOSIS | GOBP_SPINDLE_ASSEMBLY_INVOLVED_IN_FEMALE_MEIOSIS | 12 | 0.68798065 | 1.39744067 | 0.04420138 | 0.35771353 | 0.32741132 | 2960 | tags=58%, list=20%, signal=[illegible] | [illegible] |
| GOBP_CARBOHYDRATE_TRANSPORT | GOBP_CARBOHYDRATE_TRANSPORT | 95 | -0.4710456 | -1.3912872 | 0.04424389 | 0.35771353 | 0.32741132 | 1235 | tags=16%, list=8%, signal=[illegible] | [illegible] |
| GOBP_DE_NOVO_CENTRIOLE_ASSEMBLY_INVOLVED_IN_MULTI_CILIAT | GOBP_DE_NOVO_CENTRIOLE_ASSEMBLY_INVOLVED_IN_MULTI_CIL | 3 | 0.89992057 | 1.37297139 | 0.04450105 | 0.35863453 | 0.32825429 | 1451 | tags=67%, list=10%, signal=[illegible] | [illegible] |
| GOBP_POSITIVE_REGULATION_OF_INTRACELLULAR_PROTEIN_TRANSP | GOBP_POSITIVE_REGULATION_OF_INTRACELLULAR_PROTEIN_TRAN | 89 | 0.45166361 | 1.36182961 | 0.04446895 | 0.35863453 | 0.32825429 | 2390 | tags=22%, list=16%, signal=[illegible] | [illegible] |
| GOBP_CELLULAR_RESPONSE_TO_HORMONE_STIMULUS | GOBP_CELLULAR_RESPONSE_TO_HORMONE_STIMULUS | 536 | -0.3357203 | -1.2237711 | 0.04441504 | 0.35863453 | 0.32825429 | 869 | tags=11%, list=6%, signal=[illegible] | [illegible] |
| GOBP_HEMATOPOIETIC_STEM_CELL_HOMEOSTASIS | GOBP_HEMATOPOIETIC_STEM_CELL_HOMEOSTASIS | 26 | 0.58727208 | 1.39726448 | 0.04458599 | 0.35893392 | 0.32852833 | 1169 | tags=23%, list=8%, signal=[illegible] | [illegible] |
| GOBP_7_METHYLGUANOSINE_CAP_HYPERMETHYLATION | GOBP_7_METHYLGUANOSINE_CAP_HYPERMETHYLATION | 10 | 0.71792071 | 1.39842994 | 0.04470518 | 0.35948102 | 0.32902908 | 1584 | tags=50%, list=11%, signal=[illegible] | [illegible] |
| GOBP_PRENYLATION | GOBP_PRENYLATION | 9 | 0.73312891 | 1.39262467 | 0.04474967 | 0.35948102 | 0.32902908 | 1848 | tags=44%, list=13%, signal=[illegible] | [illegible] |
| GOBP_SKELETAL_MUSCLE_SATELLITE_CELL_ACTIVATION | GOBP_SKELETAL_MUSCLE_SATELLITE_CELL_ACTIVATION | 6 | -0.8860961 | -1.5535272 | 0.04490058 | 0.36015574 | 0.32964665 | 257 | tags=50%, list=2%, signal=[illegible] | [illegible] |
| GOBP_ALPHA_KETOGLUTARATE_TRANSPORT | GOBP_ALPHA_KETOGLUTARATE_TRANSPORT | 2 | -0.9686717 | -1.4049298 | 0.04496165 | 0.36015574 | 0.32964665 | 274 | tags=100%, list=2%, signal=[illegible] | [illegible] |
| GOBP_EMBRYO_DEVELOPMENT_ENDING_IN_BIRTH_OR_EGG_HATCHI | GOBP_EMBRYO_DEVELOPMENT_ENDING_IN_BIRTH_OR_EGG_HATC | 588 | 0.31155182 | 1.18302667 | 0.04497751 | 0.36015574 | 0.32964665 | 2575 | tags=21%, list=18%, signal=[illegible] | [illegible] |
| GOBP_POSITIVE_REGULATION_OF_DNA_METABOLIC_PROCESS | GOBP_POSITIVE_REGULATION_OF_DNA_METABOLIC_PROCESS | 257 | 0.35803162 | 1.25762823 | 0.04514851 | 0.36114004 | 0.33054756 | 5191 | tags=37%, list=35%, signal=[illegible] | [illegible] |
| GOBP_MITOTIC_DNA_REPLICATION | GOBP_MITOTIC_DNA_REPLICATION | 17 | 0.63994998 | 1.39863727 | 0.04532004 | 0.36212643 | 0.3314504 | 2553 | tags=71%, list=17%, signal=[illegible] | [illegible] |
| GOBP_AUDITORY_RECEPTOR_CELL_DEVELOPMENT | GOBP_AUDITORY_RECEPTOR_CELL_DEVELOPMENT | 21 | 0.61147258 | 1.39670032 | 0.04572154 | 0.36494634 | 0.33403143 | 4864 | tags=52%, list=33%, signal=[illegible] | [illegible] |
| GOBP_REGULATION_OF_CARBOHYDRATE_CATABOLIC_PROCESS | GOBP_REGULATION_OF_CARBOHYDRATE_CATABOLIC_PROCESS | 53 | -0.5382329 | -1.4421991 | 0.04592315 | 0.36616642 | 0.33514816 | 1033 | tags=21%, list=7%, signal=[illegible] | [illegible] |
| GOBP_CELL_SURFACE_RECEPTOR_SIGNALING_PATHWAY_VIA_JAK_STAT | GOBP_CELL_SURFACE_RECEPTOR_SIGNALING_PATHWAY_VIA_JAK_S | 94 | 0.44494636 | 1.35585429 | 0.04599816 | 0.36637559 | 0.3353396 | 4370 | tags=39%, list=30%, signal=[illegible] | [illegible] |
| GOBP_POSITIVE_REGULATION_OF_PEROXISOME_PROLIFERATOR_ACTI | GOBP_POSITIVE_REGULATION_OF_PEROXISOME_PROLIFERATOR_AC | 9 | 0.73206151 | 1.39059707 | 0.04622655 | 0.36762821 | 0.33648611 | 2804 | tags=33%, list=19%, signal=[illegible] | [illegible] |
| GOBP_PROTEIN_LOCALIZATION_TO_ENDOPLASMIC_RETICULUM | GOBP_PROTEIN_LOCALIZATION_TO_ENDOPLASMIC_RETICULUM | 30 | 0.56576572 | 1.38670428 | 0.04625332 | 0.36762821 | 0.33648611 | 1677 | tags=20%, list=11%, signal=[illegible] | [illegible] |
| GOBP_SPHINGOLIPID_BIOSYNTHETIC_PROCESS | GOBP_SPHINGOLIPID_BIOSYNTHETIC_PROCESS | 88 | 0.45039841 | 1.35468214 | 0.04644759 | 0.36839263 | 0.33718578 | 4185 | tags=40%, list=29%, signal=[illegible] | [illegible] |
| GOBP_DIGESTIVE_SYSTEM_DEVELOPMENT | GOBP_DIGESTIVE_SYSTEM_DEVELOPMENT | 101 | 0.43688992 | 1.34806034 | 0.04639941 | 0.36839263 | 0.33718578 | 5271 | tags=48%, list=36%, signal=[illegible] | [illegible] |
| GOBP_DNA_GEOMETRIC_CHANGE | GOBP_DNA_GEOMETRIC_CHANGE | 8 | 0.74990334 | 1.3869082 | 0.04669604 | 0.36997249 | 0.33863181 | 3493 | tags=75%, list=24%, signal=[illegible] | [illegible] |
| GOBP_G2_MI_TRANSITION_OF_MEIOTIC_CELL_CYCLE | GOBP_G2_MI_TRANSITION_OF_MEIOTIC_CELL_CYCLE | 8 | 0.74975313 | 1.38663039 | 0.04684288 | 0.37035459 | 0.33898154 | 2717 | tags=75%, list=19%, signal=[illegible] | [illegible] |
| GOBP_MORPHOGENESIS_OF_A_POLARIZED_EPITHELIUM | GOBP_MORPHOGENESIS_OF_A_POLARIZED_EPITHELIUM | 46 | 0.52055796 | 1.38411438 | 0.04679675 | 0.37035459 | 0.33898154 | 6410 | tags=67%, list=44%, signal=[illegible] | [illegible] |
| GOBP_CAMKK_AMPK_SIGNALING_CASCADE | GOBP_CAMKK_AMPK_SIGNALING_CASCADE | 9 | -0.8209368 | -1.5455155 | 0.04704426 | 0.37116536 | 0.33972363 | 389 | tags=33%, list=3%, signal=[illegible] | [illegible] |
| GOBP_CELLULAR_RESPONSE_TO_BIOTIC_STIMULUS | GOBP_CELLULAR_RESPONSE_TO_BIOTIC_STIMULUS | 192 | 0.38040456 | 1.28457758 | 0.04703357 | 0.37116536 | 0.33972363 | 2381 | tags=21%, list=16%, signal=[illegible] | [illegible] |
| GOBP_CENP_A_CONTAINING_CHROMATIN_ASSEMBLY | GOBP_CENP_A_CONTAINING_CHROMATIN_ASSEMBLY | 11 | 0.69804306 | 1.39020695 | 0.04716002 | 0.37168824 | 0.34020222 | 3666 | tags=82%, list=25%, signal=[illegible] | [illegible] |
| GOBP_MITOTIC_G1_S_TRANSITION_CHECKPOINT_SIGNALING | GOBP_MITOTIC_G1_S_TRANSITION_CHECKPOINT_SIGNALING | 16 | 0.64890613 | 1.39961785 | 0.04726484 | 0.37208233 | 0.34056292 | 1622 | tags=38%, list=11%, signal=[illegible] | [illegible] |
| GOBP_POSITIVE_REGULATION_OF_T_CELL_CYTOKINE_PRODUCTION | GOBP_POSITIVE_REGULATION_OF_T_CELL_CYTOKINE_PRODUCTION | 21 | 0.60955536 | 1.39232108 | 0.0473091 | 0.37208233 | 0.34056292 | 3985 | tags=48%, list=27%, signal=[illegible] | [illegible] |
| GOBP_SPLEEN_DEVELOPMENT | GOBP_SPLEEN_DEVELOPMENT | 35 | 0.54869074 | 1.38301651 | 0.04766734 | 0.3729473 | 0.34135462 | 3049 | tags=34%, list=21%, signal=[illegible] | [illegible] |
| GOBP_PARACRINE_SIGNALING | GOBP_PARACRINE_SIGNALING | 7 | 0.76782536 | 1.37969763 | 0.04752187 | 0.3729473 | 0.34135462 | 1497 | tags=29%, list=10%, signal=[illegible] | [illegible] |
| GOBP_PROTEIN_LOCALIZATION_TO_CHROMATIN | GOBP_PROTEIN_LOCALIZATION_TO_CHROMATIN | 49 | 0.51067512 | 1.37548262 | 0.04761905 | 0.3729473 | 0.34135462 | 1899 | tags=22%, list=13%, signal=[illegible] | [illegible] |
| GOBP_REGULATION_OF_HEPATIC_STELLATE_CELL_ACTIVATION | GOBP_REGULATION_OF_HEPATIC_STELLATE_CELL_ACTIVATION | 6 | 0.78796611 | 1.37122501 | 0.04750145 | 0.3729473 | 0.34135462 | 121 | tags=17%, list=1%, signal=[illegible] | [illegible] |
| GOBP_REGULATION_OF_PROTEIN_CONTAINING_COMPLEX_ASSEMBLY | GOBP_REGULATION_OF_PROTEIN_CONTAINING_COMPLEX_ASSEMB | 372 | -0.3549621 | -1.2521647 | 0.04761905 | 0.3729473 | 0.34135462 | 726 | tags=7%, list=5%, signal=[illegible] | [illegible] |
| GOBP_POSITIVE_REGULATION_OF_T_HELPER_2_CELL_DIFFERENTIATIO | GOBP_POSITIVE_REGULATION_OF_T_HELPER_2_CELL_DIFFERENTIA | 6 | 0.78774554 | 1.37084117 | 0.04793725 | 0.37466873 | 0.34293023 | 2608 | tags=83%, list=18%, signal=[illegible] | [illegible] |
| GOBP_PURINE_NUCLEOTIDE_SALVAGE | GOBP_PURINE_NUCLEOTIDE_SALVAGE | 17 | -0.7097946 | -1.5096348 | 0.04803128 | 0.37501343 | 0.34324572 | 187 | tags=18%, list=1%, signal=[illegible] | [illegible] |
| GOBP_NEGATIVE_REGULATION_OF_CAMP_PKA_SIGNAL_TRANSDUCTIO | GOBP_NEGATIVE_REGULATION_OF_CAMP_PKA_SIGNAL_TRANSDUC | 25 | -0.6461479 | -1.4850173 | 0.04810192 | 0.37517503 | 0.34339363 | 1510 | tags=28%, list=10%, signal=[illegible] | [illegible] |
| GOBP_MITOCHONDRIAL_TRNA_PROCESSING | GOBP_MITOCHONDRIAL_TRNA_PROCESSING | 17 | 0.63688121 | 1.39193035 | 0.04843482 | 0.37698856 | 0.34505354 | 553 | tags=18%, list=4%, signal=[illegible] | [illegible] |
| GOBP_DORSAL_VENTRAL_PATTERN_FORMATION | GOBP_DORSAL_VENTRAL_PATTERN_FORMATION | 58 | 0.49284597 | 1.37466052 | 0.04839558 | 0.37698856 | 0.34505354 | 4599 | tags=40%, list=31%, signal=[illegible] | [illegible] |
| GOBP_KERATINIZATION | GOBP_KERATINIZATION | 17 | 0.63674394 | 1.39163033 | 0.04859056 | 0.37756668 | 0.34558269 | 4788 | tags=71%, list=33%, signal=[illegible] | [illegible] |
| GOBP_DETOXIFICATION | GOBP_DETOXIFICATION | 118 | -0.4472318 | -1.3711675 | 0.04860964 | 0.37756668 | 0.34558269 | 999 | tags=19%, list=7%, signal=[illegible] | [illegible] |
| GOBP_REGULATION_OF_AUTOPHAGY_OF_MITOCHONDRION | GOBP_REGULATION_OF_AUTOPHAGY_OF_MITOCHONDRION | 50 | -0.5426706 | -1.4416982 | 0.04882629 | 0.37885772 | 0.34676436 | 1588 | tags=36%, list=11%, signal=[illegible] | [illegible] |
| GOBP_INTRACELLULAR_SODIUM_ION_HOMEOSTASIS | GOBP_INTRACELLULAR_SODIUM_ION_HOMEOSTASIS | 15 | -0.7274941 | -1.5065999 | 0.04894104 | 0.37898681 | 0.34688252 | 696 | tags=20%, list=5%, signal=[illegible] | [illegible] |
| GOBP_NEGATIVE_REGULATION_OF_SIGNAL_TRANSDUCTION_BY_P53_ | GOBP_NEGATIVE_REGULATION_OF_SIGNAL_TRANSDUCTION_BY_PS | 42 | 0.52920017 | 1.38118878 | 0.04894384 | 0.37898681 | 0.34688252 | 4613 | tags=40%, list=32%, signal=[illegible] | [illegible] |
| GOBP_CELLULAR_RESPONSE_TO_FOLLICLE_STIMULATING_HORMONE_ | GOBP_CELLULAR_RESPONSE_TO_FOLLICLE_STIMULATING_HORMON | 9 | 0.72874958 | 1.38430585 | 0.04932802 | 0.38014577 | 0.3479433 | 1519 | tags=33%, list=10%, signal=[illegible] | [illegible] |
| GOBP_REGULATION_OF_ACTIN_NUCLEATION | GOBP_REGULATION_OF_ACTIN_NUCLEATION | 33 | 0.555228 | 1.38412177 | 0.04939719 | 0.38014577 | 0.3479433 | 1412 | tags=30%, list=10%, signal=[illegible] | [illegible] |
| GOBP_CELL_SURFACE_RECEPTOR_PROTEIN_TYROSINE_PHOSPHATASE_ | GOBP_CELL_SURFACE_RECEPTOR_PROTEIN_TYROSINE_PHOSPHATAS | 7 | 0.7662741 | 1.37691018 | 0.04927114 | 0.38014577 | 0.3479433 | 1938 | tags=43%, list=13%, signal=[illegible] | [illegible] |
| GOBP_REGULATION_OF_TUBE_SIZE | GOBP_REGULATION_OF_TUBE_SIZE | 105 | -0.4584786 | -1.3751828 | 0.04925146 | 0.38014577 | 0.3479433 | 408 | tags=8%, list=3%, signal=[illegible] | [illegible] |
| GOBP_REGULATION_OF_DEVELOPMENTAL_GROWTH | GOBP_REGULATION_OF_DEVELOPMENTAL_GROWTH | 243 | -0.3825815 | -1.2903435 | 0.04935275 | 0.38014577 | 0.3479433 | 781 | tags=9%, list=5%, signal=[illegible] | [illegible] |
| GOBP_CARBOHYDRATE_DERIVATIVE_METABOLIC_PROCESS | GOBP_CARBOHYDRATE_DERIVATIVE_METABOLIC_PROCESS | 835 | -0.3138489 | -1.1837538 | 0.04933969 | 0.38014577 | 0.3479433 | 1042 | tags=11%, list=7%, signal=[illegible] | [illegible] |
| GOBP_REGULATION_OF_LEUKOCYTE_DIFFERENTIATION | GOBP_REGULATION_OF_LEUKOCYTE_DIFFERENTIATION | 280 | 0.34970084 | 1.23925218 | 0.04945718 | 0.38021787 | 0.34800929 | 4344 | tags=33%, list=30%, signal=[illegible] | [illegible] |
| GOBP_MYELOID_LEUKOCYTE_ACTIVATION | GOBP_MYELOID_LEUKOCYTE_ACTIVATION | 194 | 0.37735055 | 1.27735657 | 0.04963987 | 0.38123215 | 0.34893766 | 4334 | tags=35%, list=30%, signal=[illegible] | [illegible] |
| GOBP_NUCLEOTIDE_SUGAR_BIOSYNTHETIC_PROCESS | GOBP_NUCLEOTIDE_SUGAR_BIOSYNTHETIC_PROCESS | 28 | 0.57334102 | 1.38531491 | 0.04972831 | 0.38152125 | 0.34920226 | 3154 | tags=43%, list=22%, signal=[illegible] | [illegible] |
| GOBP_BONE_REMODELING | GOBP_BONE_REMODELING | 70 | 0.4705834 | 1.36105799 | 0.04982143 | 0.38184566 | 0.34949919 | 4572 | tags=49%, list=31%, signal=[illegible] | [illegible] |
| GOBP_GLYCOSAMINOGLYCAN_CATABOLIC_PROCESS | GOBP_GLYCOSAMINOGLYCAN_CATABOLIC_PROCESS | 19 | 0.62053441 | 1.38944158 | 0.05011783 | 0.38275192 | 0.35032868 | 3308 | tags=53%, list=23%, signal=[illegible] | [illegible] |
| GOBP_POSITIVE_REGULATION_OF_BINDING | GOBP_POSITIVE_REGULATION_OF_BINDING | 33 | 0.55482847 | 1.38312579 | 0.05006698 | 0.38275192 | 0.35032868 | 3174 | tags=36%, list=22%, signal=[illegible] | [illegible] |
| GOBP_AMINO_SUGAR_METABOLIC_PROCESS | GOBP_AMINO_SUGAR_METABOLIC_PROCESS | 37 | 0.54004317 | 1.37659927 | 0.05014351 | 0.38275192 | 0.35032868 | 3919 | tags=51%, list=27%, signal=[illegible] | [illegible] |
| GOBP_T_CELL_PROLIFERATION | GOBP_T_CELL_PROLIFERATION | 157 | 0.3936524 | 1.29274637 | 0.0500956 | 0.38275192 | 0.35032868 | 4580 | tags=37%, list=31%, signal=[illegible] | [illegible] |
| GOBP_VESICLE_MEDIATED_TRANSPORT | GOBP_VESICLE_MEDIATED_TRANSPORT | 1355 | 0.27810274 | 1.11581175 | 0.05022295 | 0.38296913 | 0.35052749 | 2957 | tags=21%, list=20%, signal=[illegible] | [illegible] |
| GOBP_DEOXYRIBONUCLEOSIDE_MONOPHOSPHATE_BIOSYNTHETIC_PR | GOBP_DEOXYRIBONUCLEOSIDE_MONOPHOSPHATE_BIOSYNTHETIC_ | 9 | 0.72723879 | 1.38143601 | 0.05036184 | 0.3836387 | 0.35114034 | 3775 | tags=78%, list=26%, signal=[illegible] | [illegible] |
| GOBP_PLASMA_MEMBRANE_ORGANIZATION | GOBP_PLASMA_MEMBRANE_ORGANIZATION | 127 | -0.4361582 | -1.3528366 | 0.05057956 | 0.38426835 | 0.35171665 | 454 | tags=8%, list=3%, signal=[illegible] | [illegible] |
| GOBP_CELLULAR_RESPONSE_TO_METAL_ION | GOBP_CELLULAR_RESPONSE_TO_METAL_ION | 132 | -0.4325683 | -1.347652 | 0.05052854 | 0.38426835 | 0.35171665 | 734 | tags=11%, list=5%, signal=[illegible] | [illegible] |
| GOBP_NEPHRON_EPITHELIUM_DEVELOPMENT | GOBP_NEPHRON_EPITHELIUM_DEVELOPMENT | 94 | 0.44140832 | 1.34507305 | 0.05059798 | 0.38426835 | 0.35171665 | 4167 | tags=41%, list=28%, signal=[illegible] | [illegible] |
| GOBP_ENDOLYSOSOMAL_TOLL_LIKE_RECEPTOR_SIGNALING_PATHWAY | GOBP_ENDOLYSOSOMAL_TOLL_LIKE_RECEPTOR_SIGNALING_PATHW | 41 | 0.52760629 | 1.37191509 | 0.05071967 | 0.38480348 | 0.35220646 | 3681 | tags=32%, list=25%, signal=[illegible] | [illegible] |
| GOBP_NADPLUS_BIOSYNTHETIC_PROCESS | GOBP_NADPLUS_BIOSYNTHETIC_PROCESS | 12 | 0.68164767 | 1.38457699 | 0.05094396 | 0.38494979 | 0.35234037 | 2096 | tags=25%, list=14%, signal=[illegible] | [illegible] |
| GOBP_REGULATION_OF_RECEPTOR_SIGNALING_PATHWAY_VIA_JAK_ST | GOBP_REGULATION_OF_RECEPTOR_SIGNALING_PATHWAY_VIA_JAK | 56 | 0.49521065 | 1.37093801 | 0.05090973 | 0.38494979 | 0.35234037 | 5189 | tags=57%, list=35%, signal=[illegible] | [illegible] |
| GOBP_NEGATIVE_REGULATION_OF_DEFENSE_RESPONSE | GOBP_NEGATIVE_REGULATION_OF_DEFENSE_RESPONSE | 216 | 0.36836251 | 1.26552828 | 0.05084413 | 0.38494979 | 0.35234037 | 4334 | tags=33%, list=30%, signal=[illegible] | [illegible] |
| GOBP_EMBRYONIC_ORGAN_DEVELOPMENT | GOBP_EMBRYONIC_ORGAN_DEVELOPMENT | 370 | 0.33215913 | 1.21241022 | 0.05085797 | 0.38494979 | 0.35234037 | 4262 | tags=36%, list=29%, signal=[illegible] | [illegible] |
| GOBP_POSITIVE_REGULATION_OF_INTERLEUKIN_12_PRODUCTION | GOBP_POSITIVE_REGULATION_OF_INTERLEUKIN_12_PRODUCTION | 35 | 0.54590905 | 1.37600505 | 0.05121704 | 0.3866243 | 0.35387303 | 2744 | tags=34%, list=19%, signal=[illegible] | [illegible] |
| GOBP_PRE_MIRNA_PROCESSING | GOBP_PRE_MIRNA_PROCESSING | 14 | 0.66090524 | 1.3879755 | 0.05140329 | 0.38764071 | 0.35480334 | 788 | tags=14%, list=5%, signal=[illegible] | [illegible] |
| GOBP_RESPONSE_TO_INTERLEUKIN_4 | GOBP_RESPONSE_TO_INTERLEUKIN_4 | 35 | 0.54475602 | 1.37309874 | 0.05189317 | 0.38938023 | 0.3563955 | 4436 | tags=46%, list=30%, signal=[illegible] | [illegible] |
| GOBP_NEGATIVE_REGULATION_OF_VITAMIN_METABOLIC_PROCESS | GOBP_NEGATIVE_REGULATION_OF_VITAMIN_METABOLIC_PROCESS | 5 | 0.81337899 | 1.3654048 | 0.05184969 | 0.38938023 | 0.3563955 | 2562 | tags=60%, list=18%, signal=[illegible] | [illegible] |
| GOBP_ADENOSINE_TO_INOSINE_EDITING | GOBP_ADENOSINE_TO_INOSINE_EDITING | 3 | 0.89115586 | 1.35959944 | 0.05184297 | 0.38938023 | 0.3563955 | 1537 | tags=67%, list=11%, signal=[illegible] | [illegible] |
| GOBP_REGULATION_OF_MEIOSIS_I | GOBP_REGULATION_OF_MEIOSIS_I | 3 | 0.89094797 | 1.35928227 | 0.05184297 | 0.38938023 | 0.3563955 | 1583 | tags=67%, list=11%, signal=[illegible] | [illegible] |
| GOBP_REGULATION_OF_GROWTH | GOBP_REGULATION_OF_GROWTH | 494 | -0.3381881 | -1.223747 | 0.05182927 | 0.38938023 | 0.3563955 | 781 | tags=7%, list=5%, signal=[illegible] | [illegible] |
| GOBP_ISOPENTENYL_DIPHOSPHATE_BIOSYNTHETIC_PROCESS_MEVALO | GOBP_ISOPENTENYL_DIPHOSPHATE_BIOSYNTHETIC_PROCESS_MEV | 5 | 0.8129983 | 1.36476574 | 0.05199534 | 0.38975748 | 0.35674079 | 294 | tags=40%, list=2%, signal=[illegible] | [illegible] |
| GOBP_IN_UTERO_EMBRYONIC_DEVELOPMENT | GOBP_IN_UTERO_EMBRYONIC_DEVELOPMENT | 355 | 0.33484775 | 1.21683704 | 0.05207688 | 0.38997947 | 0.35694398 | 3023 | tags=24%, list=21%, signal=[illegible] | [illegible] |
| GOBP_NEGATIVE_REGULATION_OF_MULTICELLULAR_ORGANISM_GRO | GOBP_NEGATIVE_REGULATION_OF_MULTICELLULAR_ORGANISM_G | 12 | 0.68029867 | 1.38183687 | 0.05229248 | 0.39120399 | 0.35806477 | 1488 | tags=33%, list=10%, signal=[illegible] | [illegible] |
| GOBP_AMP_METABOLIC_PROCESS | GOBP_AMP_METABOLIC_PROCESS | 21 | -0.6690292 | -1.4811062 | 0.05238995 | 0.39154323 | 0.35837527 | 187 | tags=24%, list=1%, signal=[illegible] | [illegible] |
| GOBP_CIRCADIAN_SLEEP_WAKE_CYCLE_SLEEP | GOBP_CIRCADIAN_SLEEP_WAKE_CYCLE_SLEEP | 10 | -0.7992657 | -1.5338572 | 0.05261548 | 0.39233149 | 0.35909676 | 427 | tags=30%, list=3%, signal=[illegible] | [illegible] |
| GOBP_PYRIMIDINE_RIBONUCLEOSIDE_TRIPHOSPHATE_METABOLIC_PR | GOBP_PYRIMIDINE_RIBONUCLEOSIDE_TRIPHOSPHATE_METABOLIC_ | 15 | 0.65304998 | 1.38973256 | 0.05270436 | 0.39233149 | 0.35909676 | 2528 | tags=40%, list=17%, signal=[illegible] | [illegible] |
| GOBP_NEURON_FATE_COMMITMENT | GOBP_NEURON_FATE_COMMITMENT | 33 | 0.55308972 | 1.37879127 | 0.0525787 | 0.39233149 | 0.35909676 | 2575 | tags=30%, list=18%, signal=[illegible] | [illegible] |
| GOBP_RNA_PROCESSING | GOBP_RNA_PROCESSING | 918 | 0.29043629 | 1.13965217 | 0.05269008 | 0.39233149 | 0.35909676 | 5673 | tags=35%, list=39%, signal=[illegible] | [illegible] |
| GOBP_CELLULAR_RESPONSE_TO_WATER_STIMULUS | GOBP_CELLULAR_RESPONSE_TO_WATER_STIMULUS | 5 | -0.8953614 | -1.5204757 | 0.05293367 | 0.39325897 | 0.35994567 | 135 | tags=40%, list=1%, signal=[illegible] | [illegible] |
| GOBP_REGULATION_OF_CELL_DIVISION | GOBP_REGULATION_OF_CELL_DIVISION | 146 | 0.40019372 | 1.30054256 | 0.052902 | 0.39325897 | 0.35994567 | 3614 | tags=29%, list=25%, signal=[illegible] | [illegible] |
| GOBP_PHYSIOLOGICAL_CARDIAC_MUSCLE_HYPERTROPHY | GOBP_PHYSIOLOGICAL_CARDIAC_MUSCLE_HYPERTROPHY | 18 | -0.6890153 | -1.4780577 | 0.05318853 | 0.3935951 | 0.36025333 | 1400 | tags=39%, list=10%, signal=[illegible] | [illegible] |
| GOBP_NEGATIVE_REGULATION_OF_ACTION_POTENTIAL | GOBP_NEGATIVE_REGULATION_OF_ACTION_POTENTIAL | 2 | -0.9637141 | -1.3977395 | 0.05316054 | 0.3935951 | 0.36025333 | 160 | tags=100%, list=1%, signal=[illegible] | [illegible] |
| GOBP_DERMATAN_SULFATE_PROTEOGLYCAN_METABOLIC_PROCESS | GOBP_DERMATAN_SULFATE_PROTEOGLYCAN_METABOLIC_PROCESS | 17 | 0.63314326 | 1.3837609 | 0.05310699 | 0.3935951 | 0.36025333 | 3162 | tags=35%, list=22%, signal=[illegible] | [illegible] |
| GOBP_MESODERMAL_CELL_DIFFERENTIATION | GOBP_MESODERMAL_CELL_DIFFERENTIATION | 26 | 0.57781477 | 1.37476324 | 0.05307856 | 0.3935951 | 0.36025333 | 2318 | tags=23%, list=16%, signal=[illegible] | [illegible] |
| GOBP_POSITIVE_REGULATION_OF_CALCIUM_ION_IMPORT_ACROSS_PL | GOBP_POSITIVE_REGULATION_OF_CALCIUM_ION_IMPORT_ACROSS_ | 6 | -0.8773245 | -1.5381487 | 0.05388069 | 0.39402359 | 0.36064552 | 461 | tags=50%, list=3%, signal=[illegible] | [illegible] |
| GOBP_MITOCHONDRIAL_RNA_MODIFICATION | GOBP_MITOCHONDRIAL_RNA_MODIFICATION | 16 | 0.64302335 | 1.38692935 | 0.05392841 | 0.39402359 | 0.36064552 | 553 | tags=19%, list=4%, signal=[illegible] | [illegible] |
| GOBP_ANAPHASE_PROMOTING_COMPLEX_DEPENDENT_CATABOLIC_P | GOBP_ANAPHASE_PROMOTING_COMPLEX_DEPENDENT_CATABOLIC | 24 | 0.58867162 | 1.37879082 | 0.05347938 | 0.39402359 | 0.36064552 | 2002 | tags=29%, list=14%, signal=[illegible] | [illegible] |
| GOBP_MAINTENANCE_OF_PROTEIN_LOCALIZATION_IN_ENDOPLASMIC_ | GOBP_MAINTENANCE_OF_PROTEIN_LOCALIZATION_IN_ENDOPLASM | 10 | 0.70760879 | 1.37834347 | 0.05376504 | 0.39402359 | 0.36064552 | 1677 | tags=50%, list=11%, signal=[illegible] | [illegible] |
| GOBP_SPHINGOID_BIOSYNTHETIC_PROCESS | GOBP_SPHINGOID_BIOSYNTHETIC_PROCESS | 11 | 0.69177411 | 1.37772186 | 0.053537 | 0.39402359 | 0.36064552 | 3972 | tags=55%, list=27%, signal=[illegible] | [illegible] |
| GOBP_CEREBELLAR_GRANULAR_LAYER_DEVELOPMENT | GOBP_CEREBELLAR_GRANULAR_LAYER_DEVELOPMENT | 9 | 0.72398039 | 1.37524649 | 0.0534633 | 0.39402359 | 0.36064552 | 487 | tags=33%, list=3%, signal=[illegible] | [illegible] |
| GOBP_GLYCOSAMINOGLYCAN_BIOSYNTHETIC_PROCESS | GOBP_GLYCOSAMINOGLYCAN_BIOSYNTHETIC_PROCESS | 26 | 0.57714255 | 1.37316386 | 0.05356851 | 0.39402359 | 0.36064552 | 3437 | tags=46%, list=23%, signal=[illegible] | [illegible] |
| GOBP_HOMOLOGOUS_CHROMOSOME_PAIRING_AT_MEIOSIS | GOBP_HOMOLOGOUS_CHROMOSOME_PAIRING_AT_MEIOSIS | 29 | 0.56343775 | 1.37090079 | 0.05391591 | 0.39402359 | 0.36064552 | 3244 | tags=41%, list=22%, signal=[illegible] | [illegible] |
| GOBP_DE_NOVO_CENTRIOLE_ASSEMBLY | GOBP_DE_NOVO_CENTRIOLE_ASSEMBLY | 4 | 0.85080189 | 1.36637086 | 0.0537414 | 0.39402359 | 0.36064552 | 1451 | tags=50%, list=10%, signal=[illegible] | [illegible] |
| GOBP_SKIN_DEVELOPMENT | GOBP_SKIN_DEVELOPMENT | 201 | 0.37221248 | 1.26525154 | 0.05358194 | 0.39402359 | 0.36064552 | 4788 | tags=49%, list=33%, signal=[illegible] | [illegible] |
| GOBP_MICROTUBULE_BASED_TRANSPORT | GOBP_MICROTUBULE_BASED_TRANSPORT | 213 | 0.36865134 | 1.26393432 | 0.05341712 | 0.39402359 | 0.36064552 | 3131 | tags=29%, list=21%, signal=[illegible] | [illegible] |
| GOBP_NEGATIVE_REGULATION_OF_RESPONSE_TO_EXTERNAL_STIMULU | GOBP_NEGATIVE_REGULATION_OF_RESPONSE_TO_EXTERNAL_STIM | 305 | 0.34205576 | 1.22659932 | 0.05378286 | 0.39402359 | 0.36064552 | 4278 | tags=35%, list=29%, signal=[illegible] | [illegible] |
| GOBP_RESPONSE_TO_VIRUS | GOBP_RESPONSE_TO_VIRUS | 350 | 0.33468594 | 1.21563325 | 0.05389344 | 0.39402359 | 0.36064552 | 4833 | tags=35%, list=33%, signal=[illegible] | [illegible] |
| GOBP_RRNA_3_END_PROCESSING | GOBP_RRNA_3_END_PROCESSING | 12 | 0.67817743 | 1.37752816 | 0.0540905 | 0.39482386 | 0.36137799 | 1334 | tags=33%, list=9%, signal=[illegible] | [illegible] |
| GOBP_NEGATIVE_REGULATION_OF_VASCULAR_ENDOTHELIAL_CELL_PR | GOBP_NEGATIVE_REGULATION_OF_VASCULAR_ENDOTHELIAL_CELL_ | 5 | -0.8946299 | -1.5192335 | 0.05484694 | 0.39616186 | 0.36260266 | 252 | tags=60%, list=2%, signal=[illegible] | [illegible] |
| GOBP_REGULATION_OF_SODIUM_ION_EXPORT_ACROSS_PLASMA_MEN | GOBP_REGULATION_OF_SODIUM_ION_EXPORT_ACROSS_PLASMA_N | 5 | -0.8946099 | -1.5191994 | 0.05484694 | 0.39616186 | 0.36260266 | 696 | tags=40%, list=5%, signal=[illegible] | [illegible] |
| GOBP_POSITIVE_REGULATION_OF_AUTOPHAGY_OF_MITOCHONDRION | GOBP_POSITIVE_REGULATION_OF_AUTOPHAGY_OF_MITOCHONDRI | 24 | -0.6467402 | -1.4695249 | 0.05455983 | 0.39616186 | 0.36260266 | 1726 | tags=46%, list=12%, signal=[illegible] | [illegible] |
| GOBP_INTERLEUKIN_17_MEDIATED_SIGNALING_PATHWAY | GOBP_INTERLEUKIN_17_MEDIATED_SIGNALING_PATHWAY | 12 | 0.67769845 | 1.37655526 | 0.05498951 | 0.39616186 | 0.36260266 | 3316 | tags=50%, list=23%, signal=[illegible] | [illegible] |
| GOBP_HOMEOSTASIS_OF_NUMBER_OF_CELLS_WITHIN_A_TISSUE | GOBP_HOMEOSTASIS_OF_NUMBER_OF_CELLS_WITHIN_A_TISSUE | 32 | 0.5542717 | 1.37254272 | 0.05497995 | 0.39616186 | 0.36260266 | 4148 | tags=47%, list=28%, signal=[illegible] | [illegible] |
| GOBP_IMMATURE_T_CELL_PROLIFERATION | GOBP_IMMATURE_T_CELL_PROLIFERATION | 9 | 0.72201928 | 1.37152123 | 0.05464481 | 0.39616186 | 0.36260266 | 285 | tags=22%, list=2%, signal=[illegible] | [illegible] |
| GOBP_HEMATOPOIETIC_STEM_CELL_MIGRATION | GOBP_HEMATOPOIETIC_STEM_CELL_MIGRATION | 9 | 0.72166052 | 1.37083974 | 0.0547925 | 0.39616186 | 0.36260266 | 126 | tags=11%, list=1%, signal=[illegible] | [illegible] |
| GOBP_POSITIVE_REGULATION_OF_FIBROBLAST_PROLIFERATION | GOBP_POSITIVE_REGULATION_OF_FIBROBLAST_PROLIFERATION | 45 | 0.51630397 | 1.36608549 | 0.05481813 | 0.39616186 | 0.36260266 | 3589 | tags=33%, list=25%, signal=[illegible] | [illegible] |
| GOBP_CELLULAR_RESPONSE_TO_TOXIC_SUBSTANCE | GOBP_CELLULAR_RESPONSE_TO_TOXIC_SUBSTANCE | 106 | -0.4526329 | -1.3601016 | 0.05443505 | 0.39616186 | 0.36260266 | 999 | tags=19%, list=7%, signal=[illegible] | [illegible] |
| GOBP_MAST_CELL_ACTIVATION | GOBP_MAST_CELL_ACTIVATION | 54 | 0.49500232 | 1.35884459 | 0.05501222 | 0.39616186 | 0.36260266 | 5678 | tags=50%, list=39%, signal=[illegible] | [illegible] |
| GOBP_NEGATIVE_REGULATION_OF_ESTABLISHMENT_OF_PROTEIN_LO | GOBP_NEGATIVE_REGULATION_OF_ESTABLISHMENT_OF_PROTEIN_ | 106 | -0.4520935 | -1.3584808 | 0.05486879 | 0.39616186 | 0.36260266 | 195 | tags=6%, list=1%, signal=[illegible] | [illegible] |
| GOBP_ADAPTIVE_IMMUNE_RESPONSE_BASED_ON_SOMATIC_RECOMB | GOBP_ADAPTIVE_IMMUNE_RESPONSE_BASED_ON_SOMATIC_RECO | 257 | 0.35345046 | 1.24153635 | 0.05485149 | 0.39616186 | 0.36260266 | 4875 | tags=37%, list=33%, signal=[illegible] | [illegible] |
| GOBP_NEGATIVE_REGULATION_OF_IMMUNE_SYSTEM_PROCESS | GOBP_NEGATIVE_REGULATION_OF_IMMUNE_SYSTEM_PROCESS | 423 | 0.32383965 | 1.19810179 | 0.05444305 | 0.39616186 | 0.36260266 | 2821 | tags=22%, list=19%, signal=[illegible] | [illegible] |
| GOBP_POSITIVE_REGULATION_OF_GENE_EXPRESSION | GOBP_POSITIVE_REGULATION_OF_GENE_EXPRESSION | 939 | 0.28859057 | 1.13402133 | 0.05489154 | 0.39616186 | 0.36260266 | 4273 | tags=30%, list=29%, signal=[illegible] | [illegible] |
| GOBP_FOREBRAIN_DEVELOPMENT | GOBP_FOREBRAIN_DEVELOPMENT | 310 | 0.34072533 | 1.2239169 | 0.05507363 | 0.39622419 | 0.36265971 | 4544 | tags=35%, list=31%, signal=[illegible] | [illegible] |
| GOBP_TRANSPOSABLE_ELEMENT_SILENCING_BY_PIRNA_MEDIATED_HI | GOBP_TRANSPOSABLE_ELEMENT_SILENCING_BY_PIRNA_MEDIATED | 2 | 0.93415801 | 1.33736691 | 0.05529658 | 0.39744745 | 0.36377934 | 936 | tags=50%, list=6%, signal=[illegible] | [illegible] |
| GOBP_GONADOTROPIN_SECRETION | GOBP_GONADOTROPIN_SECRETION | 12 | 0.67741881 | 1.37598723 | 0.05543902 | 0.3980903 | 0.36436774 | 3364 | tags=50%, list=23%, signal=[illegible] | [illegible] |
| GOBP_POSITIVE_REGULATION_OF_RESPONSE_TO_CYTOKINE_STIMULU | GOBP_POSITIVE_REGULATION_OF_RESPONSE_TO_CYTOKINE_STIMU | 60 | 0.48291045 | 1.35524665 | 0.05550661 | 0.39819497 | 0.36446354 | 5009 | tags=40%, list=34%, signal=[illegible] | [illegible] |
| GOBP_MISMATCH_REPAIR | GOBP_MISMATCH_REPAIR | 29 | 0.56161786 | 1.36647281 | 0.05556472 | 0.39823147 | 0.36449694 | 3856 | tags=41%, list=26%, signal=[illegible] | [illegible] |
| GOBP_IMP_SALVAGE | GOBP_IMP_SALVAGE | 5 | -0.8927884 | -1.5161062 | 0.05580357 | 0.39880174 | 0.3650189 | 23 | tags=40%, list=0%, signal=[illegible] | [illegible] |
| GOBP_REGULATION_OF_HEART_RATE_BY_CHEMICAL_SIGNAL | GOBP_REGULATION_OF_HEART_RATE_BY_CHEMICAL_SIGNAL | 5 | -0.8927679 | -1.5160714 | 0.05580357 | 0.39880174 | 0.3650189 | 19 | tags=40%, list=0%, signal=[illegible] | [illegible] |
| GOBP_NEGATIVE_REGULATION_OF_BLOOD_CIRCULATION | GOBP_NEGATIVE_REGULATION_OF_BLOOD_CIRCULATION | 14 | -0.7346659 | -1.501036 | 0.05571677 | 0.39880174 | 0.3650189 | 427 | tags=29%, list=3%, signal=[illegible] | [illegible] |
| GOBP_NEGATIVE_REGULATION_OF_DNA_DAMAGE_RESPONSE_SIGNAL | GOBP_NEGATIVE_REGULATION_OF_DNA_DAMAGE_RESPONSE_SIG | 15 | 0.64913038 | 1.38139139 | 0.05593116 | 0.39933361 | 0.36550572 | 2839 | tags=47%, list=19%, signal=[illegible] | [illegible] |
| GOBP_REGULATION_OF_ARP2_3_COMPLEX_MEDIATED_ACTIN_NUCLE | GOBP_REGULATION_OF_ARP2_3_COMPLEX_MEDIATED_ACTIN_NUC | 27 | 0.57205681 | 1.3726136 | 0.05614842 | 0.39992649 | 0.36604838 | 1412 | tags=33%, list=10%, signal=[illegible] | [illegible] |
| GOBP_THIRD_VENTRICLE_DEVELOPMENT | GOBP_THIRD_VENTRICLE_DEVELOPMENT | 4 | 0.84796471 | 1.36181441 | 0.05608435 | 0.39992649 | 0.36604838 | 380 | tags=25%, list=3%, signal=[illegible] | [illegible] |
| GOBP_NEGATIVE_REGULATION_OF_CHEMOKINE_C_X_C_MOTIF_LIGAN | GOBP_NEGATIVE_REGULATION_OF_CHEMOKINE_C_X_C_MOTIF_LIG | 4 | 0.84771667 | 1.36141607 | 0.05623078 | 0.39992649 | 0.36604838 | 1779 | tags=50%, list=12%, signal=[illegible] | [illegible] |
| GOBP_DUCTUS_ARTERIOSUS_CLOSURE | GOBP_DUCTUS_ARTERIOSUS_CLOSURE | 2 | 0.93290058 | 1.33556674 | 0.05626105 | 0.39992649 | 0.36604838 | 984 | tags=100%, list=7%, signal=[illegible] | [illegible] |
| GOBP_REGULATION_OF_LYMPHOCYTE_DIFFERENTIATION | GOBP_REGULATION_OF_LYMPHOCYTE_DIFFERENTIATION | 193 | 0.37391001 | 1.26464998 | 0.05628043 | 0.39992649 | 0.36604838 | 4184 | tags=36%, list=29%, signal=[illegible] | [illegible] |
| GOBP_NEGATIVE_REGULATION_OF_ENDOPLASMIC_RETICULUM_CALCI | GOBP_NEGATIVE_REGULATION_OF_ENDOPLASMIC_RETICULUM_CA | 6 | -0.8744821 | -1.5331653 | 0.0574086 | 0.39996035 | 0.36607937 | 7 | tags=33%, list=0%, signal=[illegible] | [illegible] |
| GOBP_PRIMARY_ALCOHOL_CATABOLIC_PROCESS | GOBP_PRIMARY_ALCOHOL_CATABOLIC_PROCESS | 10 | -0.794877 | -1.5254349 | 0.05750994 | 0.39996035 | 0.36607937 | 794 | tags=30%, list=5%, signal=[illegible] | [illegible] |
| GOBP_NEGATIVE_REGULATION_OF_CALCIUM_ION_TRANSMEMBRANE_ | GOBP_NEGATIVE_REGULATION_OF_CALCIUM_ION_TRANSMEMBRA | 4 | -0.9145647 | -1.4903911 | 0.05735897 | 0.39996035 | 0.36607937 | 1071 | tags=75%, list=7%, signal=[illegible] | [illegible] |
| GOBP_PHOSPHOCREATINE_METABOLIC_PROCESS | GOBP_PHOSPHOCREATINE_METABOLIC_PROCESS | 3 | -0.9404521 | -1.4493488 | 0.05709135 | 0.39996035 | 0.36607937 | 341 | tags=67%, list=2%, signal=[illegible] | [illegible] |
| GOBP_MYELOID_DENDRITIC_CELL_DIFFERENTIATION | GOBP_MYELOID_DENDRITIC_CELL_DIFFERENTIATION | 16 | 0.63805735 | 1.37621824 | 0.05749264 | 0.39996035 | 0.36607937 | 2808 | tags=38%, list=19%, signal=[illegible] | [illegible] |
| GOBP_FIBRINOLYSIS | GOBP_FIBRINOLYSIS | 12 | 0.67578878 | 1.37267629 | 0.05693737 | 0.39996035 | 0.36607937 | 3320 | tags=42%, list=23%, signal=[illegible] | [illegible] |
| GOBP_REGULATION_OF_MEIOTIC_CELL_CYCLE_PHASE_TRANSITION | GOBP_REGULATION_OF_MEIOTIC_CELL_CYCLE_PHASE_TRANSITION | 10 | 0.70300469 | 1.36937518 | 0.05747809 | 0.39996035 | 0.36607937 | 2717 | tags=70%, list=19%, signal=[illegible] | [illegible] |
| GOBP_CLEAVAGE_FURROW_FORMATION | GOBP_CLEAVAGE_FURROW_FORMATION | 8 | 0.73988329 | 1.36837662 | 0.05668135 | 0.39996035 | 0.36607937 | 1721 | tags=38%, list=12%, signal=[illegible] | [illegible] |
| GOBP_NEGATIVE_REGULATION_OF_NF_KAPPAB_TRANSCRIPTION_FACT | GOBP_NEGATIVE_REGULATION_OF_NF_KAPPAB_TRANSCRIPTION_F | 34 | 0.54579007 | 1.36731005 | 0.0568545 | 0.39996035 | 0.36607937 | 4320 | tags=41%, list=30%, signal=[illegible] | [illegible] |
| GOBP_INTRA_GOLGI_VESICLE_MEDIATED_TRANSPORT | GOBP_INTRA_GOLGI_VESICLE_MEDIATED_TRANSPORT | 32 | 0.55186165 | 1.3665747 | 0.05748663 | 0.39996035 | 0.36607937 | 3165 | tags=28%, list=22%, signal=[illegible] | [illegible] |
| GOBP_ENUCLEATE_ERYTHROCYTE_DEVELOPMENT | GOBP_ENUCLEATE_ERYTHROCYTE_DEVELOPMENT | 7 | 0.75976207 | 1.36520879 | 0.05641399 | 0.39996035 | 0.36607937 | 887 | tags=43%, list=6%, signal=[illegible] | [illegible] |
| GOBP_REGULATION_OF_KIDNEY_DEVELOPMENT | GOBP_REGULATION_OF_KIDNEY_DEVELOPMENT | 29 | 0.56046683 | 1.36367226 | 0.05688376 | 0.39996035 | 0.36607937 | 3871 | tags=52%, list=26%, signal=[illegible] | [illegible] |
| GOBP_MACROPHAGE_DIFFERENTIATION | GOBP_MACROPHAGE_DIFFERENTIATION | 55 | 0.49212404 | 1.35682736 | 0.05698304 | 0.39996035 | 0.36607937 | 2762 | tags=29%, list=19%, signal=[illegible] | [illegible] |
| GOBP_MITOTIC_G2_M_TRANSITION_CHECKPOINT | GOBP_MITOTIC_G2_M_TRANSITION_CHECKPOINT | 55 | 0.49201776 | 1.35653435 | 0.05733263 | 0.39996035 | 0.36607937 | 5169 | tags=44%, list=35%, signal=[illegible] | [illegible] |
| GOBP_REGULATION_OF_CD40_SIGNALING_PATHWAY | GOBP_REGULATION_OF_CD40_SIGNALING_PATHWAY | 5 | 0.80764375 | 1.35577714 | 0.05665599 | 0.39996035 | 0.36607937 | 1492 | tags=60%, list=10%, signal=[illegible] | [illegible] |
| GOBP_MITOCHONDRIAL_MRNA_CATABOLIC_PROCESS | GOBP_MITOCHONDRIAL_MRNA_CATABOLIC_PROCESS | 5 | 0.806684 | 1.35416604 | 0.05723857 | 0.39996035 | 0.36607937 | 1291 | tags=20%, list=9%, signal=[illegible] | [illegible] |
| GOBP_FIBROBLAST_PROLIFERATION | GOBP_FIBROBLAST_PROLIFERATION | 78 | 0.45703479 | 1.34866152 | 0.05684287 | 0.39996035 | 0.36607937 | 4453 | tags=35%, list=30%, signal=[illegible] | [illegible] |
| GOBP_POSITIVE_REGULATION_OF_DNA_RECOMBINATION | GOBP_POSITIVE_REGULATION_OF_DNA_RECOMBINATION | 69 | 0.46723282 | 1.34748192 | 0.0566542 | 0.39996035 | 0.36607937 | 5619 | tags=51%, list=38%, signal=[illegible] | [illegible] |
| GOBP_KIDNEY_MORPHOGENESIS | GOBP_KIDNEY_MORPHOGENESIS | 76 | 0.45788323 | 1.34662913 | 0.05688569 | 0.39996035 | 0.36607937 | 4019 | tags=41%, list=27%, signal=[illegible] | [illegible] |
| GOBP_SKIN_EPIDERMIS_DEVELOPMENT | GOBP_SKIN_EPIDERMIS_DEVELOPMENT | 94 | 0.43677467 | 1.33095326 | 0.05685373 | 0.39996035 | 0.36607937 | 4644 | tags=54%, list=32%, signal=[illegible] | [illegible] |
| GOBP_LYMPHOCYTE_MEDIATED_IMMUNITY | GOBP_LYMPHOCYTE_MEDIATED_IMMUNITY | 217 | 0.36439538 | 1.25352915 | 0.0572825 | 0.39996035 | 0.36607937 | 4875 | tags=36%, list=33%, signal=[illegible] | [illegible] |
| GOBP_REGULATION_OF_INFLAMMATORY_RESPONSE | GOBP_REGULATION_OF_INFLAMMATORY_RESPONSE | 316 | 0.33825229 | 1.21692904 | 0.05749235 | 0.39996035 | 0.36607937 | 4334 | tags=34%, list=30%, signal=[illegible] | [illegible] |
| GOBP_CELL_ADHESION | GOBP_CELL_ADHESION | 1141 | 0.28153375 | 1.12078784 | 0.05673599 | 0.39996035 | 0.36607937 | 4190 | tags=32%, list=29%, signal=[illegible] | [illegible] |
| GOBP_NEGATIVE_REGULATION_OF_AMIDE_METABOLIC_PROCESS | GOBP_NEGATIVE_REGULATION_OF_AMIDE_METABOLIC_PROCESS | 23 | -0.6479189 | -1.4630074 | 0.0581549 | 0.40184129 | 0.36780097 | 160 | tags=17%, list=1%, signal=[illegible] | [illegible] |
| GOBP_REGULATION_OF_VASOCONSTRICTION | GOBP_REGULATION_OF_VASOCONSTRICTION | 44 | -0.5519423 | -1.4270795 | 0.05806452 | 0.40184129 | 0.36780097 | 124 | tags=7%, list=1%, signal=[illegible] | [illegible] |
| GOBP_NEGATIVE_REGULATION_OF_TOLL_LIKE_RECEPTOR_SIGNALING_ | GOBP_NEGATIVE_REGULATION_OF_TOLL_LIKE_RECEPTOR_SIGNALI | 17 | 0.62926469 | 1.37528414 | 0.05809064 | 0.40184129 | 0.36780097 | 2892 | tags=41%, list=20%, signal=[illegible] | [illegible] |
| GOBP_POSITIVE_REGULATION_OF_VASCULAR_ENDOTHELIAL_CELL_PR | GOBP_POSITIVE_REGULATION_OF_VASCULAR_ENDOTHELIAL_CELL_ | 12 | 0.67444681 | 1.36995045 | 0.05798622 | 0.40184129 | 0.36780097 | 1826 | tags=33%, list=12%, signal=[illegible] | [illegible] |
| GOBP_DETECTION_OF_BIOTIC_STIMULUS | GOBP_DETECTION_OF_BIOTIC_STIMULUS | 25 | 0.57808558 | 1.36393187 | 0.05815465 | 0.40184129 | 0.36780097 | 2628 | tags=40%, list=18%, signal=[illegible] | [illegible] |
| GOBP_NEGATIVE_REGULATION_OF_RESPONSE_TO_BIOTIC_STIMULUS | GOBP_NEGATIVE_REGULATION_OF_RESPONSE_TO_BIOTIC_STIMUL | 104 | 0.42594717 | 1.32134059 | 0.05801668 | 0.40184129 | 0.36780097 | 4849 | tags=36%, list=33%, signal=[illegible] | [illegible] |
| GOBP_SYMBIONT_ENTRY_INTO_HOST | GOBP_SYMBIONT_ENTRY_INTO_HOST | 110 | 0.41921005 | 1.31121009 | 0.05808128 | 0.40184129 | 0.36780097 | 4380 | tags=34%, list=30%, signal=[illegible] | [illegible] |
| GOBP_FC_GAMMA_RECEPTOR_SIGNALING_PATHWAY | GOBP_FC_GAMMA_RECEPTOR_SIGNALING_PATHWAY | 31 | 0.55554336 | 1.36965982 | 0.05833472 | 0.4023435 | 0.36826064 | 4531 | tags=48%, list=31%, signal=[illegible] | [illegible] |
| GOBP_POSITIVE_REGULATION_OF_BMP_SIGNALING_PATHWAY | GOBP_POSITIVE_REGULATION_OF_BMP_SIGNALING_PATHWAY | 32 | 0.55125128 | 1.36506325 | 0.05832219 | 0.4023435 | 0.36826064 | 3945 | tags=34%, list=27%, signal=[illegible] | [illegible] |
| GOBP_CELLULAR_PROCESS_INVOLVED_IN_REPRODUCTION_IN_MULTIC | GOBP_CELLULAR_PROCESS_INVOLVED_IN_REPRODUCTION_IN_MUL | 247 | 0.35471682 | 1.24051312 | 0.05846215 | 0.40248322 | 0.36838853 | 5502 | tags=45%, list=38%, signal=[illegible] | [illegible] |
| GOBP_POSITIVE_REGULATION_OF_CELL_CELL_ADHESION | GOBP_POSITIVE_REGULATION_OF_CELL_CELL_ADHESION | 247 | 0.35468776 | 1.24041148 | 0.05846215 | 0.40248322 | 0.36838853 | 4208 | tags=35%, list=29%, signal=[illegible] | [illegible] |
| GOBP_POSITIVE_REGULATION_OF_ACTIN_FILAMENT_POLYMERIZATIO | GOBP_POSITIVE_REGULATION_OF_ACTIN_FILAMENT_POLYMERIZAT | 45 | -0.5476251 | -1.4209857 | 0.05855749 | 0.4027704 | 0.36865138 | 641 | tags=13%, list=4%, signal=[illegible] | [illegible] |
| GOBP_NEGATIVE_REGULATION_OF_AMYLOID_PRECURSOR_PROTEIN_ | GOBP_NEGATIVE_REGULATION_OF_AMYLOID_PRECURSOR_PROTEI | 17 | -0.6948605 | -1.477872 | 0.05864284 | 0.40298843 | 0.36885094 | 160 | tags=29%, list=1%, signal=[illegible] | [illegible] |
| GOBP_REGULATION_OF_CHONDROCYTE_DIFFERENTIATION | GOBP_REGULATION_OF_CHONDROCYTE_DIFFERENTIATION | 48 | 0.50560536 | 1.35805903 | 0.05885408 | 0.40407036 | 0.36984121 | 2501 | tags=31%, list=17%, signal=[illegible] | [illegible] |
| GOBP_REGULATION_OF_DENDRITIC_CELL_ANTIGEN_PROCESSING_AND | GOBP_REGULATION_OF_DENDRITIC_CELL_ANTIGEN_PROCESSING_ | 10 | -0.7933528 | -1.5225099 | 0.05903946 | 0.40422366 | 0.36998154 | 641 | tags=30%, list=4%, signal=[illegible] | [illegible] |
| GOBP_POSITIVE_REGULATION_OF_THE_FORCE_OF_HEART_CONTRACTI | GOBP_POSITIVE_REGULATION_OF_THE_FORCE_OF_HEART_CONTRA | 5 | -0.8904772 | -1.5121814 | 0.05899235 | 0.40422366 | 0.36998154 | 236 | tags=60%, list=2%, signal=[illegible] | [illegible] |
| GOBP_SNRNA_PROCESSING | GOBP_SNRNA_PROCESSING | 37 | 0.53454628 | 1.3625874 | 0.05909168 | 0.40422366 | 0.36998154 | 5426 | tags=41%, list=37%, signal=[illegible] | [illegible] |
| GOBP_RIBOFLAVIN_METABOLIC_PROCESS | GOBP_RIBOFLAVIN_METABOLIC_PROCESS | 3 | 0.88508474 | 1.35033698 | 0.05903506 | 0.40422366 | 0.36998154 | 1223 | tags=67%, list=8%, signal=[illegible] | [illegible] |
| GOBP_REGULATION_OF_OVARIAN_FOLLICLE_DEVELOPMENT | GOBP_REGULATION_OF_OVARIAN_FOLLICLE_DEVELOPMENT | 4 | 0.84433412 | 1.35598376 | 0.05915947 | 0.40431916 | 0.37006894 | 1854 | tags=50%, list=13%, signal=[illegible] | [illegible] |
| GOBP_RESPONSE_TO_INTERLEUKIN_18 | GOBP_RESPONSE_TO_INTERLEUKIN_18 | 9 | 0.71585112 | 1.35980441 | 0.05937085 | 0.40464913 | 0.37037096 | 2209 | tags=56%, list=15%, signal=[illegible] | [illegible] |
| GOBP_PEPTIDE_ANTIGEN_ASSEMBLY_WITH_MHC_CLASS_I_PROTEIN_C | GOBP_PEPTIDE_ANTIGEN_ASSEMBLY_WITH_MHC_CLASS_I_PROTEI | 5 | 0.80445694 | 1.3504275 | 0.05942324 | 0.40464913 | 0.37037096 | 1423 | tags=60%, list=10%, signal=[illegible] | [illegible] |
| GOBP_PEPTIDE_ANTIGEN_ASSEMBLY_WITH_MHC_PROTEIN_COMPLEX | GOBP_PEPTIDE_ANTIGEN_ASSEMBLY_WITH_MHC_PROTEIN_COMPL | 5 | 0.80445694 | 1.3504275 | 0.05942324 | 0.40464913 | 0.37037096 | 1423 | tags=60%, list=10%, signal=[illegible] | [illegible] |
| GOBP_CHIASMA_ASSEMBLY | GOBP_CHIASMA_ASSEMBLY | 5 | 0.80427679 | 1.35012509 | 0.05942324 | 0.40464913 | 0.37037096 | 2108 | tags=60%, list=14%, signal=[illegible] | [illegible] |
| GOBP_LENS_MORPHOGENESIS_IN_CAMERA_TYPE_EYE | GOBP_LENS_MORPHOGENESIS_IN_CAMERA_TYPE_EYE | 21 | 0.59866886 | 1.3674546 | 0.05953326 | 0.40503108 | 0.37072055 | 2723 | tags=38%, list=19%, signal=[illegible] | [illegible] |
| GOBP_POSITIVE_REGULATION_OF_RELEASE_OF_SEQUESTERED_CALCIU | GOBP_POSITIVE_REGULATION_OF_RELEASE_OF_SEQUESTERED_CAL | 23 | -0.6450922 | -1.4566248 | 0.05974095 | 0.40534259 | 0.37100568 | 333 | tags=17%, list=2%, signal=[illegible] | [illegible] |
| GOBP_NOREPINEPHRINE_TRANSPORT | GOBP_NOREPINEPHRINE_TRANSPORT | 13 | 0.66096536 | 1.36473247 | 0.05970602 | 0.40534259 | 0.37100568 | 3090 | tags=54%, list=21%, signal=[illegible] | [illegible] |
