## Supplementary material for "Integrated transcriptomic analysis identifies Paternally Expressed Gene 10 as a novel candidate biomarker associated with aggressive features in liposarcoma": Table S3

| gene | ensembl_id | logFC | logCPM | F | PValue | FDR | significant |
|---|---|---|---|---|---|---|---|
| SBSPON | ENSG00000164764 | 7.454463927 | 5.901188125 | 40.23637714 | 1.99E-07 | 0.00106618 | Yes |
| IL20RB | ENSG00000174564 | -5.628688288 | 4.88025579 | 38.21876517 | 2.10E-07 | 0.00106618 | Yes |
| FNDC4 | ENSG00000115226 | 3.709526967 | 4.626379525 | 39.50194554 | 2.19E-07 | 0.00106618 | Yes |
| CA4 | ENSG00000167434 | 4.949061669 | 5.397509984 | 36.36261745 | 3.93E-07 | 0.001135649 | Yes |
| IGHM | ENSG00000211899 | 8.600117501 | 7.996201579 | 34.31598779 | 4.70E-07 | 0.001135649 | Yes |
| RBP4 | ENSG00000138207 | 5.761796616 | 6.576396307 | 36.62382883 | 4.83E-07 | 0.001135649 | Yes |
| CNTFR | ENSG00000122756 | 4.345008408 | 4.860521201 | 35.22978223 | 5.43E-07 | 0.001135649 | Yes |
| PLIN1 | ENSG00000166819 | 6.009469899 | 4.516585932 | 35.16710472 | 7.02E-07 | 0.001285018 | Yes |
| SH3PXD2A | ENSG00000107957 | 2.480316862 | 8.744017484 | 37.68823061 | 9.36E-07 | 0.001454238 | Yes |
| DLK1 | ENSG00000185559 | 7.537945765 | 8.345192826 | 33.8546951 | 1.02E-06 | 0.001454238 | Yes |
| PI3 | ENSG00000124102 | -6.229268951 | 6.203438902 | 31.60128888 | 1.09E-06 | 0.001454238 | Yes |
| DHRS9 | ENSG00000073737 | -8.16435414 | 6.753864368 | 33.41062457 | 1.39E-06 | 0.001571359 | Yes |
| PEG10 | ENSG00000242265 | 3.275128884 | 7.779324212 | 35.54479339 | 1.51E-06 | 0.001571359 | Yes |
| IGHG3 | ENSG00000211897 | 7.125011982 | 6.34212425 | 30.35341781 | 1.59E-06 | 0.001571359 | Yes |
| SLPI | ENSG00000124107 | -7.537730194 | 8.999899199 | 32.20789197 | 1.61E-06 | 0.001571359 | Yes |
| KRT19 | ENSG00000171345 | -8.661586591 | 9.197852744 | 30.06047951 | 1.96E-06 | 0.001794029 | Yes |
| KRT17 | ENSG00000128422 | -7.630424549 | 6.771423928 | 30.16991459 | 2.13E-06 | 0.001832357 | Yes |
| SLC29A4 | ENSG00000164638 | 3.91376886 | 5.015055898 | 30.52679351 | 2.74E-06 | 0.002087397 | Yes |
| CIDEC | ENSG00000187288 | 6.99690222 | 4.318425413 | 35.03831519 | 2.94E-06 | 0.002087397 | Yes |
| FHAD1 | ENSG00000142621 | -4.724605029 | 5.175292662 | 28.56366916 | 2.97E-06 | 0.002087397 | Yes |
| ADIPOQ | ENSG00000181092 | 11.39303397 | 6.745387447 | 39.69912153 | 3.04E-06 | 0.002087397 | Yes |
| IGHA1 | ENSG00000211895 | 6.805278344 | 4.566840232 | 32.81737191 | 3.17E-06 | 0.002087397 | Yes |
| DSP | ENSG00000096696 | -7.481876216 | 6.655838039 | 28.71379659 | 3.28E-06 | 0.002087397 | Yes |
| CXCL1 | ENSG00000163739 | -4.698331839 | 4.455109808 | 27.92410014 | 3.50E-06 | 0.002132239 | Yes |
| PERP | ENSG00000112378 | -6.24578792 | 6.833745403 | 27.91060718 | 3.86E-06 | 0.002261972 | Yes |
| RET | ENSG00000165731 | 4.686005083 | 5.105716755 | 27.44113915 | 5.07E-06 | 0.002670186 | Yes |
| PLPP2 | ENSG00000141934 | -5.278717517 | 4.976789457 | 26.65024839 | 5.30E-06 | 0.002670186 | Yes |
| PLEKHG4B | ENSG00000153404 | 4.731972791 | 3.919069104 | 26.49849225 | 5.46E-06 | 0.002670186 | Yes |
| DEFB1 | ENSG00000164825 | -5.23618512 | 6.135255251 | 27.04983681 | 5.58E-06 | 0.002670186 | Yes |
| AQP7 | ENSG00000165269 | 4.786271493 | 6.585622156 | 29.03225802 | 5.65E-06 | 0.002670186 | Yes |
| LAMB3 | ENSG00000196878 | -5.009597153 | 4.341455907 | 26.82194882 | 5.66E-06 | 0.002670186 | Yes |
| TACSTD2 | ENSG00000184292 | -8.803611848 | 6.961765986 | 29.70162631 | 5.94E-06 | 0.002718988 | Yes |
| CPLX1 | ENSG00000168993 | 3.978156795 | 3.428713765 | 25.5356221 | 6.90E-06 | 0.002775342 | Yes |
| HOXC12 | ENSG00000123407 | 7.612312934 | 3.835755519 | 33.48486711 | 7.02E-06 | 0.002775342 | Yes |
| ANGPT1 | ENSG00000154188 | 3.93597601 | 5.25794764 | 27.5993131 | 7.04E-06 | 0.002775342 | Yes |
| CFAP53 | ENSG00000172361 | -6.214424508 | 5.260858908 | 26.45262981 | 7.21E-06 | 0.002775342 | Yes |
| HLA.DRB5 | NA | -5.430785567 | 5.236404816 | 25.66752605 | 7.22E-06 | 0.002775342 | Yes |
| THRSP | ENSG00000151365 | 5.543253028 | 4.563536075 | 25.65541245 | 7.73E-06 | 0.002775342 | Yes |
| CLDN5 | ENSG00000184113 | 3.448779684 | 6.012362772 | 28.37701371 | 7.83E-06 | 0.002775342 | Yes |
| C7 | ENSG00000112936 | 5.64183161 | 4.187971473 | 26.60888772 | 7.86E-06 | 0.002775342 | Yes |
| ELF3 | ENSG00000163435 | -6.383525324 | 6.483145455 | 25.34232243 | 8.03E-06 | 0.002775342 | Yes |
| IGHG1 | ENSG00000211896 | 5.506157582 | 4.067477342 | 26.47551079 | 8.06E-06 | 0.002775342 | Yes |
| PLIN5 | ENSG00000214456 | 4.555259084 | 5.955736916 | 27.2087003 | 8.28E-06 | 0.002775342 | Yes |
| FZD4 | ENSG00000174804 | 3.337473008 | 7.756994745 | 28.69012221 | 8.34E-06 | 0.002775342 | Yes |
| PNMA3 | ENSG00000183837 | 4.430866344 | 3.957784157 | 24.82131535 | 8.96E-06 | 0.002910754 | Yes |
| WDR54 | ENSG00000005448 | -4.003813885 | 5.064843016 | 25.54831483 | 9.29E-06 | 0.002910754 | Yes |
| MRAP | ENSG00000170262 | 8.644874875 | 4.802423366 | 32.6958608 | 9.35E-06 | 0.002910754 | Yes |
| MDFI | ENSG00000112559 | 3.232089041 | 4.737806894 | 26.39610642 | 1.03E-05 | 0.003063734 | Yes |
| VSTM2L | ENSG00000132821 | -6.414908417 | 4.3351662 | 28.75120248 | 1.03E-05 | 0.003063734 | Yes |
| KLF15 | ENSG00000163884 | 4.429324192 | 3.152116539 | 25.35073542 | 1.05E-05 | 0.003063734 | Yes |
| HOXC11 | ENSG00000123388 | 5.850991148 | 3.540219523 | 28.2013489 | 1.09E-05 | 0.003063734 | Yes |
| HSD11B2 | ENSG00000176387 | 3.058530723 | 6.99332174 | 27.61885005 | 1.09E-05 | 0.003063734 | Yes |
| IGLC2 | ENSG00000211677 | 6.623032762 | 4.976835634 | 25.92116221 | 1.11E-05 | 0.003063734 | Yes |
| SIX3 | ENSG00000138083 | -5.610556415 | 5.039474436 | 24.53822775 | 1.13E-05 | 0.003063734 | Yes |
| DGAT2 | ENSG00000062282 | 4.539739171 | 6.117013286 | 26.22955159 | 1.17E-05 | 0.003109543 | Yes |
| CLDN7 | ENSG00000181885 | -4.374839985 | 5.679655951 | 25.00866303 | 1.22E-05 | 0.003200289 | Yes |
| PDZD4 | ENSG00000067840 | 3.635438161 | 2.985133855 | 23.72251434 | 1.26E-05 | 0.003229355 | Yes |
| WFDC2 | ENSG00000101443 | -6.391659578 | 7.797417547 | 25.0641979 | 1.28E-05 | 0.003229355 | Yes |
| NPY1R | ENSG00000164128 | 6.721361151 | 3.800078665 | 28.98172538 | 1.34E-05 | 0.00332425 | Yes |
| COL6A6 | ENSG00000206384 | 4.936807965 | 4.641874725 | 23.56782817 | 1.43E-05 | 0.003480886 | Yes |
| RXRG | ENSG00000143171 | 4.13012054 | 3.124738493 | 23.7209148 | 1.46E-05 | 0.003508731 | Yes |
| VMO1 | ENSG00000182853 | -5.165065992 | 6.248585824 | 24.09178108 | 1.54E-05 | 0.003607616 | Yes |
| LRRIQ1 | ENSG00000133640 | -5.495609595 | 5.977157402 | 23.53512241 | 1.55E-05 | 0.003607616 | Yes |
| ZBTB16 | ENSG00000109906 | 3.522220839 | 5.023784286 | 24.80244946 | 1.68E-05 | 0.003836148 | Yes |
| DNALI1 | ENSG00000163879 | -4.562208191 | 4.760812162 | 23.10806331 | 1.71E-05 | 0.00385074 | Yes |
| ADAMTS19 | ENSG00000145808 | 4.156380269 | 4.408425893 | 23.24253098 | 1.79E-05 | 0.003944449 | Yes |
| GPD1 | ENSG00000167588 | 4.49044042 | 6.530394432 | 25.13439749 | 1.81E-05 | 0.003944449 | Yes |
| SLC17A8 | ENSG00000179520 | 6.624758991 | 3.380956042 | 27.62617706 | 2.01E-05 | 0.004322826 | Yes |
| CLDN4 | ENSG00000189143 | -4.887869178 | 5.630602199 | 22.7032736 | 2.19E-05 | 0.004624714 | Yes |
| PNPLA2 | ENSG00000177666 | 2.288833582 | 7.429160185 | 25.12946471 | 2.23E-05 | 0.004624714 | Yes |
| NRG2 | ENSG00000158458 | 3.324543227 | 3.902351629 | 22.72356397 | 2.27E-05 | 0.004624714 | Yes |
| RAPGEF3 | ENSG00000079337 | 3.251883078 | 5.060475616 | 24.04572446 | 2.27E-05 | 0.004624714 | Yes |
| TMEM132C | ENSG00000181234 | 4.944452839 | 4.063585276 | 22.32476636 | 2.33E-05 | 0.004666782 | Yes |
| SLC44A4 | ENSG00000204385 | -5.196496022 | 4.933144475 | 22.06399824 | 2.48E-05 | 0.004902533 | Yes |
| ADIRF | ENSG00000148671 | 2.958804564 | 6.825954172 | 24.58891459 | 2.53E-05 | 0.004938397 | Yes |
| TCF15 | ENSG00000125878 | 3.296818931 | 4.617458891 | 23.02504806 | 2.73E-05 | 0.005196473 | Yes |
| PARM1 | ENSG00000169116 | 2.909552976 | 4.974621437 | 23.61563993 | 2.73E-05 | 0.005196473 | Yes |
| CDS1 | ENSG00000163624 | -4.561150192 | 5.658113937 | 22.26802362 | 2.80E-05 | 0.005196473 | Yes |
| FABP4 | ENSG00000170323 | 4.81376598 | 10.55770962 | 24.31705508 | 2.81E-05 | 0.005196473 | Yes |
| GPT | ENSG00000167701 | 5.513868757 | 3.557114315 | 23.73534575 | 2.84E-05 | 0.005196473 | Yes |
| LIPE | ENSG00000079435 | 3.889514582 | 6.925718355 | 23.90624553 | 2.95E-05 | 0.005210748 | Yes |
| HOXA4 | ENSG00000197576 | 3.461921886 | 4.722897246 | 22.71090076 | 2.97E-05 | 0.005210748 | Yes |
| IL1RN | ENSG00000136689 | -5.157282112 | 6.688644407 | 22.52819417 | 2.97E-05 | 0.005210748 | Yes |
| CD300LG | ENSG00000161649 | 3.437595775 | 5.548158282 | 23.41900847 | 2.99E-05 | 0.005210748 | Yes |
| GDPD1 | ENSG00000153982 | -3.259889384 | 2.912321878 | 21.46438813 | 3.14E-05 | 0.005408777 | Yes |
| P3H2 | ENSG00000090530 | 2.810670918 | 6.021254594 | 23.57357056 | 3.26E-05 | 0.005552306 | Yes |
| MMP13 | ENSG00000137745 | -5.06085386 | 4.419113823 | 21.39848366 | 3.40E-05 | 0.005713613 | Yes |
| SGK2 | ENSG00000101049 | 3.996356066 | 4.108279691 | 21.1349976 | 3.49E-05 | 0.005811256 | Yes |
| AGT | ENSG00000135744 | 3.77523877 | 6.513300478 | 23.18439715 | 3.54E-05 | 0.005828867 | Yes |
| SAA2 | ENSG00000134339 | -8.762308787 | 6.383884267 | 24.51682672 | 3.76E-05 | 0.006043671 | Yes |
| OPLAH | ENSG00000178814 | 2.945560006 | 4.345764616 | 21.98815714 | 3.77E-05 | 0.006043671 | Yes |
| BTNL9 | ENSG00000165810 | 3.51956749 | 7.527368883 | 23.21209811 | 3.80E-05 | 0.006043671 | Yes |
| ADAMTS9 | ENSG00000163638 | 2.915755475 | 6.213579254 | 23.02501158 | 3.87E-05 | 0.006087512 | Yes |
| LAMC2 | ENSG00000058085 | -4.150474442 | 4.944927023 | 20.97665891 | 3.95E-05 | 0.006119195 | Yes |
| ALCAM | ENSG00000170017 | -3.985757296 | 6.902621476 | 22.57631787 | 4.01E-05 | 0.006119195 | Yes |
| MYH11 | ENSG00000133392 | 4.264720249 | 5.152106065 | 21.50103067 | 4.01E-05 | 0.006119195 | Yes |
| TMEM240 | ENSG00000205090 | -3.922637698 | 3.437012917 | 20.57085713 | 4.47E-05 | 0.006737562 | Yes |
| SMIM10L2A | ENSG00000178947 | 3.237030594 | 3.143420366 | 20.01742362 | 4.66E-05 | 0.006932622 | Yes |
| POU3F3 | ENSG00000198914 | 4.813228354 | 3.557361141 | 20.72567791 | 4.73E-05 | 0.006932622 | Yes |
| MBOAT7 | ENSG00000125505 | -1.607795019 | 6.222335746 | 22.41743877 | 4.74E-05 | 0.006932622 | Yes |
| EGR3 | ENSG00000179388 | 3.725556905 | 3.802147567 | 20.13537418 | 4.85E-05 | 0.006972564 | Yes |
| PNMA6F | ENSG00000225110 | 6.16605946 | 4.000323283 | 22.47692393 | 4.86E-05 | 0.006972564 | Yes |
| LINC00348 | ENSG00000226846 | 8.593901744 | 4.028789956 | 25.39203887 | 4.98E-05 | 0.00707964 | Yes |
| PON3 | ENSG00000105852 | 3.493104051 | 4.517705542 | 20.85130604 | 5.07E-05 | 0.007088143 | Yes |
| CFAP43 | ENSG00000197748 | -5.184051336 | 4.655252309 | 20.16219757 | 5.08E-05 | 0.007088143 | Yes |
| NDRG4 | ENSG00000103034 | 3.279714671 | 4.532764619 | 20.91412396 | 5.29E-05 | 0.007298209 | Yes |
| ZBTB7C | ENSG00000184828 | -3.432779191 | 4.452244367 | 20.16286829 | 5.45E-05 | 0.007403745 | Yes |
| DSC3 | ENSG00000134762 | -4.27310229 | 3.763768878 | 19.97961075 | 5.46E-05 | 0.007403745 | Yes |
| FOSB | ENSG00000125740 | 4.785164649 | 8.891978713 | 21.88733649 | 5.66E-05 | 0.007596285 | Yes |
| RAB27B | ENSG00000041353 | -8.297533332 | 3.645759141 | 25.41030762 | 5.98E-05 | 0.007955447 | Yes |
| TBC1D2 | ENSG00000095383 | -3.737758553 | 3.936810486 | 19.5163587 | 6.05E-05 | 0.007972234 | Yes |
| ST6GALNAC2 | ENSG00000070731 | -4.24987488 | 3.92188921 | 19.44516752 | 6.39E-05 | 0.008356339 | Yes |
| CDKN1C | ENSG00000129757 | 1.988388778 | 6.254839912 | 21.41528554 | 6.50E-05 | 0.008424036 | Yes |
| CAPS | ENSG00000105519 | -3.743225398 | 7.065780023 | 20.97324636 | 6.90E-05 | 0.008853303 | Yes |
| CFAP91 | ENSG00000183833 | -5.165085146 | 4.344096987 | 19.43883364 | 7.18E-05 | 0.00896231 | Yes |
| TEX22 | ENSG00000226174 | 6.682831232 | 2.400321403 | 22.07350538 | 7.21E-05 | 0.00896231 | Yes |
| LRRC10B | ENSG00000204950 | -4.078350685 | 4.457582286 | 19.09560374 | 7.21E-05 | 0.00896231 | Yes |
| S100A9 | ENSG00000163220 | -5.149781822 | 8.061138388 | 20.70262273 | 7.23E-05 | 0.00896231 | Yes |
| GABRB1 | ENSG00000163288 | 4.75820841 | 3.362704035 | 19.62268896 | 7.71E-05 | 0.00940716 | Yes |
| PCK1 | ENSG00000124253 | 8.342633935 | 3.787172509 | 23.45817602 | 7.74E-05 | 0.00940716 | Yes |
| DOCK3 | ENSG00000088538 | 6.574299841 | 3.920502764 | 21.76292276 | 7.78E-05 | 0.00940716 | Yes |
| ECHDC3 | ENSG00000134463 | 4.634459043 | 3.254064103 | 19.3644644 | 7.97E-05 | 0.009467484 | Yes |
| KLB | ENSG00000134962 | 4.502630339 | 3.78326098 | 18.6990182 | 8.08E-05 | 0.009467484 | Yes |
| SCNN1A | ENSG00000111319 | -4.791124506 | 4.465613834 | 18.75065672 | 8.22E-05 | 0.009467484 | Yes |
| LINC01305 | ENSG00000231453 | 4.257629492 | 3.414101054 | 18.64155786 | 8.31E-05 | 0.009467484 | Yes |
| RYR2 | ENSG00000198626 | 5.081058072 | 2.870106386 | 20.55244436 | 8.31E-05 | 0.009467484 | Yes |
| CBLN1 | ENSG00000102924 | 4.308389197 | 2.739404578 | 19.56337758 | 8.32E-05 | 0.009467484 | Yes |
| FAM13A | ENSG00000138640 | 2.514731075 | 7.417296689 | 20.65126376 | 8.33E-05 | 0.009467484 | Yes |
| S100B | ENSG00000160307 | 5.752022224 | 2.652187477 | 21.3758923 | 8.34E-05 | 0.009467484 | Yes |
| TFCP2L1 | ENSG00000115112 | -6.47730457 | 3.649832659 | 22.42249837 | 8.55E-05 | 0.009621349 | Yes |
| LGALS12 | ENSG00000133317 | 5.677543432 | 3.149043893 | 20.97914356 | 8.69E-05 | 0.009705052 | Yes |
| SOX2 | ENSG00000181449 | -5.004694167 | 5.310058457 | 18.55916674 | 8.81E-05 | 0.009761066 | Yes |
| ENSG0000024344 | ENSG00000243444 | 3.290330397 | 3.985251625 | 18.88775445 | 8.87E-05 | 0.009761066 | Yes |
| DNAI4 | ENSG00000152763 | -3.372915378 | 4.606197329 | 18.9125076 | 8.95E-05 | 0.009768391 | Yes |
| SPAG8 | ENSG00000137098 | -4.203574047 | 4.176919842 | 18.43174216 | 9.06E-05 | 0.009768391 | Yes |
| SLC2A1 | ENSG00000117394 | -3.64502038 | 5.834828027 | 19.58001623 | 9.08E-05 | 0.009768391 | Yes |
| AR | ENSG00000169083 | 3.832177318 | 5.01801776 | 19.31327122 | 9.17E-05 | 0.009802175 | Yes |
| S100A2 | ENSG00000196754 | -6.133961612 | 8.780108464 | 19.80302744 | 9.33E-05 | 0.009861632 | Yes |
| SFN | ENSG00000175793 | -12.32851346 | 7.53478032 | 23.70587822 | 9.37E-05 | 0.009861632 | Yes |
| IL11RA | ENSG00000137070 | 2.199835613 | 8.071188381 | 20.23261588 | 9.53E-05 | 0.00993558 | Yes |
| RASEF | ENSG00000165105 | -3.821388845 | 3.331446302 | 18.442842 | 9.57E-05 | 0.00993558 | Yes |
| PLIN4 | ENSG00000167676 | 3.331749355 | 6.616373475 | 19.93894565 | 0.00010001 | 0.010308824 | Yes |
| ZSCAN1 | ENSG00000152467 | 3.255380099 | 3.13113919 | 17.94213067 | 0.000102244 | 0.010465326 | Yes |
| CAMK2B | ENSG00000058404 | 2.8693139 | 4.766460272 | 19.30056754 | 0.000103749 | 0.010540019 | Yes |
| SPATA22 | ENSG00000141255 | 6.915088611 | 2.530437013 | 21.07108678 | 0.000104414 | 0.010540019 | Yes |
| ANHX | ENSG00000227059 | 7.747942706 | 3.293769107 | 21.80946626 | 0.000107006 | 0.010727745 | Yes |
| SCUBE3 | ENSG00000146197 | 4.192944811 | 3.111287243 | 18.00065446 | 0.000112453 | 0.011197131 | Yes |
| PTX3 | ENSG00000163661 | 3.351220025 | 7.75805319 | 19.60952768 | 0.000114897 | 0.011307161 | Yes |
| ENSG0000027421 | ENSG00000274213 | -3.503753556 | 3.292262674 | 17.8280735 | 0.000115103 | 0.011307161 | Yes |
| CLDN1 | ENSG00000163347 | -4.299065744 | 3.529524124 | 18.03438496 | 0.000119203 | 0.01161591 | Yes |
| MYOC | ENSG00000034971 | 5.022149077 | 4.851938537 | 17.68509151 | 0.000119833 | 0.01161591 | Yes |
| ELAPOR2 | ENSG00000164659 | 2.992404109 | 4.345718817 | 18.46061877 | 0.000122976 | 0.011842132 | Yes |
| PIERCE1 | ENSG00000160345 | -4.717691807 | 5.647348121 | 17.83706031 | 0.000127296 | 0.01217801 | Yes |
| PLA2R1 | ENSG00000153246 | 3.812905488 | 5.224276406 | 18.35964091 | 0.000133518 | 0.012669301 | Yes |
| ENSG0000025062 | ENSG00000250620 | 5.380412417 | 3.486470282 | 18.74064877 | 0.000134163 | 0.012669301 | Yes |
| KRT8 | ENSG00000170421 | -4.707052953 | 5.704564578 | 17.63826292 | 0.000138588 | 0.013003296 | Yes |
| NT5M | ENSG00000205309 | 3.263111225 | 3.32351459 | 17.14659811 | 0.000140883 | 0.013134424 | Yes |
| DNER | ENSG00000187957 | -5.182194907 | 3.395843255 | 18.94303051 | 0.00014279 | 0.013224392 | Yes |
| STARD8 | ENSG00000130052 | 2.751349936 | 4.232755027 | 18.02485016 | 0.000144576 | 0.013224392 | Yes |
| SLC25A18 | ENSG00000182902 | 7.289899046 | 2.891226206 | 20.31229001 | 0.000144873 | 0.013224392 | Yes |
| CPXM1 | ENSG00000088882 | 3.470394805 | 5.351871165 | 18.35241035 | 0.000145462 | 0.013224392 | Yes |
| HOXA3 | ENSG00000105997 | 3.085198415 | 5.559268565 | 18.5122691 | 0.00014805 | 0.013376628 | Yes |
| NWD1 | ENSG00000188039 | -4.639518556 | 4.502922131 | 17.11579966 | 0.000149718 | 0.013422967 | Yes |
| CLXN | ENSG00000034239 | -4.482202377 | 4.376677994 | 17.10020595 | 0.000150397 | 0.013422967 | Yes |
| ENSG0000026653 | ENSG00000266538 | 2.562090197 | 5.35469695 | 18.46389856 | 0.000153564 | 0.01362249 | Yes |
| RIPK4 | ENSG00000183421 | -5.726598306 | 3.986833929 | 18.51618514 | 0.000155493 | 0.013710511 | Yes |
| CDH6 | ENSG00000113361 | 3.59832659 | 4.713003889 | 17.67958309 | 0.000156769 | 0.013740269 | Yes |
| CCDC146 | ENSG00000135205 | -3.913235637 | 5.872379575 | 17.77474763 | 0.000161557 | 0.01407569 | Yes |
| INHBA | ENSG00000122641 | -3.406194564 | 4.147775473 | 16.99008861 | 0.00016352 | 0.014135936 | Yes |
| CSTA | ENSG00000121552 | -3.823991806 | 5.24841672 | 17.40038397 | 0.00016418 | 0.014135936 | Yes |
| HRK | ENSG00000135116 | -3.578450837 | 4.135782257 | 16.87525887 | 0.00016716 | 0.014242483 | Yes |
| RAI2 | ENSG00000131831 | 4.246423409 | 4.289005646 | 16.88992734 | 0.000167364 | 0.014242483 | Yes |
| MMP19 | ENSG00000123342 | 2.910147391 | 4.689963288 | 17.6708833 | 0.000176442 | 0.014928185 | Yes |
| FAM133A | ENSG00000179083 | -3.876597021 | 3.545769885 | 16.65710842 | 0.000182513 | 0.015353119 | Yes |
| TMEM116 | ENSG00000198270 | -3.265948071 | 3.539070849 | 16.511174 | 0.000186173 | 0.015526375 | Yes |
| LRRC73 | ENSG00000204052 | -4.173182874 | 3.257008151 | 16.91867293 | 0.000189729 | 0.015526375 | Yes |
| SCARNA17 | ENSG00000267322 | 2.492982028 | 4.447196894 | 17.47658248 | 0.000190185 | 0.015526375 | Yes |
| PDE3B | ENSG00000152270 | 3.610961716 | 4.748412035 | 17.13821453 | 0.00019147 | 0.015526375 | Yes |
| OPRK1 | ENSG00000082556 | 7.823396761 | 3.372687862 | 19.72623736 | 0.000191555 | 0.015526375 | Yes |
| HAAO | ENSG00000162882 | 2.66166371 | 3.562443947 | 16.77288923 | 0.000191914 | 0.015526375 | Yes |
| PLD5 | ENSG00000180287 | 6.655483022 | 2.41166952 | 18.68836767 | 0.000193413 | 0.015526375 | Yes |
| LEPR | ENSG00000116678 | 2.75325746 | 6.649407732 | 17.94720835 | 0.000194292 | 0.015526375 | Yes |
| ENSG0000018508 | ENSG00000185087 | 6.147578727 | 2.560737941 | 18.71077932 | 0.000194788 | 0.015526375 | Yes |
| TTC9 | ENSG00000133985 | -4.140613568 | 4.342990148 | 16.42060741 | 0.00019518 | 0.015526375 | Yes |
| ADORA1 | ENSG00000163485 | 7.006000149 | 2.646224091 | 18.95266823 | 0.000198816 | 0.015730145 | Yes |
| ADAMTS18 | ENSG00000140873 | 4.187093854 | 2.528239058 | 17.00257642 | 0.000200153 | 0.015750767 | Yes |
| MT1A | ENSG00000205362 | 2.6985927 | 3.853909175 | 16.83720388 | 0.000203244 | 0.015881877 | Yes |
| ITGA10 | ENSG00000143127 | -4.383839618 | 4.578090427 | 16.31021312 | 0.000203989 | 0.015881877 | Yes |
| TRARG1 | ENSG00000184811 | 6.1019728 | 2.609664509 | 18.50056991 | 0.0002053 | 0.015899353 | Yes |
| BBOF1 | ENSG00000119636 | -3.750167787 | 5.077090245 | 16.68734014 | 0.000207667 | 0.015997971 | Yes |
| ALDH3A1 | ENSG00000108602 | -9.369716335 | 5.499322409 | 20.10421302 | 0.000208936 | 0.016011466 | Yes |
| MICAL2 | ENSG00000133816 | -3.466692678 | 4.534440358 | 16.44875169 | 0.000214012 | 0.016274889 | Yes |
| S100A8 | ENSG00000143546 | -4.084198694 | 5.654789867 | 16.76383959 | 0.000214597 | 0.016274889 | Yes |
| DGAT1 | ENSG00000185000 | 2.183480813 | 5.850263732 | 17.474664 | 0.000224348 | 0.016872289 | Yes |
| ENSG0000023373 | ENSG00000233730 | -4.766120567 | 4.459442214 | 16.0714978 | 0.000227276 | 0.016872289 | Yes |
| HS6ST3 | ENSG00000185352 | 6.59325859 | 2.391540825 | 18.08823267 | 0.000228266 | 0.016872289 | Yes |
| RIN1 | ENSG00000174791 | -2.689152158 | 4.060590741 | 16.33846624 | 0.000230171 | 0.016872289 | Yes |
| DNAH12 | ENSG00000174844 | -4.51629344 | 4.175733052 | 16.04187367 | 0.000230543 | 0.016872289 | Yes |
| COL25A1 | ENSG00000188517 | 4.468417593 | 2.561146652 | 16.96033176 | 0.000231071 | 0.016872289 | Yes |
| ST14 | ENSG00000149418 | -3.694801078 | 4.739961122 | 16.24370545 | 0.000231673 | 0.016872289 | Yes |
| CYP4B1 | ENSG00000142973 | -4.695054628 | 6.184564052 | 16.49391818 | 0.000233976 | 0.016872289 | Yes |
| RGS11 | ENSG00000076344 | 3.691696495 | 3.223796797 | 15.80411006 | 0.000235369 | 0.016872289 | Yes |
| NTNG1 | ENSG00000162631 | 3.429203028 | 4.287814205 | 16.37322106 | 0.000235521 | 0.016872289 | Yes |
| TUFT1 | ENSG00000143367 | -2.623840559 | 5.586781713 | 17.0603789 | 0.000236261 | 0.016872289 | Yes |
| FZD8 | ENSG00000177283 | 2.34671716 | 3.714382099 | 16.4643205 | 0.00023762 | 0.016872289 | Yes |
| MEST | ENSG00000106484 | 2.271820085 | 6.915751339 | 17.36210315 | 0.000238678 | 0.016872289 | Yes |
| SPOCD1 | ENSG00000134668 | -4.243955382 | 5.713504701 | 16.41112055 | 0.000239231 | 0.016872289 | Yes |
| MEOX1 | ENSG00000005102 | 3.663019964 | 3.212882979 | 15.75093983 | 0.000239765 | 0.016872289 | Yes |
| PPARG | ENSG00000132170 | 2.87083211 | 4.865269607 | 16.79834604 | 0.000247433 | 0.017289082 | Yes |
| CCDC190 | ENSG00000185860 | -5.692803262 | 4.23029635 | 16.74853324 | 0.000248351 | 0.017289082 | Yes |
| PHLDA2 | ENSG00000181649 | -3.515660196 | 5.306410669 | 16.44398807 | 0.000249231 | 0.017289082 | Yes |
| NCALD | ENSG00000104490 | 2.378283247 | 5.183920036 | 16.98380545 | 0.000251872 | 0.017389869 | Yes |
| C10orf143 | ENSG00000237489 | 5.133833836 | 2.445472866 | 17.27746502 | 0.000254837 | 0.017511936 | Yes |
| BMPER | ENSG00000164619 | 3.312198168 | 4.97502152 | 16.5119271 | 0.000264345 | 0.018080474 | Yes |
| RFLNA | ENSG00000178882 | 6.960177795 | 2.61500388 | 17.91506966 | 0.000268485 | 0.018241075 | Yes |
| ITGB4 | ENSG00000132470 | -3.884339164 | 5.063675046 | 15.90671602 | 0.000269825 | 0.018241075 | Yes |
| C19orf38 | ENSG00000214212 | 3.172565075 | 3.996713175 | 15.94959744 | 0.000270479 | 0.018241075 | Yes |
| CLIC6 | ENSG00000159212 | -3.865117649 | 4.085480339 | 15.56764623 | 0.000271848 | 0.018241075 | Yes |
| COL18A1 | ENSG00000182871 | 1.490731743 | 7.551241599 | 16.96859972 | 0.000273866 | 0.018241075 | Yes |
| RRAD | ENSG00000166592 | -3.386354428 | 4.939532376 | 16.04484545 | 0.000274171 | 0.018241075 | Yes |
| UAP1 | ENSG00000117143 | -2.394313819 | 6.300193935 | 16.81498648 | 0.000276189 | 0.018292218 | Yes |
| ENSG0000025614 | ENSG00000256146 | 6.272643113 | 3.415231191 | 17.5884449 | 0.00028288 | 0.018650998 | Yes |
| GRID2 | ENSG00000152208 | 4.320847357 | 2.833792409 | 15.90357032 | 0.000292051 | 0.019169292 | Yes |
| PLAAT3 | ENSG00000176485 | 1.620099182 | 7.122508518 | 16.71367124 | 0.000298079 | 0.019278364 | Yes |
| SEPTIN6 | ENSG00000125354 | 1.662719622 | 7.392300478 | 16.71514031 | 0.000298138 | 0.019278364 | Yes |
| MNS1 | ENSG00000138587 | -2.976182468 | 6.094349074 | 16.42993429 | 0.000300402 | 0.019278364 | Yes |
| ZG16B | ENSG00000162078 | -3.899353753 | 4.463096876 | 15.37867696 | 0.000301406 | 0.019278364 | Yes |
| DRC3 | ENSG00000171962 | -3.679440673 | 3.787191429 | 15.27860693 | 0.000303138 | 0.019278364 | Yes |
| NBL1 | ENSG00000158747 | -2.095937878 | 7.240941943 | 16.63941168 | 0.000303361 | 0.019278364 | Yes |
| RFX8 | ENSG00000196460 | -4.055112253 | 3.5138172 | 15.42906518 | 0.000303632 | 0.019278364 | Yes |
| RAB11FIP1 | ENSG00000156675 | -3.807945467 | 6.883322671 | 16.40314198 | 0.00030425 | 0.019278364 | Yes |
| NEFL | ENSG00000277586 | 3.960880085 | 4.411238286 | 15.48706211 | 0.000308255 | 0.019437563 | Yes |
| TUBA4A | ENSG00000127824 | -2.892972795 | 4.357387925 | 15.63164412 | 0.000309464 | 0.019437563 | Yes |
| CDKN2B | ENSG00000147883 | -3.313920255 | 5.214752398 | 15.88864026 | 0.000310746 | 0.019437563 | Yes |
| TMEM131L | ENSG00000121210 | 1.778966088 | 5.311865169 | 16.41756607 | 0.000316703 | 0.019654569 | Yes |
| BMP6 | ENSG00000153162 | 4.851158303 | 1.517066439 | 16.07558816 | 0.000316901 | 0.019654569 | Yes |
| TMEM163 | ENSG00000152128 | 3.529966189 | 3.089351263 | 15.00050243 | 0.000321792 | 0.019873707 | Yes |
| ENSG0000028188 | ENSG00000281881 | 3.496759275 | 4.276185897 | 15.41909045 | 0.00033193 | 0.020413678 | Yes |
| ENSG0000026638 | ENSG00000266385 | 2.584124027 | 2.936730265 | 14.90408583 | 0.000338521 | 0.020731951 | Yes |
| BEGAIN | ENSG00000183092 | 3.545391303 | 3.713708342 | 15.03419323 | 0.000342963 | 0.020916467 | Yes |
| MAL2 | ENSG00000147676 | -4.484763864 | 5.460438238 | 15.20744027 | 0.000344869 | 0.020945417 | Yes |
| MARCHF9 | ENSG00000139266 | 2.774686653 | 6.122066666 | 16.11430056 | 0.000354056 | 0.021358988 | Yes |
| ADAMTS9.AS1 | NA | 6.911057376 | 3.174938602 | 17.28733534 | 0.000354894 | 0.021358988 | Yes |
| ENSG0000026122 | ENSG00000261227 | 8.36823634 | 3.909841137 | 17.79378986 | 0.000356056 | 0.021358988 | Yes |
| CSMD1 | ENSG00000183117 | 3.841944767 | 2.873370254 | 14.90645909 | 0.000358885 | 0.021440839 | Yes |
| GPIHBP1 | ENSG00000277494 | 2.714402174 | 3.989286195 | 15.36654915 | 0.000360677 | 0.021457039 | Yes |
| UCKL1.AS1 | NA | -3.666643724 | 3.238011832 | 14.92230603 | 0.00036288 | 0.021457039 | Yes |
| EGFLAM | ENSG00000164318 | 3.27054739 | 5.031785026 | 15.65008337 | 0.000363554 | 0.021457039 | Yes |
| ENSG0000023305 | ENSG00000233058 | 3.292085099 | 2.890579079 | 14.64988092 | 0.000366409 | 0.021538662 | Yes |
| CD79A | ENSG00000105369 | 4.248774207 | 2.283513724 | 15.51970675 | 0.000372242 | 0.02172535 | Yes |
| FOS | ENSG00000170345 | 3.099185781 | 7.257462969 | 16.03360272 | 0.000372553 | 0.02172535 | Yes |
| CEBPA | ENSG00000245848 | 2.406934297 | 6.757355655 | 16.00101589 | 0.000377212 | 0.021909746 | Yes |
| NUP107 | ENSG00000111581 | 3.074602941 | 7.94818431 | 16.00306283 | 0.000379158 | 0.021935691 | Yes |
| BRSK2 | ENSG00000174672 | 4.699241648 | 2.524169127 | 15.73583825 | 0.000382711 | 0.022054114 | Yes |
| FMO1 | ENSG00000010932 | 3.44109002 | 4.483724669 | 15.16732311 | 0.00038548 | 0.022126527 | Yes |
| MGLL | ENSG00000074416 | 1.843392869 | 7.323325615 | 15.91469157 | 0.000391579 | 0.02234373 | Yes |
| CFAP61 | ENSG00000089101 | -3.721338557 | 2.768389078 | 15.07448245 | 0.000392884 | 0.02234373 | Yes |
| ZNF365 | ENSG00000138311 | -4.269248382 | 2.625755386 | 15.78354987 | 0.000393843 | 0.02234373 | Yes |
| DNAAF4 | ENSG00000256061 | -3.183175232 | 4.165420552 | 14.75549786 | 0.000400426 | 0.022629453 | Yes |
| TIAM2 | ENSG00000146426 | 2.542951166 | 4.79356449 | 15.47759677 | 0.000404784 | 0.022787774 | Yes |
| CTXN1 | ENSG00000178531 | -3.29603004 | 4.379958413 | 14.74983165 | 0.000409258 | 0.022951382 | Yes |
| MAGEL2 | ENSG00000254585 | 3.740880239 | 2.404503174 | 14.72234578 | 0.000417273 | 0.023239723 | Yes |
| EBF2 | ENSG00000221818 | 3.045626682 | 7.847173579 | 15.72038554 | 0.000417829 | 0.023239723 | Yes |
| DNAH11 | ENSG00000105877 | -6.323814921 | 3.526023062 | 16.81149229 | 0.000419528 | 0.023239723 | Yes |

| | | | | | | | |
|---|---|---|---|---|---|---|---|
| SLC4A11 | ENSG00000088836 | -4.059551041 | 4.377331496 | 14.4886788 | 0.000420751 | 0.023239723 | Yes |
| MAOB | ENSG00000069535 | 3.16818967 | 3.133615278 | 14.23741394 | 0.000438219 | 0.024113553 | Yes |
| LRRC36 | ENSG00000159708 | 2.97229151 | 4.04104925 | 14.74070966 | 0.000442455 | 0.024173735 | Yes |
| GRHL1 | ENSG00000134317 | -2.875494664 | 3.42837968 | 14.34118683 | 0.000442615 | 0.024173735 | Yes |
| WNT9A | ENSG00000143816 | -2.810324812 | 3.109668791 | 14.28504981 | 0.000445711 | 0.024187589 | Yes |
| GPRC5A | ENSG00000013588 | -3.914426182 | 6.15993809 | 15.06824005 | 0.000447644 | 0.024187589 | Yes |
| CHODL | ENSG00000154645 | 5.32812286 | 2.22137214 | 15.67498522 | 0.000447827 | 0.024187589 | Yes |
| NFIB | ENSG00000147862 | 1.834778205 | 9.56070057 | 15.49353962 | 0.000453238 | 0.024389854 | Yes |
| ENSG0000027213 | ENSG00000272130 | 4.550010222 | 3.305330985 | 14.5955783 | 0.00045556 | 0.024425049 | Yes |
| ETV4 | ENSG00000175832 | -3.530874235 | 2.836908535 | 14.43041162 | 0.00046844 | 0.025013612 | Yes |
| C6orf132 | ENSG00000188112 | -2.718692249 | 5.704730664 | 15.11579845 | 0.000469956 | 0.025013612 | Yes |
| ESR2 | ENSG00000140009 | 3.873540936 | 3.822492246 | 14.1306679 | 0.000474011 | 0.025138036 | Yes |
| WFDC3 | ENSG00000124116 | -4.852779023 | 2.014826843 | 16.01136641 | 0.000476998 | 0.025205111 | Yes |
| TMEM154 | ENSG00000170006 | -3.27406272 | 2.782177475 | 14.24203886 | 0.000482623 | 0.025410603 | Yes |
| LHX9 | ENSG00000143355 | 3.944384317 | 3.750046032 | 13.98641872 | 0.000498676 | 0.026161725 | Yes |
| PRR13 | ENSG00000205352 | -1.343224091 | 6.635353182 | 15.15192373 | 0.000506923 | 0.026399583 | Yes |
| ANKRD35 | ENSG00000198483 | -3.263060509 | 3.721973869 | 14.01723916 | 0.000507152 | 0.026399583 | Yes |
| SLC35E3 | ENSG00000175782 | 2.922078796 | 8.327046675 | 15.16311991 | 0.000508621 | 0.026399583 | Yes |
| CCDC17 | ENSG00000159588 | -4.109841515 | 4.282624319 | 13.97279525 | 0.000514611 | 0.026507136 | Yes |
| CFAP184 | ENSG00000173013 | -4.348759838 | 3.123895687 | 14.53427275 | 0.000515999 | 0.026507136 | Yes |
| ENSG0000023333 | ENSG00000233330 | 3.373657631 | 3.686346328 | 13.98015218 | 0.000516126 | 0.026507136 | Yes |
| RIMBP2 | ENSG00000060709 | 4.530524339 | 2.187238086 | 14.8427577 | 0.000519013 | 0.026562196 | Yes |
| ITGA2 | ENSG00000164171 | -3.325124996 | 4.572625446 | 14.20290443 | 0.000522806 | 0.026615222 | Yes |
| DHCR24 | ENSG00000116133 | -3.122489956 | 7.339308552 | 15.01906385 | 0.000523685 | 0.026615222 | Yes |
| SOX1.OT | NA | 7.102966699 | 2.799600047 | 15.87468186 | 0.000526253 | 0.026653191 | Yes |
| SLC49A3 | ENSG00000169026 | 2.895593141 | 2.673117681 | 13.69510246 | 0.000531316 | 0.026816785 | Yes |
| SPRR2A | ENSG00000241794 | -5.005400644 | 3.229542378 | 15.00525943 | 0.000534266 | 0.026873024 | Yes |
| G0S2 | ENSG00000123689 | 2.88091622 | 6.859947142 | 14.97014759 | 0.000538201 | 0.026887184 | Yes |
| S100A6 | ENSG00000197956 | -2.093361178 | 10.56022622 | 15.00269436 | 0.000538221 | 0.026887184 | Yes |
| CMKLR2 | ENSG00000183671 | -3.951901694 | 2.534139375 | 14.6734587 | 0.000541011 | 0.026934614 | Yes |
| C1QTNF4 | ENSG00000172247 | 3.878736362 | 2.205264853 | 14.22835091 | 0.00055522 | 0.027548344 | Yes |
| AQP3 | ENSG00000165272 | -3.291876726 | 4.729623532 | 14.08021265 | 0.000566726 | 0.027992054 | Yes |
| AHNAK2 | ENSG00000185567 | -3.193311584 | 6.646976455 | 14.70491876 | 0.000567988 | 0.027992054 | Yes |
| FAHD2B | ENSG00000144199 | 2.607741192 | 2.890427758 | 13.6065249 | 0.000570337 | 0.028013496 | Yes |
| OSCP1 | ENSG00000116885 | -2.774389933 | 4.538758799 | 14.09981781 | 0.00058521 | 0.028647867 | Yes |
| MTA1 | ENSG00000182979 | 1.394146868 | 6.696037087 | 14.74562188 | 0.000588528 | 0.028714282 | Yes |
| AREG | ENSG00000109321 | -4.006674327 | 4.475115464 | 13.66457096 | 0.0005927 | 0.028821776 | Yes |
| SDC4 | ENSG00000124145 | -2.497357552 | 6.099238041 | 14.56568199 | 0.000598481 | 0.029006496 | Yes |
| DOCK6 | ENSG00000130158 | 1.424427955 | 5.969232044 | 14.66172713 | 0.000601404 | 0.029051981 | Yes |
| SERPINE2 | ENSG00000135919 | 2.496883371 | 8.974272601 | 14.67578751 | 0.000604573 | 0.029108992 | Yes |
| ACACB | ENSG00000076555 | 2.96347996 | 8.175828008 | 14.65341319 | 0.000608757 | 0.029214351 | Yes |
| HOMER3 | ENSG00000051128 | -1.436557475 | 7.283890635 | 14.61256836 | 0.000616686 | 0.029498133 | Yes |
| LAD1 | ENSG00000159166 | -7.740412445 | 3.951534601 | 16.14308869 | 0.000621838 | 0.029553848 | Yes |
| GRIA2 | ENSG00000120251 | 5.854259139 | 1.757602234 | 14.46726055 | 0.000621889 | 0.029553848 | Yes |
| NRAV | ENSG00000248008 | -5.750829419 | 3.020100374 | 15.43276675 | 0.000627233 | 0.029592952 | Yes |
| RFXAP | ENSG00000133111 | 2.346582553 | 3.392146964 | 13.71184301 | 0.000628584 | 0.029592952 | Yes |
| CFAP157 | ENSG00000160401 | -3.974227051 | 3.718418787 | 13.50902054 | 0.000628777 | 0.029592952 | Yes |
| FBXO15 | ENSG00000141665 | -3.979393543 | 3.62511421 | 13.50102835 | 0.000636319 | 0.029851919 | Yes |
| SYDE2 | ENSG00000097096 | 2.512759409 | 4.469790624 | 14.08952071 | 0.000646758 | 0.030244699 | Yes |
| BEST4 | ENSG00000142959 | -4.445018923 | 2.870796068 | 14.2570443 | 0.000652557 | 0.030418725 | Yes |
| SLC19A1 | ENSG00000173638 | 2.222313266 | 4.852485447 | 14.20874596 | 0.000658675 | 0.030549313 | Yes |
| SEMA6A | ENSG00000092421 | 2.667260467 | 5.972879418 | 14.34571398 | 0.000659533 | 0.030549313 | Yes |
| FOXJ1 | ENSG00000129654 | -4.273261111 | 3.924747082 | 13.39030341 | 0.000666045 | 0.030753614 | Yes |
| OXTR | ENSG00000180914 | -3.050571348 | 3.146918663 | 13.28491847 | 0.000674191 | 0.031020888 | Yes |
| SLCO1C1 | ENSG00000139155 | 3.855522624 | 4.671274869 | 13.62752431 | 0.000676768 | 0.031020888 | Yes |
| WDR31 | ENSG00000148225 | -2.748987011 | 3.640835875 | 13.37347758 | 0.000678191 | 0.031020888 | Yes |
| DPCD | ENSG00000166171 | -2.09061134 | 4.838095307 | 14.00606711 | 0.000680561 | 0.031032315 | Yes |
| RSPH9 | ENSG00000172426 | -3.673529541 | 4.317341284 | 13.32396188 | 0.000688619 | 0.031147095 | Yes |
| AGBL2 | ENSG00000165923 | -5.990823551 | 3.226954915 | 15.17749484 | 0.000690571 | 0.031147095 | Yes |
| FRMD1 | ENSG00000153303 | 6.457867479 | 2.241854972 | 14.59702509 | 0.000692326 | 0.031147095 | Yes |
| PPP1R14B.AS1 | NA | -3.113836285 | 2.777005624 | 13.29241591 | 0.000693314 | 0.031147095 | Yes |
| NHSL3 | ENSG00000162522 | -2.588420149 | 4.268562037 | 13.60564649 | 0.000693718 | 0.031147095 | Yes |
| CCL19 | ENSG00000172724 | 3.4805788 | 3.331119973 | 13.13150492 | 0.00070143 | 0.031397039 | Yes |
| USHBP1 | ENSG00000130307 | 3.358956481 | 3.353576042 | 13.11423436 | 0.000706322 | 0.03151962 | Yes |
| SYT1 | ENSG00000067715 | 4.232403039 | 2.782644238 | 13.48413898 | 0.000711342 | 0.031569252 | Yes |
| EDARADD | ENSG00000186197 | -3.642651355 | 3.471702413 | 13.18344338 | 0.000713604 | 0.031569252 | Yes |
| DCX | ENSG00000077279 | 3.927360163 | 2.647967288 | 13.40608064 | 0.000715145 | 0.031569252 | Yes |
| MORN2 | ENSG00000188010 | -2.696970618 | 5.31929432 | 13.87728835 | 0.000716061 | 0.031569252 | Yes |
| TP73 | ENSG00000078900 | -3.372636015 | 3.985444146 | 13.1888944 | 0.000721812 | 0.03166544 | Yes |
| LINC01638 | ENSG00000233521 | 3.223773929 | 2.603693427 | 13.01455344 | 0.000724406 | 0.03166544 | Yes |
| DCP1A | ENSG00000272886 | 1.648997696 | 5.763804307 | 14.12472213 | 0.000724733 | 0.03166544 | Yes |
| ENSG0000027976 | ENSG00000279762 | 6.932276095 | 2.67049026 | 14.72808942 | 0.00073866 | 0.032177896 | Yes |
| DMRTA1 | ENSG00000176399 | -3.074040431 | 3.099099022 | 13.01184631 | 0.000756979 | 0.032878048 | Yes |
| PRKAR1B | ENSG00000188191 | 2.148539484 | 5.552496378 | 13.94536587 | 0.000761365 | 0.032970694 | Yes |
| CENPV | ENSG00000166582 | 1.859413889 | 5.489288969 | 13.94196851 | 0.000765645 | 0.033058232 | Yes |
| KLF5 | ENSG00000102554 | -4.028849407 | 5.976936855 | 13.48903742 | 0.00077853 | 0.033453415 | Yes |
| LRRTM2 | ENSG00000146006 | 4.37559635 | 2.29149337 | 13.64332423 | 0.000779368 | 0.033453415 | Yes |
| SYTL5 | ENSG00000147041 | -6.907984243 | 2.453090314 | 15.13014281 | 0.0007916 | 0.033879075 | Yes |
| RGS6 | ENSG00000182732 | 5.760892076 | 1.707107733 | 13.73465886 | 0.000795862 | 0.033962171 | Yes |
| TRUB2 | ENSG00000167112 | 1.63810041 | 5.70292229 | 13.85465947 | 0.000798553 | 0.033977951 | Yes |
| IP6K3 | ENSG00000161896 | -5.499501634 | 2.098249114 | 14.64661408 | 0.000808297 | 0.034213684 | Yes |
| CCDC148 | ENSG00000153237 | -4.173840094 | 2.692179346 | 13.61554829 | 0.000808768 | 0.034213684 | Yes |
| TSGA10 | ENSG00000135951 | -2.961502849 | 3.730622623 | 12.90837089 | 0.000814707 | 0.034365591 | Yes |
| HYDIN | ENSG00000157423 | -4.206447879 | 3.725773107 | 12.90470836 | 0.000822424 | 0.03454219 | Yes |
| NFATC2 | ENSG00000101096 | 2.368529081 | 5.907158646 | 13.75125253 | 0.000823613 | 0.03454219 | Yes |
| OLFM4 | ENSG00000102837 | -4.217135135 | 4.320242969 | 12.78842051 | 0.000837432 | 0.035021394 | Yes |
| PRRT4 | ENSG00000224940 | 3.239604608 | 2.648380933 | 12.66563705 | 0.000840335 | 0.035042686 | Yes |
| KIF9 | ENSG00000088727 | -3.412869562 | 4.67302723 | 12.99607133 | 0.000848108 | 0.035118313 | Yes |
| PECR | ENSG00000115425 | 3.126208852 | 3.628193019 | 12.82329413 | 0.000850107 | 0.035118313 | Yes |
| GNG7 | ENSG00000176533 | 2.838326556 | 4.295569005 | 13.22309212 | 0.000851166 | 0.035118313 | Yes |
| DNAAF11 | ENSG00000129295 | -2.977159485 | 4.738775569 | 13.15056244 | 0.000851746 | 0.035118313 | Yes |
| KAZALD1 | ENSG00000107821 | 2.246935022 | 4.985177142 | 13.52712154 | 0.000858509 | 0.03529776 | Yes |
| ENSG0000026106 | ENSG00000261068 | -3.662332278 | 2.914875502 | 12.87960951 | 0.000873502 | 0.035654819 | Yes |
| ANXA2 | ENSG00000182718 | -1.914353765 | 9.352352807 | 13.65350072 | 0.000873583 | 0.035654819 | Yes |
| ENSG0000027948 | ENSG00000279483 | -3.641090219 | 5.386115408 | 13.10225394 | 0.000874502 | 0.035654819 | Yes |
| JCHAIN | ENSG00000132465 | 4.267842504 | 5.54408569 | 13.18318214 | 0.000877842 | 0.03568369 | Yes |
| ITGB6 | ENSG00000115221 | -3.344799849 | 3.903244316 | 12.69468705 | 0.000880086 | 0.03568369 | Yes |
| MGP | ENSG00000111341 | 2.242391981 | 9.640812186 | 13.61626591 | 0.000885477 | 0.035764108 | Yes |
| ARHGEF15 | ENSG00000198844 | 3.010782623 | 4.053397902 | 12.95371532 | 0.000886956 | 0.035764108 | Yes |
| NRK | ENSG00000123572 | 3.180807185 | 2.888339504 | 12.50291029 | 0.000893379 | 0.035916662 | Yes |
| SCARF1 | ENSG00000074660 | 3.092665385 | 3.383348445 | 12.61319082 | 0.000895647 | 0.035916662 | Yes |
| CTU1 | ENSG00000142544 | 1.875628183 | 4.6021832 | 13.33120713 | 0.000909132 | 0.036357816 | Yes |
| TMEM184A | ENSG00000164855 | -2.935273458 | 3.587378009 | 12.60350572 | 0.000912941 | 0.03641066 | Yes |
| OBSL1 | ENSG00000124006 | 1.629022624 | 7.418098649 | 13.52116748 | 0.000917027 | 0.036474265 | Yes |
| VILL | ENSG00000136059 | -2.714061191 | 3.633480842 | 12.60821491 | 0.000930808 | 0.036844773 | Yes |
| VSIG8 | ENSG00000243284 | -3.769574082 | 2.700187169 | 12.94014396 | 0.000932368 | 0.036844773 | Yes |
| SHD | ENSG00000105251 | 5.286858568 | 1.380547959 | 13.08412487 | 0.000933894 | 0.036844773 | Yes |
| CHDH | ENSG00000016391 | 2.928013611 | 2.977724766 | 12.35672612 | 0.000952853 | 0.037491673 | Yes |
| GIPC3 | ENSG00000179855 | 3.54291765 | 2.501077173 | 12.54798071 | 0.000959964 | 0.03764526 | Yes |
| NHERF1 | ENSG00000109062 | -2.805370034 | 5.832549031 | 13.18685729 | 0.000963573 | 0.03764526 | Yes |
| SPHK1 | ENSG00000176170 | -2.747367803 | 4.431942537 | 12.78249899 | 0.000964472 | 0.03764526 | Yes |
| STRADB | ENSG00000082146 | 2.477975567 | 4.627999717 | 13.0777655 | 0.000970763 | 0.037760286 | Yes |
| PNMA2 | ENSG00000240694 | 2.566733194 | 2.620459741 | 12.27602933 | 0.000972578 | 0.037760286 | Yes |
| MZB1 | ENSG00000170476 | 6.132712846 | 2.473701653 | 13.72815511 | 0.000987207 | 0.038226851 | Yes |
| ATP5IF1 | ENSG00000130770 | -1.754880394 | 7.587200292 | 13.30476987 | 0.000990978 | 0.038271625 | Yes |
| SNCG | ENSG00000173267 | 3.031302764 | 3.27033633 | 12.32184123 | 0.000997457 | 0.038420484 | Yes |
| KCNMB4 | ENSG00000135643 | 3.470749171 | 5.680223036 | 13.06939082 | 0.001002626 | 0.038453201 | Yes |
| KCNK5 | ENSG00000164626 | -3.54786669 | 2.461980385 | 12.78017027 | 0.001004523 | 0.038453201 | Yes |
| CRACDL | ENSG00000196872 | -3.289528359 | 3.611040956 | 12.34345559 | 0.001006188 | 0.038453201 | Yes |
| EFHB | ENSG00000163576 | -3.001807857 | 3.557505986 | 12.32731007 | 0.001019878 | 0.038874886 | Yes |
| CSRNP3 | ENSG00000178662 | 2.762098131 | 4.938060405 | 12.95255815 | 0.001030189 | 0.03916593 | Yes |
| CTNNBL1 | ENSG00000132792 | -1.518409241 | 6.157919108 | 13.14080864 | 0.001038607 | 0.039260401 | Yes |
| CAVIN2 | ENSG00000168497 | 2.185890539 | 4.771477555 | 12.98174789 | 0.00103934 | 0.039260401 | Yes |
| PTP4A3 | ENSG00000184489 | 1.57995949 | 5.708267461 | 13.13785349 | 0.001040721 | 0.039260401 | Yes |
| BMP2 | ENSG00000125845 | -2.404625964 | 3.414326022 | 12.31324121 | 0.001048293 | 0.039444388 | Yes |
| MKRN3 | ENSG00000179455 | 3.96681104 | 2.924211619 | 12.35634636 | 0.001054812 | 0.039563172 | Yes |
| TENT5B | ENSG00000158246 | -2.955811114 | 2.291584195 | 12.41187297 | 0.001057554 | 0.039563172 | Yes |
| PRSS12 | ENSG00000164099 | -3.11295387 | 3.403340417 | 12.21128058 | 0.001059559 | 0.039563172 | Yes |
| MBD6 | ENSG00000166987 | 2.570955149 | 7.857115947 | 13.10094372 | 0.00106998 | 0.039850615 | Yes |
| TAGLN2 | ENSG00000158710 | -1.940181138 | 10.43936071 | 13.0938374 | 0.001073133 | 0.039866604 | Yes |
| CACNA2D2 | ENSG00000007402 | 3.662699181 | 2.736155513 | 12.20741381 | 0.001084365 | 0.040181897 | Yes |
| DEGS2 | ENSG00000168350 | -3.136725482 | 3.856731273 | 12.19326407 | 0.001096723 | 0.040537222 | Yes |
| C14orf180 | ENSG00000184601 | 3.823976071 | 2.860587517 | 12.18192501 | 0.001105601 | 0.040676317 | Yes |
| DIPK1B | ENSG00000165716 | 2.544776077 | 3.865303582 | 12.46356033 | 0.001106045 | 0.040676317 | Yes |
| SLC26A7 | ENSG00000147606 | 3.416010475 | 4.183083214 | 12.36186413 | 0.001109241 | 0.040691625 | Yes |
| SLC6A1 | ENSG00000157103 | 4.310195895 | 1.815058961 | 12.65856329 | 0.001120229 | 0.040900396 | Yes |
| NR1H3 | ENSG00000025434 | 2.044228158 | 5.009868379 | 12.82828052 | 0.001126451 | 0.040900396 | Yes |
| GSE1 | ENSG00000131149 | 1.557999746 | 6.343868231 | 12.9494593 | 0.001127419 | 0.040900396 | Yes |
| DIO3 | ENSG00000197406 | 3.408790926 | 3.82688177 | 12.16345799 | 0.001128807 | 0.040900396 | Yes |
| LOC105372725 | ENSG00000230226 | 7.985014439 | 3.598228408 | 13.91627689 | 0.001130006 | 0.040900396 | Yes |
| GABBR1 | ENSG00000204681 | 2.028980242 | 6.392491754 | 12.93245679 | 0.001131698 | 0.040900396 | Yes |
| STPG3.AS1 | NA | 3.368847851 | 2.208968278 | 12.14003391 | 0.001139552 | 0.041053494 | Yes |
| ENSG0000017468 | ENSG00000174680 | 3.706913251 | 3.079788013 | 11.98022841 | 0.001141543 | 0.041053494 | Yes |
| CD83 | ENSG00000112149 | 2.069333479 | 4.536639194 | 12.68407306 | 0.001146179 | 0.041119158 | Yes |
| LRP5 | ENSG00000162337 | 1.802046099 | 4.887348943 | 12.77254078 | 0.001151969 | 0.041225826 | Yes |
| PALD1 | ENSG00000107719 | 2.659682671 | 4.171118961 | 12.43839231 | 0.001155315 | 0.041244761 | Yes |
| CXADR | ENSG00000154639 | -3.438774376 | 4.465079524 | 12.13629913 | 0.0011676 | 0.041542896 | Yes |
| C5orf63 | ENSG00000164241 | 2.29256645 | 4.209934233 | 12.50335409 | 0.001171076 | 0.041542896 | Yes |
| SPRR2E | ENSG00000203785 | -4.726659939 | 2.439470831 | 13.20196259 | 0.00117358 | 0.041542896 | Yes |
| PLCXD3 | ENSG00000182836 | 4.087933752 | 3.321183626 | 11.97327957 | 0.001176303 | 0.041542896 | Yes |
| TRMT2A | ENSG00000099899 | 1.635465205 | 5.219371278 | 12.76518907 | 0.001177858 | 0.041542896 | Yes |
| RASIP1 | ENSG00000105538 | 2.133921863 | 5.29430126 | 12.71315943 | 0.001191406 | 0.041919735 | Yes |
| CCBE1 | ENSG00000183287 | 3.293231448 | 3.147005886 | 11.83461257 | 0.001200058 | 0.042089902 | Yes |
| ENSG0000023381 | ENSG00000233818 | -5.901734751 | 1.762912656 | 13.30233095 | 0.001202038 | 0.042089902 | Yes |
| SPINT2 | ENSG00000167642 | -3.236891647 | 4.710213441 | 12.1944028 | 0.001204869 | 0.042089902 | Yes |
| PLXNA4 | ENSG00000221866 | 2.975663993 | 3.83029633 | 12.11904842 | 0.001214058 | 0.042309934 | Yes |
| CABP1 | ENSG00000157782 | 2.803303209 | 3.519840672 | 12.01500042 | 0.001220012 | 0.042400538 | Yes |
| ECM1 | ENSG00000143369 | -2.376383392 | 5.810936074 | 12.609104 | 0.00122505 | 0.042400538 | Yes |
| CHEK2 | ENSG00000183765 | -2.991587603 | 4.344422273 | 12.09993832 | 0.001225349 | 0.042400538 | Yes |
| LOC728485 | ENSG00000267260 | 4.522533772 | 1.93809622 | 12.50371243 | 0.00123341 | 0.042578816 | Yes |
| IL7R | ENSG00000168685 | 3.838120693 | 4.490490698 | 12.03356202 | 0.001244936 | 0.042729937 | Yes |
| APLP2 | ENSG00000084234 | -1.639219761 | 9.04511836 | 12.69515733 | 0.001245363 | 0.042729937 | Yes |
| CDH3 | ENSG00000062038 | -3.873848386 | 5.756437453 | 12.25613341 | 0.001248976 | 0.042729937 | Yes |
| S100A16 | ENSG00000188643 | -2.053814711 | 7.113979943 | 12.66904199 | 0.001249465 | 0.042729937 | Yes |
| RIC3 | ENSG00000166405 | 3.762853716 | 2.179327993 | 12.0679081 | 0.001266653 | 0.043216768 | Yes |
| GFRA2 | ENSG00000168546 | 2.670325364 | 2.46889098 | 11.61889118 | 0.001272904 | 0.043329073 | Yes |
| LEP | ENSG00000174697 | 3.635270225 | 2.897175232 | 11.7147883 | 0.001293519 | 0.04392864 | Yes |
| CPEB2 | ENSG00000137449 | -2.522062637 | 4.490369727 | 12.12721665 | 0.001300185 | 0.043979197 | Yes |
| HEPN1 | ENSG00000221932 | 4.782537463 | 3.32173589 | 12.12900457 | 0.001301017 | 0.043979197 | Yes |
| RIBC2 | ENSG00000128408 | -3.487163809 | 3.476802492 | 11.72510767 | 0.001306395 | 0.044059218 | Yes |
| IQCD | ENSG00000166578 | -3.861114305 | 2.670016935 | 12.14525936 | 0.001313195 | 0.044186748 | Yes |
| MUC4 | ENSG00000145113 | -3.86492095 | 4.264431284 | 11.71604466 | 0.001324243 | 0.044456285 | Yes |
| TUBB4B | ENSG00000188229 | -2.05150836 | 8.464465956 | 12.49091104 | 0.001344187 | 0.044985035 | Yes |
| DAW1 | ENSG00000123977 | -4.931256879 | 3.067911993 | 12.56779021 | 0.00134614 | 0.044985035 | Yes |
| GREB1 | ENSG00000196208 | 2.595763359 | 3.379291074 | 11.74787242 | 0.001353865 | 0.045140138 | Yes |
| C15orf48 | ENSG00000166920 | -4.557913737 | 3.127023896 | 12.1625416 | 0.001376481 | 0.045789891 | Yes |
| PRXL2A | ENSG00000122378 | 1.794487965 | 6.86527283 | 12.40537593 | 0.001386495 | 0.046018424 | Yes |
| TOM1L1 | ENSG00000141198 | -4.242165143 | 2.461935214 | 12.4172582 | 0.00139182 | 0.046090655 | Yes |
| LGALS3 | ENSG00000131981 | -1.57479252 | 6.572279294 | 12.35692692 | 0.001403045 | 0.046357503 | Yes |
| TP63 | ENSG00000073282 | -6.418269231 | 2.076764432 | 13.11057024 | 0.001417328 | 0.046723948 | Yes |
| ABCC8 | ENSG00000006071 | 3.267933926 | 3.523557295 | 11.56847302 | 0.001421222 | 0.046740933 | Yes |
| BMAL2 | ENSG00000029153 | -2.762852376 | 2.8317346 | 11.50078977 | 0.001425395 | 0.046740933 | Yes |
| BAIAP2L1 | ENSG00000006453 | -3.063358537 | 4.317570573 | 11.70294879 | 0.001430267 | 0.046740933 | Yes |
| CSTB | ENSG00000160213 | -1.660205011 | 7.587535102 | 12.32340406 | 0.001430617 | 0.046740933 | Yes |
| GRM4 | ENSG00000124493 | 3.334972113 | 2.464867936 | 11.45531985 | 0.001439663 | 0.04682557 | Yes |
| CD36 | ENSG00000135218 | 2.82434225 | 10.30487749 | 12.30542267 | 0.00144222 | 0.04682557 | Yes |
| DDX43 | ENSG00000080007 | 3.097398406 | 3.618662354 | 11.59409291 | 0.001442805 | 0.04682557 | Yes |
| VAMP8 | ENSG00000118640 | -2.810246275 | 6.361414254 | 12.19624423 | 0.001448356 | 0.046901755 | Yes |
| BAALC | ENSG00000164929 | -2.700885099 | 4.120252459 | 11.68414904 | 0.001455343 | 0.047023978 | Yes |
| EPCAM.DT | NA | 3.703992715 | 1.689942224 | 11.78732899 | 0.00145999 | 0.047070212 | Yes |
| KRT7 | ENSG00000135480 | -4.536184435 | 5.975215095 | 11.7386837 | 0.001466187 | 0.047166106 | Yes |
| NDRG1 | ENSG00000104419 | -2.080678042 | 8.087384074 | 12.21706846 | 0.001490194 | 0.047833275 | Yes |
| PDE3A | ENSG00000172572 | 2.380997905 | 4.242253808 | 11.86653291 | 0.001504885 | 0.048199134 | Yes |
| FTH1 | ENSG00000167996 | -1.670424333 | 7.942205852 | 12.18370448 | 0.001509902 | 0.04825424 | Yes |
| COLEC11 | ENSG00000118004 | 3.304122599 | 2.268197762 | 11.36953444 | 0.00152959 | 0.048624372 | Yes |
| SLC39A11 | ENSG00000133195 | -2.235040096 | 5.28297765 | 11.96527 | 0.00153234 | 0.048624372 | Yes |
| CSAD | ENSG00000139631 | 2.255770376 | 7.012722444 | 12.13636913 | 0.001534099 | 0.048624372 | Yes |
| CPB1 | ENSG00000153002 | -2.982702755 | 4.937226057 | 11.7347534 | 0.001534772 | 0.048624372 | Yes |
| CPM | ENSG00000135678 | 2.16739552 | 6.20283324 | 12.09489743 | 0.001546994 | 0.048778907 | Yes |
| ENSG0000027244 | ENSG00000272440 | 2.603476898 | 3.490202544 | 11.48983663 | 0.001547031 | 0.048778907 | Yes |
| PFKP | ENSG00000067057 | -1.67106441 | 6.312806211 | 12.08257441 | 0.001549648 | 0.048778907 | Yes |
| BASP1 | ENSG00000176788 | -1.413375897 | 8.319699053 | 12.09970067 | 0.001559915 | 0.048996715 | Yes |
| SNRPG | ENSG00000143977 | -2.055876221 | 5.441481934 | 11.95980766 | 0.001563526 | 0.049005002 | Yes |
| OTX1 | ENSG00000115507 | -3.661679261 | 3.216514877 | 11.36438182 | 0.001569935 | 0.049100727 | Yes |
| FGF12 | ENSG00000114279 | 3.317942834 | 2.306784303 | 11.33197227 | 0.001576475 | 0.049200147 | Yes |
| SPA17 | ENSG00000064199 | -3.342373711 | 4.385467043 | 11.39241394 | 0.001594171 | 0.04964654 | Yes |
