## Supplementary material for "Integrated transcriptomic analysis identifies Paternally Expressed Gene 10 as a novel candidate biomarker associated with aggressive features in liposarcoma": Table S4_H

| ID | Description | setSize | enrichmentS | NES | pvalue | p.adjust | qvalue | rank | leading_edg | core_enrichment |
|---|---|---|---|---|---|---|---|---|---|---|
| HALLMARK_ADIPOGENESIS | HALLMARK_ADIPOGENESIS | 196 | 0.57050053 | 1.73118681 | 4.2667E-06 | 0.00021334 | 0.0001572 | 1613 | tags=23%, list | ADIPOQ/MRAP/FZD4/ANGPT1/FABP4/LIPE/DGAT1/PPARG/MGLL/SLC19A1/CAVIN2/SNCG/CD36/LEP/LPL/PTGER3/MTARC2/IDH1/SPARCL1/ANGPTL4/SORBS1/ECH1/SLC25A10/CIDEA/ACAA2/ITIH5/ITSN1/CHCHD10/IDH3G/PGM1/COL15A1/COL4A1/ABCB8/HIBCH/CS/ECHS1/ADIPOR2/SULT1A1/ETFB/PDCD4/OMD/ACADS/AGPAT3/CAVIN1/BCKDHA |
| HALLMARK_P53_PATHWAY | HALLMARK_P53_PATHWAY | 190 | -0.5287838 | -1.6523492 | 6.0561E-05 | 0.00151403 | 0.0011156 | 1030 | tags=19%, list | SDC1/MXD1/DNTTIP2/FAM162A/GM2A/BAK1/TNFSF9/VDR/FUCA1/IER5/CTSD/FDXR/EPS8L2/HSPA4L/CD82/TRIAP1/RPS27L/ISCU/TM7SF3/HINT1/SAT1/DRAM1/POLH/S100A10/VAMP8/NDRG1/BMP2/SPHK1/TP63/ZNF365/CDKN2B/ITGB4/RRAD/ST14/SFN/PERP/KRT17 |
| HALLMARK_MTORC1_SIGNALING | HALLMARK_MTORC1_SIGNALING | 197 | -0.4919366 | -1.5421663 | 0.00041933 | 0.00698891 | 0.00514973 | 3231 | tags=37%, list | DDX39A/ETF1/MCM4/TCEA1/NIBAN1/GBE1/ARPC5L/GPI/RIT1/SQSTM1/HMGCR/CCT6A/DDIT4/SLC6A6/CANX/MCM2/IMMT/PDK1/EPRS1/AURKA/ALDOA/TUBG1/PDAP1/CTSC/SLC1A4/GSR/TPI1/QDPR/TBK1/SDF2L1/SERP1/CDKN1A/ACTR3/PSPH/SLC7A11/NMT1/MLLT11/ATP6V1D/YKT6/LDHA/XBP1/ACSL3/PSMD12/SYTL2/RDH11/BCAT1/NFKBIB/POLR3G/HSPA5/GAPDH/PSME3/ATP2A2/BUB1/TES/PGK1/SRD5A1/PSMA4/SLC1A5/FDXR/ENO1/PIK3R3/UBE2D3/EIF2S2/TXNRD1/SC5D/SLC7A5/EBP/PRDX1/CD9/NHERF1/DHCR24/TUBA4A/SLC2A1 |
| HALLMARK_ANDROGEN_RESPONSE | HALLMARK_ANDROGEN_RESPONSE | 91 | -0.5944968 | -1.6894014 | 0.00073506 | 0.00918829 | 0.00677032 | 2524 | tags=32%, list | B2M/SPCS3/GNAI3/RPS6KA3/PA2G4/GSR/MYL12A/PMEPA1/ACSL3/STK39/ABCC4/XRCC6/ACTN1/PTK2B/SLC26A2/ITGAV/INPP4B/PDLIM5/ALDH1A3/ADRM1/ELL2/SAT1/SMS/MAP7/NDRG1/DHCR24/UAP1/KRT8/KRT19 |
| HALLMARK_REACTIVE_OXYGEN_SPECIES_PATHWAY | HALLMARK_REACTIVE_OXYGEN_SPECIES_PATHWAY | 44 | -0.6880758 | -1.7307115 | 0.00095082 | 0.00950819 | 0.00700604 | 3032 | tags=45%, list | PTPA/GLRX2/TXN/SBNO2/ATOX1/GSR/NDUFA6/PRDX4/SOD1/MBP/GCLM/NDUFS2/LAMTOR5/CDKN2D/SELENOS/PDLIM1/TXNRD1/NQO1/PRDX1/PFKP |
| HALLMARK_MYC_TARGETS_V1 | HALLMARK_MYC_TARGETS_V1 | 200 | -0.478152 | -1.501495 | 0.00136047 | 0.01133726 | 0.00835377 | 3535 | tags=34%, list | DDX18/VDAC3/HSPE1/AIMP2/CDK2/EIF4E/PSMC4/ACP1/ETF1/MCM4/PWP1/CLNS1A/PCNA/TARDBP/CANX/MCM2/UBE2L3/EPRS1/PSMD8/PSMB3/GSPT1/RAD23B/PHB1/SNRPD3/CBX3/PSMB2/SMARCC1/PSMD3/PA2G4/SET/SSBP1/RRM1/ERH/GOT2/PRDX4/HDGF/EIF4G2/YWHAQ/SSB/SNRPB2/XPOT/LDHA/PSMD1/CSTF2/XRCC6/POLE3/PTGES3/KPNB1/PGK1/NAP1L1/TFDP1/PSMA4/UBE2E1/SRPK1/EIF1AX/EIF2S1/DEK/RPLP0/EIF2S2/KPNA2/CCT5/PSMA7/RAN/PRPS2/H2AZ1/COX5A/SNRPG |
| HALLMARK_E2F_TARGETS | HALLMARK_E2F_TARGETS | 200 | -0.4697549 | -1.4751264 | 0.00235867 | 0.01684764 | 0.01241405 | 4779 | tags=49%, list | DEPDC1/CIT/MLH1/UBR7/RNASEH2A/SNRPB/MRE11/SRSF1/SRSF2/CDC25A/RFC3/BUB1B/TFRC/MCM6/PSMC3IP/POLD1/GINS3/PLK4/RANBP1/MCM3/TK1/CKS2/RBBP7/TMPO/CDC20/NME1/NUP205/SLBP/TUBB/PPP1R8/SPC24/WEE1/EED/DUT/MTHFD2/HMGB3/PRPS1/RAD50/KIF18B/SPAG5/RAD51AP1/PHF5A/RRM2/RACGAP1/ASF1A/DDX39A/ANP32E/MCM4/PTTG1/SMC3/PCNA/MCM2/GINS1/TCF19/CCP110/DONSON/AURKA/MELK/RAD1/CDKN2C/SUV39H1/GSPT1/TUBG1/CCNB2/DNMT1/CDKN2A/PA2G4/SMC4/SSRP1/POP7/PRDX4/HUS1/CDKN1A/SMC1A/TIPIN/RAD51C/HMGA1/CDCA8/XRCC6/BIRC5/UBE2T/RPA2/CENPE/CSE1L/SMC6/POLD3/ASF1B/NAP1L1/POLE4/CDC25B/JPT1/EIF2S1/DEK/KPNA2/CENPM/RAN/H2AZ1/CHEK2 |
| HALLMARK_ESTROGEN_RESPONSE_EARLY | HALLMARK_ESTROGEN_RESPONSE_EARLY | 176 | -0.4642846 | -1.4367422 | 0.00454752 | 0.02842201 | 0.02094253 | 1287 | tags=23%, list | CD44/GJA1/SLC26A2/FRK/MLPH/WWC1/MYOF/TTC39A/ARL3/INPP5F/MYB/ENDOD1/CLIC3/DYNLT3/PEX11A/SLC16A1/FHL2/RBBP8/FOXC1/SEC14L2/SIAH2/SLC7A5/MUC1/ALDH3B1/TFF3/REEP1/SLC39A6/MICB/KCNK5/NHERF1/AREG/AQP3/LAD1/NBL1/KRT8/SCNN1A/SLC2A1/SFN/CLDN7/ELF3/KRT19 |
| HALLMARK_ESTROGEN_RESPONSE_LATE | HALLMARK_ESTROGEN_RESPONSE_LATE | 167 | -0.4680727 | -1.4365365 | 0.00567408 | 0.03152267 | 0.02322723 | 895 | tags=20%, list | LLGL2/MYOF/ARL3/MYB/TST/CLIC3/DYNLT3/UGDH/SLC16A1/HSPA4L/RBBP8/SERPINA1/FOXC1/SIAH2/SLC7A5/ALDH3B1/LSR/CD9/TFF3/FABP5/MICB/KCNK5/NHERF1/AREG/ST14/NBL1/SCNN1A/ST6GALNAC2/S100A9/LAMC2/SFN/PERP/KRT19 |
| HALLMARK_PROTEIN_SECRETION | HALLMARK_PROTEIN_SECRETION | 94 | -0.5237112 | -1.4956126 | 0.00792461 | 0.03962305 | 0.02919593 | 3820 | tags=38%, list | PAM/AP3B1/COPE/RER1/CD63/ADAM10/DOP1A/VPS45/ANP32E/BNIP3/RAB22A/SNAP23/VPS4B/CLTA/ABCA1/TMX1/CLN5/RPS6KA3/GOSR2/CTSC/LAMP2/KIF1B/SCAMP3/GOLGA4/ATP1A1/AP2B1/CLCN3/YKT6/SEC22B/SOD1/TMED10/CLTC/EGFR/AP2M1/DST/TOM1L1 |
| HALLMARK_COMPLEMENT | HALLMARK_COMPLEMENT | 166 | -0.4420528 | -1.3577143 | 0.01619798 | 0.07362716 | 0.05425159 | 1652 | tags=19%, list | ACTN2/PPP2CB/PDP1/GPD2/HSPA5/FN1/CD59/CASP4/PSEN1/PHEX/IRF2/CTSH/S100A13/MMP14/TIMP2/CASP3/CTSB/USP16/CTSD/SERPINB2/CP/SERPINA1/LYN/F3/C3/CFB/KYNU/LGALS3/S100A9/MMP13/CXCL1 |
| HALLMARK_APOPTOSIS | HALLMARK_APOPTOSIS | 143 | -0.4464281 | -1.3493222 | 0.02188813 | 0.09120053 | 0.06720039 | 3285 | tags=34%, list | CASP8/NEFH/ETF1/LMNA/IRF1/SQSTM1/BID/CD38/TSPO/GCH1/FAS/CCND1/BCL2L1/RNASEL/GADD45A/PSEN2/TAP1/DNAJC3/IER3/TNFSF10/IGFBP6/GSR/ERBB3/MGMT/DNAJA1/CDKN1A/GNA15/SOD1/PMAIP1/TNFRSF12A/CD44/DAP3/CASP4/PSEN1/CDC25B/HGF/TIMP2/CASP3/GPX1/FDXR/SC5D/IL18/EBP/TGFB2/SAT1/LEF1/ENO2/BMP2/LGALS3 |
| HALLMARK_APICAL_JUNCTION | HALLMARK_APICAL_JUNCTION | 173 | -0.4262558 | -1.3154226 | 0.02480655 | 0.09540982 | 0.07030198 | 2084 | tags=20%, list | CADM2/NECTIN1/MYH9/ITGA3/SRC/SHROOM2/NRAP/CTNNA1/ACTN2/MYL12B/ARPC2/ACTN1/ADAMTS5/CD99/EGFR/LDLRAP1/LAYN/ACTN4/NEGR1/LAMA3/ITGB1/PIK3R3/AMH/TJP1/MPZL2/CDH3/ITGA2/ITGB4/ITGA10/DSC3/LAMC2/CLDN4/CLDN7/LAMB3 |
| HALLMARK_COAGULATION | HALLMARK_COAGULATION | 95 | -0.4689176 | -1.3408944 | 0.03860176 | 0.13786342 | 0.10158358 | 1599 | tags=24%, list | PLAU/RAC1/PROS1/FN1/DUSP14/HTRA1/CTSK/CTSH/WDR1/S100A13/MMP14/CTSB/P2RY1/SERPINB2/SERPINA1/F3/C3/ISCU/CFB/CD9/ITGB3/GDA/ITGA2 |
| HALLMARK_KRAS_SIGNALING_DN | HALLMARK_KRAS_SIGNALING_DN | 106 | 0.472984 | 1.33223757 | 0.0443488 | 0.14782934 | 0.10892688 | 1145 | tags=16%, list | CNTFR/HSD11B2/ZBTB16/RYR2/GRID2/RGS11/COL2A1/TCF7L1/BTG2/KLHDC8A/NOS1/MYO15A/OXT/SLC16A7/GTF3C5/IFI44L/SLC25A23 |
| HALLMARK_HEME_METABOLISM | HALLMARK_HEME_METABOLISM | 163 | -0.4089201 | -1.2524718 | 0.05211141 | 0.16284816 | 0.11999338 | 2352 | tags=20%, list | OPTN/YPEL5/LAMP2/SYNJ1/TSPAN5/BPGM/HDGF/BMP2K/TOP1/SLC7A11/FN3K/CLCN3/SLC25A37/BLVRA/SMOX/UBAC1/GCLM/RNF123/LRP10/SLC25A38/DCAF11/VEZF1/GAPVD1/CTSB/HTATIP2/ENDOD1/MBOAT2/ELL2/C3/GDE1/ANK1/AQP3/SLC2A1 |
| HALLMARK_EPITHELIAL_MESENCHYMAL_TRANSITION | HALLMARK_EPITHELIAL_MESENCHYMAL_TRANSITION | 191 | -0.3901622 | -1.2203661 | 0.06251416 | 0.18386517 | 0.1354796 | 1558 | tags=21%, list | ITGB5/MXRA5/LOXL2/FAP/FN1/CD59/TNFRSF12A/CD44/TPM1/GJA1/HTRA1/SCG2/CDH2/SDC1/ITGAV/LAMA3/ITGB1/GREM1/MMP14/ANPEP/FUCA1/QSOX1/COL7A1/IGFBP2/TPM4/DST/SPP1/RGS4/SAT1/ITGB3/LRRC15/ENO2/BASP1/ECM1/OXTR/AREG/ITGA2/SDC4/INHBA/LAMC2/CXCL1 |
| HALLMARK_G2M_CHECKPOINT | HALLMARK_G2M_CHECKPOINT | 197 | -0.3823687 | -1.1986832 | 0.07760381 | 0.21556614 | 0.15883821 | 4825 | tags=43%, list | NOTCH2/YTHDC1/WRN/H2AZ2/KIF5B/CENPA/PRC1/TENT4A/SRSF1/TGFB1/SRSF2/CDC25A/E2F3/NUP98/MCM6/TNPO2/PLK4/ODC1/UBE2C/CDC7/MCM3/CKS2/PBK/RBL1/TMPO/CDC20/TTK/SLC12A2/DKC1/SLC7A1/G3BP1/HIF1A/BCL3/SMAD3/PURA/AMD1/HMGB3/DR1/CUL4A/RAD54L/RACGAP1/DDX39A/PTTG1/SLC38A1/CCND1/KNL1/NUSAP1/MCM2/CDC27/STIL/AURKA/CDKN2C/SUV39H1/GSPT1/RAD23B/SMARCC1/CCNB2/SMC4/CDC6/HUS1/NDC80/TOP1/MT2A/PRP4K/SMC1A/UPF1/DMD/HMGA1/CASP8AP2/BIRC5/RPA2/BUB1/KPNB1/SAP30/CENPE/TFDP1/NUP50/KMT5A/CDC25B/JPT1/KPNA2/SLC7A5/H2AZ1/CHAF1A/HSPA8 |
| HALLMARK_HEDGEHOG_SIGNALING | HALLMARK_HEDGEHOG_SIGNALING | 31 | 0.56672002 | 1.31341839 | 0.11194324 | 0.25660521 | 0.18907752 | 1675 | tags=26%, list | CNTFR/NRCAM/GLI1/VLDLR/THY1/PTCH1/ACHE/NF1 |
| HALLMARK_CHOLESTEROL_HOMEOSTASIS | HALLMARK_CHOLESTEROL_HOMEOSTASIS | 70 | -0.4613477 | -1.2529525 | 0.10353643 | 0.25660521 | 0.18907752 | 626 | tags=16%, list | JAG1/MVK/SC5D/EBP/CD9/FABP5/S100A11/CPEB2/LGALS3/MAL2/ALCAM |
| HALLMARK_BILE_ACID_METABOLISM | HALLMARK_BILE_ACID_METABOLISM | 82 | 0.45481545 | 1.23379148 | 0.1180384 | 0.25660521 | 0.18907752 | 1366 | tags=21%, list | LIPE/RXRG/AR/BMP6/PECR/ACSL5/IDH1/ABCA2/LCK/ISOC1/CYP46A1/PEX19/CYP7B1/RXRA/ABCA8/SLC27A5/ABCD2 |
| HALLMARK_FATTY_ACID_METABOLISM | HALLMARK_FATTY_ACID_METABOLISM | 145 | 0.41546819 | 1.21770295 | 0.10563766 | 0.25660521 | 0.18907752 | 1992 | tags=26%, list | CA4/AQP7/GPD1/MGLL/FMO1/G0S2/CD36/ACADVL/ACSL5/IDH1/ECH1/AOC3/CIDEA/ACAA2/GCDH/ACOT2/HSDL2/IDH3G/BPHL/HIBCH/ECHS1/ADIPOR2/EHHADH/ACADS/D2HGDH/ETFDH/SDHD/UROS/PPARA/GSTZ1/HADH/EPHX1/DLD/HPGD/PRDX6/GRHPR/SUCLA2 |
| HALLMARK_MYOGENESIS | HALLMARK_MYOGENESIS | 189 | 0.39134208 | 1.18219101 | 0.11596278 | 0.25660521 | 0.18907752 | 1475 | tags=17%, list | MYH11/CAMK2B/PTP4A3/CD36/FST/PC/IGF1/GNAO1/MYOM1/SORBS3/FXYD1/SORBS1/NOS1/SCD/SGCG/MYO1C/CASQ2/HRC/PICK1/NCAM1/COL4A2/TSC2/MRAS/TAGLN/PGAM2/COL15A1/KIFC3/APLNR/EPHB3/COL3A1/VIPR1/SVIL |
| HALLMARK_PEROXISOME | HALLMARK_PEROXISOME | 90 | 0.42320303 | 1.16581789 | 0.18125236 | 0.35045256 | 0.2582282 | 1512 | tags=14%, list | HSD11B2/RXRG/ESR2/ABCC8/CRABP1/ACSL5/IDH1/ECH1/ISOC1/ABCD2/ABCB1/EHHADH/ABCB9 |
| HALLMARK_PI3K_AKT_MTOR_SIGNALING | HALLMARK_PI3K_AKT_MTOR_SIGNALING | 99 | -0.4003503 | -1.1507613 | 0.1854804 | 0.35045256 | 0.2582282 | 2022 | tags=29%, list | GRK2/DAPP1/CDK2/EIF4E/PAK4/PRKAG1/MAPKAP1/RPS6KA1/RIT1/SQSTM1/PRKAR2A/PDK1/RPS6KA3/TBK1/PLA2G12A/CDKN1A/ACTR3/PTPN11/RAC1/NFKBIB/MKNK2/CLTC/EGFR/AP2M1/YWHAB/PIK3R3/UBE2D3/SLC2A1/SFN |
| HALLMARK_DNA_REPAIR | HALLMARK_DNA_REPAIR | 148 | -0.3756492 | -1.138578 | 0.17139665 | 0.35045256 | 0.2582282 | 2345 | tags=21%, list | POLR3GL/ELOA/NT5C3A/FEN1/SSRP1/POLR1H/POLR2E/CCNO/BRF2/POLA1/ZWINT/RALA/MPC2/TAF6/RPA2/DAD1/UPF3B/NUDT9/POLD3/GSDME/SNAPC5/POLE4/SUPT4H1/POLR2I/GTF2A2/TAF13/ADRM1/STX3/RAE1/POLH/CETN2 |
| HALLMARK_TNFA_SIGNALING_VIA_NFKB | HALLMARK_TNFA_SIGNALING_VIA_NFKB | 182 | -0.3581768 | -1.1127365 | 0.18924438 | 0.35045256 | 0.2582282 | 874 | tags=13%, list | NFE2L2/PDLIM5/FOSL1/TNFSF9/IER5/JAG1/SERPINB2/GFPT2/IL18/F3/B4GALT5/ZC3H12A/SAT1/DRAM1/KYNU/BMP2/SPHK1/AREG/SDC4/PHLDA2/INHBA/LAMB3/CXCL1 |
| HALLMARK_IL2_STAT5_SIGNALING | HALLMARK_IL2_STAT5_SIGNALING | 177 | -0.3563549 | -1.1037424 | 0.20749149 | 0.35774394 | 0.2636008 | 1026 | tags=23%, list | BCL2L1/LRIG1/UMPS/PHLDA1/PLSCR1/S100A1/CAPG/TNFSF10/ADAM19/SWAP70/PTRH2/SYNGR2/SYT11/PRAF2/CDC6/TNFRSF9/RHOH/XBP1/IL4R/CD44/MXD1/ITGAV/MAP6/TNFRSF21/CASP3/SLC1A5/RABGAP1L/MYO1E/NCS1/GSTO1/PTGER2/SPP1/ARL4A/MUC1/CDCP1/NDRG1/BMP2/ECM1/ETV4/ALCAM |
| HALLMARK_MITOTIC_SPINDLE | HALLMARK_MITOTIC_SPINDLE | 198 | -0.3518164 | -1.1030583 | 0.20300068 | 0.35774394 | 0.2636008 | 2780 | tags=20%, list | NUSAP1/CDC27/SEPTIN9/DOCK4/AURKA/CDC42/CLASP1/ARHGEF2/CCNB2/PPP4R2/ARHGAP5/KIF1B/SASS6/SMC4/CKAP5/MYH9/CAPZB/MYO9B/NDC80/SHROOM2/PCM1/SMC1A/ABI1/BIRC5/KATNB1/CLIP1/BUB1/CENPE/ACTN4/PDLIM5/STAU1/RALBP1/SORBS2/MYO1E/ARF6/PXN/DST/KIF3B/EZR/TUBA4A |
| HALLMARK_PANCREAS_BETA_CELLS | HALLMARK_PANCREAS_BETA_CELLS | 19 | 0.54118479 | 1.13260108 | 0.30394338 | 0.48092998 | 0.35436946 | 2055 | tags=42%, list | DCX/ABCC8/STXBP1/PCSK2/FOXO1/PAK3/LMO2/AKT3 |
| HALLMARK_MYC_TARGETS_V2 | HALLMARK_MYC_TARGETS_V2 | 57 | 0.42295998 | 1.08940607 | 0.30779519 | 0.48092998 | 0.35436946 | 2237 | tags=19%, list | SLC19A1/WDR74/UNG/TMEM97/CDK4/TCOF1/PPRC1/RRP12/MYBBP1A/MCM5/DUSP2 |
| HALLMARK_INFLAMMATORY_RESPONSE | HALLMARK_INFLAMMATORY_RESPONSE | 151 | -0.3490227 | -1.0613966 | 0.30040053 | 0.48092998 | 0.35436946 | 2745 | tags=28%, list | SLC31A2/ABCA1/EMP3/ADORA2B/GNAI3/P2RX7/TNFSF10/ATP2B1/TPBG/SRI/KIF1B/RNF144B/CDKN1A/TNFRSF9/SGMS2/GNA15/ABI1/IL4R/ATP2A2/OSMR/PTAFR/DCBLD2/PSEN1/CLEC5A/NMI/MXD1/AXL/MMP14/TNFSF9/SELENOS/ADRM1/CD40/BDKRB1/CD82/IL18/LYN/PTGER2/F3/MET/ITGB3/SPHK1/INHBA |
| HALLMARK_UV_RESPONSE_DN | HALLMARK_UV_RESPONSE_DN | 143 | 0.35765952 | 1.04601884 | 0.34239825 | 0.51878522 | 0.3822628 | 1388 | tags=15%, list | PPARG/MGLL/NFIB/MTA1/PRKAR2B/CCN1/MAGI2/CITED2/KIT/KALRN/VLDLR/FBLN5/DLC1/APBB2/CAP2/DBP/PLPP3/DMAC2L/ARHGEF9/RXRA/HAS2/COL3A1 |
| HALLMARK_UV_RESPONSE_UP | HALLMARK_UV_RESPONSE_UP | 142 | 0.35263315 | 1.03158424 | 0.37639117 | 0.55351643 | 0.40785421 | 782 | tags=10%, list | RET/FOSB/CDKN1C/DGAT1/FOS/FMO1/COL2A1/GRINA/BTG1/OLFM1/BTG2/ATF3/TYRO3/LHX2 |
| HALLMARK_GLYCOLYSIS | HALLMARK_GLYCOLYSIS | 182 | -0.3178572 | -0.987477 | 0.47787611 | 0.66096691 | 0.48702825 | 2727 | tags=24%, list | CYB5A/SDC2/CHST6/AURKA/TXN/ADORA2B/IER3/ALDOA/B4GALT4/GNPDA1/ARPP19/TPBG/TALDO1/TPI1/GOT2/CHPF2/FUT8/POLR3K/LDHA/TGFA/PPP2CB/SRD5A3/IL13RA1/SOD1/HSPA5/CD44/SAP30/EGFR/SLC16A3/SDC1/PGK1/SPAG4/PPFIA4/FAM162A/DSC2/QSOX1/ENO1/ARTN/TFF3/MET/PKM/ENO2/PFKP/ELF3 |
| HALLMARK_UNFOLDED_PROTEIN_RESPONSE | HALLMARK_UNFOLDED_PROTEIN_RESPONSE | 111 | -0.3374496 | -0.9837659 | 0.47764757 | 0.66096691 | 0.48702825 | 2527 | tags=19%, list | DNAJC3/SPCS3/GOSR2/YIF1A/SLC1A4/SDAD1/SERP1/NFYA/XPOT/XBP1/SPCS1/HSPA5/BAG3/CHAC1/DNAJA4/EIF2S1/CNOT6/POP4/TARS1/SLC7A5/IMP3 |
| HALLMARK_APICAL_SURFACE | HALLMARK_APICAL_SURFACE | 36 | 0.41032509 | 0.97592398 | 0.48911551 | 0.66096691 | 0.48702825 | 2133 | tags=33%, list | NTNG1/SLC2A4/EPHB4/CD160/SRPX/THY1/ADIPOR2/AKAP7/CX3CL1/MDGA1/GAS1/FLOT2 |
| HALLMARK_KRAS_SIGNALING_UP | HALLMARK_KRAS_SIGNALING_UP | 165 | 0.32792108 | 0.97544333 | 0.5088352 | 0.66952 | 0.49333052 | 2382 | tags=25%, list | RBP4/G0S2/IL7R/PEG3/SPRY2/SPON1/TNFAIP3/CBX8/DUSP6/SPARCL1/ANGPTL4/GABRA3/PECAM1/CIDEA/PRDM1/ANO1/SEMA3B/TRIB2/FLT4/TSPAN7/ENG/ABCB1/PCSK1N/SNAP25/ATG10/CBR4/KIF5C/GUCY1A1/RGS16/CAB39L/ACE/MMD/NRP1/TRIB1/ALDH1A2/AKAP12/PTGS2/CPE/CSF2RA/CD37/MAP3K1 |
| HALLMARK_SPERMATOGENESIS | HALLMARK_SPERMATOGENESIS | 77 | -0.3437978 | -0.950867 | 0.55209657 | 0.70781611 | 0.52154872 | 1700 | tags=19%, list | TSN/AURKA/AGFG1/CCNB2/TALDO1/CCT6B/COIL/BUB1/TNNI3/NPHP1/HSPA4L/PCSK4/ARL4A/CLGN/MAP7 |
| HALLMARK_XENOBIOTIC_METABOLISM | HALLMARK_XENOBIOTIC_METABOLISM | 153 | -0.3118696 | -0.9485423 | 0.58309038 | 0.72886297 | 0.53705693 | 2293 | tags=24%, list | ARPP19/PTGES/GSR/CYP2J2/COMT/ADH1C/GSS/MT2A/NMT1/AKR1C2/CBR1/NINJ1/SSR3/BCAT1/PROS1/SMOX/SLC35B1/ATP2A2/PTGES3/SLC46A3/TDO2/NFS1/SERTAD1/PTS/PDLIM5/SLC1A5/NPC1/UGDH/GSTO1/NQO1/HES6/CFB/TGFB2/KYNU/GCNT2/SPINT2/ALDH3A1 |
| HALLMARK_HYPOXIA | HALLMARK_HYPOXIA | 180 | -0.2925986 | -0.9081761 | 0.72850987 | 0.82785212 | 0.6099963 | 1357 | tags=14%, list | HSPA5/GAPDH/SAP30/EGFR/TES/PGK1/JMJD6/PPFIA4/FAM162A/NEDD4L/ENO1/CP/MT1E/PRKCA/F3/SIAH2/HAS1/TMEM45A/GCNT2/ENO2/PFKP/NDRG1/ANXA2/SDC4/SLC2A1 |
| HALLMARK_ALLOGRAFT_REJECTION | HALLMARK_ALLOGRAFT_REJECTION | 149 | 0.3045297 | 0.89624709 | 0.70793247 | 0.82785212 | 0.6099963 | 1976 | tags=15%, list | CD79A/CCL19/ZAP70/CD96/LTB/IRF4/RPL39/IRF7/LCK/CD8A/DYRK3/CD7/INHBB/THY1/IL16/ACHE/IL12RB1/PRF1/CFP/STAB1/ITGAL/CD3D |
| HALLMARK_IL6_JAK_STAT3_SIGNALING | HALLMARK_IL6_JAK_STAT3_SIGNALING | 71 | 0.32763892 | 0.87436862 | 0.70722505 | 0.82785212 | 0.6099963 | 516 | tags=6%, list= | CNTFR/LEPR/CD36/LTB |
| HALLMARK_ANGIOGENESIS | HALLMARK_ANGIOGENESIS | 28 | -0.3621349 | -0.8322086 | 0.72648714 | 0.82785212 | 0.6099963 | 1257 | tags=18%, list | SLCO2A1/ITGAV/TNFRSF21/JAG1/SPP1 |
| HALLMARK_WNT_BETA_CATENIN_SIGNALING | HALLMARK_WNT_BETA_CATENIN_SIGNALING | 39 | 0.33704059 | 0.81463998 | 0.77680312 | 0.86311458 | 0.63597916 | 2139 | tags=18%, list | FZD8/KAT2A/NOTCH4/DLL1/PTCH1/WNT5B/GNAI1 |
| HALLMARK_TGF_BETA_SIGNALING | HALLMARK_TGF_BETA_SIGNALING | 51 | 0.28984903 | 0.73429632 | 0.92133971 | 0.98084977 | 0.72273141 | 479 | tags=4%, list= | CDKN1C/SPTBN1 |
| HALLMARK_NOTCH_SIGNALING | HALLMARK_NOTCH_SIGNALING | 32 | -0.2982403 | -0.7029774 | 0.92199878 | 0.98084977 | 0.72273141 | 3437 | tags=38%, list | WNT5A/RBX1/FZD5/HES1/FBXW11/CCND1/PSEN2/APH1A/SAP30/JAG1/PRKCA/PSENEN |
| HALLMARK_INTERFERON_GAMMA_RESPONSE | HALLMARK_INTERFERON_GAMMA_RESPONSE | 177 | -0.2597422 | -0.8045028 | 0.96549376 | 0.99906033 | 0.73614971 | 3090 | tags=20%, list | IRF1/CFH/CD38/GCH1/OGFR/PTPN1/FAS/TRIM26/PSMB8/PLSCR1/TAP1/B2M/BATF2/TNFSF10/PSMB2/SRI/TDRD7/BPGM/SPPL2A/CDKN1A/MT2A/IDO1/MVP/IL4R/CASP4/NMI/IRF2/CASP3/TRIM14/CD40/IL7/ZNFX1/ARL4A/CFB/PFKP/VAMP8 |
| HALLMARK_OXIDATIVE_PHOSPHORYLATION | HALLMARK_OXIDATIVE_PHOSPHORYLATION | 199 | -0.2439522 | -0.7654421 | 0.99614163 | 0.99906033 | 0.73614971 | 3259 | tags=23%, list | ACAA1/COX17/NDUFA5/FDX1/NDUFS3/COX5B/SDHA/GPI/ATP5PD/TIMM10/ATP6V1G1/ATP6V0B/IMMT/NDUFA1/CYB5A/COX6C/MFN2/NDUFC1/COX7A2L/COX7A2/NDUFA6/GOT2/ATP5MG/NDUFA9/NDUFB2/UQCRQ/ATP6V1D/NDUFC2/LDHA/PDP1/ATP6V1F/ATP6V1E1/ATP5PB/NDUFS2/HCCS/ATP5PF/MRPS30/UQCR11/TCIRG1/ATP6V0E1/ISCU/COX5A/MRPL34/ATP1B1/ATP5F1E |
| HALLMARK_INTERFERON_ALPHA_RESPONSE | HALLMARK_INTERFERON_ALPHA_RESPONSE | 91 | 0.2284294 | 0.63055826 | 0.99906033 | 0.99906033 | 0.73614971 | 1964 | tags=12%, list | IRF7/OASL/IFI44L/LY6E/SELL/IFITM2/TRIM5/IFI35/LGALS3BP/IFITM3/TENT5A |
