## Supplementary material for "Integrated transcriptomic analysis identifies Paternally Expressed Gene 10 as a novel candidate biomarker associated with aggressive features in liposarcoma": Table S4_GOBP

| ID | Description | setSize | enrichmentS | NES | pvalue | p.adjust | qvalue | rank | leading_edg | core_enrichment |
|---|---|---|---|---|---|---|---|---|---|---|
| GOBP_CILIUM_MOVEMENT | GOBP_CILIUM_MOVEMENT | 145 | -0.7527455 | -2.2740545 | 1E-10 | 1.2518E-07 | 1.1598E-07 | 1708 | tags=46%, list | [illegible] |
| GOBP_CILIUM_OR_FLAGELLUM_DEPENDENT_CELL_MOTILITY | GOBP_CILIUM_OR_FLAGELLUM_DEPENDENT_CELL_MOTILITY | 122 | -0.752237 | -2.2205809 | 1E-10 | 1.2518E-07 | 1.1598E-07 | 1075 | tags=36%, list | [illegible] |
| GOBP_MICROTUBULE_BASED_MOVEMENT | GOBP_MICROTUBULE_BASED_MOVEMENT | 363 | -0.6592637 | -2.1916643 | 1E-10 | 1.2518E-07 | 1.1598E-07 | 1837 | tags=33%, list | [illegible] |
| GOBP_CILIUM_ORGANIZATION | GOBP_CILIUM_ORGANIZATION | 380 | -0.6299657 | -2.1059414 | 1E-10 | 1.2518E-07 | 1.1598E-07 | 1840 | tags=34%, list | [illegible] |
| GOBP_CELL_PROJECTION_ASSEMBLY | GOBP_CELL_PROJECTION_ASSEMBLY | 547 | -0.5623486 | -1.9288346 | 1E-10 | 1.2518E-07 | 1.1598E-07 | 2616 | tags=30%, list | [illegible] |
| GOBP_MICROTUBULE_BASED_PROCESS | GOBP_MICROTUBULE_BASED_PROCESS | 834 | -0.5193704 | -1.8310559 | 1E-10 | 1.2518E-07 | 1.1598E-07 | 2542 | tags=29%, list | [illegible] |
| GOBP_ORGANELLE_ASSEMBLY | GOBP_ORGANELLE_ASSEMBLY | 958 | -0.4407651 | -1.5661308 | 2.9573E-10 | 3.1732E-07 | 2.94E-07 | 2768 | tags=26%, list | [illegible] |
| GOBP_AXONEME_ASSEMBLY | GOBP_AXONEME_ASSEMBLY | 57 | -0.826004 | -2.178599 | 6.8811E-10 | 6.4605E-07 | 5.9856E-07 | 1125 | tags=58%, list | [illegible] |
| GOBP_MICROTUBULE_BASED_TRANSPORT | GOBP_MICROTUBULE_BASED_TRANSPORT | 213 | -0.6146522 | -1.9445072 | 2.3582E-09 | 1.9681E-06 | 1.8234E-06 | 2156 | tags=35%, list | [illegible] |
| GOBP_MOTILE_CILIUM_ASSEMBLY | GOBP_MOTILE_CILIUM_ASSEMBLY | 61 | -0.79794 | -2.1275048 | 2.9752E-09 | 2.2347E-06 | 2.0705E-06 | 1497 | tags=44%, list | [illegible] |
| GOBP_MICROTUBULE_BUNDLE_FORMATION | GOBP_MICROTUBULE_BUNDLE_FORMATION | 85 | -0.7456643 | -2.0932433 | 6.3446E-09 | 4.3322E-06 | 4.0138E-06 | 1125 | tags=41%, list | [illegible] |
| GOBP_MICROTUBULE_CYTOSKELETON_ORGANIZATION | GOBP_MICROTUBULE_CYTOSKELETON_ORGANIZATION | 578 | -0.4699766 | -1.6182394 | 2.7217E-08 | 1.7036E-05 | 1.5784E-05 | 2542 | tags=26%, list | [illegible] |
| GOBP_CYTOSKELETON_ORGANIZATION | GOBP_CYTOSKELETON_ORGANIZATION | 1276 | -0.3976322 | -1.4363953 | 4.3554E-08 | 2.5164E-05 | 2.3314E-05 | 1747 | tags=17%, list | [illegible] |
| GOBP_EPIDERMIS_DEVELOPMENT | GOBP_EPIDERMIS_DEVELOPMENT | 233 | -0.5474287 | -1.7493244 | 4.6816E-07 | 0.00025117 | 0.00023271 | 963 | tags=18%, list | [illegible] |
| GOBP_RESPONSE_TO_ENDOGENOUS_STIMULUS | GOBP_RESPONSE_TO_ENDOGENOUS_STIMULUS | 1201 | 0.40924173 | 1.42316207 | 5.0924E-07 | 0.000255 | 0.00023625 | 1561 | tags=16%, list | [illegible] |
| GOBP_EXTRACELLULAR_TRANSPORT | GOBP_EXTRACELLULAR_TRANSPORT | 36 | -0.8230319 | -2.0031457 | 9.1891E-07 | 0.00040691 | 0.000377 | 1125 | tags=58%, list | [illegible] |
| GOBP_REGULATION_OF_LIPID_METABOLIC_PROCESS | GOBP_REGULATION_OF_LIPID_METABOLIC_PROCESS | 261 | 0.55430982 | 1.73979138 | 1.0157E-06 | 0.00040691 | 0.000377 | 1289 | tags=17%, list | [illegible] |
| GOBP_ENZYME_LINKED_RECEPTOR_PROTEIN_SIGNALING_PATHWAY | GOBP_ENZYME_LINKED_RECEPTOR_PROTEIN_SIGNALING_PATHWAY | 830 | 0.43335961 | 1.47880519 | 9.3374E-07 | 0.00040691 | 0.000377 | 1870 | tags=19%, list | [illegible] |
| GOBP_CELLULAR_RESPONSE_TO_ENDOGENOUS_STIMULUS | GOBP_CELLULAR_RESPONSE_TO_ENDOGENOUS_STIMULUS | 1014 | 0.41252296 | 1.42315993 | 1.0293E-06 | 0.00040691 | 0.000377 | 1561 | tags=15%, list | [illegible] |
| GOBP_SYNAPTIC_SIGNALING | GOBP_SYNAPTIC_SIGNALING | 557 | 0.46115746 | 1.53511303 | 1.1409E-06 | 0.00042845 | 0.00039696 | 1445 | tags=17%, list | [illegible] |
| GOBP_NEUTRAL_LIPID_METABOLIC_PROCESS | GOBP_NEUTRAL_LIPID_METABOLIC_PROCESS | 101 | 0.67399028 | 1.89305548 | 1.677E-06 | 0.00059979 | 0.00055571 | 1316 | tags=25%, list | [illegible] |
| GOBP_PROTEIN_LOCALIZATION_TO_CILIUM | GOBP_PROTEIN_LOCALIZATION_TO_CILIUM | 77 | -0.7039087 | -1.9490197 | 1.7922E-06 | 0.00061187 | 0.00056689 | 2156 | tags=35%, list | [illegible] |
| GOBP_KERATINOCYTE_DIFFERENTIATION | GOBP_KERATINOCYTE_DIFFERENTIATION | 85 | -0.6919263 | -1.9423889 | 2.2944E-06 | 0.00074927 | 0.0006942 | 756 | tags=25%, list | [illegible] |
| GOBP_SKIN_DEVELOPMENT | GOBP_SKIN_DEVELOPMENT | 201 | -0.5616622 | -1.7643436 | 2.5495E-06 | 0.00079787 | 0.00073923 | 756 | tags=18%, list | [illegible] |
| GOBP_IMMUNE_RESPONSE_REGULATING_CELL_SURFACE_RECEPTOR_S | GOBP_IMMUNE_RESPONSE_REGULATING_CELL_SURFACE_RECEPTOR_S | 289 | 0.53157287 | 1.68309987 | 3.3835E-06 | 0.00101654 | 0.00094183 | 1549 | tags=18%, list | [illegible] |
| GOBP_EPIDERMAL_CELL_DIFFERENTIATION | GOBP_EPIDERMAL_CELL_DIFFERENTIATION | 138 | -0.604657 | -1.8135305 | 3.5512E-06 | 0.0010259 | 0.0009505 | 756 | tags=19%, list | [illegible] |
| GOBP_SPERM_FLAGELLUM_ASSEMBLY | GOBP_SPERM_FLAGELLUM_ASSEMBLY | 38 | -0.8061024 | -1.9801951 | 4.1697E-06 | 0.00115211 | 0.00106743 | 1450 | tags=45%, list | [illegible] |
| GOBP_TRIGLYCERIDE_METABOLIC_PROCESS | GOBP_TRIGLYCERIDE_METABOLIC_PROCESS | 81 | 0.70045284 | 1.90812825 | 4.2949E-06 | 0.00115211 | 0.00106743 | 708 | tags=22%, list | [illegible] |
| GOBP_CELL_CELL_SIGNALING | GOBP_CELL_CELL_SIGNALING | 885 | 0.42006684 | 1.43924098 | 5.4366E-06 | 0.00140807 | 0.00130458 | 1476 | tags=16%, list | [illegible] |
| GOBP_B_CELL_RECEPTOR_SIGNALING_PATHWAY | GOBP_B_CELL_RECEPTOR_SIGNALING_PATHWAY | 54 | 0.76229199 | 1.9379885 | 6.8039E-06 | 0.00170348 | 0.00157828 | 982 | tags=22%, list | [illegible] |
| GOBP_AXONEMAL_DYNEIN_COMPLEX_ASSEMBLY | GOBP_AXONEMAL_DYNEIN_COMPLEX_ASSEMBLY | 18 | -0.8921486 | -1.8792238 | 9.4996E-06 | 0.00205788 | 0.00190663 | 1059 | tags=67%, list | [illegible] |
| GOBP_RESPONSE_TO_PURINE_CONTAINING_COMPOUND | GOBP_RESPONSE_TO_PURINE_CONTAINING_COMPOUND | 111 | 0.65465365 | 1.85924977 | 9.2086E-06 | 0.00205788 | 0.00190663 | 1539 | tags=25%, list | [illegible] |
| GOBP_ADENYLATE_CYCLASE_MODULATING_G_PROTEIN_COUPLED_RE | GOBP_ADENYLATE_CYCLASE_MODULATING_G_PROTEIN_COUPLED_RE | 148 | 0.60941294 | 1.79160732 | 9.2491E-06 | 0.00205788 | 0.00190663 | 1452 | tags=27%, list | [illegible] |
| GOBP_SPERMATID_DIFFERENTIATION | GOBP_SPERMATID_DIFFERENTIATION | 138 | -0.5916 | -1.7743692 | 9.5894E-06 | 0.00205788 | 0.00190663 | 1688 | tags=27%, list | [illegible] |
| GOBP_VASCULATURE_DEVELOPMENT | GOBP_VASCULATURE_DEVELOPMENT | 660 | 0.43919139 | 1.47894242 | 9.2513E-06 | 0.00205788 | 0.00190663 | 1834 | tags=21%, list | [illegible] |
| GOBP_CELL_MOTILITY | GOBP_CELL_MOTILITY | 1389 | -0.3589501 | -1.301679 | 1.0067E-05 | 0.00210044 | 0.00194606 | 1708 | tags=16%, list | [illegible] |
| GOBP_INTRACILIARY_TRANSPORT | GOBP_INTRACILIARY_TRANSPORT | 51 | -0.7360218 | -1.9074492 | 1.2049E-05 | 0.002446 | 0.00226622 | 2454 | tags=51%, list | [illegible] |
| GOBP_NEGATIVE_REGULATION_OF_LIPID_METABOLIC_PROCESS | GOBP_NEGATIVE_REGULATION_OF_LIPID_METABOLIC_PROCESS | 76 | 0.70436075 | 1.90053391 | 1.4235E-05 | 0.00271386 | 0.00251439 | 708 | tags=21%, list | [illegible] |
| GOBP_REGULATION_OF_KETONE_METABOLIC_PROCESS | GOBP_REGULATION_OF_KETONE_METABOLIC_PROCESS | 105 | 0.65162778 | 1.83750023 | 1.4453E-05 | 0.00271386 | 0.00251439 | 643 | tags=16%, list | [illegible] |
| GOBP_INTRACELLULAR_TRANSPORT | GOBP_INTRACELLULAR_TRANSPORT | 1303 | -0.3613932 | -1.3070667 | 1.3943E-05 | 0.00271386 | 0.00251439 | 3424 | tags=29%, list | [illegible] |
| GOBP_ANTIGEN_RECEPTOR_MEDIATED_SIGNALING_PATHWAY | GOBP_ANTIGEN_RECEPTOR_MEDIATED_SIGNALING_PATHWAY | 164 | 0.58676372 | 1.74703352 | 1.6435E-05 | 0.00294345 | 0.00272711 | 1454 | tags=16%, list | [illegible] |
| GOBP_FATTY_ACID_METABOLIC_PROCESS | GOBP_FATTY_ACID_METABOLIC_PROCESS | 310 | 0.50237006 | 1.60284071 | 1.6459E-05 | 0.00294345 | 0.00272711 | 1895 | tags=20%, list | [illegible] |
| GOBP_LIPID_CATABOLIC_PROCESS | GOBP_LIPID_CATABOLIC_PROCESS | 245 | 0.53226521 | 1.6593218 | 1.708E-05 | 0.00298352 | 0.00276423 | 1577 | tags=22%, list | [illegible] |
| GOBP_REGULATION_OF_FATTY_ACID_METABOLIC_PROCESS | GOBP_REGULATION_OF_FATTY_ACID_METABOLIC_PROCESS | 83 | 0.69008674 | 1.88847672 | 1.9158E-05 | 0.00327034 | 0.00302998 | 643 | tags=18%, list | [illegible] |
| GOBP_RESPONSE_TO_INSULIN | GOBP_RESPONSE_TO_INSULIN | 235 | 0.532204 | 1.65140788 | 2.0079E-05 | 0.00335145 | 0.00310512 | 1245 | tags=15%, list | [illegible] |
| GOBP_NEGATIVE_REGULATION_OF_CELLULAR_RESPONSE_TO_VASCUL | GOBP_NEGATIVE_REGULATION_OF_CELLULAR_RESPONSE_TO_VASCUL | 12 | 0.95072229 | 1.81864292 | 2.165E-05 | 0.003535 | 0.00327518 | 329 | tags=25%, list | [illegible] |
| GOBP_NEGATIVE_REGULATION_OF_VASCULATURE_DEVELOPMENT | GOBP_NEGATIVE_REGULATION_OF_VASCULATURE_DEVELOPMENT | 90 | 0.66439625 | 1.83618675 | 2.5794E-05 | 0.0040363 | 0.00373963 | 1154 | tags=22%, list | [illegible] |
| GOBP_RESPONSE_TO_ORGANOPHOSPHORUS | GOBP_RESPONSE_TO_ORGANOPHOSPHORUS | 93 | 0.65825596 | 1.82699393 | 2.577E-05 | 0.0040363 | 0.00373963 | 1539 | tags=27%, list | [illegible] |
| GOBP_REGULATION_OF_LIPID_CATABOLIC_PROCESS | GOBP_REGULATION_OF_LIPID_CATABOLIC_PROCESS | 48 | 0.76626079 | 1.9109411 | 2.6773E-05 | 0.00410392 | 0.00380228 | 708 | tags=27%, list | [illegible] |
| GOBP_POSITIVE_REGULATION_OF_LIPID_METABOLIC_PROCESS | GOBP_POSITIVE_REGULATION_OF_LIPID_METABOLIC_PROCESS | 104 | 0.64051826 | 1.80434174 | 3.2675E-05 | 0.0048122 | 0.00445851 | 1612 | tags=25%, list | [illegible] |
| GOBP_CIRCULATORY_SYSTEM_DEVELOPMENT | GOBP_CIRCULATORY_SYSTEM_DEVELOPMENT | 984 | 0.39445266 | 1.35903341 | 3.2444E-05 | 0.0048122 | 0.00445851 | 1834 | tags=19%, list | [illegible] |
| GOBP_LIPID_OXIDATION | GOBP_LIPID_OXIDATION | 100 | 0.64484443 | 1.8087952 | 3.5105E-05 | 0.00488285 | 0.00452396 | 1832 | tags=30%, list | [illegible] |
| GOBP_TEMPERATURE_HOMEOSTASIS | GOBP_TEMPERATURE_HOMEOSTASIS | 153 | 0.58241303 | 1.72067958 | 3.4973E-05 | 0.00488285 | 0.00452396 | 1996 | tags=27%, list | [illegible] |
| GOBP_CELL_MORPHOGENESIS_INVOLVED_IN_NEURON_DIFFERENTIATI | GOBP_CELL_MORPHOGENESIS_INVOLVED_IN_NEURON_DIFFERENTIAT | 481 | 0.44776534 | 1.47702199 | 3.5032E-05 | 0.00488285 | 0.00452396 | 1938 | tags=22%, list | [illegible] |
| GOBP_POSITIVE_REGULATION_OF_MULTICELLULAR_ORGANISMAL_PR | GOBP_POSITIVE_REGULATION_OF_MULTICELLULAR_ORGANISMAL_PR | 1292 | 0.38000678 | 1.32684218 | 3.5762E-05 | 0.0048838 | 0.00452485 | 1576 | tags=15%, list | [illegible] |
| GOBP_DIACYLGLYCEROL_METABOLIC_PROCESS | GOBP_DIACYLGLYCEROL_METABOLIC_PROCESS | 19 | 0.87914595 | 1.84758753 | 3.8679E-05 | 0.00509682 | 0.0047222 | 116 | tags=21%, list | [illegible] |
| GOBP_REGULATION_OF_TRANS_SYNAPTIC_SIGNALING | GOBP_REGULATION_OF_TRANS_SYNAPTIC_SIGNALING | 366 | 0.47470627 | 1.53423679 | 3.8378E-05 | 0.00509682 | 0.0047222 | 1818 | tags=21%, list | [illegible] |
| GOBP_CARBOHYDRATE_BIOSYNTHETIC_PROCESS | GOBP_CARBOHYDRATE_BIOSYNTHETIC_PROCESS | 149 | 0.58242985 | 1.71296502 | 3.9854E-05 | 0.00516103 | 0.0047817 | 1320 | tags=19%, list | [illegible] |
| GOBP_BLOOD_VESSEL_MORPHOGENESIS | GOBP_BLOOD_VESSEL_MORPHOGENESIS | 546 | 0.43794172 | 1.45685505 | 5.1113E-05 | 0.00650693 | 0.00602867 | 1935 | tags=23%, list | [illegible] |
| GOBP_SKELETAL_SYSTEM_DEVELOPMENT | GOBP_SKELETAL_SYSTEM_DEVELOPMENT | 445 | 0.45673667 | 1.49906722 | 6.2525E-05 | 0.00782714 | 0.00725185 | 1928 | tags=20%, list | [illegible] |
| GOBP_RESPONSE_TO_CAMP | GOBP_RESPONSE_TO_CAMP | 63 | 0.6996682 | 1.831345 | 7.554E-05 | 0.00930128 | 0.00861764 | 1539 | tags=32%, list | [illegible] |
| GOBP_TUBE_DEVELOPMENT | GOBP_TUBE_DEVELOPMENT | 921 | 0.39741159 | 1.36385814 | 7.9065E-05 | 0.00957831 | 0.00887431 | 1935 | tags=20%, list | [illegible] |
| GOBP_MONOSACCHARIDE_BIOSYNTHETIC_PROCESS | GOBP_MONOSACCHARIDE_BIOSYNTHETIC_PROCESS | 81 | 0.6551467 | 1.78470819 | 8.484E-05 | 0.00986831 | 0.009143 | 1151 | tags=21%, list | [illegible] |
| GOBP_NERVOUS_SYSTEM_PROCESS | GOBP_NERVOUS_SYSTEM_PROCESS | 697 | 0.41235851 | 1.39370387 | 8.54E-05 | 0.00986831 | 0.009143 | 1270 | tags=15%, list | [illegible] |
| GOBP_REGULATION_OF_MULTICELLULAR_ORGANISMAL_DEVELOPME | GOBP_REGULATION_OF_MULTICELLULAR_ORGANISMAL_DEVELOPME | 1142 | 0.38202924 | 1.32603694 | 8.3098E-05 | 0.00986831 | 0.009143 | 1818 | tags=17%, list | [illegible] |
| GOBP_RESPONSE_TO_PEPTIDE_HORMONE | GOBP_RESPONSE_TO_PEPTIDE_HORMONE | 370 | 0.47408572 | 1.5330637 | 9.3825E-05 | 0.01067759 | 0.0098928 | 1560 | tags=17%, list | [illegible] |
| GOBP_ESTABLISHMENT_OF_SKIN_BARRIER | GOBP_ESTABLISHMENT_OF_SKIN_BARRIER | 17 | -0.8810248 | -1.8377668 | 9.75E-05 | 0.01093023 | 0.01012687 | 479 | tags=47%, list | [illegible] |
| GOBP_POSITIVE_REGULATION_OF_SMALL_MOLECULE_METABOLIC_PR | GOBP_POSITIVE_REGULATION_OF_SMALL_MOLECULE_METABOLIC_PR | 124 | 0.5884498 | 1.69723273 | 9.9498E-05 | 0.01099019 | 0.01018242 | 996 | tags=16%, list | [illegible] |
| GOBP_POSITIVE_REGULATION_OF_FATTY_ACID_METABOLIC_PROCESS | GOBP_POSITIVE_REGULATION_OF_FATTY_ACID_METABOLIC_PROCESS | 32 | 0.80338541 | 1.86351618 | 0.00010828 | 0.01161882 | 0.01076485 | 643 | tags=22%, list | [illegible] |
| GOBP_LEARNING | GOBP_LEARNING | 112 | 0.61613113 | 1.75195442 | 0.00010794 | 0.01161882 | 0.01076485 | 1270 | tags=22%, list | [illegible] |
| GOBP_TUBE_MORPHOGENESIS | GOBP_TUBE_MORPHOGENESIS | 754 | 0.40866897 | 1.38774774 | 0.00011162 | 0.0116443 | 0.01078845 | 1834 | tags=21%, list | [illegible] |
| GOBP_RESPONSE_TO_OXYGEN_CONTAINING_COMPOUND | GOBP_RESPONSE_TO_OXYGEN_CONTAINING_COMPOUND | 1250 | 0.37296288 | 1.29951037 | 0.00011139 | 0.0116443 | 0.01078845 | 1561 | tags=13%, list | [illegible] |
| GOBP_RESPONSE_TO_HORMONE | GOBP_RESPONSE_TO_HORMONE | 720 | 0.40920756 | 1.38533554 | 0.00011937 | 0.01228202 | 0.0113793 | 1561 | tags=17%, list | [illegible] |
| GOBP_CELL_PROJECTION_MORPHOGENESIS | GOBP_CELL_PROJECTION_MORPHOGENESIS | 538 | 0.42817979 | 1.4241719 | 0.00012112 | 0.01229377 | 0.01139018 | 1938 | tags=21%, list | [illegible] |
| GOBP_NEGATIVE_REGULATION_OF_FATTY_ACID_METABOLIC_PROCESS | GOBP_NEGATIVE_REGULATION_OF_FATTY_ACID_METABOLIC_PROCESS | 31 | 0.79476821 | 1.83532318 | 0.00012535 | 0.01245075 | 0.01153563 | 514 | tags=23%, list | [illegible] |
| GOBP_RENAL_SYSTEM_DEVELOPMENT | GOBP_RENAL_SYSTEM_DEVELOPMENT | 272 | 0.49689639 | 1.56559397 | 0.00012598 | 0.01245075 | 0.01153563 | 1928 | tags=27%, list | [illegible] |
| GOBP_GLUCOSE_METABOLIC_PROCESS | GOBP_GLUCOSE_METABOLIC_PROCESS | 168 | 0.54707422 | 1.63369736 | 0.00013133 | 0.01271277 | 0.01177839 | 1151 | tags=16%, list | [illegible] |
| GOBP_CYTOSKELETON_DEPENDENT_INTRACELLULAR_TRANSPORT | GOBP_CYTOSKELETON_DEPENDENT_INTRACELLULAR_TRANSPORT | 202 | -0.5032881 | -1.5817217 | 0.00013202 | 0.01271277 | 0.01177839 | 2315 | tags=30%, list | [illegible] |
| GOBP_ADIPOSE_TISSUE_DEVELOPMENT | GOBP_ADIPOSE_TISSUE_DEVELOPMENT | 70 | 0.66734792 | 1.77416377 | 0.00015873 | 0.01509172 | 0.01398249 | 815 | tags=13%, list | [illegible] |
| GOBP_REGULATION_OF_GLUCOSE_METABOLIC_PROCESS | GOBP_REGULATION_OF_GLUCOSE_METABOLIC_PROCESS | 86 | 0.62757211 | 1.72463314 | 0.00016147 | 0.01515999 | 0.01404575 | 996 | tags=17%, list | [illegible] |
| GOBP_REGULATION_OF_MICROTUBULE_BASED_MOVEMENT | GOBP_REGULATION_OF_MICROTUBULE_BASED_MOVEMENT | 30 | -0.7831079 | -1.8347355 | 0.00017083 | 0.01584096 | 0.01467666 | 1785 | tags=37%, list | [illegible] |
| GOBP_TRIGLYCERIDE_CATABOLIC_PROCESS | GOBP_TRIGLYCERIDE_CATABOLIC_PROCESS | 21 | 0.84490682 | 1.81479913 | 0.00017539 | 0.01598523 | 0.01481032 | 946 | tags=48%, list | [illegible] |
| GOBP_SPERM_AXONEME_ASSEMBLY | GOBP_SPERM_AXONEME_ASSEMBLY | 21 | -0.8352024 | -1.8130842 | 0.00017877 | 0.01598523 | 0.01481032 | 1075 | tags=52%, list | [illegible] |
| GOBP_ORGANIC_ACID_CATABOLIC_PROCESS | GOBP_ORGANIC_ACID_CATABOLIC_PROCESS | 187 | 0.53588794 | 1.62125188 | 0.00017726 | 0.01598523 | 0.01481032 | 1868 | tags=26%, list | [illegible] |
| GOBP_REGULATION_OF_CAMP_PKA_SIGNAL_TRANSDUCTION | GOBP_REGULATION_OF_CAMP_PKA_SIGNAL_TRANSDUCTION | 40 | 0.75661373 | 1.83104648 | 0.00019913 | 0.0174441 | 0.01616198 | 1657 | tags=40%, list | [illegible] |
| GOBP_INTRINSIC_APOPTOTIC_SIGNALING_PATHWAY | GOBP_INTRINSIC_APOPTOTIC_SIGNALING_PATHWAY | 304 | -0.455155 | -1.4878503 | 0.00020083 | 0.0174441 | 0.01616198 | 1844 | tags=18%, list | [illegible] |
| GOBP_POSITIVE_REGULATION_OF_DEVELOPMENTAL_PROCESS | GOBP_POSITIVE_REGULATION_OF_DEVELOPMENTAL_PROCESS | 1049 | 0.38298634 | 1.32429948 | 0.00020206 | 0.0174441 | 0.01616198 | 1822 | tags=18%, list | [illegible] |
| GOBP_ADAPTIVE_THERMOGENESIS | GOBP_ADAPTIVE_THERMOGENESIS | 145 | 0.55777753 | 1.6365354 | 0.00020469 | 0.01747097 | 0.01618686 | 1996 | tags=25%, list | [illegible] |
| GOBP_ACYLGLYCEROL_HOMEOSTASIS | GOBP_ACYLGLYCEROL_HOMEOSTASIS | 30 | 0.78895184 | 1.8137253 | 0.000212 | 0.01789175 | 0.01657672 | 643 | tags=30%, list | [illegible] |
| GOBP_MALE_GAMETE_GENERATION | GOBP_MALE_GAMETE_GENERATION | 449 | -0.4153133 | -1.4049735 | 0.00022151 | 0.01848638 | 0.01712765 | 1725 | tags=18%, list | [illegible] |
| GOBP_NEGATIVE_REGULATION_OF_LIPID_CATABOLIC_PROCESS | GOBP_NEGATIVE_REGULATION_OF_LIPID_CATABOLIC_PROCESS | 20 | 0.84302853 | 1.78945758 | 0.0002285 | 0.01886001 | 0.01747381 | 708 | tags=45%, list | [illegible] |
| GOBP_REPRODUCTIVE_PROCESS | GOBP_REPRODUCTIVE_PROCESS | 1083 | -0.3581837 | -1.281674 | 0.00024504 | 0.02000552 | 0.01853513 | 1844 | tags=17%, list | [illegible] |
| GOBP_ORGANIC_ACID_METABOLIC_PROCESS | GOBP_ORGANIC_ACID_METABOLIC_PROCESS | 699 | 0.40405157 | 1.3656203 | 0.00025118 | 0.02028629 | 0.01879526 | 1872 | tags=18%, list | [illegible] |
| GOBP_NEUTRAL_LIPID_CATABOLIC_PROCESS | GOBP_NEUTRAL_LIPID_CATABOLIC_PROCESS | 29 | 0.80648569 | 1.84420804 | 0.00026475 | 0.02115467 | 0.01959982 | 1218 | tags=41%, list | [illegible] |
| GOBP_BEHAVIOR | GOBP_BEHAVIOR | 475 | 0.44210816 | 1.45650844 | 0.00027026 | 0.02136725 | 0.01979677 | 1911 | tags=20%, list | [illegible] |
| GOBP_NEGATIVE_REGULATION_OF_SMALL_MOLECULE_METABOLIC_PI | GOBP_NEGATIVE_REGULATION_OF_SMALL_MOLECULE_METABOLIC_PI | 82 | 0.6269746 | 1.71148492 | 0.00030814 | 0.02410889 | 0.02233691 | 514 | tags=12%, list | [illegible] |
| GOBP_CELL_SURFACE_RECEPTOR_PROTEIN_TYROSINE_KINASE_SIGNAL | GOBP_CELL_SURFACE_RECEPTOR_PROTEIN_TYROSINE_KINASE_SIGNAL | 541 | 0.42059439 | 1.39903808 | 0.00033608 | 0.02602338 | 0.02411068 | 1643 | tags=17%, list | [illegible] |
| GOBP_CELLULAR_RESPONSE_TO_CAMP | GOBP_CELLULAR_RESPONSE_TO_CAMP | 41 | 0.74593785 | 1.8117212 | 0.00036953 | 0.02832213 | 0.02624047 | 1460 | tags=27%, list | [illegible] |
| GOBP_CEREBELLAR_GRANULAR_LAYER_MORPHOGENESIS | GOBP_CEREBELLAR_GRANULAR_LAYER_MORPHOGENESIS | 6 | 0.9745039 | 1.62504943 | 0.00037878 | 0.02873744 | 0.02662526 | 169 | tags=50%, list | [illegible] |
| GOBP_REGULATION_OF_CELL_DEVELOPMENT | GOBP_REGULATION_OF_CELL_DEVELOPMENT | 660 | 0.40638755 | 1.36847804 | 0.00038344 | 0.02879997 | 0.0266832 | 1818 | tags=17%, list | [illegible] |
| GOBP_NEGATIVE_REGULATION_OF_GLUCONEOGENESIS | GOBP_NEGATIVE_REGULATION_OF_GLUCONEOGENESIS | 12 | 0.91342727 | 1.74730102 | 0.00039023 | 0.02902003 | 0.02688708 | 303 | tags=25%, list | [illegible] |
| GOBP_RESPONSE_TO_GROWTH_FACTOR | GOBP_RESPONSE_TO_GROWTH_FACTOR | 596 | 0.41747753 | 1.39690848 | 0.00039575 | 0.02914201 | 0.0270001 | 1867 | tags=18%, list | [illegible] |
| GOBP_HORMONE_MEDIATED_SIGNALING_PATHWAY | GOBP_HORMONE_MEDIATED_SIGNALING_PATHWAY | 179 | 0.52045904 | 1.56566281 | 0.00040286 | 0.02937779 | 0.02721855 | 2007 | tags=22%, list | [illegible] |
| GOBP_FATTY_ACID_BETA_OXIDATION | GOBP_FATTY_ACID_BETA_OXIDATION | 68 | 0.66054517 | 1.74861717 | 0.00043029 | 0.03034313 | 0.02811293 | 1832 | tags=35%, list | [illegible] |
| GOBP_RESPONSE_TO_LITHIUM_ION | GOBP_RESPONSE_TO_LITHIUM_ION | 10 | 0.92946659 | 1.70938993 | 0.00042139 | 0.03034313 | 0.02811293 | 140 | tags=20%, list | [illegible] |
| GOBP_REGULATION_OF_SYNAPSE_STRUCTURE_OR_ACTIVITY | GOBP_REGULATION_OF_SYNAPSE_STRUCTURE_OR_ACTIVITY | 265 | 0.48212676 | 1.51437708 | 0.00042537 | 0.03034313 | 0.02811293 | 1760 | tags=20%, list | [illegible] |
| GOBP_AXON_DEVELOPMENT | GOBP_AXON_DEVELOPMENT | 413 | 0.43872979 | 1.43058708 | 0.00044438 | 0.03034313 | 0.02811293 | 1938 | tags=22%, list | [illegible] |
| GOBP_CIRCULATORY_SYSTEM_PROCESS | GOBP_CIRCULATORY_SYSTEM_PROCESS | 456 | 0.43059522 | 1.41491334 | 0.00043939 | 0.03034313 | 0.02811293 | 1317 | tags=17%, list | [illegible] |
| GOBP_EPITHELIAL_CELL_DIFFERENTIATION | GOBP_EPITHELIAL_CELL_DIFFERENTIATION | 487 | -0.4048702 | -1.377468 | 0.00043286 | 0.03034313 | 0.02811293 | 1029 | tags=14%, list | [illegible] |
| GOBP_CHEMICAL_HOMEOSTASIS | GOBP_CHEMICAL_HOMEOSTASIS | 780 | 0.39083232 | 1.32930699 | 0.00044236 | 0.03034313 | 0.02811293 | 1358 | tags=14%, list | [illegible] |
| GOBP_DETECTION_OF_CHEMICAL_STIMULUS | GOBP_DETECTION_OF_CHEMICAL_STIMULUS | 36 | 0.74303486 | 1.76176201 | 0.00046577 | 0.03090213 | 0.02863084 | 1204 | tags=31%, list | [illegible] |
| GOBP_NEGATIVE_REGULATION_OF_VASCULAR_PERMEABILITY | GOBP_NEGATIVE_REGULATION_OF_VASCULAR_PERMEABILITY | 17 | 0.84585951 | 1.73577552 | 0.00046902 | 0.03090213 | 0.02863084 | 507 | tags=35%, list | [illegible] |
| GOBP_REGULATION_OF_POSTSYNAPTIC_MEMBRANE_POTENTIAL | GOBP_REGULATION_OF_POSTSYNAPTIC_MEMBRANE_POTENTIAL | 79 | 0.63040708 | 1.71456639 | 0.00046314 | 0.03090213 | 0.02863084 | 1042 | tags=24%, list | [illegible] |
| GOBP_RESPONSE_TO_NITROGEN_COMPOUND | GOBP_RESPONSE_TO_NITROGEN_COMPOUND | 851 | 0.38990855 | 1.33195806 | 0.0004635 | 0.03090213 | 0.02863084 | 1911 | tags=16%, list | [illegible] |
| GOBP_NEGATIVE_REGULATION_OF_SYNAPTIC_TRANSMISSION | GOBP_NEGATIVE_REGULATION_OF_SYNAPTIC_TRANSMISSION | 40 | 0.74020658 | 1.79134028 | 0.00049671 | 0.03177085 | 0.02943571 | 1270 | tags=30%, list | [illegible] |
| GOBP_REGULATION_OF_FATTY_ACID_OXIDATION | GOBP_REGULATION_OF_FATTY_ACID_OXIDATION | 33 | 0.76293012 | 1.78071151 | 0.00051182 | 0.03177085 | 0.02943571 | 580 | tags=27%, list | [illegible] |
| GOBP_PROSTATE_GLAND_DEVELOPMENT | GOBP_PROSTATE_GLAND_DEVELOPMENT | 39 | 0.72144771 | 1.73719864 | 0.00050618 | 0.03177085 | 0.02943571 | 2447 | tags=46%, list | [illegible] |
| GOBP_NEGATIVE_REGULATION_OF_SPROUTING_ANGIOGENESIS | GOBP_NEGATIVE_REGULATION_OF_SPROUTING_ANGIOGENESIS | 13 | 0.88988863 | 1.72977476 | 0.000507 | 0.03177085 | 0.02943571 | 176 | tags=15%, list | [illegible] |
| GOBP_REGULATION_OF_CARBOHYDRATE_BIOSYNTHETIC_PROCESS | GOBP_REGULATION_OF_CARBOHYDRATE_BIOSYNTHETIC_PROCESS | 80 | 0.6214015 | 1.69181726 | 0.00049796 | 0.03177085 | 0.02943571 | 996 | tags=15%, list | [illegible] |
| GOBP_DETECTION_OF_STIMULUS | GOBP_DETECTION_OF_STIMULUS | 127 | 0.55790515 | 1.61269908 | 0.00049929 | 0.03177085 | 0.02943571 | 1313 | tags=23%, list | [illegible] |
| GOBP_NEURON_PROJECTION_GUIDANCE | GOBP_NEURON_PROJECTION_GUIDANCE | 190 | 0.51676867 | 1.56872868 | 0.00050855 | 0.03177085 | 0.02943571 | 2249 | tags=32%, list | [illegible] |
| GOBP_FATTY_ACID_CATABOLIC_PROCESS | GOBP_FATTY_ACID_CATABOLIC_PROCESS | 92 | 0.60297648 | 1.67082218 | 0.00052311 | 0.03220536 | 0.02983829 | 1832 | tags=27%, list | [illegible] |
| GOBP_SKELETAL_SYSTEM_MORPHOGENESIS | GOBP_SKELETAL_SYSTEM_MORPHOGENESIS | 195 | 0.50725931 | 1.5426327 | 0.00053544 | 0.03269688 | 0.03029369 | 1928 | tags=23%, list | [illegible] |
| GOBP_TRICARBOXYLIC_ACID_METABOLIC_PROCESS | GOBP_TRICARBOXYLIC_ACID_METABOLIC_PROCESS | 13 | 0.88781789 | 1.72574963 | 0.00054421 | 0.03296422 | 0.03054138 | 1252 | tags=46%, list | [illegible] |
| GOBP_POSITIVE_REGULATION_OF_BMP_SIGNALING_PATHWAY | GOBP_POSITIVE_REGULATION_OF_BMP_SIGNALING_PATHWAY | 32 | 0.77022704 | 1.78660271 | 0.00058463 | 0.03457576 | 0.03203447 | 1827 | tags=38%, list | [illegible] |
| GOBP_COMPLEMENT_ACTIVATION_CLASSICAL_PATHWAY | GOBP_COMPLEMENT_ACTIVATION_CLASSICAL_PATHWAY | 22 | 0.82422623 | 1.78236052 | 0.00058193 | 0.03457576 | 0.03203447 | 364 | tags=23%, list | [illegible] |
| GOBP_POSITIVE_REGULATION_OF_SIGNALING | GOBP_POSITIVE_REGULATION_OF_SIGNALING | 1469 | 0.35713034 | 1.25242869 | 0.00058135 | 0.03457576 | 0.03203447 | 1681 | tags=14%, list | [illegible] |
| GOBP_CELLULAR_RESPONSE_TO_HORMONE_STIMULUS | GOBP_CELLULAR_RESPONSE_TO_HORMONE_STIMULUS | 536 | 0.41330957 | 1.37361171 | 0.0005961 | 0.03497922 | 0.03240828 | 1561 | tags=15%, list | [illegible] |
| GOBP_NEGATIVE_REGULATION_OF_CELL_COMMUNICATION | GOBP_NEGATIVE_REGULATION_OF_CELL_COMMUNICATION | 1245 | 0.36224635 | 1.26191066 | 0.00063526 | 0.03670328 | 0.03400562 | 1689 | tags=14%, list | [illegible] |
| GOBP_CELL_PROJECTION_ORGANIZATION | GOBP_CELL_PROJECTION_ORGANIZATION | 1335 | -0.3377822 | -1.2222792 | 0.00063405 | 0.03670328 | 0.03400562 | 2620 | tags=23%, list | [illegible] |
| GOBP_EXTERNAL_ENCAPSULATING_STRUCTURE_ORGANIZATION | GOBP_EXTERNAL_ENCAPSULATING_STRUCTURE_ORGANIZATION | 272 | 0.47570675 | 1.49883081 | 0.00064305 | 0.03686996 | 0.03416005 | 1766 | tags=24%, list | [illegible] |
| GOBP_INTERLEUKIN_11_MEDIATED_SIGNALING_PATHWAY | GOBP_INTERLEUKIN_11_MEDIATED_SIGNALING_PATHWAY | 7 | 0.95461802 | 1.63365497 | 0.00065754 | 0.03741526 | 0.03466527 | 80 | tags=29%, list | [illegible] |
| GOBP_SYNAPSE_ASSEMBLY | GOBP_SYNAPSE_ASSEMBLY | 216 | 0.49498811 | 1.52305756 | 0.00067032 | 0.03785562 | 0.03507326 | 1620 | tags=21%, list | [illegible] |
| GOBP_POSITIVE_REGULATION_OF_T_CELL_DIFFERENTIATION_IN_THYM | GOBP_POSITIVE_REGULATION_OF_T_CELL_DIFFERENTIATION_IN_THYM | 9 | 0.94232561 | 1.6948093 | 0.00070426 | 0.03836062 | 0.03554114 | 344 | tags=33%, list | [illegible] |
| GOBP_RESPONSE_TO_OSMOTIC_STRESS | GOBP_RESPONSE_TO_OSMOTIC_STRESS | 75 | -0.6101161 | -1.6825158 | 0.0007048 | 0.03836062 | 0.03554114 | 1881 | tags=31%, list | [illegible] |
| GOBP_DESMOSOME_ORGANIZATION | GOBP_DESMOSOME_ORGANIZATION | 10 | -0.9029085 | -1.6691649 | 0.00069864 | 0.03836062 | 0.03554114 | 624 | tags=50%, list | [illegible] |
| GOBP_REGULATION_OF_SMALL_MOLECULE_METABOLIC_PROCESS | GOBP_REGULATION_OF_SMALL_MOLECULE_METABOLIC_PROCESS | 266 | 0.47771582 | 1.50163195 | 0.00069436 | 0.03836062 | 0.03554114 | 804 | tags=11%, list | [illegible] |
| GOBP_PROCESS_UTILIZING_AUTOPHAGIC_MECHANISM | GOBP_PROCESS_UTILIZING_AUTOPHAGIC_MECHANISM | 559 | -0.3961633 | -1.3628618 | 0.00069263 | 0.03836062 | 0.03554114 | 3241 | tags=27%, list | [illegible] |
| GOBP_LIPID_METABOLIC_PROCESS | GOBP_LIPID_METABOLIC_PROCESS | 1026 | 0.37310348 | 1.28849741 | 0.00072197 | 0.03901218 | 0.03614482 | 1895 | tags=17%, list | [illegible] |
| GOBP_SINGLE_FERTILIZATION | GOBP_SINGLE_FERTILIZATION | 91 | -0.5731606 | -1.6272896 | 0.00073063 | 0.03919852 | 0.03631746 | 1585 | tags=20%, list | [illegible] |
| GOBP_POSTSYNAPTIC_SPECIALIZATION_ASSEMBLY | GOBP_POSTSYNAPTIC_SPECIALIZATION_ASSEMBLY | 31 | 0.75468781 | 1.74276728 | 0.0007394 | 0.0393874 | 0.03649246 | 1509 | tags=42%, list | [illegible] |
| GOBP_POSITIVE_REGULATION_OF_CELL_DEVELOPMENT | GOBP_POSITIVE_REGULATION_OF_CELL_DEVELOPMENT | 348 | 0.45071587 | 1.45121996 | 0.00075982 | 0.04019033 | 0.03723638 | 1818 | tags=17%, list | [illegible] |
| GOBP_SPECIFICATION_OF_SYMMETRY | GOBP_SPECIFICATION_OF_SYMMETRY | 103 | -0.5749707 | -1.6662372 | 0.00076837 | 0.04035844 | 0.03739213 | 1708 | tags=27%, list | [illegible] |
| GOBP_NEUTRAL_LIPID_BIOSYNTHETIC_PROCESS | GOBP_NEUTRAL_LIPID_BIOSYNTHETIC_PROCESS | 41 | 0.73010207 | 1.77325954 | 0.00077788 | 0.04057375 | 0.03759161 | 1134 | tags=29%, list | [illegible] |
| GOBP_REGULATION_OF_NERVOUS_SYSTEM_DEVELOPMENT | GOBP_REGULATION_OF_NERVOUS_SYSTEM_DEVELOPMENT | 341 | 0.45026925 | 1.44652084 | 0.00082727 | 0.04285265 | 0.03970301 | 1817 | tags=21%, list | [illegible] |
| GOBP_SYNAPSE_ORGANIZATION | GOBP_SYNAPSE_ORGANIZATION | 462 | 0.42796999 | 1.40793119 | 0.00083452 | 0.04293223 | 0.03977675 | 1925 | tags=20%, list | [illegible] |
| GOBP_CONNECTIVE_TISSUE_DEVELOPMENT | GOBP_CONNECTIVE_TISSUE_DEVELOPMENT | 259 | 0.47100883 | 1.47656821 | 0.00088737 | 0.04503379 | 0.04172384 | 1411 | tags=15%, list | [illegible] |
| GOBP_POSITIVE_REGULATION_OF_CELL_DIFFERENTIATION | GOBP_POSITIVE_REGULATION_OF_CELL_DIFFERENTIATION | 685 | 0.39557076 | 1.33537127 | 0.00088384 | 0.04503379 | 0.04172384 | 1822 | tags=18%, list | [illegible] |
| GOBP_REGULATION_OF_MICROTUBULE_BASED_PROCESS | GOBP_REGULATION_OF_MICROTUBULE_BASED_PROCESS | 225 | -0.4689783 | -1.4906835 | 0.00090168 | 0.04515002 | 0.04183153 | 2939 | tags=30%, list | [illegible] |
| GOBP_PROTEIN_MATURATION | GOBP_PROTEIN_MATURATION | 486 | -0.4011397 | -1.3649159 | 0.0009016 | 0.04515002 | 0.04183153 | 2581 | tags=23%, list | [illegible] |
| GOBP_CELLULAR_RESPONSE_TO_NITROGEN_COMPOUND | GOBP_CELLULAR_RESPONSE_TO_NITROGEN_COMPOUND | 520 | 0.41277024 | 1.36916264 | 0.00091387 | 0.04545737 | 0.04211629 | 1460 | tags=13%, list | [illegible] |
| GOBP_REGULATION_OF_CILIUM_DEPENDENT_CELL_MOTILITY | GOBP_REGULATION_OF_CILIUM_DEPENDENT_CELL_MOTILITY | 9 | -0.9267946 | -1.6725983 | 0.00092819 | 0.04584404 | 0.04247454 | 160 | tags=44%, list | [illegible] |
| GOBP_RHYTHMIC_PROCESS | GOBP_RHYTHMIC_PROCESS | 235 | 0.47697118 | 1.48002266 | 0.00093385 | 0.04584404 | 0.04247454 | 1320 | tags=15%, list | [illegible] |
| GOBP_NEGATIVE_REGULATION_OF_LIPID_LOCALIZATION | GOBP_NEGATIVE_REGULATION_OF_LIPID_LOCALIZATION | 33 | 0.74669044 | 1.74280742 | 0.00099291 | 0.04780622 | 0.0442925 | 543 | tags=24%, list | [illegible] |
| GOBP_REGULATION_OF_GLUCONEOGENESIS | GOBP_REGULATION_OF_GLUCONEOGENESIS | 45 | 0.7027994 | 1.73569905 | 0.00098935 | 0.04780622 | 0.0442925 | 996 | tags=18%, list | [illegible] |
| GOBP_CELLULAR_RESPONSE_TO_LITHIUM_ION | GOBP_CELLULAR_RESPONSE_TO_LITHIUM_ION | 8 | 0.94750275 | 1.66393148 | 0.00098073 | 0.04780622 | 0.0442925 | 140 | tags=25%, list | [illegible] |
| GOBP_POSTSYNAPTIC_DENSITY_ASSEMBLY | GOBP_POSTSYNAPTIC_DENSITY_ASSEMBLY | 24 | 0.79360717 | 1.75012992 | 0.00101249 | 0.04843833 | 0.04487815 | 841 | tags=38%, list | [illegible] |
| GOBP_REGULATION_OF_SYNAPSE_ASSEMBLY | GOBP_REGULATION_OF_SYNAPSE_ASSEMBLY | 145 | 0.52945504 | 1.55343639 | 0.00106063 | 0.05032293 | 0.04662423 | 1760 | tags=23%, list | [illegible] |
| GOBP_REGULATION_OF_CELL_JUNCTION_ASSEMBLY | GOBP_REGULATION_OF_CELL_JUNCTION_ASSEMBLY | 232 | 0.47381753 | 1.46876763 | 0.00106528 | 0.05032293 | 0.04662423 | 1760 | tags=23%, list | [illegible] |
| GOBP_LEUKOCYTE_DIFFERENTIATION | GOBP_LEUKOCYTE_DIFFERENTIATION | 526 | 0.41591787 | 1.38068977 | 0.0010763 | 0.05052533 | 0.04681176 | 1315 | tags=13%, list | [illegible] |
| GOBP_POSITIVE_REGULATION_OF_COLD_INDUCED_THERMOGENESIS | GOBP_POSITIVE_REGULATION_OF_COLD_INDUCED_THERMOGENESIS | 85 | 0.60971205 | 1.67304569 | 0.00112354 | 0.05145654 | 0.04767452 | 1996 | tags=31%, list | [illegible] |
| GOBP_INTRINSIC_APOPTOTIC_SIGNALING_PATHWAY_BY_P53_CLASS_I | GOBP_INTRINSIC_APOPTOTIC_SIGNALING_PATHWAY_BY_P53_CLASS_ | 83 | -0.5922418 | -1.6566955 | 0.00110797 | 0.05145654 | 0.04767452 | 1909 | tags=25%, list | [illegible] |
| GOBP_COGNITION | GOBP_COGNITION | 234 | 0.47495239 | 1.47271524 | 0.00111526 | 0.05145654 | 0.04767452 | 1270 | tags=17%, list | [illegible] |
| GOBP_NEGATIVE_REGULATION_OF_DEVELOPMENTAL_PROCESS | GOBP_NEGATIVE_REGULATION_OF_DEVELOPMENTAL_PROCESS | 775 | 0.38353761 | 1.30439378 | 0.00112075 | 0.05145654 | 0.04767452 | 1767 | tags=17%, list | [illegible] |
| GOBP_NEGATIVE_REGULATION_OF_ADENYLATE_CYCLASE_ACTIVATING | GOBP_NEGATIVE_REGULATION_OF_ADENYLATE_CYCLASE_ACTIVATING | 12 | 0.89554458 | 1.71309312 | 0.00114731 | 0.05222694 | 0.0483883 | 713 | tags=50%, list | [illegible] |
| GOBP_NEGATIVE_REGULATION_OF_TRIGLYCERIDE_METABOLIC_PROCE | GOBP_NEGATIVE_REGULATION_OF_TRIGLYCERIDE_METABOLIC_PROCE | 12 | 0.89520153 | 1.71243689 | 0.00116597 | 0.05275641 | 0.04887885 | 708 | tags=33%, list | [illegible] |
| GOBP_REGULATION_OF_POSTSYNAPTIC_SPECIALIZATION_ASSEMBLY | GOBP_REGULATION_OF_POSTSYNAPTIC_SPECIALIZATION_ASSEMBLY | 20 | 0.80981844 | 1.71896406 | 0.00122758 | 0.05443567 | 0.05043469 | 1509 | tags=50%, list | [illegible] |
| GOBP_CELLULAR_RESPONSE_TO_CALCIUM_ION | GOBP_CELLULAR_RESPONSE_TO_CALCIUM_ION | 60 | 0.66039545 | 1.71581243 | 0.00121499 | 0.05443567 | 0.05043469 | 1245 | tags=23%, list | [illegible] |
| GOBP_NUTRIENT_STORAGE | GOBP_NUTRIENT_STORAGE | 76 | 0.62168948 | 1.67746703 | 0.00123106 | 0.05443567 | 0.05043469 | 279 | tags=17%, list | [illegible] |
| GOBP_FATTY_ACID_HOMEOSTASIS | GOBP_FATTY_ACID_HOMEOSTASIS | 14 | 0.84964691 | 1.67348211 | 0.0012477 | 0.05443567 | 0.05043469 | 643 | tags=36%, list | [illegible] |
| GOBP_TRIGLYCERIDE_RICH_LIPOPROTEIN_PARTICLE_REMODELING | GOBP_TRIGLYCERIDE_RICH_LIPOPROTEIN_PARTICLE_REMODELING | 7 | 0.94767411 | 1.62177174 | 0.00124552 | 0.05443567 | 0.05043469 | 576 | tags=57%, list | [illegible] |
| GOBP_ADENYLATE_CYCLASE_ACTIVATING_G_PROTEIN_COUPLED_RECE | GOBP_ADENYLATE_CYCLASE_ACTIVATING_G_PROTEIN_COUPLED_RECE | 102 | 0.57435439 | 1.61647551 | 0.00123689 | 0.05443567 | 0.05043469 | 1545 | tags=25%, list | [illegible] |
| GOBP_REGULATION_OF_CARBOHYDRATE_METABOLIC_PROCESS | GOBP_REGULATION_OF_CARBOHYDRATE_METABOLIC_PROCESS | 145 | 0.52658007 | 1.54500115 | 0.00126106 | 0.05443567 | 0.05043469 | 2281 | tags=23%, list | [illegible] |
| GOBP_CELLULAR_RESPONSE_TO_INSULIN_STIMULUS | GOBP_CELLULAR_RESPONSE_TO_INSULIN_STIMULUS | 187 | 0.49919593 | 1.51024548 | 0.00125826 | 0.05443567 | 0.05043469 | 1245 | tags=15%, list | [illegible] |
| GOBP_FEMALE_SEX_DIFFERENTIATION | GOBP_FEMALE_SEX_DIFFERENTIATION | 95 | 0.57809833 | 1.60820652 | 0.00129337 | 0.05551138 | 0.05143134 | 1289 | tags=24%, list | [illegible] |
| GOBP_POSTSYNAPSE_ASSEMBLY | GOBP_POSTSYNAPSE_ASSEMBLY | 73 | 0.62247071 | 1.66949798 | 0.0013572 | 0.05791994 | 0.05366287 | 1853 | tags=25%, list | [illegible] |
| GOBP_REGULATION_OF_ANATOMICAL_STRUCTURE_MORPHOGENESIS | GOBP_REGULATION_OF_ANATOMICAL_STRUCTURE_MORPHOGENESIS | 702 | 0.38905803 | 1.31530806 | 0.00137856 | 0.05849924 | 0.05419959 | 1853 | tags=17%, list | [illegible] |
| GOBP_REGULATION_OF_CILIUM_MOVEMENT | GOBP_REGULATION_OF_CILIUM_MOVEMENT | 14 | -0.8527115 | -1.7030506 | 0.00144016 | 0.05981175 | 0.05541563 | 1270 | tags=50%, list | [illegible] |
| GOBP_NEGATIVE_REGULATION_OF_FATTY_ACID_OXIDATION | GOBP_NEGATIVE_REGULATION_OF_FATTY_ACID_OXIDATION | 14 | 0.84525823 | 1.66483808 | 0.00143532 | 0.05981175 | 0.05541563 | 498 | tags=43%, list | [illegible] |
| GOBP_B_CELL_MEDIATED_IMMUNITY | GOBP_B_CELL_MEDIATED_IMMUNITY | 99 | 0.58312674 | 1.63361948 | 0.0014392 | 0.05981175 | 0.05541563 | 675 | tags=9%, list | [illegible] |
| GOBP_EMBRYONIC_SKELETAL_SYSTEM_DEVELOPMENT | GOBP_EMBRYONIC_SKELETAL_SYSTEM_DEVELOPMENT | 109 | 0.56571482 | 1.6040385 | 0.00144134 | 0.05981175 | 0.05541563 | 1357 | tags=17%, list | [illegible] |
| GOBP_RENAL_SYSTEM_PROCESS | GOBP_RENAL_SYSTEM_PROCESS | 95 | 0.57512875 | 1.59994547 | 0.0014818 | 0.0608456 | 0.0563735 | 1320 | tags=22%, list | [illegible] |
| GOBP_LOCOMOTORY_BEHAVIOR | GOBP_LOCOMOTORY_BEHAVIOR | 151 | 0.52084022 | 1.53565466 | 0.00148246 | 0.0608456 | 0.0563735 | 1823 | tags=23%, list | [illegible] |
| GOBP_RESPONSE_TO_ALCOHOL | GOBP_RESPONSE_TO_ALCOHOL | 199 | 0.49531608 | 1.51069257 | 0.00154626 | 0.06311946 | 0.05848023 | 1555 | tags=19%, list | [illegible] |
| GOBP_EMBRYONIC_SKELETAL_SYSTEM_MORPHOGENESIS | GOBP_EMBRYONIC_SKELETAL_SYSTEM_MORPHOGENESIS | 76 | 0.61782363 | 1.66703606 | 0.00157541 | 0.0639617 | 0.05926057 | 562 | tags=14%, list | [illegible] |
| GOBP_REGULATION_OF_POSTSYNAPSE_ORGANIZATION | GOBP_REGULATION_OF_POSTSYNAPSE_ORGANIZATION | 128 | 0.53996648 | 1.56240282 | 0.00160919 | 0.064982 | 0.06020588 | 1719 | tags=20%, list | [illegible] |
| GOBP_SPERM_EGG_RECOGNITION | GOBP_SPERM_EGG_RECOGNITION | 23 | -0.7698618 | -1.7008762 | 0.00164685 | 0.06614681 | 0.06128507 | 706 | tags=26%, list | [illegible] |
| GOBP_RESPONSE_TO_ETHANOL | GOBP_RESPONSE_TO_ETHANOL | 88 | 0.58106451 | 1.60208407 | 0.0016814 | 0.06717532 | 0.06223798 | 1354 | tags=20%, list | [illegible] |
| GOBP_HOMEOSTATIC_PROCESS | GOBP_HOMEOSTATIC_PROCESS | 1369 | 0.35572346 | 1.24459539 | 0.00170945 | 0.06793486 | 0.06294171 | 1555 | tags=15%, list | [illegible] |
| GOBP_SKIN_EPIDERMIS_DEVELOPMENT | GOBP_SKIN_EPIDERMIS_DEVELOPMENT | 94 | -0.5677684 | -1.6212983 | 0.0017997 | 0.07069219 | 0.06549637 | 479 | tags=17%, list | [illegible] |
| GOBP_ODONTOGENESIS | GOBP_ODONTOGENESIS | 89 | -0.5632774 | -1.5937399 | 0.00180707 | 0.07069219 | 0.06549637 | 656 | tags=27%, list | [illegible] |
| GOBP_ANATOMICAL_STRUCTURE_FORMATION_INVOLVED_IN_MORPH | GOBP_ANATOMICAL_STRUCTURE_FORMATION_INVOLVED_IN_MORPH | 1016 | 0.36464565 | 1.2584515 | 0.00179494 | 0.07069219 | 0.06549637 | 1629 | tags=16%, list | [illegible] |
| GOBP_ACTIVATION_OF_IMMUNE_RESPONSE | GOBP_ACTIVATION_OF_IMMUNE_RESPONSE | 471 | 0.41696313 | 1.37366841 | 0.00182445 | 0.07100248 | 0.06578386 | 1855 | tags=15%, list | [illegible] |
| GOBP_REGULATION_OF_BLOOD_PRESSURE | GOBP_REGULATION_OF_BLOOD_PRESSURE | 129 | 0.53979627 | 1.56386268 | 0.00183967 | 0.07122545 | 0.06599043 | 1468 | tags=19%, list | [illegible] |
| GOBP_REGULATION_OF_POSTSYNAPTIC_DENSITY_ASSEMBLY | GOBP_REGULATION_OF_POSTSYNAPTIC_DENSITY_ASSEMBLY | 16 | 0.84487877 | 1.70992198 | 0.001868 | 0.07132587 | 0.06608347 | 591 | tags=44%, list | [illegible] |
| GOBP_GLYCEROLIPID_CATABOLIC_PROCESS | GOBP_GLYCEROLIPID_CATABOLIC_PROCESS | 51 | 0.65706682 | 1.65241614 | 0.00189013 | 0.07132587 | 0.06608347 | 1218 | tags=27%, list | [illegible] |
| GOBP_DNA_STRAND_RESECTION_INVOLVED_IN_REPLICATION_FORK_P | GOBP_DNA_STRAND_RESECTION_INVOLVED_IN_REPLICATION_FORK_F | 10 | -0.8851179 | -1.6362762 | 0.00189924 | 0.07132587 | 0.06608347 | 487 | tags=30%, list | [illegible] |
| GOBP_POSITIVE_REGULATION_OF_SYNAPTIC_TRANSMISSION | GOBP_POSITIVE_REGULATION_OF_SYNAPTIC_TRANSMISSION | 112 | 0.55883461 | 1.58903311 | 0.00186393 | 0.07132587 | 0.06608347 | 1121 | tags=21%, list | [illegible] |
| GOBP_VIRAL_LIFE_CYCLE | GOBP_VIRAL_LIFE_CYCLE | 241 | -0.4465362 | -1.4301072 | 0.0018872 | 0.07132587 | 0.06608347 | 2939 | tags=27%, list | [illegible] |
| GOBP_RESPONSE_TO_LIPID | GOBP_RESPONSE_TO_LIPID | 696 | 0.38694248 | 1.307445 | 0.00189477 | 0.07132587 | 0.06608347 | 1561 | tags=14%, list | [illegible] |
| GOBP_REGULATION_OF_TRIGLYCERIDE_METABOLIC_PROCESS | GOBP_REGULATION_OF_TRIGLYCERIDE_METABOLIC_PROCESS | 36 | 0.70959662 | 1.68247873 | 0.00191735 | 0.07164798 | 0.06638191 | 218 | tags=19%, list | [illegible] |
| GOBP_ADENYLATE_CYCLASE_INHIBITING_G_PROTEIN_COUPLED_RECE | GOBP_ADENYLATE_CYCLASE_INHIBITING_G_PROTEIN_COUPLED_RECE | 52 | 0.66477574 | 1.68051967 | 0.00195142 | 0.0725601 | 0.0672269 | 1367 | tags=31%, list | [illegible] |
| GOBP_LEARNING_OR_MEMORY | GOBP_LEARNING_OR_MEMORY | 199 | 0.49057199 | 1.49622329 | 0.00202255 | 0.07483425 | 0.06933399 | 1342 | tags=18%, list | [illegible] |
| GOBP_MONOSACCHARIDE_METABOLIC_PROCESS | GOBP_MONOSACCHARIDE_METABOLIC_PROCESS | 220 | 0.47676219 | 1.46941483 | 0.00205287 | 0.07558373 | 0.07002839 | 1218 | tags=14%, list | [illegible] |
| GOBP_MONOCARBOXYLIC_ACID_CATABOLIC_PROCESS | GOBP_MONOCARBOXYLIC_ACID_CATABOLIC_PROCESS | 104 | 0.56281027 | 1.5854381 | 0.00211947 | 0.07765539 | 0.07194778 | 1868 | tags=27%, list | [illegible] |
| GOBP_MAPK_CASCADE | GOBP_MAPK_CASCADE | 610 | 0.39126254 | 1.31024013 | 0.00214175 | 0.07809069 | 0.07235109 | 1818 | tags=17%, list | [illegible] |
| GOBP_POSITIVE_REGULATION_OF_MUSCLE_CELL_DIFFERENTIATION | GOBP_POSITIVE_REGULATION_OF_MUSCLE_CELL_DIFFERENTIATION | 55 | 0.64800695 | 1.65526062 | 0.00236701 | 0.08588702 | 0.0795744 | 1822 | tags=33%, list | [illegible] |
| GOBP_ASSOCIATIVE_LEARNING | GOBP_ASSOCIATIVE_LEARNING | 65 | 0.6227389 | 1.63788849 | 0.0024087 | 0.08697944 | 0.08058652 | 841 | tags=23%, list | [illegible] |
| GOBP_SYNAPTIC_TRANSMISSION_GLUTAMATERGIC | GOBP_SYNAPTIC_TRANSMISSION_GLUTAMATERGIC | 70 | 0.60890321 | 1.61878681 | 0.0024512 | 0.08809068 | 0.08161609 | 956 | tags=23%, list | [illegible] |
| GOBP_REGULATION_OF_LIPID_LOCALIZATION | GOBP_REGULATION_OF_LIPID_LOCALIZATION | 114 | 0.5450406 | 1.55205691 | 0.00255362 | 0.09133444 | 0.08462143 | 736 | tags=15%, list | [illegible] |
| GOBP_EXTRACELLULAR_MATRIX_ASSEMBLY | GOBP_EXTRACELLULAR_MATRIX_ASSEMBLY | 41 | 0.69859952 | 1.69674666 | 0.00257421 | 0.09163464 | 0.08489957 | 1567 | tags=37%, list | [illegible] |
| GOBP_CEREBROSPINAL_FLUID_CIRCULATION | GOBP_CEREBROSPINAL_FLUID_CIRCULATION | 9 | -0.9077342 | -1.6381996 | 0.00261173 | 0.09253173 | 0.08573072 | 1075 | tags=89%, list | [illegible] |
| GOBP_NEURON_DEVELOPMENT | GOBP_NEURON_DEVELOPMENT | 915 | 0.36834971 | 1.26378749 | 0.00264509 | 0.09327361 | 0.08641807 | 1870 | tags=18%, list | [illegible] |
| GOBP_REGULATION_OF_CELL_DIFFERENTIATION | GOBP_REGULATION_OF_CELL_DIFFERENTIATION | 1246 | 0.35417816 | 1.2339224 | 0.00271084 | 0.09514554 | 0.08815241 | 1822 | tags=16%, list | [illegible] |
| GOBP_OUTER_DYNEIN_ARM_ASSEMBLY | GOBP_OUTER_DYNEIN_ARM_ASSEMBLY | 9 | -0.9057396 | -1.6346 | 0.0027894 | 0.0969962 | 0.08986705 | 1059 | tags=78%, list | [illegible] |
| GOBP_CELL_CYCLE_CHECKPOINT_SIGNALING | GOBP_CELL_CYCLE_CHECKPOINT_SIGNALING | 191 | -0.4678159 | -1.461196 | 0.00278623 | 0.0969962 | 0.08986705 | 3574 | tags=31%, list | [illegible] |
| GOBP_POSITIVE_REGULATION_OF_IMMUNE_SYSTEM_PROCESS | GOBP_POSITIVE_REGULATION_OF_IMMUNE_SYSTEM_PROCESS | 859 | 0.37050896 | 1.26718491 | 0.00281602 | 0.09747063 | 0.09030661 | 1831 | tags=14%, list | [illegible] |
| GOBP_FEEDING_BEHAVIOR | GOBP_FEEDING_BEHAVIOR | 54 | 0.65090817 | 1.65481544 | 0.00297276 | 0.10242394 | 0.09489586 | 1528 | tags=24%, list | [illegible] |
| GOBP_CARBOHYDRATE_HOMEOSTASIS | GOBP_CARBOHYDRATE_HOMEOSTASIS | 196 | 0.47875492 | 1.45750056 | 0.00304114 | 0.10430155 | 0.09663547 | 1317 | tags=15%, list | [illegible] |
| GOBP_ORGANIC_HYDROXY_COMPOUND_TRANSPORT | GOBP_ORGANIC_HYDROXY_COMPOUND_TRANSPORT | 176 | 0.50163139 | 1.50638718 | 0.00307332 | 0.10492585 | 0.09721388 | 833 | tags=14%, list | [illegible] |
| GOBP_POSITIVE_REGULATION_OF_GLUCOSE_METABOLIC_PROCESS | GOBP_POSITIVE_REGULATION_OF_GLUCOSE_METABOLIC_PROCESS | 40 | 0.69313381 | 1.6774216 | 0.0031074 | 0.10513369 | 0.09740645 | 2182 | tags=35%, list | [illegible] |
| GOBP_NEGATIVE_REGULATION_OF_CELL_CYCLE_PROCESS | GOBP_NEGATIVE_REGULATION_OF_CELL_CYCLE_PROCESS | 288 | -0.4209093 | -1.3701966 | 0.00310087 | 0.10513369 | 0.09740645 | 2693 | tags=22%, list | [illegible] |
| GOBP_REGULATION_OF_SMALL_GTPASE_MEDIATED_SIGNAL_TRANSD | GOBP_REGULATION_OF_SMALL_GTPASE_MEDIATED_SIGNAL_TRANSD | 293 | 0.44802434 | 1.42182146 | 0.00312337 | 0.10520025 | 0.09746812 | 2147 | tags=21%, list | [illegible] |
| GOBP_HEMOPOIESIS | GOBP_HEMOPOIESIS | 807 | 0.37399185 | 1.27411986 | 0.00316106 | 0.10599415 | 0.09820366 | 1324 | tags=12%, list | [illegible] |
| GOBP_RESPONSE_TO_CALCIUM_ION | GOBP_RESPONSE_TO_CALCIUM_ION | 103 | 0.55480417 | 1.56168497 | 0.00317995 | 0.10615372 | 0.0983515 | 1245 | tags=20%, list | [illegible] |
| GOBP_LONG_TERM_SYNAPTIC_POTENTIATION | GOBP_LONG_TERM_SYNAPTIC_POTENTIATION | 74 | 0.59692487 | 1.60434021 | 0.00320271 | 0.1064406 | 0.0986173 | 1121 | tags=23%, list | [illegible] |
| GOBP_VASCULAR_PROCESS_IN_CIRCULATORY_SYSTEM | GOBP_VASCULAR_PROCESS_IN_CIRCULATORY_SYSTEM | 206 | 0.47840909 | 1.46451206 | 0.00323102 | 0.10655714 | 0.09872528 | 1122 | tags=18%, list | [illegible] |
| GOBP_REGULATION_OF_SYSTEM_PROCESS | GOBP_REGULATION_OF_SYSTEM_PROCESS | 404 | 0.41853387 | 1.36199664 | 0.00323459 | 0.10655714 | 0.09872528 | 1355 | tags=17%, list | [illegible] |
| GOBP_LYMPHOCYTE_DIFFERENTIATION | GOBP_LYMPHOCYTE_DIFFERENTIATION | 359 | 0.42471664 | 1.37012338 | 0.00324897 | 0.10656351 | 0.09873118 | 1478 | tags=14%, list | [illegible] |
| GOBP_SMALL_MOLECULE_METABOLIC_PROCESS | GOBP_SMALL_MOLECULE_METABOLIC_PROCESS | 1353 | 0.34838032 | 1.2185789 | 0.0032733 | 0.1068947 | 0.09903802 | 1613 | tags=13%, list | [illegible] |
| GOBP_RESPONSE_TO_XENOBIOTIC_STIMULUS | GOBP_RESPONSE_TO_XENOBIOTIC_STIMULUS | 268 | 0.45502692 | 1.43197171 | 0.00331181 | 0.10768396 | 0.09976927 | 1561 | tags=18%, list | [illegible] |
| GOBP_TRANS_SYNAPTIC_SIGNALING_MODULATING_SYNAPTIC_TRANS | GOBP_TRANS_SYNAPTIC_SIGNALING_MODULATING_SYNAPTIC_TRANS | 17 | 0.81942709 | 1.68153395 | 0.00346444 | 0.11167982 | 0.10347144 | 1748 | tags=47%, list | [illegible] |
| GOBP_FAT_CELL_PROLIFERATION | GOBP_FAT_CELL_PROLIFERATION | 17 | 0.81636283 | 1.67524585 | 0.00346444 | 0.11167982 | 0.10347144 | 360 | tags=24%, list | [illegible] |
| GOBP_NEGATIVE_REGULATION_OF_CAMP_PKA_SIGNAL_TRANSDUCTIC | GOBP_NEGATIVE_REGULATION_OF_CAMP_PKA_SIGNAL_TRANSDUCTIC | 25 | 0.76175602 | 1.69264303 | 0.00358423 | 0.11407266 | 0.10568841 | 1657 | tags=44%, list | [illegible] |
| GOBP_NON_CANONICAL_WNT_SIGNALING_PATHWAY | GOBP_NON_CANONICAL_WNT_SIGNALING_PATHWAY | 58 | 0.62299291 | 1.60972709 | 0.00357546 | 0.11407266 | 0.10568841 | 1753 | tags=28%, list | [illegible] |
| GOBP_POSITIVE_REGULATION_OF_NERVOUS_SYSTEM_DEVELOPMENT | GOBP_POSITIVE_REGULATION_OF_NERVOUS_SYSTEM_DEVELOPMENT | 215 | 0.47133189 | 1.44976322 | 0.00357015 | 0.11407266 | 0.10568841 | 1817 | tags=21%, list | [illegible] |
| GOBP_REGULATION_OF_CELLULAR_RESPONSE_TO_GROWTH_FACTOR_ | GOBP_REGULATION_OF_CELLULAR_RESPONSE_TO_GROWTH_FACTOR_ | 271 | 0.45137337 | 1.42097871 | 0.00365472 | 0.11582521 | 0.10731215 | 1827 | tags=19%, list | [illegible] |
| GOBP_SYMBIONT_ENTRY_INTO_HOST | GOBP_SYMBIONT_ENTRY_INTO_HOST | 110 | -0.5288464 | -1.5425943 | 0.00368045 | 0.11615068 | 0.1076137 | 2170 | tags=25%, list | [illegible] |
| GOBP_RETROGRADE_TRANS_SYNAPTIC_SIGNALING_BY_LIPID | GOBP_RETROGRADE_TRANS_SYNAPTIC_SIGNALING_BY_LIPID | 4 | -0.9756689 | -1.4813295 | 0.00372505 | 0.11657867 | 0.10801023 | 288 | tags=50%, list | [illegible] |
| GOBP_RESPONSE_TO_METAL_ION | GOBP_RESPONSE_TO_METAL_ION | 242 | 0.4584308 | 1.42705541 | 0.00371813 | 0.11657867 | 0.10801023 | 1308 | tags=17%, list | [illegible] |
| GOBP_POSITIVE_REGULATION_OF_CARBOHYDRATE_METABOLIC_PROC | GOBP_POSITIVE_REGULATION_OF_CARBOHYDRATE_METABOLIC_PROC | 65 | 0.60756224 | 1.59797181 | 0.00376506 | 0.11734177 | 0.10871725 | 2182 | tags=32%, list | [illegible] |
| GOBP_DETECTION_OF_STIMULUS_INVOLVED_IN_SENSORY_PERCEPTIO | GOBP_DETECTION_OF_STIMULUS_INVOLVED_IN_SENSORY_PERCEPTIO | 54 | 0.64535814 | 1.64070551 | 0.00378295 | 0.11741208 | 0.10878239 | 1167 | tags=26%, list | [illegible] |
| GOBP_WHITE_FAT_CELL_DIFFERENTIATION | GOBP_WHITE_FAT_CELL_DIFFERENTIATION | 18 | 0.82034643 | 1.70399731 | 0.00384304 | 0.11781669 | 0.10915726 | 946 | tags=28%, list | [illegible] |
| GOBP_POSITIVE_REGULATION_OF_LIPID_CATABOLIC_PROCESS | GOBP_POSITIVE_REGULATION_OF_LIPID_CATABOLIC_PROCESS | 19 | 0.80485262 | 1.69145484 | 0.00384149 | 0.11781669 | 0.10915726 | 1366 | tags=37%, list | [illegible] |
| GOBP_CYTOPLASMIC_MICROTUBULE_ORGANIZATION | GOBP_CYTOPLASMIC_MICROTUBULE_ORGANIZATION | 66 | -0.5879699 | -1.5891625 | 0.00384123 | 0.11781669 | 0.10915726 | 2639 | tags=35%, list | [illegible] |
| GOBP_RESPONSE_TO_EPINEPHRINE | GOBP_RESPONSE_TO_EPINEPHRINE | 12 | 0.87094634 | 1.66603898 | 0.00386965 | 0.11782427 | 0.10916428 | 713 | tags=33%, list | [illegible] |
| GOBP_MONONUCLEAR_CELL_DIFFERENTIATION | GOBP_MONONUCLEAR_CELL_DIFFERENTIATION | 441 | 0.40981838 | 1.3431949 | 0.00387466 | 0.11782427 | 0.10916428 | 1315 | tags=13%, list | [illegible] |
| GOBP_NEGATIVE_REGULATION_OF_MULTICELLULAR_ORGANISMAL_PI | GOBP_NEGATIVE_REGULATION_OF_MULTICELLULAR_ORGANISMAL_PI | 868 | 0.3678995 | 1.25866864 | 0.00391175 | 0.11847242 | 0.10976479 | 1821 | tags=17%, list | [illegible] |
| GOBP_OVULATION_FROM_OVARIAN_FOLLICLE | GOBP_OVULATION_FROM_OVARIAN_FOLLICLE | 9 | 0.90853088 | 1.63402817 | 0.00394109 | 0.11864904 | 0.10992843 | 263 | tags=22%, list | [illegible] |
| GOBP_SIGNAL_RELEASE | GOBP_SIGNAL_RELEASE | 340 | 0.429123 | 1.37892635 | 0.00394918 | 0.11864904 | 0.10992843 | 1036 | tags=14%, list | [illegible] |
| GOBP_PHOSPHATIDYLINOSITOL_ACYL_CHAIN_REMODELING | GOBP_PHOSPHATIDYLINOSITOL_ACYL_CHAIN_REMODELING | 4 | -0.9674887 | -1.4689098 | 0.00402685 | 0.12050055 | 0.11164385 | 34 | tags=50%, list | [illegible] |
| GOBP_COMPLEMENT_ACTIVATION | GOBP_COMPLEMENT_ACTIVATION | 37 | 0.69678124 | 1.66295405 | 0.00404624 | 0.12060051 | 0.11173647 | 364 | tags=16%, list | [illegible] |
| GOBP_PROTEIN_CONTAINING_COMPLEX_DISASSEMBLY | GOBP_PROTEIN_CONTAINING_COMPLEX_DISASSEMBLY | 244 | -0.4367273 | -1.4004245 | 0.00413034 | 0.12262034 | 0.11360784 | 3234 | tags=33%, list | [illegible] |
| GOBP_CELLULAR_RESPONSE_TO_PEPTIDE_HORMONE_STIMULUS | GOBP_CELLULAR_RESPONSE_TO_PEPTIDE_HORMONE_STIMULUS | 282 | 0.44612832 | 1.41016713 | 0.00416233 | 0.12308373 | 0.11403717 | 1347 | tags=15%, list | [illegible] |
| GOBP_REGULATION_OF_FLAGELLATED_SPERM_MOTILITY | GOBP_REGULATION_OF_FLAGELLATED_SPERM_MOTILITY | 4 | -0.9658701 | -1.4664524 | 0.0042186 | 0.12425846 | 0.11512556 | 160 | tags=100%, lis | [illegible] |
| GOBP_REGULATION_OF_PRESYNAPTIC_MEMBRANE_POTENTIAL | GOBP_REGULATION_OF_PRESYNAPTIC_MEMBRANE_POTENTIAL | 9 | 0.91298859 | 1.64204553 | 0.00425974 | 0.12498001 | 0.11579407 | 435 | tags=56%, list | [illegible] |
| GOBP_POSITIVE_REGULATION_OF_CELL_MORPHOGENESIS | GOBP_POSITIVE_REGULATION_OF_CELL_MORPHOGENESIS | 32 | 0.71931861 | 1.66851658 | 0.00430866 | 0.12592342 | 0.11666815 | 1104 | tags=19%, list | [illegible] |
| GOBP_IMMUNE_RESPONSE_REGULATING_SIGNALING_PATHWAY | GOBP_IMMUNE_RESPONSE_REGULATING_SIGNALING_PATHWAY | 457 | 0.40535173 | 1.33192791 | 0.00436315 | 0.12702173 | 0.11768573 | 1551 | tags=14%, list | [illegible] |
| GOBP_DEVELOPMENT_OF_PRIMARY_FEMALE_SEXUAL_CHARACTERISTI | GOBP_DEVELOPMENT_OF_PRIMARY_FEMALE_SEXUAL_CHARACTERISTI | 84 | 0.57710843 | 1.58240005 | 0.00439101 | 0.12733925 | 0.11797991 | 1289 | tags=25%, list | [illegible] |
| GOBP_ACETYLCHOLINE_SECRETION | GOBP_ACETYLCHOLINE_SECRETION | 1 | -0.997677 | -1.3320158 | 0.00454007 | 0.13115567 | 0.12151583 | 36 | tags=100%, lis | [illegible] |
| GOBP_VENTRICULAR_COMPACT_MYOCARDIUM_MORPHOGENESIS | GOBP_VENTRICULAR_COMPACT_MYOCARDIUM_MORPHOGENESIS | 5 | -0.9507742 | -1.5120946 | 0.00458628 | 0.13198293 | 0.12228229 | 10 | tags=40%, list | [illegible] |
| GOBP_CELLULAR_RESPONSE_TO_OXYGEN_CONTAINING_COMPOUND | GOBP_CELLULAR_RESPONSE_TO_OXYGEN_CONTAINING_COMPOUND | 920 | 0.36413865 | 1.24943098 | 0.00467581 | 0.13404585 | 0.12419358 | 1840 | tags=14%, list | [illegible] |
| GOBP_POSITIVE_REGULATION_OF_LIPID_BIOSYNTHETIC_PROCESS | GOBP_POSITIVE_REGULATION_OF_LIPID_BIOSYNTHETIC_PROCESS | 76 | 0.58349001 | 1.57439573 | 0.00469925 | 0.13420552 | 0.12434152 | 2287 | tags=34%, list | [illegible] |
| GOBP_REGULATION_OF_POSTSYNAPTIC_DENSITY_ORGANIZATION | GOBP_REGULATION_OF_POSTSYNAPTIC_DENSITY_ORGANIZATION | 21 | 0.78380035 | 1.68354682 | 0.00481444 | 0.13697456 | 0.12690704 | 591 | tags=33%, list | [illegible] |
| GOBP_INTRINSIC_APOPTOTIC_SIGNALING_PATHWAY_IN_RESPONSE_TI | GOBP_INTRINSIC_APOPTOTIC_SIGNALING_PATHWAY_IN_RESPONSE_T | 48 | -0.6340856 | -1.6257417 | 0.00490938 | 0.13914847 | 0.12892117 | 1909 | tags=29%, list | [illegible] |
| GOBP_REGULATION_OF_CELL_CYCLE_PHASE_TRANSITION | GOBP_REGULATION_OF_CELL_CYCLE_PHASE_TRANSITION | 425 | -0.3890183 | -1.310404 | 0.00493218 | 0.13926926 | 0.12903308 | 3254 | tags=27%, list | [illegible] |
| GOBP_PROTEIN_LOCALIZATION_TO_CILIARY_TRANSITION_ZONE | GOBP_PROTEIN_LOCALIZATION_TO_CILIARY_TRANSITION_ZONE | 20 | -0.773283 | -1.6654509 | 0.00496032 | 0.13931496 | 0.12907542 | 1580 | tags=50%, list | [illegible] |
| GOBP_CAMP_PKA_SIGNAL_TRANSDUCTION | GOBP_CAMP_PKA_SIGNAL_TRANSDUCTION | 51 | 0.63329264 | 1.59262794 | 0.00498945 | 0.13931496 | 0.12907542 | 1657 | tags=37%, list | [illegible] |
| GOBP_NEGATIVE_REGULATION_OF_STEROL_TRANSPORT | GOBP_NEGATIVE_REGULATION_OF_STEROL_TRANSPORT | 7 | 0.92654286 | 1.58560945 | 0.00498235 | 0.13931496 | 0.12907542 | 543 | tags=29%, list | [illegible] |
| GOBP_REGULATION_OF_LEUKOCYTE_DIFFERENTIATION | GOBP_REGULATION_OF_LEUKOCYTE_DIFFERENTIATION | 280 | 0.43737036 | 1.3815496 | 0.00502426 | 0.13976733 | 0.12949454 | 1315 | tags=14%, list | [illegible] |
| GOBP_NEGATIVE_REGULATION_OF_ALCOHOL_BIOSYNTHETIC_PROCESS | GOBP_NEGATIVE_REGULATION_OF_ALCOHOL_BIOSYNTHETIC_PROCESS | 11 | -0.8605178 | -1.6216353 | 0.00509404 | 0.14118584 | 0.13080879 | 182 | tags=18%, list | [illegible] |
| GOBP_BIOLOGICAL_PROCESS_INVOLVED_IN_SYMBIOTIC_INTERACTION | GOBP_BIOLOGICAL_PROCESS_INVOLVED_IN_SYMBIOTIC_INTERACTION | 156 | -0.4711202 | -1.4328928 | 0.00520951 | 0.14385534 | 0.13328208 | 2627 | tags=28%, list | [illegible] |
| GOBP_REGULATION_OF_EXCITATORY_SYNAPSE_ASSEMBLY | GOBP_REGULATION_OF_EXCITATORY_SYNAPSE_ASSEMBLY | 19 | 0.79880262 | 1.67874033 | 0.00525677 | 0.14421428 | 0.13361464 | 925 | tags=42%, list | [illegible] |
| GOBP_KERATINIZATION | GOBP_KERATINIZATION | 17 | -0.7973435 | -1.6632123 | 0.00529931 | 0.14421428 | 0.13361464 | 479 | tags=41%, list | [illegible] |
| GOBP_METANEPHROS_DEVELOPMENT | GOBP_METANEPHROS_DEVELOPMENT | 73 | 0.58643793 | 1.57285625 | 0.00528501 | 0.14421428 | 0.13361464 | 1711 | tags=29%, list | [illegible] |
| GOBP_REGULATION_OF_LIPID_BIOSYNTHETIC_PROCESS | GOBP_REGULATION_OF_LIPID_BIOSYNTHETIC_PROCESS | 148 | 0.50209753 | 1.47611174 | 0.00527849 | 0.14421428 | 0.13361464 | 1203 | tags=17%, list | [illegible] |
| GOBP_PEPTIDYL_LYSINE_ACETYLATION | GOBP_PEPTIDYL_LYSINE_ACETYLATION | 14 | 0.83833541 | 1.65120273 | 0.00532569 | 0.14440877 | 0.13379484 | 868 | tags=21%, list | [illegible] |
| GOBP_POSITIVE_REGULATION_OF_RESPIRATORY_BURST | GOBP_POSITIVE_REGULATION_OF_RESPIRATORY_BURST | 6 | 0.93808632 | 1.56432072 | 0.00539531 | 0.1447292 | 0.13409171 | 204 | tags=33%, list | [illegible] |
| GOBP_LIPID_DROPLET_FUSION | GOBP_LIPID_DROPLET_FUSION | 6 | 0.93728781 | 1.56298915 | 0.00539531 | 0.1447292 | 0.13409171 | 610 | tags=50%, list | [illegible] |
| GOBP_POSITIVE_REGULATION_OF_IMMUNE_RESPONSE | GOBP_POSITIVE_REGULATION_OF_IMMUNE_RESPONSE | 591 | 0.38326435 | 1.28142443 | 0.00537372 | 0.1447292 | 0.13409171 | 1855 | tags=14%, list | [illegible] |
| GOBP_MULTICELLULAR_ORGANISMAL_RESPONSE_TO_STRESS | GOBP_MULTICELLULAR_ORGANISMAL_RESPONSE_TO_STRESS | 60 | 0.60367617 | 1.56844673 | 0.00545626 | 0.14576912 | 0.13505521 | 1042 | tags=17%, list | [illegible] |
| GOBP_REGULATION_OF_HORMONE_LEVELS | GOBP_REGULATION_OF_HORMONE_LEVELS | 361 | 0.41702078 | 1.34592228 | 0.00547289 | 0.14576912 | 0.13505521 | 1629 | tags=18%, list | [illegible] |
| GOBP_SMALL_MOLECULE_CATABOLIC_PROCESS | GOBP_SMALL_MOLECULE_CATABOLIC_PROCESS | 273 | 0.44062623 | 1.38842676 | 0.00554401 | 0.14714143 | 0.13632665 | 1868 | tags=23%, list | [illegible] |
| GOBP_SARCOMERE_ORGANIZATION | GOBP_SARCOMERE_ORGANIZATION | 45 | -0.6280366 | -1.5876889 | 0.00565919 | 0.1491445 | 0.1381825 | 2512 | tags=38%, list | [illegible] |
| GOBP_CELLULAR_RESPONSE_TO_ABIOTIC_STIMULUS | GOBP_CELLULAR_RESPONSE_TO_ABIOTIC_STIMULUS | 292 | -0.4144087 | -1.3511961 | 0.00565638 | 0.1491445 | 0.1381825 | 1888 | tags=19%, list | [illegible] |
| GOBP_VENTRICULAR_SYSTEM_DEVELOPMENT | GOBP_VENTRICULAR_SYSTEM_DEVELOPMENT | 29 | -0.7125737 | -1.6580014 | 0.00591113 | 0.15523948 | 0.1438295 | 517 | tags=21%, list | [illegible] |
| GOBP_RESPONSE_TO_BACTERIAL_LIPOPROTEIN | GOBP_RESPONSE_TO_BACTERIAL_LIPOPROTEIN | 7 | 0.92304123 | 1.57961705 | 0.00602034 | 0.15615848 | 0.14468096 | 420 | tags=29%, list | [illegible] |
| GOBP_ENERGY_HOMEOSTASIS | GOBP_ENERGY_HOMEOSTASIS | 71 | 0.58663654 | 1.56525241 | 0.0060241 | 0.15615848 | 0.14468096 | 1517 | tags=20%, list | [illegible] |
| GOBP_FORMATION_OF_PRIMARY_GERM_LAYER | GOBP_FORMATION_OF_PRIMARY_GERM_LAYER | 101 | -0.5295203 | -1.5290063 | 0.00602929 | 0.15615848 | 0.14468096 | 1866 | tags=24%, list | [illegible] |
| GOBP_MITOTIC_CELL_CYCLE_CHECKPOINT_SIGNALING | GOBP_MITOTIC_CELL_CYCLE_CHECKPOINT_SIGNALING | 140 | -0.4917638 | -1.4780663 | 0.00598802 | 0.15615848 | 0.14468096 | 2839 | tags=26%, list | [illegible] |
| GOBP_MONOCARBOXYLIC_ACID_METABOLIC_PROCESS | GOBP_MONOCARBOXYLIC_ACID_METABOLIC_PROCESS | 489 | 0.3963444 | 1.3098725 | 0.00612076 | 0.15798294 | 0.14637132 | 2026 | tags=18%, list | [illegible] |
| GOBP_VACUOLAR_ACIDIFICATION | GOBP_VACUOLAR_ACIDIFICATION | 35 | -0.6751705 | -1.6347517 | 0.0061665 | 0.15861831 | 0.14695999 | 3579 | tags=60%, list | [illegible] |
| GOBP_REGULATION_OF_ADENYLATE_CYCLASE_ACTIVATING_G_PROTEI | GOBP_REGULATION_OF_ADENYLATE_CYCLASE_ACTIVATING_G_PROTEI | 19 | 0.78972674 | 1.65966672 | 0.00626769 | 0.15958179 | 0.14785265 | 1049 | tags=42%, list | [illegible] |
| GOBP_SPONTANEOUS_NEUROTRANSMITTER_SECRETION | GOBP_SPONTANEOUS_NEUROTRANSMITTER_SECRETION | 4 | 0.97494203 | 1.51567246 | 0.00626697 | 0.15958179 | 0.14785265 | 200 | tags=25%, list | [illegible] |
| GOBP_POST_GOLGI_VESICLE_MEDIATED_TRANSPORT | GOBP_POST_GOLGI_VESICLE_MEDIATED_TRANSPORT | 114 | -0.5122496 | -1.5029557 | 0.00626215 | 0.15958179 | 0.14785265 | 3518 | tags=35%, list | [illegible] |
| GOBP_HUMORAL_IMMUNE_RESPONSE_MEDIATED_BY_CIRCULATING_I | GOBP_HUMORAL_IMMUNE_RESPONSE_MEDIATED_BY_CIRCULATING_ | 30 | 0.71896803 | 1.65283916 | 0.00629673 | 0.15977961 | 0.14803594 | 364 | tags=17%, list | [illegible] |
| GOBP_ESTABLISHMENT_OF_PROTEIN_LOCALIZATION | GOBP_ESTABLISHMENT_OF_PROTEIN_LOCALIZATION | 1490 | -0.3207344 | -1.1646245 | 0.00632911 | 0.16006052 | 0.1482962 | 2600 | tags=22%, list | [illegible] |
| GOBP_POSITIVE_REGULATION_OF_TRANSCRIPTION_OF_NOTCH_RECEP | GOBP_POSITIVE_REGULATION_OF_TRANSCRIPTION_OF_NOTCH_RECEP | 7 | 0.92069736 | 1.57560594 | 0.00643554 | 0.16166337 | 0.14978124 | 949 | tags=43%, list | [illegible] |
| GOBP_UREA_CYCLE | GOBP_UREA_CYCLE | 7 | 0.91830823 | 1.57151738 | 0.00643554 | 0.16166337 | 0.14978124 | 257 | tags=29%, list | [illegible] |
| GOBP_FERROPTOSIS | GOBP_FERROPTOSIS | 30 | -0.6999802 | -1.6399764 | 0.00650407 | 0.16229911 | 0.15037025 | 2380 | tags=43%, list | [illegible] |
| GOBP_WHITE_FAT_CELL_PROLIFERATION | GOBP_WHITE_FAT_CELL_PROLIFERATION | 5 | 0.95200891 | 1.53525796 | 0.00650031 | 0.16229911 | 0.15037025 | 360 | tags=60%, list | [illegible] |
| GOBP_EPITHELIUM_DEVELOPMENT | GOBP_EPITHELIUM_DEVELOPMENT | 894 | -0.3423843 | -1.212632 | 0.00664636 | 0.16530065 | 0.15315119 | 1066 | tags=13%, list | [illegible] |
| GOBP_CEREBELLAR_GRANULAR_LAYER_FORMATION | GOBP_CEREBELLAR_GRANULAR_LAYER_FORMATION | 4 | 0.97391056 | 1.51406891 | 0.00668477 | 0.16533669 | 0.15318457 | 143 | tags=50%, list | [illegible] |
| GOBP_LIPID_LOCALIZATION | GOBP_LIPID_LOCALIZATION | 324 | 0.42157127 | 1.35065668 | 0.00669183 | 0.16533669 | 0.15318457 | 736 | tags=10%, list | [illegible] |
| GOBP_INTRACILIARY_RETROGRADE_TRANSPORT | GOBP_INTRACILIARY_RETROGRADE_TRANSPORT | 18 | -0.7752191 | -1.6329233 | 0.00672202 | 0.16553812 | 0.1533712 | 1708 | tags=39%, list | [illegible] |
| GOBP_REGULATION_OF_IMMUNE_SYSTEM_PROCESS | GOBP_REGULATION_OF_IMMUNE_SYSTEM_PROCESS | 1233 | 0.34682103 | 1.20781873 | 0.00676997 | 0.16617404 | 0.15396038 | 1831 | tags=14%, list | [illegible] |
| GOBP_NON_MOTILE_CILIUM_ASSEMBLY | GOBP_NON_MOTILE_CILIUM_ASSEMBLY | 58 | -0.5891536 | -1.557212 | 0.00682594 | 0.16700203 | 0.15472752 | 2810 | tags=41%, list | [illegible] |
| GOBP_OXALOACETATE_METABOLIC_PROCESS | GOBP_OXALOACETATE_METABOLIC_PROCESS | 10 | 0.87387764 | 1.60715582 | 0.00685216 | 0.16709925 | 0.15481759 | 197 | tags=20%, list | [illegible] |
| GOBP_GOLGI_VESICLE_TRANSPORT | GOBP_GOLGI_VESICLE_TRANSPORT | 301 | -0.4102589 | -1.3402535 | 0.00688673 | 0.16739868 | 0.15509501 | 3836 | tags=31%, list | [illegible] |
| GOBP_REGULATION_OF_LONG_TERM_SYNAPTIC_DEPRESSION | GOBP_REGULATION_OF_LONG_TERM_SYNAPTIC_DEPRESSION | 13 | 0.83787851 | 1.62867694 | 0.00714432 | 0.17211689 | 0.15946644 | 841 | tags=31%, list | [illegible] |
| GOBP_REGULATION_OF_TRIGLYCERIDE_CATABOLIC_PROCESS | GOBP_REGULATION_OF_TRIGLYCERIDE_CATABOLIC_PROCESS | 11 | 0.86166396 | 1.62117991 | 0.00714577 | 0.17211689 | 0.15946644 | 1218 | tags=45%, list | [illegible] |
| GOBP_REGULATION_OF_PLASMA_LIPOPROTEIN_PARTICLE_LEVELS | GOBP_REGULATION_OF_PLASMA_LIPOPROTEIN_PARTICLE_LEVELS | 60 | 0.59930148 | 1.55708058 | 0.00714958 | 0.17211689 | 0.15946644 | 730 | tags=20%, list | [illegible] |
| GOBP_POSITIVE_REGULATION_OF_OSTEOCLAST_DIFFERENTIATION | GOBP_POSITIVE_REGULATION_OF_OSTEOCLAST_DIFFERENTIATION | 19 | 0.78473497 | 1.64917617 | 0.00727861 | 0.17325005 | 0.1605163 | 754 | tags=16%, list | [illegible] |
| GOBP_RESPONSE_TO_DIACYL_BACTERIAL_LIPOPEPTIDE | GOBP_RESPONSE_TO_DIACYL_BACTERIAL_LIPOPEPTIDE | 4 | 0.97292788 | 1.5125412 | 0.00731147 | 0.17325005 | 0.1605163 | 238 | tags=25%, list | [illegible] |
| GOBP_DIACYLGLYCEROL_CATABOLIC_PROCESS | GOBP_DIACYLGLYCEROL_CATABOLIC_PROCESS | 4 | 0.97270151 | 1.51218927 | 0.00731147 | 0.17325005 | 0.1605163 | 45 | tags=25%, list | [illegible] |
| GOBP_THIRD_VENTRICLE_DEVELOPMENT | GOBP_THIRD_VENTRICLE_DEVELOPMENT | 4 | -0.9597916 | -1.4572235 | 0.00728667 | 0.17325005 | 0.1605163 | 133 | tags=50%, list | [illegible] |
| GOBP_T_CELL_DIFFERENTIATION | GOBP_T_CELL_DIFFERENTIATION | 270 | 0.43747978 | 1.37676196 | 0.00731198 | 0.17325005 | 0.1605163 | 1478 | tags=16%, list | [illegible] |
| GOBP_LONG_TERM_SYNAPTIC_DEPRESSION | GOBP_LONG_TERM_SYNAPTIC_DEPRESSION | 25 | 0.74023416 | 1.64482086 | 0.00736758 | 0.17347308 | 0.16072295 | 867 | tags=28%, list | [illegible] |
| GOBP_ADAPTIVE_IMMUNE_RESPONSE | GOBP_ADAPTIVE_IMMUNE_RESPONSE | 346 | 0.41497095 | 1.3350123 | 0.0073668 | 0.17347308 | 0.16072295 | 1372 | tags=12%, list | [illegible] |
| GOBP_CELLULAR_RESPONSE_TO_EPINEPHRINE_STIMULUS | GOBP_CELLULAR_RESPONSE_TO_EPINEPHRINE_STIMULUS | 10 | 0.87295937 | 1.60546702 | 0.00747508 | 0.17436444 | 0.16154879 | 713 | tags=40%, list | [illegible] |
| GOBP_ENDODERMAL_CELL_DIFFERENTIATION | GOBP_ENDODERMAL_CELL_DIFFERENTIATION | 39 | -0.6469925 | -1.5972673 | 0.00746114 | 0.17436444 | 0.16154879 | 1689 | tags=33%, list | [illegible] |
| GOBP_REPRODUCTIVE_SYSTEM_DEVELOPMENT | GOBP_REPRODUCTIVE_SYSTEM_DEVELOPMENT | 226 | 0.44972471 | 1.39119251 | 0.00744153 | 0.17436444 | 0.16154879 | 1307 | tags=18%, list | [illegible] |
| GOBP_LIPID_HOMEOSTASIS | GOBP_LIPID_HOMEOSTASIS | 138 | 0.49818361 | 1.45663283 | 0.00764192 | 0.17770425 | 0.16464312 | 1164 | tags=16%, list | [illegible] |
| GOBP_CIRCADIAN_SLEEP_WAKE_CYCLE_NON_REM_SLEEP | GOBP_CIRCADIAN_SLEEP_WAKE_CYCLE_NON_REM_SLEEP | 3 | 0.98139055 | 1.45430689 | 0.00776169 | 0.17939067 | 0.1662056 | 91 | tags=33%, list | [illegible] |
| GOBP_BILE_ACID_SECRETION | GOBP_BILE_ACID_SECRETION | 2 | -0.9886799 | -1.3641091 | 0.00776221 | 0.17939067 | 0.1662056 | 152 | tags=100%, lis | [illegible] |
| GOBP_MESENCHYMAL_CELL_MIGRATION | GOBP_MESENCHYMAL_CELL_MIGRATION | 52 | 0.61727026 | 1.56042821 | 0.0078304 | 0.18041157 | 0.16715146 | 2053 | tags=31%, list | [illegible] |
| GOBP_HORMONE_TRANSPORT | GOBP_HORMONE_TRANSPORT | 223 | 0.44785948 | 1.38177922 | 0.00785574 | 0.18044178 | 0.16717945 | 1036 | tags=15%, list | [illegible] |
| GOBP_PEPTIDE_CROSS_LINKING | GOBP_PEPTIDE_CROSS_LINKING | 7 | -0.9025505 | -1.539826 | 0.00790743 | 0.18107522 | 0.16776633 | 65 | tags=43%, list | [illegible] |
| GOBP_NEGATIVE_REGULATION_OF_BLOOD_PRESSURE | GOBP_NEGATIVE_REGULATION_OF_BLOOD_PRESSURE | 27 | 0.73089138 | 1.64924277 | 0.00794439 | 0.18136872 | 0.16803826 | 1042 | tags=26%, list | [illegible] |
| GOBP_NUCLEOSIDE_TRANSPORT | GOBP_NUCLEOSIDE_TRANSPORT | 6 | 0.93212844 | 1.55438555 | 0.00809297 | 0.1829089 | 0.16946524 | 91 | tags=33%, list | [illegible] |
| GOBP_NEGATIVE_REGULATION_OF_TRIGLYCERIDE_CATABOLIC_PROCES | GOBP_NEGATIVE_REGULATION_OF_TRIGLYCERIDE_CATABOLIC_PROCE | 6 | 0.93172125 | 1.55370654 | 0.00809297 | 0.1829089 | 0.16946524 | 708 | tags=50%, list | [illegible] |
| GOBP_POSITIVE_REGULATION_OF_NF_KAPPAB_TRANSCRIPTION_FACTI | GOBP_POSITIVE_REGULATION_OF_NF_KAPPAB_TRANSCRIPTION_FACT | 64 | -0.5674968 | -1.5261153 | 0.00810926 | 0.1829089 | 0.16946524 | 2425 | tags=22%, list | [illegible] |
| GOBP_CELL_CYCLE | GOBP_CELL_CYCLE | 1455 | -0.3185335 | -1.1564013 | 0.00804954 | 0.1829089 | 0.16946524 | 2888 | tags=22%, list | [illegible] |
| GOBP_INTRACELLULAR_TRIGLYCERIDE_HOMEOSTASIS | GOBP_INTRACELLULAR_TRIGLYCERIDE_HOMEOSTASIS | 9 | 0.88848223 | 1.59796988 | 0.00829703 | 0.18589744 | 0.17223412 | 39 | tags=22%, list | [illegible] |
| GOBP_POSITIVE_REGULATION_OF_MYELOID_CELL_APOPTOTIC_PROCE | GOBP_POSITIVE_REGULATION_OF_MYELOID_CELL_APOPTOTIC_PROCE | 8 | 0.89736728 | 1.57588742 | 0.00831601 | 0.18589744 | 0.17223412 | 2 | tags=12%, list | [illegible] |
| GOBP_D_GLUCOSE_IMPORT | GOBP_D_GLUCOSE_IMPORT | 59 | 0.59709269 | 1.54632864 | 0.00830502 | 0.18589744 | 0.17223412 | 747 | tags=19%, list | [illegible] |
| GOBP_G_PROTEIN_COUPLED_RECEPTOR_SIGNALING_PATHWAY_COUI | GOBP_G_PROTEIN_COUPLED_RECEPTOR_SIGNALING_PATHWAY_COUI | 17 | 0.79239405 | 1.62605987 | 0.00835541 | 0.186224 | 0.17253669 | 1452 | tags=41%, list | [illegible] |
| GOBP_BIOLOGICAL_PROCESS_INVOLVED_IN_INTERACTION_WITH_HOS | GOBP_BIOLOGICAL_PROCESS_INVOLVED_IN_INTERACTION_WITH_HOS | 139 | -0.4772667 | -1.4326374 | 0.0084332 | 0.1869988 | 0.17325454 | 2627 | tags=28%, list | [illegible] |
| GOBP_PROTEIN_LOCALIZATION_TO_ORGANELLE | GOBP_PROTEIN_LOCALIZATION_TO_ORGANELLE | 809 | -0.346287 | -1.2185329 | 0.00843997 | 0.1869988 | 0.17325454 | 3426 | tags=26%, list | [illegible] |
| GOBP_CEREBELLAR_GRANULAR_LAYER_DEVELOPMENT | GOBP_CEREBELLAR_GRANULAR_LAYER_DEVELOPMENT | 9 | 0.88745185 | 1.5961167 | 0.00850446 | 0.18741175 | 0.17363713 | 169 | tags=33%, list | [illegible] |
| GOBP_SMALL_MOLECULE_BIOSYNTHETIC_PROCESS | GOBP_SMALL_MOLECULE_BIOSYNTHETIC_PROCESS | 461 | 0.39405243 | 1.29642198 | 0.00850851 | 0.18741175 | 0.17363713 | 1387 | tags=13%, list | [illegible] |
| GOBP_ALDITOL_BIOSYNTHETIC_PROCESS | GOBP_ALDITOL_BIOSYNTHETIC_PROCESS | 5 | 0.94501575 | 1.52398043 | 0.00859719 | 0.18881139 | 0.1749339 | 263 | tags=40%, list | [illegible] |
| GOBP_VASCULOGENESIS | GOBP_VASCULOGENESIS | 75 | 0.56573495 | 1.52322396 | 0.00865638 | 0.189006 | 0.17511421 | 1495 | tags=24%, list | [illegible] |
| GOBP_NEGATIVE_REGULATION_OF_MITOTIC_CELL_CYCLE | GOBP_NEGATIVE_REGULATION_OF_MITOTIC_CELL_CYCLE | 216 | -0.4240208 | -1.3433179 | 0.00865013 | 0.189006 | 0.17511421 | 1943 | tags=17%, list | [illegible] |
| GOBP_CARDIAC_MUSCLE_CELL_CARDIAC_MUSCLE_CELL_ADHESION | GOBP_CARDIAC_MUSCLE_CELL_CARDIAC_MUSCLE_CELL_ADHESION | 6 | -0.922193 | -1.517179 | 0.00868223 | 0.18901889 | 0.17512615 | 761 | tags=67%, list | [illegible] |
| GOBP_REGULATION_OF_INTRINSIC_APOPTOTIC_SIGNALING_PATHWAY | GOBP_REGULATION_OF_INTRINSIC_APOPTOTIC_SIGNALING_PATHWA | 167 | -0.4543312 | -1.3970188 | 0.0087073 | 0.18901889 | 0.17512615 | 3321 | tags=33%, list | [illegible] |

| | | | | | | | | | | |
|---|---|---|---|---|---|---|---|---|---|---|
| GOBP_MITOTIC_CELL_CYCLE | GOBP_MITOTIC_CELL_CYCLE | 839 | -0.3444449 | -1.2149675 | 0.00877915 | 0.19002937 | 0.17606236 | 3042 | tags=24%, list | SMC3/TMOD3/PTPA/DYNLRB1/CENPI/PCNA/BID/HSPA1B/DAPK3/TMEM14B/NAT9/VPS4B/RHOC/PSMB1/HES1/SMARCA4/FBXW11/CCND1/ERCC3/DDX3X/BBS4/KNL1/NUSAP1/MCM2/DNM2/GINS1/CKAP2/CDC27/STIL/TADA3/NFE2L1/DONS [illegible] |
| GOBP_ALANINE_METABOLIC_PROCESS | GOBP_ALANINE_METABOLIC_PROCESS | 2 | 0.99021839 | 1.3813486 | 0.00889831 | 0.19095763 | 0.17692239 | 46 | tags=50%, list | GPT |
| GOBP_MESENCHYME_DEVELOPMENT | GOBP_MESENCHYME_DEVELOPMENT | 262 | 0.43596249 | 1.36830674 | 0.00888811 | 0.19095763 | 0.17692239 | 1817 | tags=21%, list | RET/AGT/TCF15/MEOX1/SEMA6A/FGF12/SPRY2/GSC/OLFM1/PRICKLE1/IGF1/HOXA5/EFNB1/MDM4/BCL2/EDNRA/NOTCH4/SMO/MDM2/CITED2/SEMA3G/TIAM1/ROBO2/FRZB/EDNRB/CUL7/SEMA3B/NCAM1/QKI/SEMA4A/SFRP1/EOMES/SEMA [illegible] |
| GOBP_ENDOMEMBRANE_SYSTEM_ORGANIZATION | GOBP_ENDOMEMBRANE_SYSTEM_ORGANIZATION | 520 | -0.369216 | -1.26256 | 0.00888776 | 0.19095763 | 0.17692239 | 3378 | tags=29%, list | LPCAT3/ARL8B/BLOC1S2/SERINC5/RAB8A/TRAPPC8/GET1/TRAM2/LMNB2/VMP1/DMKN/SNF8/LMNA/EMC9/ALS2/CHMP5/SLC9A8/CSNK1A1/EMC6/SQSTM1/RAB22A/MYO5A/RAB6B/CCDC47/CLN3/TPST2/HGS/TMF1/SNAP23/TARDBP/STX2/ [illegible] |
| GOBP_BRAINSTEM_DEVELOPMENT | GOBP_BRAINSTEM_DEVELOPMENT | 7 | 0.90879848 | 1.55524316 | 0.00892672 | 0.19102159 | 0.17698165 | 8 | tags=14%, list | CNTFR |
| GOBP_REGULATION_OF_D_GLUCOSE_IMPORT | GOBP_REGULATION_OF_D_GLUCOSE_IMPORT | 43 | 0.64994702 | 1.59195706 | 0.00901765 | 0.19241926 | 0.1782766 | 747 | tags=21%, list | ADIPOQ/KLF15/LEP/IGF1/IRS2/SORBS1/GPC3/PRKAG2/PIK3R1 |
| GOBP_CARBOHYDRATE_METABOLIC_PROCESS | GOBP_CARBOHYDRATE_METABOLIC_PROCESS | 450 | 0.39561064 | 1.29861615 | 0.00908021 | 0.19320523 | 0.1790048 | 1375 | tags=13%, list | ADIPOQ/RBP4/NPY1R/DGAT2/GPD1/PCK1/KLB/LEPR/ACACB/LRP5/PDK4/LEP/KAT2A/GYG2/SLC2A4/PC/B3GAT1/IGF1/GALT/KL/IRS2/IDH1/PGGHG/FBP2/SORBS1/PPP1R1A/PRKAG2/PLA2G4A/SLC25A10/MLXIPL/PHKA1/ADCYAP1R1/GPLD1/ [illegible] |
| GOBP_REGULATION_OF_LIPID_STORAGE | GOBP_REGULATION_OF_LIPID_STORAGE | 36 | 0.68120853 | 1.61516958 | 0.00914931 | 0.19412559 | 0.17985751 | 279 | tags=25%, list | PLIN5/PNPLA2/MEST/PPARG/ACACB/NR1H3/CD36/LEP/LPL |
| GOBP_NEURON_CELL_CELL_ADHESION | GOBP_NEURON_CELL_CELL_ADHESION | 14 | 0.8222596 | 1.6195395 | 0.00921753 | 0.19447443 | 0.18018071 | 659 | tags=21%, list | RET/NCAM2/NRXN2 |
| GOBP_RESPONSE_TO_ACTIVITY | GOBP_RESPONSE_TO_ACTIVITY | 66 | 0.58455881 | 1.54206097 | 0.00921746 | 0.19447443 | 0.18018071 | 926 | tags=18%, list | ADIPOQ/RBP4/AGT/RYR2/BMP6/FOS/LEP/KL/OXT/KDM6B/ANGPT2/COL4A2 |
| GOBP_NEGATIVE_REGULATION_OF_TYPE_II_INTERFERON_PRODUCTIO | GOBP_NEGATIVE_REGULATION_OF_TYPE_II_INTERFERON_PRODUCTIO | 26 | -0.7092158 | -1.6138328 | 0.00924437 | 0.19449436 | 0.18019917 | 767 | tags=23%, list | TNFSF4/AXL/IL20RA/ZC3H12A/INHBA/IL20RB |
| GOBP_CELLULAR_RESPONSE_TO_METAL_ION | GOBP_CELLULAR_RESPONSE_TO_METAL_ION | 132 | 0.49913339 | 1.45031823 | 0.00932529 | 0.19537802 | 0.18101789 | 1245 | tags=17%, list | FABP4/FOSB/MT1A/BMP6/FOS/CEBPA/NFATC2/SYT1/ITPKB/RYR3/ABCB6/RASGRP2/CPNE5/CPNE7/PRND/AQP1/CAMK2D/JUN/SLC25A23/HESX1/ASPH/INHBB |
| GOBP_ANIMAL_ORGAN_MORPHOGENESIS | GOBP_ANIMAL_ORGAN_MORPHOGENESIS | 785 | 0.36046253 | 1.22603834 | 0.0093384 | 0.19537802 | 0.18101789 | 1836 | tags=19%, list | RBP4/NPY1R/CLDN5/HOXC11/MDFI/ADAMTS19/AGT/TCF15/ADAMTS9/HOXA4/NDRG4/RYR2/AR/HOXA3/RFLNA/COL18A1/BMP6/NFIB/CSMD1/SEMA6A/LHX9/MGP/LRP5/DIO3/FST/COL2A1/PTN/SPRY2/GSC/PDZD7/HOXA2/THRA/SMARCD3/ [illegible] |
| GOBP_REGULATION_OF_SKELETAL_MUSCLE_CELL_PROLIFERATION | GOBP_REGULATION_OF_SKELETAL_MUSCLE_CELL_PROLIFERATION | 12 | 0.83838039 | 1.60374336 | 0.00936864 | 0.19546617 | 0.18109956 | 438 | tags=17%, list | ANGPT1/EPHB1 |
| GOBP_PROTEIN_TRANSPORT | GOBP_PROTEIN_TRANSPORT | 1237 | -0.3241886 | -1.1687328 | 0.00954939 | 0.19868547 | 0.18408224 | 2600 | tags=22%, list | SCAMP2/PFKM/PPP1R10/PIK3R2/TAP1/PSMD9/STX1A/TUBA1A/STX8/RAB10/B2M/CHMP2B/UEVLD/ANKLE1/ALDOA/P2RX7/BCAP29/CHMP2A/IFT25/GOPC/WASHC2C/CHMP3/ARHGEF2/GOSR2/YIF1A/ACSL4/ARF3/OPTN/KPNA6/ARRDC4/SPC [illegible] |
| GOBP_OVULATION_CYCLE | GOBP_OVULATION_CYCLE | 49 | 0.6252076 | 1.56045945 | 0.00962649 | 0.19973641 | 0.18505594 | 1320 | tags=29%, list | FZD4/OPRK1/GABRB1/PTX3/MMP19/LEP/PTN/EGR1/NOTCH4/ROBO2/TYRO3/SLIT3/STAT5B/HAS2 |
| GOBP_RETINA_VASCULATURE_DEVELOPMENT_IN_CAMERA_TYPE_EYE | GOBP_RETINA_VASCULATURE_DEVELOPMENT_IN_CAMERA_TYPE_EYE | 19 | 0.77507548 | 1.62887607 | 0.00970481 | 0.20080673 | 0.18604759 | 1351 | tags=32%, list | FZD4/ARHGEF15/LRP5/LAMA1/COL4A1/ACVR2B |
| GOBP_DISTAL_CONVOLUTED_TUBULE_DEVELOPMENT | GOBP_DISTAL_CONVOLUTED_TUBULE_DEVELOPMENT | 2 | 0.98942368 | 1.38023998 | 0.00974576 | 0.20110007 | 0.18631937 | 76 | tags=50%, list | POU3F3 |
| GOBP_MANCHETTE_ASSEMBLY | GOBP_MANCHETTE_ASSEMBLY | 9 | -0.8660038 | -1.5628883 | 0.00984366 | 0.20201018 | 0.18716259 | 1688 | tags=67%, list | SPAG17/HOOK1/CCDC181/IFT56/CCDC146/CFAP53 |
| GOBP_POSITIVE_REGULATION_OF_PLASMINOGEN_ACTIVATION | GOBP_POSITIVE_REGULATION_OF_PLASMINOGEN_ACTIVATION | 7 | -0.8986028 | -1.5330909 | 0.00983607 | 0.20201018 | 0.18716259 | 1381 | tags=71%, list | MELTF/PLGRKT/ENO1/S100A10/ANXA2 |
| GOBP_MITOTIC_SISTER_CHROMATID_SEGREGATION | GOBP_MITOTIC_SISTER_CHROMATID_SEGREGATION | 196 | -0.4365822 | -1.3687499 | 0.0099728 | 0.20410276 | 0.18910136 | 3835 | tags=37%, list | CEP55/SPC24/CENPK/WAPL/BAZ1B/MAPRE1/ANAPC7/SPDL1/KIF18B/SPAG5/NCAPG2/NUP62/RACGAP1/MAD2L2/CDC16/CHMP5/BOD1/SMC3/DYNLRB1/CENPI/HSPA1B/NAT9/VPS4B/KNL1/NUSAP1/MAD2L1BP/SKA3/CLASP1/ANAPC11/CH [illegible] |
| GOBP_POSITIVE_REGULATION_OF_EPITHELIAL_CELL_DIFFERENTIATION | GOBP_POSITIVE_REGULATION_OF_EPITHELIAL_CELL_DIFFERENTIATION | 45 | -0.6099219 | -1.5418947 | 0.01006078 | 0.20534388 | 0.19025127 | 832 | tags=27%, list | VEZF1/RFX3/VDR/PLAAT4/FOXC1/CD24/LEF1/BMP2/TP73/FOXJ1/ETV4/SFN |
| GOBP_NEGATIVE_REGULATION_OF_LONG_TERM_SYNAPTIC_POTENTIA | GOBP_NEGATIVE_REGULATION_OF_LONG_TERM_SYNAPTIC_POTENTIA | 12 | 0.83680725 | 1.60073409 | 0.0101833 | 0.20616378 | 0.1910109 | 418 | tags=25%, list | ADORA1/PTN/FAM107A |
| GOBP_PROTEIN_DEPOLYMERIZATION | GOBP_PROTEIN_DEPOLYMERIZATION | 108 | -0.503372 | -1.4650523 | 0.01013149 | 0.20616378 | 0.1910109 | 2595 | tags=33%, list | SLN/PPP1R9B/CAPG/CLASP1/WASHC2C/ARHGEF2/SWAP70/CAMSAP2/HDGFL3/SYNJ1/CKAP5/CAPZB/WDR47/ACTN2/KATNB1/CLTC/CARMIL1/LMOD2/TMOD4/TRIM54/KIF21A/KIF24/TWF1/RDX/MTPN/WDR1/CAPZA1/CIB1/TRPV4/DSTN/KIF [illegible] |
| GOBP_SIGNAL_TRANSDUCTION_BY_P53_CLASS_MEDIATOR | GOBP_SIGNAL_TRANSDUCTION_BY_P53_CLASS_MEDIATOR | 184 | -0.4431853 | -1.3791295 | 0.0101626 | 0.20616378 | 0.1910109 | 1909 | tags=20%, list | PSMD10/CDKN2A/COPS3/ZMPSTE24/ING2/SESN2/CDKN1A/ANKRD1/TP53BP1/RRS1/SMYD2/PYCARD/TMEM109/PMAIP1/MORN3/RNF34/CD44/PTTG1IP/USP2/ZNF385B/IFI16/MYO6/URI1/GREM1/KMT5A/HIPK1/TRIAP1/RPS27L/MUC1/BDK [illegible] |
| GOBP_L_ASCORBIC_ACID_METABOLIC_PROCESS | GOBP_L_ASCORBIC_ACID_METABOLIC_PROCESS | 10 | -0.8541946 | -1.5791098 | 0.01021982 | 0.20634701 | 0.19118067 | 559 | tags=30%, list | GSTO2/GSTO1/SLC2A1 |
| GOBP_REGULATION_OF_DENDRITE_MORPHOGENESIS | GOBP_REGULATION_OF_DENDRITE_MORPHOGENESIS | 50 | 0.61925141 | 1.5519814 | 0.01038661 | 0.20915241 | 0.19377987 | 1104 | tags=14%, list | FZD4/CAMK2B/OBSL1/PPP3CA/RAB21/CUL7/ANAPC2 |
| GOBP_INTERCELLULAR_TRANSPORT | GOBP_INTERCELLULAR_TRANSPORT | 3 | 0.97861239 | 1.45019 | 0.01048878 | 0.21008321 | 0.19464226 | 7 | tags=33%, list | PEG10 |
| GOBP_NUCLEAR_RECEPTOR_MEDIATED_SIGNALING_PATHWAY | GOBP_NUCLEAR_RECEPTOR_MEDIATED_SIGNALING_PATHWAY | 176 | 0.46683833 | 1.40190444 | 0.01046742 | 0.21008321 | 0.19464226 | 2288 | tags=23%, list | PLIN5/RXRG/AR/PPARG/ESR2/PDE3A/LEP/THRA/IGF1/GREB1L/LMO3/CYP27B1/CITED2/PRAME/ZNF366/PGRMC2/KDM5D/PER1/SFRP1/ASXL1/CYP26B1/CYP7B1/PIAS2/RARA/RXRA/KDM1A/LATS1/PHB2/PPARA/TBX1/ESR1/THRB/NR1H2/TCF [illegible] |
| GOBP_HYPOTONIC_RESPONSE | GOBP_HYPOTONIC_RESPONSE | 13 | -0.8132238 | -1.5939621 | 0.01058201 | 0.21138692 | 0.19585015 | 1423 | tags=46%, list | STK39/SLC12A6/AQP5/TRPV4/ITGA2/SLC4A11 |
| GOBP_SMOOTH_MUSCLE_CELL_DIFFERENTIATION | GOBP_SMOOTH_MUSCLE_CELL_DIFFERENTIATION | 60 | 0.5883746 | 1.52869081 | 0.01072437 | 0.2130971 | 0.19743463 | 1561 | tags=30%, list | NFATC2/SMARCD3/RAMP2/EDNRA/NOTCH4/COMP/KIT/EDNRB/MRTFB/QKI/ITGA8/NFATC1/ENG/EFEMP2/RBPMS2/APLNR/PDCD4/GATA6 |
| GOBP_REGULATION_OF_AXONOGENESIS | GOBP_REGULATION_OF_AXONOGENESIS | 119 | 0.50924213 | 1.46016504 | 0.01071297 | 0.2130971 | 0.19743463 | 1817 | tags=22%, list | RET/BRSK2/CHODL/NEFL/TIAM2/EFNB3/CXCL12/SEMA3G/TIAM1/ROBO2/PLXND1/GDI1/TRAK2/ARHGAP4/ANAPC2/NTRK2/PLXNA3/THY1/EPHB3/PAK3/LRRC4C/EFNA1/TRIM46/EPHA7/ZFYVE27/ROBO1 |
| GOBP_MULTICELLULAR_ORGANISM_GROWTH | GOBP_MULTICELLULAR_ORGANISM_GROWTH | 146 | 0.48392513 | 1.42011091 | 0.01078611 | 0.21375842 | 0.19804735 | 1938 | tags=23%, list | NPY1R/CDKN1C/AR/CACNA2D2/DIO3/KAT2A/PRICKLE1/IGF1/HOXA5/ATN1/BCL2/FLVCR1/ATF5/SMO/CTC1/COMP/ARID5B/FXN/ADRB1/HESX1/RARA/STAT5B/COL3A1/GHR/GNAS/PTCH1/ATM/HSF1/FOXS1/SLITRK6/APBA1/EXT1/ADARB1 |
| GOBP_BINDING_OF_SPERM_TO_ZONA_PELLUCIDA | GOBP_BINDING_OF_SPERM_TO_ZONA_PELLUCIDA | 19 | -0.7527723 | -1.6013182 | 0.01087816 | 0.21445117 | 0.19868917 | 706 | tags=26%, list | CCT8/CCT5/PCSK4/CLGN/SPA17 |
| GOBP_ENERGY_RESERVE_METABOLIC_PROCESS | GOBP_ENERGY_RESERVE_METABOLIC_PROCESS | 82 | 0.54936142 | 1.49962021 | 0.0108716 | 0.21445117 | 0.19868917 | 1158 | tags=18%, list | LEPR/LEP/GYG2/IGF1/KL/IRS2/SORBS1/PPP1R1A/PRKAG2/PHKA1/AKR1E2/PYGL/ADGRF5/PGM1/EPM2A |
| GOBP_AUTOCRINE_SIGNALING | GOBP_AUTOCRINE_SIGNALING | 9 | -0.8610768 | -1.5539964 | 0.01100174 | 0.2163195 | 0.20042018 | 74 | tags=44%, list | FZD1/CX3CR1/S100A8/S100A9 |
| GOBP_EXCITATORY_SYNAPSE_ASSEMBLY | GOBP_EXCITATORY_SYNAPSE_ASSEMBLY | 36 | 0.67461837 | 1.59954408 | 0.01109597 | 0.2176027 | 0.20160907 | 925 | tags=28%, list | CBLN1/GRID2/LRRTM2/CASKIN1/PRICKLE1/LRFN1/LRRC4B/NRXN2/SHANK3/SEMA4A |
| GOBP_FERTILIZATION | GOBP_FERTILIZATION | 112 | -0.4967117 | -1.4524295 | 0.01125785 | 0.22020233 | 0.20401764 | 2013 | tags=21%, list | MYH9/CCNO/UBE2Q1/REC8/RAB3A/MFGE8/ANTXR2/PARK7/YBX3/IFTAP/CCT8/CFAP119/CCT5/PCSK4/CLGN/CD9/SPAG1/SPA17/BBOF1/CCDC146/SPAG8/LRRIQ1/WDR54 |
| GOBP_SUPRAMOLECULAR_FIBER_ORGANIZATION | GOBP_SUPRAMOLECULAR_FIBER_ORGANIZATION | 677 | -0.3514726 | -1.222818 | 0.01140281 | 0.22245847 | 0.20610795 | 2588 | tags=22%, list | CDC42/PPP1R9B/PIK3R2/AGFG1/TUBA1A/CAPG/B2M/MYPN/CLASP1/TOGARAM1/ALDOA/DIAPH1/WASHC2C/MYL2/ARHGEF2/ACTC1/AKAP9/TUBG1/MYBPC1/SWAP70/PPFIBP1/ACTA1/RND2/CDKN2A/KATNBL1/ACTG1/CYRIB/HSPA1A/CAMS [illegible] |
| GOBP_MUCOCILIARY_CLEARANCE | GOBP_MUCOCILIARY_CLEARANCE | 9 | -0.8580809 | -1.5485898 | 0.01158078 | 0.2253451 | 0.20878241 | 1075 | tags=67%, list | VANGL1/SPAG16/ODAD4/DNAH9/NEK10/CFAP43 |
| GOBP_SKELETAL_MUSCLE_CELL_PROLIFERATION | GOBP_SKELETAL_MUSCLE_CELL_PROLIFERATION | 17 | 0.78588997 | 1.61271295 | 0.01161606 | 0.22544759 | 0.20887737 | 438 | tags=18%, list | ANGPT1/FOS/EPHB1 |
| GOBP_POSITIVE_REGULATION_OF_RECEPTOR_RECYCLING | GOBP_POSITIVE_REGULATION_OF_RECEPTOR_RECYCLING | 13 | -0.8092899 | -1.5862515 | 0.01175779 | 0.22702508 | 0.21033892 | 1383 | tags=46%, list | NSF/PSEN1/NSG1/INPP5F/ECE1/ANXA2 |
| GOBP_ESTABLISHMENT_OF_LEFT_RIGHT_ASYMMETRY | GOBP_ESTABLISHMENT_OF_LEFT_RIGHT_ASYMMETRY | 2 | -0.9847018 | -1.3586203 | 0.01173798 | 0.22702508 | 0.21033892 | 61 | tags=100%, list | PIERCE1 |
| GOBP_NEUTROPHIL_CHEMOTAXIS | GOBP_NEUTROPHIL_CHEMOTAXIS | 62 | -0.5620846 | -1.5026782 | 0.01180004 | 0.22725672 | 0.21055353 | 381 | tags=11%, list | SAA1/TGFB2/CXADR/LGALS3/S100A8/S100A9/PERP |
| GOBP_PROTEIN_CONTAINING_COMPLEX_REMODELING | GOBP_PROTEIN_CONTAINING_COMPLEX_REMODELING | 22 | 0.75249872 | 1.62725229 | 0.01186407 | 0.22728712 | 0.21058169 | 576 | tags=23%, list | AGT/GPIHBP1/LPL/ANGPTL4/LCAT |
| GOBP_NEGATIVE_REGULATION_OF_FERROPTOSIS | GOBP_NEGATIVE_REGULATION_OF_FERROPTOSIS | 19 | -0.7483545 | -1.5919205 | 0.01186709 | 0.22728712 | 0.21058169 | 1924 | tags=47%, list | CAMK2A/SLC7A11/ADGRG1/PRKAA1/NFE2L2/GSTP1/SC5D/NQO1/FTH1 |
| GOBP_BUNDLE_OF_HIS_CELL_TO_PURKINJE_MYOCYTE_COMMUNICAT | GOBP_BUNDLE_OF_HIS_CELL_TO_PURKINJE_MYOCYTE_COMMUNICAT | 11 | -0.8339245 | -1.5715205 | 0.01195141 | 0.22728712 | 0.21058169 | 761 | tags=27%, list | DSC2/DSG2/DSP |
| GOBP_STEROL_METABOLIC_PROCESS | GOBP_STEROL_METABOLIC_PROCESS | 122 | 0.50467055 | 1.45316829 | 0.01195292 | 0.22728712 | 0.21058169 | 1387 | tags=18%, list | DGAT2/LIPE/AGT/LEPR/CEBPA/LRP5/SCARF1/LEP/ACADVL/ABCA2/LCAT/PRKAG2/ACAA2/EBPL/VLDLR/QKI/CYP46A1/CCDC3/CYP7B1/GPR146/GBA2/MVD |
| GOBP_REGULATION_OF_MEMBRANE_POTENTIAL | GOBP_REGULATION_OF_MEMBRANE_POTENTIAL | 297 | 0.4186247 | 1.33037766 | 0.01193978 | 0.22728712 | 0.21058169 | 1121 | tags=15%, list | AGT/RYR2/GABRB1/CBLN1/ADORA1/MYOC/GRID2/BEGAIN/GRIA2/PRKAR1B/SYT1/KCNMB4/GABBR1/CD36/ABCC8/CACNA1C/FGF12/SCN4A/SLC1A7/KCNJ3/NRCAM/GRIN2B/SLC4A3/KCNIP2/ASIC2/FXYD1/GABRA3/CLCN1/BCL2/NOS1/GRIN2 [illegible] |
| GOBP_REGULATION_OF_HEMOPOIESIS | GOBP_REGULATION_OF_HEMOPOIESIS | 357 | 0.40307601 | 1.29943016 | 0.01200686 | 0.2277362 | 0.21099776 | 1762 | tags=17%, list | ADIPOQ/DLK1/ZBTB16/PCK1/EGR3/TMEM131L/FOS/CCL19/CD83/PRXL2A/IL7R/ITPKB/PTN/KAT2A/TESPA1/SMARCD3/HOXA5/ZAP70/BRD1/IRF4/IRF7/PRDM1/CTLA4/MEIS2/LCK/PIK3R1/PPP3CA/ACIN1/ZBTB46/QKI/TAL1/SFRP1/NFAM1/DTX [illegible] |
| GOBP_CHEMICAL_SYNAPTIC_TRANSMISSION_POSTSYNAPTIC | GOBP_CHEMICAL_SYNAPTIC_TRANSMISSION_POSTSYNAPTIC | 69 | 0.56973873 | 1.51171035 | 0.01204593 | 0.22790162 | 0.21115103 | 841 | tags=17%, list | CBLN1/ADORA1/GRID2/BEGAIN/PRKAR1B/SYT1/SLC1A7/GRIN2B/GRIN2C/PPP3CA/GRIN2A/SHANK3 |
| GOBP_TERPENOID_BIOSYNTHETIC_PROCESS | GOBP_TERPENOID_BIOSYNTHETIC_PROCESS | 21 | -0.7346342 | -1.5947677 | 0.01215866 | 0.22945653 | 0.21259166 | 1045 | tags=29%, list | RBP1/RDH10/ALDH1A3/MVK/DPM3/DHRS9 |
| GOBP_REGULATION_OF_DEVELOPMENTAL_GROWTH | GOBP_REGULATION_OF_DEVELOPMENTAL_GROWTH | 243 | 0.43449256 | 1.35300981 | 0.01222971 | 0.23021885 | 0.21329794 | 1801 | tags=20%, list | RBP4/NPY1R/AR/ACACB/SYT1/CACNA2D2/DIO3/PLXNA4/LEP/OLFM1/IGF1/NRCAM/LIN7A/DUSP6/CPNE5/BCL2/FLVCR1/SMO/CXCL12/GLI1/SEMA3G/RAB21/GDI1/SPAAR/FXN/ARHGAP4/ADRB1/ANAPC2/PLXNA3/ZFPM2/STAT5B/LATS2/MAPK [illegible] |
| GOBP_REGULATION_OF_EPIDERMIS_DEVELOPMENT | GOBP_REGULATION_OF_EPIDERMIS_DEVELOPMENT | 49 | -0.5952708 | -1.533119 | 0.01227121 | 0.23031223 | 0.21338446 | 730 | tags=22%, list | VDR/PLAAT4/MYCL/TMEM79/CD109/FOXC1/HEY2/AQP3/GRHL1/ETV4/SFN |
| GOBP_CELLULAR_RESPONSE_TO_MINERALOCORTICOID_STIMULUS | GOBP_CELLULAR_RESPONSE_TO_MINERALOCORTICOID_STIMULUS | 8 | -0.8720545 | -1.5310609 | 0.01232666 | 0.23031223 | 0.21338446 | 857 | tags=38%, list | SCNN1D/RAN/SCNN1A |
| GOBP_REGULATION_OF_RENAL_SODIUM_EXCRETION | GOBP_REGULATION_OF_RENAL_SODIUM_EXCRETION | 4 | 0.9611372 | 1.49421108 | 0.01232505 | 0.23031223 | 0.21338446 | 56 | tags=25%, list | AGT |
| GOBP_CELL_CELL_JUNCTION_ORGANIZATION | GOBP_CELL_CELL_JUNCTION_ORGANIZATION | 158 | -0.453126 | -1.3822623 | 0.01239856 | 0.2310808 | 0.21409654 | 1218 | tags=18%, list | GJA1/OCLN/IRX3/CDH2/[illegible]/RDX/WDR1/HIPK1/NPHP1/NPHP4/DSG2/CLDN12/TRPV4/CDHR3/TJP1/PRKCA/F11R/LSR/CD9/TGFB2/MARVELD2/CXADR/CDH3/GRHL1/CLDN1/CLDN7/PERP/DSP |
| GOBP_MONOSACCHARIDE_TRANSMEMBRANE_TRANSPORT | GOBP_MONOSACCHARIDE_TRANSMEMBRANE_TRANSPORT | 86 | 0.54312295 | 1.49255811 | 0.01253508 | 0.23132919 | 0.21432668 | 810 | tags=16%, list | ADIPOQ/KLF15/TRARG1/LEP/SLC2A4/IGF1/IRS2/SORBS1/GPC3/EDNRA/PRKAG2/FFAR4/PIK3R1/SLC45A1 |
| GOBP_CARBOHYDRATE_TRANSPORT | GOBP_CARBOHYDRATE_TRANSPORT | 95 | 0.52967637 | 1.47350192 | 0.01253039 | 0.23132919 | 0.21432668 | 833 | tags=17%, list | ADIPOQ/AQP7/KLF15/TRARG1/LEP/SLC2A4/IGF1/IRS2/SORBS1/GPC3/EDNRA/PRKAG2/FFAR4/PIK3R1/SLC45A1/AQP1 |
| GOBP_PLASMA_MEMBRANE_ORGANIZATION | GOBP_PLASMA_MEMBRANE_ORGANIZATION | 127 | -0.4776216 | -1.4189127 | 0.01246992 | 0.23132919 | 0.21432668 | 2503 | tags=28%, list | CHMP2B/P2RX7/CHMP2A/CHMP3/OSBPL2/MICALL1/SYT11/MYH9/SERP1/UGCG/PRX/RAB3A/SOD1/NSF/SYT7/ATP2A2/CSRP3/IQGAP1/MYOF/MAFB/XKR6/ABCA7/SERINC2/CLPTM1L/CD9/CHMP4B/TGFB2/TLCD1/A4GALT/ANK3/S100A10/BA [illegible] |
| GOBP_SEX_DIFFERENTIATION | GOBP_SEX_DIFFERENTIATION | 203 | 0.44789377 | 1.36973264 | 0.01250447 | 0.23132919 | 0.21432668 | 1289 | tags=18%, list | RBP4/CNTFR/FZD4/PTX3/AR/MMP19/BMP6/NUP107/CSMD1/LHX9/LEP/FST/DMRT3/HSD17B3/IDH1/ATN1/GREB1L/BCL2/NOTCH4/CITED2/ARID5B/KIT/ADCYAP1R1/EIF2B4/ROBO2/TYRO3/SLIT3/SFRP1/RHOBTB3/BOK/HESX1/TESC/RARA/ZFP [illegible] |
| GOBP_POSITIVE_REGULATION_OF_LEUKOCYTE_APOPTOTIC_PROCESS | GOBP_POSITIVE_REGULATION_OF_LEUKOCYTE_APOPTOTIC_PROCESS | 26 | -0.6948869 | -1.5812271 | 0.01266077 | 0.23260434 | 0.2155081 | 456 | tags=23%, list | P2RX7/CDKN2A/IDO1/LYN/TGFB2/PERP |
| GOBP_NITRIC_OXIDE_CGMP_MEDIATED_SIGNALING | GOBP_NITRIC_OXIDE_CGMP_MEDIATED_SIGNALING | 10 | 0.85898926 | 1.57977447 | 0.01266611 | 0.23260434 | 0.2155081 | 238 | tags=20%, list | AGT/CD36 |
| GOBP_SENSORY_ORGAN_DEVELOPMENT | GOBP_SENSORY_ORGAN_DEVELOPMENT | 459 | 0.38792616 | 1.2753621 | 0.01272179 | 0.23305707 | 0.21592756 | 1428 | tags=16%, list | RBP4/FZD4/SLC17A8/TCF15/ADAMTS9/CDKN1C/ADAMTS18/PLAAT3/BMPER/BMP6/FOS/CEBPA/NFIB/DCX/ARHGEF15/LRP5/DIO3/LGR5/PXDN/CACNA1C/COL2A1/LAMB2/SPRY2/GSC/PDZD7/HOXA2/SMARCD3/PRICKLE1/IGF1/SMOC1/PAX5/ [illegible] |
| GOBP_RESPONSE_TO_FATTY_ACID | GOBP_RESPONSE_TO_FATTY_ACID | 48 | 0.61502262 | 1.53377546 | 0.01284509 | 0.23455497 | 0.21731536 | 1188 | tags=21%, list | ADIPOQ/DGAT2/CD36/PDK4/ASS1/LPL/FOXO1/PRKCD/GIPR/CDK4 |
| GOBP_MICROTUBULE_ORGANIZING_CENTER_ORGANIZATION | GOBP_MICROTUBULE_ORGANIZING_CENTER_ORGANIZATION | 148 | -0.4592428 | -1.3896141 | 0.01286602 | 0.23455497 | 0.21731536 | 2902 | tags=27%, list | VPS4B/CROCC/FBXW11/BBS4/DNM2/STIL/GADD45A/CCP110/AURKA/TUBA1A/POC1B/CLASP1/CHMP2B/CHMP2A/CHMP3/CEP135/PDE4DIP/SASS6/WDR62/CKAP5/TRIM37/[illegible]/NDC80/POC1A/CEP295/PCM1/GEN1/C2CD3/CDK11B/O [illegible] |
| GOBP_OUTFLOW_TRACT_MORPHOGENESIS | GOBP_OUTFLOW_TRACT_MORPHOGENESIS | 75 | 0.5527111 | 1.48815764 | 0.01317275 | 0.23956546 | 0.22195758 | 2373 | tags=36%, list | NPY1R/CLDN5/PRICKLE1/GREB1L/EDNRA/CITED2/ROBO2/PLXND1/JUN/ENG/RARA/ZFPM2/SOX17/WNT11/GATA6/SOX11/SFRP2/ILK/TBX1/ROBO1/PITX2/NRP1/FZD2/EYA1/SMAD4/ELN/FGFR2 |
| GOBP_CELLULAR_RESPONSE_TO_OSMOTIC_STRESS | GOBP_CELLULAR_RESPONSE_TO_OSMOTIC_STRESS | 54 | -0.5834427 | -1.5267324 | 0.01321032 | 0.23966833 | 0.22205289 | 1881 | tags=46%, list | CLN3/USP15/TSPO/DDX3X/BAD/EFHD1/ARHGEF2/ERRFI1/ATP1A1/NINJ1/PYCARD/WNK3/STK39/XRCC6/YBX3/LRRC8E/CASP3/SLC12A6/AQP5/TRPV4/VPS13A/LRRC8D/BDKRB2/SLC4A11/SLC2A1 |
| GOBP_POSITIVE_REGULATION_OF_LONG_TERM_SYNAPTIC_DEPRESSIO | GOBP_POSITIVE_REGULATION_OF_LONG_TERM_SYNAPTIC_DEPRESSIO | 7 | 0.89797796 | 1.53672581 | 0.01349388 | 0.2442229 | 0.2262727 | 143 | tags=29%, list | CBLN1/GRID2 |
| GOBP_POSITIVE_REGULATION_OF_LIPID_LOCALIZATION | GOBP_POSITIVE_REGULATION_OF_LIPID_LOCALIZATION | 72 | 0.55962748 | 1.49759233 | 0.01355422 | 0.2447253 | 0.22673818 | 736 | tags=17%, list | ADIPOQ/PLIN5/PLA2R1/PPARG/BMP6/ACACB/NR1H3/CD36/LPL/ACSL5/PLA2G4A/OXT |
| GOBP_SEXUAL_REPRODUCTION | GOBP_SEXUAL_REPRODUCTION | 774 | -0.3421844 | -1.20088 | 0.01359434 | 0.24486121 | 0.2268641 | 1844 | tags=15%, list | SRC/NDRG3/HUS1/UBE2Q1/ACTR3/NDC80/GGT1/ZWINT/YTHDF2/SIRT7/POC1A/SPAG17/REC8/CDYL/ZNF35/SMC1A/BMAL1/SKA2/RAB3A/PHC2/[illegible]/RAD51C/LGR4/HOOK1/IQCG/MFGE8/TMEM119/ANTXR2/SLC4A2/SOD1/CCDC181/TUBA [illegible] |
| GOBP_PARANODAL_JUNCTION_ASSEMBLY | GOBP_PARANODAL_JUNCTION_ASSEMBLY | 5 | -0.92528 | -1.4715491 | 0.01375884 | 0.24723118 | 0.22905988 | 311 | tags=40%, list | UGT8/CD9 |
| GOBP_INTRACILIARY_ANTEROGRADE_TRANSPORT | GOBP_INTRACILIARY_ANTEROGRADE_TRANSPORT | 18 | -0.748706 | -1.577076 | 0.01383946 | 0.24733631 | 0.22915729 | 2454 | tags=61%, list | IFT25/IFT27/IFT20/IFT80/[illegible]/IFT46/TRAF3IP1/IFT57/IFT56/IFT22/IFT81 |
| GOBP_GOLGI_TO_VACUOLE_TRANSPORT | GOBP_GOLGI_TO_VACUOLE_TRANSPORT | 23 | -0.7086974 | -1.5657441 | 0.01386099 | 0.24733631 | 0.22915729 | 2094 | tags=35%, list | VTI1A/CLN3/AP3D1/LAMP1/AP1M1/VTI1B/EHD3/CCDC91 |
| GOBP_KETONE_METABOLIC_PROCESS | GOBP_KETONE_METABOLIC_PROCESS | 168 | 0.46525651 | 1.38936966 | 0.01389641 | 0.24733631 | 0.22915729 | 643 | tags=12%, list | ADIPOQ/HSD11B2/PLIN5/DGAT2/PPARG/BMP6/GPIHBP1/FMO1/ACACB/NR1H3/PDK4/EGR1/HSD17B3/ACADVL/IRS2/FMO2/ANGPTL4/GATD1/PRKAG2/MLXIPL |
| GOBP_TRANSMEMBRANE_TRANSPORT | GOBP_TRANSMEMBRANE_TRANSPORT | 1056 | 0.34499365 | 1.19268833 | 0.01388465 | 0.24733631 | 0.22915729 | 1553 | tags=14%, list | ADIPOQ/RBP4/SLC29A4/AQP7/SLC17A8/C7/KLF15/AGT/POU3F3/RYR2/SLC25A18/OPRK1/GABRB1/TRARG1/GRID2/SLC35E3/TMEM163/GRIA2/SLC19A1/SLC49A3/SLCO1C1/SYT1/CCL19/KCNMB4/SLC6A1/SLC26A7/CD36/CACNA2D2/LEP/ABC [illegible] |
| GOBP_REGULATION_OF_MACROPHAGE_DERIVED_FOAM_CELL_DIFFER | GOBP_REGULATION_OF_MACROPHAGE_DERIVED_FOAM_CELL_DIFFER | 23 | 0.73274096 | 1.6025858 | 0.01393312 | 0.24740348 | 0.22921952 | 279 | tags=26%, list | ADIPOQ/AGT/PPARG/NR1H3/CD36/LPL |
| GOBP_PROTEIN_ACTIVATION_CASCADE | GOBP_PROTEIN_ACTIVATION_CASCADE | 15 | -0.7744831 | -1.5705212 | 0.01408451 | 0.24798163 | 0.22975518 | 1394 | tags=33%, list | MVP/FN1/F3/HPSE/ITGB3 |
| GOBP_MAMMARY_GLAND_INVOLUTION | GOBP_MAMMARY_GLAND_INVOLUTION | 8 | -0.8668132 | -1.5218588 | 0.01406009 | 0.24798163 | 0.22975518 | 730 | tags=25%, list | VDR/ELF3 |
| GOBP_RESPONSE_TO_OXYGEN_GLUCOSE_DEPRIVATION | GOBP_RESPONSE_TO_OXYGEN_GLUCOSE_DEPRIVATION | 8 | -0.865776 | -1.5200378 | 0.01406009 | 0.24798163 | 0.22975518 | 269 | tags=38%, list | BNIP3/DRAM1/INHBA |
| GOBP_REGULATION_OF_NERVOUS_SYSTEM_PROCESS | GOBP_REGULATION_OF_NERVOUS_SYSTEM_PROCESS | 76 | 0.54891424 | 1.48110201 | 0.01409774 | 0.24798163 | 0.22975518 | 922 | tags=17%, list | S100B/CBLN1/ADORA1/MGLL/PRKAR1B/FGF12/IGF1/GRIN2B/CDK18/GRIN2C/GRIN2A/SHANK3/QKI |
| GOBP_CORTICAL_CYTOSKELETON_ORGANIZATION | GOBP_CORTICAL_CYTOSKELETON_ORGANIZATION | 54 | -0.5789295 | -1.5149225 | 0.01425875 | 0.24964453 | 0.23129586 | 1813 | tags=31%, list | ASB7/PPFIBP1/PDCD6IP/IQGAP3/RAC1/PTK2B/IQGAP1/KIF21A/RHOQ/LLGL2/WDR1/FMNL1/KCNC3/ARF6/TRPV4/PPFIA1/EZR |
| GOBP_REGULATION_OF_HORMONE_SECRETION | GOBP_REGULATION_OF_HORMONE_SECRETION | 174 | 0.46208903 | 1.38596242 | 0.01423937 | 0.24964453 | 0.23129586 | 1629 | tags=20%, list | ADIPOQ/RBP4/RAPGEF3/AGT/OPRK1/ADORA1/PPARG/BMP6/BRSK2/GABBR1/LRP5/LEP/ABCC8/GNAO1/IRS2/PDE8B/MLXIPL/GPLD1/FFAR4/ADCYAP1/PICK1/ITSN1/VAMP2/FOXO1/GIPR/INHBB/C1QTNF1/GNAZ/PLA2G6/GNAS/SNAP25/SIRT3 [illegible] |
| GOBP_REGULATION_OF_G_PROTEIN_COUPLED_RECEPTOR_SIGNALING | GOBP_REGULATION_OF_G_PROTEIN_COUPLED_RECEPTOR_SIGNALING | 116 | 0.50234586 | 1.43425509 | 0.01429632 | 0.2497202 | 0.23136596 | 1049 | tags=17%, list | MRAP/PDE3B/MGLL/RGS11/RGS6/GNG7/PDE3A/PDE4B/KCTD12/RAMP2/PDE2A/GRK3/NMT2/NOS1/PDE4D/ADCYAP1/PPP3CA/SLIT3/GIPR/RACK1 |
| GOBP_RENAL_FILTRATION | GOBP_RENAL_FILTRATION | 25 | 0.7149932 | 1.58719293 | 0.01453604 | 0.25273196 | 0.23415636 | 1317 | tags=36%, list | IGHA1/ADORA1/JCHAIN/EDNRA/PPP3CA/AQP1/ADGRF5/SULF1/CYBA |
| GOBP_INTERMEDIATE_FILAMENT_BASED_PROCESS | GOBP_INTERMEDIATE_FILAMENT_BASED_PROCESS | 40 | -0.6161529 | -1.5267463 | 0.01452885 | 0.25273196 | 0.23415636 | 546 | tags=18%, list | MTM1/DST/FAM83H/KRT7/DSP/KRT19/KRT17 |
| GOBP_RESPONSE_TO_BACTERIUM | GOBP_RESPONSE_TO_BACTERIUM | 447 | 0.38557698 | 1.2655301 | 0.01460957 | 0.25342377 | 0.23479732 | 1526 | tags=14%, list | ADIPOQ/IGHM/IGHA1/IGHG3/IGHG1/THRSP/FABP4/PCK1/ADAMTS9/OPRK1/PLAAT3/BMP6/FOS/NFIB/FMO1/C1QTNF4/JCHAIN/NR1H3/CD36/IL7R/ASS1/ABCC8/PDE4B/LPL/IGHG4/MMRN2/SHARPIN/SSC5D/TNFAIP3/CD96/GSDMB/RPL30/G [illegible] |
| GOBP_REGULATION_OF_BEHAVIOR | GOBP_REGULATION_OF_BEHAVIOR | 44 | 0.62720239 | 1.54115569 | 0.01480485 | 0.25618883 | 0.23735915 | 2385 | tags=27%, list | OPRK1/ADORA1/LEPR/CNTNAP4/ADRB1/MC1R/PCSK1N/NAPEPLD/DLG4/POMC/MGRN1/CCL3 |
| GOBP_UROGENITAL_SYSTEM_DEVELOPMENT | GOBP_UROGENITAL_SYSTEM_DEVELOPMENT | 52 | 0.59974006 | 1.5161275 | 0.01489687 | 0.25618883 | 0.23735915 | 2681 | tags=42%, list | AR/IGF1/GLI1/SFRP1/SULF1/CYP7B1/RARA/ID4/EPHB3/WNT11/HOXA10/PTCH1/HOXA9/ESR1/PRLR/FRS2/SERPINF1/FGFR2/PLAG1/STAT5A/OSR2/OSR1 |
| GOBP_RESPONSE_TO_CGMP | GOBP_RESPONSE_TO_CGMP | 5 | 0.93674469 | 1.51064209 | 0.01488782 | 0.25618883 | 0.23735915 | 507 | tags=40%, list | PDE3A/PDE2A |
| GOBP_SEPTUM_PRIMUM_DEVELOPMENT | GOBP_SEPTUM_PRIMUM_DEVELOPMENT | 5 | -0.9237738 | -1.4691537 | 0.01490541 | 0.25618883 | 0.23735915 | 303 | tags=60%, list | ACVR1/TGFB2/SOX4 |
| GOBP_REGULATION_OF_RECEPTOR_BINDING | GOBP_REGULATION_OF_RECEPTOR_BINDING | 6 | 0.91616902 | 1.52777218 | 0.01494086 | 0.25621186 | 0.23738048 | 80 | tags=17%, list | IL11RA |
| GOBP_NEGATIVE_REGULATION_OF_ENDOTHELIAL_CELL_MIGRATION | GOBP_NEGATIVE_REGULATION_OF_ENDOTHELIAL_CELL_MIGRATION | 44 | 0.62578387 | 1.53767011 | 0.01499712 | 0.25659075 | 0.23773153 | 881 | tags=9%, list | ADAMTS9/PPARG/MMRN2/ANGPT2 |
| GOBP_RENAL_SYSTEM_PROCESS_INVOLVED_IN_REGULATION_OF_SYST | GOBP_RENAL_SYSTEM_PROCESS_INVOLVED_IN_REGULATION_OF_SYST | 17 | 0.77712686 | 1.59473029 | 0.0150805 | 0.2566359 | 0.23777336 | 1317 | tags=29%, list | HSD11B2/AGT/ADORA1/EDNRB/CYBA |
| GOBP_RESPONSE_TO_GLUCAGON | GOBP_RESPONSE_TO_GLUCAGON | 24 | 0.72085881 | 1.58969907 | 0.01513643 | 0.2566359 | 0.23777336 | 1908 | tags=33%, list | PRKAR1B/ASS1/PRKAR2B/ADCY4/FUT1/CDO1/GNAS/ADCY6 |
| GOBP_REGULATION_OF_D_GLUCOSE_TRANSMEMBRANE_TRANSPORT | GOBP_REGULATION_OF_D_GLUCOSE_TRANSMEMBRANE_TRANSPORT | 60 | 0.57763666 | 1.50079192 | 0.01505174 | 0.2566359 | 0.23777336 | 747 | tags=18%, list | ADIPOQ/KLF15/LEP/IGF1/IRS2/SORBS1/GPC3/EDNRA/PRKAG2/FFAR4/PIK3R1 |
| GOBP_OOGENESIS | GOBP_OOGENESIS | 67 | 0.56295966 | 1.48731699 | 0.01512573 | 0.2566359 | 0.23777336 | 1660 | tags=21%, list | PTX3/LGR5/PDE3A/PTN/PIWIL2/IGF1/BCL2/EDNRA/RPS6KA2/INHBB/YTHDC2/IHO1/ATM/TAF4B |
| GOBP_REGULATION_OF_PROTEIN_FOLDING | GOBP_REGULATION_OF_PROTEIN_FOLDING | 12 | -0.8086994 | -1.5567205 | 0.01531815 | 0.25913197 | 0.24008597 | 1619 | tags=33%, list | GRN/HSPA5/DNAJB1/PDCL3 |
| GOBP_INNER_DYNEIN_ARM_ASSEMBLY | GOBP_INNER_DYNEIN_ARM_ASSEMBLY | 8 | -0.8623123 | -1.5139566 | 0.01540832 | 0.26003793 | 0.24092535 | 1019 | tags=50%, list | CCDC40/DNAI3/DNAAF11/DNAAF4 |
| GOBP_MULTICELLULAR_ORGANISMAL_LEVEL_HOMEOSTASIS | GOBP_MULTICELLULAR_ORGANISMAL_LEVEL_HOMEOSTASIS | 695 | 0.36088124 | 1.21958958 | 0.01544094 | 0.26003793 | 0.24092535 | 1545 | tags=15%, list | ADIPOQ/RBP4/DLK1/CLDN5/ANGPT1/FABP4/ADORA1/LEPR/BMP6/CEBPA/EBF2/NFIB/FMO1/G0S2/ACACB/C1QTNF4/PRKAR1B/NR1H3/CD36/IL7R/PDK4/LEP/ABCC8/PDE4B/COL2A1/ITPKB/EGR1/PTGER3/HOXC10/ACOT11/PRKAR2B/THRA/AL [illegible] |
| GOBP_RECEPTOR_GUANYLYL_CYCLASE_SIGNALING_PATHWAY | GOBP_RECEPTOR_GUANYLYL_CYCLASE_SIGNALING_PATHWAY | 8 | 0.88347261 | 1.55148667 | 0.01559252 | 0.26141827 | 0.24220423 | 507 | tags=25%, list | GNG7/PDE2A |
| GOBP_MIDGUT_DEVELOPMENT | GOBP_MIDGUT_DEVELOPMENT | 8 | 0.88328997 | 1.55116593 | 0.01559252 | 0.26141827 | 0.24220423 | 836 | tags=50%, list | RET/ASS1/SMO/EDNRB |
| GOBP_MEMBRANE_RAFT_ORGANIZATION | GOBP_MEMBRANE_RAFT_ORGANIZATION | 28 | -0.6722894 | -1.5553315 | 0.01575758 | 0.26286276 | 0.24354255 | 1029 | tags=21%, list | IQGAP1/NPC1/MAL/ABCA7/S100A10/ANXA2 |
| GOBP_REGULATION_OF_KERATINOCYTE_PROLIFERATION | GOBP_REGULATION_OF_KERATINOCYTE_PROLIFERATION | 30 | -0.6590246 | -1.5440219 | 0.01585366 | 0.26286276 | 0.24354255 | 730 | tags=17%, list | VDR/CD109/CDH3/AREG/SFN |
| GOBP_ZYMOGEN_ACTIVATION | GOBP_ZYMOGEN_ACTIVATION | 35 | -0.633866 | -1.5347434 | 0.01582734 | 0.26286276 | 0.24354255 | 1759 | tags=37%, list | GGT1/PLAU/MELTF/PLGRKT/PGK1/CTSH/RUNX1/MMP14/ENO1/S100A10/PRSS12/ANXA2/DHCR24 |
| GOBP_POSITIVE_REGULATION_OF_ORGANELLE_ASSEMBLY | GOBP_POSITIVE_REGULATION_OF_ORGANELLE_ASSEMBLY | 81 | -0.5224271 | -1.4579063 | 0.01582549 | 0.26286276 | 0.24354255 | 2902 | tags=31%, list | NUP62/UBAP2L/ATG5/VPS4B/CROCC/BBS4/STIL/SEPTIN9/CCP110/POC1B/CEP135/IFT20/SEPTIN7/SASS6/SRC/PDCD6IP/CEP295/ULK1/SDC1/SNX7/PIP4K2C/CNOT6/GPSM2/SDC4/MNS1 |
| GOBP_SENSORY_SYSTEM_DEVELOPMENT | GOBP_SENSORY_SYSTEM_DEVELOPMENT | 323 | 0.40434602 | 1.295344 | 0.01578045 | 0.26286276 | 0.24354255 | 2272 | tags=24%, list | RBP4/FZD4/SLC17A8/ADAMTS9/CDKN1C/ADAMTS18/PLAAT3/BMP6/FOS/NFIB/DCX/ARHGEF15/LRP5/DIO3/PXDN/CACNA1C/LAMB2/SPRY2/SMARCD3/PRICKLE1/SMOC1/EPHB1/TMEM135/MYOM1/TMEM132E/BCL2/CRYGS/CITED2/PRDM1/S [illegible] |
| GOBP_ACTIVATION_OF_PROTEIN_KINASE_ACTIVITY | GOBP_ACTIVATION_OF_PROTEIN_KINASE_ACTIVITY | 41 | 0.63315753 | 1.53780228 | 0.01600771 | 0.26309397 | 0.24375677 | 516 | tags=17%, list | ANGPT1/PRKAR1B/STRADB/LEP/PRKAR2B/IGF1/LTB |
| GOBP_NEURON_MATURATION | GOBP_NEURON_MATURATION | 43 | 0.6273242 | 1.53654554 | 0.01592479 | 0.26309397 | 0.24375677 | 867 | tags=26%, list | RET/CDKN1C/NDRG4/MYOC/SCARF1/NRCAM/KCNIP2/BCL2/EDNRA/EDNRB/PICK1 |
| GOBP_FATTY_ACID_DERIVATIVE_METABOLIC_PROCESS | GOBP_FATTY_ACID_DERIVATIVE_METABOLIC_PROCESS | 58 | 0.58310937 | 1.50667357 | 0.01599548 | 0.26309397 | 0.24375677 | 1994 | tags=21%, list | DGAT2/DGAT1/PECR/ACSL5/GCDH/SLC27A5/ADHFE1/BCKDHA/CBR4/DLD/HPGD/ACSF3 |
| GOBP_CELL_ADHESION | GOBP_CELL_ADHESION | 1141 | 0.34016033 | 1.18066645 | 0.01594049 | 0.26309397 | 0.24375677 | 1963 | tags=21%, list | ADIPOQ/FZD4/CLDN5/ANGPT1/RET/PLEKHG4B/ZBTB16/COL6A6/PCK1/ADAMTS9/S100B/EGR3/CBLN1/MYOC/CDH6/PDE3B/ADAMTS18/COL18A1/TMEM131L/NTNG1/BMP6/GRID2/EGFLAM/SERPINE2/SEMA6A/CCL19/LRP5/CD83/SCARF1/CD [illegible] |
| GOBP_RESPONSE_TO_RETINOIC_ACID | GOBP_RESPONSE_TO_RETINOIC_ACID | 80 | 0.53742409 | 1.46318178 | 0.01606276 | 0.2634222 | 0.24406087 | 1698 | tags=26%, list | RBP4/FZD4/RET/MEST/BMP6/LEP/HOXA2/TIE1/OXT/AQP1/ASXL1/CYP26B1/TESC/RARA/RXRA/WNT11/YAP1/PTCH1/GATA6/ATM/TEAD2 |
| GOBP_REGULATION_OF_ACUTE_INFLAMMATORY_RESPONSE_TO_ANTI | GOBP_REGULATION_OF_ACUTE_INFLAMMATORY_RESPONSE_TO_ANTI | 10 | -0.8338852 | -1.5415646 | 0.01639028 | 0.26820785 | 0.24849479 | 1498 | tags=50%, list | FCGR2B/PARK7/SELENOS/C3/IL20RB |
| GOBP_DIACYLGLYCEROL_BIOSYNTHETIC_PROCESS | GOBP_DIACYLGLYCEROL_BIOSYNTHETIC_PROCESS | 6 | 0.9135173 | 1.52335025 | 0.01660095 | 0.27047672 | 0.2505969 | 39 | tags=33%, list | DGAT2/PNPLA2 |
| GOBP_NEGATIVE_REGULATION_OF_MONOCYTE_CHEMOTAXIS | GOBP_NEGATIVE_REGULATION_OF_MONOCYTE_CHEMOTAXIS | 6 | -0.9033365 | -1.4861566 | 0.01659271 | 0.27047672 | 0.2505969 | 865 | tags=33%, list | GREM1/NBL1 |
| GOBP_INTRACELLULAR_PH_REDUCTION | GOBP_INTRACELLULAR_PH_REDUCTION | 49 | -0.5804369 | -1.4949142 | 0.01663894 | 0.27050875 | 0.25062657 | 3579 | tags=55%, list | WDR7/VMA21/CLIC4/SLC9A8/CLN3/GPR89B/ATP6V0B/SLC4A7/CLN5/SNAPIN/LAMP2/DMXL2/RAB38/CREG1/VMA22/LAMP1/ATP6V1D/CLCN3/GRN/ATP6V0E2/ATP6V1F/ATP6V1A/ATP6AP2/ROGDI/TCIRG1/SLC9A7/ATP6V0E1 |
| GOBP_PROSTATE_GLAND_MORPHOGENESIS | GOBP_PROSTATE_GLAND_MORPHOGENESIS | 22 | 0.73570067 | 1.59092708 | 0.01669013 | 0.27075495 | 0.25085467 | 2447 | tags=45%, list | AR/IGF1/SFRP1/SULF1/CYP7B1/ID4/ESR1/FRS2/FGFR2/STAT5A |
| GOBP_NEPHRON_DEVELOPMENT | GOBP_NEPHRON_DEVELOPMENT | 128 | 0.48853257 | 1.41357788 | 0.0167025 | 0.2709828 | 0.25106578 | 1555 | tags=25%, list | ANGPT1/RET/KLF15/AGT/POU3F3/NUP107/EGR1/LAMB2/GREB1L/ADAMTS16/GPC3/BCL2/MAGI2/EDNRA/SMO/PECAM1/NUP160/PPP3CA/AQP1/EDNRB/ANGPT2/NID1/CITED1/ASXL1/NOTCH3/SULF1/PLCE1/WNT11/DLL1/ENPEP/YAP1/PTC [illegible] |
| GOBP_NEUROTRANSMITTER_TRANSPORT | GOBP_NEUROTRANSMITTER_TRANSPORT | 148 | 0.46975424 | 1.38102602 | 0.01692756 | 0.27228901 | 0.25227598 | 956 | tags=14%, list | SLC29A4/SLC17A8/CPLX1/SYT1/KCNMB4/SLC6A1/SNCG/ABCC8/GRM4/SLC1A7/UNC13C/SV2B/LIN7A/SNCAIP/SLC18A2/STXBP1/NOS1/NRXN2/PPP3CA/VAMP2/CACNB4 |
| GOBP_POSITIVE_REGULATION_OF_HEMOPOIESIS | GOBP_POSITIVE_REGULATION_OF_HEMOPOIESIS | 161 | 0.46394552 | 1.37917312 | 0.01687309 | 0.27228901 | 0.25227598 | 1315 | tags=14%, list | DLK1/ZBTB16/PCK1/EGR3/FOS/CCL19/CD83/IL7R/ITPKB/TESPA1/SMARCD3/ZAP70/LCK/PPP3CA/ACIN1/ZBTB46/ZBTB7B/TESC/RARA/STAT5B/MALT1/SOX12 |
| GOBP_REGULATION_OF_CELL_CYCLE | GOBP_REGULATION_OF_CELL_CYCLE | 971 | -0.3283605 | -1.166392 | 0.0169297 | 0.27228901 | 0.25227598 | 2939 | tags=22%, list | HSPA1B/DAPK3/TMEM14B/NAT9/VPS4B/PSMB1/HES1/MORF4L1/SMARCA4/PHACTR4/CCND1/ERCC3/DDX3X/BBS4/KNL1/BCL2L1/PAF1/PRR11/NUSAP1/[illegible]/VPS72/CDC27/STIL/[illegible]/TADA3/GADD45A/CCP110/NFE2L1/DONSON/MAD2 [illegible] |
| GOBP_REGULATION_OF_ORGANIC_ACID_TRANSPORT | GOBP_REGULATION_OF_ORGANIC_ACID_TRANSPORT | 45 | 0.61632789 | 1.52214094 | 0.01701396 | 0.2724772 | 0.25245036 | 971 | tags=27%, list | SLC17A8/ADORA1/PLA2R1/GABBR1/LEP/ACSL5/IRS2/PLA2G4A/STXBP1/OXT/SLC43A1/SEPTIN2 |
| GOBP_IMMUNE_EFFECTOR_PROCESS | GOBP_IMMUNE_EFFECTOR_PROCESS | 477 | 0.37972447 | 1.25160385 | 0.01698867 | 0.2724772 | 0.25245036 | 1565 | tags=13%, list | RBP4/IGHA1/IGHG3/ANGPT1/C7/IGHG1/IGLC2/PCK1/PTX3/MZB1/CCL19/CD36/IL7R/LEP/COLEC11/DNASE1/IGHG4/IGKC/PRAM1/CD96/GAB2/IRF4/SLC18A2/IRF7/STXBP1/UNG/KIT/ITM2A/PIK3R1/CD160/CD8A/KMO/VAMP2/SEMA4A/E [illegible] |
| GOBP_NEURAL_CREST_CELL_DIFFERENTIATION | GOBP_NEURAL_CREST_CELL_DIFFERENTIATION | 85 | 0.52909019 | 1.45181986 | 0.01722846 | 0.27462746 | 0.25444256 | 2236 | tags=29%, list | RET/SEMA6A/FGF12/GSC/EFNB1/EDNRA/SMO/CITED2/SEMA3G/FRZB/EDNRB/SEMA3B/SEMA4A/SFRP1/SEMA6B/ENG/TCOF1/SEMA3D/SEMA3A/TBX1/EXT1/PITX2/NRP1/ALDH1A2/SMAD4 |
| GOBP_FOREBRAIN_DORSAL_VENTRAL_PATTERN_FORMATION | GOBP_FOREBRAIN_DORSAL_VENTRAL_PATTERN_FORMATION | 4 | -0.9432515 | -1.4321111 | 0.01725791 | 0.27462746 | 0.25444256 | 24 | tags=50%, list | GLI3/SIX3 |
| GOBP_CELLULAR_RESPONSE_TO_CHEMICAL_STRESS | GOBP_CELLULAR_RESPONSE_TO_CHEMICAL_STRESS | 280 | -0.3956779 | -1.2835353 | 0.01725768 | 0.27462746 | 0.25444256 | 2985 | tags=30%, list | PCNA/USP15/PRKRA/HSPA1B/STX2/TSPO/GCH1/FAS/DDX3X/BAD/NFE2L1/PYCR2/PAWR/TXN/STK26/TRPM2/PINK1/EFHD1/ARHGEF2/ABL2/SIN3A/ADPRS/CHCHD2/ATG7/HSPA1A/GSR/GDF15/OSER1/ENDOG/ERMP1/VRK2/SESN2/ERRFI1/BR [illegible] |
| GOBP_SEROTONIN_METABOLIC_PROCESS | GOBP_SEROTONIN_METABOLIC_PROCESS | 12 | 0.82241709 | 1.57320705 | 0.01731161 | 0.27477106 | 0.25457561 | 2042 | tags=50%, list | MAOB/SULT1A4/GRIN2A/PDE1B/ALDH2/MDGA1 |
| GOBP_POSITIVE_REGULATION_OF_NEUROEPITHELIAL_CELL_DIFFEREN | GOBP_POSITIVE_REGULATION_OF_NEUROEPITHELIAL_CELL_DIFFEREN | 7 | -0.8782078 | -1.4982953 | 0.01735776 | 0.27477106 | 0.25457561 | 236 | tags=43%, list | CTNNB1/LEF1/BMP2 |
| GOBP_ADULT_BEHAVIOR | GOBP_ADULT_BEHAVIOR | 106 | 0.50666759 | 1.43086091 | 0.01737668 | 0.27477106 | 0.25457561 | 1829 | tags=24%, list | FOSB/OPRK1/SNCG/LEP/FGF12/SHANK2/DMRT3/PAK5/PREX2/EFNB3/SEZ6L2/CXCL12/NRXN2/SHANK3/FXN/CACNB4/SEPTIN5/UCHL1/SLITRK5/PPP1R1B/SLITRK6/PPARA/PUM1/NAGLU/TPGS1 |
| GOBP_NERVE_DEVELOPMENT | GOBP_NERVE_DEVELOPMENT | 64 | 0.56580166 | 1.48543639 | 0.01749436 | 0.27603329 | 0.25574506 | 2053 | tags=34%, list | HOXA3/COL25A1/SERPINE2/PLXNA4/EPHB1/EDNRA/CITED2/SULF1/PLXNA3/RPL24/VCAM1/SEMA3A/SLITRK6/ILK/TBX1/NAGLU/ARK2C/B4GALNT1/EXT1/ADARB1/NGFR/NRP1 |
| GOBP_CELL_MORPHOGENESIS | GOBP_CELL_MORPHOGENESIS | 799 | 0.35315348 | 1.20245159 | 0.01753001 | 0.27603329 | 0.25574506 | 1870 | tags=19%, list | FZD4/RET/PLEKHG4B/S100B/AR/CAMK2B/CDH6/COL18A1/COL25A1/NTNG1/BRSK2/CHODL/NFIB/NEFL/TIAM2/SEMA6A/LHX9/OBSL1/SYT1/STRADB/PLXNA4/IL7R/PTN/LAMB2/PDZD7/HOXA2/OLFM1/PRICKLE1/NRCAM/PREX2/MAP4K4/EPHB [illegible] |
| GOBP_REGULATION_OF_KERATINOCYTE_DIFFERENTIATION | GOBP_REGULATION_OF_KERATINOCYTE_DIFFERENTIATION | 28 | -0.6673561 | -1.5439184 | 0.01757576 | 0.27617472 | 0.25587609 | 730 | tags=32%, list | ERRFI1/MACROH2A1/VDR/PLAAT4/CD109/FOXC1/AQP3/GRHL1/ETV4 |
| GOBP_POSITIVE_REGULATION_OF_T_CELL_APOPTOTIC_PROCESS | GOBP_POSITIVE_REGULATION_OF_T_CELL_APOPTOTIC_PROCESS | 12 | -0.8010927 | -1.5420779 | 0.01767478 | 0.27657355 | 0.25624561 | 303 | tags=25%, list | IDO1/TGFB2/PERP |
| GOBP_POSITIVE_REGULATION_OF_KIDNEY_DEVELOPMENT | GOBP_POSITIVE_REGULATION_OF_KIDNEY_DEVELOPMENT | 6 | 0.90894476 | 1.51572524 | 0.01763851 | 0.27657355 | 0.25624561 | 754 | tags=33%, list | RET/PPP3CA |
| GOBP_AMINO_ACID_BETAINE_BIOSYNTHETIC_PROCESS | GOBP_AMINO_ACID_BETAINE_BIOSYNTHETIC_PROCESS | 5 | 0.93095762 | 1.50130956 | 0.01782344 | 0.27831992 | 0.25786362 | 235 | tags=20%, list | CHDH |
| GOBP_PLASMA_MEMBRANE_REPAIR | GOBP_PLASMA_MEMBRANE_REPAIR | 27 | -0.6714628 | -1.5397109 | 0.01791826 | 0.27922003 | 0.25869758 | 3374 | tags=52%, list | ARL8B/CHMP5/VPS4B/CHMP2B/CHMP2A/CHMP3/SYT11/MYH9/RAB3A/SYT7/MYOF/CHMP4B/S100A10/ANXA2 |
| GOBP_SENSORY_PERCEPTION_OF_PAIN | GOBP_SENSORY_PERCEPTION_OF_PAIN | 50 | 0.59857585 | 1.50016384 | 0.0180804 | 0.2794275 | 0.25889003 | 970 | tags=24%, list | NPY1R/OPRK1/ADORA1/MGLL/EPHB1/CXCL12/ANO1/OPRL1/KCND2/GRIN2A/AQP1/TMEM120A |
| GOBP_INNATE_IMMUNE_RESPONSE_ACTIVATING_CELL_SURFACE_RECE | GOBP_INNATE_IMMUNE_RESPONSE_ACTIVATING_CELL_SURFACE_RECE | 77 | 0.53791518 | 1.45566999 | 0.01803833 | 0.2794275 | 0.25889003 | 1944 | tags=23%, list | C1QTNF4/NR1H3/CD36/COLEC11/RAB11FIP2/TNFAIP3/PIK3R1/CREBBP/ZNRF1/CYBA/IRF3/PAK3/MASP1/FCN3/IFI35/ECSIT/NAGLU/TNIP3 |
| GOBP_CENTRAL_NERVOUS_SYSTEM_NEURON_DIFFERENTIATION | GOBP_CENTRAL_NERVOUS_SYSTEM_NEURON_DIFFERENTIATION | 152 | 0.46799286 | 1.38149497 | 0.01801477 | 0.2794275 | 0.25889003 | 2053 | tags=25%, list | GABRB1/CBLN1/GRID2/NFIB/PLXNA4/LEP/HOXC10/DMRT3/BTG2/EPHB1/UNC5D/ATF5/LHX6/SMO/CDH11/ROBO2/LHX8/SHANK3/SLIT3/TAL1/SFRP1/EOMES/NTRK2/PLXNA3/ID4/LMX1A/CRK/EPHB3/GBA2/PTCH1/LBX1/SFRP2/ROBO1/NOVA [illegible] |
| GOBP_CELL_CHEMOTAXIS | GOBP_CELL_CHEMOTAXIS | 232 | -0.4040831 | -1.2902501 | 0.0180778 | 0.2794275 | 0.25889003 | 1278 | tags=11%, list | RAC1/STK39/TNFSF14/PTK2B/SCG2/TMSB4X/ACKR2/GREM1/HGF/TRPV4/GSTP1/HSD3B7/LYN/C3/SAA1/SBDS/TGFB2/MET/LEF1/CXADR/LGALS3/NBL1/S100A8/S100A9/PERP/CXCL1 |
| GOBP_REGULATION_OF_CHOLESTEROL_STORAGE | GOBP_REGULATION_OF_CHOLESTEROL_STORAGE | 15 | 0.78530677 | 1.57083652 | 0.01820041 | 0.2794617 | 0.25892148 | 279 | tags=27%, list | PPARG/NR1H3/CD36/LPL |
| GOBP_POSITIVE_REGULATION_OF_CAMP_PKA_SIGNAL_TRANSDUCTIO | GOBP_POSITIVE_REGULATION_OF_CAMP_PKA_SIGNAL_TRANSDUCTIO | 12 | 0.81921782 | 1.56708715 | 0.01812627 | 0.2794617 | 0.25892148 | 745 | tags=33%, list | ADIPOQ/MYOM1/SPATC1L/ADCYAP1 |
| GOBP_SYNAPTIC_VESICLE_LUMEN_ACIDIFICATION | GOBP_SYNAPTIC_VESICLE_LUMEN_ACIDIFICATION | 15 | -0.7640303 | -1.5493249 | 0.01819249 | 0.2794617 | 0.25892148 | 1653 | tags=53%, list | ATP6V1G1/ATP6V1D/CLCN3/ATP6V1F/ATP6V1A/ATP6V1E1/ATP6AP2/SLC17A7 |
| GOBP_STEM_CELL_FATE_COMMITMENT | GOBP_STEM_CELL_FATE_COMMITMENT | 7 | 0.88891483 | 1.52121592 | 0.01826863 | 0.2794617 | 0.25892148 | 1293 | tags=57%, list | GSC/EDNRA/SFRP1/SOX17 |
| GOBP_POSITIVE_REGULATION_OF_MAPKKK_CASCADE_BY_FIBROBLAST | GOBP_POSITIVE_REGULATION_OF_MAPKKK_CASCADE_BY_FIBROBLAST | 3 | 0.96943711 | 1.43659329 | 0.01825047 | 0.2794617 | 0.25892148 | 431 | tags=67%, list | KLB/KL |
| GOBP_REGULATION_OF_MYELOID_CELL_APOPTOTIC_PROCESS | GOBP_REGULATION_OF_MYELOID_CELL_APOPTOTIC_PROCESS | 21 | 0.73444869 | 1.577543 | 0.01845537 | 0.27980055 | 0.25923543 | 614 | tags=24%, list | ADIPOQ/ITPKB/THRA/BCL2/IRF7 |
| GOBP_REGULATION_OF_LONG_CHAIN_FATTY_ACID_IMPORT_ACROSS_ | GOBP_REGULATION_OF_LONG_CHAIN_FATTY_ACID_IMPORT_ACROSS_ | 6 | 0.90644538 | 1.51155737 | 0.01846856 | 0.27980055 | 0.25923543 | 491 | tags=33%, list | ACSL5/IRS2 |
| GOBP_REGULATION_OF_PLATELET_DERIVED_GROWTH_FACTOR_RECEP | GOBP_REGULATION_OF_PLATELET_DERIVED_GROWTH_FACTOR_RECEP | 5 | 0.92873028 | 1.49771765 | 0.01845251 | 0.27980055 | 0.25923543 | 2 | tags=20%, list | ADIPOQ |
| GOBP_DIGESTIVE_SYSTEM_DEVELOPMENT | GOBP_DIGESTIVE_SYSTEM_DEVELOPMENT | 101 | 0.50993603 | 1.43227168 | 0.01847704 | 0.27980055 | 0.25923543 | 1980 | tags=32%, list | RET/CDKN1C/ASS1/NPHP3/IGF1/HOXA5/BCL2/SMO/PRDM1/KIT/GLI1/EDNRB/SFRP1/SOX17/RBPMS2/EPHB3/ACVR2B/WNT11/COL3A1/YAP1/GATA6/WDPCP/NR5A2/SOX11/HLX/PKD1/SFRP2/DCHS1/RCBTB2/EXT1/TCF21/PITX2 |
| GOBP_G_PROTEIN_COUPLED_RECEPTOR_SIGNALING_PATHWAY | GOBP_G_PROTEIN_COUPLED_RECEPTOR_SIGNALING_PATHWAY | 480 | 0.3784088 | 1.24774656 | 0.01846615 | 0.27980055 | 0.25923543 | 1463 | tags=16%, list | MRAP/NPY1R/FZD4/RAPGEF3/AGT/OPRK1/ADORA1/PDE3B/PPARG/FZD8/MGLL/RGS11/PRKAR1B/RGS6/GNG7/CCL19/GABBR1/LGR5/PDE3A/PDE4B/GRM4/PTGER3/KCTD12/LPAR4/PRKAR2B/ADGRL1/RAMP2/PREX2/GNAO1/ADCY4/PDE2A/ [illegible] |
