## Supplementary material for "Integrated transcriptomic analysis identifies Paternally Expressed Gene 10 as a novel candidate biomarker associated with aggressive features in liposarcoma": Table S5

List of genes used to annotate tumor cell clusters, from Gruel, N., Quignot, C., Lesage, L. et al. Cellular origin and clonal evolution of human dedifferentiated liposarcoma. Nat Commun 15, 7941 (2024). https://doi.org/10.1038/s41467-024-52067-1

| Tumor cell cluster | Gene |
|---|---|
| Adipo diff | *CXCL14* |
| Adipo diff | *PTGDS* |
| Adipo diff | *CFD* |
| Adipo diff | *TSPAN8* |
| Adipo diff | *IGF1* |
| Adipo diff | *MT1M* |
| Adipo diff | *MT2A* |
| Adipo diff | *MT1A* |
| Adipo diff | *MGP* |
| Adipo diff | *MT1X* |
| Adipo diff | *GSN* |
| Adipo diff | *APOD* |
| Adipo diff | *C7* |
| Adipo diff | *CYR61* |
| Adipo diff | *GPX3* |
| Adipo diff | *EPHX1* |
| Adipo diff | *ZFP36* |
| Adipo diff | *DCN* |
| Adipo diff | *VIM* |
| Adipo diff | *RASD1* |
| Angiogenesis | *SMOC2* |
| Angiogenesis | *IGF2* |
| Angiogenesis | *CLEC14A* |
| Angiogenesis | *SCN7A* |
| Angiogenesis | *CLEC11A* |
| Angiogenesis | *HSPA1A* |
| Angiogenesis | *MOXD1* |
| Angiogenesis | *IGFBP2* |
| Angiogenesis | *NDUFA12* |
| Angiogenesis | *TMEM106C* |
| Angiogenesis | *RGS16* |
| Angiogenesis | *CAPN6* |
| Angiogenesis | *ATP5F1B* |
| Angiogenesis | *MMP11* |
| Angiogenesis | *COL15A1* |
| Angiogenesis | *SERPINE2* |
| Angiogenesis | *ARL1* |
| Angiogenesis | *CRLF1* |
| Angiogenesis | *DLK1* |
| Angiogenesis | *SPRY1* |
| ECM remodeling | *VCAN* |
| ECM remodeling | *COL1A1* |
| ECM remodeling | *EMILIN1* |
| ECM remodeling | *LTBP3* |
| ECM remodeling | *COL5A1* |
| ECM remodeling | *SLIT3* |
| ECM remodeling | *C3* |
| ECM remodeling | *ELN* |
| ECM remodeling | *LRP1* |
| ECM remodeling | *HSPG2* |
| ECM remodeling | *CHPF* |
| ECM remodeling | *PAPPA* |
| ECM remodeling | *MMP2* |
| ECM remodeling | *MTRNR2L12* |
| ECM remodeling | *KCNQ1OT1* |
| ECM remodeling | *MTRNR2L8* |
| ECM remodeling | *BTG1* |
| ECM remodeling | *MRC2* |
| ECM remodeling | *COL1A2* |
| ECM remodeling | *COL11A1* |
| Hypoxia | *NEAT1* |
| Hypoxia | *NKTR* |
| Hypoxia | *MALAT1* |
| Hypoxia | *WSB1* |
| Hypoxia | *KCNQ1OT1* |
| Hypoxia | *XIST* |
| Hypoxia | *MEG3* |
| Hypoxia | *MEG8* |
| Hypoxia | *HIF3A* |
| Hypoxia | *PLAGL1* |
| Hypoxia | *DDX17* |
| Hypoxia | *LUC7L3* |
| Hypoxia | *FTX* |
| Hypoxia | *SYNE1* |
| Hypoxia | *CSAD* |
| Hypoxia | *GABPB1-AS1* |
| Hypoxia | *IGF2BP2* |
| Hypoxia | *ARGLU1* |
| Hypoxia | *N4BP2L2* |
| Hypoxia | *ZRANB2* |
| Invasion | *C1QB* |
| Invasion | *IGKC* |
| Invasion | *RNASE1* |
| Invasion | *C1QA* |
| Invasion | *CD24* |
| Invasion | *RPL18A* |
| Invasion | *AC245595.1* |
| Invasion | *LDHA* |
| Invasion | *S100A11* |
| Invasion | *TAGLN2* |
| Invasion | *CD74* |
| Invasion | *CDK4* |
| Invasion | *AC123567.2* |
| Invasion | *HMGN3* |
| Invasion | *TSG101* |
| Invasion | *AGT* |
| Invasion | *HLA-DRA* |
| Invasion | *POSTN* |
| Invasion | *PDE4DIP* |
| Invasion | *IFI27* |
| Stemness | *PRG4* |
| Stemness | *IGFBP6* |
| Stemness | *PI16* |
| Stemness | *SEMA3C* |
| Stemness | *ACKR3* |
| Stemness | *MFAP5* |
| Stemness | *CLEC3B* |
| Stemness | *SFRP4* |
| Stemness | *SLPI* |
| Stemness | *HTRA3* |
| Stemness | *S100A4* |
| Stemness | *FBN1* |
| Stemness | *PCOLCE2* |
| Stemness | *PROCR* |
| Stemness | *S100A10* |
| Stemness | *TNXB* |
| Stemness | *C17orf58* |
| Stemness | *SCARA5* |
| Stemness | *CD248* |
| Stemness | *UAP1* |
